# Identification of PKCα-dependent phosphoproteins in mouse retina

**DOI:** 10.1101/589184

**Authors:** Colin M. Wakeham, Phillip A. Wilmarth, Jennifer M. Cunliffe, John E. Klimek, Gaoying Ren, Larry L. David, Catherine W. Morgans

## Abstract

Adjusting to a wide range of light intensities is an essential feature of retinal rod bipolar cell (RBC) function. While persuasive evidence suggests this modulation involves phosphorylation by protein kinase C-alpha (PKCα), the targets of PKCα phosphorylation in the retina have not been identified. PKCα activity and phosphorylation in RBCs was examined by immunofluorescence confocal microscopy using a conformation-specific PKCα antibody and antibodies to phosphorylated PKC motifs. PKCα activity was dependent on light and expression of TRPM1, and RBC dendrites were the primary sites of light-dependent phosphorylation. PKCα-dependent retinal phosphoproteins were identified using a phosphoproteomics approach to compare total protein and phosphopeptide abundance between phorbol ester-treated wild type and PKCα knockout (PKCα-KO) mouse retinas. Phosphopeptide mass spectrometry identified over 1100 phosphopeptides in mouse retina, with 12 displaying significantly greater phosphorylation in WT compared to PKCα-KO samples. The differentially phosphorylated proteins fall into the following functional groups: cytoskeleton/trafficking (4 proteins), ECM/adhesion (2 proteins), signaling (2 proteins), transcriptional regulation (3 proteins), and homeostasis/metabolism (1 protein). Two strongly differentially expressed phosphoproteins, BORG4 and TPBG, were localized to the synaptic layers of the retina, and may play a role in PKCα-dependent modulation of RBC physiology. Data are available via ProteomeXchange with identifier PXD012906.

**Significance:** Retinal rod bipolar cells (RBCs), the second-order neurons of the mammalian rod visual pathway, are able to modulate their sensitivity to remain functional across a wide range of light intensities, from starlight to daylight. Evidence suggests that this modulation requires the serine/threonine kinase, PKCα, though the specific mechanism by which PKCα modulates RBC physiology is unknown. This study examined PKCα phosophorylation patterns in mouse rod bipolar cells and then used a phosphoproteomics approach to identify PKCα-dependent phosphoproteins in the mouse retina. A small number of retinal proteins showed significant PKCα-dependent phosphorylation, including BORG4 and TPBG, suggesting a potential contribution to PKCα-dependent modulation of RBC physiology.

**Highlights:**

- PKCα is a major source of phosphorylation in retinal RBC dendrites and its activity in RBCs is light dependent.
- Proteins showing differential phosphorylation between phorbol ester-treated wild type and PKCα-KO retinas belong to the following major functional groups: cytoskeleton/trafficking (4 proteins), ECM/adhesion (2 proteins), signaling (2 proteins), transcriptional regulation (3 proteins), and homeostasis/metabolism (1 protein).
- The PKCα-dependent phosphoproteins, BORG4 and TPBG, are present in the synaptic layers of the retina and may be involved in PKCα-dependent modulation of RBC physiology.

## 1. Introduction

Rod bipolar cells (RBCs) are key retinal interneurons in the rod visual pathway that receive light-driven synaptic input from rod photoreceptors and drive retinal output via synapses onto AII amacrine cells. RBCs serve different visual functions depending on luminance conditions. When dark adapted, they are able to transmit single photon responses [1,2], allowing for useful vision in starlight. RBCs are also able to transmit contrast changes against dim background light [3,4], and have recently been shown to influence vision in daylight [5]. Little is known about how RBCs adjust their sensitivity and gain to transition between these modes, but compelling evidence suggests phosphorylation by protein kinase C-alpha (PKCα) may play a role.

PKCα is a serine/threonine protein kinase that undergoes calcium-dependent translocation from the cytosol to the plasma membrane, where it is activated upon binding to diacylglycerol (DAG). It is a powerful modulator of signal transduction pathways and is so abundant in RBCs that it is used as a cell marker to identify RBCs with retinal immunofluorescence [6–8]. The light response of RBCs is reflected in the b-wave of the dark-adapted electroretinogram (ERG), and comparison of ERGs from wild type (WT) and PKCα knockout (PKCα-KO) mice reveal that genetic deletion of PKCα results in increases in both amplitude and duration of the scotopic b-wave [9,10]. This effect is particularly dramatic at brighter light intensities, suggesting that PKCα modulates the light response in an intensity-dependent manner. If the effect of PKCα on the light response is due to its kinase activity, then RBC proteins phosphorylated by PKCα are also likely to be involved in the modulation of RBC activity.

We used a multiplexed tandem mass tag (TMT; Thompson et al., 2003) mass spectrometry-based phosphoproteomics approach to identify proteins that were differentially phosphorylated between WT and PKCα-KO retinas in order to gain insight into the biochemical mechanisms and pathways that mediate the effect of PKCα in RBCs. Phosphopeptide abundance is expected to be dynamic, so TMT acquisition methods that have improved accuracy and wider dynamic ranges are necessary [12]. The larger number of replicates available with high-resolution instruments and TMT tags require improved data normalization and statistical testing methods, and we have successfully applied analysis techniques developed for large-scale protein expression studies [13] to phosphopeptide abundance data. Since PKCα-dependent changes in protein phosphorylation may be due to changes in total protein abundance, we also identified proteins that were differentially expressed between WT and PKCα-KO retinas.

## 2. Experimental Procedures

### Experimental design and statistical rationale

For quantification of immunofluorescence (Figure 2C), we used retina sections from 4 WT and 3 TRPM-1-KO mice, each with 4 technical replicates. For the multiplexed TMT mass spectroscopy experiments, we used 4 WT and 5 PKC*α*-KO mice. The 5 × 5 study design (one WT sample was lost) was the maximum number of samples accommodated by 10-plex TMT. For both total protein and phosphopeptide experiments, all nine samples were pooled after isobaric labeling and run simultaneously to reduce variability between samples. The statistical tests used are described within the *Statistical analysis of differential expression* section.

### Animals

All studies were approved by the Institutional Animal Care and Use Committee at Oregon Health and Science University. Wild type mice used were C57BL/6J (Jackson Laboratory; Bar Harbor, ME, USA; Cat# 000664). The PKC*α*-KO mouse strain was B6;129-Pkrca^tm1Jmk^/J (Jackson Laboratory; Cat# 009068; [10]). The TRPM1-KO mouse strain was TRPM1^tm1Lex^ (Texas A&M Institute of Genomic Medicine; College Station, TX, USA; [14]).

### PKCα activation by PMA treatment for immunoblotting or immunofluorescence

Freshly dissected retinas from WT and PKCα-KO mice were incubated at 37°C in 1 μM phorbol 12-myristate 13-acetate (PMA; Sigma-Aldrich; St. Louis, MO, USA; Cat# P8139) diluted in bicarbonate buffered Ames medium (US Biological Life Sciences; Salem, MA, USA; Cat# A1372-25) for 0 min, 15 min, 30 min, or 60 min. The retinas were then washed three times with Ames medium before being either fixed for cryo-sectioning or processed for western blotting.

### Immunoblotting

Retinas were suspended in chilled lysis buffer (50 mM Tris pH 7.4, 150 mM NaCl, 1 mM EDTA, 1% Triton X-100, 1% deoxycholate, 0.1% SDS) with 1X protease/phosphatase inhibitor cocktail (Cell Signaling Technology; Danvers, MA, USA; Cat# 5872). The retinas were then homogenized and incubated on ice for 1 hour. Lysates were centrifuged for 15 min at 28,000 × g and 4°C, and the pellets discarded. Lysates were stored at −20°C. Equal quantities of WT and PKCα-KO retinal proteins were subjected to electrophoresis on precast NuPAGE 1mm 4-12% Bis-Tris gels (Thermo Scientific; Waltham, MA, USA; Cat# NP0322BOX). The separated proteins were transferred to polyvinylidene difluoride (PVDF) membranes and then blocked with Odyssey Blocking Buffer (LI-COR Biosciences; Lincoln, NE, USA; Cat# 927-50003) for 1 hr. The blots were probed with primary antibody for 1 hr at room temperature. The membranes were washed 3 × 5 min with TBST (Tris-buffered saline with 0.1% Tween-20) and then incubated for 1 hr at room temperature with secondary antibody and washed 3 × 5 min with TBST. Immunoreactive bands were visualized with a LI-COR Odyssey CLx Imaging System at 700 or 800 nm.

Primary antibodies used for immunoblotting were rabbit anti-PKC motif phosphoserine [(R/KXpSX(R/K)] MultiMab mAb mix (1:250; Cell Signaling Technology; Cat# 6967), rabbit anti-Borg4 (1:200; Bethyl Laboratories; Montgomery, TX, USA; Cat# A302-379A), rabbit anti-5T4 (1:5000; Abcam; Cambridge, UK; Cat# ab129058), rabbit anti-NHERF1 (1:5000; Abcam; Cambridge, UK; Cat# ab3452). Secondary antibodies used were 680RD anti-rabbit (1:10,000; LI-COR Biosciences; Cat# 925-68071) and 800CW anti-mouse (1:10,000; LI-COR Biosciences; Cat# 925-32212).

### Retinal immunofluorescence

Mouse eyecups were prepared by cutting the sclera behind the ora serrata and removing the cornea and lens. Eyecups were fixed for 30 min by immersion in 4% paraformaldehyde, followed by washing in PBS. The fixed tissue was cryoprotected via sequential immersion in 10, 20, and 30% sucrose, and then embedded in Tissue-Tek O.C.T. Compound (Sakura Finetek; Torrance, CA, USA; Cat# 4583) and frozen. Sections were cut at 25 μm thickness on a cryostat and mounted onto glass slides, then air dried and stored at −20° C. Thawed retina sections were blocked at room temperature for 1 hr in Antibody Incubation Solution (AIS: 3% normal horse serum, 0.5% Triton X-100, 0.025% NaN_3_ in PBS). The sections were then incubated in primary antibody diluted in AIS for 1 hr at room temperature. After washing with 3x with PBS, the sections were incubated for 1 hr at room temperature in secondary antibody diluted 1:1000 in AIS. The slides were washed again 3x in PBS and coverslips applied with Lerner Aqua-Mount (Thermo Scientific; Cat# 13800).

Primary antibodies used for immunofluorescence were the same as used for immunoblotting unless indicated: rabbit anti-PKC motif phosphoserine [(R/KXpSX(R/K)] MultiMab mAb mix; 1:250), rabbit anti-PKCα (1:5000; Sigma-Aldrich; Cat# P4334), mouse anti-PKCα clone MC5 (1:5000; Sigma-Aldrich; Cat# P5704), rabbit anti-Borg4 (1:500), rabbit anti-5T4 (1:500;), rabbit anti-NHERF1 (1:100). Secondary antibodies used were anti-rabbit-AF488 (1:1000; Jackson ImmunoResearch Labs; West Grove, PA, USA; Cat# 11-545-144) and anti-mouse-Cy3 (1:1000 Jackson ImmunoResearch Labs; Cat# 115-165-003).

### Scanning confocal imaging

Confocal immunofluorescence images were taken with a Leica TCS SP8 X confocal microscope (Leica; Wetzlar, Germany) using a Leica HC PL APO CS2 63x/1.40 oil immersion objective (Leica; Cat#15506350) and Leica HyD hybrid detectors, or with an Olympus Fluoview 1000 microscope (Olympus; Tokyo, Japan) using a 60x/1.42 oil immersion objective. Laser lines used were AF488 (499 nm), and Cy3 (554 nm). Detection windows used were AF488 (509-544nm) and Cy3 (564-758) nm. Brightness and contrast were adjusted using Olympus Fluoview software, Leica LAS X software, or ImageJ [15,16].

### Preparation of retinas for TMT analysis

To maximize the difference between groups, retinas from wild type (n = 4) and PKCα-KO (n = 5) mice were extracted and treated for 1 hour at 37°C with 1 μM PMA diluted in bicarbonate buffered Ames medium. Following treatment, retinas were washed three times with Ames medium before being placed in chilled lysis buffer (50 mM HEPES, pH 8.5, 8 M urea, 1 mM NaF, 1 mM sodium orthovanadate, 10 mM sodium pyrophosphate, 1 mM beta-glycerophosphate), and lysed by probe sonication (Sonic Dismembrator 60; Thermo Scientific) 3 × 15 seconds at a setting of 4 with cooling on ice for 30 seconds between treatments. Protein concentrations were determined using the Pierce BCA Protein Assay Kit (Thermo Scientific; Cat# 223227) and approximately 500 µg of protein in 250 µL of lysis buffer was used for further processing. Protein disulfides were reduced with 5 µL of 1.25 M dithiothreitol at 55° C for 30 min, then alkylated by adding 22 µL of 1 M iodoacetamide and incubation at room temperature in the dark for 15 min, followed by an additional 5 µL of 1.25 M dithiothreitol. Water was added to dilute the urea concentration to 2 M. Sequence grade modified trypsin (Promega; Madison, WI, USA; Cat# V5111) was added at a 25:1 protein:trypsin ratio and samples were incubated overnight at 37° C before being acidified with trifluoroacetic acid (TFA) to a final concentration of 1%. Remaining particulates were removed by centrifugation at 16,000 × g for 10 min.

Peptides were purified by solid phase extraction using 1 cc (50 mg) Waters Sep-Pak Vac tC18 cartridges (Waters Corporation; Milford, MA, USA; Cat# WAT054960). Briefly, the cartridges were conditioned twice with 1 mL acetonitrile (ACN) and twice with 300 µL of 50% ACN/0.5% acetic acid, then equilibrated twice with 1 mL 0.1% TFA. The samples were loaded and passed through the bed, then were washed twice with 1 mL 0.1% TFA followed by 200 µL of 0.5% acetic acid. Finally, the samples were eluted twice with 500 µL of 50% ACN/0.5% acetic acid. Peptide concentrations were determined using the Pierce Quantitative Colorimetric Peptide Assay (Thermo Scientific; Cat# 23275). Approximately 300 µg of peptide was recovered from each digest. Fifteen µg of each sample was reserved for TMT analysis of total protein abundance, and the remainder was used for the phosphopeptide enrichment experiment.

### Phosphopeptide enrichment

Phosphopeptides were enriched following previously described methods [17,18] with small modifications. Titanosphere TiO_2_ 5 µm beads (GL Biosciences; Tokyo, Japan; Cat# 5020-75000) were washed three times in 2 M lactic acid/50% ACN, then resuspended in the same solution at 24 mg/mL. Approximately 285 µg of dried peptide for each sample was resuspended in 950 µL of 2 M lactic acid/50% ACN and 100 µL (2.4 mg) of the bead suspension was added to each sample, ensuring an 8:1 ratio of beads:peptide. The bead:peptide mixture was rotated at room temperature for 1 hour, then washed twice with 500 µL 2M lactic acid/50% ACN, 0.1% TFA/50% ACN, then 0.1% TFA/25% ACN. The enriched phosphopeptides were eluted from the beads by vortexing in 100 µL of 50 mM K_2_HPO_4_ at pH 10 for 5 min. The elution step was repeated once and the 200 µL of eluate was dried by vacuum centrifugation.

The enriched phosphopeptides were then purified by solid phase extraction using UltraMicro Spin columns (The Nest Group Inc.; Southborough, MA, USA). The dried phosphopeptides were resuspended in 60 µL 1% TFA and the pH was tested to ensure the samples were acidic. The columns were conditioned three times with 100 µL 80% ACN/0.1% TFA and equilibrated three times with 50 µL of 0.1% TFA. The samples were loaded and passed through the columns three times, washed three times with 25 µL of 0.1% TFA, eluted three times with 50 µL 80% ACN/0.1% formic acid, and dried by vacuum centrifugation before TMT labeling.

### TMT labeling and mass spectrometric analysis

In preparation for TMT labeling, nine dried unfractionated peptide samples (4 WT and 5 KO) and nine phosphopeptide enriched samples (4 WT and 5 KO) were dissolved in 25 µL of 100 mM triethylammonium bicarbonate buffer, and TMT 10-plex reagents (Thermo Scientific; Cat# 90110) were dissolved at a concentration of 15 µg/µL in anhydrous ACN. Each of the samples was then labeled by adding 12 µL (180 µg) of an individual TMT reagent, followed by shaking at room temperature for 1 hr. Two µL of each of the nine labeled samples in each group were pooled, and 2 µL of 5% hydroxylamine was added. The samples were incubated for 15 min, dried by vacuum centrifugation, dissolved in 21 µL of 5% formic acid, and peptides were analyzed by a single 2-hour LC-MS/MS method using an Orbitrap Fusion as described below. The run was performed to normalize the total reporter ion intensity of each multiplexed sample and to check labeling efficiency. After the normalization and efficiency run, the remaining unmixed samples were then quenched by the addition of 2 µL 5% hydroxylamine, then combined in adjusted volumes to yield equal summed reporter ion intensities during the subsequent two-dimensional LC/MS.

Following volume-based normalization, the combined samples were dried by vacuum centrifugation, and TMT-labeled samples were reconstituted in 20 µL water. The reconstituted peptides were separated by two-dimensional nano reverse-phase liquid chromatography (Dionex NCS-3500 UltiMate RSLCnano UPLC) EasySpray NanoSource (Thermo Scientific), ionized using an EasySpray NanoSource (Thermo Scientific), and SPS MS3 data acquired with an Orbitrap Fusion Tribrid mass spectrometer (Thermo Scientific). The liquid chromatography details and mass spectrometer settings were as previously described [13] with the modification that non-enriched peptides were eluted from the first dimension high pH column using sequential injections of 20 µL volumes of 14, 17, 20, 21, 22, 23, 24, 25, 26, 27, 28, 29, 30, 35, 40, 50, and 90% ACN in 10 mM ammonium formate, pH 9, and enriched phosphopeptides were eluted by sequential 20 µL injections of 4, 6, 8, 10, 12, 18, 20, 22, 25, 30, and 60% ACN in 10 mM ammonium formate, pH 9.

### TMT data analysis

The binary instrument files were processed with Proteome Discoverer version 1.4 (Thermo Scientific) to extract fragment ion spectra, precursor information (m/z values and peptide charge state), and TMT reporter ion peak intensities. The fragment ion spectra were searched against a canonical Uniprot Swiss-Prot mouse protein database (downloaded 07/07/2016 from https://www.uniprot.org) with 16,794 entries. There were 179 common contaminant entries appended for a total of 16,973 sequences.

The SEQUEST [19] search engine was used with a parent ion mass tolerance of 1.25 Da and fragment ion tolerance of 1.0005 Da. Trypsin cleavage with up to two missed cleavages was specified. A static modification of +57.0215 Da was applied to all cysteine residues and a variable modification of +15.9949 was applied to methionine. Confident peptide-to-spectrum matches (PSMs) were obtained using the percolator [20] node, and only peptides with q values < 0.05 were accepted. Parsimonious protein inference was used in Proteome Discoverer to produce final protein lists, and the results were exported to tab-delimited files for post processing using in-house Python scripts (https://www.github.com/pwilmart/PAW_pipeline.git). For the protein expression analysis of the total protein preparations, the reporter ion intensities from all unique (matching to just one protein) peptides were summed into protein reporter ion intensities. Any contaminant proteins were excluded from further analysis. Reporter ion data from PSMs where the trimmed average reporter ion intensity did not exceed 500 were excluded. Any final protein reporter ion sums of zero were replaced with a value of 150 (the smallest non-zero reporter ion intensities observed were approximately 350) to avoid mathematical errors during visualizations and statistical testing.

The data from the phosphopeptide enrichment experiment was searched with additional variable modifications of +79.9799 on serine, threonine, or tyrosine residues, and only peptides with q < 0.01 were accepted. The phosphorylation site localization node phosphoRS [21] was configured after the search node in Proteome Discoverer. Phosphorylation enrichment experiments are peptide centric, so data aggregation was done differently than for protein expression. Peptide sequences were aggregated by summing reporter ion intensities within each channel to reduce variance and increase statistical power. All PSMs assigned to the same base peptide sequence were aggregated in the same modification state, which was determined by integral peptide MH+ mass (the peptide in a 1+ charge state). Localization information from phosphoRS was simplified to the same number of top probabilities as the number of phosphorylation modifications present in the peptides. The reporter ion intensities from the combined reports were used for differential expression (DE) testing as described below. Any contaminant protein matches were excluded from further analysis. Minimum intensity filtering and zero replacement was done similarly to the total protein analysis.

### Statistical analysis of differential expression

For immunofluorescence quantification, data are represented in text as the mean ± SEM and p-values were calculated using an unpaired t-test.

For TMT data, the table of non-contaminant reporter ion intensities for proteins or for aggregated phosphopeptides were exported to tab-delimited files and imported into R (https://www.r-project.org) for statistical analysis using the edgeR [22–24] Bioconductor package. Normalization was done using a “library size” factor and the trimmed mean of M-values normalization [25] function in edgeR was used to correct for sample loading and compositional bias. DE testing was performed pairwise using the exact test with Benjamini-Hochberg multiple testing corrections [26]. The results from edgeR analyses were exported and added to the combined data summaries. Annotation information was added from Uniprot using a script available at https://github.com/pwilmart/annotations.

### Data availability

The mass spectrometry proteomics data have been deposited to the ProteomeXchange Consortium via the PRIDE partner repository [27], with the dataset identifier PXD012906.

## 3. Results

### RBC dendrites are major sites of light-dependent PKCα phosphorylation

To visualize sites of PKCα phosphorylation in the retina, we used a monoclonal antibody mix that binds to canonical PKC substrate motifs containing a phosphorylated serine (PKC motif p-serine). Anti-PKC motif phosphoserine immunofluorescence between wild type and PKCα-KO retina sections indicates that RBC dendrites are the main sites of PKCα phosphorylation in the mouse retina (Figure 1A and B). The small immunofluorescent puncta in the outer plexiform layer (OPL) are presumptive RBC dendrites, whereas the larger patches of phosphoserine immunofluorescence (arrows) are associated with cone pedicles. The small immunofluorescent puncta are visible throughout the wild type OPL but are greatly reduced in intensity in the PKCα-KO retina, while immunofluorescence corresponding to cone pedicles is unchanged. These results suggest that PKCα phosphorylates targets in RBC dendrites of wild type adult mice, and that a different PKC isoform phosphorylates targets associated with cone pedicles.

To examine whether PKCα activity is light-dependent, we used a conformation-dependent monoclonal PKCα antibody which binds an epitope in the hinge region of PKCα that is inaccessible in the inactive state [28]. Retina sections from light-adapted and dark-adapted mice were double-labeled with the conformation-specific antibody (anti-PKCα-A) and a non-conformation-specific PKCα antibody (anti-PKCα-B; Figure 2). Both antibodies strongly label RBC cell bodies and dendrites in sections from light-adapted retina, with co-localization of the two secondary antibodies appearing white (Figure 2A, left). By contrast, only anti-PKCα-B labels RBCs in sections from dark adapted retina (Figure 2A, right). These results suggest that PKCα is active in RBC dendrites in the light-adapted state. This was supported by labeling of light- and dark-adapted retina sections for phosphorylated PKC motifs. The anti-PKC motif phosphoserine antibody labeled puncta in the OPL of light-adapted retina (Figure 2B, left), similar to the wild type immunofluorescence seen in Figure 1. Anti-PKC motif phosphoserine immunofluorescence was absent in the OPL of the dark-adapted retina (Figure 2B, right). Together, the results from Figures 1 and 2 indicate that PKCα is active in RBC dendrites in the light-adapted state.

**Figure 1:**
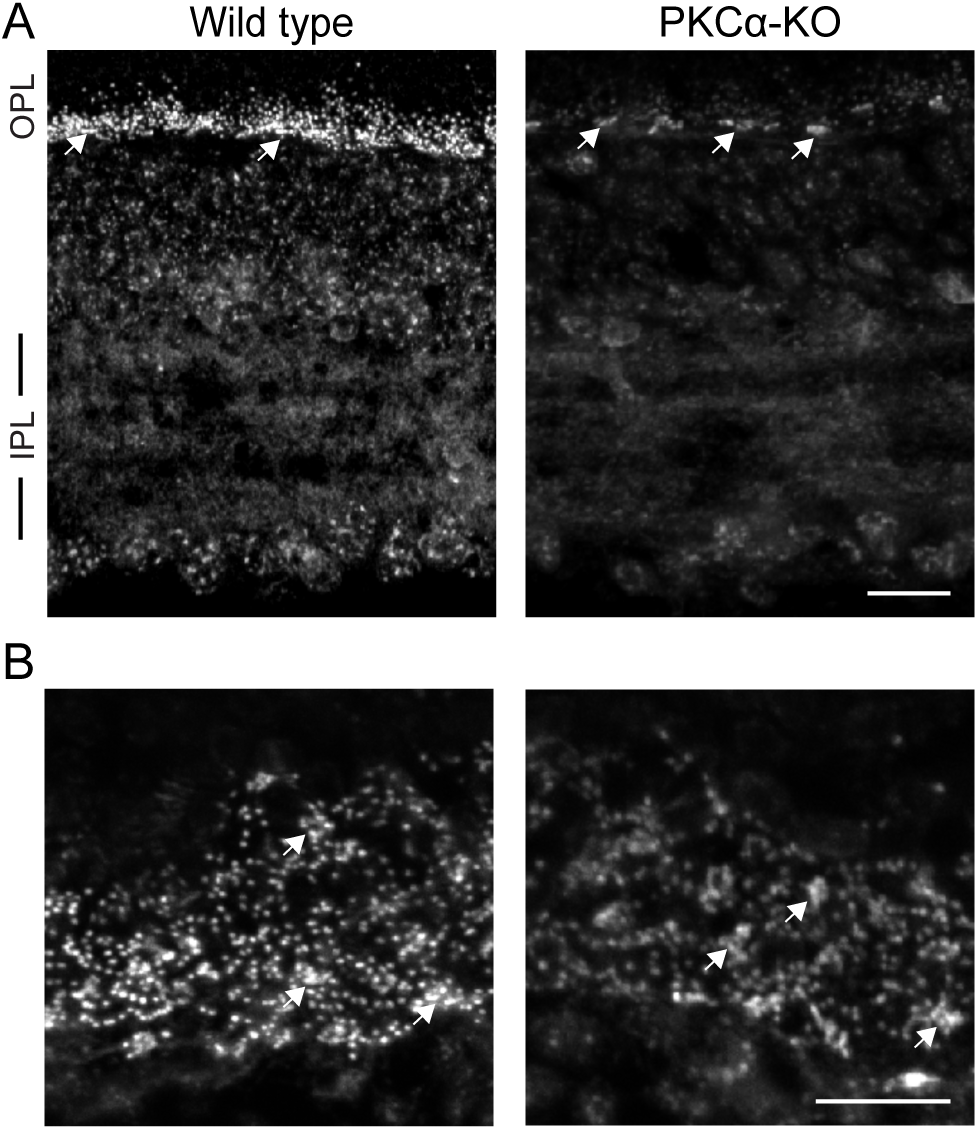
Phosphoserine labeling in the OPL is reduced in PKCα-KO retina. (A) Immunofluorescence confocal images of mouse retina sections from wild type and PKCα-KO retinas labeled with an antibody against phosphoserine residues within canonical PKC motif phosphoserine (PKC motif p-serine). (B) Images of PKC motif p-serine immunofluorescence in the outer plexiform layer of wild type and PKCα knockout (KO) retinas in obliquely cut sections. In both A and B, white arrows indicate labeling associated with presumptive cone pedicles. Scale bars: 20 μm. OPL: outer plexiform layer; IPL: inner plexiform layer.

**Figure 2:**
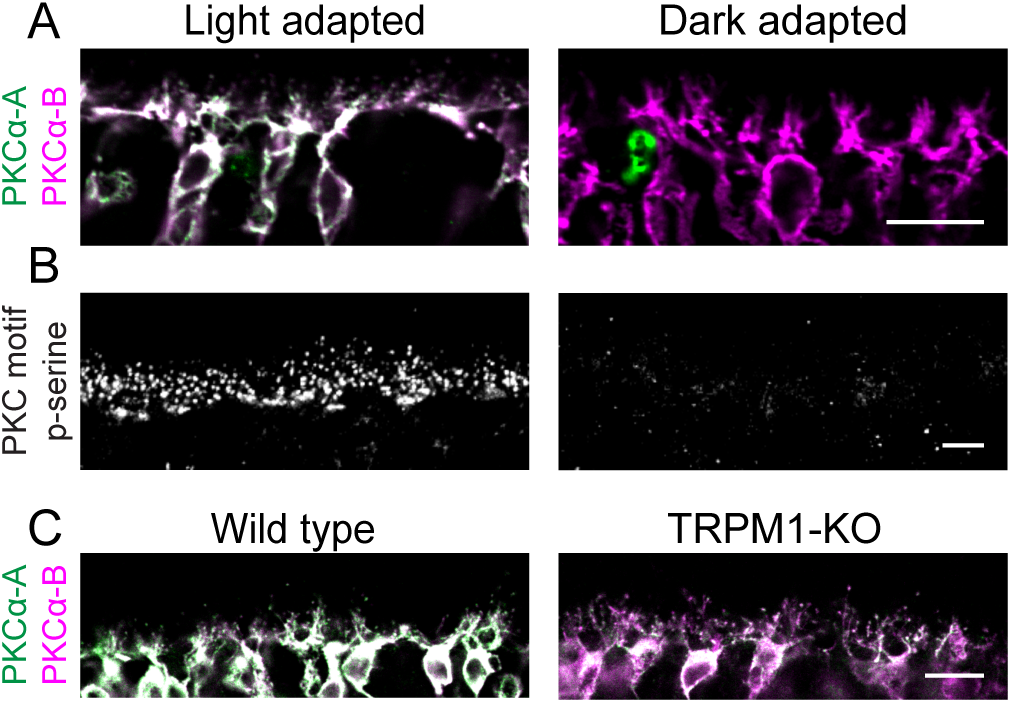
PKCα is active in light-adapted RBCs. (A) Light- and dark-adapted mouse retina sections double-labeled with two antibodies against PKCα; conformation-specific PKCα-A (green) binds only active PKCα, while PKCα-B (magenta) binds to both active and inactive PKCα. (B) Sections from light-adapted and dark-adapted mouse retina labeled with an antibody mixture against PKC motif phosphoserines (PKC motif p-serine). (C) WT and TRPM1-KO mouse retina sections double-labeled with conformation-specific PKCα-A (green) and conformation non-specific PKCα-B (magenta). Scale bars: 10 μm. OPL: outer plexiform layer.

Conventional PKC isoforms, including PKCα, require calcium for activation [29]. In RBC dendrites, a likely source of calcium is the Transient Receptor Potential cation channel subfamily M 1 (TRPM1) cation channel, which mediates an influx of sodium and calcium to generate the RBC light response [14,30,31]. The dependence of PKCα activation on TRPM1 was assessed by double-labeling retina sections from wild type and TRPM1 knockout (TRPM1-KO) mice with anti-PKCα-A and anti-PKCα-B to detect active and total PKCα, respectively, in RBC cell bodies and dendrites (Figure 2C), and the intensities of the immunofluorescence obtained with the two antibodies was compared. The average ratio of anti-PKCα-A (active) to anti-PKCα-B (total) immunofluorescence in wild type RBCs was 1.12 ± 0.12 (n = 4 mice, each with 4 technical replicates) compared to 0.70 ± 0.15 (n = 3 mice, each with 4 technical replicates) in TRPM1-KO RBCs (P < 0.0001), indicating that PKCα is less active in RBCs in the absence of TRPM1.

PKCα is a DAG-sensitive PKC isoform, and therefore PKCα phosphorylation can be potentiated by the DAG analogue phorbol 12-myristate 13-acetate (PMA). To confirm that PKCα is phosphorylating proteins in the retina *ex vivo*, we incubated freshly dissected wild type and PKCα-KO retinas in PMA before analyzing changes in protein phosphorylation by immunoblotting for PKC motif phosphoserines (PKC motif p-serine). PMA activation of other DAG-sensitive PKC isoforms expressed in the retina, such as PKCβ, should be relatively equivalent between WT and PKCα-KO samples and result in no difference in phosphorylation. PMA treatment resulted in a significant increase in intensity of several phosphoserine immunoreactive bands in the wild type samples at the 15, 30, and 60 min time-points compared to the PKCα-KO samples (Figure 3A) demonstrating that PMA treatment increases differential phosphorylation between WT and PKCα-KO retinas. To identify sites of PMA-activated PKC phosphorylation in the wild type retina, PKC motif phosphoserine immunofluorescent labeling was performed on retina sections made from PMA-treated, light-adapted WT retinas. PMA incubation resulted in an increase in PKC motif phosphoserine immunofluorescence in presumptive RBC dendritic tips (Figure 3B).

**Figure 3:**
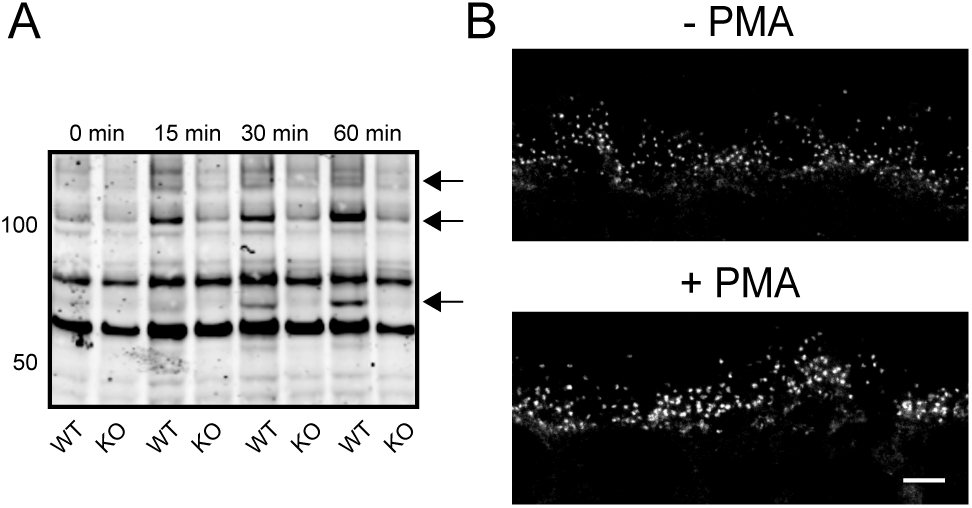
PMA increases phosphorylation by PKC isoforms in the mouse retina. (A) Wild type (WT) and PKCα knockout (PKCα-KO) retinas were incubated in PMA for 0, 15, 30, and 60 min followed by western blotting with an antibody against PKC motif phosphoserines. Arrows indicate candidate PKCα phosphorylation targets. (B) Immunofluorescent PKC motif phosphoserine labeling of wild type mouse OPL from retinas that were incubated with and without PMA for 1 hr. Scale bar: 10 μm. PMA: phorbol 12-myristate 13-acetate; OPL: outer plexiform layer.

### Differential protein abundance in wild type and PKCα-KO retina

To identify retinal proteins whose expression is dependent on PKCα, four wild type and five PKCα-KO retinas were incubated with PMA for an hour immediately after dissection and then processed for multiplexed TMT mass spectroscopy (Figure 4). Retinas were lysed and proteins digested with trypsin before TMT labeling and LCMS/MS. Peptide identification was performed with Proteome Discoverer using SEQUEST and Percolator. From 38,384 confidently identified PSMs, there were 34,969 unique peptides corresponding to 4,435 proteins (excluding contaminants; Figure 5, S1 – Total Protein and Phosphopeptide Abundance Analysis). Differential expression (DE) statistical testing was done with edgeR using the trimmed mean of M-values normalization and an exact pairwise test. After Benjamini-Hochberg p-value corrections for multiple comparisons to establish DE false discovery rates (FDRs), we grouped proteins with significant differential expression into significance groups based on FDR thresholds: not significant (FDR > 0.10 [4412 proteins]) low significance (0.10 > FDR > 0.05 [2 proteins]), medium significance (0.05 > FDR > 0.01 [5 proteins]), and high significance (FDR < 0.01 [16 proteins]).

**Figure 4:**
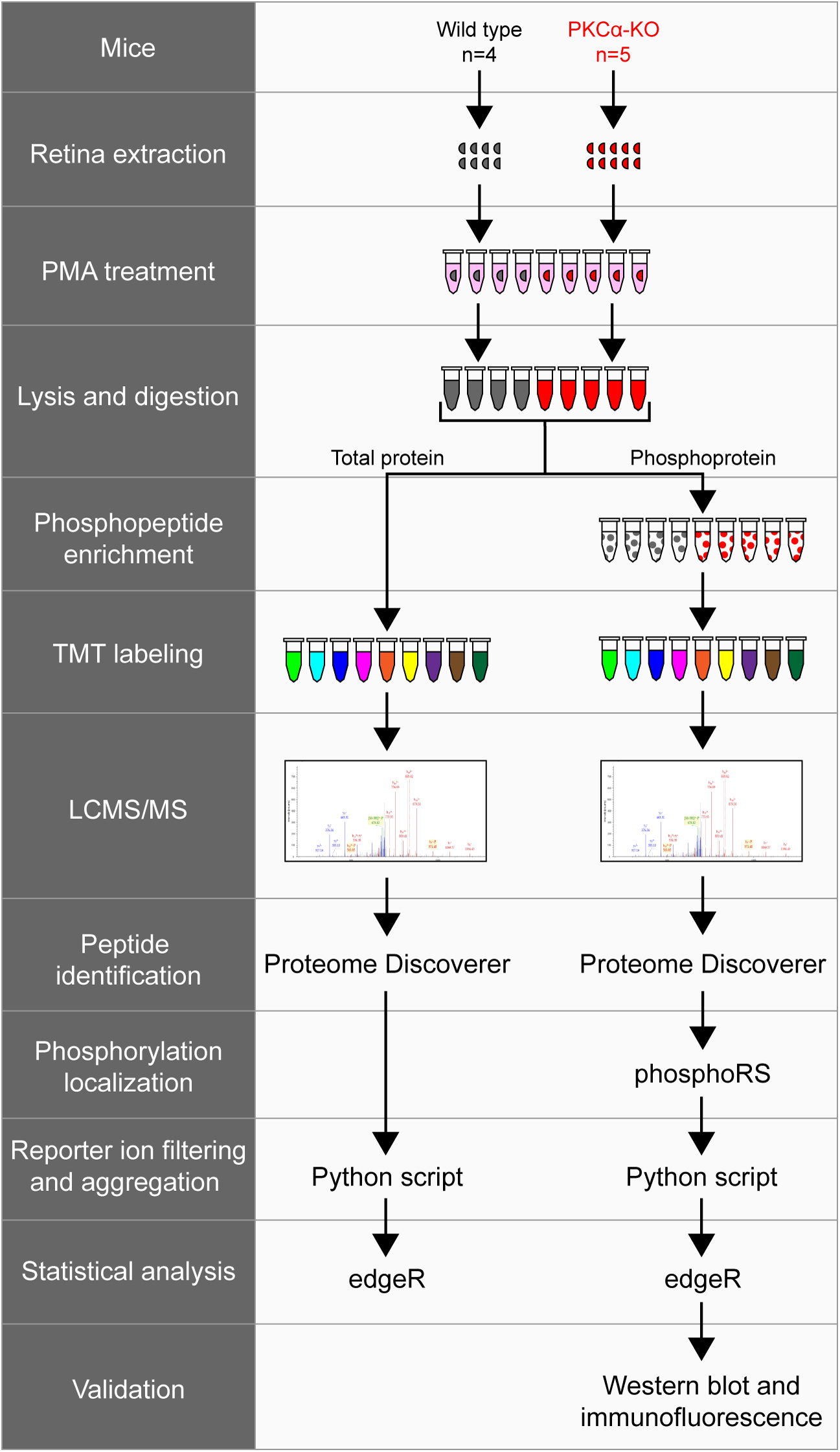
Experimental workflow of total protein and phosphopeptide identification. Wild type (n=4) and PKCα knockout (KO) (n=5) retinas were extracted and treated with PMA before lysis and trypsin digestion. A small fraction of each sample was removed for total protein analysis, while the rest of the samples underwent phosphopeptide enrichment. Following TMT labeling, samples were combined and analyzed by LCMS/MS. Tandem mass spectrometry data was collected on an Orbitrap Fusion and proteins were identified using Proteome Discoverer (SEQUEST and Percolator). Phosphorylation site localization was scored using phosphoRS, and reporter ion intensities were filtered and aggregated with an in-house Python script. TMT reporter ion intensities from total proteins or from phosphopeptides were tested for differential expression using the Bioconductor package edgeR. The presence of representative phosphoproteins was validated in the retina by western blot and confocal immunofluorescence microscopy. PMA: phorbol 12-myristate 13-acetate.

**Figure 5:**
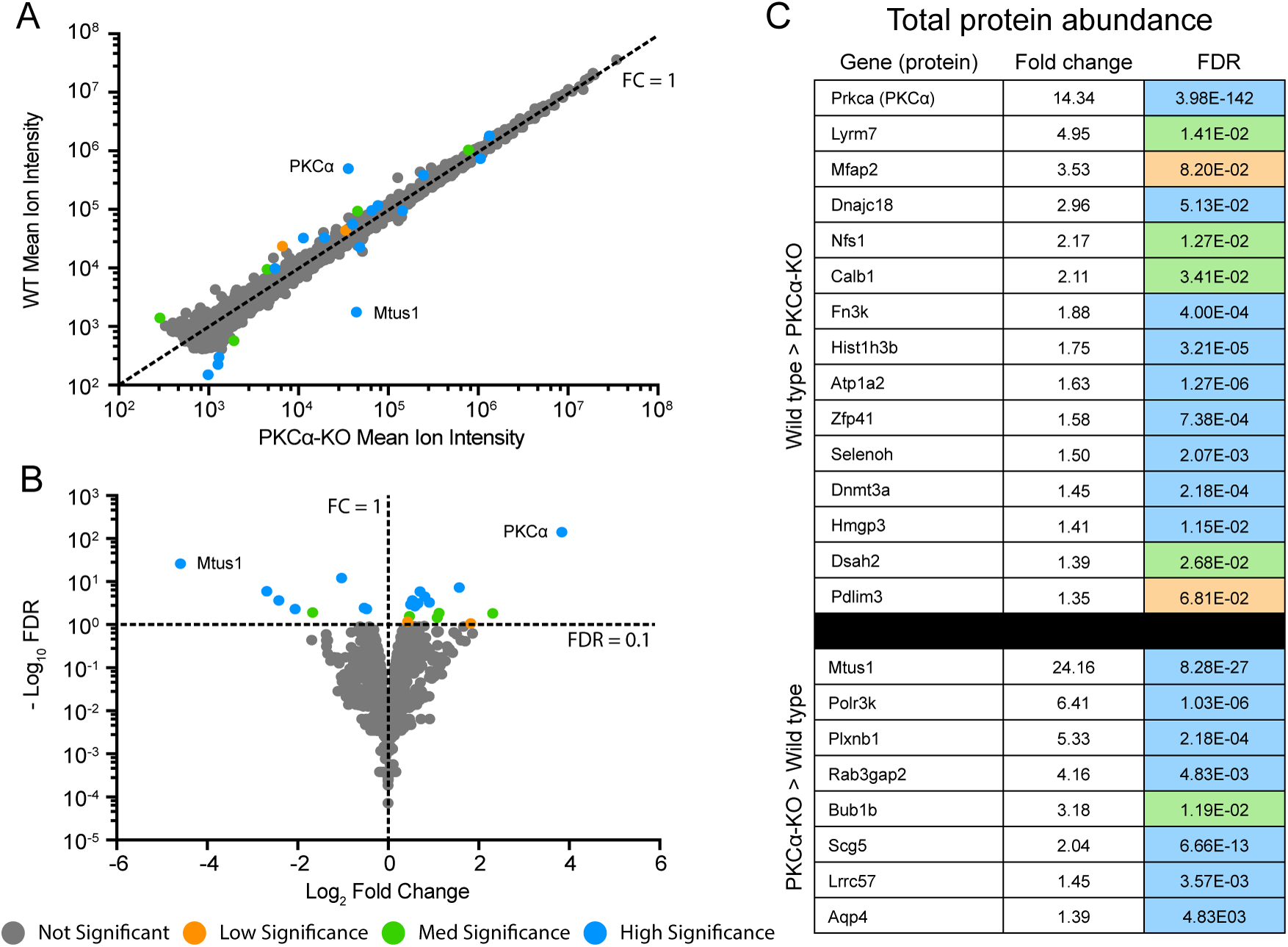
Identification of differentially expressed total proteins. (A) Scatter plot of peak reporter ion intensities from wild type (WT) and PKCα knockout (KO) total protein abundance samples. The dotted line corresponds to an FC (WT / KO) of 1, with FC > 1 corresponding to increased abundance in WT vs KO, and FC < 1 corresponding to increased abundance in KO vs WT. (B) Volcano plot of log_2_ FC and –log_10_ FDR. The dotted lines correspond to FC = 1 and FDR = 0.1. FDR < 0.1 corresponds to proteins passing the low significance threshold, and FDR > 0.1 corresponds to proteins failing the low significance threshold. The low significance (orange) threshold was 0.1, the med significance threshold (green) was 0.05, and the high significance threshold (blue) was 0.01. (C) Table of all differentially abundant proteins with an FDR lower than the low significance threshold of 0.1. Fold change was calculated by dividing the mean reporter ion intensities of each protein from the genotype with higher abundance by that with lower abundance. Colors correspond to DE FDR thresholds. DE: differential expression; FC: fold change; FDR: false discovery rate.

Plotting the mean reporter ion intensities between wild type and PKCα-KO samples (Figure 5A) revealed a small number of significantly different protein abundances in both WT (15 proteins) and KO (8 proteins). The proteins with the largest changes in abundance were KPCA (PKCα; 14-fold decrease in KO), and MTUS1 (Microtubule Associated Scaffold Protein 1; 24-fold increase in KO). A volcano plot comparing log_2_ fold change (WT / KO) with –log_10_ FDR (Figure 5B) depicts the 23 proteins passing the FDR < 0.1 cutoff for low significance. We did not attempt any isotopic corrections for reporter ions, so the degree of downregulation for PKCα is consistent with an absence of the kinase in the knockout. A list of all significant differentially expressed proteins can be seen in Figure 5C. The sample-to-sample reproducibility of the protein abundance experiment was excellent: the WT samples had a median coefficient of variance (CV) of 11.4%, the KO samples had a median CV of 15.5%, and the independent CV was 13.5%. Ninety-five percent of the expression changes were less than 1.25-fold different (S2 – Total Protein and Phosphopeptide Statistical Testing).

### Differential protein phosphorylation between wild type and PKCα-KO retina

To examine differences in protein phosphorylation between wild type and PKCα-KO retinas, phosphopeptides were enriched from the same retinal extracts used above (described in the Methods), and phosphopeptide abundance was analyzed by multiplexed TMT mass spectroscopy. Peptide identification was performed with Proteome Discoverer using SEQUEST and Percolator, and phosphorylation localization was performed using phosphoRS. Results files were exported for post processing (filtering and aggregation) using an in-house Python script (PD1.4 TMT phospho processer.py available at https://github.com/pwilmart/PAW pipeline). The aggregated phosphopeptide reporter ion data was tested for differential expression using edgeR with multiple testing corrections, and phosphopeptides were grouped into significance groups based on differential expression FDR as described for total proteins.

We identified 1137 distinct phosphopeptides in wild type or PKCα-KO retina lysates (Figure 6, S1 – Total Protein and Phosphopeptide Abundance Analysis) 1113 were not significant (FDR > 0.10), seven showed low significance (0.10 > FDR > 0.05), four medium significance (0.05 > FDR > 0.01), and thirteen were highly significant (FDR < 0.01). Of the 24 significant phosphoproteins, only two showed differential expression in the total protein abundance analysis: Dnmt3a (DNA (Cytosine-5)-Methyltransferase 3A), which was significantly more abundant in the WT samples (1.5fold), and Scg5 (Neuroendrocrine Protein 7B2), which was significantly more abundant in the KO samples (2.04-fold). Plotting WT and KO peak ion intensities (Figure 6A) highlights five phosphopeptides with much greater expression in WT than KO. A volcano plot comparing log_2_ FC (WT / KO) with –log_10_ FDR (Figure 6B) shows the 24 phosphopeptides passing the FDR < 0.1 threshold (14 increased in WT and 10 increased in KO). A full list of all significant differentially expressed phosphopeptides, along with their total protein abundance changes, can be seen in Figure 6C. The reproducibility of the peptide-centric experiment was strong: the WT samples had a median CV of 13.9%, the KO samples had a median CV of 18.7%, and the median CV independent of condition was 16.2%. Ninety-five percent of the expression changes were less than 1.4-fold different (S2 – Total Protein and Phosphopeptide Statistical Testing). Annotated fragment ion spectra for the phosphorylated peptides are shown in S3 – Annotated Phosphopeptide Spectra.

**Figure 6:**
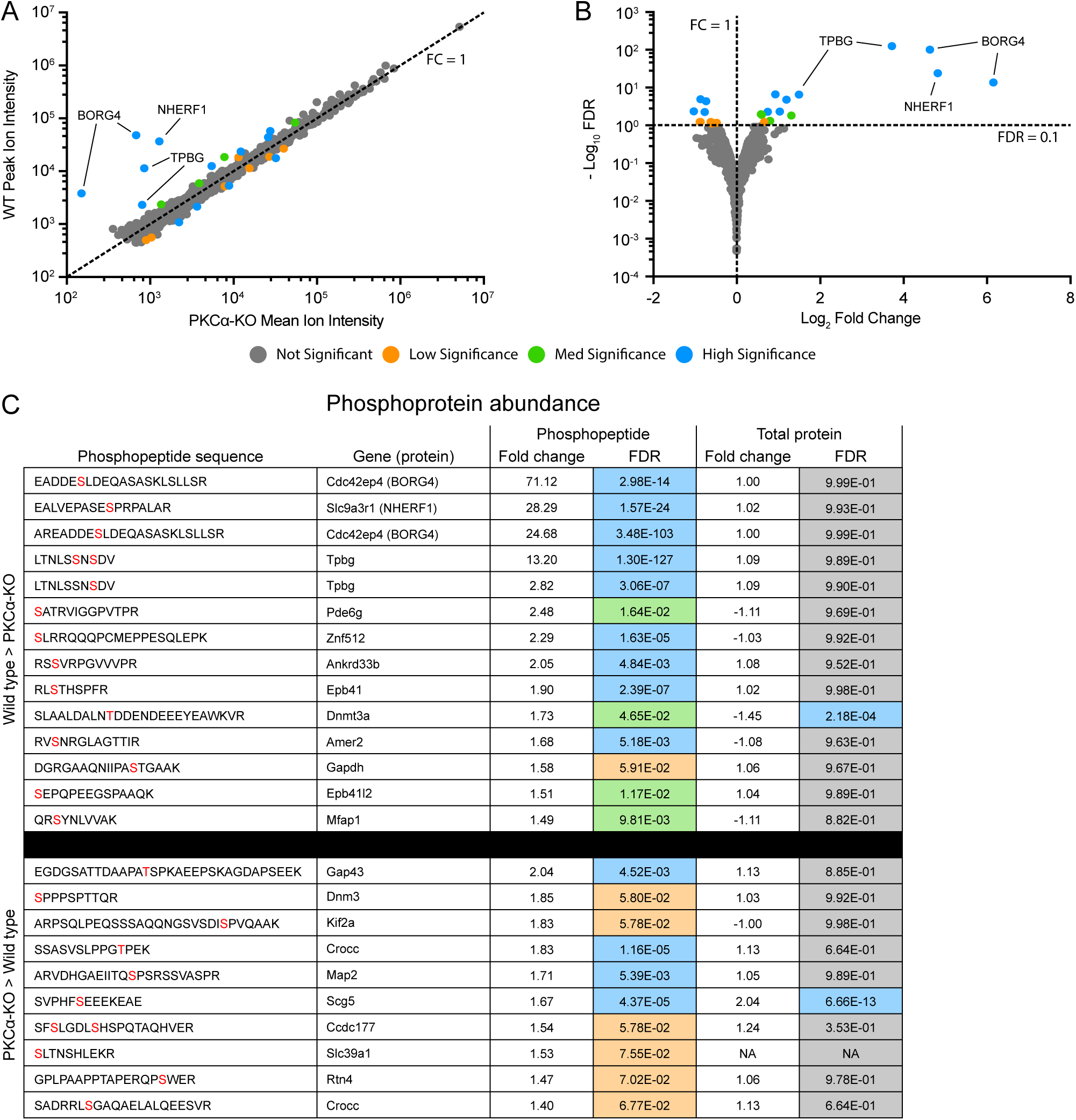
Identification of differentially expressed phosphopeptides. (A) Scatter plot of peak reporter ion intensities from wild type (WT) and PKCα knockout (KO) phosphopeptide abundance samples. The dotted line corresponds to an FC (WT / KO) of 1, with FC > 1 corresponding to increased abundance in WT vs KO, and FC < 1 corresponding to increased abundance in KO vs WT. (B) Volcano plot of log_2_ FC and –log_10_ FDR. The dotted lines correspond to FC = 1 and FDR = 0.1. FDR < 0.1 corresponds to proteins passing the low significance threshold, and FDR > 0.1 corresponds to proteins failing the low significance threshold. The low significance (orange) threshold was 0.1, the med significance threshold (green) was 0.05, and the high significance threshold (blue) was 0.01. (C) Table of all differentially abundance phosphopeptides with an FDR lower than the low significance threshold of 0.1. In the phosphopeptide sequence column, phosphorylated residues are in red. In the Total Protein Fold Change column, negative values indicate an increased abundance in the KO samples. Fold change was calculated by dividing the mean reporter ion intensities of each protein from the genotype with higher abundance by that with lower abundance. Fold change and DE FDR values were taken from the phosphopeptide abundance experiment and the total protein abundance experiment. Colors correspond to DE FDR thresholds. DE: differential expression; FC: fold change; FDR: false discovery rate.

Five phosphopeptides belonging to three proteins showed the largest and most significant differential abundance between the wild type and PKCα-KO samples (Figure 7). Two similar phosphopeptides were identified corresponding to BORG4 (Binder of Rho-GTPase, also called CDC42EP4) with phosphoserine residues observed at the peptide sites S6 (71-fold increase) and S8 (25-fold increase). Both peptides were generated by slightly different cleavage patterns of the same amino acid sequence with the same phosphorylated serine residue corresponding to site S64 in full-length BORG4 (Figure 7A). One phosphopeptide fragment was identified from NHERF1 (Na^+^/H^+^ Exchanger Regulatory Factor 1, also called SLC9A3R1 and EBP50) with a phosphorylated serine residue at the peptide S10 position (28-fold increase) corresponding to S275 in the full-length protein (Figures 7B). The last two major significant phosphopeptides correspond to the same 10 amino acid sequence of TPBG (Trophoblast Glycoprotein, also called the 5T4 antigen and WAIF1) with two identified species: a doubly-phosphorylated peptide with phosphoserines at peptide S6 and S8 (13-fold increase), and a singly-phosphorylated peptide with just phosphorylated peptide S8 (2.8-fold increase). These two serine residues correspond to S422 and S424 in the C-terminal intracellular tail of TPBG (Figure 7C).

**Figure 7:**
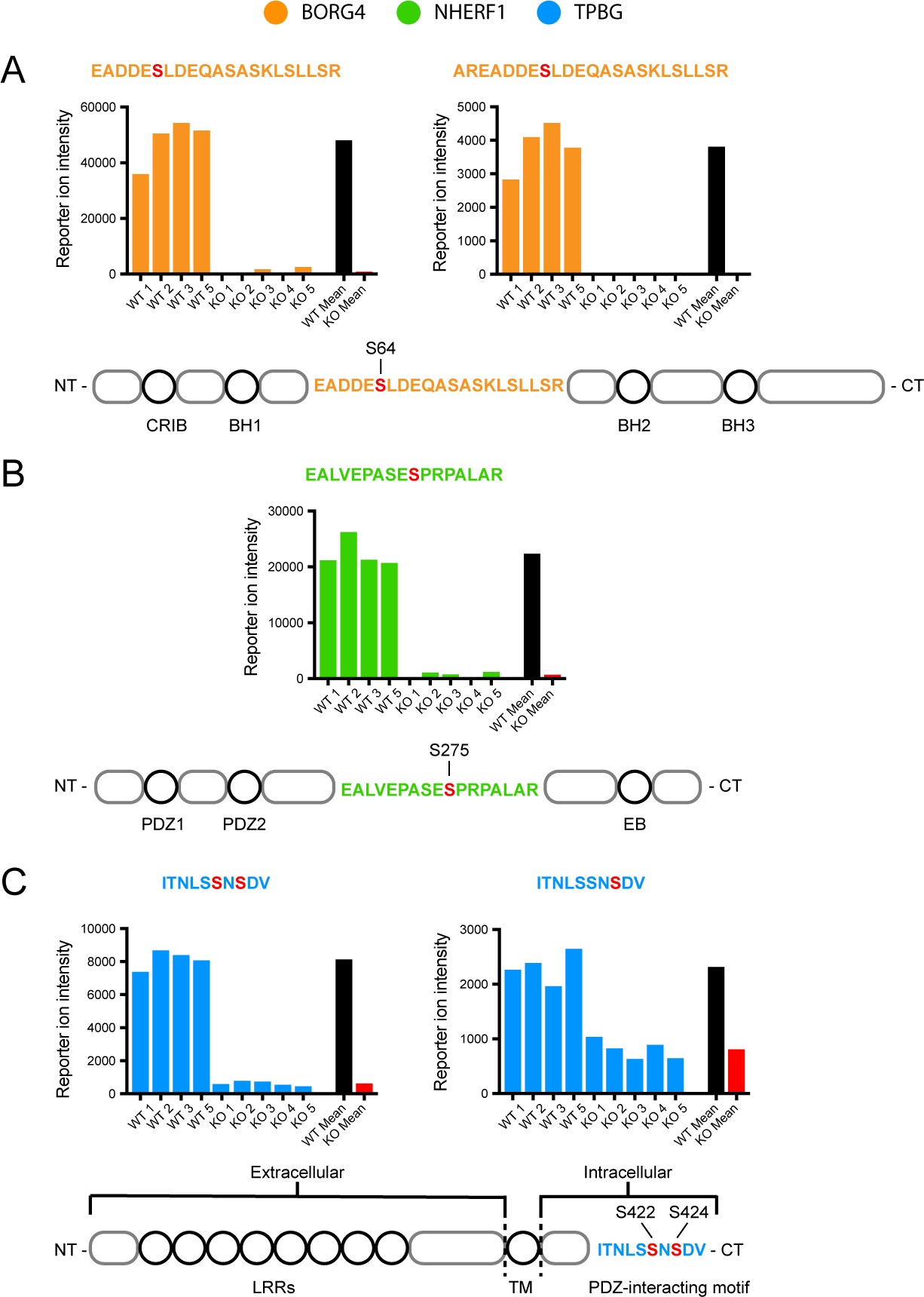
TMT data from representative phosphoproteins. Reporter ion intensity values from each TMT channel for the five phosphopeptide fragments with the largest differential expression between WT (n = 4) and PKCα-KO (n = 5): two from BORG4 (orange), one from NHERF1 (green), and two from TPBG (blue). For statistical significance of differential expression analysis of mean WT (black) and mean KO (red) reporter ion intensities, see S1 – Total Protein and Phosphopeptide Abundance Analysis. (A) Two phosphopeptide fragments from an overlapping region of BORG4, each containing a phosphorylated serine corresponding to S64 on the full-length protein. (B) One phosphopeptide fragment from NHERF1 with a phosphorylated serine corresponding to S275 on the full-length protein. (C) Two phosphopeptide fragments from the C-terminal tail of TPBG with two similar phosphorylation patterns: one with two phosphoserines corresponding to S422 and S424, and one with a single phosphoserine corresponding to S424 of the full-length protein.

### Grouping of significant protein and phosphoprotein results by biological function

Proteins identified in the total protein abundance (Figure 8A) and phosphoprotein abundance (Figure 8B) experiments were grouped into broad categories based on general biological function annotations added from Uniprot (https://github.com/pwilmart/annotations.git). PKCα-KO resulted in differential expression of proteins and phosphoproteins involved in many aspects of cellular physiology, particularly cytoskeletal rearrangement and vesicle trafficking (15 proteins), transcriptional regulation (8 proteins), and homeostasis and metabolism (5 proteins).

**Figure 8:**
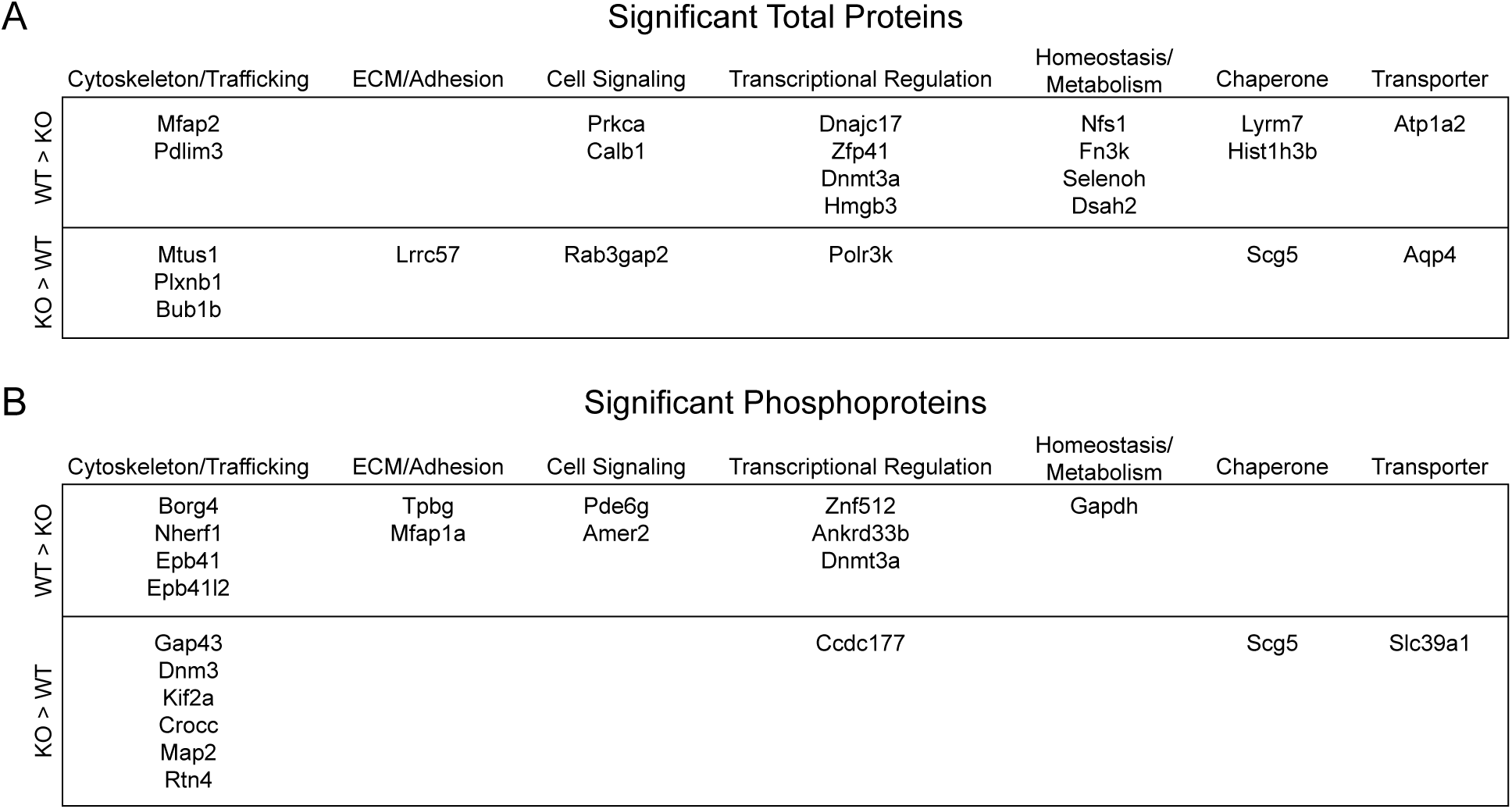
Significant total proteins and phosphoproteins grouped by biological function. Table of genes of identified total proteins (A) and phosphoproteins (B) with significant differential abundance between wild type and PKCα-KO samples grouped into broad categories based on general biological function gathered from Uniprot protein annotations.

### Localization of the major PKCα-dependent phosphoproteins in the mouse retina

We used immunoblotting and immunofluorescence confocal microscopy to examine the presence of the three most prominent PKCα-dependent phosphoprotein hits in the wild type retina: BORG4, NHERF1, and TPBG. Immunoblotting for BORG4 shows a distinct band at 38 kDa (Figure 9A) in agreement with the predicted molecular weight of BORG4. Immunofluorescence double-labeling of retina sections for BORG4 and PKCα shows punctate BORG4 labeling in the outer plexiform layer (OPL), but the BORG4 puncta are not strongly co-localized with PKCα (Figure 9B). This is consistent with labeling of either RBC or horizontal cell dendritic tips, as they are closely apposed to one another within the rod spherule invagination [32,33]. BORG4 immunofluorescence was also detected in nuclei in the inner nuclear layer (INL) and ganglion cell layer.

**Figure 9:**
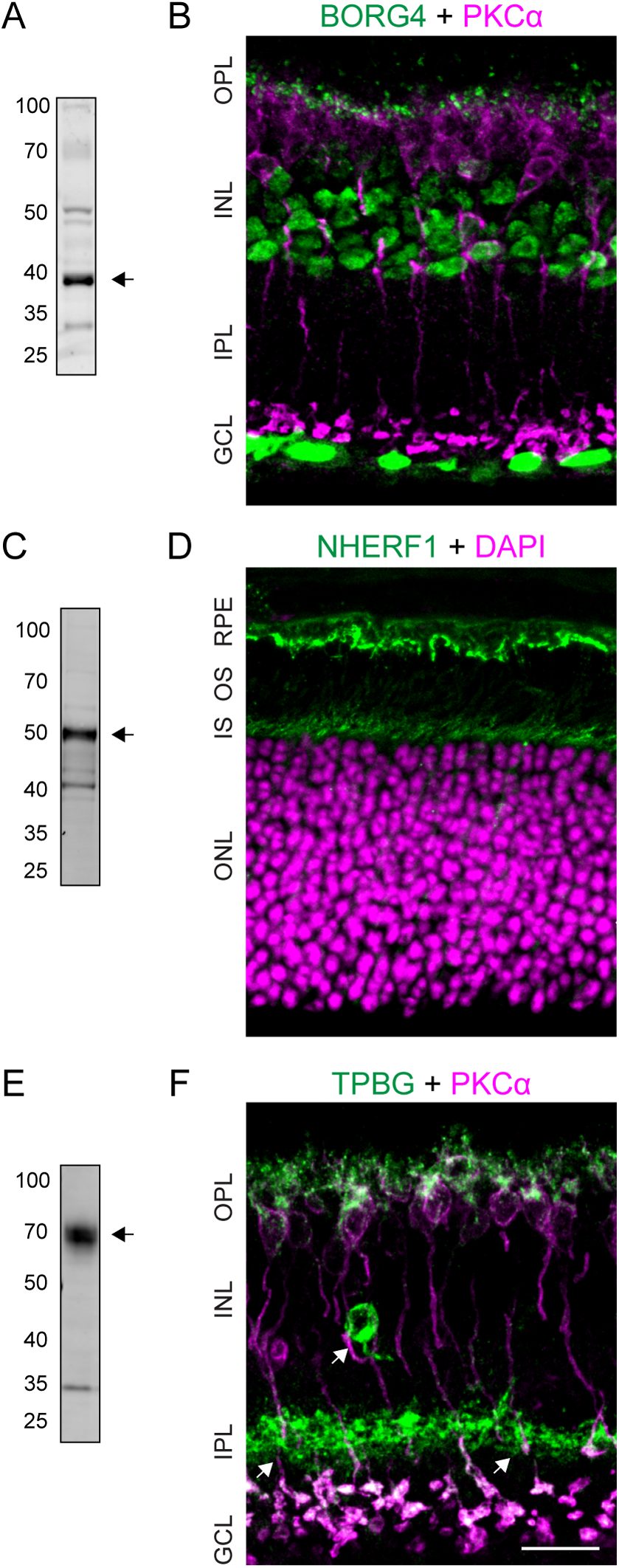
Validation of representative phosphoproteins in the mouse retina. (A) Immunoblot of retinal lysate shows a band corresponding to BORG4 at ∼38 kDa. (B) Confocal microscopy analysis of BORG4 and PKCα immunoreactivity in the retina. (C) Immunoblot of retinal lysate shows a band corresponding to NHERF1 at ∼50 kDa. (D) Confocal microscopy analysis of NHERF1 immunoreactivity in the retina with retinal layers labeled with DAPI. (E) Immunoblot of retinal lysate shows a smear corresponding to glycosylated TPBG at ∼72 kDa. (F) Confocal microscopy analysis of TPBG and PKCα immunoreactivity in the retina. Scale bars: 20 μm. RPE: retinal pigment epithelium; OS: outer segments; IS: inner segments; ONL: outer nuclear layer; OPL: outer plexiform layer; INL: inner nuclear layer; IPL: inner plexiform layer; GCL: ganglion cell layer.

Immunoblotting of retinal proteins for NHERF1 detected a strong band at ∼50 kDa, the predicted molecular weight of NHERF1 (Figure 9C). Immunofluorescent labeling of retina sections revealed NHERF1 immunoreactivity at the level of the photoreceptor inner segments and in the retinal pigment epithelium (RPE; Figure 9D). It was not possible to determine whether NERF1 at the level of the inner segments is due to its presence in the photoreceptors themselves or the apical microvilli of the RPE, which surround the photoreceptor outer and inner segments. No NHERF1 immunofluorescence was seen in the synaptic layers of the retina.

Immunoblotting of retina lysate for TPBG labels a broad band centered around 72 kDa (Figure 9E). This is higher than the predicted molecular weight of 42 kDa, but is consistent with extensive glycosylation of TPBG [34]. Immunofluorescent localization of TPBG revealed strong immunoreactivity in the OPL as well as in large synaptic terminals at sublamina 5 of the inner plexiform layer (IPL; Figure 9F). In the OPL, TPBG labeling co-localizes with PKCα in RBC dendrites and cell bodies. In the IPL, TPBG immunoreactivity overlaps with all PKCα-positive RBC synaptic terminals. The immunofluorescence localization of TPBG to RBCs corroborates previous findings that TPBG mRNA is expressed primarily in RBCs [35]. The TPBG antibody also labeled a population of amacrine cells with cell bodies in the inner INL and dense dendritic projections to the middle of the IPL (Figure 9F arrows). The existence of a TPBG-positive amacrine cell population in the retina was observed by Imamura et al. in 2006 [36], but its identity has not been established.

## 4. Discussion

RBCs are able to modulate their responses to changing light conditions and evidence suggests this process is regulated in part by PKCα [9,10,37]. The transient physiological effect of PKCα on the RBC dendrites is presumably mediated by its kinase activity, whereby PKCα-dependent phosphorylation changes the activity of downstream proteins. We show that RBC dendrites are the main sites of light- and PKCα-dependent phosphorylation (Figure 1). Using a conformation-specific PKCα antibody and an antibody mix that recognizes phosphorylated PKC substrate motifs, our results suggest that PKCα is active in light-adapted RBC dendrites. In the dark-adapted retina, PKCα is inactive and PKC substrate phosphorylation in the OPL is significantly reduced (Figures 2A and B). Comparison of light-adapted wild type and PKCα-KO retinas confirmed that phosphorylation in RBC dendrites requires PKCα, and also indicated that phosphorylation in presumed cone bipolar cell dendrites is mediated by a different kinase. Deletion of TRPM1, a calcium-permeable cation channel responsible for signal transduction in ON-bipolar cell dendrites, also results in a significant reduction in RBC labeling with the conformation-specific PKCα antibody, suggesting that TRPM1-mediated calcium influx during the RBC light response is a primary source of calcium required for PKCα activation in RBC dendrites.

Differential abundance of retinal phosphopeptides between WT and PKCα-KO retinas could potentially be due to altered expression of proteins caused by the deletion of PKCα, and not due to changes in PKCα kinase activity. To identify proteins whose phosphorylation status is dependent on PKCα, but whose expression levels are unaffected, we compared both phosphoprotein and total protein abundance between WT and PKCα-KO mouse retinas using isobaric tagging and mass spectrometry. We identified over 4000 total proteins in either WT or PKCα-KO samples with 23 showing significantly different expression between groups. These proteins can be clustered into several general categories corresponding to biological ontology, most notably proteins involved in regulating cell shape and vesicle transport, transcriptional regulation, and homeostasis and metabolism. PKCα has been implicated in a variety of diverse cell signaling processes, including cell proliferation and morphology, inflammation, and tumorgenesis. Furthermore, PKCα activity has a major impact on gene expression by modulating transcription factors such as CREB, NF-κB, and c-REL [38]. Identifying proteins that show altered expression levels in PKCα-KO mouse retina could be valuable to the examination of many PKCα-dependent functions in RBCs, such as regulation of RBC morphology and development.

To identify RBC proteins that display PKCα-dependent phosphorylation, total retinal phosphopeptides were compared between wild type and PKCα-KO mice using isobaric tagging. Normalizations and differential expression statistical testing for isobaric tagged phosphopeotides differs from traditional phosphoproteomic experiments where replicate numbers are smaller. When replicate numbers are greater than one or two, ratios are no longer an appropriate framework for the analysis. We extended our approach of working directly with aggregated reporter ion intensities for differential protein expression to phosphopeptide datasets. Our approach enables the use of robust statistical testing from genomics packages such as edgeR [23] or Limma [39]. The background of unchanged phosphorylation levels in the enrichment experiment, like that of the total protein experiment, was sufficient in these samples to use standard normalization approaches. Our normalizations and statistical testing for both experiments demonstrates that the same analysis steps used for total protein abundance can be successfully applied to the phosphopeptide datasets (S2 – Total Protein and Phosphopeptide Statistical Testing).

Of over 1100 distinct phosphopeptides identified by multiplex TMT mass spectroscopy, 14 displayed significantly greater phosphorylation in wild type compared to PKCα-KO samples (Figure 6), suggesting their phosphorylation state is dependent on PKCα. These putative PKCα-dependent phosphoproteins may be phosphorylated directly by PKCα or may be phosphorylated by a different kinase whose activity is dependent on PKCα. Only one phosphoprotein, Dnmt3a, showed a significant decrease in abundance in PKCα-KO compared to WT in both the total protein and phosphopeptide experiments (Figures 5 and 6), indicating that reduced abundance of the Dnmt3a phosphopeptide may be due to downregulation of the protein in the KO. Three proteins, BORG4, NHERF1, and TPBG, displayed a particularly striking increase in phosphorylation in wild type compared to PKCα-KO samples (Figures 6 and 7). Of these proteins, BORG4 and NHERF1 are known substrates of PKCα [40,41].

Surprisingly, several of the major PKCα-dependent phosphoproteins identified in this study, including TPBG, do not conform to a strong consensus PKCα substrate motif, in which a phosphorylated serine is flanked by positively charged arginine and lysine residues (Kinexus Database; Vancouver, Canada; http://www.kinexusnet.ca). However, kinase substrate motifs are not always linear, contiguous sequences, but can also be formed structurally by bending of flexible loops that brings positively charged amino acids into proximity to the phosphorylated serine or threonine residues [42]. In the case of TPBG, the cytoplasmic domain is predicted to be unstructured and flexible, possibly allowing the phosphorylated serines at the C-terminal tail (S422 and S424) to be brought close to upstream pairs of lysines and arginines (R384 K385 and K388 and K389) to form a structural PKCα substrate motif. Alternatively, TPBG and other PKCα-dependent non-consensus phosphopeptides may be phosphorylated by a downstream kinase that is dependent on PKCα activity. For example, casein kinase 2 (CK2) is a serine/threonine kinase that is activated by PKCα [43,44], and whose consensus substrate motif is a serine flanked by acidic residues (Kinexus Database). Two of the phosphopeptides that showed increased abundance in wild type samples compared to PKCα-KO contain a CK2 substrate motif (TPBG and Mfap2). The same site in Mfap2 has previously been demonstrated to be phosphorylated *in vitro*. The C-terminals of the NR2B subunit of the NMDA receptor is similar in sequence to the C-terminus of TPBG (LSSIESDV compared to LSSNSDV), and CK2 phosphorylation of the C-terminal serine in NR2B has been demonstrated to regulate trafficking of the receptor [45]. Ten phosphopeptides were more abundant in PKCα-KO samples than in wild type. These are likely to be phosphorylated by kinases that are inhibited by PKCα, such as GSK3 [46,47]. The consensus substrate motif for GSK3 kinase is a serine residue with a neighboring proline (Kinexus Database), and several of the phosphopeptides that are increased in the PKCα-KO samples fit this motif (examples: Dyn3, Crocc, Map2, and Rtn4).

We used immunofluorescence to localize three phosphoproteins that displayed the greatest differential phosphorylation between wild type and PKCα-KO samples. Immunofluorescence labeling of BORG4 resulted in bright puncta in the OPL (Figure 9) consistent with BORG4 localization to either RBC or horizontal cell dendrites, as well as immunofluorescence over most nuclei. BORG4 belongs to a protein family (BORG1-5) that bind to the Rho GTPase CDC42 as well as to septins, a family of GTP-binding cytoskeletal proteins that are involved in regulation of cell morphology through modulation of cytoskeletal rearrangement. BORG4 contains an N-terminal CCD42/Rac Interactive Binding Motif (CRIB) and three BORG Homology (BHs) domains that are conserved across all BORG family proteins [48]. Our proteomics data indicates that the serine residue S64 in BORG4, which is located between BH1 and BH2, is phosphorylated in a PKCα-dependent manner. Phosphorylation of BORG4 S64 has been previously recognized in large-scale analyses experiments of tissue-specific phosphorylation patterns [49,50] though not in retina. BORG4 and Septin-4 mRNAs have been previously found to be expressed in horizontal cells [51]. Our immunofluorescence labeling of BORG4 in retina resulted in bright puncta in the OPL consistent with BORG4 localization to either RBC or horizontal cell dendrites.

We detected NHERF1 immunofluorescence at the level of the photoreceptor inner segments in the region of the connecting cilia and in the retina pigment epithelium (RPE, Figure 9). NHERF1 is a scaffolding protein containing tandem PDZ domains and an Ezrin/Radixin/Moesin Binding (EB) domain. Our detection of strong NHERF1 immunofluorescence in the RPE consistent with previous reports localizing NHERF1 to the RPE apical microvilli [52]. NHERF1 interacts with ezrin to maintain the structure of apical microvilli on epithelia. In the RPE, NHERF1 has been implicated in retinoid recycling [52] through its interactions with CRALBP [53,54]. PKCα is also expressed in the RPE [55] where it is involved in proliferation and migration [56], phagocytosis [57], and melanin production [58]; however, the specific role of PKCα-mediated phosphorylation of NHERF1 in the RPE is unknown.

Immunofluorescence confocal microscopy localized TPBG immunoreactivity to RBC dendrites and synaptic terminals, as well as to a class of amacrine cells (Figure 9). TPBG is a heavily glycosylated type-1 transmembrane protein with a large extracellular N-terminal domain and a short C-terminal intracellular tail. The N-terminal domain contains eight leucine-rich repeats (LRRs) and seven N-linked glycosylation sites. The C-terminal tail ends with the class-1 PDZ-interacting motif (S/T X Φ) SDV [59], which our proteomics data suggests is phosphorylated in a PKCα-dependent manner. Since phosphorylation of a PDZ-interacting motif typically prevents binding of a PDZ protein [60], PKCα might be regulating interactions between TPBG and PDZ proteins by stimulating phosphorylation of its C-terminal tail. As an oncofetal antigen, TPBG is present primarily during embryonic development [61,62], but is also expressed in many carcinomas (Southall et al., 1990). In the adult, TPBG is expressed in the brain, retina, and ovaries [65,66]. In the embryo and in cancer tissue, TPBG is involved in regulating actin polymerization [67] filopodia formation [68], and chemotaxis [69,70], and its expression in tumors is linked to increased metastatic malignancy and poor survival outcomes in cancer patients [71,72]. In the adult olfactory bulb, TPBG is required to stimulate the development of input-dependent dendritic arborization and synaptogenesis of newborn granule cells [36,73–75]. The role of TPBG in the retina is not yet understood; however, a recent transcriptomic classification of retinal cell types identified TPBG mRNA as being highly enriched in RBCs [35]. This is consistent with our immunofluorescent analysis which localized TPBG to RBC dendrites and synaptic terminals.

## 5. Conclusions

The molecular mechanisms of PKCα-mediated modulation of the RBC light response have not been thoroughly explored. In this study, we have shown that PKCα phosphorylation in the retina occurs predominately in RBC dendrites in the light. Using a phosphoproteomics approach, we have identified a small number of phosphoproteins with significantly increased PKCα-dependent phosphorylation in the PMA-treated retina. These differentially phosphorylated proteins fall into several broad functional groups, including cytoskeleton/trafficking (4 proteins), structure and adhesion (2 proteins), cell signaling (2 proteins), transcriptional regulation (3 proteins), and homeostasis/metabolism (1 protein). Two strongly differentially expressed phosphoproteins, BORG4 and TPBG, are localized to the synaptic layers of the retina, and may play a role in PKCα-dependent modulation of RBC function.

## Supporting information

S1 - Total Protein and Phosphopeptide Results

S2 - Total Protein and Phosphopeptide Data Analysis

S3 - Annotated Phosphopeptide Spectra

## Acknowledgments

The authors would like to thank Tammie L. Haley for producing mouse retina sections for immunofluorescence experiments. This work was supported by the National Institutes of Health grants R01EY022369 and 5P30EY010572 and an OHSU University Shared Resources Core Pilot Fund grant.

## Author contributions

study design: CMW, LLD, CWM; experimentation: CMW, JMC, PAW, GR, CWM; PAW Pipeline script development: PAW; data analysis: CMW, PAW, JEK, LLD, CWM; figure and manuscript preparation: CMW, JMC, PAW, JEK, LLD, CWM; editing and review: CMW, JMC, PAW, GR, JEK, LLD, CWM; repository preparation: CMW, PAW; supplemental files preparation: CMW, PAW

## Abbreviations

BORG4: Binder of Rho GTPase 4
DAG: diacylglycerol
ERG: electroretinogram
DE: differential expression
FC: fold change
FDR: false discovery rate
GCL: ganglion cell layer
INL: inner nuclear layer
IPL: inner plexiform layer
KO: knockout
LC-MS/MS: liquid chromatography tandem mass spectroscopy
NHERF1: Na^+^/H^+^ Exchange Regulatory Factor 1
OPL: outer plexiform layer
PKCα/Prkca: Protein Kinase C-alpha
PMA: phorbol 12-myristate 13-acetate
PSM: peptide spectrum match
RBC: rod bipolar cell
RPE: retinal pigment epithelium
TMT: tandem mass tag
TPBG: Trophoblast Glycoprotein
TRPM1: Transient Receptor Potential cation channel subfamily M member 1

