## Supplementary material for "Identification of PKCα-dependent phosphoproteins in mouse retina": S2 - Total Protein and Phosphopeptide Data Analysis

whole\_protein


### TMT Whole Protein and Phosphopeptide Analysis¶

##### Submitted to ...¶

Mice retina were treated with PMA (an activator of DAG-sensitive protein kinase C isoforms such as PKCa), cells lysed, protein extracted, and digested with trypsin. The mice were either wild-type (n=4) or protein kinase C-alpha (PKCa) knock out animals (n=5). Some of the resulting peptides were directly labeled with 10-plex TMT reagents {Thompson 2003}. Remaining peptides were enriched for phosphopepides, labeled with 10-plex TMT for a second set of labeled samples. After combining equal portions of labeled peptide samples (either whole protein or enriched phosphopeptides), they were separated by high pH reverse phase/low pH reverse phase liquid chromatography and analyzed on a Thermo Fusion Tribrid Orbitrap mass spectrometer {Senko 2013}. Fragment ions were generated using CID and the ion trap analyzer; the reporter ions were fragmented using high energy collision dissociation of (synchronous precursor selection) SPS selected fragment ions at high resolution in the Orbitrap analyzer.

Peptides and proteins were identified using Proteome Discoverer v1.4 and SEQUEST {Eng 1994}. A wider 1.25 Da parent ion mass tolerance was used, TMT labels and alkylated cysteine were specified as static modifications, oxidation of methionine (+15.9949 Da) was specified as a variable modification, trypsin enzyme specificity was used, and a canonical UniProt Swiss-Prot mouse protein database was used. Fragment ion tolerance was set at 1.0005 Da. Confident peptide identifications were obtained using Percolator and the target/decoy method. PSM information (peptide sequences, q-values, masses, and reporter ions) was exported to tab-delimited files for processing with PAW pipeline modules.

The labeled phosphopeptides were analyzed using similar SEQUEST search parameters with the addition of variable phosphorylation (+79.9799) on serine, threonine, or tyrosine residues. Proteome Discoverer was also configured with a phosphRS {Taus 2011} node for site localizations.

The PAW pipeline Python scripts take the files exported from Proteome Discoverer, does some additional accurate peptide mass and minimum reporter ion intensity filtering. The exported data from Proteome Discoverer has already undergone PSM confidence filtering (Percolator q-values) and parsimonious protein inference. Reporter ions are peak heights of the most confident (closest to expected m/z value) centroided peak within 20 ppm of the expected position.

For differential protein expression, the reporter ions from all peptide-spectrum-matches (PSMs) from unique peptides associated with each protein were summed into protein total reporter ions for each TMT channel (the biological samples). This aggregation step dramatically reduces dataset dimensionality (10 to 20 fold) and improves data quality. Normalizations and statistical testing were performed using the Bioconductor {Gentleman 2004} package edgeR {Robinson 2010} as detailed in the first half of the notebook below. A Jupyter notebook with an R kernel was used to execute R commands and visualize the results.

Phosphopeptide enrichment is very different compared to protein expression. The measurements are peptide-centric rather than protein centric. Protein inference cannot be used as a noise filter to reduce the negative impact of incorrect PSMs. The positive effect of aggregated data to the protein level is also greatly compromised. We did a couple of things to address these issues. We increased the q-value cutoff for PSM filtering from 5% to 1%. We also considered phospho group localization to be less reliable than determination of the base peptide sequence and total number of phospho groups present. To get some degree of data aggregation to reduce the multiple testing impact and improve the data quality, we combined all PSMs from the same base peptide sequence and the same number of phosphogroups. This aggregation step reduces the number of data points in the quantitative analysis by about a factor of 2. The aggregation is not uniform across the phosphopeptides. Many peptides only have a single PSM, others may have many combined PSMs. We carried forward localization information in a limited way. The localization of the most confident PSM (smallest q-value) is reported and annotated as consistent (all PSMs has the same site localization) or variable. Analysis of the phosphopeptide data is in the second half of this notebook.

> Thompson, A., Schäfer, J., Kuhn, K., Kienle, S., Schwarz, J., Schmidt, G., Neumann, T. and Hamon, C., 2003. Tandem mass tags: a novel quantification strategy for comparative analysis of complex protein mixtures by MS/MS. Analytical chemistry, 75(8), pp.1895-1904.
>
> Senko, M.W., Remes, P.M., Canterbury, J.D., Mathur, R., Song, Q., Eliuk, S.M., Mullen, C., Earley, L., Hardman, M., Blethrow, J.D. and Bui, H., 2013. Novel parallelized quadrupole/linear ion trap/Orbitrap tribrid mass spectrometer improving proteome coverage and peptide identification rates. Analytical chemistry, 85(24), pp.11710-11714.
>
> Eng, J.K., McCormack, A.L. and Yates, J.R., 1994. An approach to correlate tandem mass spectral data of peptides with amino acid sequences in a protein database. Journal of the American Society for Mass Spectrometry, 5(11), pp.976-989.
>
> McAlister, G.C., Nusinow, D.P., Jedrychowski, M.P., Wühr, M., Huttlin, E.L., Erickson, B.K., Rad, R., Haas, W. and Gygi, S.P., 2014. MultiNotch MS3 enables accurate, sensitive, and multiplexed detection of differential expression across cancer cell line proteomes. Analytical chemistry, 86(14), pp.7150-7158.
>
> Wilmarth, P.A., Riviere, M.A. and David, L.L., 2009. Techniques for accurate protein identification in shotgun proteomic studies of human, mouse, bovine, and chicken lenses. Journal of ocular biology, diseases, and informatics, 2(4), pp.223-234.
>
> Taus, T., Köcher, T., Pichler, P., Paschke, C., Schmidt, A., Henrich, C. and Mechtler, K., 2011. Universal and confident phosphorylation site localization using phosphoRS. Journal of proteome research, 10(12), pp.5354-5362.
>
> Gentleman, R.C., Carey, V.J., Bates, D.M., Bolstad, B., Dettling, M., Dudoit, S., Ellis, B., Gautier, L., Ge, Y., Gentry, J. and Hornik, K., 2004. Bioconductor: open software development for computational biology and bioinformatics. Genome biology, 5(10), p.R80.
>
> Robinson, M.D., McCarthy, D.J. and Smyth, G.K., 2010. edgeR: a Bioconductor package for differential expression analysis of digital gene expression data. Bioinformatics, 26(1), pp.139-140.

In [1]:

```
# load the libraries
library(tidyverse)
library(stringr)
library(edgeR)
library(limma)
library(psych)
```

```
── Attaching packages ─────────────────────────────────────── tidyverse 1.2.1 ──
✔ ggplot2 3.1.0     ✔ purrr   0.2.5
✔ tibble  1.4.2     ✔ dplyr   0.7.8
✔ tidyr   0.8.2     ✔ stringr 1.3.1
✔ readr   1.1.1     ✔ forcats 0.3.0
── Conflicts ────────────────────────────────────────── tidyverse_conflicts() ──
✖ dplyr::filter() masks stats::filter()
✖ dplyr::lag()    masks stats::lag()
Loading required package: limma
Warning message:
“package ‘limma’ was built under R version 3.5.1”
Attaching package: ‘psych’

The following objects are masked from ‘package:ggplot2’:

    %+%, alpha
```

### Whole protein¶

#### Read in the data that was prepped in Excel¶

The TMT data was labeled by sample for respective channels, and any contaminants were excluded. The rows have accessions and description in case we need those. Once the data is read in, we will extract the TMT columns and the accessions (like gene names). We know that the first 4 channels are the wild type (WT) and that the next 5 are the knock out (KO).

In [2]:

```
# read in the data export file
data_all <- read_tsv("whole_protein.txt")
counts <- data_all %>% select(starts_with("TotInt_"))
accessions <- data_all$Accession
WT  <- 1:4
KO <- 5:9
```

```
Parsed with column specification:
cols(
  Counter = col_integer(),
  Accession = col_character(),
  Description = col_character(),
  `# Used PSMs` = col_integer(),
  `TotInt_127-N_WT-1` = col_integer(),
  `TotInt_128-N_WT-2` = col_integer(),
  `TotInt_129-N_WT-3` = col_integer(),
  `TotInt_130-N_WT-5` = col_integer(),
  `TotInt_126C_PKC-KO-1` = col_integer(),
  `TotInt_127C_PKC-KO-2` = col_integer(),
  `TotInt_128C_PKC-KO-3` = col_integer(),
  `TotInt_129C_PKC-KO-4` = col_integer(),
  `TotInt_131-N_PKC-KO-5` = col_integer()
)
```

#### Load the data into edgeR to normalize¶

We will eventually be using edgeR for the statistical testing. The first thing we need to do is normalize the data. EdgeR has a normalization function that corrects for total signal in each channel (like a library size correction) and does a compositional bias correction called a trimmed mean of M-values (TMM) normalization.

> Robinson, M.D. and Oshlack, A., 2010. A scaling normalization method for differential expression analysis of RNA-seq data. Genome biology, 11(3), p.R25.

In [3]:

```
# load into edgeR DGEList object and normalize the data
group = c(rep("WT", 4), rep("KO", 5))
yw <- DGEList(counts = counts, group = group, gene = accessions)
yw <- calcNormFactors(yw)
yw$samples
```

|  | group | lib.size | norm.factors |
| --- | --- | --- | --- |
| TotInt\_127-N\_WT-1 | WT | 862499506 | 0.9892065 |
| TotInt\_128-N\_WT-2 | WT | 908475057 | 0.9826495 |
| TotInt\_129-N\_WT-3 | WT | 931588739 | 0.9966336 |
| TotInt\_130-N\_WT-5 | WT | 903720499 | 0.9903727 |
| TotInt\_126C\_PKC-KO-1 | KO | 1051577135 | 1.0285432 |
| TotInt\_127C\_PKC-KO-2 | KO | 924259426 | 1.0090856 |
| TotInt\_128C\_PKC-KO-3 | KO | 830710814 | 0.9969902 |
| TotInt\_129C\_PKC-KO-4 | KO | 851461610 | 0.9878139 |
| TotInt\_131-N\_PKC-KO-5 | KO | 924743408 | 1.0196793 |

#### Check the normalization¶

EdgeR does not export the actual normalized data, but we can get the factors and do the calculations in R. We can compare box plots of the total protein intensities of the starting data and after the TMM normalization. The normalization factors should be near 1.0 and the boxes in the boxplots should be horizontally aligned. The notches are the medians and the heights of the boxes are the interquartile ranges. Notches and boxes should be in a very tight horizontal line when the normalization has worked well.

In [4]:

```
# Compute the normalized intensities (start with data_raw)
# sample loading adjusts each channel to the same average total
lib_facs <- mean(colSums(counts)) / colSums(counts)
print("Library size normalization factors")
round(lib_facs, 4)

# the TMM factors are library adjustment factors (so divide by them)
norm_facs <- lib_facs / yw$samples$norm.factors
print("Combined Lib+TMM normalization factors")
round(norm_facs, 4)

# compute the normalized data as a new data frame
data_norm <- sweep(counts, 2, norm_facs, FUN = "*")
colnames(data_norm) <- str_c(colnames(counts), "_TMMNorm")

# check norm before and after TMM
colors = c(rep('red', 4), rep('blue', 5))
boxplot(log10(counts), col = colors,
        xlab = 'TMT samples', ylab = 'log10 Intensity', 
        main = 'Whole protein starting data', notch = TRUE)
boxplot(log10(data_norm), col = colors,
        xlab = 'TMT samples', ylab = 'log10 Intensity', 
        main = 'Whole protein TMM Normalized data', notch = TRUE)
```

```
[1] "Library size normalization factors"
```

TotInt\_127-N\_WT-1
:   1.0549

TotInt\_128-N\_WT-2
:   1.0016

TotInt\_129-N\_WT-3
:   0.9767

TotInt\_130-N\_WT-5
:   1.0068

TotInt\_126C\_PKC-KO-1
:   0.8653

TotInt\_127C\_PKC-KO-2
:   0.9845

TotInt\_128C\_PKC-KO-3
:   1.0953

TotInt\_129C\_PKC-KO-4
:   1.0686

TotInt\_131-N\_PKC-KO-5
:   0.9839

```
[1] "Combined Lib+TMM normalization factors"
```

TotInt\_127-N\_WT-1
:   1.0665

TotInt\_128-N\_WT-2
:   1.0192

TotInt\_129-N\_WT-3
:   0.98

TotInt\_130-N\_WT-5
:   1.0166

TotInt\_126C\_PKC-KO-1
:   0.8413

TotInt\_127C\_PKC-KO-2
:   0.9756

TotInt\_128C\_PKC-KO-3
:   1.0986

TotInt\_129C\_PKC-KO-4
:   1.0818

TotInt\_131-N\_PKC-KO-5
:   0.965

#### Do samples cluster by biological condition?¶

The normalization above looks pretty good. Boxplots can be deceiving and a clustering view can make sure that things are really behaving correctly. We will also compute the variance trends and look at that, too.

In [5]:

```
# check the clustering (set colors by condition)
plotMDS(yw, col = colors, main = "Whole protein: WT (red) and KP (blue)")

# compute dispersions and plot BCV
yw <- estimateDisp(yw)
plotBCV(yw, main = "Whole protein: variance trend")
```

```
Design matrix not provided. Switch to the classic mode.
```

#### Data looks ready for statistical testing¶

Samples are separating by biological condition and the variance is small until we get down to small TMT intensities. We can perform an exact pairwise test in edgeR and see how many differential expression (DE) candidates we have at an FDR of 10%. We can list the DE candidates (we do this in two passes to figure out that we have 23 candidates). We will also save the testing results to eventually export.

In [6]:

```
# the exact test object has columns like fold-change, CPM, and p-values
# we have already loaded the data into "y" above, normalized, and est. dispersion
etw <- exactTest(yw, pair = c("WT", "KO"))

# this counts up, down, and unchanged genes (here it is proteins)
summary(decideTestsDGE(etw, p.value = 0.10))
topTags(etw, n = 23)

# the topTags function adds the BH FDR values to an exactTest data frame 
# make sure we do not change the row order!
ttw <- topTags(etw, n = Inf, sort.by = "none")
ttw <- ttw$table    # ttw is a list. We just need the data frame table
```

```
       WT+KO
Down      15
NotSig  4422
Up         8
```

$table
:   |  | genes | logFC | logCPM | PValue | FDR |
    | --- | --- | --- | --- | --- | --- |
    | 2967 | P20444 | -3.8414878 | 8.07110886 | 1.023811e-145 | 4.550838e-142 |
    | 2168 | Q5HZI1 | 4.5944720 | 4.78221565 | 5.111884e-30 | 1.136116e-26 |
    | 2383 | P12961 | 1.0300598 | 5.33867570 | 4.002880e-16 | 5.930933e-13 |
    | 1179 | Q91WT4 | -1.5677618 | 4.53341044 | 5.136010e-11 | 5.707391e-08 |
    | 1160 | Q9CQZ7 | 2.6803928 | -0.56372875 | 1.196139e-09 | 1.063368e-06 |
    | 3600 | Q6PIE5 | -0.7010443 | 8.40378942 | 1.700588e-09 | 1.259852e-06 |
    | 1767 | P84228 | -0.8085652 | 4.80747505 | 5.206346e-08 | 3.306030e-05 |
    | 1127 | O88508 | -0.5340565 | 5.68958367 | 3.945843e-07 | 2.192409e-04 |
    | 2710 | Q8CJH3 | 2.4154671 | -0.19343522 | 4.603523e-07 | 2.273629e-04 |
    | 1474 | Q9ER35 | -0.9073843 | 3.02357914 | 1.149515e-06 | 5.109596e-04 |
    | 4393 | Q02526 | -0.6576858 | 6.70202176 | 1.849027e-06 | 7.471751e-04 |
    | 1729 | O54879 | -0.4967832 | 10.73663686 | 3.053464e-06 | 1.131054e-03 |
    | 3421 | Q3UQA7 | -0.5883120 | 6.44559443 | 6.035174e-06 | 2.063565e-03 |
    | 1998 | Q9D1G5 | 0.5313280 | 7.05368110 | 1.102095e-05 | 3.499153e-03 |
    | 424 | P55088 | 0.4768787 | 9.97015032 | 1.629114e-05 | 4.827609e-03 |
    | 3139 | Q8BMG7 | 2.0580847 | -0.08212585 | 1.796787e-05 | 4.991699e-03 |
    | 2216 | Q9Z1S0 | 1.6701794 | 0.51364408 | 4.707243e-05 | 1.230806e-02 |
    | 989 | Q9Z1J3 | -1.1176160 | 2.87805201 | 5.729663e-05 | 1.414909e-02 |
    | 895 | Q9DA03 | -2.3067647 | -0.18052633 | 6.223601e-05 | 1.455995e-02 |
    | 2295 | Q99LD8 | -0.4724754 | 9.94587730 | 1.211718e-04 | 2.693044e-02 |
    | 635 | P12658 | -1.0775398 | 6.20659355 | 1.672232e-04 | 3.539558e-02 |
    | 2565 | O70209 | -0.4281642 | 5.39567284 | 3.340195e-04 | 6.748713e-02 |
    | 2150 | P55002 | -1.8216902 | 3.98354729 | 4.398869e-04 | 8.501292e-02 |

$adjust.method
:   'BH'

$comparison
:   1. 'WT'
    2. 'KO'

$test
:   'exact'

#### Visualize the candidates and check test p-values¶

EdgeR has a convenient MA plot function. We can also plot the distribution of p-values. We expect the plot to have two superimposed distributions. One is a uniform (flat) distribution from 0 to 1 from the non-DE proteins. We may also have some small p-values that cause a "spike" at p-values near zero associated with the true DE candidates.

In [7]:

```
# make an MD plot
plotMD(etw, p.value = 0.10, main = "Whole protein: KO vs WT")
abline(h = c(-1, 1), col = "black")

# check the p-value distribution
ggplot(ttw, aes(PValue)) + 
  geom_histogram(bins = 100, fill = "white", color = "black") + 
  geom_hline(yintercept = mean(hist(etw$table$PValue, breaks = 100, 
                                    plot = FALSE)$counts[26:100])) +
  ggtitle("Whole protein: KO vs WT p-value distribution")
```

In [8]:

```
de_plots <- function(tt, x, y, title) {
  temp <- data.frame(log10((tt[x] + tt[y])/2), 
                     log2(tt[y] / tt[x]), 
                     tt$candidate,
                     -log10(tt$FDR))
  colnames(temp) <- c("A", "M", "candidate", "P")
    
  ma_lines <- list(geom_hline(yintercept = 0.0, color = "black"),
                   geom_hline(yintercept = 1.0, color = "black", linetype = "dotted"),
                   geom_hline(yintercept = -1.0, color = "black", linetype = "dotted"))

  scatter_lines <- list(geom_abline(intercept = 0.0, slope = 1.0, color = "black"),
                        geom_abline(intercept = 0.301, slope = 1.0, color = "black", 
                                    linetype = "dotted"),
                        geom_abline(intercept = -0.301, slope = 1.0, color = "black", 
                                    linetype = "dotted"),
                        scale_y_log10(), scale_x_log10())
    
  # make main MA plot
  first  <- ggplot(temp, aes(x = A, y = M)) +
    geom_point(aes(color = candidate, shape = candidate)) +
    scale_y_continuous(paste0("logFC (", y, "/", x, ")")) +
    scale_x_continuous("Ave_intensity") +
    ggtitle(title) + 
    ma_lines
    
  # make separate MA plots
  second <- ggplot(temp, aes(x = A, y = M)) +
    geom_point(aes(color = candidate, shape = candidate)) +
    scale_y_continuous(paste0("log2 FC (", y, "/", x, ")")) +
    scale_x_continuous("log10 Ave_intensity") +
    ma_lines +
    facet_wrap(~ candidate) +
    ggtitle(paste(title, "(separated)", sep=" "))

  # make main scatter plot
  third <- ggplot(tt, aes_string(x, y)) +
    geom_point(aes(color = candidate, shape = candidate)) +
    ggtitle(title) + 
    scatter_lines

  # make separate scatter plots
  fourth <- ggplot(tt, aes_string(x, y)) +
    geom_point(aes(color = candidate, shape = candidate)) +
    scatter_lines +
    facet_wrap(~ candidate) +
    ggtitle(paste(title, "(separated)", sep=" ")) 

  # make volcano plot
  fifth <- ggplot(temp, aes(x = M, y = P)) +
    geom_point(aes(color = candidate, shape = candidate)) +
    xlab("log2 FC") +
    ylab("-log10 FDR") +
    ggtitle(paste(title, "Volcano Plot", sep = ' '))
    
  print(first) # still need to get the plots to appear
  print(second)
  print(third)
  print(fourth)
  print(fifth)
}
```

#### Make some more elaborate candidate visualizations¶

The above function uses ggplot2 to make some nice MA and scatter plots with the DE candidates highlighted. We can also make separate plots for each candidate class. We can also do a traditional volcano plot. We need some condition average columns for plotting. We can add those and the candidate status to the edgeR test results.

In [9]:

```
# get the averages within each condition
ttw$ave_WT <- rowMeans(data_norm[WT])
ttw$ave_KO <- rowMeans(data_norm[KO])

# add the cadidate status column
ttw <- ttw %>%
  mutate(candidate = cut(FDR, breaks = c(-Inf, 0.01, 0.05, 0.10, 1.0),
  labels = c("high", "med", "low", "no")))

# make the DE plots
de_plots(ttw, "ave_WT", "ave_KO", "Whole protein WT versus KO")

# label columns so we know what the comparison was
colnames(ttw) <- str_c(colnames(ttw), '_WT_KO')
```

#### Gather up results and export¶

We need to save the results so we can add them back to our main summary spreadsheet. We should also log the session info for our records.

In [10]:

```
# export the results: TMM normed data, 
#   statistical test results (includes gene names), averages, and candidate status
data_export <- cbind(data_norm, ttw)
write.table(data_export, "whole_protein_results.txt", sep = "\t", row.names = FALSE)
```

### Phosphopeptide data¶

#### Read in the data exported from Excel¶

##### Load data into edgeR and normalize¶

In most protein expression studies, the assumption of the majority of the proteins not having any changes in expression levels is central to normalization strategies. This concept is much less tested in phosphopeptide enrichment studies. We need to pay close attention to the sizes of the normalization factors and the alignment of the boxplots to verify that the normalization methods used for the whole protein analysis above are still valid.

In [11]:

```
# read in the data export file
data_all_pep <- read_tsv("phosphopeptide.txt")
counts_pep <- data_all_pep %>% select(starts_with("TotInt_"))
keys <- data_all_pep$Key
WT  <- 1:4
KO <- 5:9
```

```
Parsed with column specification:
cols(
  Key = col_character(),
  `TotInt_127-N_WT-1` = col_integer(),
  `TotInt_128-N_WT-2` = col_integer(),
  `TotInt_129-N_WT-3` = col_integer(),
  `TotInt_130-N_WT-5` = col_integer(),
  `TotInt_126C_PKC-KO-1` = col_integer(),
  `TotInt_127C_PKC-KO-2` = col_integer(),
  `TotInt_128C_PKC-KO-3` = col_integer(),
  `TotInt_129C_PKC-KO-4` = col_integer(),
  `TotInt_131-N_PKC-KO-5` = col_integer()
)
```

In [12]:

```
# load into edgeR DGEList object and normalize the data
group = c(rep("WT", 4), rep("KO", 5))
yp <- DGEList(counts = counts_pep, group = group, gene = keys)
yp <- calcNormFactors(yp)
yp$samples
```

|  | group | lib.size | norm.factors |
| --- | --- | --- | --- |
| TotInt\_127-N\_WT-1 | WT | 29068200 | 0.9829834 |
| TotInt\_128-N\_WT-2 | WT | 35413203 | 0.9639961 |
| TotInt\_129-N\_WT-3 | WT | 35341792 | 0.9992645 |
| TotInt\_130-N\_WT-5 | WT | 34030649 | 0.9778852 |
| TotInt\_126C\_PKC-KO-1 | KO | 26694056 | 1.0325551 |
| TotInt\_127C\_PKC-KO-2 | KO | 30742340 | 1.0631632 |
| TotInt\_128C\_PKC-KO-3 | KO | 34018111 | 1.0037429 |
| TotInt\_129C\_PKC-KO-4 | KO | 35862091 | 0.9579204 |
| TotInt\_131-N\_PKC-KO-5 | KO | 32571932 | 1.0231637 |

In [13]:

```
# Compute the normalized intensities (start with data_raw)
# sample loading adjusts each channel to the same average total
lib_facs_pep <- mean(colSums(counts_pep)) / colSums(counts_pep)
print("Library size normalization factors")
round(lib_facs_pep, 4)

# the TMM factors are library adjustment factors (so divide by them)
norm_facs_pep <- lib_facs_pep / yp$samples$norm.factors
print("Combined Lib+TMM normalization factors")
round(norm_facs_pep, 4)

# compute the normalized data as a new data frame
data_norm_pep <- sweep(counts_pep, 2, norm_facs_pep, FUN = "*")
colnames(data_norm_pep) <- str_c(colnames(counts_pep), "_TMMNorm")

# check norm before and after TMM
colors = c(rep('red', 4), rep('blue', 5))
boxplot(log10(counts_pep), col = colors,
        xlab = 'TMT samples', ylab = 'log10 Intensity', 
        main = 'Phosphopeptide starting data', notch = TRUE)
boxplot(log10(data_norm_pep), col = colors,
        xlab = 'TMT samples', ylab = 'log10 Intensity', 
        main = 'Phosphopeptide TMM Normalized data', notch = TRUE)
```

```
[1] "Library size normalization factors"
```

TotInt\_127-N\_WT-1
:   1.1228

TotInt\_128-N\_WT-2
:   0.9216

TotInt\_129-N\_WT-3
:   0.9235

TotInt\_130-N\_WT-5
:   0.9591

TotInt\_126C\_PKC-KO-1
:   1.2227

TotInt\_127C\_PKC-KO-2
:   1.0617

TotInt\_128C\_PKC-KO-3
:   0.9594

TotInt\_129C\_PKC-KO-4
:   0.9101

TotInt\_131-N\_PKC-KO-5
:   1.002

```
[1] "Combined Lib+TMM normalization factors"
```

TotInt\_127-N\_WT-1
:   1.1422

TotInt\_128-N\_WT-2
:   0.9561

TotInt\_129-N\_WT-3
:   0.9242

TotInt\_130-N\_WT-5
:   0.9808

TotInt\_126C\_PKC-KO-1
:   1.1841

TotInt\_127C\_PKC-KO-2
:   0.9986

TotInt\_128C\_PKC-KO-3
:   0.9559

TotInt\_129C\_PKC-KO-4
:   0.9501

TotInt\_131-N\_PKC-KO-5
:   0.9793

#### Normalization factors are close to 1 and boxplots look okay¶

The normalization checks seem fine, so we should be in good shape for statistical testing.

#### Check that samples cluster by condition and check variance¶

In [14]:

```
# check the clustering (set colors by condition)
plotMDS(yp, col = colors, main = "WT (red) and KP (blue)")

# compute dispersions and plot BCV
yp <- estimateDisp(yp)
plotBCV(yp, main = "Phosphopeptide Variance Trend")
```

```
Design matrix not provided. Switch to the classic mode.
```

#### Data are ready for statistical testing¶

Perform and exact test and look at the significant peptides.

In [15]:

```
# the exact test object has columns like fold-change, CPM, and p-values
# we have already loaded the data into "y" above, normalized, and est. dispersion
etp <- exactTest(yp, pair = c("WT", "KO"))

# this counts up, down, and unchanged genes (here it is proteins)
summary(decideTestsDGE(etp, p.value = 0.10))
topTags(etp, n = 24)

# the topTags function adds the BH FDR values to an exactTest data frame 
# make sure we do not change the row order!
ttp <- topTags(etp, n = Inf, sort.by = "none")
ttp <- ttp$table    # ttp is a list. We just need the data frame table
```

```
       WT+KO
Down      14
NotSig  1113
Up        10
```

$table
:   |  | genes | logFC | logCPM | PValue | FDR |
    | --- | --- | --- | --- | --- | --- |
    | 571 | Q9Z0L0 LTNLSS\*NS\*DV | -3.7222390 | 7.404119 | 1.050848e-130 | 1.194814e-127 |
    | 891 | Q9JM96 AREADDES\*LDEQASASKLSLLSR | -4.6255349 | 5.748031 | 6.675282e-106 | 3.794898e-103 |
    | 297 | P70441 EALVEPASES\*PRPALAR | -4.8218420 | 9.023062 | 3.936186e-27 | 1.491814e-24 |
    | 248 | Q9JM96 EADDES\*LDEQASASKLSLLSR | -6.1518683 | 9.360071 | 1.014892e-16 | 2.884830e-14 |
    | 293 | P48193 RLS\*THSPFR | -0.9273138 | 9.041728 | 1.019000e-09 | 2.317206e-07 |
    | 933 | Q9Z0L0 LTNLSSNS\*DV | -1.4935835 | 5.509251 | 1.621250e-09 | 3.072268e-07 |
    | 217 | Q8CJ40 SSASVSLPPGT\*PEK | 0.8702826 | 9.618360 | 6.752824e-08 | 1.096851e-05 |
    | 443 | Q69Z99 S\*LRRQQQPCMEPPESQLEPK | -1.1937569 | 8.031428 | 1.140487e-07 | 1.620917e-05 |
    | 476 | P12961 SVPHFS\*EEEKEAE | 0.7384766 | 7.797799 | 3.162974e-07 | 3.995891e-05 |
    | 899 | P06837 EGDGSATTDAAPAT\*SPKAEEPSKAGDAPSEEK | 1.0309266 | 5.717600 | 3.964396e-05 | 4.507518e-03 |
    | 136 | Q3U0L2 RSS\*VRPGVVVPR | -1.0345986 | 10.300574 | 4.685181e-05 | 4.842774e-03 |
    | 170 | Q8CCJ4 RVS\*NRGLAGTTIR | -0.7444850 | 10.013772 | 5.430078e-05 | 5.144999e-03 |
    | 766 | P20357 ARVDHGAEIITQS\*PSRSSVASPR | 0.7700313 | 6.511836 | 6.027953e-05 | 5.272140e-03 |
    | 607 | Q9CQU1 SLAALDALNT\*DDENDEEEYEAWKVR | -0.5797711 | 7.192916 | 1.208066e-04 | 9.811222e-03 |
    | 88 | O70318 QRS\*YNLVVAK | -0.5971405 | 11.006645 | 1.536077e-04 | 1.164346e-02 |
    | 362 | P09174 S\*ATRVIGGPVTPR | -1.3122038 | 8.630354 | 2.288319e-04 | 1.626137e-02 |
    | 887 | O88508 SEPQPEEGS\*PAAGQK | -0.7968379 | 5.781389 | 6.955553e-04 | 4.652037e-02 |
    | 509 | Q3UHB8 SFS\*LGDLS\*HSPQTAQHVER | 0.6229243 | 7.668709 | 9.319972e-04 | 5.593092e-02 |
    | 1086 | P28740 ARPSQLPEQSSSAQQNGSVSDIS\*PVQAAK | 0.8723021 | 4.479188 | 9.346416e-04 | 5.593092e-02 |
    | 1064 | Q8BZ98 S\*PPPSPTTQR | 0.8878245 | 4.661802 | 9.947677e-04 | 5.655254e-02 |
    | 337 | P16858 DGRGAAQNIIPAS\*TGAAK | -0.6597538 | 8.809705 | 1.090669e-03 | 5.905195e-02 |
    | 346 | Q8CJ40 SADRRLS\*GAQAELALQEESVR | 0.4803619 | 8.718833 | 1.273092e-03 | 6.579568e-02 |
    | 238 | Q99P72 GPLPAAPPTAPERQPS\*WER | 0.5536111 | 9.488092 | 1.410821e-03 | 6.974363e-02 |
    | 169 | Q8K2P7 S\*LTNSHLEKR | 0.6158267 | 10.027889 | 1.603114e-03 | 7.594753e-02 |

$adjust.method
:   'BH'

$comparison
:   1. 'WT'
    2. 'KO'

$test
:   'exact'

#### Check MA plot and p-value distribution¶

In [16]:

```
# make an MD plot
plotMD(etp, p.value = 0.10 , main = "Phosphopeptide: KO vs WT")
abline(h = c(-1, 1), col = "black")

# check the p-value distribution
ggplot(ttp, aes(PValue)) + 
  geom_histogram(bins = 100, fill = "white", color = "black") + 
  geom_hline(yintercept = mean(hist(etp$table$PValue, breaks = 100, 
                                    plot = FALSE)$counts[26:100])) +
  ggtitle("Phosphopeptide: KO vs WT p-value distribution")
```

#### Can also do some more candidate visualizations¶

In [17]:

```
# get the averages within each condition
ttp$ave_WT <- rowMeans(data_norm_pep[WT])
ttp$ave_KO <- rowMeans(data_norm_pep[KO])

# add the cadidate status column
ttp <- ttp %>%
  mutate(candidate = cut(FDR, breaks = c(-Inf, 0.01, 0.05, 0.10, 1.0),
  labels = c("high", "med", "low", "no")))

# make the DE plots
de_plots(ttp, "ave_WT", "ave_KO", "Phosphopeptide: WT versus KO")

# label columns so we know what the comparison was
colnames(ttp) <- str_c(colnames(ttp), '_WT_KO')
```

#### Export the phophopeptide results and log session¶

In [18]:

```
# export the results: TMM normed data, 
#   statistical test results (includes gene names), averages, and candidate status
data_export_pep <- cbind(data_norm_pep, ttp)
write.table(data_export_pep, "phosphopeptide_results.txt", sep = "\t", row.names = FALSE)
```

In [19]:

```
# log the session
sessionInfo()
```

```
R version 3.5.0 (2018-04-23)
Platform: x86_64-apple-darwin15.6.0 (64-bit)
Running under: macOS  10.14.1

Matrix products: default
BLAS: /Library/Frameworks/R.framework/Versions/3.5/Resources/lib/libRblas.0.dylib
LAPACK: /Library/Frameworks/R.framework/Versions/3.5/Resources/lib/libRlapack.dylib

locale:
[1] en_US.UTF-8/en_US.UTF-8/en_US.UTF-8/C/en_US.UTF-8/en_US.UTF-8

attached base packages:
[1] stats     graphics  grDevices utils     datasets  methods   base     

other attached packages:
 [1] bindrcpp_0.2.2  psych_1.8.10    edgeR_3.22.2    limma_3.36.3   
 [5] forcats_0.3.0   stringr_1.3.1   dplyr_0.7.8     purrr_0.2.5    
 [9] readr_1.1.1     tidyr_0.8.2     tibble_1.4.2    ggplot2_3.1.0  
[13] tidyverse_1.2.1

loaded via a namespace (and not attached):
 [1] pbdZMQ_0.3-3         locfit_1.5-9.1       tidyselect_0.2.5    
 [4] repr_0.17            splines_3.5.0        haven_1.1.2         
 [7] lattice_0.20-38      colorspace_1.3-2     htmltools_0.3.6     
[10] base64enc_0.1-3      rlang_0.3.0.1        pillar_1.3.0        
[13] foreign_0.8-71       glue_1.3.0           withr_2.1.2         
[16] modelr_0.1.2         readxl_1.1.0         uuid_0.1-2          
[19] bindr_0.1.1          plyr_1.8.4           munsell_0.5.0       
[22] gtable_0.2.0         cellranger_1.1.0     rvest_0.3.2         
[25] evaluate_0.12        labeling_0.3         parallel_3.5.0      
[28] broom_0.5.0          IRdisplay_0.6        Rcpp_1.0.0          
[31] scales_1.0.0         backports_1.1.2      IRkernel_0.8.12.9000
[34] jsonlite_1.5         mnormt_1.5-5         hms_0.4.2           
[37] digest_0.6.18        stringi_1.2.4        grid_3.5.0          
[40] cli_1.0.1            tools_3.5.0          magrittr_1.5        
[43] lazyeval_0.2.1       crayon_1.3.4         pkgconfig_2.0.2     
[46] xml2_1.2.0           lubridate_1.7.4      assertthat_0.2.0    
[49] httr_1.3.1           rstudioapi_0.8       R6_2.3.0            
[52] nlme_3.1-137         compiler_3.5.0
```

###### Notebook prepared by Phil Wilmarth, OHSU, 11/23/2018¶
