## Supplementary material for "Identification of PKCα-dependent phosphoproteins in mouse retina": S3 - Annotated Phosphopeptide Spectra

Supplement-3\_Annotated-Spectra


### Phosphopeptide fragment ion spectrum plotter script¶

##### Phil Wilmarth, OHSU¶

##### March 1, 2019¶

In support of the data analysis for the MORG-339 project.

Python 3 script uses the spectrum\_utils package for spectrum annotations. The input data is a phosphopeptide combined summary table from the PAW pipeline. This version of the script uses MGF files created by MSConvert.

#### Import libraries¶

In [1]:

```
# notebook for reading in PAW phosphopeptude results file and plotting spectra
# uses https://github.com/bittremieux/spectrum_utils.git

# imports
import os
import numpy as np
import inspect
from pyteomics import mgf
import matplotlib.pyplot as plt
from spectrum_utils import plot
from spectrum_utils import spectrum

# set some plotting parameters
plt.rc('text', usetex=True)
plt.rc('font', family='sans-serif', size=14)
```

#### Read in the combined phosphopeptide summary table¶

In [2]:

```
# get the combined results table
with open("MORG-339_phosphopeptide_TMT_8%ACN_032017_OT_psm_filtered_combined.txt", 'rt') as fin:
    buffer = fin.readlines()

# strip EOL, get header map from first row, trim bottom of table
buffer = [x.strip() for x in buffer]    
col_map = {header: i for (i, header) in enumerate(buffer[0].split('\t'))}
buffer = [x for x in buffer if len(x.split('\t')) == len(col_map)]
```

#### Make an object (Class) to hold some of the information¶

Each row has 60 items. We only need a few for plotting the spectrum. We will need to do some wrangling of the information and we will do that with some bound methods. PhosphoRS can have localization ties, so there can be some ambiguity of where to put the phospho group. We will just take the first residue in those cases.

We have to specify static mods like the TMT tags and alkylated Cys residues in order to get the correct fragment ion masses. The spectrum\_utilities does not have explicit static modifications, so we have to do them like variable modifications. We have oxidized Met and phosphorylation of STY as variable mods. The N-term is complicated. We will have a TMT tag and the first residue could be modified. We have to add the masses for those cases.

In [3]:

```
# define an object to hold some values from each row
class PlotMe:
    """Container for some row attributes."""
    def __init__(self, col_map, row):
        self.moverz = float(row.split('\t')[col_map['m/z [Da]']])
        self.MHplus = float(row.split('\t')[col_map['MH+ [Da]']])
        self.charge = row.split('\t')[col_map['Charge']]
        self.first_scan = row.split('\t')[col_map['First Scan']]
        self.last_scan = row.split('\t')[col_map['Last Scan']]
        self.spectrum_file = row.split('\t')[col_map['Spectrum File']]
        self.sequence = row.split('\t')[col_map['Group String']].split('_')[0]
        self.PD_mods = row.split('\t')[col_map['New Modifications']]
        self.new_sequence = ''
        self.sites = int(row.split('\t')[col_map['Number Phospho Sites']])
        self.probabilities = row.split('\t')[col_map['New Site Prob Peptide']]
        self.status = row.split('\t')[col_map['Localization Status']]
        self.psms = len(row.split('\t')[col_map['Grouped PSMs']].split(';'))
        self.nlpeak = self.moverz - (97.9769 / float(self.charge))
        self.mods = {}
        self.set_static_mods()
        self.set_variable_mods()
        
    def _print(self):
        """Diagnostic print of attributes."""
        for i in inspect.getmembers(self):
            # from https://stackoverflow.com/questions/2675028/list-attributes-of-an-object/31967014
            # Ignores non-simple attributes
            if i[0].startswith('_') or inspect.ismethod(i[1]) or isinstance(i[1], (list, tuple)):
                continue
            print(i)
            
    def set_static_mods(self):
        """Figures out modifications and alters peptide string."""
        # alykylation of Cys: 57.0215
        TMT = 229.1629 # TMT tags on N-term and K
        self.new_sequence = ''
        for i, char in enumerate(self.sequence):
            if i == 0:
                self.mods['N-term'] = TMT
                char = char.lower()
            if char in 'CK':
                if char == 'C':
                    self.mods[i] = 57.0215
                else:
                    self.mods[i] = TMT
                char = char.lower()
            self.new_sequence += char

    def set_variable_mods(self):
        """Parses variable modifications."""
        # phosphorylation of STY:79.9663
        # oxidation of Met: 15.9949
        # see if oxidized Met
        for mod in self.PD_mods.split('; '):            
            if 'Oxidation' in mod:
                site = int(mod.split('(')[0][1:])
                if site == 1:
                    if 'N-term' in self.mods:
                        self.mods['N-term'] += 15.9949
                else:
                    self.mods[site-1] = 15.9949
                self.new_sequence = (self.new_sequence[:site-1] + self.new_sequence[site-1].lower() +
                                     self.new_sequence[site:])
                
        # Phospho sites
        for site_prob in self.probabilities.split(';'):
            site = site_prob.split(':')[0]
            if ',' not in site:                
                residue = int(site.split()[1][1:-1])
            else:
                residue = int(site.split('(')[1].split(',')[0])
                
            if residue == 1:
                if 'N-term' in self.mods:
                    self.mods['N-term'] += 79.9663
            else:
                self.mods[residue-1] = 79.9663
                
            self.new_sequence = (self.new_sequence[:residue-1] + self.new_sequence[residue-1].lower() +
                                 self.new_sequence[residue:])
```

#### Load all of the table rows into PlotMe objects¶

In [4]:

```
# make a list of row objects
rows = []
for i, line in enumerate(buffer[1:]):
    rows.append(PlotMe(col_map, line))
```

#### Peptide strings are reformatted to indicate modifications¶

We have both static and variable modifications to indicate. TMT tags will be at the peptide N-terminus and at all lysine residues. All cysteine residues have been alkylated. Phosphorylation can be located at serine, threonine, or tyrosine residues. Modified residues will be in lower case and unmodified residues in upper case.

In the peptide below, the first "E" is lower case because of the TMT tag and the "S" (third residue from the C-terminus) is phosphorylated.

In [5]:

```
rows[100].sequence, rows[100].new_sequence
```

Out[5]:

```
('EEHGGLIRSPR', 'eEHGGLIRsPR')
```

#### Gets the data into an MsmsSpectrum object¶

In [6]:

```
def load_spectrum(mgf_in, row):
    """Loads an MsmsSpectrum object."""
    # some parsed items from the MGF file
    mz = mgf_in['m/z array']
    intensity = mgf_in['intensity array']
    retention_time = float(mgf_in['params']['rtinseconds'])
    precursor_mz = mgf_in['params']['pepmass'][0]
    precursor_charge = mgf_in['params']['charge'][0]
    identifier = mgf_in['params']['title'].split()[0]
    
    # some data from the row
    peptide = row.sequence
    modifications = row.mods

    spec = spectrum.MsmsSpectrum(identifier, precursor_mz, precursor_charge,
                                 mz, intensity, retention_time=retention_time,
                                 peptide=peptide, modifications = modifications)

    # set some plot attributes
    min_mz, max_mz = 0, 2000
    fragment_tol_mass, fragment_tol_mode = 0.5, 'Da'
    min_intensity, max_num_peaks = 0.001, 150
    scaling = 'root'
    ion_types = 'aby'

    (spec.set_mz_range(min_mz, max_mz)
         .remove_precursor_peak(fragment_tol_mass, fragment_tol_mode)
         .filter_intensity(min_intensity, max_num_peaks)
         .scale_intensity(scaling)
         .annotate_peaks(fragment_tol_mass, fragment_tol_mode, ion_types)
         .set_mz_range(min_mz, max_mz))
    
    return spec
```

#### Iterate over rows and make plots¶

The plots have titles that contain the RAW file base name, starting scan number, ending scan number, and peptide charge. That is also used for the PDF filenames. The subtitle is the peptide string along with the observed m/z value (Da) and the peptide charge state.

Y-ions (within the 05 Da tolerance) are highlighted in red, b-ions in blue, a-ions in green, and labels are placed at the locations where precursor neutral loss of 98 Da would be. The labels are always at the 80% line. The labels contain the expected m/z location.

The spectra have the intensities scaled with a square root transformation. This has the effect of attenuating the neutral loss peaks so that the other peaks can be more easily seen. The spectra are numbered to match the rows in the summary table in the `MORG-339_phosphopeptide_TMT_8%ACN_032017_OT_psm_filtered_combined.txt` file.

We will also print a short table with the masses of the annotated peaks. The tolerance for matching is 0.5 Da. The spectrum has been filtered to 150 peaks.

In [7]:

```
def print_annotations(spec):
    """Prints short annotation mass table"""
    print('\n   Ion         m/z   Intensity')
    for triple in [x for x in zip(spec.mz, spec.intensity, spec.annotation) if x[2] != None]:
        mz = triple[0]
        intensity = triple[1]
        ion = triple[2].ion_type
        index = triple[2].ion_index
        charge = int(triple[2].charge)
        label = '%s%s%s' % (ion, index, charge * '+')
        print('   %-8s %7.2f %10.1f' % (label, mz, intensity))
```

In [8]:

```
for i, row in enumerate(rows):
    mgf_file = os.path.abspath('../mgf_files/' + row.spectrum_file.replace('.raw', '.mgf'))
    
    # this is slow to open and close each MGF file for each spectrum
    # should sort row objects by RAW file and access the MGF files once
    with mgf.MGF(mgf_file) as f:
        row.title = ".".join([row.spectrum_file[:-4], row.first_scan, 
                              row.last_scan, row.charge])
        row.mgf_title = row.title + (' File:"%s", NativeID:"controllerType=0 controllerNumber=1 scan=%s"' % 
                 (row.spectrum_file, row.first_scan))
        mgf_in = f[row.mgf_title]
    
    # load spectrum object
    ms_spec = load_spectrum(mgf_in, row)

    # build plot title and nl label
    row.plot_title = row.title.replace('_', '\_')
    row.plot_title = row.plot_title.replace('%', '\%')
    title = '%d. %s\n %s (%0.2f, %s+)' % (i+1, row.plot_title, row.new_sequence, row.moverz, row.charge)
    label = '%0.2f - neutral loss' % row.nlpeak
    xpos = row.nlpeak + 10.0

    # create the plot
    print('%d. %s (%s)' % (i+1, row.title, row.new_sequence))
    print_annotations(ms_spec)
    plt.figure(figsize=(14, 8))
    plt.suptitle(title)
    plot.spectrum(ms_spec)
    plt.text(xpos, 0.81, label)
    plt.grid(True)
    plt.savefig(os.path.join('./plots', row.title + '.pdf'))
    plt.show()
    plt.close()
```

```
1. MORG-339_phosphopeptide_TMT_25%ACN_032017_OT.3429.3429.4 (kkEPAISSQNsPEAR)

   Ion         m/z   Intensity
   a5+++     328.55       71.2
   b4++      471.47       62.1
   y8++      484.94       75.0
   y9++      528.70       69.1
   b6++      563.23       60.3
   b11+++    570.48      228.6
   y10++     585.24      165.8
   a12+++    593.41       81.5
   y14+++    608.56      217.9
   y11++     620.19      109.6
   y5+       639.32       99.8
   b14+++    669.12       69.0
   a9++      700.33       60.9
   b9++      714.62       61.1
   y6+       753.37      169.9
   y14++     911.90       82.3
   y8+       968.44      104.7
   y9+      1055.41      125.4
   y12+     1336.63       73.8
```

```
2. MORG-339_phosphopeptide_TMT_10%ACN_032017_OT.8252.8252.3 (kLEkEEEEGISQEsSEEEQ)

   Ion         m/z   Intensity
   a8++      723.22       22.4
   y6+       788.63       23.1
   a11++     852.32       27.0
   y15++     909.43       17.6
   a4+       929.25       18.8
   b4+       958.05       40.9
   a13++     980.41       24.1
   y8+      1044.84       28.5
   y9+      1132.44       23.0
   a6+      1187.86       23.7
   a18++    1301.08       16.4
   y11+     1302.46       18.0
   a7+      1316.32       30.6
   y12+     1431.82       14.0
```

```
3. MORG-339_phosphopeptide_TMT_30%ACN_032017_OT.12764.12764.3 (kAsGPPVSELITk)

   Ion         m/z   Intensity
   y1+       376.32      134.4
   y2+       477.40      268.2
   a3+       568.65       45.0
   y3+       590.50      239.9
   y9++      607.05      143.8
   a10++     624.28       46.7
   y10++     635.62       63.7
   b10++     638.56      127.2
   a11++     680.50       65.2
   y4+       703.84      265.8
   y12++     754.36       56.3
   y5+       832.57      163.3
   a7+       918.75       69.1
   y6+       919.65      289.5
   y8+      1115.60      101.3
   y9+      1212.75      136.2
   y10+     1269.85      193.1
   y11+     1436.68       45.6
```

```
4. MORG-339_phosphopeptide_TMT_18%ACN_032017_OT.5246.5246.3 (vkVDGPRsPsYGR)

   Ion         m/z   Intensity
   y3+       395.32       67.4
   b4++      450.99      408.1
   y4+       562.29       77.8
   y9++      568.86      123.5
   b7++      606.10       99.7
   y10++     626.40       82.5
   a2+       658.01       58.3
   y5+       659.52      162.8
   b8++      689.26       57.8
   a9++      723.47       71.1
   y12++     854.13      118.8
   b4+       900.58      242.5
   b11++     902.74       65.3
   b12++     931.37       44.6
   y9+      1136.43      217.8
```

```
5. MORG-339_phosphopeptide_TMT_25%ACN_032017_OT.4667.4667.3 (kkEPAIsSQNsPEAR)

   Ion         m/z   Intensity
   y2+       246.22       36.2
   y4+       472.45      121.2
   y5+       639.44       30.1
   y6+       753.48       53.1
   b9++      754.37       34.5
   y13++     772.94       40.6
   a10++     797.45       74.4
   b10++     811.21       61.9
   a3+       816.44       45.2
   y14++     951.85       49.7
   y8+       968.42       58.4
   y9+      1135.40       37.2
   y10+     1248.56       26.1
   y11+     1319.61       42.7
   y12+     1416.58       51.4
```

```
6. MORG-339_phosphopeptide_TMT_10%ACN_032017_OT.4001.4001.3 (sRsPsPAPEkk)

   Ion         m/z   Intensity
   y2++      367.30       42.0
   y1+       376.36       31.2
   a5++      478.84       22.1
   y4++      480.55      124.2
   y5++      516.05       22.8
   b2+       553.38       26.4
   y6++      564.55      153.6
   b7++      576.54       21.8
   y7++      647.45       97.3
   a9++      675.35       22.7
   a3+       692.12       28.3
   y8++      696.25       37.4
   b3+       720.22       27.8
   y2+       733.55       23.8
   y9++      779.70       35.9
   a4+       789.49       30.6
   a10++     854.73       39.1
   y10++     858.10       41.6
   b10++     868.55       48.4
   y4+       959.64       49.7
   b5+       984.43       22.4
   y6+      1127.81       26.1
   b7+      1152.38       58.0
   b9+      1378.63       21.3
```

```
7. MORG-339_phosphopeptide_TMT_10%ACN_032017_OT.9742.9742.3 (kLEkEEEEGISQEssEEEQ)

   Ion         m/z   Intensity
   a6++      594.01       21.1
   a7++      659.03       19.2
   a8++      723.63       21.7
   a11++     851.88       41.1
   a12++     916.05       26.0
   y15++     950.12       14.8
   b4+       957.56       35.6
   a13++     980.47       64.2
   b13++     994.93       20.0
   y7+       996.97      144.6
   y8+      1125.05       17.2
   y9+      1212.29       20.2
   y11+     1382.44       19.4
```

```
8. MORG-339_phosphopeptide_TMT_25%ACN_032017_OT.6991.6991.4 (sAsSPkPDTkVPQATAEAk)

   Ion         m/z   Intensity
   y1+       376.45       48.3
   b2+       388.39       43.6
   y2+       447.56       79.9
   y10+++    500.64       87.6
   y8++      523.04       57.9
   a3+       527.52       62.5
   a6++      535.10       72.1
   b6++      549.13      253.1
   y3+       576.49       75.3
   b10+++    590.15       81.6
   b7++      597.22       47.5
   a11+++    612.98      100.1
   b11+++    623.04      120.9
   b4+       642.42       62.0
   y4+       647.58       78.8
   b12+++    654.88      113.2
   b13+++    698.06       89.5
   a14+++    711.76       61.4
   a15+++    746.10       50.6
   y5+       748.61       99.0
   y10++     751.17      132.3
   y15+++    757.22      863.5
   b16+++    779.11       81.9
   y11++     801.78      261.0
   y6+       819.56       91.5
   b17+++    822.04      105.4
   y17+++    841.79       53.6
   b18+++    845.79       66.5
   b10++     884.46       61.2
   y13++     907.80      396.8
   a11++     919.48      129.6
   b11++     933.85       79.6
   y7+       947.67       46.5
   y8+      1044.62      154.9
   b13++    1045.62      115.3
```

```
9. MORG-339_phosphopeptide_TMT_10%ACN_032017_OT.10218.10218.4 (gkQADVADQQTTELPAENGETENQsPAsEEEk)

   Ion         m/z   Intensity
   y1+       376.35       91.4
   b5++      479.92       60.1
   y6++      501.01       95.7
   y2+       505.42       86.4
   y7++      549.84      112.8
   a7++      550.47       61.4
   b7++      565.13       59.5
   b8++      622.41       66.5
   y13+++    623.12       65.4
   y8++      633.55       98.9
   y3+       634.45       97.6
   b14+++    648.46       54.8
   b9++      686.59       60.5
   y9++      697.34       60.8
   b10++     750.80      103.4
   y10++     754.30       91.6
   y4+       763.52       74.7
   b11++     801.22       99.4
   y11++     819.02       69.3
   b12++     851.80       98.8
   y12++     869.37       91.6
   b13++     915.97      194.2
   a5+       930.45      103.3
   a23+++    953.49       59.7
   b5+       958.70       56.7
   y14++     962.30       81.0
   b23+++    962.90       97.3
   b14++     972.78      457.5
   y6+      1001.61       91.2
   b24+++   1005.54      198.3
   y15++    1019.70       96.4
   a25+++   1051.12       68.6
   b16++    1056.79      122.6
   b6+      1057.65       89.7
   y7+      1098.54      112.7
   y17++    1119.22      120.3
   b17++    1121.36      168.4
   y18++    1167.85      179.7
   y29+++   1169.68      109.7
   b18++    1178.40      174.2
   b19++    1206.94       87.2
   a8+      1215.85       66.0
   y19++    1224.76      141.7
   b8+      1243.74       55.2
   y8+      1265.52      149.5
   b20++    1271.48      210.1
   y20++    1289.37       66.3
   b31+++   1301.88       61.9
   b21++    1321.95      144.9
   b22++    1386.47      219.0
   b23++    1443.60      145.7
   b10+     1499.94       55.2
   y10+     1507.58      172.3
```

```
10. MORG-339_phosphopeptide_TMT_14%ACN_032017_OT.3887.3887.3 (kEPAISSQNsPEAR)

   Ion         m/z   Intensity
   y2+       246.12       50.3
   b1+       358.71       58.0
   b6++      428.69       27.9
   y4+       472.27      121.4
   y11++     620.70       17.8
   y5+       639.09       21.5
   b4+       655.38       23.8
   a10++     662.31       28.3
   y12++     668.66       48.4
   b10++     675.98       45.4
   b11++     724.65       18.3
   y13++     733.02       59.4
   y6+       753.39       24.1
   b6+       855.14       93.3
   y7+       881.43       22.8
   a7+       914.53       18.5
   y8+       968.41       25.1
   y9+      1055.39       31.9
   y10+     1168.51       20.1
   y11+     1239.61       35.1
   y12+     1336.61       42.7
```

```
11. MORG-339_phosphopeptide_TMT_10%ACN_032017_OT.12563.12563.3 (dIIRQPsEEEIIk)

   Ion         m/z   Intensity
   b1+       345.22       11.8
   a4++      350.14        9.4
   b4++      364.18       15.5
   y1+       376.31       11.7
   b5++      428.52       33.3
   a2+       430.24       10.0
   b2+       458.39        8.4
   y2+       489.35       23.7
   b3+       571.41       11.2
   y3+       602.51       25.9
   a9++      674.91       11.9
   b9++      689.05       27.2
   y9++      690.88       24.4
   b4+       727.61       10.4
   y4+       731.61       17.8
   b10++     753.73       19.9
   y10++     769.43       12.8
   a11++     796.68       16.5
   b11++     810.70       23.6
   b5+       855.55       11.2
   y5+       860.54       11.3
   b12++     867.25       22.1
   y6+       989.66       17.2
   y7+      1156.77       16.2
   y8+      1253.62       17.9
```

```
12. MORG-339_phosphopeptide_TMT_12%ACN_032017_OT.4755.4755.3 (sksPPPPEEEAk)

   Ion         m/z   Intensity
   b1+       317.31       11.5
   a2++      323.80       14.9
   b2++      337.85       16.5
   y1+       376.37       16.7
   a3++      406.97       11.9
   b3++      421.09       34.3
   y2+       447.43       25.8
   a4++      455.45       26.3
   y6++      466.17       13.9
   b4++      469.53       26.7
   a5++      504.19       21.1
   y7++      514.99       58.1
   b5++      518.50       22.9
   a6++      552.94       14.7
   y8++      563.45       40.0
   y3+       576.43       22.6
   y9++      612.03      143.5
   b7++      615.39       11.7
   b2+       674.59       18.9
   b8++      679.53       27.7
   y10++     695.49       21.1
   y4+       705.45       26.0
   b9++      744.51       57.0
   b10++     808.76       22.7
   y5+       834.48       16.1
   b3+       841.49       20.2
   b11++     844.62       18.1
   y6+       931.52       36.3
   y7+      1028.53       42.9
   y8+      1125.61       24.8
   y9+      1222.65       75.4
```

```
13. MORG-339_phosphopeptide_TMT_10%ACN_032017_OT.6992.6992.3 (asSPPVEVGEPR)

   Ion         m/z   Intensity
   y1+       175.13        7.3
   y2+       272.18       33.8
   a1+       273.08       19.7
   b3++      277.73        6.3
   y5++      279.18        7.9
   b4++      326.40        8.0
   y6++      343.86       26.8
   a5++      361.41       18.2
   b5++      375.54       25.9
   y3+       401.24       21.9
   a6++      410.66       35.1
   b6++      424.94       69.1
   a2+       440.51       44.0
   y8++      441.45       16.8
   y4+       458.26       71.5
   b2+       468.09       10.3
   a7++      474.91       10.0
   b7++      489.53       73.3
   y9++      490.41       43.5
   a8++      525.05       30.9
   b8++      539.01       93.3
   b3+       555.30       13.4
   y5+       557.33       47.6
   b9++      567.43       40.3
   a10++     617.93        8.7
   b10++     632.09       46.0
   b4+       652.38        8.5
   b11++     680.47        8.0
   y6+       686.38       55.1
   b5+       749.41       10.7
   y7+       785.41       10.7
   b6+       848.40       30.7
   b7+       977.43       26.2
   y9+       979.50       12.9
```

```
14. MORG-339_phosphopeptide_TMT_20%ACN_032017_OT.2436.2436.3 (gAkEEHGGLIRsPR)

   Ion         m/z   Intensity
   y2+       272.15       27.5
   b4++      423.20       17.4
   b5++      487.50       39.0
   y9++      536.58       14.4
   a6++      542.11       12.9
   b6++      555.69       18.7
   b7++      584.60       19.7
   y4+       595.16       14.6
   a8++      598.52       23.0
   y10++     601.43       14.1
   y11++     665.98       19.7
   b9++      669.83       50.1
   a3+       687.24       11.5
   a10++     712.24       15.1
   b10++     726.14       24.0
   b11++     803.91       26.3
   a4+       816.83       11.6
   b4+       844.48       22.7
   y7+       878.57       12.5
   y13++     879.80       13.4
   b12++     887.59       17.3
   b13++     935.79       16.8
   a5+       945.78       25.7
   b5+       973.54       20.7
   y9+      1072.55       14.6
   y10+     1201.71       14.0
```

```
15. MORG-339_phosphopeptide_TMT_4%ACN_032017_OT.5447.5447.3 (dkEVsDDEAEEk)

   Ion         m/z   Intensity
   b1+       345.25       11.6
   y1+       376.24       14.8
   b3++      416.27       19.1
   b4++      465.81       21.0
   y2+       505.47       16.7
   b6++      606.95       27.7
   y8++      616.14       15.7
   y3+       634.44       20.4
   b7++      664.29       33.4
   b2+       702.48       15.4
   y4+       705.50       25.3
   a8++      715.30       11.1
   b8++      728.99        8.8
   a9++      750.38        7.4
   b9++      764.29       18.3
   b10++     829.19       19.5
   y5+       834.53       31.5
   b11++     893.60       14.8
   a4+       902.60        9.8
   y11++     909.37       26.4
   y6+       949.55       30.9
   y7+      1064.53       17.8
```

```
16. MORG-339_phosphopeptide_TMT_12%ACN_032017_OT.3486.3486.3 (kAEGAGTEEEGtPk)

   Ion         m/z   Intensity
   y1+       376.54       43.0
   b7++      423.03       25.0
   y2+       473.72      250.5
   b8++      487.15       37.9
   a9++      537.60       56.0
   a4+       587.78       38.7
   a10++     602.16      106.7
   y3+       654.56       22.5
   a5+       658.61       31.8
   b5+       686.14      165.2
   y4+       711.43       49.9
   b6+       743.60       23.4
   a13++     769.83       32.0
   y13++     792.67       72.5
   y5+       840.67       31.9
   b7+       844.56       28.7
   y6+       969.42       27.2
   b8+       973.75       30.6
   a10+     1203.70       22.9
   y9+      1256.64       21.8
   y11+     1384.65       26.3
```

```
17. MORG-339_phosphopeptide_TMT_20%ACN_032017_OT.5448.5448.3 (kEVQsPEQVk)

   Ion         m/z   Intensity
   a2++      230.20       16.0
   b1+       358.08       15.2
   y1+       376.33       43.5
   y5++      415.28       34.1
   a6++      475.40       44.3
   y7++      562.75       28.6
   b3+       586.64       26.7
   y3+       603.39       57.7
   a8++      604.44       33.7
   y8++      612.09       21.8
   b8++      618.40       36.6
   b4+       714.60       20.8
   y4+       732.78       47.9
   y5+       829.52      163.3
   b5+       881.48       17.4
   b6+       978.52       24.9
   y6+       996.57       24.1
   b7+      1107.63       16.0
```

```
18. MORG-339_phosphopeptide_TMT_10%ACN_032017_OT.4211.4211.3 (sRsPPPVSk)

   Ion         m/z   Intensity
   a2++      262.84       17.4
   y4++      330.28       34.3
   b3++      360.33       26.3
   y1+       376.37       19.1
   y5++      378.82       55.9
   a4++      395.54       10.4
   b1+       396.96       14.1
   b4++      409.38       16.3
   y6++      427.42      159.9
   a5++      443.59       16.5
   b5++      457.94       40.8
   y2+       463.34       48.4
   a6++      492.44      318.1
   b6++      506.59       55.2
   y7++      510.60       19.3
   a7++      541.62       25.1
   b7++      555.84       43.3
   y3+       562.40       24.0
   y8++      588.89       10.2
   y4+       659.43       51.8
   b3+       720.36       39.0
   y5+       756.49       27.0
   b4+       817.13       15.6
   y6+       853.57       28.8
```

```
19. MORG-339_phosphopeptide_TMT_6%ACN_032017_OT.9766.9766.4 (eSDDkPEIEDVGsDEEEEEk)

   Ion         m/z   Intensity
   y3++      317.45       13.5
   y1+       376.29       48.7
   b6+++     377.35       17.8
   y4++      382.31       11.2
   y5++      446.63       12.6
   y10+++    487.34       17.8
   b9+++     501.48       12.6
   y2+       505.37       56.9
   y6++      511.24       10.2
   b5++      517.39       56.1
   b3+       561.44       11.9
   b11+++    572.85       14.7
   a7++      616.44       17.0
   b7++      630.54       32.9
   y3+       634.43       47.4
   a8++      672.51       11.6
   a14+++    676.36       15.5
   y9++      681.07       16.1
   b8++      687.14       39.6
   b15+++    728.45       14.3
   y10++     730.51       18.0
   a9++      737.36       10.8
   b9++      751.40       49.4
   y4+       763.41       19.4
   b10++     809.14       41.0
   b17+++    814.81       33.6
   a18+++    848.78       20.8
   y12++     851.99       20.1
   b18+++    857.78       47.4
   b11++     858.65       84.9
   y18+++    877.24       12.8
   b12++     887.14       34.5
   y5+       892.47       11.9
   b19+++    901.05       47.0
   y19+++    906.62       11.7
   y14++     973.29       11.3
   y15++    1022.15        9.9
   b15++    1092.77       52.2
   b16++    1157.25       68.2
   y16++    1200.39       13.6
   a17++    1207.45       10.4
   b17++    1221.81       49.2
   b18++    1286.49       15.7
```

```
20. MORG-339_phosphopeptide_TMT_6%ACN_032017_OT.1889.1889.3 (rEsPsPAPkPR)

   Ion         m/z   Intensity
   y2+       272.22       12.5
   y4++      363.91       37.5
   a4++      375.84       16.3
   b1+       386.37       19.6
   y6++      447.85       70.9
   a5++      459.94       12.5
   a2+       487.62       11.8
   b2+       515.42       24.7
   b6++      522.19       21.6
   y7++      530.88       57.0
   b7++      557.68       17.5
   y8++      580.16       58.5
   a8++      592.49       16.0
   y3+       629.92       15.8
   a3+       654.40       12.1
   y9++      663.52       22.0
   b3+       682.38       19.1
   y4+       726.52       24.0
   y10++     727.50       14.3
   a4+       751.43       14.4
   b9++      785.03       32.7
   y5+       797.62       12.0
   b10++     833.68       13.6
   y6+       894.57       31.7
   b5+       946.46       10.7
   b7+      1114.44       18.5
```

```
21. MORG-339_phosphopeptide_TMT_10%ACN_032017_OT.2040.2040.3 (rGsRSsIEPk)

   Ion         m/z   Intensity
   y1+       376.41       34.7
   b4++      383.96       50.5
   b1+       386.39       39.5
   a5++      412.91       55.9
   b5++      427.35       42.3
   y2+       473.43      183.9
   b6++      510.79       49.9
   a7++      553.48       66.5
   y7++      563.19       54.6
   b7++      567.25       89.8
   y3+       602.42       90.1
   a8++      617.74       38.3
   b8++      631.78      168.4
   y9++      675.38       36.4
   y4+       715.51       92.9
   y5+       882.51       47.1
   y6+       969.58       37.7
```

```
22. MORG-339_phosphopeptide_TMT_10%ACN_032017_OT.4794.4794.4 (eEDEEAEsPPEkk)

   Ion         m/z   Intensity
   y4+++     320.74        9.6
   a1+       330.78       12.9
   y5+++     353.12       45.4
   b7+++     354.02       10.8
   y2++      366.97       10.2
   y1+       376.16       17.4
   y6+++     408.64       19.4
   b5++      431.11       15.4
   y3++      431.83       14.3
   b9+++     442.35        9.7
   y7+++     451.84       24.7
   a6++      453.07       10.9
   b6++      466.79       29.4
   y8+++     475.40       25.7
   y4++      480.51       20.8
   y9+++     518.41       17.3
   y5++      529.10       37.9
   b7++      531.49       23.1
   y10+++    561.52       21.6
   y11+++    599.98       12.2
   b3+       602.95       11.6
   y6++      612.67       30.9
   b8++      614.60       14.0
   b12+++    636.90       16.5
   y7++      677.15       26.3
   y8++      712.71       21.4
   b4+       732.36       17.7
   y2+       733.56       10.1
   y9++      776.99       10.7
   b5+       861.39       21.9
   y3+       862.34       23.4
   b6+       932.39       19.1
   b7+      1061.47       11.6
```

```
23. MORG-339_phosphopeptide_TMT_18%ACN_032017_OT.8484.8484.4 (fSSQQASTkQSNASsDVEVEEk)

   Ion         m/z   Intensity
   y4++      367.41       10.7
   y1+       376.39       20.1
   y2+       505.34       29.1
   y11+++    506.39       12.1
   y3+       634.42       34.4
   b15+++    693.28       11.2
   b9++      712.32       17.1
   a16+++    722.30       21.4
   y15+++    730.36       10.7
   b16+++    731.38       12.8
   y4+       733.44       22.2
   a17+++    755.76       33.3
   y11++     758.12       18.9
   y16+++    759.17       30.4
   a10++     762.31       23.3
   b17+++    764.91       44.9
   b10++     776.79       30.2
   a5+       779.64       27.2
   a18+++    798.63       13.3
   y12++     802.25       21.3
   b18+++    807.74       23.0
   b11++     820.19       17.9
   a19+++    831.76       13.5
   b19+++    841.03       49.0
   a6+       850.15       12.2
   a12++     862.53       21.1
   y13++     865.67       15.7
   b12++     877.14       20.1
   b6+       878.14       20.6
   b20+++    883.89       39.8
   y20+++    897.63       13.4
   b13++     912.78       21.8
   a21+++    917.48       15.1
   y21+++    926.06       11.0
   b21+++    927.05       33.4
   b14++     956.36       12.1
   b16++    1097.25       15.3
   b17++    1146.74       33.6
   b18++    1211.24       33.4
   b19++    1261.10       16.2
```

```
24. MORG-339_phosphopeptide_TMT_30%ACN_032017_OT.9766.9766.3 (rktSGPPVSELITk)

   Ion         m/z   Intensity
   a1+       358.48       29.9
   y1+       376.34       39.5
   y2+       477.41       63.1
   b4++      506.33       30.7
   b5++      534.84       30.7
   y3+       590.41       59.8
   y9++      606.97       32.7
   a8++      667.66       40.5
   b8++      681.77       76.9
   y4+       703.66       53.3
   a9++      711.23       35.3
   a2+       715.79       64.4
   b2+       743.62       44.3
   y12++     769.78       33.2
   a10++     775.92       35.4
   b10++     789.87       65.8
   y5+       832.45       56.6
   b11++     846.13       80.2
   a3+       896.10       34.2
   b12++     902.85       80.6
   y6+       919.67       60.7
   y13++     948.38       43.4
   b5+      1068.72       39.4
   y9+      1212.76       34.9
   y10+     1269.73       76.0
```

```
25. MORG-339_phosphopeptide_TMT_22%ACN_032017_OT.12399.12399.3 (dLsLEEIQkk)

   Ion         m/z   Intensity
   b1+       345.27       51.9
   y2++      367.43       32.3
   y1+       376.37       41.8
   y3++      431.49       54.8
   b2+       458.39       37.8
   y4++      488.08      102.6
   y5++      552.55       95.8
   a8++      605.51       34.7
   y6++      617.16       69.1
   b8++      619.08      202.7
   b3+       624.98      699.7
   y7++      673.59      140.4
   y2+       733.54       33.8
   y8++      757.13       51.4
   b9++      798.19       52.1
   y9++      813.37       57.8
   y3+       861.72       40.1
   b5+       867.49       55.8
   b6+       996.48       51.2
   b7+      1109.58       30.0
```

```
26. MORG-339_phosphopeptide_TMT_16%ACN_032017_OT.1254.1254.3 (rEsPSPAPkPR)

   Ion         m/z   Intensity
   y2+       272.23       34.1
   b3++      341.55       68.1
   y4++      363.79       88.5
   a4++      375.77       58.3
   b1+       386.21       83.7
   y5++      399.42       51.3
   b5++      433.92       37.9
   y6++      447.96      219.6
   y7++      491.61       40.2
   a7++      503.98       39.7
   b2+       515.31       84.3
   b7++      517.77       51.4
   y8++      539.80      239.9
   a8++      551.98       32.7
   b8++      566.27      203.1
   y9++      623.44      120.2
   b3+       682.26       36.3
   y10++     688.01       40.4
   y4+       726.53       67.1
   b9++      745.39       78.6
   a4+       751.29       43.6
   b10++     793.40       52.7
   y6+       894.60      135.7
   b7+      1034.59       46.3
   y8+      1078.61       52.5
```

```
27. MORG-339_phosphopeptide_TMT_12%ACN_032017_OT.4496.4496.3 (eTNVSkEDtDQEEk)

   Ion         m/z   Intensity
   b1+       359.40       12.9
   y1+       376.38       20.0
   y2+       505.43       25.8
   b6++      559.32       19.6
   b3+       574.49       14.9
   b7++      623.96       36.5
   y3+       634.46       26.2
   a4+       645.33       11.9
   b4+       673.47       20.5
   b8++      681.37       74.7
   a9++      757.84       13.2
   b5+       760.45       16.4
   y4+       762.49       70.9
   b9++      772.14       25.1
   b10++     829.52       66.1
   y9++      830.37       29.9
   y10++     874.06       24.1
   y5+       877.45       48.9
   a11++     879.50       15.2
   b11++     893.71       24.3
   y11++     923.52       18.8
   a12++     943.90       17.6
   b12++     958.05       37.4
   y12++     980.65       15.1
   b13++    1022.64       38.7
   y13++    1031.26       23.0
   y6+      1058.43       29.0
   y7+      1173.59       17.0
```

```
28. MORG-339_phosphopeptide_TMT_16%ACN_032017_OT.3535.3535.3 (ksPGDEDDkDck)

   Ion         m/z   Intensity
   y4++      504.93       36.4
   y2+       536.32       26.3
   y5++      562.40       23.2
   y6++      620.08       17.8
   b3+       621.95       10.6
   b4+       679.57       12.7
   y7++      684.53       34.4
   a9++      742.03       23.1
   b9++      755.56       28.4
   y9++      770.49       24.6
   y10++     819.15       47.9
   a11++     879.38       23.2
   y4+      1008.69       11.4
   y5+      1123.84       14.1
```

```
29. MORG-339_phosphopeptide_TMT_10%ACN_032017_OT.11287.11287.4 (qADVADQQTTELPAENGETENQsPAsEEEkEAk)

   Ion         m/z   Intensity
   b8++      543.45      104.3
   b3+       544.38      106.0
   y3+       576.44       66.9
   b9++      594.02      164.2
   b4+       643.43      129.3
   b10++     644.45       82.2
   a18+++    700.11      254.8
   b11++     708.81       91.9
   b18+++    709.60       89.6
   b5+       714.48      141.4
   y8++      744.57       67.2
   y15+++    765.64      134.2
   y9++      779.77      105.6
   b13++     814.31       77.8
   y10++     828.62      145.9
   b6+       829.48      161.1
   y11++     912.12      175.8
   a23+++    913.06      101.2
   y20+++    932.77      194.0
   y4+       933.67       67.7
   b7+       957.52      126.9
   y21+++    965.24      288.5
   y13++    1033.22      178.3
   b26+++   1034.23       98.8
   a18++    1050.25       70.4
   a8+      1057.64       86.5
   b27+++   1077.70      142.7
   b8+      1085.58      137.0
   y15++    1148.27      152.5
   b9+      1186.64      127.6
   y16++    1212.89       92.3
   y17++    1241.32      172.8
   a10+     1259.72       59.3
   y29+++   1260.64       69.1
   b10+     1287.74       79.9
   b22++    1299.70      156.0
   y7+      1320.59       61.3
   b32+++   1349.51       61.2
   y32+++   1355.13       71.1
   y20++    1398.47      207.3
   b11+     1416.69      181.7
   a24++    1417.69      165.1
   b12+     1529.80      277.8
   y23++    1568.15       61.7
   a14+     1669.95       66.9
   b15+     1826.92       61.6
```

```
30. MORG-339_phosphopeptide_TMT_22%ACN_032017_OT.4453.4453.4 (rAGDVLEDsPkRPk)

   Ion         m/z   Intensity
   y3++      315.32      164.5
   y4+++     329.71       62.1
   b8+++     362.13      261.0
   b5++      364.61       50.2
   b1+       386.42       51.1
   b9+++     417.88      205.9
   b6++      421.32       65.9
   a2+       429.56       49.4
   y7+++     456.02      101.4
   a7++      472.22       65.4
   y2+       473.43      169.2
   a3+       485.98      112.4
   y4++      493.72       59.9
   y8+++     499.17       47.3
   b3+       514.16       46.4
   y5++      542.53      276.3
   b8++      543.51      352.3
   a11+++    560.31     1464.3
   y10+++    569.89      235.1
   a9++      612.36       78.9
   b12+++    621.63      166.9
   y6++      626.03      257.3
   y12+++    626.94       79.4
   y3+       629.41      123.1
   y13+++    650.69       65.0
   y7++      683.42       72.4
   y9++      804.91      159.9
   a6+       813.82       55.9
   b6+       841.55       49.7
   y10++     854.17      187.0
   b7+       970.56      115.6
   b8+      1085.61      168.2
```

```
31. MORG-339_phosphopeptide_TMT_22%ACN_032017_OT.11708.11708.3 (eRsPALksPLQSVVVR)

   Ion         m/z   Intensity
   y4+       472.41       35.3
   b2+       515.42       31.2
   y5+       559.46       48.1
   a7++      646.71       82.7
   b7++      661.09       68.5
   y6+       687.49       41.6
   y10++     711.22       75.5
   a8++      730.59      388.7
   b8++      744.18       43.5
   a4+       751.66       47.7
   y11++     767.94       32.2
   y12++     803.37       99.5
   b10++     849.62       47.5
   b5+       850.68       40.5
   y13++     851.92       81.2
   y8+       897.63      118.2
   a11++     899.71       39.8
   b11++     913.86       58.8
   y14++     935.70       32.7
   b6+       963.53       43.4
   b13++    1006.93       32.9
   y15++    1013.78       44.4
   b14++    1056.45       64.1
   y9+      1064.82       59.7
   b15++    1105.88       79.4
```

```
32. MORG-339_phosphopeptide_TMT_18%ACN_032017_OT.3531.3531.3 (kRSPsPsPTPEAk)

   Ion         m/z   Intensity
   y1+       376.48       39.6
   y2+       447.42       56.7
   a3+       573.04       36.7
   b9++      664.76       32.7
   a4+       670.95      133.4
   y4+       673.47      150.9
   a10++     699.22       54.3
   y10++     700.00       49.9
   y5+       774.51       39.9
   b12++     812.60       76.6
   y12++     822.26       32.3
   y6+       871.54      164.6
   y7+      1038.54       64.4
   y8+      1135.61       33.9
```

```
33. MORG-339_phosphopeptide_TMT_30%ACN_032017_OT.12724.12724.3 (kTsGPPVSELITk)

   Ion         m/z   Intensity
   a5++      376.29       41.2
   y2+       477.44       71.7
   b8++      532.15       26.5
   y3+       590.46       71.9
   a3+       598.17       36.7
   y9++      606.93       31.8
   b3+       626.23       19.6
   b10++     653.68       32.3
   a4+       655.66       45.3
   b4+       683.57       30.7
   a11++     695.94       65.1
   y4+       703.52       37.1
   y11++     719.20       69.9
   y12++     769.00       29.1
   y5+       832.64       43.3
   y6+       919.58       73.7
   a7+       948.74       23.2
   y8+      1115.71       22.5
   y9+      1212.79       35.1
   y10+     1269.79       52.7
```

```
34. MORG-339_phosphopeptide_TMT_22%ACN_032017_OT.12301.12301.4 (qVDTEEAGMVTAATASNVkAsPk)

   Ion         m/z   Intensity
   b2+       457.36       63.3
   b6++      466.35       46.8
   y2+       473.43      265.2
   a13+++    501.85       76.8
   b8++      530.51      198.3
   y5++      535.05      246.3
   a14+++    536.10       69.9
   b3+       572.48       55.7
   y6++      584.53      179.4
   b9++      595.99      219.8
   y14+++    628.56       44.4
   b17+++    636.14      106.4
   y3+       640.49      120.7
   y7++      641.48      221.1
   a4+       645.52      266.5
   a18+++    659.45       60.7
   y15+++    672.28       57.1
   b4+       673.68       61.5
   y8++      685.16      372.0
   b11++     696.15      228.1
   y4+       711.51      230.7
   y9++      720.64      310.4
   b12++     731.44       96.3
   y18+++    757.79      114.6
   b19+++    788.37      305.7
   a20+++    802.46       51.7
   a14++     803.66       55.3
   y11++     806.67      338.1
   b20+++    812.21      159.7
   b14++     817.70      247.5
   y20+++    834.09       71.6
   y12++     842.15      214.2
   b15++     853.17      109.1
   b21+++    867.65       80.5
   y13++     892.74      290.6
   b16++     896.68      223.8
   b6+       931.43       71.1
   y14++     942.25      135.1
   b17++     953.80       65.1
   b18++    1003.22      372.3
   y15++    1007.69       63.5
   y16++    1036.28       75.4
   b8+      1059.56      142.8
   b19++    1181.80       85.1
   b9+      1190.55      114.0
   b10+     1289.64      102.2
   b11+     1390.71       45.7
   b12+     1461.80       51.6
```

```
35. MORG-339_phosphopeptide_TMT_14%ACN_032017_OT.2551.2551.3 (esPSPAPkPR)

   Ion         m/z   Intensity
   y2+       272.21       15.7
   a1+       331.35       11.1
   b4++      355.29       16.6
   b1+       359.35       23.9
   y4++      363.69       40.0
   y5++      399.41       23.6
   b6++      439.51       19.7
   y6++      447.92      113.5
   a7++      474.08       12.7
   y7++      491.34       46.4
   a2+       498.18       15.2
   b2+       526.29       78.0
   y8++      539.73       14.7
   a3+       595.39       14.3
   y9++      623.46       31.4
   y3+       629.66       15.8
   b8++      666.71       27.6
   b4+       710.43       26.5
   b9++      715.77       17.0
   y4+       726.47       48.8
   y5+       797.62       23.5
   b5+       807.44       18.0
   b6+       878.44       18.0
   y6+       894.59       39.7
   y7+       981.64       20.9
```

```
36. MORG-339_phosphopeptide_TMT_8%ACN_032017_OT.3577.3577.2 (esPsPAPkPR)

   Ion         m/z   Intensity
   y4+       726.44       16.0
   b4+       790.53        9.3
   y5+       797.03       15.2
   y6+       894.68       18.0
   b6+       958.49        8.5
   b7+      1055.41        7.1
   y7+      1061.46       12.0
   y8+      1158.58       12.2
   y9+      1325.75        9.3
   b8+      1412.78       10.6
```

```
37. MORG-339_phosphopeptide_TMT_14%ACN_032017_OT.4004.4004.3 (sEEEQSSASVkk)

   Ion         m/z   Intensity
   y2++      367.33       17.5
   y1+       376.31       24.7
   b1+       397.29       24.1
   y3++      416.98       20.5
   y4++      460.47       32.2
   y5++      495.89       26.9
   b2+       526.31       15.6
   y6++      539.51       22.5
   y7++      583.09       38.9
   y8++      647.14       59.7
   b3+       655.21       46.8
   a10++     658.51       25.8
   b10++     672.25       21.5
   y9++      711.72       52.3
   y2+       733.44       30.1
   y10++     776.14       57.9
   b4+       784.34       29.3
   y3+       832.61       26.0
   y11++     840.74       61.5
   b11++     851.07       30.0
   b5+       912.76       69.8
   y4+       919.75       27.0
   y5+       990.75       15.5
```

```
38. MORG-339_phosphopeptide_TMT_22%ACN_032017_OT.13229.13229.2 (sYNLVVAk)

   Ion         m/z   Intensity
   y1+       376.33       40.3
   b1+       397.20       36.2
   y2+       447.34       58.6
   y3+       546.44       74.0
   y4+       645.50       64.3
   a3+       646.43       26.7
   b3+       674.38       33.8
   y5+       758.64       39.0
   a4+       759.49       27.4
   b4+       787.47       53.0
   a5+       858.46       21.1
   y6+       872.63       42.5
   b5+       886.42       65.1
   a6+       957.55       20.5
   b6+       985.50       60.8
   y7+      1035.63       65.3
   b7+      1056.62       42.1
```

```
39. MORG-339_phosphopeptide_TMT_25%ACN_032017_OT.11597.11597.3 (aAFSkDEskEPIVEVR)

   Ion         m/z   Intensity
   y2+       274.24       37.2
   b6++      540.00       90.0
   y8++      599.92       52.2
   b7++      604.33       34.4
   y6+       712.53       72.1
   y10++     748.03       56.7
   y11++     805.43       68.1
   y7+       841.31       85.0
   b10++     931.12      112.8
   a5+       935.63       33.5
   b11++     979.23       44.5
   y13++    1028.01       35.6
   b6+      1078.67       90.1
   b13++    1085.95       45.9
   y14++    1101.47      142.8
   b14++    1150.48      117.3
   y8+      1198.88       31.7
   b15++    1199.92       58.8
   b7+      1207.91       55.8
```

```
40. MORG-339_phosphopeptide_TMT_22%ACN_032017_OT.5174.5174.4 (sRPLNAVSQDGkETk)

   Ion         m/z   Intensity
   y1+       376.48       28.2
   b4++      382.26       26.1
   b5++      439.39       40.4
   y8+++     451.07       25.0
   a6++      461.23       37.4
   b6++      474.79       54.6
   y2+       477.33       38.7
   y4++      482.35       28.6
   y5++      510.86      192.5
   b7++      524.32       70.2
   a2+       524.94       43.3
   y11+++    545.43       32.1
   a8++      553.36       31.0
   y6++      568.26       86.9
   y12+++    583.71       81.7
   y3+       606.42       28.5
   y13+++    615.87       32.8
   a3+       622.66       30.2
   b9++      631.78       53.9
   y7++      632.60       72.0
   b13+++    640.52      160.5
   y14+++    667.88       55.9
   y8++      675.90      118.9
   b10++     689.56      123.9
   a11++     704.08       29.0
   b11++     718.13       28.6
   y9++      725.53       84.0
   y10++     761.15       56.6
   y11++     818.11       50.1
   y13++     923.24       58.7
```

```
41. MORG-339_phosphopeptide_TMT_10%ACN_032017_OT.4097.4097.4 (rRsPsPAPPPPPPPPPPR)

   Ion         m/z   Intensity
   y4++      233.88       14.5
   y5++      282.42       36.9
   y6++      330.92       62.2
   y3+       369.44       17.6
   a7+++     372.13       20.0
   y7++      379.54      114.2
   b7+++     381.40       44.4
   a4++      389.48       16.4
   a8+++     404.30       12.4
   b8+++     413.78      122.2
   y8++      428.11      106.6
   a9+++     436.82       59.4
   y13+++    438.42       36.2
   b9+++     446.12      212.0
   y4+       466.45       54.0
   a10+++    469.15       69.3
   a5++      473.55       28.4
   y9++      476.64       75.8
   b10+++    478.42      292.0
   b5++      487.35       22.4
   a11+++    501.39       54.2
   b11+++    510.83      242.7
   a6++      521.83       24.0
   y10++     525.10       52.8
   y15+++    526.80       14.6
   a12+++    533.62       32.6
   b6++      535.91       15.5
   b2+       542.47       29.9
   b12+++    543.25      157.3
   a7++      557.46       58.7
   y5+       563.40      116.7
   b13+++    575.39       18.1
   b14+++    607.86       28.4
   b8++      619.82       74.1
   a9++      654.44       21.8
   y13++     657.58       27.7
   y6+       660.43      184.2
   b9++      668.27       72.6
   b16+++    672.68       19.5
   a10++     702.99       29.7
   b10++     716.83      117.7
   a11++     751.38       26.3
   y7+       757.47      158.3
   b11++     765.53      156.7
   a12++     800.08       12.5
   b12++     814.00       98.3
   y8+       854.54       75.7
   b13++     862.58       34.0
   y9+       951.57       28.3
   b7+      1141.61       17.9
```

```
42. MORG-339_phosphopeptide_TMT_6%ACN_032017_OT.3674.3674.3 (vDGPRsPsYGRsR)

   Ion         m/z   Intensity
   b5++      377.88       21.1
   y6++      443.16       21.5
   b2+       444.32       67.7
   b6++      461.28       47.4
   y7++      491.84       15.2
   b3+       501.29       13.5
   y4+       555.32       41.4
   y8++      575.03       21.1
   b8++      592.78       25.9
   y9++      653.45       16.3
   y5+       718.37       24.6
   y11++     730.38       73.2
   a11++     767.02       12.6
   b11++     781.43       19.9
   y12++     787.72       13.1
   y6+       885.54       19.3
   y7+       982.38       40.9
```

```
43. MORG-339_phosphopeptide_TMT_14%ACN_032017_OT.6153.6153.2 (eVQsPEQVk)

   Ion         m/z   Intensity
   b1+       359.31       17.3
   y1+       376.34       25.6
   a2+       430.34       15.6
   b2+       458.30       19.7
   y2+       475.38       16.8
   b3+       586.49       23.8
   y3+       603.39       28.2
   a4+       725.48       16.5
   y4+       732.17       17.3
   b4+       753.35       32.0
   y5+       829.50       71.7
   b5+       850.57       18.6
   b6+       979.46       25.8
   y6+       996.55       18.8
   b7+      1107.59       30.8
   y7+      1124.19       14.0
   b8+      1206.64       30.1
   y8+      1223.61       16.3
```

```
44. MORG-339_phosphopeptide_TMT_14%ACN_032017_OT.5814.5814.3 (sRTAsLTSAASIDGSR)

   Ion         m/z   Intensity
   a3++      273.44       15.8
   b1+       317.12       14.2
   y3+       319.22       72.5
   b5++      406.71       22.5
   y4+       434.18       30.3
   y10++     482.89       15.5
   a7++      499.50       20.3
   b7++      513.89       29.7
   y11++     539.40       49.7
   y5+       547.33       25.8
   b8++      557.53       23.8
   b3+       574.78       30.0
   a9++      578.37       17.1
   b9++      593.29       29.9
   a10++     614.48       35.5
   a4+       617.85       26.9
   y12++     622.45       19.1
   y6+       634.43       18.4
   b11++     672.15       20.1
   y7+       705.39       60.4
   a12++     714.58       22.2
   b12++     728.07       47.7
   y8+       776.66       81.5
   b13++     786.01       96.6
   y15++     786.95       29.0
   a14++     799.97       16.5
   b14++     814.66       17.6
   y9+       863.43       77.5
   y10+      964.51       55.0
   y11+     1077.52       47.7
   y12+     1244.63       14.6
```

```
45. MORG-339_phosphopeptide_TMT_30%ACN_032017_OT.5422.5422.4 (gksASSPkPDTkVPQATAEAk)

   Ion         m/z   Intensity
   y1+       376.39       35.0
   y2+       447.39       53.4
   y8++      523.01       37.2
   y3+       576.53       41.1
   a11+++    599.00       31.6
   y4+       647.52       45.5
   a12+++    718.81       40.0
   y14+++    724.86       31.8
   b12+++    728.12       76.3
   y5+       748.56       60.0
   a13+++    751.26       68.4
   b8++      755.95       88.0
   y15+++    757.15      447.3
   b13+++    761.10      108.0
   a14+++    783.96       33.3
   y16+++    786.22       41.7
   b14+++    793.26       78.9
   y11++     801.74      154.3
   b9++      804.77       50.0
   b3+       811.29       43.0
   y17+++    814.86       46.3
   y6+       819.58       62.2
   a4+       854.67       45.3
   b16+++    859.63       51.7
   b10++     861.92       73.9
   a11++     898.66       49.9
   y13++     907.74      187.7
   b11++     912.45       36.3
   b18+++    917.08       51.6
   a5+       941.57       39.4
   y7+       947.51       37.5
   b19+++    959.63      151.4
   b20+++    983.31       31.1
   a6+      1028.56       56.1
   y8+      1044.58      115.9
   b6+      1056.31      110.1
   y15++    1134.87      111.1
   b13++    1140.84       51.4
   y11+     1602.08       31.0
```

```
46. MORG-339_phosphopeptide_TMT_4%ACN_032017_OT.6543.6543.3 (tGEPDEEEGTFRSsIR)

   Ion         m/z   Intensity
   y2+       288.26       19.6
   b2+       388.29        7.2
   y8++      502.28       10.6
   b3+       517.48       23.0
   y4+       542.29        7.3
   b8++      558.55        9.7
   y9++      566.77        9.3
   a9++      573.36        9.9
   y10++     631.32        9.9
   b10++     637.97       15.9
   y11++     695.89       15.2
   a11++     696.90       12.4
   y5+       698.58       10.0
   b5+       729.42       26.7
   y12++     752.98       19.6
   b12++     789.51       20.9
   y13++     801.83       15.3
   b13++     833.04       17.0
   y6+       845.54       13.6
   b6+       858.45       25.0
   y14++     866.83        8.4
   b14++     916.34       15.3
   b7+       987.49       16.9
   y8+      1003.60        9.0
   b8+      1116.54       17.6
```

```
47. MORG-339_phosphopeptide_TMT_20%ACN_032017_OT.8884.8884.4 (tDHGAEIVYkSPVVSGDtSPR)

   Ion         m/z   Intensity
   y2+       272.21       79.7
   b3++      292.20       43.3
   b4++      320.74       73.4
   a8+++     342.20       36.2
   y6++      356.29       95.4
   a6++      406.95       55.1
   b6++      420.86      215.5
   b2+       446.31       64.5
   a7++      463.40      115.7
   b7++      477.42      246.3
   y9++      498.79       35.7
   a8++      512.94       79.9
   b10+++    525.29      226.5
   b8++      526.81      171.3
   y4+       540.33       72.9
   a11+++    544.75       49.8
   y10++     548.05      128.3
   b11+++    554.32      202.6
   y13+++    568.31       34.0
   b3+       583.34       76.0
   b12+++    586.63       50.7
   y11++     591.42       51.5
   a9++      594.34       43.5
   y14+++    601.32       51.7
   b9++      608.51      135.8
   a13+++    610.27       45.2
   b13+++    619.63      143.4
   y15+++    638.48       38.3
   b4+       640.33       39.9
   a14+++    642.69       40.9
   b14+++    652.66      135.2
   y5+       655.36       63.6
   a15+++    671.99       35.6
   y16+++    681.56       46.9
   b5+       711.42       56.8
   y6+       712.34       97.7
   b17+++    739.01      128.4
   y19+++    770.13      150.7
   b10++     786.99      115.4
   a18+++    789.76      142.7
   y7+       799.36      123.7
   b19+++    827.95       44.2
   b11++     830.70      165.3
   b6+       840.43       59.0
   y13++     851.66      263.8
   y8+       898.45      102.6
   y14++     901.19      236.5
   b13++     928.83      103.2
   b7+       953.49       54.7
   y15++     957.78      129.5
   b14++     978.10       66.1
   y9+       997.43       75.0
   y16++    1022.27       99.6
   b8+      1052.55       51.9
   y17++    1057.79       51.5
   y18++    1086.30       58.8
   y10+     1094.46      139.2
   y11+     1181.49      101.9
```

```
48. MORG-339_phosphopeptide_TMT_12%ACN_032017_OT.5994.5994.4 (dkEVsDDEAEEkEEk)

   Ion         m/z   Intensity
   b1+       345.25       76.9
   a3++      402.34       80.6
   b3++      416.45      148.7
   b4++      466.01      132.4
   y4++      496.54      152.5
   a9+++     500.54      183.4
   y2+       505.40       72.2
   b9+++     509.43       57.9
   y5++      561.08      140.8
   b6++      606.79      156.1
   y11+++    616.08      111.0
   y6++      625.61       74.1
   y3+       634.34       63.5
   y7++      661.08      150.3
   b7++      664.61      164.4
   b12+++    715.35       98.5
   y8++      725.61      272.0
   b8++      729.05       87.3
   b13+++    758.38       85.9
   b9++      764.43       55.5
   y9++      783.12      317.8
   b14+++    801.24       85.4
   y14+++    811.62      182.5
   b10++     829.08       77.4
   b3+       831.67       84.2
   y10++     840.48      177.4
   b11++     893.50       92.8
   y11++     924.33      180.8
   y12++     973.36       70.3
```

```
49. MORG-339_phosphopeptide_TMT_10%ACN_032017_OT.8328.8328.4 (dksPVREPIDNLTPEER)

   Ion         m/z   Intensity
   y2+       304.17       29.0
   a2++      337.80       15.1
   b1+       345.26       16.6
   b2++      352.18       11.2
   b6+++     408.27       10.8
   b3++      435.17       21.8
   a7+++     441.64       10.7
   b7+++     451.08       20.3
   a9+++     511.68       12.5
   a5++      519.59       12.2
   b9+++     520.76       17.8
   y4+       530.32       41.4
   b5++      533.70       12.1
   y14+++    555.36       10.6
   b10+++    559.52       62.0
   a11+++    588.45       11.7
   y10++     591.88       18.6
   a6++      597.49       42.7
   b6++      611.16       15.6
   a12+++    625.48       41.3
   y11++     656.88       14.2
   a13+++    659.84       11.9
   b13+++    668.91       28.9
   b7++      675.76       12.2
   b8++      724.88       14.6
   y16+++    730.59       22.8
   y12++     735.29       24.1
   y6+       744.31       49.3
   b16+++    787.28       18.9
   a10++     824.34       28.1
   y14++     833.23       74.5
   b10++     838.60       20.5
   y7+       858.55       38.2
   b11++     895.77       23.7
   b12++     951.75       14.3
   y8+       973.49       13.1
   b13++    1002.82       14.5
```

```
50. MORG-339_phosphopeptide_TMT_10%ACN_032017_OT.4100.4100.4 (aETSESSGSAPAVPEAsASPkQRR)

   Ion         m/z   Intensity
   a8+++     317.28       11.7
   y11+++    512.75       20.2
   b9++      533.48       24.7
   a10++     554.35       11.3
   y13+++    569.02       22.9
   y14+++    601.84       53.3
   a11++     603.42       11.3
   b4+       618.17       10.9
   b17+++    623.64       12.3
   y15+++    625.42       19.6
   a12++     639.28       11.2
   b12++     653.25       17.6
   y9++      655.00       16.3
   a19+++    666.63       10.8
   y17+++    673.76       19.9
   b19+++    675.86       10.7
   a13++     688.44       19.4
   a20+++    698.78       15.0
   y18+++    702.29       42.6
   b20+++    708.67       27.9
   a5+       719.08       25.8
   y10++     720.06       35.4
   b5+       747.53       19.7
   y11++     768.55       45.5
   y4+       816.59       11.8
   a21+++    817.86       18.0
   b21+++    826.98       17.1
   b6+       834.40       12.6
   y13++     853.23       24.5
   y14++     902.24      101.0
   b7+       921.62       16.8
   y15++     937.87       14.3
   b8+       978.58       12.5
   y16++     981.45       18.6
   y6+      1000.96       14.6
   y17++    1009.64       20.2
   b9+      1065.63       13.2
   b10+     1136.56       33.5
```

```
51. MORG-339_phosphopeptide_TMT_22%ACN_032017_OT.6471.6471.3 (sLSsSPVkkADAR)

   Ion         m/z   Intensity
   y2+       246.19       47.8
   b1+       397.11       22.9
   y6++      573.77       26.8
   y7++      623.22       40.7
   y8++      672.37      166.3
   b8++      702.53       60.9
   y9++      715.45       44.3
   y5+       789.30       36.6
   y10++     799.10       23.4
   a5+       823.16       26.3
   y11++     842.90       28.0
   b9++      881.75       35.1
   y12++     898.96       39.8
   a10++     902.66       23.4
   b10++     916.78       58.0
   a11++     960.77       30.4
   b11++     974.66      149.4
   a7+      1019.78       30.1
   b7+      1047.39       23.5
```

```
52. MORG-339_phosphopeptide_TMT_10%ACN_032017_OT.18261.18261.3 (dEkkEEsEEsDDDmGFGLFD)

   Ion         m/z   Intensity
   b1+       345.25       33.2
   b4++      594.94       41.8
   y6+       655.47       41.4
   b5++      659.71       47.6
   b6++      724.27       78.7
   y7+       802.46       56.3
   a3+       803.47       41.2
   b3+       831.07       36.7
   b8++      871.85       42.3
   y8+       917.47       48.7
   a9++      922.06       47.2
   b9++      936.84      191.4
   b10++    1020.09       32.7
   y9+      1032.28       57.8
   b11++    1077.30       45.8
   b12++    1134.87       63.5
   b13++    1192.69       89.6
   b14++    1266.19       52.9
   b15++    1294.76       36.1
   y11+     1314.38       36.1
   b16++    1368.45       65.7
   b17++    1396.92       28.9
   y19++    1421.49       42.3
   b18++    1453.45       27.6
```

```
53. MORG-339_phosphopeptide_TMT_25%ACN_032017_OT.11182.11182.3 (sGILVPSEkk)

   Ion         m/z   Intensity
   y2++      367.38       61.4
   y1+       376.30       24.1
   b1+       397.21       56.1
   y3++      431.77       22.4
   b2+       454.34       19.3
   y4++      475.29       23.7
   y5++      523.97      161.5
   a3+       539.38       41.3
   b3+       567.23       80.5
   y6++      573.60       81.2
   y7++      629.86       98.9
   b4+       680.38       60.8
   y8++      686.55       59.3
   y9++      715.13      137.5
   b9++      725.37       31.9
   a5+       751.32       31.7
   b5+       779.47       87.1
   a6+       848.67       21.8
   y5+      1046.72       63.4
   b8+      1092.51       29.8
```

```
54. MORG-339_phosphopeptide_TMT_20%ACN_032017_OT.5610.5610.3 (skEsLQEAGk)

   Ion         m/z   Intensity
   b1+       317.35       15.7
   b2++      337.89       19.6
   y1+       376.43       15.1
   a3++      388.20       19.3
   b3++      402.56       45.4
   y2+       433.44       44.6
   b4++      486.05       17.5
   y3+       504.47       57.4
   b5++      542.48       15.9
   y8++      585.61       45.6
   a6++      592.09       28.2
   b6++      606.27       52.8
   y4+       633.42       54.9
   a7++      657.30       38.8
   b7++      671.08       68.6
   b8++      706.38       38.7
   b9++      735.06       19.4
   y5+       761.48       39.7
   y9++      764.59       15.6
   b3+       803.72       17.3
   y6+       874.52       56.8
   y7+      1041.59       20.4
```

```
55. MORG-339_phosphopeptide_TMT_10%ACN_032017_OT.20841.20841.3 (dEkkEEsEEsDDDMGFGLFD)

   Ion         m/z   Intensity
   b3++      416.37       17.0
   b4++      594.93       22.5
   y5+       598.37       13.4
   y6+       655.35       14.8
   b5++      659.61       19.3
   b6++      724.18       28.3
   y7+       786.46       29.5
   a8++      858.21       13.7
   b8++      872.19       24.4
   y8+       901.50       18.4
   a9++      922.21       40.1
   y15++     926.40       49.2
   b9++      936.35       20.2
   y9+      1016.41       19.3
   b10++    1020.40       15.4
   b11++    1077.32       24.1
   b12++    1134.93       25.5
   a4+      1160.40       24.6
   y17++    1169.68       12.7
   b4+      1188.83       13.3
   b13++    1192.26       33.5
   b14++    1258.15       38.5
   b15++    1286.80       25.7
   y11+     1298.58       12.8
   b16++    1360.49       36.9
   b17++    1388.87       17.4
   y12+     1427.51       17.2
   b18++    1445.54       20.5
```

```
56. MORG-339_phosphopeptide_TMT_16%ACN_032017_OT.7859.7859.3 (eksPELPEPSVR)

   Ion         m/z   Intensity
   a1+       331.27       12.5
   a2++      344.64       18.4
   b1+       358.86       31.3
   a3++      428.21       15.8
   b3++      442.38       33.5
   y4+       458.29       65.3
   y9++      512.38       15.5
   a5++      541.02       17.2
   b5++      555.36       47.3
   y5+       587.65       19.2
   y10++     595.53       10.4
   a6++      597.57       67.1
   b6++      611.69       63.1
   y6+       684.45       61.7
   a8++      710.93       14.1
   b8++      724.98       82.2
   a9++      759.09       14.3
   y11++     774.56       19.8
   y7+       797.47       40.9
   b10++     817.21       22.7
   a3+       855.42       10.6
   b11++     866.25       25.4
   y9+      1023.51      113.6
   y10+     1190.57       10.4
```

```
57. MORG-339_phosphopeptide_TMT_30%ACN_032017_OT.11604.11604.3 (sIFREEsPLRIk)

   Ion         m/z   Intensity
   y1+       376.34       29.3
   y5++      428.48       65.1
   b5++      431.73       28.0
   a6++      482.43       23.5
   y2+       489.47       34.2
   b6++      496.41       55.8
   y6++      511.99       37.8
   b8++      628.16       36.1
   y8++      641.10       26.4
   y3+       645.20      100.5
   a4+       705.22       23.5
   b4+       733.43       27.7
   a10++     748.97       22.7
   y4+       758.71       18.9
   b10++     763.09       43.2
   y10++     792.37       25.7
   b11++     819.28       26.9
   a5+       834.96       36.0
   y5+       855.67       29.9
   b5+       862.55       41.8
   b6+       991.56       60.9
   y6+      1022.66       32.9
```

```
58. MORG-339_phosphopeptide_TMT_14%ACN_032017_OT.8094.8094.3 (dEskEPIVEVR)

   Ion         m/z   Intensity
   y1+       175.18        9.9
   y2+       274.21       50.0
   a1+       317.27       10.4
   b1+       345.24       26.9
   b2+       474.26       22.1
   b4++      499.76       16.1
   y4+       502.43       15.6
   a5++      550.35       13.8
   b5++      564.32       77.7
   a6++      598.74       21.9
   y8++      600.00       63.3
   b3+       641.43       12.7
   a7++      655.36       16.9
   b7++      669.59       42.1
   y9++      683.38       27.0
   a8++      705.06       17.2
   y6+       712.46       68.3
   b8++      719.03       34.0
   y10++     747.92       29.5
   a9++      769.56       17.8
   b9++      783.57      109.9
   b10++     832.94       36.5
   y7+       841.30       33.4
   b5+      1127.51       17.4
```

```
59. MORG-339_phosphopeptide_TMT_14%ACN_032017_OT.8786.8786.4 (dVQDAIAQsPEkEAk)

   Ion         m/z   Intensity
   a1+       317.34       11.4
   b4++      344.45        8.7
   b8+++     357.37       11.1
   y1+       376.30       28.0
   b5++      379.89       25.9
   y6+++     387.30       31.3
   b9+++     413.45       10.3
   a6++      422.41       12.2
   b6++      436.22       53.8
   y7+++     442.96       14.2
   b2+       444.53       11.4
   y2+       447.30       35.0
   y4++      467.32       19.1
   b7++      471.73       42.3
   y8+++     485.60       35.1
   y9+++     509.35       59.0
   a8++      521.97       10.6
   y5++      532.10       11.6
   b8++      535.74       13.3
   y10+++    547.13       27.0
   y11+++    570.40       50.0
   b3+       572.00       18.0
   y3+       576.50       30.2
   y6++      580.39       62.2
   b12+++    607.69       15.0
   y12+++    608.61        8.9
   b9++      619.50       23.6
   a13+++    641.72       22.8
   b13+++    650.77       49.1
   y13+++    651.72       16.6
   y7++      663.98       59.3
   a14+++    665.06       19.4
   b14+++    674.47       43.5
   b4+       687.38       24.4
   y8++      728.01       42.3
   b11++     732.80        9.7
   b5+       758.37       48.1
   y9++      763.49       73.1
   y10++     820.16       32.8
   b6+       871.51       49.4
   b12++     910.82       12.4
   b7+       942.48       30.0
   b8+      1070.56       13.0
   b9+      1237.54        9.7
```

```
60. MORG-339_phosphopeptide_TMT_12%ACN_032017_OT.1974.1974.3 (rVsRtPEPk)

   Ion         m/z   Intensity
   y4++      350.49       28.9
   y1+       376.48       29.8
   b1+       386.36       34.3
   b4++      404.96       34.0
   y2+       473.36      110.8
   b2+       485.45       34.9
   b5++      495.18       73.5
   y6++      519.19       60.2
   b6++      543.67       45.8
   a7++      594.53       22.6
   y3+       602.56       43.0
   b7++      608.43      115.6
   y4+       699.44      153.2
```

```
61. MORG-339_phosphopeptide_TMT_20%ACN_032017_OT.9048.9048.4 (sLsPGkENINSQEVEk)

   Ion         m/z   Intensity
   y3++      302.82       25.8
   b1+       317.28       14.2
   y4++      367.28       13.5
   y1+       376.30       49.8
   y5++      431.40       16.0
   y6++      474.74       16.8
   b9+++     488.80       24.7
   y2+       505.38       94.2
   a10+++    517.89       12.5
   b10+++    526.90       26.0
   y7++      532.07       19.0
   b6++      554.96       27.3
   b11+++    556.15       27.9
   y8++      588.68       21.8
   a12+++    589.73       12.3
   y11+++    593.01       37.1
   b3+       597.33       33.5
   b12+++    598.80       39.5
   y3+       604.41       67.2
   a7++      605.52       41.3
   y12+++    611.68       27.7
   b7++      619.46       45.5
   b13+++    641.94      105.6
   y13+++    644.26       17.7
   a8++      662.50       17.0
   a14+++    665.49       31.7
   a4+       666.26       13.6
   b14+++    674.84      112.9
   b8++      676.47       36.6
   b4+       693.98       15.8
   b15+++    717.87       58.5
   a9++      718.51       28.6
   y4+       733.15       76.1
   b10++     790.07       62.0
   b11++     833.74       33.8
   y5+       861.54       15.2
   y11++     888.65       30.5
   b12++     897.61       46.7
   b13++     962.20       83.2
   b14++    1011.75       20.5
```

```
62. MORG-339_phosphopeptide_TMT_10%ACN_032017_OT.22563.22563.3 (kEEsEEsEDDmGFGLFD)

   Ion         m/z   Intensity
   y2+       281.09      108.3
   a3++      294.31       29.1
   y3+       394.21       30.6
   y4+       451.32       80.9
   y5+       598.34       61.2
   y6+       655.44      109.6
   y12++     729.70       67.6
   b10++     783.79       73.6
   y7+       802.39       64.0
   y8+       917.45       59.4
   a14++     974.25       41.1
   y9+      1032.41       61.0
   b15++    1045.23       47.4
   a16++    1103.97      148.4
   y10+     1161.19      141.6
   y11+     1328.45       55.9
```

```
63. MORG-339_phosphopeptide_TMT_18%ACN_032017_OT.1339.1339.3 (rksPENTEGk)

   Ion         m/z   Intensity
   y1+       376.33       14.0
   b1+       386.27       19.1
   y2+       433.32       38.3
   b3++      455.82       16.1
   y7++      502.43       35.5
   a5++      554.64       16.6
   y3+       562.39       41.3
   b5++      569.10       51.6
   y8++      586.00       43.4
   a6++      611.87      139.0
   b6++      626.07       67.2
   y4+       663.45       38.5
   b7++      676.58       18.1
   a2+       715.64       26.9
   b8++      740.84       33.8
   y9++      764.53       23.1
   b9++      769.43       19.2
   y5+       777.46       44.7
   y7+      1003.56       67.7
```

```
64. MORG-339_phosphopeptide_TMT_18%ACN_032017_OT.12502.12502.3 (asSPPDRIDIFGR)

   Ion         m/z   Intensity
   a1+       273.30       10.6
   b1+       301.31       10.7
   b5++      374.81        9.1
   y3+       379.25       10.7
   y7++      439.12       61.5
   a2+       440.30       16.2
   y4+       492.32       55.2
   a7++      496.25       21.2
   b7++      510.72       18.2
   a3+       527.36       13.4
   y9++      544.73       20.8
   b3+       555.48       30.5
   b8++      566.98       30.7
   y10++     593.48      201.6
   y5+       607.32       20.8
   b9++      624.92       52.2
   b4+       652.22        9.1
   a10++     667.68       12.1
   b10++     681.46       18.0
   y6+       720.47       23.5
   b11++     755.01       25.8
   b12++     783.54       19.9
   b6+       864.41       18.2
```

```
65. MORG-339_phosphopeptide_TMT_4%ACN_032017_OT.6575.6575.3 (aEEDEILNRsPR)

   Ion         m/z   Intensity
   y1+       175.16        6.1
   y2+       272.19       45.8
   a1+       273.12       22.0
   y5++      355.28        5.2
   y6++      411.84        9.0
   b2+       430.26        7.3
   y3+       439.20        8.0
   b6++      459.04        6.5
   y7++      468.51       13.3
   a7++      501.02        6.2
   b7++      515.23        6.3
   a3+       531.27        5.8
   y8++      533.00       30.4
   a8++      558.32       12.4
   b3+       559.31       14.8
   b8++      572.16        7.0
   y9++      590.62        6.9
   b9++      650.64       14.9
   y10++     654.65        6.4
   b4+       674.36       28.1
   y5+       709.36       12.5
   a10++     720.05       13.1
   b10++     734.13       47.9
   a5+       775.40        7.2
   b11++     782.49        5.7
   b5+       803.44       19.9
   y6+       822.37        9.4
   b6+       916.50       14.9
   b7+      1029.57        7.2
   b8+      1143.49        5.2
```

```
66. MORG-339_phosphopeptide_TMT_6%ACN_032017_OT.7258.7258.3 (eIsDDEAEEEkGEk)

   Ion         m/z   Intensity
   b1+       359.24       18.5
   y1+       376.27       14.4
   y4++      460.51       28.7
   b2+       472.32       15.1
   y2+       505.41       11.5
   y5++      524.94       26.8
   y3+       562.45       17.9
   y6++      589.34       25.4
   y7++      654.12       18.0
   y8++      689.44       27.2
   b4+       754.16      106.9
   y10++     811.69       67.0
   b5+       869.27       52.7
   b11++     907.75       17.8
   y4+       919.69       10.9
   y12++     952.15       28.0
   b6+       998.44       17.4
   b13++    1000.71       23.0
   b7+      1069.42       11.6
   b8+      1198.52       12.0
   b10+     1456.53       11.1
```

```
67. MORG-339_phosphopeptide_TMT_10%ACN_032017_OT.9094.9094.4 (gkQADVADQQTTELPAENGETENQSPAsEEEk)

   Ion         m/z   Intensity
   y1+       376.36      108.6
   y6++      501.04      127.9
   y2+       505.41      120.9
   y7++      549.93      181.2
   a7++      550.88       87.9
   b7++      564.91       72.1
   y8++      593.47      197.0
   a8++      608.54       68.8
   a2+       616.49       73.2
   b8++      622.53      128.4
   y3+       634.43      134.3
   b14+++    648.30       92.1
   y9++      657.59      108.2
   b9++      686.62      103.9
   y10++     714.51      162.0
   b10++     750.61      126.0
   y4+       763.52      125.2
   y11++     779.17      122.2
   b11++     801.22      110.3
   y12++     829.52      150.0
   b12++     851.66      169.9
   y13++     894.10       73.7
   b13++     916.29      356.9
   y14++     922.61      111.2
   a5+       930.53       87.4
   b5+       958.31       87.3
   b23+++    962.70       77.5
   b14++     972.74      593.6
   y6+      1001.55      118.4
   a15++    1006.77      180.9
   a6+      1029.73       83.5
   a26+++   1057.11       87.3
   b6+      1057.77       72.5
   y17++    1079.52      106.2
   y7+      1098.50      195.0
   a7+      1100.57       79.8
   b17++    1120.91      213.0
   b7+      1128.73      300.1
   b18++    1177.94       68.6
   a29+++   1179.11      186.2
   y19++    1184.72      113.0
   y8+      1185.53      235.9
   b8+      1243.62      108.2
   b31+++   1275.14       84.2
   y9+      1313.52       76.0
   y10+     1427.66       73.9
   b10+     1499.65       78.6
```

```
68. MORG-339_phosphopeptide_TMT_4%ACN_032017_OT.9147.9147.3 (kETEsEAEDDNLDDLER)

   Ion         m/z   Intensity
   b4++      358.98        8.4
   y3+       417.30        9.1
   y4+       532.37       12.0
   y5+       647.39        8.0
   b4+       717.43        5.6
   b10++     722.39        7.2
   y6+       760.45        6.6
   b11++     779.63       16.8
   y13++     800.76       11.1
   y7+       874.48       11.9
   b13++     893.86        6.7
   a14++     937.11        5.2
   b14++     950.97        9.6
   a6+       985.80        5.6
   y8+       989.56       14.1
   a7+      1056.76        8.3
   y9+      1104.56       12.3
   y10+     1233.56       10.7
   y11+     1304.52       10.8
   a10+     1415.70        5.4
   y12+     1433.72        5.1
```

```
69. MORG-339_phosphopeptide_TMT_25%ACN_032017_OT.11385.11385.4 (eRsPALkSPLQSVVVR)

   Ion         m/z   Intensity
   y6+++     230.22       40.3
   a2++      244.30       55.0
   b2++      258.27       86.7
   y7+++     267.34       46.5
   b5+++     284.09       31.4
   a3++      327.86      110.7
   a1+       331.36       36.6
   b3++      341.83      255.0
   b1+       359.32       64.4
   y3+       373.37       47.5
   a4++      376.47       73.3
   a5++      411.81       96.2
   b5++      425.79      115.4
   a7+++     431.65       51.7
   b7+++     441.09       75.3
   y8++      449.37       32.1
   a8+++     460.67       33.4
   a6++      468.53       76.2
   b8+++     470.12       68.5
   y4+       472.42       55.1
   b6++      482.49      121.7
   a10+++    530.87       55.7
   b10+++    540.30      114.1
   y13+++    541.95      329.4
   y5+       559.39      106.1
   b11+++    582.98       59.4
   b12+++    612.03       64.6
   a13+++    635.64       53.0
   b13+++    645.14       82.5
   a7++      647.09       35.3
   y15+++    649.57       40.0
   b7++      660.92       39.2
   a14+++    668.57       42.0
   y10++     671.65      202.7
   b14+++    678.02      132.2
   b3+       682.36       40.6
   y6+       687.50      108.2
   a8++      690.21       35.5
   b15+++    710.97      117.7
   y11++     728.19      204.5
   y12++     763.65       51.3
   y7+       800.61       48.4
   b10++     809.51       39.2
   y13++     812.24      594.8
   y8+       897.58      111.1
   y9+       984.59       73.3
```

```
70. MORG-339_phosphopeptide_TMT_14%ACN_032017_OT.4138.4138.3 (aRVDHGAEIITQsPSR)

   Ion         m/z   Intensity
   b4++      336.31       80.5
   y3+       359.23       41.3
   b5++      404.99       14.5
   a7++      454.75       14.0
   b7++      468.92       17.6
   a8++      519.60       16.6
   y4+       525.93       28.7
   b8++      533.45       50.6
   b3+       556.40       16.9
   a9++      576.09       27.2
   b9++      590.06       42.4
   y11++     619.72       17.4
   a10++     632.72       21.6
   a4+       643.51       37.1
   b10++     646.36       36.2
   y5+       653.97       30.9
   b4+       671.21       64.2
   y12++     688.58       22.3
   y6+       755.26       33.2
   b12++     761.23       26.7
   y14++     795.71       93.5
   b13++     844.63       13.9
   y7+       868.41       16.2
   a14++     878.76       24.9
   b7+       936.75       15.0
   y8+       981.66       17.8
   y12+     1375.64       40.1
```

```
71. MORG-339_phosphopeptide_TMT_18%ACN_032017_OT.3325.3325.3 (rYsPPIQR)

   Ion         m/z   Intensity
   y1+       175.11       14.5
   b1++      193.79        8.8
   y4++      257.13       13.3
   a2++      261.20       14.3
   b2++      275.23       12.4
   y2+       303.29       12.2
   y5++      305.78       67.1
   a3++      344.72       26.2
   b3++      358.71       69.7
   b1+       386.38        8.0
   y6++      389.10       10.6
   a4++      393.68        6.2
   b4++      406.99        8.6
   y3+       416.09       54.2
   a6++      498.28       14.8
   b6++      512.48       26.2
   y4+       513.35       39.2
   a2+       520.99       12.1
   a7++      562.56       11.4
   b7++      576.46       40.4
   y5+       610.41      112.4
   b3+       716.35       17.7
```

```
72. MORG-339_phosphopeptide_TMT_18%ACN_032017_OT.7787.7787.3 (eEsPLRIk)

   Ion         m/z   Intensity
   a2++      230.20       25.3
   y3++      323.39       65.6
   b3++      328.39       19.6
   b1+       359.35       35.9
   b4++      376.33       56.1
   y4++      379.87       20.8
   y5++      428.48      274.0
   b2+       488.32       59.2
   y2+       489.37      103.3
   y6++      511.44       74.1
   a7++      553.87       23.3
   b7++      567.70       49.3
   y7++      576.57       33.7
   a3+       627.65       55.3
   b3+       655.30       37.9
   a5+       837.56       22.7
   b5+       865.41       44.6
```

```
73. MORG-339_phosphopeptide_TMT_10%ACN_032017_OT.22572.22572.3 (kEEsEEsDDDmGFGLFD)

   Ion         m/z   Intensity
   y2+       281.15       84.5
   y3+       394.27       29.2
   y4+       451.30       68.3
   y5+       598.44       49.1
   y6+       655.41       77.9
   y11++     657.31       27.0
   a9++      706.19       56.1
   a10++     763.08      102.8
   b10++     777.13       58.2
   y7+       802.58       70.5
   b11++     850.34       44.2
   y8+       917.53       53.2
   a14++     967.47       21.4
   y9+      1032.47       33.4
   b15++    1038.33       33.0
   a16++    1097.07      119.2
   y10+     1147.31       23.3
   y11+     1314.50       35.1
```

```
74. MORG-339_phosphopeptide_TMT_8%ACN_032017_OT.17510.17510.3 (aVDsssEEIEVEVPVVDRR)

   Ion         m/z   Intensity
   y2+       331.29       14.1
   y6++      371.39       17.2
   b2+       400.24        7.0
   a5++      411.22        7.2
   y7++      420.92       14.9
   y8++      485.43       15.8
   b3+       515.41       11.9
   y9++      535.16       17.6
   y10++     599.56       26.8
   y11++     656.15       13.6
   a9++      680.51       16.5
   b9++      694.32        7.2
   y12++     720.83       13.3
   y6+       741.61       10.4
   a10++     745.04        7.0
   b10++     758.74       11.7
   y13++     785.25        8.5
   b11++     808.36        8.8
   y15++     952.32       21.0
   y16++    1035.60       14.2
   y17++    1093.41        9.7
   y18++    1142.96        8.5
   b8+      1274.26       10.8
   b9+      1387.58       13.9
   b12+     1744.63        6.8
   a13+     1815.78        6.7
   b13+     1843.70       11.4
```

```
75. MORG-339_phosphopeptide_TMT_14%ACN_032017_OT.6430.6430.3 (rkVMDsDEDDADY)

   Ion         m/z   Intensity
   y2+       297.18       17.1
   b3++      421.80       18.5
   a4++      473.43       13.9
   b4++      487.51       19.3
   b5++      544.99       62.4
   y9++      562.54       14.0
   y5+       598.19       14.0
   b6++      628.52       16.3
   a7++      671.97       27.5
   y11++     677.67       24.9
   b7++      685.64       42.9
   a8++      736.60       14.6
   b2+       743.76       17.6
   b8++      750.61       27.9
   b9++      808.08       42.8
   b3+       842.71       20.1
   y12++     856.60       14.2
   b10++     865.61       46.4
   b11++     901.11       56.4
   b12++     958.67       62.1
   b4+       973.81       23.1
   y8+      1009.33       14.3
```

```
76. MORG-339_phosphopeptide_TMT_10%ACN_032017_OT.12327.12327.3 (lEkEEEEGISQEssEEEQ)

   Ion         m/z   Intensity
   b3++      415.25       32.1
   b4++      480.00       44.3
   b5++      544.58       69.4
   b6++      609.09       95.0
   b7++      673.65      134.0
   b8++      702.15       68.1
   a9++      744.77       48.6
   b9++      758.69      362.5
   b10++     802.21      141.2
   b11++     866.27      138.6
   y6+       868.29       56.8
   y14++     885.67       34.6
   a4+       930.81      151.7
   y7+       997.28       62.0
   a13++    1000.15       29.6
   b14++    1097.38       31.4
   y8+      1125.29       61.1
   y16++    1128.77       58.8
   b15++    1162.33       85.7
   y17++    1193.42       73.5
   a16++    1212.33      192.8
   b16++    1226.88      230.2
   b17++    1291.42      275.6
   y11+     1382.52       61.9
   y12+     1511.49       39.5
```

```
77. MORG-339_phosphopeptide_TMT_22%ACN_032017_OT.4492.4492.3 (rAVVVsPk)

   Ion         m/z   Intensity
   a2++      215.19       15.3
   y2++      237.26       17.7
   a3++      264.80       23.1
   b3++      278.70       28.7
   a4++      314.32       46.4
   b4++      328.21       52.8
   a5++      363.93       35.8
   y1+       376.43       33.0
   b5++      377.79       73.4
   b1+       386.70       32.1
   a6++      446.92       18.7
   y2+       473.37      147.6
   b3+       556.53       18.6
   y3+       640.39       35.1
   b4+       655.63       23.5
   y4+       739.50       33.8
   b5+       754.48       16.1
```

```
78. MORG-339_phosphopeptide_TMT_25%ACN_032017_OT.13845.13845.4 (aQTLPTSVVTITSESsPGkR)

   Ion         m/z   Intensity
   b5+++     247.22       17.0
   b4++      322.17       16.7
   y4++      343.88       44.4
   y5++      427.34       54.4
   b11+++    447.47       16.6
   y6++      470.89       43.3
   b8++      514.46       38.2
   y12+++    524.66       22.4
   b3+       530.33       23.5
   y7++      535.45       43.5
   b9++      563.97       61.4
   y8++      578.97       78.4
   y14+++    586.76       17.3
   a10++     600.70       16.1
   b10++     614.55       61.0
   a4+       615.47       23.0
   y15+++    620.44       45.3
   y9++      629.49      101.0
   b16+++    638.19       21.1
   b4+       643.43       45.6
   b11++     671.12       37.2
   y4+       686.15       76.4
   y17+++    690.46       15.1
   a5+       712.36       19.5
   b12++     721.71       75.7
   y11++     736.66       85.7
   b13++     765.25       21.7
   y12++     786.12       88.2
   b14++     829.77       17.9
   y13++     835.68       29.1
   b15++     873.24       57.2
   y14++     879.19       23.9
   b7+       928.66       15.9
   y15++     929.67       24.2
   b16++     956.75       15.5
   y16++     978.26       45.9
   b8+      1027.60       17.8
   b11+     1340.93       18.5
```

```
79. MORG-339_phosphopeptide_TMT_30%ACN_032017_OT.2851.2851.4 (kRkPsPsPPPPTATESR)

   Ion         m/z   Intensity
   y2+       262.25       49.1
   y8++      429.91       67.3
   y9++      478.56      128.8
   b4++      484.62       94.3
   y10++     526.90      150.5
   y14+++    527.57      120.2
   b10+++    564.72       30.7
   y11++     610.14       61.5
   a13+++    644.55       45.8
   y15+++    646.73      386.5
   b13+++    654.10       74.8
   y12++     658.73       71.1
   y16+++    699.08       32.8
   y13++     742.12       60.2
   b16+++    759.73       35.8
   y7+       761.35       64.4
   y14++     790.80       33.5
   b9++      797.69       33.7
   y8+       858.44       77.4
   y9+       955.46       70.7
   y10+     1052.55      178.6
   y11+     1219.61       29.4
   y12+     1316.65       44.2
```

```
80. MORG-339_phosphopeptide_TMT_18%ACN_032017_OT.8474.8474.2 (sVVFTSAR)

   Ion         m/z   Intensity
   y1+       175.09       17.2
   y2+       246.11       17.3
   y3+       333.18       30.6
   b1+       397.25       53.2
   y4+       434.23       53.6
   a2+       468.30       16.5
   b2+       496.38       22.9
   a3+       567.35       48.9
   y5+       581.30       79.1
   b3+       595.35       43.8
   y6+       680.40       68.8
   a4+       714.52       21.7
   b4+       742.39       59.0
   y7+       779.45       85.1
   b5+       843.48       20.0
   a6+       902.44       40.9
   b7+      1001.46       24.0
```

```
81. MORG-339_phosphopeptide_TMT_14%ACN_032017_OT.7210.7210.4 (vDHGAEIITQsPSRSSVAsPR)

   Ion         m/z   Intensity
   y2+       272.20       60.8
   a3++      277.26       23.5
   b3++      291.14       26.1
   b4++      319.84       48.9
   a5++      341.35       27.1
   b8+++     355.30       57.0
   a6++      405.90       27.4
   b6++      419.87      117.6
   b2+       444.31       41.5
   a7++      462.45       65.5
   b7++      476.46      119.4
   a12+++    510.22       25.2
   y9++      513.54       55.2
   a8++      518.99       36.8
   b8++      532.98       78.8
   y14+++    544.98       20.4
   y10++     562.51       72.3
   b3+       581.40       50.5
   y15+++    582.42       33.4
   b9++      583.49       49.0
   b14+++    601.25       41.3
   y5+       609.43       20.0
   y16+++    626.00       22.2
   b15+++    630.08       31.9
   a10++     633.49       23.3
   b4+       637.87       63.7
   a5+       681.26       23.0
   a17+++    683.08       35.9
   b17+++    692.29       27.6
   y6+       696.43       25.0
   a18+++    706.34       78.4
   y12++     709.58       29.7
   y13++     760.53       89.6
   b19+++    771.48       19.3
   y7+       783.33       35.3
   a13++     808.93       21.7
   y14++     817.10      103.6
   b6+       838.45       38.0
   y15++     873.63       62.9
   b14++     901.06       32.6
   a15++     930.16       21.8
   y16++     938.16       38.2
   b7+       951.53       32.1
   a16++     973.52       30.4
   y18++    1002.22       31.5
   b8+      1064.69       25.1
```

```
82. MORG-339_phosphopeptide_TMT_4%ACN_032017_OT.9317.9317.3 (kEtEsEAEDDNLDDLER)

   Ion         m/z   Intensity
   y4+       532.31        6.6
   a6++      533.45        4.2
   y5+       647.49        6.6
   y6+       760.50        7.5
   a11++     805.24        5.1
   b11++     819.61        8.0
   y14++     865.15        8.6
   b13++     933.79        7.3
   a5+       936.56        4.3
   a14++     976.83        5.0
   y8+       989.61        5.3
   b14++     990.57        8.6
   a15++    1033.49        5.6
   y9+      1104.53        6.6
   y11+     1304.55        7.6
   y12+     1433.49        6.8
```

```
83. MORG-339_phosphopeptide_TMT_10%ACN_032017_OT.1299.1299.2 (rRsPsPYYSR)

   Ion         m/z   Intensity
   b1+       386.37       25.4
   b2+       542.52       36.8
   y5+       685.43       75.1
   b3+       709.47       43.3
   a4+       778.37       20.8
   b4+       806.15       30.8
   y6+       852.44       37.2
   y7+       949.44       59.5
   b5+       973.46       21.0
   y8+      1116.44       29.5
   b7+      1233.56       24.0
   y9+      1272.56       22.3
   b8+      1396.66       29.9
```

```
84. MORG-339_phosphopeptide_TMT_20%ACN_032017_OT.9147.9147.3 (rAssQDGLETLkk)

   Ion         m/z   Intensity
   y1+       376.43       14.1
   b1+       386.30       14.8
   y3++      424.00       17.0
   y4++      474.67       26.5
   y5++      539.10       18.2
   b3+       624.18       73.4
   y8++      681.51       26.3
   a10++     703.66       15.4
   b10++     718.01       14.7
   y9++      745.63       40.6
   b4+       791.78       17.4
   y10++     829.18       23.5
   y3+       846.53       14.0
   b12++     953.18       20.3
   b6+      1034.46       38.0
   a7+      1063.51       20.4
```

```
85. MORG-339_phosphopeptide_TMT_22%ACN_032017_OT.8110.8110.4 (dAsPPPAFkPEPPkAR)

   Ion         m/z   Intensity
   a3++      277.71       39.2
   b6+++     291.86       20.2
   b1+       345.27       34.3
   y4++      350.79       20.8
   y5++      399.44      179.7
   b2+       416.20       20.1
   y8+++     461.18       24.9
   b7++      473.52       24.6
   y9+++     510.23       38.9
   y7++      512.54       53.0
   y10+++    533.86       20.4
   b8++      546.95       26.9
   y11+++    566.31      102.7
   a12+++    582.39       39.3
   b3+       583.29       50.7
   y12+++    598.44      114.6
   y3+       603.28       29.4
   y13+++    631.05      317.5
   b4+       680.71       45.1
   y14+++    686.55       18.3
   y8++      691.29       35.6
   y15+++    710.35       50.8
   a9++      711.36       21.9
   b9++      725.63       46.5
   b14+++    743.07       26.6
   a5+       748.87       32.5
   a15+++    757.77       18.3
   y9++      764.69       51.6
   b15+++    766.72       32.2
   b10++     774.06       36.0
   y5+       797.58       41.5
   y10++     799.90       42.1
   b11++     838.65      143.8
   y11++     848.80      140.2
   y12++     897.28       65.8
   y7+      1023.64       42.3
```

```
86. MORG-339_phosphopeptide_TMT_22%ACN_032017_OT.9260.9260.3 (aEEksPVSINVk)

   Ion         m/z   Intensity
   y1+       376.28       31.8
   b2+       430.36       21.6
   y6++      445.01       17.6
   b4++      458.80       25.0
   y2+       475.49       25.0
   y7++      493.51      167.8
   b5++      542.22       35.1
   b3+       559.33       30.7
   y3+       589.42       37.1
   b6++      590.55       33.7
   a7++      626.12       37.7
   b7++      640.59       70.6
   a8++      669.66       39.4
   b8++      684.15       24.3
   y4+       702.50       41.3
   b9++      740.69       18.1
   y9++      755.68       34.6
   y5+       789.56       52.3
   b10++     797.18       18.5
   y10++     820.26       23.3
   b11++     847.25       34.2
   y6+       888.62       15.4
   y7+       985.68      106.3
```

```
87. MORG-339_phosphopeptide_TMT_10%ACN_032017_OT.15519.15519.3 (aVDssSEEIEVEVPVVDRR)

   Ion         m/z   Intensity
   y5++      322.82       17.4
   y2+       331.27       38.3
   y6++      371.35       60.5
   a2+       372.25       15.7
   b2+       400.56       16.0
   a5++      411.26       40.6
   y7++      421.04       42.1
   y8++      485.49       34.6
   b3+       515.42       36.3
   y9++      535.01       68.2
   y4+       545.35       17.5
   y10++     599.65       38.9
   a4+       654.24       20.4
   y11++     655.85       33.5
   b10++     719.21       25.7
   y12++     720.58       51.7
   y6+       741.52       36.0
   a11++     754.72       16.0
   b11++     768.37       22.8
   y13++     785.06       40.0
   a12++     818.42       24.5
   a5+       821.50       16.4
   y14++     828.81       34.4
   y15++     911.85       28.6
   y16++     995.66       50.2
   b7+      1065.27       25.6
   y9+      1068.77       18.8
   y18++    1102.82       26.9
   b8+      1194.48       44.6
   b9+      1307.67       30.5
   b10+     1436.55       26.6
   b11+     1535.78       31.5
   y13+     1568.75       24.1
   b12+     1664.82       29.8
   b13+     1763.77       32.2
```

```
88. MORG-339_phosphopeptide_TMT_25%ACN_032017_OT.9373.9373.3 (qRsYNLVVAk)

   Ion         m/z   Intensity
   y3++      273.35       27.4
   b3++      341.49       50.6
   b1+       358.27       25.6
   y1+       376.28       60.4
   a4++      408.66       21.3
   b4++      422.55       23.8
   y2+       447.32      116.0
   a5++      465.77       27.6
   b5++      479.97      121.9
   b2+       514.06       52.6
   y7++      518.28       24.0
   a6++      522.59      125.8
   b6++      536.50      201.2
   y3+       546.37      242.7
   b7++      586.03      199.4
   a8++      621.38       36.7
   b8++      635.60      141.2
   y4+       645.48      119.6
   a3+       653.03       29.6
   b9++      671.14       79.2
   y9++      680.32       32.4
   y5+       758.62       62.3
   y6+       872.69       35.7
   y7+      1035.65       46.1
```

```
89. MORG-339_phosphopeptide_TMT_14%ACN_032017_OT.5701.5701.3 (aLsPGREsPk)

   Ion         m/z   Intensity
   a1+       273.25        9.4
   b1+       301.33        9.4
   y1+       376.17       18.9
   y4++      385.11        9.8
   b2+       414.45       21.0
   a6++      432.28       14.0
   b6++      446.32       46.5
   y5++      463.25       23.6
   y2+       473.36      104.8
   y6++      491.45       90.5
   a7++      496.42       77.3
   b7++      511.03       49.6
   y7++      540.41      154.1
   a8++      580.35       16.8
   b3+       581.26       26.1
   b8++      594.19       25.9
   y8++      623.50       16.7
   y3+       640.32       26.0
   y9++      680.46       11.5
   b5+       735.21       13.2
   y4+       769.42       25.3
```

```
90. MORG-339_phosphopeptide_TMT_12%ACN_032017_OT.3620.3620.4 (aVkRPNEDsDEDEEk)

   Ion         m/z   Intensity
   b6+++     375.33       12.1
   y1+       376.32       35.4
   b3++      378.98       18.4
   a7+++     409.41       11.7
   b7+++     418.82       23.8
   b4++      457.11       59.4
   b5++      505.38       36.8
   y10+++    506.40       16.8
   b9+++     512.81       17.9
   b10+++    551.12       36.8
   b6++      562.84       18.5
   a11+++    584.44       17.9
   b11+++    593.96       46.2
   a7++      612.99       35.2
   a12+++    622.69       18.5
   a13+++    665.70       17.7
   b13+++    675.58       39.7
   b14+++    718.48       40.6
   y4+       749.41       32.2
   b3+       757.97       16.8
   y10++     759.65       25.2
   y11++     808.40       18.1
   y5+       878.50       31.7
   b11++     890.67       15.5
   y6+       993.45       21.6
   a14++    1062.51       18.3
   y7+      1160.43       19.9
```

```
91. MORG-339_phosphopeptide_TMT_18%ACN_032017_OT.5292.5292.3 (sRPLNAVSQDGk)

   Ion         m/z   Intensity
   y4++      338.80       17.6
   y1+       376.30       25.3
   y5++      382.42       20.1
   b1+       397.21       22.3
   a5++      425.68       29.5
   y2+       433.35       93.6
   b5++      439.38       42.9
   a6++      461.01       33.2
   y7++      466.96       20.5
   b6++      474.76       54.5
   a7++      510.20       61.6
   y8++      524.43      117.9
   a2+       525.46       30.5
   y3+       548.37       78.0
   b2+       553.27       36.2
   a8++      554.21       21.0
   b8++      567.90       44.5
   y9++      580.46       28.4
   a9++      617.99       20.4
   y10++     629.40       45.8
   b9++      631.86       58.1
   b3+       650.78       19.7
   a10++     675.57       26.0
   y4+       676.40       53.6
   b10++     689.31       88.5
   y11++     707.57       48.6
   b11++     718.16       21.7
   y5+       763.42      111.8
   y6+       862.59       43.3
   y7+       933.60       31.9
   b6+       948.58       18.4
   y8+      1047.54       31.5
```

```
92. MORG-339_phosphopeptide_TMT_10%ACN_032017_OT.6518.6518.3 (rVDLsDEEAk)

   Ion         m/z   Intensity
   b3++      300.61       33.3
   b4++      357.15       12.5
   a1+       357.80        9.1
   y1+       376.42       19.9
   b5++      440.65       11.5
   y2+       447.33       38.3
   b6++      498.38       45.8
   a7++      548.67       16.3
   y7++      550.42       11.5
   y3+       576.42       29.5
   b8++      627.46       44.0
   y9++      658.05        8.4
   b9++      662.98       27.3
   y4+       705.41       44.9
   y5+       820.46       29.7
   y6+       987.41        8.0
   y7+      1100.57       12.6
```

```
93. MORG-339_phosphopeptide_TMT_20%ACN_032017_OT.1407.1407.3 (rRsPSLSSk)

   Ion         m/z   Intensity
   b2++      271.86       35.3
   b3++      354.97       19.3
   y1+       376.37       31.8
   b1+       386.42       24.7
   y6++      424.45       49.9
   b5++      447.02       29.6
   y2+       463.39       56.9
   a6++      489.73       53.4
   b6++      503.95       59.6
   y7++      508.19       80.9
   a2+       514.19       19.5
   b2+       542.08       31.2
   b7++      547.64       22.8
   y3+       550.42       68.8
   y8++      585.77       27.3
   b8++      590.75       16.2
   y4+       663.50       59.7
   b3+       709.45       23.6
   y5+       750.46       25.4
   y6+       847.57       52.1
   y7+      1014.53       22.7
```

```
94. MORG-339_phosphopeptide_TMT_12%ACN_032017_OT.9267.9267.4 (eSDDkPEIEDVGsDEEEEEkk)

   Ion         m/z   Intensity
   y2++      367.59       34.4
   y1+       376.50       23.6
   y3++      432.15       39.8
   y4++      496.69       45.4
   b5++      517.60       54.6
   y5++      560.80       46.3
   b3+       561.68       90.1
   a7++      616.43       22.5
   y6++      625.82       55.2
   b8++      686.77       29.8
   y13+++    687.39       40.8
   y7++      689.97       90.8
   y2+       733.69       23.9
   y8++      747.51       63.7
   b9++      751.40       49.4
   y16+++    800.22       28.9
   b10++     808.94       91.5
   y9++      831.27      107.0
   b19+++    900.22       23.4
   y11++     908.96       79.8
   y17+++    920.17      145.1
   y18+++    958.54       94.5
   y12++     966.72       46.5
   b13++     970.30       29.8
   y4+       991.70       24.4
   b20+++   1020.16       43.3
   y20+++   1025.72       35.1
   b14++    1028.22       65.0
   y13++    1031.10       23.5
   b5+      1033.61       35.0
   y14++    1087.73       61.0
   b15++    1092.77       83.0
   a6+      1102.86       25.6
   a16++    1142.86       28.7
   y15++    1151.88       71.8
   b16++    1156.85       24.2
   y16++    1200.81       45.4
   b17++    1221.72       32.2
   b18++    1286.32       26.9
   b19++    1350.91       28.0
```

```
95. MORG-339_phosphopeptide_TMT_10%ACN_032017_OT.3418.3418.2 (sGYSSPGsPGTPGSR)

   Ion         m/z   Intensity
   b1+       317.32        9.3
   y3+       319.15        7.5
   y4+       416.31       22.4
   a3+       509.30        9.2
   y5+       517.36       10.5
   b3+       537.36        8.5
   y6+       574.62        8.1
   b4+       624.47       12.8
   y7+       671.38       45.0
   a5+       683.29        7.6
   b5+       711.48       15.2
   a6+       780.78       16.5
   b6+       808.52       18.8
   a7+       837.59       14.9
   y8+       838.53       21.1
   y10+      992.55       17.3
   y11+     1079.44        8.8
   b11+     1287.43       12.9
```

```
96. MORG-339_phosphopeptide_TMT_18%ACN_032017_OT.7439.7439.4 (sEsPPPLSEPkQAk)

   Ion         m/z   Intensity
   a6+++     292.52       14.9
   a3++      293.46        8.9
   b3++      306.64       17.3
   b1+       317.20       16.3
   y5+++     343.98       33.5
   b4++      355.26       19.5
   y1+       376.28       17.9
   y6+++     386.91       16.1
   a5++      390.15       16.9
   b5++      403.99       31.9
   y7+++     416.03       15.0
   a6++      438.62       12.6
   b2+       446.28       21.3
   y2+       447.34       14.5
   b6++      452.62       21.7
   y8+++     453.32        8.6
   y4++      467.15       13.5
   y9+++     486.07       36.6
   a7++      495.37       31.1
   b7++      509.37       46.6
   y5++      515.45       81.5
   y10+++    518.41       28.9
   a8++      538.29       16.2
   y11+++    550.76       45.6
   b8++      552.67       22.0
   y3+       575.29       14.6
   y6++      579.95       32.8
   a12+++    597.08       16.5
   b12+++    605.51       12.3
   y12+++    606.16       11.6
   b3+       613.31       20.9
   b9++      617.24       21.3
   y7++      623.54       64.4
   b13+++    629.75       34.7
   b10++     665.51       11.5
   y8++      680.11       28.1
   b4+       710.33       14.9
   y9++      728.56       54.3
   y10++     777.06       17.2
   b5+       807.39       22.6
   b11++     844.48        9.7
   b7+      1017.56       18.0
   b9+      1233.61       17.0
```

```
97. MORG-339_phosphopeptide_TMT_14%ACN_032017_OT.4416.4416.3 (sRsIsPcPkVDSR)

   Ion         m/z   Intensity
   y2+       262.17       44.6
   y6++      465.97       29.9
   b2+       473.46       16.3
   b7++      589.45       27.3
   y8++      594.49       84.5
   a8++      623.95       36.1
   b8++      637.99      208.8
   b3+       640.27       18.9
   y9++      678.32       35.7
   y10++     734.67       19.5
   b4+       753.41       15.4
   b9++      816.79       23.2
   y11++     817.71       16.5
   a10++     851.92       15.8
   b10++     866.13       24.0
   a5+       892.09       37.0
   a11++     909.22       16.8
   b5+       920.43       22.5
   b11++     923.61      122.9
   y6+       930.62       18.6
   b12++     966.81       18.1
   b7+      1177.52       16.1
```

```
98. MORG-339_phosphopeptide_TMT_20%ACN_032017_OT.6914.6914.2 (gRLsPVPVPR)

   Ion         m/z   Intensity
   y2+       272.22       31.5
   b2+       443.43       18.7
   y4+       468.38       69.7
   b3+       556.44       31.1
   y5+       567.10       32.4
   y6+       664.33      119.4
   b4+       723.67       33.1
   y7+       831.47       32.6
   a6+       891.54       25.0
   b6+       919.52       80.7
   y8+       944.58       25.8
   b8+      1115.60       65.6
```

```
99. MORG-339_phosphopeptide_TMT_18%ACN_032017_OT.12356.12356.4 (kRDsEsEsDETPTVPQLk)

   Ion         m/z   Intensity
   b1+       357.88      161.4
   y1+       376.35       71.9
   b7+++     407.62       43.5
   y2+       489.38       87.0
   y7++      506.28       52.2
   y8++      557.15       23.3
   y3+       617.54       56.8
   y14+++    649.83       55.5
   a14+++    668.51       18.8
   b14+++    677.47       24.5
   a15+++    700.54       26.9
   y15+++    706.13       48.8
   b15+++    709.83       21.8
   y4+       714.65      230.1
   a17+++    780.58       29.1
   b17+++    790.03       22.8
   y5+       813.13       46.0
   b11++     867.08      116.8
   a12++     901.60       23.0
   y6+       914.86       27.0
   b5+       925.27       30.9
   y7+      1011.73       25.2
   y15++    1058.84       19.7
   y8+      1112.64       37.8
   y16++    1116.34       19.6
```

```
100. MORG-339_phosphopeptide_TMT_25%ACN_032017_OT.12549.12549.4 (aEkQVDTEEAGMVTAATASNVkAsPk)

   Ion         m/z   Intensity
   y1+       376.47       49.1
   y2+       473.44      192.9
   b5++      508.08       70.6
   y5++      535.25       49.3
   b6++      565.53      207.4
   y15+++    672.14       61.0
   b8++      680.66      110.4
   y8++      685.26       71.6
   y4+       711.50       59.5
   y9++      720.67       57.1
   a17+++    722.01       79.5
   b17+++    731.17       60.0
   a18+++    744.98      129.6
   y18+++    757.71      110.5
   y10++     771.19       86.3
   b10++     780.69      175.2
   y11++     806.69      105.5
   b11++     809.18      210.4
   y20+++    834.74       55.4
   y12++     842.25       98.4
   b21+++    854.99       64.5
   a12++     860.75       92.8
   y21+++    872.90       86.4
   b12++     874.72      236.2
   a4+       887.56       60.8
   y13++     892.91      173.9
   b13++     924.22      229.8
   y14++     942.22       97.0
   b14++     974.28      216.6
   a23+++    988.07       53.9
   b23+++    997.85       84.3
   b15++    1010.27      132.6
   b5+      1014.76      125.7
   y16++    1036.33      104.1
   b16++    1045.86      183.7
   y24+++   1067.77       57.4
   y5+      1068.63       59.2
   y17++    1071.86       65.5
   b17++    1096.31      126.2
   a6+      1101.92       66.9
   b6+      1129.63       59.0
   b18++    1131.89       94.4
   y18++    1136.42       60.6
   y6+      1167.41       49.5
   b19++    1174.98       62.6
   a7+      1202.50      148.8
   b20++    1232.40       86.2
   y20++    1251.39      129.8
   y7+      1281.89      120.7
```

```
101. MORG-339_phosphopeptide_TMT_10%ACN_032017_OT.1935.1935.3 (eEHGGLIRsPR)

   Ion         m/z   Intensity
   y1+       175.12       20.4
   a2++      230.31        9.3
   y2+       272.17      100.0
   a3++      299.52       13.8
   b3++      313.20       16.9
   b4++      341.55       11.6
   y5++      354.86       10.9
   b1+       359.24       15.9
   b5++      370.26       17.1
   a6++      412.70       19.8
   b6++      426.65       28.5
   y7++      439.35       12.0
   a2+       460.49       14.7
   y8++      468.30       30.7
   a7++      469.44       16.5
   b7++      483.25       25.3
   b2+       488.11      256.3
   y9++      536.88       27.8
   b8++      561.35       28.9
   a3+       596.98       58.0
   y10++     601.58       15.3
   b3+       625.42       18.5
   a9++      630.96       10.1
   b9++      644.86       46.5
   a4+       654.49       16.9
   y5+       708.34       27.6
   b5+       739.04       16.1
   y6+       821.40        9.3
   b7+       965.60       14.0
```

```
102. MORG-339_phosphopeptide_TMT_18%ACN_032017_OT.7991.7991.3 (kSsVEGLEPAENk)

   Ion         m/z   Intensity
   y1+       376.31       21.7
   y5++      394.25       42.1
   y2+       490.38       31.9
   y7++      514.98       15.8
   y3+       619.52       35.7
   a4+       683.84       24.1
   y4+       690.47       35.8
   a11++     705.13       24.6
   y11++     741.30       15.1
   y12++     784.68       32.6
   y5+       787.47      117.2
   y6+       916.52       47.3
   y7+      1029.55       32.2
   y8+      1086.66       50.8
   y9+      1215.73       21.3
   y10+     1314.79       22.8
```

```
103. MORG-339_phosphopeptide_TMT_4%ACN_032017_OT.12114.12114.4 (qADVADQQTTELPAENGETENQsPAsEEEk)

   Ion         m/z   Intensity
   y1+       376.27       16.2
   a12+++    500.80       14.8
   y2+       505.35       17.7
   b8++      543.46       28.7
   b3+       544.37       19.5
   y7++      549.81       28.1
   b9++      593.96       50.0
   y8++      633.52       23.2
   y3+       634.48       21.2
   b4+       643.51       26.0
   b10++     644.39       15.2
   y9++      697.60       19.8
   a18+++    700.06       81.1
   b11++     708.66       37.8
   b18+++    709.46       24.3
   b5+       714.44       36.9
   y10++     754.29       20.5
   y4+       763.54       17.7
   b12++     765.62       62.6
   a21+++    814.70       15.4
   y11++     818.77       17.0
   b6+       829.42       24.6
   b14++     849.85       19.1
   y12++     869.31       25.4
   a23+++    913.29       19.5
   b15++     913.96       17.1
   a7+       929.25       31.0
   y5+       930.34       22.8
   y13++     933.92       54.3
   b24+++    954.64       23.7
   b7+       957.52       19.3
   y15++    1019.78       24.0
   b18++    1063.67       14.9
   b8+      1085.56       33.7
   a28+++   1110.79       31.7
   y17++    1119.69       68.6
   b28+++   1120.42       29.7
   a9+      1158.79       14.2
   y29+++   1169.55       61.1
   b20++    1179.41       14.1
   b9+      1186.60       28.1
   y8+      1265.44       18.7
   b11+     1416.78       30.4
   a24++    1417.84       20.5
   b12+     1529.80       41.8
```

```
104. MORG-339_phosphopeptide_TMT_14%ACN_032017_OT.11736.11736.3 (kLEkEEEEGIsQEssEEEQ)

   Ion         m/z   Intensity
   a6++      593.96       45.2
   a7++      658.73       63.2
   b7++      672.40       27.9
   a8++      723.67       47.7
   y6+       868.27       23.9
   y14++     925.23       24.9
   b4+       957.91      218.6
   y15++     989.70       53.2
   y7+       997.20       38.8
   y17++    1233.02       24.2
   y9+      1292.30       50.6
   a7+      1316.57       23.4
   a18++    1381.21       35.3
   a8+      1445.52       50.2
   y11+     1462.51       37.4
   y12+     1591.55       24.5
   y13+     1720.41       23.1
```

```
105. MORG-339_phosphopeptide_TMT_4%ACN_032017_OT.7934.7934.3 (sLDsDEsEDEDDDYQQkR)

   Ion         m/z   Intensity
   b2+       430.36       14.2
   y5++      476.50       61.2
   y6++      534.07       51.8
   b3+       545.42       49.1
   y7++      591.61       58.9
   y8++      649.12       34.7
   y9++      713.71       64.2
   y10++     771.16       44.1
   b5+       827.39       42.9
   y11++     835.43       24.5
   a13++     907.38       15.3
   y12++     919.18       28.1
   y5+       951.67       14.0
   b6+       956.44       25.6
   y13++     983.63       32.9
   y14++    1041.17       33.2
   b15++    1066.69       24.4
   y15++    1124.66       45.0
   y16++    1182.19       16.8
   a8+      1224.38       14.5
   y17++    1238.65       13.6
   b8+      1252.61       14.5
   b17++    1309.27       13.3
   b9+      1367.47       19.6
   b10+     1496.67       15.0
   b11+     1611.54       18.9
   b12+     1726.68       16.3
   b13+     1841.57       16.8
```

```
106. MORG-339_phosphopeptide_TMT_18%ACN_032017_OT.11631.11631.3 (kVVDYSQFQEsDDADEDYGR)

   Ion         m/z   Intensity
   y3+       395.26       58.4
   y4+       510.38       37.0
   y5+       639.33       64.4
   a10++     713.65       81.2
   y6+       754.37       44.1
   y13++     814.04       28.1
   y7+       825.39       73.7
   y14++     877.78       36.7
   b6+       921.65       24.7
   b13++     926.24       31.4
   y8+       940.31       66.2
   y9+      1055.40       49.1
   b16++    1083.83       25.6
   b18++    1222.39       42.3
   a19++    1236.54       32.0
   b9+      1324.98       23.6
   y11+     1351.38       22.3
   a10+     1425.88       33.5
   y13+     1626.62       30.9
```

```
107. MORG-339_phosphopeptide_TMT_10%ACN_032017_OT.4646.4646.3 (rLsPSAsPPR)

   Ion         m/z   Intensity
   y1+       175.11       14.5
   b1++      193.72       22.7
   a2++      236.29       12.7
   b2++      250.28       10.6
   a3++      319.64       16.3
   b3++      333.72       61.5
   y6++      347.35       26.0
   a4++      367.88       24.3
   y3+       369.25       97.5
   b1+       386.34        9.9
   y7++      396.26       47.4
   a5++      412.01       14.1
   b5++      426.04       22.8
   a6++      447.63      101.4
   b6++      461.40       16.8
   b2+       499.45       13.7
   y4+       536.31       22.6
   b7++      544.83       17.9
   a8++      579.32       11.2
   b8++      592.89        9.3
   y5+       607.33       23.4
   b3+       666.38       20.7
   y6+       694.42       27.1
   y7+       791.36       69.4
```

```
108. MORG-339_phosphopeptide_TMT_14%ACN_032017_OT.6165.6165.3 (lQPEQGsPk)

   Ion         m/z   Intensity
   y2++      237.23       29.6
   b3++      285.02        7.9
   y3++      320.73       12.3
   a4++      335.19       11.8
   b4++      349.25       29.6
   y1+       376.35       17.3
   b5++      413.44       47.6
   b6++      441.89       26.1
   a2+       443.06        6.3
   b2+       471.36       32.2
   y2+       473.34       42.5
   y6++      477.40       26.3
   y7++      526.13       29.0
   a3+       540.04        8.9
   y8++      590.42        8.4
   y3+       640.27        9.6
   b4+       697.36       39.2
   b5+       825.45       28.1
   b6+       882.55       17.0
   b7+      1049.55       12.6
   y7+      1051.59        6.8
```

```
109. MORG-339_phosphopeptide_TMT_20%ACN_032017_OT.6342.6342.3 (csSkENILR)

   Ion         m/z   Intensity
   y1+       175.17       16.7
   y2+       288.24       14.1
   y3+       401.31       19.1
   b2+       500.01       13.6
   y6++      501.47       56.4
   b5++      537.16       35.8
   a6++      580.41       13.3
   y5+       644.40       20.2
```

```
110. MORG-339_phosphopeptide_TMT_10%ACN_032017_OT.5308.5308.4 (sAkDsDDEEEVVHVDR)

   Ion         m/z   Intensity
   y1+       175.07       18.6
   y4++      263.69       40.5
   y2+       290.22       20.8
   y5++      313.14       29.3
   y6++      362.79       33.9
   b3++      373.27       26.4
   y3+       389.24       50.5
   y7++      427.37       35.8
   b4++      430.88       59.4
   y8++      491.93       31.4
   b5++      514.53       15.3
   y4+       526.32       93.8
   b10+++    548.66       21.5
   y9++      556.38       37.4
   b6++      571.95       63.5
   a11+++    572.98       81.7
   b11+++    581.87       33.3
   y10++     613.95       52.6
   a7++      615.07       15.1
   y5+       625.38       81.3
   b7++      629.42       60.4
   a13+++    651.53       25.0
   y14+++    660.83       59.3
   y11++     671.47       34.6
   a8++      679.77       23.0
   y15+++    684.79       22.1
   b8++      693.94       78.3
   y6+       724.48       43.9
   b15+++    732.14       69.1
   a9++      744.57       19.4
   b3+       745.70       16.4
   y12++     754.96       38.9
   b9++      758.41       87.2
   a10++     809.03       23.3
   b10++     823.03      144.3
   a4+       832.30       20.7
   y7+       853.45       30.9
   a11++     858.59       30.0
   b11++     872.59      102.5
   b12++     922.02       68.3
   y8+       982.50       24.3
```

```
111. MORG-339_phosphopeptide_TMT_4%ACN_032017_OT.5316.5316.2 (dYDDMsPR)

   Ion         m/z   Intensity
   y2+       272.20      111.9
   b1+       345.25       26.8
   y3+       439.30       23.4
   b2+       508.34       19.4
   y4+       570.23       88.7
   b3+       623.31       95.6
   y5+       685.26       58.3
   b4+       738.27      156.8
   b5+       869.46       34.1
   y7+       963.26       29.5
   b6+      1036.32       33.1
```

```
112. MORG-339_phosphopeptide_TMT_8%ACN_032017_OT.4282.4282.4 (tGEPDEEEGTFRsSIRR)

   Ion         m/z   Intensity
   y3++      222.75       12.7
   a3++      245.19       13.0
   b3++      259.11       17.6
   y2+       331.34       11.4
   b5++      365.38       16.7
   y9+++     387.33       17.9
   b2+       388.29       18.0
   b9+++     391.84       18.2
   b10+++    425.16       16.6
   y6++      427.98       19.0
   y10+++    429.98       38.3
   y11+++    473.41       27.6
   b11+++    474.31       12.4
   a3+       489.67       12.2
   b7++      494.38       39.8
   y7++      501.40       39.8
   y12+++    516.48       35.1
   b3+       517.32       96.1
   y4+       531.51       67.5
   a8++      544.44       20.9
   y8++      551.99       55.8
   y13+++    554.42       38.3
   b8++      558.79       38.2
   y9++      580.53       88.3
   b13+++    582.11       17.6
   b9++      587.15       78.1
   b4+       614.57       18.4
   b10++     638.05       19.5
   y10++     645.06      105.6
   a16+++    691.66       17.3
   b16+++    700.55       35.2
   a5+       701.40       23.0
   y11++     709.57       80.1
   b11++     711.36       27.1
   b5+       729.35       56.4
   y12++     774.11       52.9
   a12++     774.94       23.0
   y13++     831.44       31.6
   a13++     858.39       55.9
   y14++     880.25       23.4
   b7+       987.49       36.2
   b8+      1116.52       13.3
```

```
113. MORG-339_phosphopeptide_TMT_16%ACN_032017_OT.4699.4699.3 (eRsPPLTPk)

   Ion         m/z   Intensity
   y2++      237.27       18.1
   b2++      258.22       22.6
   y3++      287.90       17.4
   a3++      327.74       22.9
   b3++      341.60       59.9
   b1+       359.32       23.7
   y1+       376.30       35.5
   b4++      390.16       15.8
   y5++      392.84       30.5
   a5++      424.89       16.5
   b5++      438.75       13.6
   y6++      441.49      229.5
   y2+       473.41       93.2
   a6++      481.38       38.6
   a2+       487.20       12.5
   b6++      495.16      120.1
   b2+       515.68       36.6
   a7++      531.69       14.1
   b7++      545.94       32.9
   y3+       574.42       61.8
   y8++      602.72       17.2
   a3+       654.21       25.9
   b3+       682.36       32.6
   y4+       687.52       31.3
   y5+       784.53       31.8
   y6+       881.57       64.3
   b6+       989.54       11.7
```

```
114. MORG-339_phosphopeptide_TMT_10%ACN_032017_OT.5838.5838.3 (rSsVSGISLEDNR)

   Ion         m/z   Intensity
   a2++      223.14       11.0
   y2+       289.21       60.0
   a4++      355.78       26.8
   b4++      370.33       10.4
   b1+       386.31       10.4
   a5++      399.63       11.0
   y3+       404.66       53.1
   b5++      413.78       25.3
   b6++      442.21       23.4
   a2+       445.32       10.1
   a7++      484.83       47.9
   b7++      498.86       53.7
   a8++      528.03       15.1
   y4+       533.30       47.1
   b8++      542.29       27.3
   b9++      598.89       15.9
   b3+       640.49       10.6
   y5+       646.36       31.1
   a10++     649.64       10.6
   b10++     663.21       43.3
   y12++     671.99       85.1
   b11++     721.01      108.5
   y6+       733.38       61.7
   b4+       739.54       13.7
   b12++     778.12       20.6
   y7+       846.46       30.0
   y8+       903.47       50.4
   y9+       990.52       43.2
   b7+       996.55       12.9
   y10+     1089.58       24.6
```

```
115. MORG-339_phosphopeptide_TMT_25%ACN_032017_OT.10144.10144.4 (tITVGRPDkAEPRAPsPAPAVsPk)

   Ion         m/z   Intensity
   y1+       376.32       30.8
   y5++      405.34       30.5
   y2+       473.38      139.0
   y7++      489.44       41.9
   b8++      535.51       81.8
   y8++      538.45       29.7
   b11+++    542.68       35.9
   b13+++    627.23       36.7
   y3+       640.39       31.0
   a14+++    641.52       47.0
   b14+++    651.34       94.7
   b5+       701.48       32.0
   y4+       739.24       81.4
   b10++     749.66       40.6
   a17+++    761.86       32.8
   y17+++    779.06      149.1
   y12++     784.28       85.6
   a18+++    785.67       48.7
   b18+++    795.51      243.0
   a11++     800.22       89.0
   y5+       810.53       84.2
   y18+++    811.75       57.6
   b11++     813.96       96.8
   y13++     832.32       36.4
   b20+++    850.95       37.3
   b6+       857.97       38.7
   b12++     862.67       38.4
   a21+++    874.88      177.7
   b21+++    884.50       84.7
   y14++     896.76       30.9
   y6+       907.55      296.5
   b22+++    940.05       53.3
   b13++     940.82       38.8
   b14++     976.19       43.2
   y7+       978.64       67.7
   y23+++    987.65       31.2
   b8+      1069.62       37.0
   y8+      1075.68      101.4
   b16++    1108.33       46.6
   y16++    1110.95       32.6
   y10+     1339.58       36.7
```

```
116. MORG-339_phosphopeptide_TMT_18%ACN_032017_OT.8899.8899.3 (vGSLtPPsSPkTQR)

   Ion         m/z   Intensity
   y2+       303.25       13.8
   b2+       386.34       13.0
   y3+       404.33       16.9
   y5++      429.89       73.1
   b3+       473.39       12.4
   y7++      556.95       27.7
   b4+       586.45       17.6
   y8++      604.99       92.1
   y9++      653.96      190.3
   b10++     656.78       13.1
   y10++     744.43       12.0
   b5+       767.43       39.1
   b11++     835.59       30.9
   y5+       858.72       17.7
   a12++     872.23       12.9
   y13++     873.02       20.2
   b13++     950.18       31.1
```

```
117. MORG-339_phosphopeptide_TMT_6%ACN_032017_OT.5038.5038.2 (sGYSSPGsPGtPGSR)

   Ion         m/z   Intensity
   y3+       319.28        6.2
   y4+       416.22       25.6
   b3+       537.26        9.2
   b4+       624.42       10.7
   y6+       654.45       15.0
   b5+       711.42        7.2
   y7+       751.38        7.7
   b7+       865.13       16.7
   y8+       918.45        5.7
   y9+       975.47       11.7
   y12+     1246.51       10.3
   b11+     1367.47       11.3
   y14+     1466.47        7.8
```

```
118. MORG-339_phosphopeptide_TMT_22%ACN_032017_OT.5703.5703.4 (kAEGEPQEEsPLkSk)

   Ion         m/z   Intensity
   y1+       376.54       25.8
   y3++      411.12       43.8
   y2+       463.77       56.6
   y5++      516.30      241.5
   y9+++     528.61       59.4
   a8++      535.93       32.0
   b8++      550.12       31.0
   y10+++    561.28       45.9
   a4+       587.82       26.3
   y6++      599.68       58.2
   a9++      600.40       58.8
   b4+       615.46       45.2
   b13+++    654.35       28.4
   y8++      728.39       30.3
   a11++     732.27       21.2
   a12++     788.51       25.5
   y9++      792.29       97.3
   b12++     802.83       26.5
   y3+       820.53       26.9
   b6+       841.12       69.8
   y12++     934.18       45.0
```

```
119. MORG-339_phosphopeptide_TMT_10%ACN_032017_OT.7295.7295.3 (sRtAsLTSAASIDGSR)

   Ion         m/z   Intensity
   y6++      317.24       11.3
   y3+       319.19       21.4
   b3++      327.70       18.3
   b4++      363.26       10.8
   y4+       434.24       20.7
   b5++      446.91        9.2
   a7++      539.47       13.7
   y5+       547.40       15.1
   b7++      553.64       12.9
   b8++      597.05       11.7
   a9++      618.50       82.1
   y12++     622.69       14.5
   b9++      632.92       23.3
   y6+       634.30       28.4
   b10++     668.49       22.1
   y7+       705.39       24.4
   b11++     711.59       10.9
   b12++     768.10       16.4
   y8+       776.65       31.4
   b13++     825.88       27.0
   y15++     826.56       14.4
   a14++     840.11       13.6
   b14++     854.39       10.7
   y9+       863.44       29.3
   a5+       864.42       15.2
   y10+      964.41       27.5
   y11+     1077.55       24.1
   a7+      1078.54       11.2
```

```
120. MORG-339_phosphopeptide_TMT_10%ACN_032017_OT.10854.10854.3 (lLPEGEETVEsDDDkDER)

   Ion         m/z   Intensity
   y2+       304.19       26.9
   y4++      388.76       22.2
   y5++      446.37       18.5
   b2+       456.32       20.4
   y6++      503.86       25.3
   y7++      561.56       27.2
   b4+       682.54       15.1
   b5+       739.60       14.5
   b12++     804.53       15.3
   y11++     809.34       20.5
   a6+       840.57       74.9
   a13++     848.37       27.7
   b13++     862.36       20.6
   a14++     905.56       14.9
   y13++     938.77       12.9
   y14++     966.45       17.6
   b7+       997.56       14.3
   y15++    1031.30       30.7
   y16++    1080.29       25.3
   b8+      1098.53       19.7
   y17++    1136.30       16.1
   b16++    1156.34       55.7
   b9+      1197.73       15.9
   b17++    1220.82       21.6
   b10+     1326.69       14.8
   b14+     1838.71       15.4
```

```
121. MORG-339_phosphopeptide_TMT_4%ACN_032017_OT.1204.1204.3 (kREtDDEGEDD)

   Ion         m/z   Intensity
   y2+       249.05       31.6
   b1+       357.93       12.5
   y3+       378.13       10.3
   y4+       435.19       22.5
   b6++      527.64       36.9
   y5+       564.19       38.5
   a7++      577.84      102.9
   b7++      591.84       42.2
   a8++      606.56       12.8
   b3+       643.33       29.2
   y6+       679.24       25.3
   b10++     743.25       12.7
   y7+       794.33       21.8
   a4+       796.23       11.4
```

```
122. MORG-339_phosphopeptide_TMT_8%ACN_032017_OT.6040.6040.3 (sRsRtPLLPR)

   Ion         m/z   Intensity
   y2+       272.14       25.6
   y5++      298.31       18.9
   y3+       385.29       27.8
   b4++      438.92       19.4
   y7++      466.82       26.6
   y4+       498.36       32.2
   b5++      529.13       59.9
   b6++      577.47       17.9
   y5+       595.43      132.5
   y9++      628.47       21.1
   b7++      634.38       35.9
   a8++      676.80       19.1
   b8++      690.96       52.7
```

```
123. MORG-339_phosphopeptide_TMT_22%ACN_032017_OT.9832.9832.3 (aSsQDGLETLkk)

   Ion         m/z   Intensity
   b1+       301.25       15.1
   y2++      367.29       16.3
   y1+       376.38       17.2
   y3++      423.87       16.6
   y4++      474.27       36.7
   y5++      539.03       25.1
   b8++      549.47       25.6
   a9++      585.45       13.0
   y6++      595.70       16.6
   b9++      599.74       14.9
   y7++      623.92       90.1
   a4+       655.62       89.0
   y2+       733.62       19.1
   y9++      745.58       47.4
   a5+       770.15       19.2
   b5+       798.36       48.8
   y10++     828.81       15.6
   b11++     834.78       21.5
   y4+       947.56       14.1
   b7+       968.52       18.8
   b8+      1097.59       20.6
```

```
124. MORG-339_phosphopeptide_TMT_18%ACN_032017_OT.4653.4653.3 (gFsDSGGGPPAkQR)

   Ion         m/z   Intensity
   y6++      463.47       34.8
   y7++      491.84       18.5
   y8++      520.41       28.5
   y9++      549.05       36.1
   b10++     584.62       19.0
   y10++     592.50      115.7
   b3+       601.23      340.9
   a11++     606.42       38.8
   b11++     620.01       24.0
   y11++     649.99       56.3
   y3+       660.49       14.7
   b4+       716.36       55.4
   y12++     733.09       17.7
   a12++     785.28       13.2
   b12++     799.07       16.8
   b5+       803.52       17.2
   y5+       828.71       14.1
   b13++     863.14       19.1
   b8+       974.44       22.2
```

```
125. MORG-339_phosphopeptide_TMT_14%ACN_032017_OT.14139.14139.2 (dLsLEEIQk)

   Ion         m/z   Intensity
   a1+       317.31        8.6
   b1+       345.24       16.9
   y1+       376.33       15.6
   a2+       430.35        7.1
   b2+       458.34       11.1
   y2+       504.31       14.2
   y3+       617.46       21.6
   y4+       746.55       25.2
   b5+       867.49       15.2
   y5+       875.64       19.6
   y6+       988.60       30.3
   b6+       996.51       13.9
   b7+      1109.60       22.0
   y7+      1155.65       10.5
   b8+      1237.68       18.0
   y8+      1268.72        9.7
```

```
126. MORG-339_phosphopeptide_TMT_22%ACN_032017_OT.8232.8232.2 (ssLVTSk)

   Ion         m/z   Intensity
   b1+       317.36       20.8
   y1+       376.33       23.8
   y2+       463.31       27.9
   b2+       484.25       25.3
   y3+       564.42       51.6
   a3+       569.51       19.3
   b3+       596.89       23.5
   y4+       663.42       44.7
   a4+       668.29       18.0
   b4+       696.38       42.2
   a5+       768.96       11.4
   y5+       776.58       53.3
   y6+       943.41       15.7
```

```
127. MORG-339_phosphopeptide_TMT_8%ACN_032017_OT.7602.7602.2 (eVIEAsDk)

   Ion         m/z   Intensity
   b1+       359.16        8.5
   y1+       376.29       19.1
   a2+       430.17        6.1
   b2+       458.40       10.8
   y2+       491.30       13.8
   a3+       543.21        5.4
   b3+       571.41       14.5
   y3+       658.44        5.7
   a4+       672.13        9.9
   b4+       700.50       22.4
   y4+       729.40        8.2
   b5+       771.36        7.2
   y6+       971.44        8.6
   b7+      1053.50       16.7
   y7+      1070.66        7.7
```

```
128. MORG-339_phosphopeptide_TMT_20%ACN_032017_OT.8321.8321.3 (kVELsEsEEDkGSk)

   Ion         m/z   Intensity
   y4++      439.38       39.3
   y6++      561.20       19.4
   y8++      709.61       16.5
   a5+       838.34       28.4
   y10++     856.89       48.6
   b5+       866.74       32.5
   a6+       967.86       14.2
   y5+       992.37       16.3
   y13++    1028.26       24.8
```

```
129. MORG-339_phosphopeptide_TMT_16%ACN_032017_OT.3740.3740.3 (rRsPPEDAIPk)

   Ion         m/z   Intensity
   y5++      386.45       14.7
   y2+       473.38       71.3
   y7++      499.83       14.9
   b6++      516.74       25.7
   b2+       542.40       27.2
   y8++      548.48       82.8
   a7++      560.09       18.5
   b7++      574.45       88.8
   y3+       586.36       41.0
   a8++      596.12       17.9
   b8++      610.09       30.3
   a9++      652.49       27.7
   y4+       657.48       62.0
   b9++      666.50       81.7
   y10++     709.47       30.6
   y5+       772.49       19.9
   y8+      1095.77       18.2
```

```
130. MORG-339_phosphopeptide_TMT_60%ACN_032017_OT.8626.8626.3 (kSsLVTSkLAGGQVE)

   Ion         m/z   Intensity
   y5+       489.30       22.8
   y6+       560.37       21.9
   a8++      671.04       20.0
   y7+       673.26       30.9
   a4+       697.22       19.5
   y12++     716.17       35.2
   b4+       725.36      267.9
   b9++      742.25       35.9
   y13++     799.88       35.0
   y14++     843.07       50.9
   b7+      1013.00       31.8
   y8+      1030.64       21.6
   y9+      1117.63       36.7
   y10+     1218.69       56.2
   y11+     1317.87       26.0
   y12+     1430.87       32.3
```

```
131. MORG-339_phosphopeptide_TMT_25%ACN_032017_OT.6373.6373.3 (kEVQsPEQVkSEk)

   Ion         m/z   Intensity
   y6++      588.85       25.8
   y8++      702.13       97.4
   y9++      785.15       31.6
   y10++     849.88       27.5
   a11++     876.29       27.7
   b5+       881.04       29.3
   b11++     890.18       24.7
   y11++     899.03       35.1
   a6+       950.73       30.2
   b12++     954.43       35.5
```

```
132. MORG-339_phosphopeptide_TMT_25%ACN_032017_OT.15103.15103.3 (kFsEEPEVAANFTk)

   Ion         m/z   Intensity
   y1+       376.28       28.1
   a2+       477.37       47.9
   y9++      603.47       35.9
   y3+       624.47       42.0
   a3+       644.48       44.3
   a10++     685.56       18.5
   y4+       738.50       60.6
   b11++     756.32      250.6
   a4+       773.04       39.7
   y5+       809.45       59.3
   a12++     815.44       18.6
   y6+       880.52       67.4
   y13++     889.38       18.4
   b5+       930.52       20.7
   y7+       979.54       40.0
   y9+      1205.71       93.7
   y10+     1334.77       24.9
   y11+     1463.82       22.4
```

```
133. MORG-339_phosphopeptide_TMT_12%ACN_032017_OT.6002.6002.2 (aGsPDVLR)

   Ion         m/z   Intensity
   y1+       175.12       13.5
   a1+       273.38        9.6
   y2+       288.25       20.6
   b1+       301.34        8.2
   b2+       358.32        7.6
   y3+       387.29       44.1
   y4+       502.24       12.1
   b3+       525.19       22.2
   y5+       599.38       87.9
   b4+       622.59       11.9
   b5+       737.39       43.3
   a6+       808.43       10.7
   y7+       823.50       13.5
   b6+       836.49       33.6
   b7+       949.45       23.2
```

```
134. MORG-339_phosphopeptide_TMT_22%ACN_032017_OT.6969.6969.4 (kEksPELPEPSVR)

   Ion         m/z   Intensity
   a2++      230.15       18.6
   y5++      294.26       18.2
   y6++      342.80       32.7
   y7++      399.39       21.8
   y4+       458.29       82.8
   a2+       459.29       40.1
   b8+++     483.72       19.5
   y9++      512.55       88.7
   a5++      540.72       20.6
   a10+++    549.10       32.2
   b5++      554.90       29.9
   b11+++    587.46       28.3
   b12+++    620.80       25.0
   b7++      675.85       18.0
   y6+       684.42      106.0
   a9++      774.60       20.5
   y7+       797.48       61.9
   y9+      1023.58      123.1
```

```
135. MORG-339_phosphopeptide_TMT_22%ACN_032017_OT.6317.6317.3 (rVPSPtPVPk)

   Ion         m/z   Intensity
   y2++      237.29       18.9
   b3++      291.76       15.7
   y4++      335.29       41.1
   y1+       376.25       16.4
   b1+       386.32       12.0
   y5++      425.38       50.0
   a2+       457.41       14.0
   a6++      460.42       18.0
   y2+       473.34       83.4
   y6++      474.38       42.5
   b2+       485.44       35.0
   a7++      509.07       28.4
   y7++      517.51       34.1
   b7++      522.79       29.3
   a8++      558.14       20.1
   y8++      566.49       38.2
   y3+       572.42       57.4
   b3+       582.29       14.4
   y9++      615.64       12.2
   b9++      621.33       14.8
   a4+       641.13       18.9
   y4+       669.47      110.3
   y5+       850.69       14.4
   y8+      1131.60       11.9
```

```
136. MORG-339_phosphopeptide_TMT_20%ACN_032017_OT.3406.3406.4 (rSsVRPGVVVPR)

   Ion         m/z   Intensity
   y1+       175.20       10.5
   b1++      193.73       11.2
   a2++      223.29        7.9
   y4++      235.76       10.0
   b2++      237.19       12.4
   y2+       272.10       45.4
   a5+++     290.21       11.1
   b5+++     299.33       30.2
   a3++      307.10        9.8
   b3++      320.62       12.1
   y9+++     327.35       14.8
   b6+++     331.95       23.2
   b7+++     350.73       22.9
   y7++      362.24       13.9
   b4++      370.07       11.8
   y3+       371.38       40.3
   a8+++     374.33       36.7
   b8+++     383.71       37.3
   b1+       386.27       17.9
   y11+++    411.96        7.5
   b9+++     416.82       20.1
   y8++      440.50       13.7
   a2+       445.58        9.9
   b5++      448.35       20.8
   b10+++    449.80       20.3
   y4+       470.33       34.3
   y9++      490.04       37.8
   a7++      511.45        8.9
   b7++      525.50       35.2
   a8++      560.82       18.5
   y5+       569.49        9.4
   b8++      575.04       65.1
   a9++      610.57       14.3
   y11++     616.39       10.4
   b9++      624.59       55.0
   y6+       626.42        7.8
   b10++     674.10       13.4
   y7+       723.47       18.2
```

```
137. MORG-339_phosphopeptide_TMT_25%ACN_032017_OT.16390.16390.4 (eLEkRAsGQAFELILSPR)

   Ion         m/z   Intensity
   y2+       272.19       22.2
   b1+       359.24       24.4
   b5+++     372.47       19.7
   b7+++     451.20       12.5
   b2+       472.34       39.6
   a9+++     504.39       34.3
   b9+++     513.35       51.7
   a10+++    527.51       17.5
   b10+++    537.13       42.7
   a5++      544.03       16.0
   a11+++    576.82       22.9
   a6++      578.92       20.9
   y5+       585.38       36.4
   b11+++    586.18       45.7
   y10++     587.16       20.9
   b3+       601.42       24.9
   y11++     615.42       31.3
   b12+++    628.93       80.1
   a13+++    657.20       17.9
   y15+++    661.38       29.9
   b13+++    666.82       59.9
   a14+++    695.18       31.5
   y6+       698.53       33.8
   y12++     699.56       21.4
   b14+++    704.64       41.1
   b8++      705.55       15.1
   b15+++    742.17       30.7
   b9++      769.69       17.3
   b16+++    771.11       24.5
   b10++     805.14       19.5
   y7+       827.53       29.8
   b11++     878.54       13.6
   b12++     942.97       27.6
   y8+       974.53       39.8
   b13++     999.52       27.7
   a14++    1042.16       13.4
   y9+      1045.73       31.2
   b14++    1056.23       14.3
   y11+     1230.74       22.6
```

```
138. MORG-339_phosphopeptide_TMT_20%ACN_032017_OT.8990.8990.4 (tVSDNsLsSSkEGkPELR)

   Ion         m/z   Intensity
   y2+       288.30       28.0
   a6++      442.97       16.3
   b6++      456.88       25.9
   y6++      465.06       32.5
   y8+++     472.44       13.3
   a7++      499.43       23.2
   y9+++     501.58       24.1
   y4+       514.42       22.6
   y7++      529.56       16.5
   y10+++    530.61       30.0
   b8++      597.43       70.1
   b4+       632.51       13.8
   b13+++    637.49       16.9
   b9++      641.15       18.5
   a10++     670.17       43.5
   y13+++    679.39       30.3
   b10++     683.79       24.7
   b5+       746.68       15.6
   y9++      751.17       18.1
   b14+++    757.00       35.8
   a15+++    779.31       17.8
   y16+++    784.66       14.1
   y10++     794.78       17.0
   b16+++    832.39       53.8
   b11++     863.37       14.2
   b17+++    869.90       29.1
   y11++     878.59       25.9
   a12++     913.63       14.0
```

```
139. MORG-339_phosphopeptide_TMT_8%ACN_032017_OT.3691.3691.3 (gTDDsPkNSQEDLQDR)

   Ion         m/z   Intensity
   b3+       503.30       39.2
   y4+       531.39       58.5
   b4+       618.36       53.5
   b7++      620.48       28.7
   y5+       646.37       28.9
   a8++      663.21       12.2
   b8++      677.48       26.6
   y10++     731.62       29.2
   a5+       757.07       13.1
   a10++     771.17       52.6
   b11++     849.21       29.0
   a6+       854.65       33.1
   y12++     863.50       53.8
   y7+       903.45       16.4
   b12++     907.04       52.9
   y13++     921.07       41.1
   b13++     963.60       16.2
   y14++     978.41       14.3
   y8+       990.45       27.3
   b14++    1027.53       31.6
   y15++    1028.52       49.3
   b15++    1085.14       51.1
   y9+      1104.50       31.5
```

```
140. MORG-339_phosphopeptide_TMT_22%ACN_032017_OT.14245.14245.2 (lATTVsAPDLk)

   Ion         m/z   Intensity
   b1+       343.28       22.3
   y1+       376.35       24.2
   b2+       414.33       20.6
   y2+       489.42       67.4
   b3+       515.40       29.4
   y3+       604.32       12.6
   b4+       616.53       26.2
   y4+       701.49       61.7
   b5+       715.50       45.3
   y5+       772.55       61.2
   a6+       854.23       16.1
   y6+       939.51       38.4
   b7+       953.49       21.0
   y7+      1038.50       16.7
   b8+      1050.62       12.5
   y8+      1139.65       22.5
   b9+      1165.58       50.6
   y9+      1240.68       19.4
   b10+     1278.65       28.8
   y10+     1311.70       24.4
```

```
141. MORG-339_phosphopeptide_TMT_20%ACN_032017_OT.7787.7787.4 (nRPPLAAGANSkGPPDFsSDEER)

   Ion         m/z   Intensity
   y2+       304.16       44.8
   y3+       433.28       50.6
   b2+       500.39       23.5
   b9++      539.75       24.2
   b12+++    546.15       64.8
   y4+       548.24       47.8
   b13+++    565.15       73.8
   b10++     596.58       26.5
   b3+       597.50       24.6
   y10++     629.74       32.7
   y5+       635.64       36.3
   b16+++    668.37      187.1
   a17+++    707.39       54.7
   b17+++    717.37       80.8
   a18+++    763.31       48.2
   y20+++    785.21       23.2
   y6+       801.77       59.0
   b5+       807.67      116.7
   b12++     818.27       28.1
   a20+++    831.15       29.4
   a13++     833.44       34.4
   y12++     836.93       24.6
   b20+++    840.26       90.8
   b13++     847.27       33.7
   a6+       850.67       91.3
   b21+++    883.30       90.0
   b22+++    926.29       66.6
   b7+       949.34       64.3
   b16++    1001.46       25.7
   y9+      1161.44       23.2
   b20++    1259.49       27.2
```

```
142. MORG-339_phosphopeptide_TMT_25%ACN_032017_OT.17065.17065.3 (lkQLSVPAsDEEDEVPAPIPR)

   Ion         m/z   Intensity
   b2++      350.81       26.5
   b3++      414.87       19.6
   b4++      471.74       33.0
   y4+       482.43       36.4
   a5++      500.86       49.8
   b5++      514.60       37.3
   a6++      550.44       23.4
   b6++      564.47      100.2
   y6+       650.46       69.3
   y7+       749.44       55.8
   b10++     789.56       24.0
   y14++     802.64       66.6
   y15++     851.73       63.3
   y8+       878.55       46.7
   b13++     976.09       48.7
   y9+       993.36       47.3
   a14++    1026.60       21.8
   b5+      1028.77       43.7
   b15++    1090.34       65.8
   a6+      1099.75       33.5
   y10+     1122.47       24.4
   b6+      1127.79       68.6
   b17++    1174.27       23.3
   y11+     1251.70       31.3
   b19++    1279.25       55.2
   y12+     1366.98       19.9
   y14+     1604.86       69.8
```

```
143. MORG-339_phosphopeptide_TMT_10%ACN_032017_OT.5406.5406.3 (skPAAADsEGEEEEEDTAk)

   Ion         m/z   Intensity
   b2++      337.70       11.2
   y1+       376.30       11.0
   b5++      457.37       14.1
   y3+       548.45       25.9
   b7++      550.48       37.8
   y4+       663.46       20.5
   b2+       674.60       17.9
   b11++     791.22       17.1
   y5+       792.30       25.9
   b4+       842.30       11.4
   b12++     855.72       24.2
   a5+       885.33       23.6
   y13++     888.89      120.5
   a13++     906.01       25.6
   b5+       913.48       13.4
   b14++     985.12       24.4
   b15++    1049.21       22.3
   y7+      1050.39       26.0
   b7+      1099.67       17.1
   b16++    1106.85       20.2
   y8+      1179.48       12.7
   b8+      1266.79       13.0
   y10+     1365.69       14.9
   y11+     1494.72       11.9
   y12+     1661.71       13.7
```

```
144. MORG-339_phosphopeptide_TMT_18%ACN_032017_OT.7670.7670.3 (tSsLPEsSPSkTAk)

   Ion         m/z   Intensity
   y1+       376.38       14.6
   y2+       447.43       14.1
   y6++      545.41       86.5
   b7++      546.46       20.2
   y3+       548.50       17.7
   y8++      672.43       21.9
   b10++     681.28       26.3
   b4+       698.38       18.3
   y9++      737.18       41.7
   y10++     785.43       45.6
   y11++     842.11       37.2
   a11++     846.54       17.8
   b11++     860.38       15.9
   b12++     911.08       14.8
   b6+       924.46       19.5
   y12++     925.53       14.0
   a7+      1063.49       20.5
```

```
145. MORG-339_phosphopeptide_TMT_8%ACN_032017_OT.1745.1745.3 (sPERPTGDLRER)

   Ion         m/z   Intensity
   y1+       175.22       17.0
   y4++      287.20       34.3
   a4++      376.29       12.2
   b4++      390.26       42.1
   b1+       397.17       40.2
   y8++      472.57       27.7
   a7++      503.39       22.4
   y9++      550.33       52.0
   a8++      561.33       15.8
   b3+       623.26       27.7
   y11++     663.52       79.8
   y5+       688.46       14.2
   a10++     695.47       31.2
   b10++     709.97       20.8
   y6+       745.24       13.9
   a11++     760.67       21.1
   b11++     774.48       81.1
   b4+       779.36       18.6
   y8+       943.47       33.8
   a6+       949.47       12.2
   b8+      1149.57       24.4
```

```
146. MORG-339_phosphopeptide_TMT_14%ACN_032017_OT.8262.8262.3 (aksPTPsLsPAR)

   Ion         m/z   Intensity
   a2++      315.70       11.6
   y3+       343.23       32.2
   b3++      413.34       30.1
   y8++      494.43       17.3
   y4+       510.24       19.8
   b5++      512.61       14.5
   y9++      543.39       18.4
   a6++      546.36       10.7
   y5+       623.38       19.5
   a2+       630.38       16.0
   b2+       658.43       10.1
   a8++      686.48       35.5
   b8++      701.12       16.0
   a9++      770.71       11.2
   b9++      784.41       10.0
   a3+       797.40       13.2
   y11++     805.39        9.9
   a10++     819.21       13.1
   b3+       825.67       12.1
   b11++     868.24       19.2
   y7+       887.42       14.0
   y8+       988.34       12.4
   b5+      1023.51       18.7
   y9+      1085.40       32.2
```

```
147. MORG-339_phosphopeptide_TMT_4%ACN_032017_OT.6527.6527.3 (rAsVcAEAYNPDEEEDDAESR)

   Ion         m/z   Intensity
   b5++      442.69        8.0
   y4+       462.36        9.6
   b6++      477.78       10.0
   a7++      528.61       17.3
   b7++      542.26       15.9
   b8++      577.47       22.8
   a9++      645.50       12.7
   y11++     646.68       17.4
   b9++      659.34       22.7
   y6+       692.42        8.2
   a10++     702.55       10.7
   b10++     716.59       41.8
   b11++     765.03        7.3
   a12++     808.52        8.1
   y7+       821.61       16.2
   b12++     822.66       22.4
   a5+       855.81        9.6
   a13++     872.58       11.3
   y15++     884.74        8.6
   b13++     887.16       11.6
   y16++     920.78        8.4
   a14++     937.10        6.7
   y8+       950.64       16.9
   b14++     951.55       15.4
   b15++    1016.26       18.6
   b16++    1073.89       29.3
   y9+      1079.54       22.4
   b7+      1083.61        9.1
   a17++    1116.96        6.9
   a8+      1126.95        8.7
   b17++    1131.44       20.0
   y20++    1169.39        8.9
   y10+     1194.55        8.7
   b19++    1231.36       24.2
   b20++    1274.53       12.8
   y11+     1291.54       33.3
   b9+      1317.51        9.7
   y12+     1405.52       18.6
   y13+     1568.64       12.7
   y14+     1639.63       11.3
   y15+     1768.99        6.9
   y16+     1839.89        8.8
```

```
148. MORG-339_phosphopeptide_TMT_12%ACN_032017_OT.3957.3957.2 (ssVPGVR)

   Ion         m/z   Intensity
   y1+       175.14       14.9
   y2+       274.04        8.1
   b1+       317.19       15.2
   y3+       331.29       10.1
   y4+       428.29       65.8
   b2+       484.13       41.0
   y5+       527.34       57.4
   a3+       555.53       11.1
   b3+       583.35       28.0
   y6+       694.37       13.5
   b5+       737.41       14.1
   a6+       808.46       11.4
   b6+       836.52       30.1
```

```
149. MORG-339_phosphopeptide_TMT_20%ACN_032017_OT.13756.13756.2 (sGDALTTVVVk)

   Ion         m/z   Intensity
   y1+       376.38       33.1
   b1+       397.12       20.2
   y2+       475.44       27.3
   b3+       569.22       43.5
   y3+       574.44       36.0
   b4+       640.35       24.8
   y4+       673.54       25.0
   a5+       725.52       18.6
   b5+       753.32       33.7
   y5+       774.62       83.6
   a6+       826.62       10.3
   y6+       875.62       45.1
   y7+       988.75       33.6
   b8+      1054.54       14.1
   y8+      1059.74       59.5
   b9+      1153.55       31.4
   y9+      1174.59       12.8
   y10+     1231.76       37.3
   b10+     1252.69       30.0
```

```
150. MORG-339_phosphopeptide_TMT_20%ACN_032017_OT.6251.6251.3 (sYsPDGkEsPSDkk)

   Ion         m/z   Intensity
   y2++      367.53       31.1
   y1+       376.30       13.9
   y4++      468.45       12.8
   y5++      516.94       44.1
   b7++      637.71       14.8
   a8++      687.66       21.1
   b8++      702.03       18.2
   y2+       733.64       29.1
   a10++     819.60       23.1
   a5+       831.57       15.8
   b10++     833.82       19.7
   y8++      843.57       37.3
   y3+       848.52       27.4
   b5+       859.42       48.3
   y9++      872.17       14.9
   b11++     877.68       14.2
   y10++     929.37       41.9
   b12++     934.83       12.7
   y4+       935.53       14.0
   y11++     977.98       34.3
   y5+      1032.67       31.3
   b13++    1113.83       16.3
   y13++    1142.76       13.1
   b8+      1402.74       13.1
   b12+     1868.75       18.8
```

```
151. MORG-339_phosphopeptide_TMT_12%ACN_032017_OT.7675.7675.3 (eIQSITDESRGsIR)

   Ion         m/z   Intensity
   y2+       288.25       30.5
   y6++      378.44       12.9
   y7++      442.68       16.5
   b6++      451.38       19.8
   y8++      500.41       14.1
   y4+       512.30       11.2
   a9++      602.94       35.2
   y10++     607.56       32.4
   b4+       687.38       18.2
   b10++     695.01       22.5
   b11++     723.11       20.3
   b5+       800.64       17.7
   b12++     807.21       15.3
   y7+       884.49       13.1
   y8+       999.25       15.0
   b7+      1016.56       29.4
   y9+      1100.67       12.4
   b8+      1145.58       15.6
```

```
152. MORG-339_phosphopeptide_TMT_4%ACN_032017_OT.8635.8635.3 (eGEEPTVYsDDEEPk)

   Ion         m/z   Intensity
   b1+       359.23        5.6
   y1+       376.27        8.0
   b2+       416.20       10.3
   y2+       473.42       50.3
   b7++      486.51        6.6
   b3+       545.35       12.3
   y7++      564.97        7.1
   b8++      567.87        5.9
   y3+       602.45       20.8
   a9++      637.47        8.4
   a4+       646.46       19.6
   b4+       674.37       29.3
   y9++      695.66       10.1
   y4+       731.68        8.7
   a5+       743.55        9.3
   y10++     745.90        9.9
   y11++     795.00       16.9
   a12++     817.01        5.5
   b12++     830.89       20.1
   a6+       844.62        6.8
   y5+       846.47       30.9
   y12++     859.42        7.8
   b6+       872.43       15.0
   b13++     895.51       39.4
   y6+       961.46        5.1
   b7+       971.54       36.0
   y7+      1128.49        6.7
   b8+      1134.53       25.0
   y8+      1291.57        9.8
   b10+     1416.69        8.7
   b11+     1531.65       12.0
   b12+     1660.72       12.0
   b13+     1789.84       11.0
```

```
153. MORG-339_phosphopeptide_TMT_18%ACN_032017_OT.7334.7334.3 (skDAsPINRWsPTR)

   Ion         m/z   Intensity
   y3+       373.25       23.4
   b3++      395.41       46.8
   y4+       540.40       15.0
   y8++      555.09       38.4
   y9++      604.07       35.6
   b7++      619.65       18.8
   b2+       674.58       47.8
   b8++      676.69       20.0
   y10++     686.90       31.1
   y11++     722.56       35.7
   y5+       726.37       48.1
   b3+       789.51       41.4
   a4+       832.79       18.0
   a10++     833.58       34.1
   b10++     847.81       29.7
   y6+       882.23       58.2
   b11++     930.56       25.2
   y7+       996.50       17.2
   y9+      1206.59       23.1
   y11+     1444.57       25.6
```

```
154. MORG-339_phosphopeptide_TMT_16%ACN_032017_OT.5400.5400.3 (aRVDHGAEIITQsPSRSsVASPR)

   Ion         m/z   Intensity
   y2+       272.15       30.2
   y9++      513.37       35.5
   b8++      533.28       25.4
   y10++     562.55       32.5
   b9++      590.01       24.4
   b10++     646.29       24.9
   b4+       671.45       89.7
   b11++     696.68       25.3
   b12++     761.05       43.5
   y7+       783.44       36.0
   b5+       808.15       28.2
   y15++     872.95      196.7
   a14++     878.46      142.6
   b14++     893.20       76.0
   a7+       908.94       25.8
   y18++    1002.06       29.9
   b16++    1014.75       62.9
   a17++    1043.71       23.8
   b8+      1065.63       23.8
   y19++    1070.72       80.3
   y10+     1123.54       43.7
   y21++    1177.82       34.4
   b9+      1178.58       29.9
   b19++    1191.46       33.5
   a20++    1212.34       57.5
   b21++    1269.85       28.2
   b10+     1291.80       32.8
   b12+     1520.90       24.4
```

```
155. MORG-339_phosphopeptide_TMT_20%ACN_032017_OT.12044.12044.2 (sGILVPSEk)

   Ion         m/z   Intensity
   y1+       376.40       18.1
   b1+       397.16       15.2
   a3+       539.34       23.6
   b3+       567.31       32.6
   a4+       652.47       11.7
   b4+       680.42       37.0
   y4+       689.43       79.9
   a5+       751.44       19.0
   b5+       779.46       64.6
   y5+       788.52       38.0
   y6+       901.57       38.0
   y7+      1014.72       16.0
   y8+      1071.67       36.7
   b8+      1092.60       13.2
```

```
156. MORG-339_phosphopeptide_TMT_12%ACN_032017_OT.6737.6737.4 (aRVDHGAEIItQSPSRSsVAsPR)

   Ion         m/z   Intensity
   y2+       272.20       48.2
   a1+       273.25       16.4
   b4++      336.32       68.4
   y3+       439.31       21.5
   b7++      468.95       14.5
   y8++      510.34       32.3
   a12+++    524.73       16.4
   y13+++    533.44       25.1
   a13+++    553.52       52.6
   b9++      590.07       31.3
   y10++     602.50       57.5
   y11++     645.83       27.1
   b10++     646.68       33.1
   a16+++    667.11       34.1
   b4+       670.99       54.2
   y17+++    675.87       33.5
   b16+++    676.87       53.4
   y18+++    694.64      114.3
   a17+++    696.39       36.4
   b17+++    706.12       40.1
   y12++     710.07       14.7
   y19+++    740.84       15.7
   a18+++    751.85       27.6
   b18+++    760.89       24.9
   y6+       776.15       57.9
   y20+++    779.30       24.2
   a5+       780.08       17.4
   a12++     786.89       28.9
   b19+++    794.64       38.7
   a20+++    808.68      118.9
   y21+++    812.19       19.3
   b20+++    818.35       49.8
   b13++     844.62       17.0
   y7+       863.35       24.2
   a21+++    864.50       21.2
   b21+++    874.07       24.4
   y17++    1013.55       19.5
   y19++    1110.70       55.9
```

```
157. MORG-339_phosphopeptide_TMT_6%ACN_032017_OT.8581.8581.3 (sksPPksPEEEGAVSS)

   Ion         m/z   Intensity
   a2++      363.31       26.5
   b2++      377.53       12.2
   y6+       549.36       14.1
   y7+       678.42       19.1
   a6++      722.74       45.6
   a2+       726.48       15.6
   b6++      737.03       35.3
   b2+       754.34       13.3
   y12++     763.24       36.2
   y8+       807.67       77.6
   y13++     811.76       56.5
   b7++      820.61       17.8
   y9+       904.39       33.5
   b3+       921.49       16.1
   b9++      933.73       29.5
   b10++     998.17       39.2
   b11++    1062.77       25.8
   y10+     1071.49       16.4
   a12++    1077.35       12.7
   b12++    1091.49       14.5
   y12+     1525.74       16.4
   y13+     1622.86       27.4
```

```
158. MORG-339_phosphopeptide_TMT_12%ACN_032017_OT.9733.9733.3 (tREtVsIEVk)

   Ion         m/z   Intensity
   a2++      230.25        5.5
   b3++      308.72        6.3
   y1+       376.38       14.4
   a4++      385.34        6.8
   a5++      434.23        8.5
   b5++      448.76       10.3
   a2+       458.83        7.0
   y2+       475.37       17.7
   b2+       487.58        6.2
   y6++      492.50        6.6
   y7++      582.47       13.7
   y3+       604.46       26.3
   b8++      653.42       25.1
   a9++      688.78        6.8
   b9++      702.84       14.7
   y4+       717.58       19.9
   y5+       884.47       19.2
   y6+       983.58        7.8
```

```
159. MORG-339_phosphopeptide_TMT_16%ACN_032017_OT.8683.8683.2 (eVQTsELk)

   Ion         m/z   Intensity
   a1+       331.23       11.9
   b1+       359.28       18.2
   y1+       376.25       21.7
   a2+       430.36        9.2
   b2+       458.35       22.3
   y2+       489.37       27.1
   a3+       558.61        8.0
   b3+       586.32       20.9
   y3+       618.37       23.2
   a4+       659.12       15.6
   b4+       687.52      227.0
   y4+       785.39       20.4
   y5+       886.46       19.2
   b6+       983.26       10.5
   y6+      1014.94        9.4
   b7+      1096.56       25.5
   y7+      1113.56       15.8
```

```
160. MORG-339_phosphopeptide_TMT_20%ACN_032017_OT.7714.7714.3 (rSSAsVSLPPGTPEk)

   Ion         m/z   Intensity
   y6++      429.32       19.4
   b6++      449.24       18.7
   y7++      477.93      106.7
   a7++      478.99       26.5
   b7++      492.67       15.7
   y8++      534.27       19.6
   a8++      535.33       22.2
   b8++      549.35       37.2
   b9++      597.83       34.9
   y3+       602.44       48.6
   a4+       603.34       19.9
   b4+       631.11       15.2
   b10++     646.82       16.4
   a11++     660.57       38.4
   b11++     674.62       30.1
   y4+       703.52       26.1
   b12++     725.52       32.1
   b5+       798.77       13.6
   b14++     838.58       15.3
   y6+       857.50       38.1
   b6+       897.56       17.6
   y7+       954.58       71.5
   a7+       956.57       14.5
   y8+      1067.77       18.1
   b8+      1097.59       66.1
   y9+      1154.74       15.1
```

```
161. MORG-339_phosphopeptide_TMT_16%ACN_032017_OT.3664.3664.4 (dkkAEGAGTEEEGtPk)

   Ion         m/z   Intensity
   y2++      237.23       48.2
   y1+       376.33       43.7
   b4+++     377.38       24.2
   a5+++     411.58       21.4
   y5++      420.75       43.4
   b7+++     463.36       22.4
   y2+       473.35      147.8
   y6++      485.35       23.9
   a3++      516.07       22.1
   y7++      549.93       64.4
   b10+++    559.28       57.7
   a11+++    592.75       27.9
   b11+++    602.22      100.6
   a5++      615.93       26.5
   y9++      629.18       56.8
   b5++      630.17       46.4
   a12+++    635.55       37.0
   b12+++    645.14      133.5
   y14+++    648.08       38.5
   a13+++    654.37       25.9
   a7++      680.52       23.9
   y11++     692.93       84.2
   b7++      694.15       29.5
   y4+       711.36       52.6
   b8++      722.81       20.1
   b14+++    724.43       51.9
   y12++     757.09       30.1
   a9++      759.26       23.5
   y15+++    767.14       38.6
   b9++      773.60       41.2
   a10++     823.68       32.7
   b10++     838.22       72.7
   b11++     902.65       91.4
   b12++     967.19       90.9
   y6+       969.38       21.1
   b13++     995.69       69.4
```

```
162. MORG-339_phosphopeptide_TMT_18%ACN_032017_OT.21715.21715.3 (sLmssPEDLTkDFEELk)

   Ion         m/z   Intensity
   y1+       376.21       10.9
   b2+       430.51       10.2
   y2+       489.36        8.0
   b3+       577.43        7.9
   y3+       618.51       11.9
   y7++      683.81       13.4
   a4+       716.46       15.3
   b10++     733.66        8.2
   y8++      734.72       17.7
   y4+       747.63       12.5
   y9++      790.96       29.0
   y10++     848.35       18.0
   y5+       894.51       22.8
   y11++     912.55      131.2
   y12++     961.82       50.2
   b12++     970.14       18.8
   y6+      1009.63       14.3
   b13++    1043.65       13.0
   y13++    1045.15       18.4
   b14++    1108.34       16.1
   a7+      1109.27        8.3
   y14++    1128.37        8.2
   b15++    1172.47        7.9
   y15++    1202.01        8.4
   b16++    1229.42       15.0
   b8+      1252.51       15.0
   y16++    1258.95        8.8
   y7+      1366.57        8.4
```

```
163. MORG-339_phosphopeptide_TMT_10%ACN_032017_OT.776.776.4 (eRPPRARPAstSPsPGTRGR)

   Ion         m/z   Intensity
   y6++      322.38       68.3
   b7+++     365.01       19.6
   y7++      405.46       55.8
   b9+++     421.32       26.4
   y8++      454.22       35.7
   a6++      454.99       28.9
   y12+++    471.70       22.4
   b10+++    476.95       19.9
   y13+++    504.35       19.8
   b2+       515.28       20.5
   a11+++    527.46       24.2
   b7++      547.07       62.6
   b12+++    566.18       28.9
   y15+++    580.19       22.5
   a8++      581.07       25.6
   a3+       583.90       32.8
   y10++     587.99       34.7
   a13+++    588.89       30.2
   b8++      594.89       46.9
   b13+++    598.16      131.3
   b3+       612.38       34.5
   a9++      617.10       26.4
   b9++      631.18       53.2
   y16+++    631.83       57.2
   y6+       643.45       32.1
   y17+++    664.10       25.2
   y12++     706.76       32.9
   y19+++    749.09       24.4
   y13++     756.02       52.4
   b18+++    791.16       38.1
   a19+++    800.50       37.0
   b19+++    809.62       36.3
   y7+       810.54       28.9
   b12++     848.55       26.7
   y15++     869.58       19.0
   a13++     882.68       46.7
   y8+       907.40       41.4
   a6+       908.40       27.2
```

```
164. MORG-339_phosphopeptide_TMT_10%ACN_032017_OT.7905.7905.3 (tASLTSAAsIDGSRSR)

   Ion         m/z   Intensity
   y2+       262.37       16.8
   y5++      281.70       48.4
   y6++      339.18       24.9
   y7++      395.81       48.7
   a2+       454.22       14.4
   b7++      471.29       27.1
   y8++      479.20       28.4
   y9++      514.81       24.6
   a3+       541.42       16.9
   y10++     550.22       27.9
   b3+       568.81       17.0
   b9++      590.04       25.5
   y11++     593.86       41.3
   a10++     632.53       23.4
   y12++     644.32       31.0
   b10++     646.83       34.1
   y13++     700.83       40.3
   b11++     704.03       35.3
   b13++     776.37       15.2
   b5+       783.53       15.0
   y8+       957.39       18.5
   b8+      1012.66       18.9
   b11+     1407.74       28.8
```

```
165. MORG-339_phosphopeptide_TMT_25%ACN_032017_OT.9783.9783.3 (eVQsPEQVkSEkLk)

   Ion         m/z   Intensity
   y6++      710.73       30.1
   y7++      760.31       26.5
   a9++      768.58       17.4
   b9++      782.61       19.7
   y8++      824.22       45.9
   b10++     826.18       20.4
   y3+       846.68       40.9
   b5+       850.50       56.5
   y10++     937.05       87.4
   b6+       979.30       22.7
   y11++    1020.22       17.8
   y5+      1062.45       21.2
   y12++    1084.83       21.9
   b7+      1107.57       18.5
   b13++    1125.58       18.0
   y13++    1134.37       19.3
```

```
166. MORG-339_phosphopeptide_TMT_20%ACN_032017_OT.11943.11943.3 (asNEDVTkAVDITTPk)

   Ion         m/z   Intensity
   y1+       376.33       22.2
   b2+       468.23       14.7
   y2+       473.32       59.1
   y3+       574.34       31.9
   y4+       675.52       40.2
   a4+       683.37       19.7
   b8++      692.32       20.9
   b4+       711.36       19.9
   a9++      714.09       15.6
   y9++      716.38       28.9
   b9++      728.27       61.4
   a10++     763.81       22.1
   y10++     766.37       30.6
   b10++     777.61       62.3
   y5+       788.51       79.0
   a5+       798.68       36.6
   b5+       826.18       20.6
   b11++     835.14       77.2
   y12++     873.46       26.1
   b12++     891.64       54.7
   a6+       897.37       14.7
   y6+       903.61       41.9
   b6+       925.64       29.7
   y13++     938.31       21.8
   b14++     992.87       22.9
   y14++     995.38       55.7
   y7+      1002.64       30.3
   y8+      1073.75       27.2
```

```
167. MORG-339_phosphopeptide_TMT_16%ACN_032017_OT.6317.6317.3 (sFSkEVEER)

   Ion         m/z   Intensity
   y1+       175.12       20.2
   y2+       304.16       54.9
   b1+       397.29       30.3
   y3+       433.18       18.8
   b4++      494.90       17.9
   y6++      509.77       23.6
   a2+       515.84       11.2
   y4+       532.29       42.2
   b2+       544.27       68.6
   a5++      545.03       19.8
   b5++      559.55       34.0
   a6++      594.56       13.1
   b6++      608.81       32.1
   y8++      627.01       54.3
   a7++      659.59       21.1
   y5+       661.43       31.1
   b7++      673.42      102.8
   a8++      724.00       12.2
   b8++      737.79       48.6
```

```
168. MORG-339_phosphopeptide_TMT_4%ACN_032017_OT.6320.6320.3 (qSNASsDVEVEEk)

   Ion         m/z   Intensity
   y3++      317.23        5.0
   b1+       358.19       15.4
   b5++      358.90        5.3
   y4++      367.24        6.0
   y1+       376.22       36.9
   y5++      431.93       15.0
   b2+       445.30        5.1
   b7++      500.57       12.1
   y2+       505.32       42.3
   a8++      535.84       24.2
   y7++      539.29        5.3
   b8++      549.82       36.6
   b3+       559.35        8.2
   a9++      600.18        8.5
   b9++      614.31       39.3
   b4+       630.50        9.3
   y3+       634.43       42.3
   b10++     663.75       32.5
   b11++     728.38       52.3
   y4+       733.55       27.0
   b12++     792.85       32.7
   y12++     801.52        5.8
   a6+       856.85        5.7
   y5+       862.51       18.2
   b7+       999.47       19.5
   b8+      1098.45       28.6
   b9+      1227.50       26.1
   b10+     1326.64       14.4
```

```
169. MORG-339_phosphopeptide_TMT_20%ACN_032017_OT.3770.3770.3 (sLTNSHLEkR)

   Ion         m/z   Intensity
   y2++      266.83       30.9
   b1+       397.19       18.7
   y5++      456.36       18.6
   y6++      499.89       24.3
   b2+       510.75       14.0
   a7++      517.29       17.4
   b7++      531.59       18.9
   y2+       532.41       29.4
   y7++      557.22       43.1
   b8++      596.46       25.7
   y8++      607.53       35.2
   b3+       611.00       15.4
   y3+       661.88       48.3
   y9++      664.09       46.5
   b4+       725.71       46.5
   b9++      774.78       29.8
   a5+       784.72       27.9
   b8+      1191.64       14.6
```

```
170. MORG-339_phosphopeptide_TMT_20%ACN_032017_OT.3204.3204.4 (rVsNRGLAGTTIR)

   Ion         m/z   Intensity
   y1+       175.12       10.8
   b1++      193.41        8.5
   y3++      195.26        8.0
   y4++      245.61        9.5
   a4+++     247.03        9.8
   b4+++     256.39       17.4
   y2+       288.24       12.1
   a5+++     298.90        7.7
   b5+++     308.27       21.4
   b6+++     327.22       19.0
   a7+++     355.70       53.0
   b7+++     365.03       38.9
   a8+++     379.39       12.5
   b4++      384.07       10.3
   b8+++     388.72       51.5
   y3+       389.38       30.1
   y8++      395.11       33.2
   a9+++     398.22       15.3
   b9+++     407.67       45.2
   y11+++    409.22       17.8
   b10+++    441.21       15.0
   y12+++    442.48       12.6
   a2+       456.91       10.2
   y9++      472.98       12.9
   a6++      476.17       13.4
   b2+       485.23        9.6
   y4+       490.25       29.3
   a12+++    503.55        7.6
   b12+++    512.36        7.2
   y10++     529.38       25.8
   a7++      532.99       22.4
   y5+       547.11       72.8
   a8++      568.39        7.0
   b8++      582.35       39.5
   b9++      611.03       30.1
   y6+       618.38       29.5
   b3+       652.53       12.2
```

```
171. MORG-339_phosphopeptide_TMT_20%ACN_032017_OT.15229.15229.3 (sAVSsLRAFQTLAR)

   Ion         m/z   Intensity
   b1+       397.28       21.0
   y8++      481.92       35.6
   y9++      538.53      102.2
   b7++      545.82       32.3
   b3+       567.48      138.1
   b8++      581.45       22.1
   y5+       588.27       35.8
   y10++     622.00       15.9
   b9++      654.83       16.2
   y11++     665.36       28.7
   y12++     715.21       16.9
   b10++     718.80       10.8
   y6+       735.42       19.2
   y13++     750.53       30.9
   y7+       806.51       34.0
   b5+       821.32       13.0
   b13++     861.40       18.7
```

```
172. MORG-339_phosphopeptide_TMT_10%ACN_032017_OT.8897.8897.4 (vDHGAEIITQSPsRsSVAsPR)

   Ion         m/z   Intensity
   y2+       272.20       31.0
   b3++      291.12       19.6
   b4++      319.75       25.9
   b8+++     355.32       31.1
   a6++      405.57       13.6
   b6++      419.80       62.4
   y3+       439.19       19.2
   b2+       444.32       23.2
   b11+++    461.40       15.5
   a7++      462.25       23.8
   b7++      476.46       63.9
   y8++      510.27       23.1
   a8++      518.81       16.9
   b8++      533.03       41.6
   y13+++    533.94       16.6
   a3+       553.43       29.1
   b3+       581.37       32.3
   b9++      583.59       23.1
   y9++      593.40       22.7
   b14+++    600.89       29.6
   y5+       609.35       19.4
   a10++     633.57       30.5
   b4+       637.92       80.2
   y10++     642.39      185.2
   b10++     647.31       52.6
   b15+++    656.97       28.3
   a11++     676.87       18.0
   y11++     685.66       16.9
   y6+       696.28       14.4
   b5+       709.49       30.8
   b17+++    718.88       20.7
   a18+++    733.00       45.2
   y19+++    740.68       16.1
   b18+++    742.69       19.0
   y13++     800.59       42.5
   a13++     808.88       19.1
   b6+       838.47       19.7
   y14++     857.12       53.8
   y7+       863.32       16.2
   b14++     901.26       19.6
   y15++     913.68       30.3
   b7+       951.60       17.7
   a15++     970.58       22.0
   y16++     978.28       26.0
   a16++    1013.53       14.1
   y18++    1041.83       13.6
   b8+      1064.59       16.9
   b17++    1077.90       13.7
```

```
173. MORG-339_phosphopeptide_TMT_20%ACN_032017_OT.9992.9992.3 (qVDTEEAGmVTAATASNVkAsPk)

   Ion         m/z   Intensity
   y1+       376.37       19.5
   y2+       473.34       83.3
   b8++      530.52       17.8
   b3+       572.36       48.3
   b9++      603.88       21.1
   y3+       640.52       21.3
   y7++      641.77       19.7
   b10++     653.50       18.0
   b4+       673.47       22.3
   y8++      685.16       18.2
   y4+       711.50       17.8
   b5+       802.46       28.7
   y13++     892.87       18.1
   b6+       931.52       39.5
   y14++     942.42       19.3
   a18++     996.80       21.7
   b7+      1002.54       37.2
   b18++    1010.58       42.9
   y15++    1015.48       24.4
   b8+      1059.51       23.8
   y5+      1068.65       22.4
   y17++    1079.85       25.6
   y18++    1143.94       18.0
   a9+      1178.82       29.8
   b19++    1189.99       54.3
   b9+      1206.57       26.4
   a20++    1211.42       31.1
   b20++    1225.36       48.0
   y7+      1281.63       18.4
   b10+     1305.69       24.9
   b21++    1308.88       34.9
   y21++    1317.07       34.1
   a22++    1342.78       21.0
   y22++    1366.23       23.6
   y8+      1368.80       23.1
```

```
174. MORG-339_phosphopeptide_TMT_14%ACN_032017_OT.2315.2315.3 (sGkNSQEDsEDSEEk)

   Ion         m/z   Intensity
   y1+       376.27       15.6
   y2+       505.33       19.1
   a6++      517.19       16.5
   b6++      531.23       13.9
   b7++      595.48       28.5
   y3+       634.44       11.6
   b8++      653.05       52.8
   y9++      689.18       12.9
   a3+       703.49       11.1
   y4+       721.47       32.8
   a9++      722.48       21.5
   b3+       731.58       15.7
   y10++     753.27       24.8
   a10++     786.40       38.4
   y11++     796.51       23.6
   b10++     800.85       38.1
   y5+       836.50       16.3
   b4+       845.32       10.5
   b11++     858.59       35.4
   y6+       965.51       18.7
   b13++     966.44       13.8
   a14++    1016.48        9.4
   b14++    1031.28       23.0
   a6+      1032.14       10.2
   b6+      1060.77       11.6
   y7+      1132.57       24.6
   y8+      1247.47       11.4
```

```
175. MORG-339_phosphopeptide_TMT_22%ACN_032017_OT.5262.5262.3 (sPAkTITPQNAPR)

   Ion         m/z   Intensity
   a1+       369.26       19.8
   b1+       397.19       70.0
   b4++      461.86       45.9
   a6++      555.13       24.6
   b3+       565.39       22.3
   b6++      569.09       31.6
   y5+       585.17       31.8
   a7++      604.87       31.1
   b7++      619.23       54.3
   a8++      654.25       20.1
   y10++     678.09       49.9
   y6+       682.38       51.0
   y11++     713.51       25.8
   a9++      717.55       19.9
   b9++      732.04       24.7
   y12++     762.13      186.8
   y7+       783.44       51.0
   b10++     788.82       27.0
   b11++     824.67       53.5
   b12++     873.19       29.9
   y8+       896.47       34.7
   y9+       997.63       39.2
```

```
176. MORG-339_phosphopeptide_TMT_16%ACN_032017_OT.6651.6651.3 (eVQsPEQVkSEk)

   Ion         m/z   Intensity
   y1+       376.35       17.3
   b2+       458.23       20.0
   y4++      475.33       22.4
   a6++      475.99       24.2
   y2+       505.52       14.9
   y5++      525.03       26.5
   b3+       586.38       25.4
   y6++      589.05       40.0
   a8++      589.90       16.4
   y3+       592.41       26.0
   y7++      652.97       41.6
   y8++      702.12      151.7
   a9++      768.61       15.2
   b9++      782.54       30.8
   y9++      785.61       18.3
   a10++     812.40       17.9
   b5+       850.27       21.5
   b11++     890.33       23.0
   y11++     899.26       30.5
   y4+       949.81       21.9
   a6+       951.79       46.3
   b6+       979.52       26.6
   b7+      1107.49       19.8
   b8+      1206.62       16.0
```

```
177. MORG-339_phosphopeptide_TMT_10%ACN_032017_OT.5353.5353.4 (aEsPETSAVESTQSTPQkGR)

   Ion         m/z   Intensity
   y5++      407.93      149.2
   b5++      412.32       35.0
   a10+++    428.47       39.7
   y6++      458.46      107.9
   b6++      462.77       72.3
   y11+++    483.74       45.9
   y7++      502.00       95.3
   b7++      506.51       28.1
   y12+++    516.52       60.6
   a13+++    533.81       40.6
   y13+++    539.97       39.5
   b8++      542.00       89.6
   b13+++    542.87       25.1
   y8++      566.05       90.9
   a3+       569.16       51.1
   a9++      577.57       27.4
   b9++      591.41      150.6
   b3+       597.03       44.4
   y15+++    602.96       85.4
   y9++      616.58       97.7
   a16+++    629.12       79.2
   b16+++    637.90       36.4
   a10++     642.11      108.1
   y16+++    646.09       44.7
   b10++     656.05       30.0
   y17+++    677.88       23.4
   b11++     699.55       50.9
   y11++     724.64      172.2
   b12++     750.06       42.8
   y12++     774.15      183.9
   a13++     799.87       59.5
   y13++     809.64       79.7
   y5+       814.10       68.8
   b19+++    819.07       41.1
   b5+       823.36       63.9
   y14++     853.13       79.5
   b14++     857.62      115.9
   y15++     903.75       58.3
   b15++     907.56       51.0
   b6+       924.39       30.2
   b7+      1011.48       26.6
   b8+      1082.52       67.3
   b9+      1181.59       37.5
   b10+     1310.64       32.1
```

```
178. MORG-339_phosphopeptide_TMT_20%ACN_032017_OT.10422.10422.2 (rIDIsPSALR)

   Ion         m/z   Intensity
   y3+       359.29        9.0
   b1+       386.26        9.4
   b2+       499.17        9.7
   y5+       543.40       34.5
   b3+       614.47       34.9
   a4+       699.37       25.7
   y6+       710.06       38.9
   b4+       727.58       14.7
   y7+       823.48       26.1
   b5+       894.53        8.2
   y8+       938.34       10.8
   y9+      1051.54       12.7
   b8+      1149.64        9.9
   b9+      1262.64       12.1
```

```
179. MORG-339_phosphopeptide_TMT_22%ACN_032017_OT.4582.4582.3 (kSQAWAEGGsPR)

   Ion         m/z   Intensity
   y2+       272.19       57.0
   a3++      273.17       22.5
   a5++      401.92       14.7
   a6++      437.19       19.4
   a7++      501.77       14.2
   a8++      530.55       22.7
   y5+       553.21       26.7
   a9++      558.87       16.6
   y10++     569.65       26.4
   a10++     642.06       21.1
   b4+       644.64       17.6
   b10++     655.85       23.5
   a11++     691.31       16.9
   b11++     704.91       17.0
   y7+       753.12       16.3
   b5+       830.24       13.9
```

```
180. MORG-339_phosphopeptide_TMT_10%ACN_032017_OT.9166.9166.3 (aPsPAPAVsPk)

   Ion         m/z   Intensity
   y2++      237.18        9.3
   b3++      282.83       11.8
   a4++      317.34        6.5
   y3++      321.00       19.4
   a5++      353.22        6.9
   b5++      367.28       17.7
   a2+       370.24       30.7
   y1+       376.31       11.0
   y5++      405.25        8.9
   b6++      415.36       17.2
   b7++      451.45       24.6
   y6++      454.32      100.5
   y2+       473.32       26.6
   a8++      486.58       22.4
   y7++      489.48        6.4
   b8++      500.74       45.2
   a3+       537.40       24.4
   y8++      538.29       29.8
   b3+       565.30        9.0
   a9++      570.53       10.6
   b9++      584.34       10.2
   a4+       634.24        5.3
   y3+       640.33       41.8
   b4+       662.35        6.8
   y10++     670.46        6.2
   b5+       733.38       76.9
   y4+       739.37       16.1
   y5+       810.49        6.0
   b7+       901.43       28.1
   y6+       907.50       22.3
   b8+      1000.51       14.6
```

```
181. MORG-339_phosphopeptide_TMT_8%ACN_032017_OT.7121.7121.2 (eIssSPTSk)

   Ion         m/z   Intensity
   a1+       331.31        9.6
   b1+       359.26       19.0
   y1+       376.41       23.5
   a2+       444.55       10.1
   y2+       463.27       20.4
   b2+       472.37       29.1
   y3+       564.41       11.0
   y4+       661.42       64.4
   y5+       748.93       29.4
   a5+       865.19        8.6
   b5+       893.39       15.3
   y6+       915.51       26.0
   a7+      1063.43        9.0
   y7+      1082.42       21.6
   b7+      1091.42        9.1
   y8+      1195.56       18.8
```

```
182. MORG-339_phosphopeptide_TMT_8%ACN_032017_OT.2616.2616.3 (lETEDTSGsPSRASR)

   Ion         m/z   Intensity
   y3+       333.26       28.1
   a4++      337.29       67.1
   y8++      449.36       19.2
   b2+       472.33       20.8
   y9++      492.86       24.4
   a8++      517.50       15.4
   y10++     543.39       54.3
   a9++      601.52       72.5
   b9++      615.02       21.0
   b4+       702.47       28.7
   y13++     715.79       16.5
   a12++     771.83       15.2
   b12++     785.50       53.7
   a13++     806.74       21.1
   b5+       817.43       76.4
   b13++     820.73       29.1
   b6+       918.42       18.6
   b7+      1005.54       21.5
   b8+      1062.52       20.7
```

```
183. MORG-339_phosphopeptide_TMT_20%ACN_032017_OT.7119.7119.3 (iERWHQLQNENHVsSD)

   Ion         m/z   Intensity
   b5++      476.39       17.2
   y4+       487.14       17.1
   b6++      540.59       21.1
   a7++      583.16       15.2
   b7++      596.67       54.6
   a3+       600.37       24.2
   y5+       624.45       22.6
   b8++      661.12       34.4
   b9++      718.27       55.7
   y6+       738.20       55.0
   y12++     743.90       33.4
   b10++     782.63       48.5
   b11++     839.73       55.9
   y7+       867.12       16.5
   a12++     894.38       27.4
   b12++     908.30       64.8
   a13++     943.72       17.7
   b13++     957.83       52.3
   y15++     979.55       45.5
   y8+       981.47       31.8
```

```
184. MORG-339_phosphopeptide_TMT_12%ACN_032017_OT.4071.4071.3 (sRtASGSSVTSLEGTR)

   Ion         m/z   Intensity
   b2++      237.43       15.0
   y5++      287.76       12.5
   b3++      327.69       27.7
   y6++      332.12       13.5
   y3+       333.22       23.1
   a4++      348.75       13.0
   b4++      363.33       13.0
   b5++      406.96       21.8
   a6++      421.40       17.9
   b6++      435.59       14.1
   y4+       462.29       18.8
   a7++      464.35       16.3
   b2+       473.34       16.2
   b7++      478.99       19.6
   b8++      522.64       19.6
   y11++     547.04       16.3
   b9++      571.98       25.6
   y5+       575.41       18.1
   y12++     591.07       53.1
   a10++     608.06       23.0
   b10++     622.48       21.8
   a3+       625.91       29.0
   y6+       662.38       27.6
   b11++     665.82       15.9
   y7+       763.48       50.8
   a13++     772.98       13.0
   b13++     787.09       39.4
   b14++     815.48       13.6
   y8+       862.55       29.3
   b15++     865.52       13.0
   y9+       949.50       33.8
   y10+     1036.63       21.1
   y11+     1093.59       41.5
   y12+     1180.68       12.5
   y13+     1251.69       18.1
```

```
185. MORG-339_phosphopeptide_TMT_12%ACN_032017_OT.2812.2812.3 (sRstTPAPk)

   Ion         m/z   Intensity
   y2++      237.22       20.0
   y3++      272.37       17.3
   y4++      321.32       48.7
   y5++      371.86       16.7
   y1+       376.37       14.9
   a4++      397.19       10.0
   a5++      447.67       15.1
   b5++      461.59       31.4
   y6++      462.46       30.5
   y2+       473.30       72.9
   a6++      496.13       19.6
   b6++      509.74       92.7
   a7++      531.66       13.8
   y3+       544.41       18.3
   y7++      545.92       52.7
   a8++      580.48       10.8
   b8++      594.17       10.6
   a3+       612.24        9.7
   b3+       640.53       16.3
   y4+       641.48       83.4
   y5+       742.51       25.2
   b4+       821.42        9.8
   b5+       922.36       31.1
   y6+       923.41       18.8
```

```
186. MORG-339_phosphopeptide_TMT_4%ACN_032017_OT.3011.3011.3 (sRsIDRGLDR)

   Ion         m/z   Intensity
   y1+       175.15       37.1
   y5++      308.76       56.1
   y6++      366.24       33.6
   y7++      422.96       36.5
   a5++      460.35       33.1
   y8++      506.21       28.2
   b2+       552.98       35.3
   y9++      584.42       25.7
   b8++      637.63       29.4
   b9++      695.46      100.4
   b5+       948.40       45.0
```

```
187. MORG-339_phosphopeptide_TMT_25%ACN_032017_OT.7629.7629.4 (rASsQDGLETLkk)

   Ion         m/z   Intensity
   b4++      356.35       18.0
   y2++      367.23       28.1
   y1+       376.41       26.3
   b5++      419.99       26.4
   y3++      423.84       33.1
   a2+       428.88       71.4
   b2+       457.48       27.7
   y4++      474.38       51.6
   b6++      477.88       97.3
   b11+++    489.58       20.1
   y9+++     497.77       19.9
   b7++      506.41       40.5
   a3+       516.54       18.8
   y5++      539.01       89.6
   b3+       543.99       29.9
   b8++      562.62       25.0
   y6++      595.69       45.5
   b12+++    609.03       27.5
   a9++      613.35       28.1
   y7++      624.12      148.1
   b9++      627.20       34.6
   a10++     663.83       20.7
   y8++      681.69       84.3
   y9++      745.72       88.8
   a5+       811.43       22.9
   y10++     829.17       22.0
```

```
188. MORG-339_phosphopeptide_TMT_4%ACN_032017_OT.7392.7392.4 (eGEEPTVYsDDEEPkDETAR)

   Ion         m/z   Intensity
   a5+++     248.21        6.0
   y3+       347.35       12.6
   b6++      436.86        7.8
   a10+++    463.34       21.0
   y6++      474.91        7.1
   y4+       476.40       11.2
   b7++      486.42       10.0
   y7++      523.53       42.0
   b3+       545.34       10.5
   b12+++    554.24        7.5
   b8++      567.75        6.9
   a13+++    588.05       19.2
   y5+       591.44        6.8
   y13+++    622.17       18.3
   b14+++    629.16        7.9
   y9++      652.69       21.8
   y14+++    655.06       15.5
   b4+       674.39       26.0
   y15+++    688.65       17.3
   a10++     694.86        7.9
   y16+++    721.11       16.7
   b15+++    749.07        6.6
   y11++     767.55       10.6
   b17+++    830.23       15.6
   b12++     830.95       17.9
   y12++     851.16       22.5
   a18+++    854.53       13.4
   b18+++    864.04       11.7
   b6+       872.44       12.8
   a13++     881.81       15.1
   b13++     895.52       18.3
   y13++     932.63       44.1
   b7+       971.60       22.4
   y14++     982.39       13.4
   y15++    1032.94        6.8
   b8+      1134.64       13.6
   y8+      1174.80        5.8
```

```
189. MORG-339_phosphopeptide_TMT_14%ACN_032017_OT.3841.3841.2 (eAsPSRLSVAR)

   Ion         m/z   Intensity
   b1+       359.27       16.7
   b2+       430.23       15.9
   y4+       432.28       12.7
   y5+       545.37       22.2
   b4+       694.44       24.8
   y6+       701.37       46.8
   y8+       885.55       39.7
   b6+       937.51       21.9
   b7+      1050.58       15.7
   y9+      1052.61       19.7
   y10+     1123.58       26.2
   b10+     1307.70       15.6
```

```
190. MORG-339_phosphopeptide_TMT_16%ACN_032017_OT.5742.5742.3 (rySPsPPPk)

   Ion         m/z   Intensity
   y3++      285.84       28.3
   y4++      334.31       62.1
   b3++      358.45       19.9
   y1+       376.46       12.6
   a4++      393.34       17.4
   y5++      417.28       24.7
   y6++      466.34       34.8
   y2+       473.44       36.8
   a5++      476.47       10.3
   b5++      490.52       37.9
   y7++      509.64       12.7
   a6++      525.44       21.9
   y3+       570.39       29.6
   a7++      573.29        9.8
   b7++      587.49       11.4
   a8++      622.73       10.6
   y8++      631.27       13.0
   b8++      636.14       10.7
   y4+       667.47       74.1
   b3+       716.31       12.2
   b5+       980.42       14.8
```

```
191. MORG-339_phosphopeptide_TMT_20%ACN_032017_OT.10640.10640.3 (sGDALTTVVVkQER)

   Ion         m/z   Intensity
   b1+       397.26       18.5
   y3+       432.21       14.6
   y6++      494.51       15.3
   y7++      544.01       13.9
   b3+       569.22       37.6
   y8++      594.62       18.6
   b4+       640.27       23.3
   y9++      645.31       21.9
   y10++     701.87       27.4
   y11++     737.27       80.2
   b5+       753.46       16.0
   y4+       789.48       18.7
   y12++     794.70       24.1
   b11++     805.39       17.8
   y13++     823.32       62.9
   y5+       888.56       22.6
   a13++     919.65       12.2
   b13++     934.26       55.7
   y6+       987.68       21.8
   y7+      1086.79       12.2
```

```
192. MORG-339_phosphopeptide_TMT_10%ACN_032017_OT.7569.7569.3 (sRsGEGEVSGLLR)

   Ion         m/z   Intensity
   b2++      237.20        9.7
   a3++      306.72        8.6
   b1+       317.24       12.4
   b3++      320.68       22.4
   a4++      335.34        7.2
   b4++      349.38        7.8
   a5++      399.65       15.8
   b5++      413.74       36.6
   y8++      416.06        9.8
   b6++      442.22       20.0
   y4+       458.04       22.2
   b2+       473.49        7.9
   y9++      480.69        7.2
   a7++      492.61       15.4
   b7++      506.82       42.4
   y10++     509.05        7.5
   a8++      542.60       27.2
   y5+       545.37       26.3
   b9++      599.50        8.3
   b10++     628.36       13.7
   y6+       644.43       27.0
   a11++     670.93       14.9
   b11++     684.87       24.5
   b12++     741.50       21.6
   y7+       773.54       19.1
   y8+       830.48       35.5
   y9+       959.50        8.1
   y10+     1016.59       20.5
   y11+     1183.50        6.6
```

```
193. MORG-339_phosphopeptide_TMT_10%ACN_032017_OT.6056.6056.2 (sRsPLAIR)

   Ion         m/z   Intensity
   y2+       288.33       16.1
   y3+       359.34       15.5
   b2+       553.32       14.4
   y5+       569.40       37.0
   b3+       720.01       15.1
   y6+       736.47       16.5
   y7+       892.57       17.2
   b5+       930.50       16.7
   b6+      1001.50       27.3
   b7+      1114.54       30.9
```

```
194. MORG-339_phosphopeptide_TMT_6%ACN_032017_OT.6036.6036.2 (tEsIEkEGE)

   Ion         m/z   Intensity
   a1+       303.09        8.5
   b1+       331.23       15.7
   a2+       432.33       11.6
   b2+       460.32       22.5
   y4+       691.42       34.7
   b4+       740.58       58.1
   y5+       820.50       35.8
   b5+       869.41       25.0
   y6+       933.53       40.6
   y7+      1100.53       23.1
```

```
195. MORG-339_phosphopeptide_TMT_4%ACN_032017_OT.10011.10011.3 (iAcEEEFsDsDEEGEGGRk)

   Ion         m/z   Intensity
   y4++      323.98        8.5
   y1+       376.42        6.5
   y6++      416.77       10.8
   y7++      481.45        7.4
   a3+       546.00       15.9
   b3+       574.39        6.2
   y9++      603.64        7.0
   b8++      638.52        9.2
   y10++     686.74       12.0
   b9++      695.89        5.6
   b4+       703.50        6.4
   y11++     744.44        8.6
   b10++     779.32       11.0
   a5+       804.88        8.3
   a11++     823.09       13.5
   y12++     828.11        8.5
   y6+       832.47        6.8
   a12++     887.26        6.2
   y13++     901.35       13.8
   a13++     951.38        7.3
   y7+       961.47       10.6
   a14++     980.25        5.6
   y15++    1030.79        5.8
   b7+      1108.57       10.1
   y17++    1175.37        7.0
   b9+      1390.52        7.8
   y13+     1801.73        5.7
```

```
196. MORG-339_phosphopeptide_TMT_25%ACN_032017_OT.8520.8520.3 (iTRkPASVSPTTPtSPTEGEAS)

   Ion         m/z   Intensity
   b8++      656.43       40.1
   y7+       690.40       51.3
   b9++      700.16       69.6
   a10++     734.43       24.7
   b10++     748.64       26.5
   b11++     799.33       55.9
   a12++     835.91       44.4
   b12++     849.47       85.5
   y17++     850.08       86.9
   a13++     884.41       39.6
   b13++     897.86       92.5
   y18++     898.86       40.7
   b4+       957.38       25.0
   a14++     974.23       26.4
   b15++    1032.45       25.5
   y10+     1055.43       48.2
   b17++    1131.50       27.2
   y11+     1156.62       26.1
   y13+     1354.53       27.0
```

```
197. MORG-339_phosphopeptide_TMT_30%ACN_032017_OT.11339.11339.3 (kVPLPGPGsPEVk)

   Ion         m/z   Intensity
   y1+       376.34       25.2
   y5++      435.06       26.1
   y2+       475.46       29.4
   a3+       526.49       25.1
   y8++      540.10       48.4
   y9++      589.15       43.4
   a10++     606.31       18.0
   a4+       639.51       16.8
   y10++     645.36       17.0
   b4+       667.16       35.1
   a11++     670.72       43.0
   y4+       701.49       67.8
   b12++     734.75       18.2
   a5+       736.58       15.8
   y12++     743.25       21.9
   y5+       868.60       15.9
   a7+       890.71       16.2
   y8+      1079.67       26.7
   y9+      1176.58       21.5
```

```
198. MORG-339_phosphopeptide_TMT_10%ACN_032017_OT.6470.6470.3 (ekEIsDDEAEEEk)

   Ion         m/z   Intensity
   a2++      344.26       14.6
   y1+       376.35       16.4
   y4++      381.90       13.5
   b3++      423.58       14.5
   y2+       505.38       15.5
   b6++      620.90       28.0
   y3+       634.41       14.6
   b7++      678.57       30.1
   y9++      680.28       16.1
   a8++      728.71       18.0
   y10++     736.98       24.4
   b8++      742.54       46.3
   y4+       763.49       42.2
   a9++      764.50       34.1
   b9++      778.60       16.0
   y5+       834.59       20.3
   b10++     843.32       16.3
   b11++     907.64       14.9
   y6+       963.49       24.9
   b12++     971.72       17.2
   y7+      1078.50       31.6
   y8+      1193.50       15.9
```

```
199. MORG-339_phosphopeptide_TMT_12%ACN_032017_OT.7629.7629.3 (eEQsPVkAEVAEk)

   Ion         m/z   Intensity
   b1+       359.28       16.5
   y1+       376.35       19.2
   b2+       488.35       23.0
   y2+       505.39       16.8
   y3+       576.31       23.0
   b3+       616.66       27.7
   y7++      617.35       30.5
   a7++      654.60       12.3
   b7++      668.97       44.0
   y4+       675.53       29.7
   b8++      704.75       48.4
   y9++      715.24      115.7
   a9++      754.70       13.1
   a4+       755.64       12.8
   b9++      769.20       28.4
   b4+       783.27       10.3
   y5+       804.59       32.2
   b10++     818.37       24.6
   a11++     839.54       13.1
   b11++     854.06       36.6
   y6+       875.49       33.0
   b12++     918.68       27.1
   y12++     927.15       10.7
   a6+       951.50       11.6
   b6+       979.34       30.5
```

```
200. MORG-339_phosphopeptide_TMT_12%ACN_032017_OT.4932.4932.2 (dAsPSSSSAVASSk)

   Ion         m/z   Intensity
   b1+       345.16       10.8
   y1+       376.46        6.8
   b2+       416.33        7.3
   y2+       463.33        6.1
   y3+       550.40       11.5
   b3+       583.11        5.5
   y4+       621.46       13.6
   y5+       720.40       11.8
   y6+       791.63        8.9
   y7+       878.29       17.2
   y8+       965.56       12.9
   b8+      1028.68        6.9
   y9+      1052.64       10.5
   b9+      1099.54        5.9
   b10+     1198.55        9.8
   y11+     1236.63       40.4
   b11+     1269.62       10.0
   a12+     1328.75        6.4
   y13+     1474.71        9.2
```

```
201. MORG-339_phosphopeptide_TMT_18%ACN_032017_OT.10433.10433.4 (tEGYAAFQEDssGDEAESPSkVkR)

   Ion         m/z   Intensity
   b5++      376.27       25.3
   y6+++     391.64       40.3
   b6++      411.89       27.5
   y7+++     420.75       68.5
   y8+++     463.95       30.8
   b7++      485.44       29.9
   y9+++     487.53       56.7
   y10+++    530.51       83.0
   y6++      586.94       23.5
   y12+++    587.93       85.7
   b9++      613.93       35.6
   y7++      630.62       52.3
   y3+       631.50       23.7
   y13+++    643.49       42.8
   b4+       680.47       43.4
   y8++      695.00       42.0
   y14+++    699.26       49.8
   y9++      730.49       43.5
   y15+++    737.57       27.6
   b5+       751.44       65.9
   a19+++    778.62       22.7
   y16+++    780.62       24.4
   a6+       794.35       24.9
   y10++     795.21       53.3
   a20+++    807.59       51.2
   b6+       822.44      104.6
   y17+++    823.44       70.0
   a12++     824.59       45.3
   b12++     837.88       30.2
   y18+++    872.31       22.7
   y12++     881.34       36.8
   y19+++    896.15       24.1
   a7+       941.59       39.1
   y13++     964.74       22.9
   b7+       969.49       70.6
   y14++    1047.64       27.7
   a8+      1069.78       26.6
   b23+++   1088.21       43.1
   b8+      1097.55       50.4
   y15++    1105.81       34.2
   a18++    1118.35       22.8
   a20++    1209.99       26.6
   b9+      1226.59       52.1
   b10+     1341.63       50.3
   y19++    1342.63       39.2
   a11+     1480.72       23.4
   b14+     1847.71       27.0
```

```
202. MORG-339_phosphopeptide_TMT_25%ACN_032017_OT.6859.6859.3 (rWQsPVTk)

   Ion         m/z   Intensity
   b2++      286.76       12.5
   y4++      337.21       50.7
   b3++      350.88       26.2
   y1+       376.43       30.2
   b1+       385.87       83.2
   b4++      434.16       31.3
   a5++      468.65       14.6
   y2+       477.39       59.0
   b5++      482.47       35.5
   y6++      484.93       13.0
   a6++      518.47       18.0
   b6++      532.39       37.9
   a7++      569.09       14.8
   b2+       572.26       11.0
   y3+       576.49       16.6
   y7++      577.76       11.9
   a3+       672.45       19.4
   y4+       673.49      118.0
   y5+       840.40       12.9
   b4+       867.53       26.2
```

```
203. MORG-339_phosphopeptide_TMT_14%ACN_032017_OT.8112.8112.4 (aQPSDNAPEkGNksPsPPPDGSPAATPEIR)

   Ion         m/z   Intensity
   a6++      407.34       21.2
   b2+       429.40       23.5
   y4+       514.44       23.3
   y12++     606.20       23.6
   y5+       615.45       23.6
   y13++     654.43       36.7
   b13+++    675.61       25.8
   y14++     703.23      128.7
   b5+       728.50       49.0
   b14+++    731.34       22.1
   y18+++    732.01       24.3
   b10++     748.91       34.4
   a15+++    754.61      138.9
   b11++     777.26       21.9
   b16+++    819.60       39.8
   y16++     835.00       53.1
   b17+++    851.82       21.1
   y8+       854.58       43.8
   b18+++    883.79       22.0
   b7+       913.58       31.4
   y9+       941.79      333.8
   a20+++    945.15       21.8
   y22+++    950.73       26.6
   b20+++    955.09       44.9
   y10+      998.56       84.9
   a13++     999.56       65.9
   y24+++   1007.03       20.8
   b13++    1012.65       38.8
   y25+++   1045.32       47.4
   a24+++   1049.83       27.4
   y26+++   1083.06       24.7
   y18++    1096.91       26.8
   a9+      1111.43       23.7
   y27+++   1112.46       25.0
   a15++    1130.96       79.9
   b9+      1139.70       40.9
   b15++    1144.91       23.5
   a28+++   1181.81       35.2
   b28+++   1191.95       42.1
   y12+     1210.63       37.6
   b16++    1228.46       25.4
   b29+++   1229.40       39.0
   y13+     1307.70       31.2
   y14+     1404.76      106.2
   y15+     1571.82       34.2
   y16+     1668.85       40.7
```

```
204. MORG-339_phosphopeptide_TMT_12%ACN_032017_OT.4473.4473.3 (rkVmDsDEDDADY)

   Ion         m/z   Intensity
   a3++      407.82       10.2
   b4++      495.59       11.3
   b5++      553.00       38.3
   b6++      636.09       16.0
   a7++      679.57       23.7
   y11++     685.75       35.8
   b7++      693.85       50.0
   y6+       727.21        9.7
   b8++      758.10       10.2
   a9++      802.25        9.6
   a3+       814.36       11.9
   b9++      815.87       28.2
   y7+       841.90        9.8
   b10++     873.65       28.4
   b11++     908.82       27.4
   b12++     966.60       38.2
```

```
205. MORG-339_phosphopeptide_TMT_25%ACN_032017_OT.12232.12232.3 (aALsPARLGsPIRSPk)

   Ion         m/z   Intensity
   y6++      463.78       31.2
   y2+       473.43       20.4
   b3+       485.52       18.5
   a9++      559.41       18.2
   y3+       560.45       18.1
   y9++      632.20       23.6
   a10++     643.76       22.4
   y10++     710.55       28.0
   y4+       716.32       24.8
   a5+       721.69       39.5
   b12++     762.04       23.7
   y12++     794.60       59.8
   a13++     826.83       18.8
   y5+       829.13       30.3
   b13++     840.59       30.4
   a14++     869.93       19.6
   y13++     877.90       26.8
   b14++     883.71       22.9
   y15++     969.97       19.6
```

```
206. MORG-339_phosphopeptide_TMT_16%ACN_032017_OT.12002.12002.2 (eDsGTFSLGk)

   Ion         m/z   Intensity
   b1+       359.32        8.2
   y1+       376.32       12.6
   y2+       433.38       18.1
   b2+       474.26       27.3
   y3+       546.46       13.9
   y4+       633.43       18.1
   b4+       698.15        9.4
   a5+       771.24       36.2
   y5+       780.83       57.9
   y6+       881.59       11.5
   y7+       938.57       36.5
   b6+       946.42       10.8
   y8+      1105.56       23.7
   b9+      1203.49       11.0
   y9+      1220.60       13.4
```

```
207. MORG-339_phosphopeptide_TMT_10%ACN_032017_OT.6851.6851.3 (eGEEPtVYSDDEEPkDETARk)

   Ion         m/z   Intensity
   b3+       545.37        9.6
   b4+       674.31       14.5
   y8++      701.82       17.3
   y9++      766.87       12.6
   y10++     831.14       24.9
   y5+       833.60       11.7
   y11++     888.76       25.2
   y12++     945.75       13.6
   a7+      1023.08       12.3
   b7+      1050.96        9.1
   b15++    1122.35        8.5
   y16++    1210.93       18.0
   b17++    1244.22        8.4
   y17++    1259.41        9.6
   b10+     1416.46        8.4
   y9+      1531.75       10.3
```

```
208. MORG-339_phosphopeptide_TMT_14%ACN_032017_OT.14162.14162.3 (eVWAsDsGHEEDcLSPEPPLRPk)

   Ion         m/z   Intensity
   y6++      468.94       28.8
   y2+       473.37       10.2
   y8++      582.06       23.0
   b7++      582.86       12.4
   y9++      625.62       14.2
   b4+       715.49       12.2
   y11++     761.96       26.7
   y12++     819.49       18.2
   y13++     884.33       15.8
   y6+       936.61       23.2
   a14++     988.56       14.6
   b6+       997.48       29.5
   b14++    1002.47       12.6
   y15++    1016.60       15.2
   a15++    1032.77       14.9
   y16++    1045.84       25.3
   y17++    1129.30       35.8
   b17++    1159.83       18.9
   y8+      1162.86       17.6
   y18++    1187.00       11.6
   b8+      1221.18       16.3
   y9+      1249.97       18.3
   b21++    1391.26       15.5
   b10+     1487.43       10.9
   b12+     1731.65       18.4
```

```
209. MORG-339_phosphopeptide_TMT_10%ACN_032017_OT.12910.12910.3 (sAsPDDDLGSSNWEAADLGNEERk)

   Ion         m/z   Intensity
   y3++      331.39       16.8
   y1+       376.37       16.4
   b2+       388.31       16.5
   y7++      537.99       72.5
   y8++      595.54       19.1
   y9++      631.08       18.7
   y3+       661.51       16.2
   y10++     666.57       38.0
   b11++     671.40       16.2
   y11++     731.05       19.4
   b5+       767.34       37.1
   y12++     824.09       17.4
   y13++     880.93       21.8
   b6+       882.41       36.5
   y5+       904.51       18.4
   y14++     924.68       24.5
   y6+       961.50       23.6
   y15++     968.14       24.7
   a7+       969.24       25.5
   y16++     996.66      106.1
   b7+       997.57       57.7
   a17++    1000.31       18.2
   b17++    1014.57       27.6
   y17++    1053.23       80.0
   a18++    1057.26       21.2
   y7+      1074.66       34.0
   y18++    1110.68       77.6
   b20++    1156.07       18.0
   y19++    1167.90       48.5
   y8+      1189.75       22.8
   y21++    1274.29      118.7
   y10+     1331.67       23.7
   b12+     1455.34       19.1
```

```
210. MORG-339_phosphopeptide_TMT_10%ACN_032017_OT.11176.11176.2 (sLsySPVER)

   Ion         m/z   Intensity
   y2+       304.27       16.1
   b1+       317.27       14.1
   b2+       430.42       20.5
   y4+       500.29       67.3
   a3+       569.37       15.0
   b3+       597.37       14.3
   y6+       830.36       46.0
   a5+       899.49       13.4
   b5+       927.55       15.5
   y7+       997.44       13.2
   y8+      1110.50       13.3
   b7+      1123.48       23.9
   b8+      1252.60       24.0
```

```
211. MORG-339_phosphopeptide_TMT_10%ACN_032017_OT.2354.2354.2 (aAsPSPQSVR)

   Ion         m/z   Intensity
   b1+       301.31       12.8
   y3+       361.25       16.5
   b2+       372.34       14.4
   y4+       489.30       11.8
   b3+       539.25       20.0
   y5+       586.40       47.8
   a4+       608.79       25.0
   b4+       635.95       30.5
   a5+       695.25       12.0
   b5+       723.45       21.9
   y7+       770.45      113.3
   a7+       920.49       14.8
   y8+       937.51       17.0
   b7+       948.41       15.6
   y9+      1008.51       15.3
   b9+      1134.65       12.3
```

```
212. MORG-339_phosphopeptide_TMT_22%ACN_032017_OT.9496.9496.3 (kLsGDLEAGAPk)

   Ion         m/z   Intensity
   y1+       376.27       16.9
   b2+       471.41       45.3
   y2+       473.37       55.8
   y3+       544.28       23.0
   b8++      562.54       16.9
   a9++      576.81       32.8
   b9++      590.85       20.7
   y4+       601.44       37.5
   a10++     612.02       16.1
   b10++     626.15       30.0
   y10++     627.05       29.0
   b3+       638.72       30.6
   a4+       667.79       11.5
   y5+       672.48       39.8
   b11++     674.63       12.6
   y6+       801.51       27.3
   y7+       914.58       40.7
   b6+       923.90       11.0
   y9+      1086.62       25.5
```

```
213. MORG-339_phosphopeptide_TMT_16%ACN_032017_OT.13495.13495.3 (asSPPVEVGEPREFVLQPAPQGR)

   Ion         m/z   Intensity
   a1+       273.38       17.0
   b1+       301.28       21.6
   a2+       440.39       25.9
   y4+       457.31      118.6
   a7++      475.37       16.4
   b3+       555.32       20.4
   y6+       625.38       34.7
   y13++     748.19      127.7
   b5+       749.17       53.1
   y7+       753.56       20.6
   b12++     758.14       16.2
   y14++     812.69       24.3
   b13++     823.25       22.3
   y15++     841.23       40.4
   b6+       848.49       19.4
   y8+       866.54       25.3
   y16++     890.76       72.0
   b15++     946.31       28.3
   y17++     955.30       28.6
   y9+       965.56       16.5
   b7+       977.48       32.7
   y18++    1004.32       16.7
   a17++    1052.79       20.0
   y19++    1053.53       31.9
   b17++    1066.76       30.8
   b8+      1076.52       20.3
   y20++    1101.93      302.4
   y10+     1112.62       26.9
   a19++    1136.41       40.9
   y21++    1145.35       27.9
   b19++    1150.95       30.7
   a20++    1184.92       15.5
   y11+     1241.85       24.6
   b10+     1262.61       52.9
   b21++    1263.61       44.4
   b22++    1292.15       19.5
   y13+     1494.89       27.0
```

```
214. MORG-339_phosphopeptide_TMT_6%ACN_032017_OT.3839.3839.4 (eRDkEVsDDEAEEk)

   Ion         m/z   Intensity
   b3++      315.83        6.1
   y3++      317.48        6.3
   b5+++     372.89       13.0
   y1+       376.31       17.8
   b6+++     406.09       12.3
   a4++      480.46       10.9
   b8+++     499.93       16.6
   y2+       505.53       31.7
   b9+++     538.41       33.8
   a5++      544.33        7.6
   b5++      558.46       11.4
   a10+++    571.98        9.8
   b10+++    581.49       31.3
   a6++      594.35       10.7
   a11+++    595.33       13.0
   a3+       602.60       11.2
   b11+++    604.90       19.1
   y3+       634.39       16.8
   b12+++    648.33       31.7
   y9++      666.21        6.1
   b13+++    691.19       21.9
   b7++      692.22       13.7
   y13+++    697.07       11.0
   y4+       705.42       21.0
   y10++     730.71       13.6
   a9++      792.71        7.5
   b9++      807.24        6.1
   y5+       834.52       16.4
   b10++     871.85        8.6
   b11++     907.15        9.3
   y11++     908.67       10.0
   y6+       949.60       12.7
   b12++     971.65        8.7
   a13++    1021.76        9.8
   y7+      1064.54       12.7
```

```
215. MORG-339_phosphopeptide_TMT_12%ACN_032017_OT.8337.8337.2 (dVYLsPR)

   Ion         m/z   Intensity
   y2+       272.16       60.5
   a1+       317.36        8.1
   b1+       345.30       13.2
   y3+       439.20       10.1
   b2+       444.28       12.3
   y4+       552.68       10.7
   b3+       607.58       18.0
   y5+       715.48        8.4
   b4+       720.53       14.6
   y6+       814.50       11.9
   b5+       887.43       13.9
```

```
216. MORG-339_phosphopeptide_TMT_20%ACN_032017_OT.8480.8480.3 (kQsSSEISLAVER)

   Ion         m/z   Intensity
   y5++      294.25       11.3
   b1+       358.38       13.4
   y4+       474.16       12.9
   b7++      535.51       13.9
   y5+       587.54       18.0
   a3+       625.59      251.8
   y11++     628.94       24.8
   b9++      635.11       16.0
   y6+       674.48       21.6
   y12++     693.17       13.1
   a11++     706.11       11.4
   b4+       740.23       10.8
   a12++     770.87        9.7
   b12++     784.71        9.6
   y7+       787.52       27.9
   a5+       799.45        9.5
   y8+       916.47       21.1
   y9+      1003.51       16.5
   y10+     1090.56       31.0
```

```
217. MORG-339_phosphopeptide_TMT_18%ACN_032017_OT.9850.9850.3 (sSASVSLPPGtPEk)

   Ion         m/z   Intensity
   y3++      301.83       18.5
   b1+       317.29       12.5
   y1+       376.32       13.0
   b7++      431.26       14.9
   y6++      469.11       16.5
   b8++      479.94       24.2
   y7++      517.98      145.5
   b9++      528.44       10.5
   b10++     557.07        8.0
   y8++      574.34       17.6
   y3+       602.43       52.5
   y9++      617.95       30.8
   b11++     647.22       11.5
   b5+       661.48       16.0
   a6+       720.19        7.7
   b6+       748.52       21.4
   b13++     760.24        7.9
   y4+       783.47       12.7
   y5+       840.43        8.0
   b7+       861.49      101.3
   a8+       930.29       10.9
   y6+       937.49       23.3
   y7+      1034.55       41.3
   b11+     1293.69        9.7
```

```
218. MORG-339_phosphopeptide_TMT_22%ACN_032017_OT.9248.9248.3 (iTRkPASVSPtTPtSPTEGEAS)

   Ion         m/z   Intensity
   y9++      479.59       29.8
   b6++      563.58       20.1
   b7++      606.92       21.0
   a8++      642.63       26.1
   b8++      656.64       61.8
   y7+       690.40       57.4
   b9++      699.95       42.6
   b10++     748.85       20.4
   y8+       777.41       24.2
   a11++     825.31       20.4
   b11++     839.10       47.7
   y17++     889.72       49.8
   a13++     924.00       39.2
   a4+       929.40       27.8
   b4+       957.53       29.4
   a14++    1014.35       53.1
   y10+     1055.38       80.0
   b15++    1072.25       23.6
   y11+     1156.43       29.6
   a17++    1157.52       26.9
   y21++    1245.94       20.3
   y13+     1434.53       21.8
   y14+     1521.57       24.6
```

```
219. MORG-339_phosphopeptide_TMT_6%ACN_032017_OT.6176.6176.3 (eVsDDEAEEkEEk)

   Ion         m/z   Intensity
   y3++      318.11        7.6
   b1+       359.31        9.4
   y1+       376.53        8.2
   b2+       458.32        7.8
   y4++      496.34       16.2
   y2+       505.51       10.3
   b7++      528.40        8.3
   y5++      561.09       17.7
   y3+       634.44        7.1
   a9++      643.35        7.9
   y7++      661.07       24.9
   y8++      725.49       37.7
   b4+       740.46       19.5
   y9++      783.12       40.7
   a5+       827.61        8.5
   b10++     835.99       10.1
   y10++     840.42       28.3
   b5+       855.57       15.5
   a11++     886.45       11.1
   b11++     900.48        7.9
   y11++     923.89       13.3
   a12++     951.35        7.2
   a6+       956.51        7.3
   b12++     964.87       11.2
   b6+       984.35        7.8
   b8+      1184.49        7.6
   b9+      1313.56        6.9
```

```
220. MORG-339_phosphopeptide_TMT_30%ACN_032017_OT.14634.14634.3 (sksVENFFLR)

   Ion         m/z   Intensity
   b1+       317.22       21.7
   a2++      323.77       10.7
   b2++      337.80       13.6
   b3++      421.18       14.4
   y3+       435.38        9.9
   a4++      456.34       12.5
   a5++      521.40       10.5
   b5++      535.33       29.4
   y8++      546.32       14.4
   a6++      578.67       16.3
   y4+       582.38       20.4
   b7++      665.96       20.5
   b2+       674.85       12.9
   y5+       696.45       26.6
   y9++      724.63       10.4
   a8++      725.52       13.0
   b8++      739.56       24.8
   a9++      781.72       13.1
   b9++      796.04       30.0
   y6+       825.42       27.0
   a4+       912.39        9.4
   y7+       924.50       33.7
   y8+      1091.42        8.8
```

```
221. MORG-339_phosphopeptide_TMT_12%ACN_032017_OT.5278.5278.3 (aLGPRGDVGsPDAPR)

   Ion         m/z   Intensity
   y2+       272.20       21.7
   a1+       273.23       10.9
   y3+       343.23       47.4
   b5++      362.94       11.0
   b7++      448.84       38.9
   b3+       471.39       11.1
   y9++      497.11       17.9
   b8++      498.32       15.4
   b9++      526.42       21.3
   a4+       540.15       10.7
   y5+       555.33       61.3
   b4+       568.63       12.2
   b10++     610.49        9.8
   y13++     680.59        9.6
   a5+       695.98        8.0
   b12++     716.53       75.7
   b5+       724.54        7.9
   a13++     737.72       11.9
   b13++     752.02       44.9
   y7+       779.43       13.4
   b6+       781.36       11.4
   b14++     800.65        8.1
   y8+       878.35       16.3
   b7+       896.55        8.8
```

```
222. MORG-339_phosphopeptide_TMT_14%ACN_032017_OT.5512.5512.3 (tASPPPPPk)

   Ion         m/z   Intensity
   y2++      237.27        4.4
   a3++      270.76        5.3
   b3++      284.67       10.2
   y3++      285.76       11.6
   a4++      319.37        4.4
   b4++      333.24        8.8
   y4++      334.27       29.1
   a5++      368.23       15.8
   y1+       376.27        9.2
   b5++      381.99       25.7
   a1+       382.87       19.6
   a6++      416.58       25.4
   y6++      431.04       31.7
   a2+       454.43        8.7
   a7++      465.11        8.2
   y2+       473.36       12.7
   b8++      527.64        5.8
   b3+       569.33       15.1
   y3+       570.48       26.5
   a4+       638.34        4.8
   b4+       666.42        6.9
   y4+       667.47       12.3
   b5+       763.33       25.2
   y5+       764.40       10.3
   b6+       860.51        8.3
   y6+       861.51        5.5
```

```
223. MORG-339_phosphopeptide_TMT_8%ACN_032017_OT.3208.3208.3 (rsPsPYYSR)

   Ion         m/z   Intensity
   b1++      193.88        8.8
   y2+       262.19       15.7
   a2++      263.21       22.2
   b2++      277.11       49.1
   y4++      294.49        9.7
   a3++      311.22       36.7
   b3++      325.76        9.4
   y5++      343.21       46.1
   a4++      395.14       15.0
   b4++      409.25       32.6
   y3+       425.30       25.7
   y6++      426.33       42.5
   b5++      457.40       11.5
   y7++      475.00      101.9
   a2+       525.30       28.4
   b6++      539.42       19.8
   b2+       553.21       22.1
   y4+       588.37       20.5
   a7++      607.00       17.7
   b7++      620.83       20.7
   a3+       622.01       12.3
   b8++      664.45       14.7
   y5+       685.32      157.4
   b4+       817.31       14.6
   y6+       852.39       17.1
   y7+       949.46       30.7
```

```
224. MORG-339_phosphopeptide_TMT_30%ACN_032017_OT.11974.11974.3 (kGsFGLHAQPEFLR)

   Ion         m/z   Intensity
   y7++      431.04       18.8
   y3+       435.29       20.7
   y8++      499.59       22.6
   y10++     584.69       30.7
   y11++     657.59       16.4
   y5+       661.30       36.3
   b10++     667.22       24.9
   a4+       700.98       32.0
   b11++     730.93       18.3
   a5+       758.68       21.6
   y6+       789.56       24.7
   b12++     805.17       17.0
   y7+       860.53       33.5
   b13++     861.48       26.6
   a6+       871.77       18.1
   b6+       899.63       15.5
   y8+       997.71       35.8
```

```
225. MORG-339_phosphopeptide_TMT_14%ACN_032017_OT.9467.9467.3 (tASLTSAASIDGSR)

   Ion         m/z   Intensity
   y2+       262.20        7.7
   y6++      317.30       14.5
   y3+       319.19       24.8
   y7++      353.23        7.8
   a1+       383.24        8.6
   y8++      388.57        9.7
   b5++      392.14       12.6
   a6++      422.13        9.6
   y9++      432.39       11.0
   y4+       434.22       31.4
   b6++      435.93        8.5
   a7++      457.62       16.9
   b7++      471.26       28.2
   b2+       481.99        8.2
   y10++     482.90       12.2
   a8++      492.32       14.1
   b8++      506.88       44.8
   a9++      536.53       13.4
   y11++     539.44       32.1
   a3+       541.31       16.1
   b10++     606.85       25.5
   y6+       634.36       38.2
   b11++     664.25       17.6
   b4+       682.38        9.0
   b12++     692.88       15.6
   y7+       705.41       26.7
   b13++     736.43       11.9
   y8+       776.50        9.7
   b5+       783.50       15.8
   y9+       863.46        9.8
   b6+       870.49        9.0
   b7+       941.54       11.2
   b8+      1012.54       14.3
   b9+      1099.62        6.3
```

```
226. MORG-339_phosphopeptide_TMT_16%ACN_032017_OT.1853.1853.3 (sPLRPPRER)

   Ion         m/z   Intensity
   y1+       175.11       22.5
   y3++      230.21        9.8
   y5++      327.82       20.3
   a4++      368.33       14.8
   b4++      382.34       63.5
   b1+       397.23       39.1
   y6++      405.96       23.6
   b5++      430.84       15.3
   y3+       459.96       21.7
   a6++      465.06       20.1
   a2+       466.10       23.1
   y8++      510.97       68.6
   y4+       557.35       20.3
   a3+       579.38       11.7
   a8++      607.53       24.4
   b8++      621.99       66.1
   y5+       654.41       47.5
   b4+       763.46       17.7
```

```
227. MORG-339_phosphopeptide_TMT_18%ACN_032017_OT.2819.2819.3 (gRAsPGGVSTSSSDGkAEk)

   Ion         m/z   Intensity
   y1+       376.38       10.3
   a3+       486.73       10.4
   b8++      495.95       41.9
   y8++      640.70        9.4
   y9++      683.87       10.3
   y10++     734.73       10.4
   a14++     764.50       13.3
   y11++     777.81       11.3
   b5+       778.63       24.8
   a15++     792.86       10.4
   b15++     806.84       24.5
   a6+       807.68       18.6
   y12++     827.61       21.2
   b6+       835.56       14.0
   y4+       933.26       55.3
   b16++     985.77       10.8
   y5+       990.65       19.6
   b8+       991.62       17.8
   y16++    1016.34       10.5
   b18++    1085.83       11.9
   a10+     1151.30        9.4
   b14+     1555.67       19.2
```

```
228. MORG-339_phosphopeptide_TMT_20%ACN_032017_OT.13911.13911.4 (vGDTSLDPNDFDFTVtGRGSPSRR)

   Ion         m/z   Intensity
   b5+++     230.15        7.7
   a8+++     329.30        8.3
   b5++      345.12        9.9
   a6++      387.82        8.9
   b6++      401.68       14.1
   b10+++    415.25       13.7
   b7++      459.52       17.5
   b3+       501.42       18.0
   b8++      507.67       12.7
   y9++      527.37       10.3
   y14+++    555.39       10.2
   y10++     577.08       11.7
   y15+++    593.43        7.7
   y5+       602.44       24.9
   a10++     608.07        8.4
   b10++     622.55        8.7
   y11++     627.51       17.0
   y16+++    631.29       23.1
   y6+       659.00       11.2
   a5+       661.27       14.6
   y17+++    663.96      128.0
   a16+++    669.61       15.3
   b16+++    678.76        8.8
   a11++     682.26       10.9
   a17+++    688.21       15.3
   b5+       689.42       51.4
   b11++     696.20       22.6
   b17+++    697.26       22.6
   y12++     700.65       21.2
   y18+++    702.10       65.6
   a18+++    740.14       13.3
   y13++     758.38       19.9
   a19+++    759.08       17.7
   y20+++    768.83       15.3
   a6+       774.06        7.9
   b6+       802.54       38.7
   y14++     832.13       34.4
   a7+       889.64       28.5
   b23+++    911.16       11.7
   b7+       917.52       68.0
   y16++     946.62       38.0
   a8+       986.31       20.9
   y17++     995.26      120.7
   y18++    1052.72       79.5
   y9+      1053.56       39.1
   a9+      1100.70        8.1
   y19++    1109.18       22.6
   a18++    1110.08       11.0
   b10+     1243.54        8.4
```

```
229. MORG-339_phosphopeptide_TMT_10%ACN_032017_OT.9252.9252.3 (aGDVLEDsPk)

   Ion         m/z   Intensity
   y2++      237.18       24.6
   a4++      272.32        5.6
   b4++      286.20        6.4
   y3++      320.53       14.9
   a5++      329.30       23.5
   b5++      343.25       12.2
   y1+       376.27       15.3
   y4++      378.28       37.4
   a6++      393.75        7.0
   b6++      407.63       28.4
   y5++      442.82       40.6
   b7++      465.22        9.1
   y2+       473.33       32.5
   y7++      548.86       17.8
   b4+       572.37       19.1
   y3+       640.28       10.7
   a5+       657.33        7.6
   b5+       685.43       42.9
   y4+       755.48       20.0
   a6+       786.25        5.5
   b6+       814.51       41.4
   y5+       884.58        9.2
   b7+       929.46       16.3
   y6+       997.62        5.5
   y7+      1096.53       15.4
```

```
230. MORG-339_phosphopeptide_TMT_8%ACN_032017_OT.2264.2264.3 (sRVsVsPGR)

   Ion         m/z   Intensity
   y1+       175.17       11.0
   b2++      237.22       13.2
   b3++      286.81       20.5
   y3+       329.28       71.8
   b4++      370.15       22.1
   a5++      405.47       81.7
   b5++      419.59       21.0
   y7++      431.22       13.2
   y4+       496.22       22.6
   b6++      503.42       20.6
   b8++      580.43       19.6
   y5+       595.28       19.6
```

```
231. MORG-339_phosphopeptide_TMT_22%ACN_032017_OT.10819.10819.4 (qEsDPEDDDVkkPALQSSVVATSk)

   Ion         m/z   Intensity
   y2+       463.43       14.8
   y3+       564.48       24.9
   y4+       635.49       27.9
   y5+       734.58       34.4
   y15+++    744.27       50.8
   b4+       769.32       37.5
   b14+++    774.60       15.1
   y16+++    782.54       46.0
   y17+++    820.96       45.9
   y6+       833.59       23.0
   a16+++    845.33       15.4
   b16+++    854.75       31.3
   y18+++    859.65       34.2
   b5+       866.48       26.0
   a17+++    874.43       34.0
   b17+++    884.30       15.2
   a18+++    903.86       15.5
   b18+++    913.49       29.8
   y20+++    934.73       77.9
   a19+++    936.88       19.7
   b19+++    945.97       22.3
   a6+       967.04       22.6
   y21+++    973.23       33.2
   b20+++    979.21       45.5
   a21+++    993.14       15.4
   b6+       995.59       16.7
   b21+++   1002.93       37.0
   y8+      1007.69       22.1
   y22+++   1028.50       28.6
   b22+++   1036.93       17.3
   b23+++   1065.44       17.1
   y14++    1066.08       15.0
   b7+      1110.43       21.9
   y15++    1115.89       14.9
   b8+      1225.19       23.2
   b9+      1340.51       17.8
```

```
232. MORG-339_phosphopeptide_TMT_14%ACN_032017_OT.5248.5248.3 (sQAQQPQkEAALSs)

   Ion         m/z   Intensity
   y2+       273.08       15.3
   b5++      386.30       12.7
   y4+       457.20       14.9
   y5+       528.33       14.8
   y8++      571.53       19.3
   a4+       616.71       11.8
   y9++      620.53       39.6
   b4+       644.50       33.2
   y6+       657.48       27.5
   b8++      678.30       15.0
   y10++     684.56       21.1
   b9++      742.65       27.2
   a5+       744.87       15.7
   b5+       772.43       30.8
   b10++     778.16       38.6
   y12++     784.34       15.4
   b11++     813.73       41.4
   a12++     855.91       13.0
   b12++     870.25       48.4
   b13++     913.81       14.6
   y7+      1014.54       12.3
```

```
233. MORG-339_phosphopeptide_TMT_25%ACN_032017_OT.17394.17394.3 (kIVETINSDsDSEFGIPk)

   Ion         m/z   Intensity
   y1+       376.33       21.3
   y2+       473.38       90.8
   y3+       586.45       19.3
   y4+       643.46       41.5
   y9++      644.47       24.4
   y5+       790.61       41.6
   b12++     799.70       19.3
   a13++     850.63       26.4
   a6+       885.83       18.7
   y7+      1006.60       37.9
   a16++    1008.76       16.8
   b16++    1022.77       16.5
   y16++    1023.66       20.5
   b17++    1071.25       17.7
   y17++    1080.14       29.6
   b8+      1114.82       20.6
   y8+      1121.52       28.3
   y9+      1288.53       26.7
```

```
234. MORG-339_phosphopeptide_TMT_10%ACN_032017_OT.14885.14885.4 (dDDDIDLFGsDDEEESEEAkkLR)

   Ion         m/z   Intensity
   y5+++     358.56       22.6
   b5++      401.85       18.5
   b2+       460.27       13.9
   y8+++     473.95       22.9
   y9+++     516.65       23.6
   y5++      537.56       19.0
   y10+++    559.72       20.4
   a8++      575.30       31.4
   y11+++    602.96       43.0
   a9++      604.16       17.9
   y12+++    641.27       57.3
   y7++      666.59       20.1
   y13+++    679.69       45.9
   b4+       690.39       44.8
   y8++      710.11       33.8
   y14+++    735.32       17.8
   y15+++    754.30       26.4
   y9++      774.26       19.6
   a5+       775.11       14.7
   b5+       803.30       55.1
   b19+++    812.34       14.4
   y10++     838.95       30.1
   y17+++    840.39       52.9
   y18+++    879.28       16.1
   y11++     903.88       38.6
   y19+++    917.18       35.5
   b6+       918.41       88.4
   a20+++    922.71       30.1
   y20+++    955.18       29.3
   y12++     961.20       41.0
   y21+++    993.46       28.2
   y13++    1018.50       26.4
   y22+++   1031.48       66.3
   b21+++   1050.78       19.3
   b16++    1053.86       26.4
   y5+      1074.14       14.0
   b22+++   1088.95       26.1
   y14++    1102.18       21.1
   y15++    1130.77       44.9
   b8+      1178.51       35.3
   a19++    1204.40       34.2
   b11+     1517.72       14.7
   b12+     1632.65       17.1
```

```
235. MORG-339_phosphopeptide_TMT_14%ACN_032017_OT.12593.12593.3 (eRLkGQEDSLASAVDATTGQEAcDsD)

   Ion         m/z   Intensity
   b4++      493.59       24.2
   b6++      586.16       22.1
   y5+       647.27       24.5
   b7++      650.71       36.3
   b8++      708.21      104.1
   b9++      751.72       27.2
   a10++     794.30       28.6
   b10++     808.11       40.2
   b11++     843.60       54.3
   b12++     887.11       42.1
   a13++     908.89       21.7
   b13++     922.50       44.5
   a14++     958.33       25.2
   y8+       961.38       34.3
   b14++     972.14       69.5
   b4+       985.94       22.9
   a15++    1015.49       32.4
   b15++    1029.55      112.4
   y9+      1062.69       67.4
   b17++    1115.54       35.5
   y10+     1163.32       33.1
   b18++    1166.42       47.3
   b19++    1194.94       40.4
   y11+     1234.43       54.2
   b20++    1258.97       65.3
   b21++    1323.48       73.7
   y12+     1349.44       36.0
   b22++    1358.60       35.1
   b8+      1414.85       23.9
   y13+     1448.51       23.6
   y25++    1467.51       19.6
   y14+     1519.46       33.5
   y15+     1606.58       23.6
   y16+     1677.69       19.9
   y18+     1877.57       27.9
```

```
236. MORG-339_phosphopeptide_TMT_18%ACN_032017_OT.7694.7694.3 (eTkEVQTsELk)

   Ion         m/z   Intensity
   y1+       376.31       16.4
   b2+       459.86       11.8
   b4++      473.92       22.4
   y2+       489.37       17.9
   a5++      509.28       13.1
   b5++      523.40       24.0
   a6++      573.11       16.3
   b6++      587.27       25.2
   y3+       618.45       25.1
   y8++      622.07       17.3
   b7++      637.96       30.7
   b8++      720.89       18.0
   b9++      785.48       16.8
   y9++      800.23       14.5
   b10++     842.53       22.9
   y10++     851.00       15.9
   y5+       886.64       13.8
   b5+      1045.69       13.1
```

```
237. MORG-339_phosphopeptide_TMT_12%ACN_032017_OT.4157.4157.3 (rVSSSRsVsGsPEPAAk)

   Ion         m/z   Intensity
   b1+       386.39        9.6
   b6++      451.80       11.3
   b7++      535.75       10.1
   b8++      584.90        8.9
   y4+       615.47       25.5
   y9++      616.34       11.4
   b4+       659.89        9.3
   a10++     683.04       17.8
   b5+       746.10       11.5
   a11++     766.12       22.9
   b11++     780.75       25.1
   y6+       841.68       32.0
   a6+       874.38       13.0
   a13++     879.69       13.5
   b13++     893.29       22.8
   y14++     915.19       13.4
   b15++     977.60       10.0
   y16++    1007.78        9.4
   y8+      1065.57        9.4
```

```
238. MORG-339_phosphopeptide_TMT_18%ACN_032017_OT.10763.10763.3 (gPLPAAPPTAPERQPsWER)

   Ion         m/z   Intensity
   y3+       490.37       25.7
   b3+       497.32       14.5
   y7++      519.94       24.5
   y9++      632.97       15.9
   y4+       657.33       15.4
   b5+       665.56       18.2
   a13++     728.60       23.7
   b6+       736.55       46.8
   b13++     742.79       36.9
   y5+       754.42       37.6
   y12++     767.12       37.7
   b14++     807.23       49.0
   y13++     815.83       26.5
   a15++     841.24       19.1
   y14++     851.43       23.0
   b15++     855.66       20.0
   y15++     886.49       24.0
   y17++     991.49       23.9
   b18++    1096.73       28.6
   y8+      1167.70       14.4
   y9+      1264.64       18.2
   b12+     1328.70       26.3
```

```
239. MORG-339_phosphopeptide_TMT_6%ACN_032017_OT.13746.13746.4 (eEsGssEEEAVLQRAk)

   Ion         m/z   Intensity
   y3++      302.42        4.9
   y4++      366.24        9.8
   y1+       376.57        5.2
   y8+++     381.97        3.8
   a7+++     383.32        3.2
   y5++      422.76       13.7
   y2+       447.35       11.3
   y6++      472.45       14.9
   b2+       488.00        3.6
   y7++      508.00       16.3
   a6++      509.38        4.2
   b6++      524.13        4.1
   a11+++    525.78        5.5
   b11+++    534.88        5.4
   a12+++    563.61       36.7
   b12+++    572.48       14.7
   y12+++    579.08        3.8
   b7++      588.12       52.1
   y13+++    598.08        6.3
   y3+       603.83       12.9
   y9++      637.00        5.6
   b8++      652.77       10.9
   y14+++    654.12        5.1
   b3+       654.97        3.7
   b14+++    667.66       15.4
   b15+++    691.73        4.4
   b9++      717.16       11.2
   b10++     752.69       15.0
   b11++     802.44       18.6
   b12++     858.92       13.4
   b5+       879.25        3.6
   b14++    1000.91        3.6
   b7+      1175.29        3.3
   b8+      1304.27        3.9
   b10+     1504.32        3.0
```

```
240. MORG-339_phosphopeptide_TMT_10%ACN_032017_OT.8605.8605.3 (sEsPPPLSEPk)

   Ion         m/z   Intensity
   y2++      237.18       22.9
   y3++      301.69        9.2
   b1+       317.36        7.9
   y4++      345.20       12.6
   b4++      355.20        8.4
   y1+       376.27       15.2
   b5++      403.89       14.2
   a6++      438.84       15.6
   b2+       446.26       11.1
   y6++      450.48        9.6
   b6++      452.65       18.2
   y2+       473.37       69.2
   a7++      495.15       30.1
   y7++      498.56        6.4
   b7++      509.38       53.3
   a8++      538.95       22.4
   y8++      547.39       19.5
   b8++      552.60       29.3
   y3+       602.38       21.5
   a9++      603.44       12.8
   b3+       613.22       10.9
   b9++      617.38       37.7
   b10++     666.02        7.3
   y4+       689.46       24.2
   b4+       710.36        8.7
   b5+       807.34       15.1
   b7+      1017.48       24.6
   b8+      1104.59       12.1
   b9+      1233.55       22.0
```

```
241. MORG-339_phosphopeptide_TMT_30%ACN_032017_OT.8521.8521.4 (vkFSSQQASTkQSNASsDVEVEEk)

   Ion         m/z   Intensity
   y1+       376.37       44.2
   y2+       505.36       68.1
   b6++      568.39       55.3
   a7++      618.28       31.0
   b11+++    627.86       36.9
   b7++      632.48       66.7
   y3+       634.39      105.5
   b8++      667.79       44.7
   b12+++    670.39       48.4
   b13+++    699.10       46.7
   b9++      711.64       56.6
   y4+       733.47       90.3
   b14+++    737.33       70.4
   b15+++    761.05      100.3
   b10++     761.80       42.6
   b16+++    790.22       42.8
   b17+++    845.36       96.3
   y5+       862.57      101.6
   y13++     865.50       46.6
   y19+++    868.69       50.2
   a18+++    874.57      377.8
   b18+++    884.03      152.4
   b19+++    917.06       75.4
   b11++     940.82       44.7
   a20+++    950.78       47.3
   b20+++    960.21      161.4
   y6+       961.45      103.3
   b21+++    992.93       90.4
   b12++    1004.62       39.3
   b22+++   1036.00       91.4
   b13++    1047.89       36.5
   y7+      1076.66       53.0
   b23+++   1079.09       86.5
   y15++    1095.11       55.1
   b14++    1105.50       57.3
   a6+      1107.75       38.1
   y16++    1139.01       36.1
   b15++    1140.78       62.0
   y17++    1174.23       50.5
   b19++    1374.99       40.6
```

```
242. MORG-339_phosphopeptide_TMT_16%ACN_032017_OT.11241.11241.3 (tVTMIsPEDEQk)

   Ion         m/z   Intensity
   a4++      317.24       20.6
   y4++      375.19        9.6
   y1+       376.27       41.3
   b2+       430.44       16.3
   y6++      487.66       47.2
   a3+       503.20       11.8
   y2+       504.32       37.7
   a7++      506.61       11.3
   b3+       531.35       15.6
   y7++      571.34       39.9
   b8++      584.90       12.3
   a9++      627.93       35.2
   y3+       633.42       21.8
   b4+       662.41       21.8
   b10++     706.72       30.3
   y10++     743.83       10.2
   y4+       748.44       26.0
   b11++     770.81       20.4
   b5+       775.48       51.8
   b6+       942.42       17.9
   y7+      1141.56       12.8
   b8+      1168.54       21.6
   b9+      1283.69       20.7
   b10+     1412.66       14.9
```

```
243. MORG-339_phosphopeptide_TMT_6%ACN_032017_OT.6880.6880.2 (sVsGsPEPAAk)

   Ion         m/z   Intensity
   b1+       317.18        6.7
   y1+       376.38       18.5
   b2+       416.24       10.0
   y2+       447.29       14.1
   a4+       612.37       11.8
   y4+       615.48       33.2
   b4+       640.39        6.6
   y5+       744.21        8.9
   a5+       779.29        6.4
   b5+       807.25       24.4
   y6+       841.49       51.0
   y7+      1008.56        8.9
   b7+      1033.37       22.9
   y8+      1065.55       15.3
   b9+      1201.50       12.0
   y9+      1232.58        8.6
   b10+     1272.53       18.0
   y10+     1331.64        6.1
```

```
244. MORG-339_phosphopeptide_TMT_22%ACN_032017_OT.7966.7966.3 (lALVPGHSGsQDTQR)

   Ion         m/z   Intensity
   b3++      264.19       17.2
   b1+       343.36       19.0
   y3+       404.28       37.2
   b2+       414.29       26.6
   a7++      445.53       35.0
   b7++      459.56       49.0
   y8++      479.97       18.7
   a8++      488.69       16.2
   a3+       499.45       30.6
   b8++      503.07       48.6
   y4+       519.41       20.0
   b3+       527.43       69.1
   b9++      531.60       23.7
   y9++      548.21       23.0
   y10++     576.58       70.6
   a4+       598.40       36.6
   a10++     600.38       24.5
   b10++     615.14       34.6
   y5+       647.38       34.6
   b11++     678.64       16.6
   b12++     736.64       52.0
   y14++     767.07       16.4
   a13++     773.32       34.6
   b6+       780.52       38.0
   b13++     786.85       15.5
   y6+       814.09       14.1
   b14++     851.27       21.7
   y7+       871.42       21.5
   b7+       917.66       14.7
   y8+       958.36       39.2
   b8+      1004.45       18.3
   y10+     1152.52       15.1
```

```
245. MORG-339_phosphopeptide_TMT_4%ACN_032017_OT.5987.5987.3 (lYERRGGGSGGGDEsEGEEVDED)

   Ion         m/z   Intensity
   b5++      474.61       14.1
   y4+       477.22        9.7
   b6++      503.11       10.3
   b7++      531.31        9.0
   y5+       606.30       12.4
   a3+       607.38       11.0
   b10++     631.92       13.6
   b3+       635.45       15.5
   b11++     660.30       10.1
   b12++     688.51        8.3
   y6+       735.68       14.2
   b13++     746.55       65.4
   y7+       792.39       11.9
   b14++     811.18       29.3
   a15++     880.70       16.9
   y18++     881.74       13.8
   y8+       921.42        9.8
   y19++     959.08       19.1
   b17++     987.73       12.3
   a18++    1037.70       16.6
   b18++    1052.19       24.2
   b20++    1166.26       30.3
   y10+     1217.43       16.1
   b21++    1223.83       17.3
   a11+     1291.41        8.9
```

```
246. MORG-339_phosphopeptide_TMT_60%ACN_032017_OT.10328.10328.3 (nkAITSLLGGGsPk)

   Ion         m/z   Intensity
   y1+       376.34       16.0
   b3++      386.94       14.0
   b4++      443.32       18.8
   y2+       473.37       75.0
   b5++      493.88       22.2
   b6++      537.55       33.5
   a7++      579.99       25.7
   b7++      594.09       67.9
   a8++      636.58       19.6
   y3+       640.38       16.3
   b8++      650.63       84.5
   y11++     670.11       23.1
   a2+       673.50       17.9
   b9++      679.13       81.7
   a10++     693.32       31.7
   b10++     707.63       13.6
   a11++     722.07       13.3
   b11++     736.16       34.3
   y5+       754.42       28.0
   b3+       772.36       16.6
   y6+       811.45       47.3
   b12++     819.68       23.6
   b4+       885.65       16.1
   y7+       924.52       27.3
   b7+      1186.79       14.3
   b8+      1299.89       13.5
```

```
247. MORG-339_phosphopeptide_TMT_22%ACN_032017_OT.10426.10426.4 (asNEDVTkAVDITTPkAAR)

   Ion         m/z   Intensity
   y5++      386.40       15.8
   y6++      436.94       17.5
   b2+       468.23       16.0
   y8++      544.12      139.5
   y12+++    577.17       17.0
   y9++      601.76       24.2
   y13+++    610.99       31.3
   b13+++    628.10       15.0
   y14+++    643.98      127.2
   y10++     650.73       38.7
   a14+++    652.50       17.9
   b14+++    661.68       21.7
   y4+       673.98       48.4
   a8++      678.68       50.6
   y15+++    682.37       38.6
   a4+       683.24       15.5
   b4+       711.42       27.0
   y16+++    725.19       21.0
   b9++      727.84       26.3
   a10++     763.43       77.4
   y5+       771.45       21.4
   b10++     777.34       34.6
   b16+++    812.97       21.8
   y18+++    818.78       20.1
   b5+       826.32       53.5
   a17+++    828.26       42.9
   b11++     835.14      126.3
   b17+++    837.22       24.7
   b18+++    861.05       50.0
   y12++     865.33       15.3
   y6+       872.64       18.2
   y7+       973.65       17.1
   y9+      1201.75       20.0
```

```
248. MORG-339_phosphopeptide_TMT_18%ACN_032017_OT.15388.15388.3 (eADDEsLDEQASASkLSLLSR)

   Ion         m/z   Intensity
   b3+       545.42       38.7
   y9++      602.53       16.5
   y10++     646.20       26.8
   b4+       660.34       38.6
   y11++     681.51       15.8
   y12++     745.87       28.0
   b5+       789.44       18.4
   y13++     810.24       79.9
   y14++     867.65       30.4
   b14++     879.76       17.7
   y15++     924.10       46.1
   y17++    1071.98       19.8
   a16++    1100.33       19.0
   b16++    1114.27       21.7
   y18++    1129.85       46.2
   y8+      1132.84       13.4
   b8+      1184.51       30.0
   y19++    1187.55       21.8
   b18++    1214.30       17.7
   y20++    1222.84       27.3
   a19++    1257.04       15.8
   b19++    1271.49       17.9
   b9+      1313.48       17.9
   b20++    1314.45       14.5
```

```
249. MORG-339_phosphopeptide_TMT_30%ACN_032017_OT.12575.12575.4 (eSVPDFPLsPPkkk)

   Ion         m/z   Intensity
   b9+++     427.96       14.1
   y5+++     429.13      158.0
   b2+       446.30       14.6
   b6++      452.83       14.3
   y6+++     484.83       32.8
   a3+       517.23       15.1
   y7+++     522.20       48.9
   a8++      544.03       13.1
   b3+       545.38       30.8
   y3++      546.39       16.3
   y8+++     554.88       40.2
   b8++      557.31       15.9
   y4++      594.17       26.7
   y9+++     603.96       88.5
   a4+       614.50       18.3
   a9++      627.45       95.9
   y11+++    674.72       25.1
   a10++     675.51       13.2
   y12+++    707.75       19.6
   a11++     724.20       17.7
   y6++      726.47       18.0
   y13+++    736.05       17.6
   b5+       757.40       75.0
   y7++      782.79       50.4
   y8++      831.73       24.7
   b6+       904.43       39.2
   y9++      905.47       33.4
```

```
250. MORG-339_phosphopeptide_TMT_14%ACN_032017_OT.14969.14969.3 (tGDLGIPPNPEDRsPsPEPIYNSEGk)

   Ion         m/z   Intensity
   b3+       503.42        8.8
   b5+       673.48       12.2
   a6+       758.53        9.7
   y5+       763.46        8.4
   y11++     765.76        6.8
   b6+       786.55       23.0
   b13++     796.74        6.8
   a15++     914.26       11.9
   y6+       926.58       14.9
   y14++     975.53       14.4
   a16++     998.44        7.1
   a17++    1046.80        7.4
   b17++    1060.45        7.3
   b9+      1094.37       25.1
   y16++    1096.97       58.8
   a18++    1111.45       23.9
   y8+      1136.70       13.9
   a10+     1163.80        7.3
   y18++    1203.08        9.6
   y19++    1251.82       19.7
   y20++    1300.40       25.5
   b11+     1320.54        8.0
   y10+     1362.74       15.8
   b12+     1435.81       11.9
   y23++    1441.98       12.0
   y11+     1529.94        7.7
   y12+     1626.81       14.8
```

```
251. MORG-339_phosphopeptide_TMT_18%ACN_032017_OT.10788.10788.3 (nENASFPRtPELVk)

   Ion         m/z   Intensity
   y1+       376.33       17.2
   b2+       473.38       15.2
   y2+       475.39       23.8
   b8++      573.30       14.7
   y7++      576.27       16.7
   b3+       587.30       14.6
   y3+       588.54       25.0
   y8++      624.54       34.8
   a4+       630.68       11.2
   a9++      650.11       20.3
   b4+       658.41       13.0
   b9++      664.04       64.1
   y9++      698.38       27.1
   b5+       745.55       14.2
   a11++     762.69       13.0
   y11++     777.16       31.8
   y5+       814.55       91.5
   a12++     819.75       13.3
   b12++     833.75       33.1
   y12++     834.76       24.8
   a13++     869.28       12.3
   b13++     883.31       21.2
   b6+       892.47       32.0
```

```
252. MORG-339_phosphopeptide_TMT_20%ACN_032017_OT.10362.10362.3 (dELTEsPkYIQk)

   Ion         m/z   Intensity
   a1+       317.18       12.3
   b1+       345.26       16.3
   y2+       504.35       16.9
   y5++      569.51       12.7
   y6++      618.01      123.6
   b4+       688.44       21.9
   y7++      701.36       47.9
   a8++      705.97       18.6
   b8++      719.70       43.1
   y4+       780.57       23.5
   b9++      801.31       18.7
   y9++      816.58       16.8
   b5+       817.54       19.4
   b10++     857.91       18.9
   y10++     873.11       18.9
   b11++     922.24       16.9
   y11++     937.67       15.5
```

```
253. MORG-339_phosphopeptide_TMT_22%ACN_032017_OT.14113.14113.2 (iVAPPGGRANITsLG)

   Ion         m/z   Intensity
   b1+       343.33       16.0
   b2+       442.38       21.7
   b3+       513.51       32.3
   b4+       610.50       11.8
   b7+       821.92       26.0
   y8+       911.25       15.3
   b8+       977.91       15.5
   y10+     1025.47       16.1
   y11+     1122.62       24.9
   y12+     1219.59       24.1
   y13+     1290.77       11.2
```

```
254. MORG-339_phosphopeptide_TMT_22%ACN_032017_OT.12056.12056.3 (tIksPcDSGYSYETIEk)

   Ion         m/z   Intensity
   y1+       376.35       22.5
   b3++      401.33       18.1
   b2+       444.46       18.5
   y2+       505.36       29.4
   y6++      506.28       18.5
   b6++      613.27       18.2
   y3+       618.52       25.1
   y8++      631.11       28.3
   b7++      670.92       44.2
   y4+       719.52       37.4
   b3+       801.69       19.5
   a10++     810.71       26.1
   b10++     824.73       37.3
   y5+       848.60       34.6
   b11++     867.90       20.0
   y13++     889.57       75.7
   a4+       940.40       16.6
   y6+      1011.64       33.0
   b13++    1013.86       29.4
   a14++    1050.32       18.1
   b14++    1064.31       18.9
   y7+      1098.61       37.8
   b15++    1121.35       20.2
   y15++    1151.81       21.7
   b16++    1185.75       26.1
   y8+      1261.70       21.7
   y9+      1318.71       22.1
   y10+     1405.75       51.9
   y13+     1777.86       35.6
```

```
255. MORG-339_phosphopeptide_TMT_16%ACN_032017_OT.6568.6568.2 (qIsQDVk)

   Ion         m/z   Intensity
   y1+       376.32       17.7
   b2+       471.37       14.7
   y2+       475.41       32.4
   y3+       590.17       15.5
   a3+       610.62       21.4
   b3+       638.55       34.0
   y4+       718.55       18.3
   a4+       738.48       10.6
   b4+       766.25       13.5
   b5+       881.42       30.9
   y5+       885.43       11.7
   b6+       980.52       17.6
```

```
256. MORG-339_phosphopeptide_TMT_16%ACN_032017_OT.6159.6159.2 (sRsDIDVNAAAGAk)

   Ion         m/z   Intensity
   y5+       646.55        9.4
   b4+       755.34        9.5
   y7+       831.63       10.3
   b5+       868.68       10.9
   y8+       930.59       26.4
   b6+       983.49       14.7
   y9+      1045.64       11.0
   a7+      1054.04        8.9
   b7+      1082.37       12.0
   y10+     1158.80       14.0
   y11+     1273.77        9.2
```

```
257. MORG-339_phosphopeptide_TMT_18%ACN_032017_OT.13243.13243.4 (aLDIYSAVDDAsHEkELPR)

   Ion         m/z   Intensity
   y2+       272.21       12.5
   a1+       273.29        8.4
   b5++      403.59       12.6
   b6++      446.70       17.1
   y9+++     459.47       12.3
   b7++      482.44       11.8
   y10+++    497.76       34.8
   a3+       501.01        9.1
   a8++      518.12        8.5
   b3+       529.35       15.1
   b8++      531.82       10.4
   y11+++    536.18       18.8
   y7++      569.06       22.3
   b9++      589.21       14.5
   a14+++    590.16        9.2
   y13+++    592.65       20.6
   a4+       614.36       13.8
   y14+++    621.93       25.2
   b4+       642.44       38.4
   b10++     647.29       29.0
   y15+++    676.46        9.5
   y9++      688.57       10.4
   a15+++    709.53        9.3
   y16+++    713.83       12.5
   y10++     746.04       14.5
   b16+++    762.05       17.3
   b17+++    799.64       23.6
   y11++     803.67       59.2
   b5+       805.42       21.8
   y12++     853.20       39.4
   y13++     888.60       28.4
   b6+       892.54       12.3
   y14++     932.12       30.9
   b7+       963.60       27.0
   y6+      1000.61        9.3
   b8+      1062.67       12.2
   b9+      1177.60       18.1
```

```
258. MORG-339_phosphopeptide_TMT_25%ACN_032017_OT.7986.7986.3 (kcSPGSPtDPNATLSk)

   Ion         m/z   Intensity
   y1+       376.37       21.3
   y2+       463.29       29.3
   y3+       576.44       27.8
   b10++     668.65       27.5
   y10++     676.85       22.6
   y4+       677.55       28.7
   b11++     726.10       42.6
   a12++     747.20       25.6
   y12++     748.44       26.9
   b5+       759.47       33.4
   b12++     761.35       34.0
   a13++     797.85       75.2
   a6+       818.15       22.3
   y14++     840.75       23.3
   a14++     854.01       27.1
   b14++     868.36       26.2
   a15++     898.30       28.2
   b15++     911.77       29.4
   a7+       915.50       24.7
   y7+       959.63       96.6
   y8+      1074.66       24.1
```

```
259. MORG-339_phosphopeptide_TMT_16%ACN_032017_OT.7207.7207.2 (sIcSLGAR)

   Ion         m/z   Intensity
   y3+       303.16       14.8
   b1+       397.22       21.8
   y4+       416.28       22.8
   y5+       503.34       23.6
   b2+       509.99       18.3
   y6+       663.37       22.4
   b3+       670.31       15.1
   a4+       729.79       15.8
   b4+       757.35       13.8
   y7+       776.48       41.8
   a6+       899.48       18.4
   b6+       927.47       18.5
   b7+       998.46       23.8
```

```
260. MORG-339_phosphopeptide_TMT_18%ACN_032017_OT.5943.5943.3 (sLTNSHLEk)

   Ion         m/z   Intensity
   y1+       376.31       32.7
   y4++      378.36       28.8
   b1+       397.19       18.2
   y5++      421.68       24.2
   a6++      461.37       55.1
   b6++      475.29       64.2
   y6++      478.84       33.0
   y2+       505.34       80.8
   b2+       509.91       14.4
   a7++      518.08       13.9
   b7++      531.85       22.1
   a8++      582.31       10.4
   y8++      585.90       30.1
   b8++      596.37       35.9
   b3+       611.42       12.3
   y3+       618.42       73.0
   a4+       697.46       12.7
   b4+       725.34       21.6
   y4+       755.53       13.1
   b5+       812.46       17.5
   b6+       949.56        9.6
```

```
261. MORG-339_phosphopeptide_TMT_10%ACN_032017_OT.6219.6219.3 (sPsPAPPPPPPPPPPRR)

   Ion         m/z   Intensity
   y6++      360.50       20.8
   a1+       369.25       15.6
   b1+       397.19       37.9
   y7++      408.62       20.8
   b5++      415.03       14.9
   y8++      457.62       37.4
   b6++      463.61       24.7
   y9++      506.04       39.4
   b7++      512.39       21.7
   y4+       525.41       14.5
   a8++      546.33       14.9
   y10++     554.60       68.6
   b8++      560.57       32.0
   y11++     603.17       39.6
   b9++      609.01       17.8
   y5+       622.43       26.0
   a3+       633.18       21.2
   a10++     643.65       25.0
   y12++     651.68      162.5
   b10++     657.67       20.4
   b3+       660.91       18.5
   y13++     686.61       31.5
   a11++     692.32       33.4
   y6+       719.49       19.1
   a4+       730.58       17.6
   y14++     735.74      106.8
   a5+       801.32       27.9
   y7+       816.57       29.6
   y15++     819.11       54.4
   b5+       829.35       86.4
   y16++     867.77       71.0
   y8+       913.54       22.4
   b6+       926.44       19.8
   y9+      1010.73       12.9
   b7+      1023.60       20.5
   b8+      1120.61       14.8
   b9+      1217.48       13.6
```

```
262. MORG-339_phosphopeptide_TMT_16%ACN_032017_OT.5616.5616.4 (sQEPISNDQkVsDDDkEk)

   Ion         m/z   Intensity
   y1+       376.33       22.6
   y3++      431.72       71.5
   y4++      489.43       57.7
   b10+++    529.05       15.2
   y5++      546.79       54.8
   b3+       574.38       33.0
   y6++      604.31       38.1
   b9++      615.27       54.4
   y10+++    654.10       17.2
   y7++      687.79       17.6
   a15+++    723.34       23.9
   y12+++    730.07       23.3
   b15+++    733.30       36.8
   y8++      737.20       63.5
   a5+       756.72       29.2
   b5+       784.49       23.7
   b10++     793.75       27.2
   y14+++    796.51       37.3
   y15+++    828.98       27.3
   a6+       843.21       32.1
   b16+++    851.74       19.3
   y3+       863.07       22.1
   a17+++    886.17       18.8
   b17+++    895.14       21.1
   b12++     926.72       20.7
   a13++     969.54       24.8
   b13++     983.82       43.7
   b7+       985.51       15.8
   b14++    1041.68       56.5
   b15++    1099.31       54.7
   b8+      1100.49       62.1
```

```
263. MORG-339_phosphopeptide_TMT_14%ACN_032017_OT.3081.3081.3 (sRsPDIQAk)

   Ion         m/z   Intensity
   b3++      320.78       24.2
   y1+       376.30       15.7
   b5++      426.79       57.0
   y2+       447.37       27.0
   y6++      450.86       29.1
   a6++      469.35       22.9
   b2+       473.55       16.5
   b6++      483.38       34.2
   a7++      533.16        8.1
   y7++      534.02       10.4
   b7++      547.42       33.3
   y3+       575.41       30.9
   b8++      582.88       17.8
   b3+       640.12        9.4
   y4+       688.51       40.6
   y6+       900.59       22.4
```

```
264. MORG-339_phosphopeptide_TMT_18%ACN_032017_OT.12098.12098.2 (sSGTSyPDVLk)

   Ion         m/z   Intensity
   y1+       376.38       13.5
   y3+       588.44       23.9
   y5+       800.49       31.3
   b6+       892.30       19.4
   b7+       989.48       10.9
   b8+      1104.50       14.8
   b10+     1316.64       13.9
```

```
265. MORG-339_phosphopeptide_TMT_25%ACN_032017_OT.19535.19535.3 (kIVETINsDsDSEFGIPk)

   Ion         m/z   Intensity
   y1+       376.38       16.2
   b5++      400.53       16.5
   y2+       473.34       73.9
   y3+       586.49       15.8
   y4+       643.47       37.8
   y9++      644.48       23.3
   y5+       790.60       37.5
   a6+       885.11       14.7
   a13++     890.17       16.7
   y13++     899.00       22.9
   b13++     904.48       96.0
   b6+       913.85       25.7
   y6+       919.53       32.0
   y7+      1006.56       36.0
   y15++    1014.34       19.4
   y16++    1064.39       15.1
   y17++    1120.17       33.5
   y8+      1121.37       36.3
   b8+      1194.47       14.8
   y9+      1288.42       21.0
```

```
266. MORG-339_phosphopeptide_TMT_18%ACN_032017_OT.2975.2975.3 (hAAPAPSAkssPSk)

   Ion         m/z   Intensity
   b2+       438.35        9.9
   b3+       509.27       22.5
   y3+       560.44       31.8
   b9++      645.42       15.8
   a5+       649.82       13.0
   b5+       677.48       22.5
   y8++      705.41       31.8
   a10++     714.98       32.1
   y9++      753.85       22.9
   y10++     789.52       19.3
   a11++     797.90       12.4
   y11++     837.95       18.3
   y12++     873.82       15.6
   y5+       894.71       12.4
   y13++     909.25       16.5
```

```
267. MORG-339_phosphopeptide_TMT_10%ACN_032017_OT.14365.14365.3 (eAPAGGSGDGLSsPsPLEYSPR)

   Ion         m/z   Intensity
   y2+       272.17       24.4
   b1+       359.23       33.0
   b2+       430.30       33.2
   y7++      431.32       17.9
   b10++     514.68       36.8
   y4+       522.28       45.1
   y9++      563.48       67.8
   b11++     571.47       59.4
   b4+       598.31       44.3
   b12++     614.97       35.6
   y10++     646.95       28.7
   y5+       651.41       29.0
   b5+       655.45       23.1
   y11++     690.53       49.1
   b6+       712.45       27.0
   a14++     732.50       31.7
   b14++     746.96       28.8
   y13++     775.59       40.5
   b7+       799.57       27.4
   a15++     816.33       25.5
   a8+       827.95       19.9
   b15++     830.08       22.6
   b8+       856.43       19.6
   y7+       861.67       26.0
   y17++     933.69       20.0
   b17++     935.72       19.6
   y18++     962.08       22.4
   b9+       971.54       34.0
   b18++     999.73       24.9
   a10+     1000.59       36.7
   b10+     1028.52       35.6
   y20++    1046.22       19.4
   a19++    1067.29       25.7
   b19++    1081.79       40.8
   b20++    1124.84       21.5
   y9+      1125.53       44.6
   b11+     1141.58       52.8
   b12+     1228.65       33.2
   y10+     1292.59       35.1
   y11+     1379.60       34.6
   b13+     1395.62       24.4
```

```
268. MORG-339_phosphopeptide_TMT_6%ACN_032017_OT.2877.2877.4 (tDRGTDDsPkNSQEDLQDR)

   Ion         m/z   Intensity
   y7+++     301.73       12.6
   b6++      438.32       48.7
   b2+       446.50       19.4
   b7++      495.93       74.4
   y4+       531.33       42.7
   b10+++    538.00       22.9
   a11+++    566.62       12.7
   b11+++    576.12       32.5
   b8++      579.20       23.6
   b12+++    605.29       10.9
   a4+       631.75       25.7
   y5+       646.50       38.0
   b13+++    647.57       24.9
   y14+++    652.80       20.6
   b4+       659.82       61.0
   b14+++    690.79       19.0
   b15+++    729.21       49.0
   y10++     731.35       19.6
   a5+       732.14       13.2
   a16+++    757.57       25.4
   b16+++    766.97       14.3
   y6+       775.40       18.6
   y11++     780.17      120.0
   b10++     806.60       19.3
   b17+++    809.40       31.9
   a6+       846.96       15.2
   b18+++    848.02       54.2
   b11++     863.62       51.4
   y7+       903.36       12.5
   y13++     921.26       38.0
   b7+       990.49       19.6
   y15++    1028.62       13.3
   y16++    1057.90       11.3
   y9+      1104.54       16.4
```

```
269. MORG-339_phosphopeptide_TMT_6%ACN_032017_OT.10264.10264.4 (qADVADQQTTELPAENGETENQSPAsEEEk)

   Ion         m/z   Intensity
   y1+       376.30       31.4
   a12+++    500.97       30.5
   y2+       505.41       33.4
   b8++      543.46       38.0
   b3+       544.46       44.2
   y7++      549.98       52.6
   b9++      593.67       82.4
   y3+       634.48       35.9
   b4+       643.40       44.1
   b10++     644.31       32.3
   a17+++    657.17       19.6
   a18+++    700.09      127.1
   b11++     709.07       64.3
   b18+++    709.91       36.1
   b5+       714.48       60.5
   y17+++    720.17       23.6
   y4+       763.49       29.0
   b12++     765.66       91.3
   y11++     779.14       24.0
   b13++     814.32       19.7
   b6+       829.51       59.0
   y13++     893.70       22.7
   b15++     914.06       63.5
   a24+++    919.23       33.2
   y14++     922.74       62.9
   a7+       929.64       32.4
   y5+       930.50       26.0
   b7+       957.49       41.5
   y6+      1001.52       23.3
   y17++    1079.75       80.6
   y27+++   1080.63       55.3
   b8+      1085.59       46.2
   b28+++   1093.48       19.4
   y7+      1098.49       42.0
   b29+++   1136.70       35.7
   a9+      1158.47       25.4
   y8+      1185.31      113.7
   b9+      1186.27       97.0
   b22++    1300.44       59.5
   b11+     1416.70       36.2
   b12+     1529.78       50.4
```

```
270. MORG-339_phosphopeptide_TMT_10%ACN_032017_OT.1342.1342.4 (hDtPDTsPPRkAR)

   Ion         m/z   Intensity
   a5+++     283.32       22.2
   y6+++     318.73       30.3
   b3++      332.25       28.1
   b1+       367.19       12.4
   y8+++     408.07       22.2
   y5++      428.86       11.9
   b5++      438.35       31.5
   b9+++     446.17       42.2
   a6++      474.28       14.6
   y6++      477.47       38.2
   y10+++    478.73       68.2
   b2+       482.41       24.8
   b6++      488.80       13.5
   a10+++    489.54       39.5
   b10+++    498.71       15.5
   y7++      560.98       18.8
   b7++      572.32       10.6
   y12+++    577.38        8.6
   y3+       603.40       12.4
   y8++      611.54       35.1
   b12+++    641.62       12.3
   a9++      655.17        8.4
   b3+       663.27       14.8
   y9++      668.62       28.1
   b9++      669.42       14.9
   y10++     717.60       43.3
   y4+       759.43        8.7
   b5+       875.36       22.7
```

```
271. MORG-339_phosphopeptide_TMT_10%ACN_032017_OT.2100.2100.3 (sTSATDTHHVELAR)

   Ion         m/z   Intensity
   y3+       359.26       23.5
   y6++      362.78       28.7
   b1+       397.11       14.1
   a6++      422.36       13.7
   y7++      431.33       23.2
   a7++      473.05       20.4
   y8++      481.94       75.1
   y4+       488.30       32.3
   y9++      539.39       14.4
   a8++      541.74       30.8
   b8++      555.79       31.7
   a3+       556.88       14.0
   b3+       585.54       74.1
   y5+       587.40      101.4
   y10++     589.55       47.9
   b9++      624.54       30.5
   y11++     625.51       13.2
   b4+       656.32       15.0
   a10++     659.89       19.2
   y12++     669.10       14.7
   b10++     674.04       32.9
   y13++     719.51       32.1
   y6+       724.45       68.7
   b11++     738.41       26.2
   b12++     795.16       22.2
   b13++     830.46       27.0
   y7+       861.56       16.7
   b6+       872.39       56.3
   b7+       973.46       20.7
   b8+      1110.46       18.4
```

```
272. MORG-339_phosphopeptide_TMT_12%ACN_032017_OT.7068.7068.3 (akPVVsDDDsEEEQEEDR)

   Ion         m/z   Intensity
   b2++      329.78       14.5
   b4++      427.96       14.0
   b5++      477.47       14.2
   y4+       548.29       10.3
   b7++      618.44       14.9
   b2+       658.58       14.8
   y5+       676.09       24.2
   a9++      719.27       10.2
   b9++      733.31       20.1
   b3+       755.40        9.4
   y6+       805.40       13.9
   b10++     816.50       10.9
   a4+       826.69       10.5
   b4+       854.14       14.6
   y13++     871.68       13.4
   b11++     881.40       60.4
   y7+       934.37       13.1
   b12++     946.07       11.9
   b5+       953.86        9.4
   a13++     996.55       10.1
   b13++    1010.15       11.7
   y8+      1063.44       16.1
   b14++    1074.55       10.7
   b16++    1203.47        9.7
   y9+      1230.42       22.0
   a17++    1247.43       12.4
   b17++    1261.00       15.0
   y10+     1345.47       16.6
   y11+     1460.46       15.5
   y12+     1575.52       10.6
```

```
273. MORG-339_phosphopeptide_TMT_8%ACN_032017_OT.5701.5701.2 (sLsDSESDDSk)

   Ion         m/z   Intensity
   y2+       463.32       22.7
   y3+       578.46       16.2
   y4+       693.40        8.6
   b4+       712.36       15.9
   a5+       771.28        9.8
   y5+       780.36       16.8
   b5+       798.94       11.5
   b6+       928.43       14.6
   y7+       996.53       29.3
   y8+      1111.48       16.4
   b8+      1130.57       18.4
   b9+      1245.58       21.4
   y9+      1278.32       12.8
   y10+     1391.57        7.3
```

```
274. MORG-339_phosphopeptide_TMT_6%ACN_032017_OT.6700.6700.3 (sEtPPHWR)

   Ion         m/z   Intensity
   y1+       175.10        7.8
   y3++      249.72       22.1
   y4++      298.29       21.6
   a3++      339.79       12.9
   y5++      346.83      104.0
   b3++      353.86        9.3
   y2+       361.20       19.7
   a4++      388.37       37.0
   b1+       397.17       18.5
   y6++      437.27       17.5
   y3+       498.27       23.1
   y7++      501.84       13.4
   a6++      505.73       16.5
   b6++      519.87       18.4
   b2+       526.27       22.1
   y4+       595.33       30.0
   a7++      598.60       10.7
   b7++      612.92       13.1
   y5+       692.41       14.5
   b3+       707.30       14.9
   b4+       804.26        7.9
```

```
275. MORG-339_phosphopeptide_TMT_18%ACN_032017_OT.15621.15621.2 (gIsPIVFDR)

   Ion         m/z   Intensity
   b1+       287.23       10.8
   y2+       290.24        9.5
   a2+       372.27        8.4
   b2+       400.36       12.7
   y3+       437.22       31.7
   y4+       536.37       36.2
   b3+       567.36       31.7
   y5+       648.92       14.6
   y6+       746.42       96.0
   a5+       749.44       17.4
   b5+       777.49       29.8
   a6+       848.36       11.8
   b6+       876.45       38.8
   y7+       913.41       17.2
   b7+      1023.45       16.1
   b8+      1138.61       42.0
```

```
276. MORG-339_phosphopeptide_TMT_10%ACN_032017_OT.5966.5966.3 (vDHGAEIItQSPSR)

   Ion         m/z   Intensity
   b3++      291.26       13.4
   b4++      319.57       19.8
   a5++      341.21       12.6
   b5++      355.31       19.0
   y3+       359.28       21.5
   y6++      377.99       16.4
   b6++      419.88       52.3
   a7++      462.31       29.4
   b7++      476.36       62.1
   y8++      491.30       11.4
   a8++      518.87       17.3
   b8++      532.93       44.5
   a3+       553.76       14.4
   y5+       574.42       81.2
   b3+       581.02       36.9
   y10++     591.33       28.8
   b4+       638.36       19.4
   b5+       709.16       13.4
   a11++     716.58       10.5
   y6+       755.33       48.8
   b6+       838.39       12.9
   y7+       868.40       36.6
   b7+       951.55       16.9
   y8+       981.56       22.3
```

```
277. MORG-339_phosphopeptide_TMT_4%ACN_032017_OT.5693.5693.3 (eRDHsPtPSVFNSDEER)

   Ion         m/z   Intensity
   y2+       304.13        8.4
   b3++      315.81        6.3
   y3+       433.26       10.8
   b5++      467.48        6.2
   y10++     590.44       14.3
   b7++      606.75        6.2
   b3+       630.47       15.2
   y5+       635.31        7.5
   y11++     680.54       11.6
   a9++      685.21        6.9
   b9++      698.99       14.4
   y12++     729.39       19.4
   a4+       739.26       14.1
   b10++     748.73       20.6
   y6+       749.46       17.6
   y13++     812.92        7.4
   y14++     881.31       12.0
   y7+       896.35       16.3
   b13++     921.99        6.9
   b14++     979.84       11.9
   y8+       995.46       12.8
   y16++    1017.00        6.1
   b15++    1044.60       13.1
   y9+      1082.45        6.5
   b16++    1109.25       11.3
   y10+     1179.52       39.9
   b7+      1212.47        6.8
   y11+     1360.60        5.9
   y12+     1457.62       11.8
   y14+     1761.77        6.1
```

```
278. MORG-339_phosphopeptide_TMT_25%ACN_032017_OT.13186.13186.4 (aSIGNIALSPsPVRksPR)

   Ion         m/z   Intensity
   y2+       272.17       16.3
   a1+       273.10       12.8
   b6++      393.21       35.5
   a7++      415.20       26.3
   b7++      428.72       43.4
   y8+++     439.00       19.8
   y9+++     471.65       38.6
   b8++      485.23       30.3
   y10+++    500.63       21.4
   b3+       501.34       16.3
   b13+++    506.54       14.0
   b9++      528.95       37.2
   y11+++    538.17       20.5
   b4+       558.44       18.5
   y12+++    562.09       32.8
   y7++      575.01       25.1
   b10++     577.72       17.2
   y13+++    599.39       41.8
   y14+++    637.62       16.0
   y8++      658.57       24.1
   b11++     661.06       18.5
   b5+       672.52       53.4
   b15+++    677.77       21.7
   y16+++    694.51       19.6
   y9++      707.10       67.0
   b12++     709.43       19.8
   a13++     744.49       15.1
   y10++     750.50       56.0
   b13++     759.12       14.5
   b6+       785.51       28.6
   y11++     807.07       65.9
   b14++     836.82       16.2
   y12++     842.62       56.1
   b7+       856.60       33.8
   y13++     899.12       13.1
   b8+       969.71       20.6
   b9+      1056.68       23.8
```

```
279. MORG-339_phosphopeptide_TMT_30%ACN_032017_OT.10337.10337.3 (kQPPVsPGTALVGSQk)

   Ion         m/z   Intensity
   b1+       358.31       21.8
   y1+       376.42       17.4
   a9++      587.59       19.9
   y3+       591.42       20.8
   y10++     593.93       23.4
   y4+       648.46       48.2
   y11++     677.56       18.1
   b11++     693.56       21.1
   a12++     729.06       24.9
   b12++     743.27       40.2
   y5+       747.61       43.5
   a5+       751.38       24.0
   a13++     757.04       33.5
   b13++     771.40      107.6
   y13++     775.20       36.4
   b5+       779.09       22.1
   a14++     801.05       30.2
   b14++     815.05       28.7
   y14++     824.16       36.0
   y6+       860.57       33.8
   b15++     879.18       25.3
   y7+       931.62       30.5
   b6+       946.98       18.0
   y10+     1186.72       48.6
```

```
280. MORG-339_phosphopeptide_TMT_18%ACN_032017_OT.7100.7100.3 (rLPSSPAsPsPk)

   Ion         m/z   Intensity
   b4++      342.17       11.0
   y4++      369.27       16.3
   a5++      371.96       10.9
   y1+       376.45       13.5
   b1+       385.94       28.0
   b6++      434.36       12.5
   b7++      469.94       19.8
   y2+       473.40       50.0
   b2+       499.44       14.9
   y7++      536.84       16.4
   a8++      539.56       13.5
   b8++      553.21       19.3
   y8++      580.43       13.7
   a9++      588.11       38.4
   b3+       596.90       35.1
   b9++      601.36       20.4
   y9++      623.48       13.9
   y3+       640.47       19.1
   a4+       655.61       11.5
   y10++     672.15       14.4
   b4+       683.80        9.2
   b10++     685.32       11.3
   a11++     720.11       14.5
   y11++     728.91       13.3
   b11++     733.76        9.3
   y4+       737.39       37.5
   b5+       770.50       12.3
   b7+       938.51        9.8
   y7+      1072.50       16.7
   b8+      1105.52       11.7
```

```
281. MORG-339_phosphopeptide_TMT_4%ACN_032017_OT.7191.7191.3 (eGEGIEEcsGTEVkEDPESR)

   Ion         m/z   Intensity
   y2+       262.22       13.6
   y4+       488.26       52.0
   y7++      545.43       18.4
   y8++      595.00       19.3
   b4+       602.33       14.0
   y5+       603.39       12.9
   y9++      659.65       14.4
   a11++     715.50       14.4
   b11++     729.95        8.5
   y11++     738.66       34.1
   b6+       844.34       19.1
   y13++     901.84       15.8
   y14++     966.91       10.9
   b7+       973.51       17.8
   b14++    1022.33       14.5
   b15++    1086.87       29.6
   y16++    1087.59       18.0
   a8+      1105.74       16.5
   y17++    1115.91       11.3
   b8+      1133.65       14.3
   b16++    1144.76       90.2
   y19++    1208.99       24.2
   b18++    1257.79       26.9
   b19++    1301.43       15.8
```

```
282. MORG-339_phosphopeptide_TMT_14%ACN_032017_OT.7445.7445.3 (sLtRsPPAIR)

   Ion         m/z   Intensity
   y1+       175.15       12.5
   y5++      277.23       22.7
   a3++      292.20        7.4
   b3++      306.19        8.0
   b1+       317.23       15.7
   y3+       359.30        7.9
   y6++      360.52       13.9
   a4++      369.88       44.4
   b4++      384.24       25.5
   y7++      438.86       24.6
   a5++      453.67       13.6
   y4+       456.27       20.1
   b5++      467.58       46.8
   a7++      550.72        8.2
   y5+       553.37      118.0
   b7++      564.81        8.2
   b8++      600.40       19.8
   b3+       611.44        8.1
   a9++      643.04       14.0
   b9++      656.95       25.8
   y6+       720.36       25.5
```

```
283. MORG-339_phosphopeptide_TMT_14%ACN_032017_OT.10921.10921.4 (nLAkPGVTSTsDsEDEDDQEGEk)

   Ion         m/z   Intensity
   y1+       376.38       16.8
   b4++      443.37       31.1
   y2+       505.47       14.2
   y7++      525.29       13.5
   y3+       562.40       34.6
   b7++      570.13       15.3
   b8++      620.45       24.9
   y9++      647.23       12.1
   y4+       691.43       16.3
   b10++     714.35       16.6
   a16+++    741.62       11.7
   b16+++    751.12       11.1
   y18+++    773.39       18.8
   a11++     783.56       10.3
   b17+++    789.30       24.3
   y11++     795.70       22.8
   b11++     797.61       16.5
   y19+++    805.58       28.5
   b4+       885.86       19.4
   a20+++    903.88       12.6
   b20+++    913.29       39.8
   b21+++    932.30       14.5
   y6+       934.76       10.6
   y13++     936.72       22.5
   a5+       954.91       14.0
   a22+++    966.15       10.8
   b22+++    974.90       13.2
   b14++    1003.77       10.4
   a6+      1011.82       17.7
   b15++    1061.12       17.1
   b16++    1125.74       19.6
   b17++    1183.08       15.0
   b18++    1240.63       19.2
   b19++    1305.02       12.1
```

```
284. MORG-339_phosphopeptide_TMT_4%ACN_032017_OT.6836.6836.3 (sPsPSPTPEAk)

   Ion         m/z   Intensity
   a3++      317.25        5.3
   y4++      337.25       38.1
   y1+       376.31       11.8
   b5++      423.16        5.6
   y6++      436.43       23.1
   y2+       447.33       17.1
   a2+       466.33        3.9
   b7++      522.24       23.2
   y8++      528.41       12.8
   a8++      556.28       11.2
   y9++      611.40        3.7
   b9++      635.53       14.6
   a10++     656.83        4.0
   y10++     660.58        6.4
   b10++     670.88       13.3
   y4+       673.45       18.7
   b4+       758.47        4.0
   b5+       845.36       19.2
   y6+       871.52        4.5
   b7+      1043.42       28.5
```

```
285. MORG-339_phosphopeptide_TMT_12%ACN_032017_OT.2309.2309.3 (qGSGREsPSLVSR)

   Ion         m/z   Intensity
   y2+       262.16       10.5
   b1+       358.36       25.0
   y3+       361.29       17.4
   a6++      408.91       15.4
   b6++      422.81       26.1
   y4+       474.47       21.2
   a7++      492.56       15.3
   b3+       502.37       14.1
   b7++      506.34       51.9
   a4+       531.58       45.4
   a8++      540.46       22.9
   y5+       561.36        9.9
   b9++      598.35       10.5
   y11++     627.52       17.3
   a10++     640.41       27.4
   y12++     656.66       18.7
   y6+       658.43       98.4
   a11++     690.16       19.7
   b11++     704.47       25.7
   b12++     747.80       12.6
   y7+       825.43       13.1
```

```
286. MORG-339_phosphopeptide_TMT_14%ACN_032017_OT.6029.6029.3 (rAGDVLEDsPk)

   Ion         m/z   Intensity
   b4++      315.33       33.0
   b5++      364.84       16.3
   y1+       376.38       14.6
   a6++      407.31       11.8
   b6++      421.44       22.5
   a7++      472.19       17.0
   y2+       473.36       54.7
   a3+       486.07       36.1
   y6++      499.56        8.4
   b3+       514.29       10.3
   b8++      543.30      161.2
   y7++      548.34       11.1
   y8++      606.27        9.2
   b9++      626.68       15.9
   b4+       629.61       18.9
   y9++      635.22        9.7
   y3+       640.38       27.0
   b5+       728.63       11.6
   y4+       755.42       13.5
   b6+       841.52       10.1
   b7+       970.74        8.9
   y6+       997.53       10.4
   y7+      1096.60       15.9
```

```
287. MORG-339_phosphopeptide_TMT_22%ACN_032017_OT.9607.9607.3 (sRHSsIsPSTLTLk)

   Ion         m/z   Intensity
   y1+       376.34       19.7
   b2+       473.39       15.5
   a6++      475.30       16.2
   y2+       489.41       20.2
   y7++      494.94       18.1
   b7++      572.73       17.8
   y3+       590.52       24.6
   b3+       610.70       18.8
   a9++      650.74       19.0
   b9++      664.88       15.4
   a4+       669.36       20.7
   b4+       697.12       88.9
   a10++     701.09       20.8
   y4+       703.54       19.3
   a11++     757.94       15.8
   y11++     762.10       15.9
   b11++     771.65       17.2
   y5+       804.54       26.6
   y12++     830.15       18.4
   a13++     864.82       17.6
   b13++     878.78       15.9
   y7+       988.63       44.3
   y8+      1155.65       22.6
```

```
288. MORG-339_phosphopeptide_TMT_18%ACN_032017_OT.13173.13173.2 (tLSIsTQDLSPR)

   Ion         m/z   Intensity
   b1+       331.10       11.6
   y3+       359.15       20.1
   b2+       444.41       17.3
   y4+       472.37       46.5
   b3+       531.38       16.9
   y5+       587.42       18.7
   b4+       644.50       19.0
   a5+       783.28       16.1
   b6+       912.83       15.3
   y8+       983.47       20.9
   b7+      1040.56       12.3
   b8+      1155.67       35.4
   y10+     1183.61       16.1
   b9+      1268.67       25.8
   b10+     1355.61       16.5
```

```
289. MORG-339_phosphopeptide_TMT_4%ACN_032017_OT.3660.3660.3 (aEGAGTEEEGtPk)

   Ion         m/z   Intensity
   y2++      237.27       12.5
   b1+       301.21        7.6
   y3++      327.76        6.4
   y4++      356.28       14.5
   y1+       376.31       16.0
   y5++      420.88       28.2
   b7++      423.46        9.8
   b2+       430.26       11.1
   a8++      473.38       56.4
   y6++      485.24       15.3
   b8++      487.72       23.8
   a4+       530.75        6.4
   a9++      538.41        9.3
   y7++      549.85       14.5
   b9++      552.22       33.8
   b4+       558.52       16.1
   a10++     566.73       32.8
   b10++     581.08       30.1
   a5+       587.27       12.8
   b5+       615.16       11.8
   y9++      629.01        7.9
   y3+       654.35       18.7
   b11++     671.01       15.6
   y4+       711.42       32.8
   b6+       716.49       21.7
   b12++     719.72        9.3
   y5+       840.42       12.2
   b7+       845.50       26.0
   b8+       974.51       30.8
   b9+      1103.57       17.0
   b10+     1160.57       10.6
   b11+     1341.33        9.2
```

```
290. MORG-339_phosphopeptide_TMT_18%ACN_032017_OT.17964.17964.3 (sIHEVDISNLEAALR)

   Ion         m/z   Intensity
   b3++      324.29       13.1
   y6++      336.21        9.5
   b4++      388.84       17.3
   b1+       397.14       42.3
   a5++      424.13       12.6
   y4+       430.46       11.9
   b5++      438.30       14.0
   b6++      495.85       30.8
   a7++      538.26       18.6
   y10++     551.45        7.8
   b7++      552.55       17.7
   y5+       559.42       22.5
   b8++      596.20       13.7
   y11++     601.10       12.2
   b3+       647.13       16.8
   y6+       672.59       13.7
   a10++     695.83        9.1
   b10++     709.37       16.0
   b11++     773.86       10.4
   y7+       786.67       13.9
   y14++     790.78        7.3
   b12++     809.70       15.4
   b13++     845.00       10.3
   y8+       873.53       29.0
   b5+       875.56       11.6
   b14++     901.60       12.2
   y9+       986.71       12.8
   b6+       990.70        9.0
   y10+     1101.62       16.8
   b7+      1103.53       11.2
   y11+     1200.69       10.9
```

```
291. MORG-339_phosphopeptide_TMT_20%ACN_032017_OT.12296.12296.3 (aEAPPLQREDsGTFSLGk)

   Ion         m/z   Intensity
   b2+       430.27       16.0
   y2+       433.36       19.5
   b3+       501.39       17.2
   b8++      546.56       21.6
   y4+       633.47       16.8
   b10++     669.07       43.5
   b11++     752.71       21.8
   y11++     753.76       13.2
   y5+       780.50       21.7
   b12++     781.35       42.8
   b13++     830.90        9.9
   y13++     874.26       10.1
   y6+       881.42       16.5
   a14++     891.29       18.6
   b14++     905.40       21.8
   y14++     922.56       18.3
   b7+       936.61       11.1
   y7+       938.59       28.6
   b15++     948.17       14.3
   y15++     971.34       20.1
   a16++     990.65       11.1
   y16++    1006.64       15.1
   b17++    1033.97       10.9
   y8+      1105.66       21.4
```

```
292. MORG-339_phosphopeptide_TMT_16%ACN_032017_OT.8739.8739.3 (rSsASVSLPPGtPEk)

   Ion         m/z   Intensity
   y1+       376.42       16.0
   b6++      449.32       17.9
   y6++      468.94       55.7
   y7++      517.93      132.6
   a8++      534.97       18.4
   b8++      549.13       24.8
   y8++      574.55       26.2
   b9++      597.78       21.4
   y3+       602.39       84.9
   y9++      617.95       24.2
   b3+       640.01       21.8
   b10++     646.09       30.5
   a11++     660.52       15.4
   y10++     667.62       32.6
   b11++     674.60       30.3
   a4+       683.36       13.6
   b12++     765.67       20.1
   y4+       783.00       13.3
   b5+       798.65       14.1
   y13++     830.29       16.2
   y5+       840.53       18.5
   b14++     878.67       13.8
   b6+       897.53       25.5
   y6+       937.51       38.2
   y7+      1034.48       45.6
   a8+      1069.54       15.2
   b8+      1097.51       82.1
```

```
293. MORG-339_phosphopeptide_TMT_22%ACN_032017_OT.3907.3907.3 (rLsTHSPFR)

   Ion         m/z   Intensity
   y1+       175.17       15.8
   b1++      193.70        8.8
   y2+       322.08        9.2
   b3++      333.71       12.0
   a4++      370.71       11.5
   y6++      372.65       10.3
   b4++      384.39        9.2
   b1+       386.34        8.8
   y3+       419.39       31.2
   b5++      452.94       33.6
   y7++      456.13       33.8
   a6++      482.46       12.5
   b6++      496.31       20.8
   b2+       499.36       11.4
   y4+       506.29       36.6
   a7++      531.08       10.5
   b7++      544.93       16.6
   a8++      604.33       18.6
   b8++      618.47       44.3
   y5+       643.36       32.1
   y6+       744.41       24.9
```

```
294. MORG-339_phosphopeptide_TMT_22%ACN_032017_OT.11680.11680.3 (dAsPPEPASPTIGLDkk)

   Ion         m/z   Intensity
   b1+       345.25       17.6
   y2++      367.37       27.6
   b2+       416.42       17.6
   y5++      509.99       14.5
   y6++      566.45       13.6
   b3+       583.29       18.2
   y8++      665.61       30.7
   y9++      709.19       21.8
   y2+       733.62       26.8
   b12++     737.06       13.3
   y10++     744.08       16.0
   b5+       777.39       15.7
   y11++     793.20       57.3
   y3+       848.45       22.0
   y12++     857.48       17.2
   a15++     865.49       16.2
   b6+       906.30       37.0
   y14++     954.72      168.6
   y5+      1018.75       27.4
   y15++    1038.00       18.6
   b16++    1058.36       17.1
   y16++    1073.85       15.7
   b8+      1074.54       15.1
   y8+      1329.91       17.4
   b15+     1757.94       20.5
```

```
295. MORG-339_phosphopeptide_TMT_4%ACN_032017_OT.6185.6185.3 (rLsPsAsPPR)

   Ion         m/z   Intensity
   y1+       175.09        9.0
   y2+       272.14        9.8
   b3++      333.63       40.2
   a4++      367.87       10.5
   y3+       369.27       74.0
   y6++      387.24       29.1
   y7++      436.26       38.0
   b5++      465.75       10.9
   a6++      486.79       74.3
   y4+       535.81      111.6
   b7++      584.72       21.5
   y5+       607.34       22.5
   b3+       666.43       18.7
   y6+       774.32       17.8
   y7+       871.30       40.7
```

```
296. MORG-339_phosphopeptide_TMT_10%ACN_032017_OT.18212.18212.3 (dEkkEEsEESDDDMGFGLFD)

   Ion         m/z   Intensity
   b4++      595.03       10.8
   b5++      659.27       11.7
   b6++      724.08       22.4
   y7+       786.46       23.6
   b3+       831.63       18.4
   b8++      871.87       14.7
   y8+       901.53       20.7
   a9++      922.16       10.9
   b9++      936.45       13.2
   b10++     980.43       17.1
   y9+      1016.35       17.5
   b12++    1095.16       34.7
   y10+     1131.80       10.5
   b13++    1152.84       29.7
   a14++    1203.64       11.9
   y11+     1218.44       27.6
   b15++    1246.60       19.4
   a16++    1306.17       10.9
   b16++    1320.29       17.2
   b17++    1348.90       12.8
   y19++    1372.83       17.6
   b18++    1405.64       17.6
```

```
297. MORG-339_phosphopeptide_TMT_10%ACN_032017_OT.8974.8974.3 (eALVEPASEsPRPALAR)

   Ion         m/z   Intensity
   y3+       359.29       10.0
   y7++      390.83       44.0
   y4+       430.24        9.9
   y8++      474.38       11.5
   y5+       527.42       24.5
   b3+       543.40       13.8
   b9++      578.26       13.0
   y10++     582.43       14.4
   y11++     617.46       27.8
   b4+       642.56       21.5
   a10++     648.23       11.5
   b10++     661.51       15.3
   y12++     666.50       30.5
   y6+       683.35       46.0
   a5+       742.99       10.8
   b5+       771.48       34.3
   a12++     774.60       11.8
   y7+       780.41       15.6
   b12++     788.33       29.3
   b6+       868.25        9.9
   b15++     928.98       12.1
   b7+       939.54       13.7
   b16++     964.55       22.9
   b9+      1155.62       12.5
```

```
298. MORG-339_phosphopeptide_TMT_4%ACN_032017_OT.3050.3050.3 (sRsYtPEYRR)

   Ion         m/z   Intensity
   y3++      247.79       18.3
   b2++      277.45        9.1
   y2+       331.15       11.4
   y5++      360.80       45.5
   b4++      442.59       12.8
   y6++      451.37       18.8
   y3+       494.31       17.6
   a2+       525.18        9.0
   y7++      532.75       17.6
   b2+       553.41       18.9
   a6++      567.26       10.8
   b6++      581.21       14.0
   b7++      645.63       14.9
   y9++      694.46       11.2
   y5+       720.42       10.7
   b9++      805.39       19.8
   b4+       883.45       14.0
   y7+      1064.43       14.1
   b7+      1290.47       11.7
```

```
299. MORG-339_phosphopeptide_TMT_20%ACN_032017_OT.8587.8587.4 (sESPtckSckSELPSSAPGGGGV)

   Ion         m/z   Intensity
   b7+++     443.25       59.5
   y7+       514.35       19.4
   y8+       601.44       11.8
   b10+++    645.44       15.4
   b7++      664.87       10.1
   b11+++    674.19       13.0
   y17+++    692.94       10.2
   b8++      708.75       18.4
   b12+++    717.23       40.1
   y14++     736.26       22.6
   y18+++    745.71       40.8
   b13+++    754.85       53.1
   y10+      785.43       38.4
   b14+++    787.37       19.8
   b9++      788.21       10.6
   a15+++    806.46       20.2
   b5+       811.28       13.5
   y15++     816.41       25.8
   a16+++    835.82       17.2
   b16+++    845.47       28.2
   y16++     859.69       18.9
   y21+++    867.84       11.3
   b17+++    869.08      161.2
   y11+      898.50       22.0
   b18+++    901.09       16.6
   a19+++    911.22       12.0
   b19+++    920.46       32.0
   b20+++    939.38       38.6
   b21+++    958.29       50.6
   a22+++    967.37       10.9
   b6+       971.28       10.7
   b22+++    977.39       60.2
   b13++    1131.76       10.6
```

```
300. MORG-339_phosphopeptide_TMT_14%ACN_032017_OT.8093.8093.3 (sHsLPNSLDYAQASER)

   Ion         m/z   Intensity
   b1+       317.47       14.5
   y3+       391.18       14.9
   a6++      459.00       13.3
   y4+       462.27       16.2
   b6++      473.41       19.8
   b7++      516.85       14.6
   y9++      527.25       13.6
   b8++      572.96       13.4
   y5+       590.50       20.6
   b3+       621.06       10.6
   b9++      631.05       34.3
   y6+       661.71       26.9
   y12++     676.18       64.0
   a10++     698.75        9.0
   b10++     712.42       18.7
   y13++     732.66       13.0
   b11++     747.95       18.9
   b12++     812.18       30.4
   y7+       824.49       18.0
   b13++     847.61       16.7
   b14++     891.22       11.7
   y8+       939.44       31.3
   a15++     941.45       11.8
   y9+      1052.56       14.9
   y10+     1139.56       14.6
```

```
301. MORG-339_phosphopeptide_TMT_18%ACN_032017_OT.10263.10263.3 (fSSQQASTkQSNAssDVEVEEk)

   Ion         m/z   Intensity
   y1+       376.35       17.4
   y2+       505.39       21.2
   y3+       634.49       27.9
   b4+       679.54       16.7
   b9++      712.66       32.2
   y4+       733.53       32.9
   b10++     776.44       34.3
   b5+       807.51       21.8
   b11++     820.24       31.5
   a12++     862.60       35.5
   b12++     877.07       36.9
   b6+       878.57       27.2
   b13++     912.86       35.5
   y6+       961.63       47.6
   b7+       965.57       21.4
   a14++     982.19       35.3
   b14++     996.29       21.3
   a8+      1038.27       16.6
   a15++    1065.46       15.4
   b8+      1066.68       33.8
   y15++    1135.24       18.5
   b16++    1136.84       24.7
   a17++    1172.32       15.3
   y16++    1178.78       22.3
   b17++    1186.79       29.5
   y17++    1214.44       23.4
   y8+      1243.51       28.8
   b18++    1251.25       34.7
   b19++    1300.90       24.5
   b20++    1365.43       23.7
   y20++    1385.92       15.7
   y21++    1429.04       16.5
   b21++    1429.98       30.9
   y10+     1481.51       21.0
   y11+     1595.71       15.3
   y12+     1682.73       18.6
```

```
302. MORG-339_phosphopeptide_TMT_20%ACN_032017_OT.2666.2666.3 (aDREQsPVSkTk)

   Ion         m/z   Intensity
   b5++      415.27        8.8
   b2+       416.28       11.8
   y3++      417.73       10.1
   y4++      461.56        9.9
   y2+       477.47       11.0
   y6++      559.56       44.4
   b3+       572.28        8.6
   b8++      596.87        9.4
   b9++      640.14        9.1
   y7++      642.38        9.3
   a5+       801.72       13.7
   a10++     804.52        8.8
   b10++     818.77        8.3
   b5+       829.53        9.7
   y3+       834.65       12.4
   y10++     849.65       18.2
   y4+       921.69       14.2
   y6+      1117.75       12.9
```

```
303. MORG-339_phosphopeptide_TMT_4%ACN_032017_OT.5561.5561.3 (dGQVINETsQHHDDLE)

   Ion         m/z   Intensity
   b5++      371.76        5.2
   y6++      383.18        4.0
   b7++      493.25        7.7
   a3+       502.33        3.9
   y8++      530.55        6.6
   b8++      543.92        6.3
   a4+       601.43        9.7
   b9++      626.82        6.0
   y5+       628.08       11.5
   b4+       629.18       24.5
   a10++     677.44       42.4
   b10++     691.62       13.7
   y11++     703.03        7.6
   b5+       742.51        8.0
   a11++     745.86        5.8
   b11++     759.87        8.2
   a6+       828.58        5.0
   b6+       856.50        5.1
   b13++     885.81        8.2
   b14++     943.48       12.0
   a15++     985.56        6.9
   b8+      1086.43        6.3
   a9+      1226.06        4.2
```

```
304. MORG-339_phosphopeptide_TMT_12%ACN_032017_OT.3242.3242.4 (eNSAAPSPGRPQsASPAk)

   Ion         m/z   Intensity
   y3++      272.81       10.2
   y4++      316.39       13.7
   y5++      351.71       12.8
   y1+       376.29        9.2
   b6++      400.12        8.9
   a7++      429.78       12.0
   b7++      443.85       19.6
   y2+       447.38       10.4
   a12+++    465.58        8.3
   y11+++    468.95       16.3
   b2+       473.48        7.3
   b12+++    474.41        9.5
   b8++      491.92       12.7
   y12+++    497.92       12.9
   y7++      498.83        8.0
   a9++      506.99       13.7
   a13+++    521.38       10.9
   y13+++    530.45       72.4
   a3+       532.17       11.9
   a14+++    544.41       38.9
   y8++      547.88       21.4
   y14+++    554.03       32.5
   b3+       560.20       19.4
   a4+       603.01        9.8
   y16+++    606.37        9.4
   y4+       631.40       20.7
   a11++     633.42        8.4
   b17+++    639.27        9.4
   b11++     647.17        8.8
   y10++     654.78       12.2
   y11++     702.56       59.0
   y12++     746.54       15.8
   y13++     794.84       35.2
   a14++     816.11        8.7
   y14++     830.60        9.7
   b7+       886.49        9.5
```

```
305. MORG-339_phosphopeptide_TMT_18%ACN_032017_OT.14580.14580.2 (sSsLGDLLR)

   Ion         m/z   Intensity
   y2+       288.22        9.9
   b1+       317.19       12.0
   y3+       401.27       41.6
   b2+       404.30       12.1
   y4+       516.39       14.2
   a3+       543.01        8.2
   b3+       570.91       63.1
   y5+       573.36       19.2
   y6+       686.41       35.8
   b5+       741.36       12.6
   y7+       853.52        8.7
   b6+       856.42       39.7
   y8+       940.51        8.2
   b7+       969.48       19.9
   b8+      1082.60       20.2
```

```
306. MORG-339_phosphopeptide_TMT_16%ACN_032017_OT.8855.8855.3 (kVLsDsEEEEk)

   Ion         m/z   Intensity
   y3++      318.17        9.0
   y1+       376.36       12.8
   y2+       505.38       17.4
   y7++      587.47        9.6
   a8++      625.65       12.0
   y3+       634.36       17.9
   b8++      638.96       10.8
   y8++      671.61       11.9
   a9++      689.60       49.6
   b9++      704.08       22.7
   a10++     753.97       20.8
   y4+       763.39       17.4
   a5+       824.70        8.7
   b5+       852.42       10.2
   y5+       892.54       18.3
   y6+      1059.44       21.9
   y7+      1174.56       14.7
   y9+      1454.61        8.9
```

```
307. MORG-339_phosphopeptide_TMT_14%ACN_032017_OT.6040.6040.3 (sFDYNYRR)

   Ion         m/z   Intensity
   y1+       175.15       13.9
   y3++      247.51       12.3
   y4++      304.76       15.7
   y2+       331.26        9.0
   a1+       369.08       12.3
   y5++      386.37      146.3
   a4++      397.27       23.8
   b4++      411.85        9.4
   y6++      443.94       24.7
   b5++      468.43       29.3
   y3+       494.31       10.2
   y7++      517.40       41.2
   b2+       544.34        9.4
   y4+       608.40       10.2
   a7++      614.48        7.3
   b7++      628.40       31.5
   b3+       659.25       83.1
   b4+       822.48       14.6
   b5+       936.38       12.1
```

```
308. MORG-339_phosphopeptide_TMT_20%ACN_032017_OT.8206.8206.4 (sPTENVNTPVGkDPGLAGRGDPk)

   Ion         m/z   Intensity
   b1+       397.23       34.4
   y2+       473.38       73.4
   b8++      576.45       21.1
   y10++     598.94      130.7
   y15+++    641.74       37.8
   y11++     656.69       17.4
   y16+++    675.29       21.6
   a16+++    706.14       19.1
   y17+++    713.38       29.0
   b4+       724.45       60.6
   a17+++    730.22       41.8
   b17+++    739.45       27.6
   y18+++    746.31       28.8
   a18+++    749.18      101.5
   b18+++    758.16       39.7
   y19+++    784.56       29.2
   b19+++    810.86       24.6
   a20+++    820.30       16.4
   y20+++    827.53       19.3
   b20+++    829.57       15.0
   y12++     835.42       47.4
   b5+       838.48       18.4
   y21+++    860.87       15.5
   y13++     863.76       21.0
   b21+++    867.75       55.5
   a12++     868.35       74.8
   b12++     881.78       36.1
   a22+++    890.73       80.3
   y22+++    893.53      112.5
   b22+++    899.73       38.4
   y14++     913.48       15.9
   y7+       929.78       15.4
   b6+       937.55       24.9
   b13++     939.68      138.6
   y15++     961.55       35.1
   a14++     973.66       25.0
   y16++    1012.25       17.6
   y17++    1069.45       18.3
   a17++    1094.46       15.1
   y10+     1196.80       21.8
```

```
309. MORG-339_phosphopeptide_TMT_10%ACN_032017_OT.6750.6750.3 (tAsDSDEQQWPEEkR)

   Ion         m/z   Intensity
   b1+       331.16       11.7
   y5++      444.54       18.8
   y6++      537.61       16.8
   y7++      601.48       11.9
   a4+       656.66       14.3
   y8++      665.69       29.3
   b4+       684.33       23.8
   a10++     715.67       15.2
   b10++     729.25       15.7
   y9++      730.30       54.3
   a5+       743.81       89.2
   a11++     763.94       30.0
   b5+       771.36       30.6
   y11++     831.23       54.9
   b12++     842.29       10.7
   b6+       886.44       21.7
   y5+       887.53       20.9
   y12++     888.69       37.0
   y13++     971.94       11.0
   y14++    1007.75       11.9
   b7+      1015.41       14.9
   b14++    1085.41       11.0
   b9+      1271.49       11.2
```

```
310. MORG-339_phosphopeptide_TMT_16%ACN_032017_OT.8364.8364.3 (lRLsPsPTSQR)

   Ion         m/z   Intensity
   y1+       175.02        8.1
   b2++      250.18       14.4
   a3++      292.79       14.0
   y2+       303.27        9.6
   b3++      306.80        7.5
   a4++      376.35       12.8
   y6++      377.95       12.1
   b4++      390.07       46.0
   a5++      424.33       39.8
   y7++      426.71       28.6
   a2+       471.04        9.4
   y4+       491.28       15.4
   y8++      510.63       15.0
   b6++      522.54       19.2
   b7++      570.42       14.9
   y5+       588.32       77.2
   b3+       612.31       23.0
   b9++      664.92       11.1
   b10++     728.65       13.8
   a4+       751.27        8.1
   y6+       755.42       19.3
   b4+       779.46       19.8
   y7+       852.35       51.0
   b5+       876.40        7.8
```

```
311. MORG-339_phosphopeptide_TMT_10%ACN_032017_OT.7640.7640.2 (sEGLSLER)

   Ion         m/z   Intensity
   b1+       397.18       14.6
   y3+       417.36       10.8
   a2+       498.18        7.3
   y4+       504.35       19.9
   b2+       526.25       12.6
   a3+       554.85        8.1
   b3+       583.42       20.2
   y5+       617.46       11.6
   a4+       668.62       13.4
   y6+       674.49       23.7
   b4+       696.28       21.9
   b5+       783.53        9.6
   y7+       803.45       21.8
   b7+      1025.47       22.8
```

```
312. MORG-339_phosphopeptide_TMT_25%ACN_032017_OT.14020.14020.4 (ykDIIRQPsEEEIIk)

   Ion         m/z   Intensity
   a4+++     317.32       18.4
   y1+       376.28       35.7
   b6+++     416.69       21.8
   b3++      433.21       50.9
   a7+++     450.11       20.9
   b7+++     459.48       54.8
   b4++      489.42       57.2
   y7++      578.80       20.3
   b10+++    590.53       29.3
   y3+       602.45       54.2
   a6++      610.68       17.3
   b6++      624.62       32.3
   y8++      627.53       47.0
   b11+++    633.73      376.6
   a12+++    667.18       18.6
   b12+++    676.77       77.1
   b7++      688.40       30.4
   a13+++    705.02       30.7
   b13+++    714.40       68.0
   y4+       731.49       44.0
   a14+++    742.73       18.5
   y14+++    746.37       18.3
   b14+++    752.09       35.3
   y10++     769.82       15.2
   y11++     825.62       30.3
   y5+       860.50       28.6
   y12++     882.75       41.8
   y6+       989.57       28.4
   b12++    1014.27       17.6
   b13++    1070.83       23.6
   y8+      1253.62       20.0
```

```
313. MORG-339_phosphopeptide_TMT_10%ACN_032017_OT.6000.6000.3 (qEDsGQNEkASDLETk)

   Ion         m/z   Intensity
   y1+       376.31       11.7
   b6++      477.30       18.1
   b3+       602.42       24.8
   y3+       606.46       11.6
   y4+       719.49       34.5
   a10++     799.28       18.8
   b10++     813.78       30.6
   b5+       826.39       24.2
   y5+       834.43       27.9
   b11++     856.78       14.7
   y11++     860.94       14.3
   y12++     889.28       14.1
   a12++     900.43       12.7
   b12++     914.58       42.1
   b13++     971.25       15.4
   y13++     973.32       30.4
   y7+       992.86       13.3
   y14++    1030.56       12.1
   b14++    1035.66       17.3
```

```
314. MORG-339_phosphopeptide_TMT_10%ACN_032017_OT.6248.6248.3 (sRTsVQTEDDQLIAGQSAR)

   Ion         m/z   Intensity
   a6++      470.94       17.7
   b6++      484.96       41.0
   y5+       518.36       43.3
   y6+       589.39       62.8
   b8++      599.87       34.7
   a9++      643.63       29.0
   b9++      657.58       59.2
   y7+       702.50       52.7
   b10++     715.17       62.4
   b4+       741.62       25.6
   a11++     764.70       62.5
   y14++     765.95       40.2
   a5+       812.56       21.5
   y15++     815.55       35.9
   a12++     821.60       32.4
   b12++     835.66       62.3
   b5+       840.42       20.1
   a13++     878.36       29.9
   b13++     892.23       62.8
   y16++     899.06       16.4
   b14++     927.74       51.3
   y9+       943.51       58.9
   b15++     956.23       18.6
   b16++    1019.80       18.6
   a7+      1041.53       18.6
   y10+     1058.56       52.0
   b18++    1099.30       21.4
   y11+     1173.61       35.1
   y12+     1302.67       24.2
   y13+     1403.68       29.9
   y14+     1531.78       21.1
```

```
315. MORG-339_phosphopeptide_TMT_8%ACN_032017_OT.4324.4324.4 (aEEGGEsEGDASEkDAk)

   Ion         m/z   Intensity
   a9+++     376.26       18.3
   a2+       401.79        7.8
   y7+++     403.22        9.0
   y2+       447.33       13.7
   y4++      460.35       16.2
   b12+++    476.36        6.6
   b7++      485.05       18.9
   y5++      524.80       18.2
   a8++      536.05        9.7
   b8++      549.94        7.4
   b3+       559.36       17.5
   y3+       562.37       23.5
   a9++      564.61        9.1
   y6++      568.51       24.2
   b9++      578.14       13.5
   a4+       587.91       17.7
   y12+++    602.41       20.8
   y7++      603.82       19.3
   b10++     635.64       14.7
   b14+++    639.18       26.1
   y14+++    640.34        6.0
   a5+       645.64        7.6
   y8++      661.77       18.3
   a15+++    668.36       15.6
   b11++     671.36       29.1
   b5+       673.43        7.6
   b15+++    677.49       20.3
   y9++      690.16       19.9
   a16+++    692.04        6.0
   b16+++    701.07       32.2
   b12++     714.90       19.1
   y10++     754.50        7.2
   a13++     765.58        6.0
   b13++     779.47        9.1
   b6+       802.39       13.5
   b14++     958.00        5.8
   b15++    1015.65        6.3
   b8+      1098.50        6.2
   b10+     1270.47        8.9
```

```
316. MORG-339_phosphopeptide_TMT_6%ACN_032017_OT.3884.3884.3 (sPGDEDDkDck)

   Ion         m/z   Intensity
   y1+       376.32        9.0
   b1+       397.15       33.0
   b5++      397.99       11.0
   a7++      499.63        8.7
   y4++      504.96       48.0
   y2+       536.39       35.4
   b3+       551.38       14.2
   y5++      562.27       38.5
   y6++      619.81       18.8
   a4+       637.94       15.3
   y3+       651.76       50.1
   b4+       666.41       41.3
   y7++      684.52       13.6
   b8++      691.59        9.4
   y8++      741.89       16.9
   b9++      749.38       33.6
   b5+       795.43       17.9
   y10++     819.06       90.5
   a6+       882.45        8.3
   b6+       910.30       20.6
   b7+      1025.46       23.7
```

```
317. MORG-339_phosphopeptide_TMT_4%ACN_032017_OT.4309.4309.2 (ePAISSQNsPEAR)

   Ion         m/z   Intensity
   a1+       331.29        5.1
   b1+       359.26       12.9
   y4+       472.25       16.0
   b3+       527.43        5.4
   y5+       639.47        8.5
   b4+       640.50       10.8
   b5+       727.63        6.3
   a6+       786.46        8.2
   b6+       814.85       15.4
   y8+       968.50        4.3
   y9+      1055.48        8.6
   b8+      1056.37        7.7
   y10+     1168.45        5.1
   y11+     1239.68        6.9
   y12+     1336.67       12.9
   b12+     1520.66        8.7
```

```
318. MORG-339_phosphopeptide_TMT_16%ACN_032017_OT.3263.3263.3 (rADLNQGIGEPQsPSRR)

   Ion         m/z   Intensity
   y6++      405.37       29.4
   y7++      454.39       22.9
   b6++      464.44       14.9
   b7++      492.85       16.4
   y4+       515.40       25.1
   a8++      535.72       12.0
   y9++      547.45       15.8
   b8++      549.52       22.4
   b3+       572.38       23.9
   b9++      578.04       14.7
   a10++     627.96       15.9
   b10++     642.54       48.8
   a11++     677.19       15.9
   y5+       681.93       46.3
   b4+       685.32       22.5
   b11++     691.05       30.2
   y12++     696.41       16.5
   y13++     753.75       11.5
   b12++     755.15       25.3
   b5+       799.90       13.6
   y14++     809.60       32.5
   y6+       810.69       23.6
   y7+       907.46       27.6
   b7+       984.51       20.5
   b16++    1008.76       32.8
   y9+      1093.46       16.5
   b8+      1097.65       13.5
   b10+     1283.75       24.0
```

```
319. MORG-339_phosphopeptide_TMT_25%ACN_032017_OT.11742.11742.3 (kAAVVAsPLLDQQR)

   Ion         m/z   Intensity
   a2+       400.91       24.5
   y3+       431.33       48.1
   b3+       499.96       34.4
   y4+       546.31       43.0
   b9++      573.35       41.0
   b4+       599.08       24.9
   y10++     604.25       20.1
   a10++     616.51       26.9
   b10++     630.25       36.1
   y11++     653.07       50.0
   y5+       659.24       42.0
   a5+       669.98       65.0
   b5+       698.55       24.1
   a6+       741.57       23.7
   b6+       769.83       29.9
   y7+       869.51       94.4
   y8+      1036.58       22.2
   y9+      1107.41       26.2
   y10+     1206.61       24.5
```

```
320. MORG-339_phosphopeptide_TMT_10%ACN_032017_OT.7831.7831.3 (aETSESSGSAPAVPEASAsPk)

   Ion         m/z   Intensity
   y1+       376.31       15.6
   y2+       473.37       64.3
   y7++      499.55       30.7
   b9++      533.38       13.9
   y8++      548.44       67.7
   b10++     568.81       19.5
   y9++      598.15       30.9
   b11++     617.55       21.4
   y10++     633.13       18.8
   y3+       640.34       39.2
   b12++     653.06       60.8
   y11++     682.10       75.5
   a13++     688.76       19.8
   b13++     702.58      115.5
   y4+       711.46       24.4
   b5+       747.43       22.8
   y14++     789.14       15.6
   y5+       798.47       41.1
   a15++     801.08       88.2
   a6+       806.50       20.5
   b15++     815.66       44.9
   b16++     851.28       25.9
   y6+       869.42       17.9
   b17++     894.69       27.6
   b7+       921.54       33.0
   b18++     929.88       14.4
   y7+       998.62       15.9
   b19++    1013.74       30.7
   a20++    1048.32       14.8
   b9+      1065.49       30.1
   y8+      1095.54      106.0
   b10+     1136.51       82.6
   y9+      1194.59       17.1
   b12+     1304.71       21.0
   y11+     1362.71       33.9
   b13+     1403.70       70.7
```

```
321. MORG-339_phosphopeptide_TMT_30%ACN_032017_OT.9984.9984.4 (vkFSSQQASTkQSNAssDVEVEEk)

   Ion         m/z   Intensity
   y2+       505.38       24.8
   b11+++    627.67       24.9
   b7++      632.87       25.0
   y3+       634.48       36.5
   b8++      668.28       24.9
   b12+++    670.13       31.4
   b13+++    699.28       29.3
   y4+       733.47       40.1
   b14+++    737.48       29.2
   b15+++    761.13       30.3
   b10++     762.23       22.6
   a16+++    807.03       35.8
   a17+++    862.53       43.1
   b17+++    871.72       35.4
   y19+++    894.64      103.8
   a18+++    901.56       38.7
   y13++     905.93       24.1
   b18+++    910.42       31.3
   b11++     940.69       21.9
   b19+++    943.79       27.3
   y21+++    953.48       24.7
   y6+       961.56       29.1
   a5+       979.68       23.1
   b20+++    986.83       56.3
   b12++    1004.48       27.6
   b5+      1007.65       30.5
   b21+++   1019.77       44.5
   b13++    1048.27       24.6
   b22+++   1062.84       28.2
   y7+      1076.79       24.8
   y14++    1084.28       18.4
   b23+++   1105.45       38.7
   y23+++   1121.35       26.8
   b15++    1140.58       29.5
   y16++    1178.75       17.0
```

```
322. MORG-339_phosphopeptide_TMT_4%ACN_032017_OT.11285.11285.3 (dEEALEDsAEEkMEESR)

   Ion         m/z   Intensity
   y2+       262.27       15.8
   b1+       345.20        9.6
   y3+       391.28       12.1
   b2+       474.32       14.0
   y6++      504.71       18.6
   y4+       520.29       13.6
   y7++      569.46       27.4
   b3+       603.32       21.7
   y8++      633.98       14.5
   y5+       651.40       14.6
   y9++      669.44       47.4
   y10++     753.01       39.4
   b5+       787.41       22.5
   y11++     810.60       35.9
   a6+       888.38       14.6
   b6+       916.47       24.1
   b12++     943.07       17.7
   y14++     966.93       14.4
   y6+      1008.48       17.8
   b7+      1031.45       43.5
   b14++    1073.13       22.2
   y16++    1096.14       13.9
   a15++    1123.31       13.1
   y7+      1137.55       51.3
   b16++    1181.25       15.8
   b10+     1398.63       14.2
   b11+     1527.58       12.0
```

```
323. MORG-339_phosphopeptide_TMT_16%ACN_032017_OT.6327.6327.3 (sRsDIDVNAAASAk)

   Ion         m/z   Intensity
   b1+       317.26       15.5
   b3++      320.99       14.2
   y1+       376.37       26.1
   b4++      378.33       13.3
   b5++      434.33       14.2
   y2+       447.24       32.9
   b6++      492.30       47.4
   a7++      527.96       16.4
   y3+       534.36       49.2
   b7++      541.91       38.3
   a8++      585.01       25.6
   y10++     594.90       11.1
   b8++      598.72       45.1
   y4+       605.37       52.5
   a3+       612.26       15.7
   a9++      620.07       14.7
   b9++      634.42       67.3
   b3+       640.45       24.2
   b10++     669.87       74.9
   y5+       676.50       46.6
   a11++     691.26       13.6
   b11++     705.46       64.6
   y12++     735.63       17.5
   y6+       747.53       28.9
   b12++     748.88       23.9
   b13++     784.45       25.1
   y7+       861.52       28.9
   b5+       868.55       11.5
   y8+       960.59       30.9
   y9+      1075.62       12.0
   b8+      1196.72       11.6
```

```
324. MORG-339_phosphopeptide_TMT_60%ACN_032017_OT.13738.13738.3 (sFkLSGFSFk)

   Ion         m/z   Intensity
   b1+       397.31       11.0
   y6++      451.03       18.4
   a4++      493.65       20.3
   y7++      507.89       23.0
   y2+       523.62       13.5
   a5++      537.19        9.5
   b6++      579.63       12.5
   y3+       610.32       28.4
   b7++      653.41       21.0
   y8++      686.82       16.1
   b8++      697.21       18.6
   y4+       757.55       12.4
   y9++      759.76       15.5
   b9++      770.68       10.3
   y5+       814.54       22.0
   y6+       901.36       16.0
   y7+      1014.59       11.9
```

```
325. MORG-339_phosphopeptide_TMT_18%ACN_032017_OT.15660.15660.3 (lkGQEDSLASAVDATTGQEAcDsD)

   Ion         m/z   Intensity
   b4++      443.42       18.7
   b5++      507.96       22.0
   b6++      565.71       55.0
   y4+       576.20       34.4
   b7++      609.22       46.9
   y5+       647.29       44.5
   a8++      651.79       28.6
   b8++      665.54       70.3
   b9++      701.30      129.9
   b10++     744.53       62.2
   y6+       776.32       53.7
   b11++     780.01       81.5
   a12++     815.55       18.8
   b12++     829.54      160.0
   b13++     887.04       95.4
   y7+       904.34       33.2
   b14++     922.60      121.5
   a15++     958.55       55.2
   y8+       961.32       82.6
   b15++     973.31      164.2
   a5+       986.30       22.0
   b5+      1014.94       77.4
   b16++    1023.90       90.3
   b17++    1052.37       61.1
   y9+      1062.36       97.0
   a18++    1102.47       20.5
   b18++    1116.08       79.9
   b6+      1129.62       19.9
   y10+     1163.36       96.6
   b19++    1180.97      112.4
   b20++    1216.07       55.7
   b7+      1216.67       77.3
   y11+     1234.41       65.8
   b21++    1296.45       60.5
   y23++    1332.46       38.0
   y12+     1349.45      122.1
   b22++    1353.52       23.0
   y13+     1448.54       56.3
   y14+     1519.50       42.2
   y15+     1606.56       55.1
   y16+     1677.65       37.6
```

```
326. MORG-339_phosphopeptide_TMT_10%ACN_032017_OT.9294.9294.3 (dVQDAIAQsPEk)

   Ion         m/z   Intensity
   y3++      301.83       18.7
   a1+       317.15        9.9
   b1+       344.73        6.9
   y1+       376.32       18.7
   b5++      379.98       16.5
   y4++      385.34       14.4
   a6++      422.48        8.4
   b6++      436.13       26.8
   y5++      449.37       42.5
   b7++      472.04       31.0
   y6++      484.95       56.3
   y2+       505.38        9.7
   a8++      521.56        8.1
   b8++      535.76       16.3
   y7++      541.58       15.4
   b3+       572.39       10.8
   y3+       602.41       34.7
   a9++      605.38       21.9
   y9++      634.50        9.3
   a10++     653.43        6.6
   a4+       659.67        6.6
   a11++     718.19        6.3
   b11++     732.61        9.1
   b5+       758.44       20.3
   y4+       769.39       27.9
   b6+       871.49       44.2
   y5+       897.58        8.2
   b7+       942.54       30.6
   y6+       968.57       12.7
   b8+      1070.66       14.9
   b9+      1237.76        8.7
```

```
327. MORG-339_phosphopeptide_TMT_22%ACN_032017_OT.12526.12526.4 (vEsEEsGDEEGkkHGGGGIVANLSEQSLk)

   Ion         m/z   Intensity
   y1+       376.23       31.1
   y2+       489.48       31.5
   y3+       576.44       43.5
   y4+       704.56       37.0
   a15+++    806.47       25.1
   y5+       833.57       35.8
   b5+       883.41       19.4
   y20+++    899.25       25.4
   y6+       920.55       50.8
   y21+++    942.40       30.9
   b20+++    944.20       32.8
   b21+++    967.50       26.3
   y22+++    980.73       28.4
   a22+++    996.56       24.9
   y23+++    999.66       27.5
   a23+++   1033.84       53.3
   b23+++   1043.52       48.2
   b6+      1050.14       30.3
   y17++    1077.41       33.2
   y25+++   1098.92       25.8
   b25+++   1115.49       51.6
   b13++    1126.39       35.2
   y8+      1147.91       34.5
   a26+++   1148.77       53.2
   b26+++   1158.30       66.0
   a14++    1181.28       49.2
   b27+++   1186.41       38.1
   y27+++   1197.65       19.5
   y9+      1218.97       67.8
   b8+      1222.39       25.9
   b28+++   1224.97       52.2
   y10+     1317.72       21.0
```

```
328. MORG-339_phosphopeptide_TMT_16%ACN_032017_OT.17634.17634.3 (eELAsDLEEMATSSAkR)

   Ion         m/z   Intensity
   b1+       359.22        8.5
   y6++      439.80       10.3
   y7++      475.31        9.3
   b2+       488.33       13.7
   y8++      540.99       11.7
   b3+       601.46       11.7
   y9++      605.55       25.2
   y10++     670.10       15.9
   y11++     726.72       38.9
   b10++     729.20        8.8
   b11++     764.05       51.1
   y5+       777.31       15.1
   y12++     784.38       41.1
   b12++     815.34        9.5
   y13++     867.71       11.0
   y6+       878.70        8.3
   y14++     902.68        9.5
   y7+       949.61        9.1
   b6+       954.43       13.9
   y15++     959.81       12.2
   b7+      1067.00       18.5
   y8+      1080.24       11.8
   b16++    1116.34       15.6
   a9+      1297.95        7.4
   y11+     1451.78        7.6
   b10+     1456.66        8.7
```

```
329. MORG-339_phosphopeptide_TMT_22%ACN_032017_OT.8994.8994.2 (tQsQPILNTk)

   Ion         m/z   Intensity
   b2+       459.30       16.2
   y2+       477.44       18.1
   y3+       591.45       26.8
   y4+       704.60       22.3
   b4+       754.69       22.8
   a5+       823.26       50.3
   y6+       914.61       41.5
   a6+       936.69       34.5
   b6+       964.14       22.9
   y7+      1042.64       39.1
   b7+      1077.50       21.8
   b8+      1191.66       17.0
   y8+      1209.69       16.2
```

```
330. MORG-339_phosphopeptide_TMT_22%ACN_032017_OT.3795.3795.3 (sSPASkPGSTPSRPssAk)

   Ion         m/z   Intensity
   y2+       447.38        8.3
   b6++      508.96        9.6
   y8++      610.04        8.5
   b10++     680.11       23.5
   y11++     732.57       11.4
   b12++     772.26        8.8
   y4+       781.10       14.5
   a13++     835.98       29.0
   b13++     849.84       13.4
   y5+       878.49       12.7
   a15++     967.82       15.3
   y14++    1002.86        9.7
   b6+      1016.65       19.5
   y15++    1038.22       10.0
   b7+      1114.01        9.5
   y7+      1121.38        8.5
   y8+      1218.59       13.5
   b10+     1358.85       11.7
```

```
331. MORG-339_phosphopeptide_TMT_16%ACN_032017_OT.12612.12612.4 (tGDLGIPPNPEDRsPsPEPIYNSEGkR)

   Ion         m/z   Intensity
   b3+       503.24       26.5
   y7++      542.06       14.1
   y9++      647.04       45.7
   b5+       673.35       27.3
   y15+++    702.97       38.9
   y10++     711.02       14.3
   y16+++    741.13       34.8
   b18+++    749.87       14.3
   a6+       758.45       28.5
   y11++     760.07       83.8
   b19+++    782.26       13.9
   y17+++    783.52       21.3
   b6+       786.46       66.1
   b13++     796.61       24.0
   y5+       805.34       22.2
   y18+++    816.39       26.3
   y12++     843.16       63.7
   y19+++    854.00       58.5
   a21+++    865.22       45.7
   a14++     866.29       40.0
   b21+++    874.35       26.7
   a15++     914.62       32.1
   y6+       919.37       77.0
   y23+++    975.37       18.1
   b8+       981.03       43.8
   b24+++    985.03       18.4
   a25+++    994.12       14.6
   y24+++   1013.80       33.1
   a9+      1066.88       13.7
   y7+      1082.66       17.2
   b26+++   1123.06       14.5
   b18++    1125.24       22.9
   y18++    1224.16       17.3
   y19++    1280.96       20.1
   b11+     1320.95       19.3
   y20++    1329.87       26.0
   y21++    1378.37       16.9
   b12+     1435.73       26.7
```

```
332. MORG-339_phosphopeptide_TMT_18%ACN_032017_OT.3827.3827.4 (sRTAsLTSAASIDGSRSR)

   Ion         m/z   Intensity
   a3++      273.80       23.0
   y5++      281.82      150.2
   b4++      323.32       30.9
   b5++      406.37       30.7
   y8++      439.49       32.2
   y9++      474.75       39.0
   y14+++    496.56       27.3
   a7++      499.45      101.2
   y4+       505.15      340.7
   b7++      513.97      127.1
   a13+++    514.90       52.9
   y15+++    520.50       39.5
   b13+++    523.84       25.0
   y16+++    553.95       83.7
   y5+       562.30       40.9
   b9++      592.94       25.9
   y12++     604.52      102.1
   y17+++    606.04       28.3
   a10++     613.84       39.6
   a16+++    614.60       29.6
   b16+++    624.07       33.3
   b10++     628.50       48.9
   a17+++    643.55       24.1
   b17+++    653.16       43.1
   y13++     661.07      138.6
   b11++     671.90       29.5
   y6+       677.28       32.1
   a12++     714.62       25.6
   b12++     728.17       74.4
   y14++     744.62       29.0
   y15++     780.06       24.7
   b13++     786.10      135.3
```

```
333. MORG-339_phosphopeptide_TMT_10%ACN_032017_OT.4512.4512.3 (rGsDELTVPR)

   Ion         m/z   Intensity
   b1++      193.38        5.7
   a2++      208.20        6.9
   y2+       272.24       18.7
   a3++      291.63        8.5
   b3++      305.66       11.2
   b4++      363.22       47.2
   y3+       371.24       18.4
   a5++      413.69       22.6
   a2+       415.25       10.3
   b5++      427.76       61.5
   b2+       443.50        9.5
   a6++      470.35       35.3
   y4+       472.34       45.8
   b6++      484.29        9.1
   y9++      527.14        9.4
   b7++      535.02       15.4
   a8++      570.56        9.0
   b8++      584.32       16.0
   y5+       585.41       40.5
   b3+       610.67        7.6
   b9++      632.90        7.2
   a4+       697.47       12.4
   y6+       714.43       43.1
   y7+       829.40       21.8
```

```
334. MORG-339_phosphopeptide_TMT_10%ACN_032017_OT.12156.12156.3 (dLEFcsTEEEk)

   Ion         m/z   Intensity
   y3++      317.36        9.9
   y1+       376.33       21.4
   y4++      382.37       11.9
   b5++      447.99        9.3
   y2+       505.41       23.7
   y6++      516.61        6.9
   b7++      581.52        8.2
   b3+       587.39        9.8
   y7++      596.24       16.0
   y3+       634.37       26.9
   b9++      711.05       28.6
   b4+       734.52       16.2
   a10++     761.00        9.9
   y4+       763.49        8.3
   b10++     775.67       26.3
   b5+       894.39       11.0
   b8+      1291.56       15.5
```

```
335. MORG-339_phosphopeptide_TMT_22%ACN_032017_OT.11437.11437.4 (nLAkPGVTStsDsEDEDDQEGEkk)

   Ion         m/z   Intensity
   b4++      443.55       25.8
   y4++      460.30       16.3
   b7++      569.85       17.8
   y6++      589.05       34.4
   b8++      620.49       27.1
   y7++      646.71       21.7
   b9++      663.97       17.7
   y9++      768.57       24.8
   y10++     826.28       16.5
   b11++     837.98       24.6
   a4+       857.15       38.6
   y3+       862.61       18.1
   y17+++    866.35       22.7
   a12++     881.62       95.8
   a19+++    887.61       21.5
   y11++     890.56       31.9
   b12++     895.07       35.7
   b19+++    897.11       14.2
   y19+++    918.84       45.6
   y4+       919.88       23.9
   a13++     965.24       20.0
   y12++     974.23       25.2
   b13++     978.49       14.5
   a22+++    992.34       19.7
   a14++    1029.29       15.0
   y13++    1031.10       16.1
   b6+      1039.72       17.3
   y21+++   1070.24       14.6
   y22+++   1093.48       19.9
   a7+      1110.28       14.3
   b18++    1280.86       15.1
   y17++    1299.94       16.2
   y19++    1377.31       26.5
   y20++    1425.83       16.1
```

```
336. MORG-339_phosphopeptide_TMT_22%ACN_032017_OT.9604.9604.4 (aEkQVDTEEAGmVTAATASNVkAsPk)

   Ion         m/z   Intensity
   y2+       473.42      114.0
   b5++      508.13       30.6
   y5++      534.83       31.2
   b6++      565.53       96.4
   b8++      680.59       57.7
   y4+       711.50       33.5
   b9++      745.11       82.9
   y10++     771.19       38.8
   b10++     780.67       77.8
   y11++     806.44       35.0
   b11++     809.04       85.2
   y12++     842.02       45.3
   a21+++    850.65       50.4
   a12++     868.28       31.0
   b12++     882.35       70.8
   a4+       887.39       50.0
   y13++     892.18      124.5
   b13++     932.29       93.9
   b22+++    979.53       82.7
   b14++     982.72       74.8
   a5+       986.34       34.5
   b23+++   1003.15       53.4
   y15++    1015.68       54.3
   b15++    1018.23       53.3
   y16++    1043.94       43.0
   b16++    1053.77       75.9
   b24+++   1058.54       46.9
   y17++    1079.83       42.4
   b17++    1104.16       52.8
   a18++    1125.39       32.7
   b18++    1139.75       63.2
   y18++    1143.84       36.3
   b19++    1183.35       30.2
   y20++    1259.37       70.8
   b21++    1289.61       36.2
```

```
337. MORG-339_phosphopeptide_TMT_20%ACN_032017_OT.7788.7788.3 (dGRGAAQNIIPAsTGAAk)

   Ion         m/z   Intensity
   b1+       345.23       17.3
   y1+       376.31       22.8
   y2+       447.33       33.0
   y7++      457.47       17.5
   b8++      500.46       31.7
   y8++      506.42       42.0
   a9++      543.17       24.5
   b9++      557.06       60.3
   b3+       558.00       21.6
   y9++      562.72       16.8
   y4+       575.41       48.7
   b10++     613.59      102.6
   a11++     648.07       20.4
   a5+       658.37       24.0
   b11++     661.79       22.2
   b5+       686.58       26.1
   b12++     697.59       43.4
   a6+       729.55       30.6
   y14++     811.34       25.4
   a14++     816.99       17.0
   y6+       843.45       19.8
   a15++     846.24       18.2
   b15++     860.40       15.8
   a16++     881.71       18.8
   b16++     895.68       48.4
   y7+       914.56       32.7
   b17++     931.24       27.5
   y17++     946.78       34.9
   b8+       999.56       15.1
   y8+      1011.57       50.5
   b9+      1112.73       21.9
   y9+      1124.53       16.2
   b10+     1225.69       49.0
```

```
338. MORG-339_phosphopeptide_TMT_16%ACN_032017_OT.6934.6934.3 (iGkQSEEsPADDGELR)

   Ion         m/z   Intensity
   y2+       288.15       17.8
   y4+       474.31       31.8
   b5++      486.86       15.8
   a6++      537.33       12.1
   b6++      551.60       30.7
   y5+       589.38       29.4
   b7++      616.06       28.1
   y12++     692.58       12.1
   b8++      699.41       14.9
   y6+       704.43       20.4
   a3+       729.70       19.3
   a9++      734.06       27.2
   b9++      747.73       24.0
   y7+       775.10       19.0
   b10++     783.56       22.1
   b11++     840.84       29.8
   y8+       872.41       49.0
   a12++     884.83       17.1
   b12++     898.68       44.9
   y15++     964.06       15.6
   b5+       972.27       12.5
   b14++     991.69       38.3
   y9+      1039.30       15.0
   b15++    1048.48       19.7
```

```
339. MORG-339_phosphopeptide_TMT_14%ACN_032017_OT.18797.18797.3 (vEAkkEEsEEsEDDmGFGLFD)

   Ion         m/z   Intensity
   b2+       458.25       16.5
   b5++      622.63       20.7
   y6+       655.42       12.5
   b6++      686.86       18.4
   b7++      751.74       38.4
   y7+       802.38       26.0
   b9++      899.46       23.0
   y8+       917.40       28.0
   a10++     949.89       21.1
   b10++     964.28       34.6
   y9+      1032.57       27.5
   a11++    1033.33       14.1
   b11++    1047.12       77.6
   b13++    1169.74       34.6
   b14++    1227.30       38.7
   b15++    1300.87       24.9
   a16++    1315.08       12.8
   b16++    1329.35       27.9
   b6+      1372.51       15.2
   y19++    1399.56       12.4
   b19++    1487.95       15.9
```

```
340. MORG-339_phosphopeptide_TMT_14%ACN_032017_OT.14624.14624.3 (sLRHEAESDFDVADLPQLER)

   Ion         m/z   Intensity
   b6++      502.71       15.2
   a7++      552.66       20.6
   b7++      566.72       19.6
   b8++      610.00       15.7
   y5+       642.46       44.4
   b3+       666.38       20.7
   b9++      667.69       35.5
   b10++     741.64       32.8
   y6+       755.43       56.3
   b11++     799.19       69.6
   b4+       803.48       16.3
   a12++     834.85       30.9
   b12++     847.97       89.5
   y15++     853.16       42.7
   y7+       870.42       37.4
   b13++     883.96       13.9
   b5+       932.64       22.5
   y8+       941.59       75.3
   a6+       975.32       13.7
   a15++     984.44       14.2
   y17++     986.17       33.1
   b15++     998.06       54.9
   y9+      1040.53       43.8
   b17++    1110.85       17.5
   y19++    1120.91       28.3
   y10+     1155.57       31.4
   a8+      1191.31       20.0
   y11+     1302.67       34.2
   y12+     1417.72       14.8
   y13+     1504.82       19.2
```

```
341. MORG-339_phosphopeptide_TMT_18%ACN_032017_OT.12221.12221.4 (gSAEGssDEEGkLVIDEPAkEk)

   Ion         m/z   Intensity
   y5++      516.09       23.5
   b4+       574.46       13.3
   y6++      580.26       25.5
   y7++      638.14       12.2
   y12+++    671.95       29.7
   y8++      694.64       25.6
   b11++     698.03       13.6
   y13+++    715.45       35.5
   y9++      744.17       19.6
   y14+++    758.29       38.5
   a6+       770.52       19.7
   y15+++    796.86       26.0
   y10++     800.39       13.8
   b18+++    807.53       13.7
   a19+++    821.16       29.5
   b19+++    830.51       24.8
   b12++     877.20       14.7
   y18+++    926.85       11.2
   y4+       933.49       19.2
   b7+       965.76       13.1
   y19+++    969.59       16.2
   b14++     982.69       17.8
   a21+++    983.60       16.9
   b21+++    992.37       12.9
   y20+++    993.50       13.8
   b15++    1039.89       19.0
   b8+      1080.29       15.5
   b16++    1096.98       17.8
```

```
342. MORG-339_phosphopeptide_TMT_18%ACN_032017_OT.5849.5849.4 (sYEsSEDcPEAASSPTRkVk)

   Ion         m/z   Intensity
   y6+++     396.30       27.1
   y10+++    501.87       33.5
   a13+++    565.27       41.4
   y12+++    577.15       70.6
   a14+++    594.41       36.3
   b14+++    603.66       33.4
   b3+       609.34       28.2
   y13+++    630.52       58.5
   y7++      637.45       23.6
   a9++      669.16       25.1
   y8++      681.24       40.6
   y15+++    711.79       34.1
   y9++      716.38       49.8
   b17+++    721.59       61.7
   y16+++    740.98       63.5
   a11++     768.83       23.6
   b4+       776.34       28.4
   b11++     783.23       27.9
   y17+++    795.97       23.5
   y11++     816.81       47.0
   a18+++    831.85       28.7
   b18+++    841.33       41.2
   a13++     847.55       25.3
   b13++     861.56       25.4
   y12++     865.20       87.7
   b19+++    874.13       30.7
   b14++     905.25       61.0
   y13++     945.35       24.3
   b6+       992.55       27.2
   y14++    1002.56       27.0
   b7+      1107.51       29.1
   b8+      1267.46       34.1
```

```
343. MORG-339_phosphopeptide_TMT_18%ACN_032017_OT.6575.6575.3 (scsIPcGSPNSkELSR)

   Ion         m/z   Intensity
   y3+       375.21       15.3
   y8++      580.25       24.8
   y10++     652.78       14.4
   a11++     714.70       18.9
   a4+       728.94       23.5
   y11++     732.28       17.2
   b4+       757.62       37.9
   y12++     781.10       32.2
   a5+       825.94       14.5
   y13++     837.73       50.0
   b5+       854.68       14.6
   y14++     921.37       16.4
   b13++     972.15       24.4
   a6+       986.66       21.1
   y15++    1001.17       18.5
   b14++    1028.80       14.5
   b7+      1071.13       17.3
   y8+      1159.86       14.6
```

```
344. MORG-339_phosphopeptide_TMT_16%ACN_032017_OT.9311.9311.3 (sPsPDFYkR)

   Ion         m/z   Intensity
   b2++      247.26        7.4
   y2++      266.71        7.9
   a3++      317.25       14.5
   y3++      348.26       11.7
   b4++      379.90        5.5
   b1+       397.20       31.3
   y4++      421.74       41.2
   a2+       466.55        6.6
   y5++      479.47        8.1
   b2+       494.44        7.2
   a6++      496.67        5.9
   b6++      510.58        5.8
   y6++      527.90       64.3
   y2+       532.64       10.8
   y7++      610.98       50.3
   a3+       633.35        5.5
   y8++      659.94       43.0
   b3+       661.12        9.0
   a4+       730.47        5.7
   b8++      770.82       11.0
   b5+       873.31       20.7
```

```
345. MORG-339_phosphopeptide_TMT_22%ACN_032017_OT.3427.3427.3 (vSHsPPPkQRSPtVTk)

   Ion         m/z   Intensity
   b1+       329.26        9.1
   y2+       477.45        8.8
   b3+       553.49        8.3
   y3+       576.54        8.2
   b4+       720.47       14.3
   a5+       789.00       10.6
   y9++      791.90       13.2
   a10++     813.15       24.2
   b10++     826.73       54.1
   y10++     840.81       35.8
   y5+       854.47       23.9
   a11++     856.44       12.3
   y11++     888.80       47.2
   y12++     937.82       63.3
   y13++    1021.20       12.5
   y14++    1089.71       11.8
   y15++    1133.35       14.3
```

```
346. MORG-339_phosphopeptide_TMT_10%ACN_032017_OT.9366.9366.4 (sADRRLsGAQAELALQEESVR)

   Ion         m/z   Intensity
   y2+       274.22       10.3
   b1+       317.27        7.4
   y3+       361.27       17.7
   y6++      374.47        8.7
   y11+++    415.19       11.8
   y7++      430.92       14.5
   y8++      466.45        9.6
   a3+       474.91        9.5
   y4+       490.26       22.4
   a12+++    508.45        6.9
   b12+++    518.09       25.6
   y9++      522.98       13.2
   b13+++    555.91       41.5
   y15+++    556.90        7.5
   a8++      562.88        8.6
   b14+++    579.50       70.7
   y10++     587.44        8.6
   a9++      598.56        8.3
   a15+++    607.97       12.1
   b9++      612.25       11.0
   b15+++    617.07       49.1
   y5+       619.30       33.0
   y11++     622.74       17.1
   a4+       631.52       25.8
   b16+++    659.89       24.9
   b17+++    702.87       45.9
   b11++     711.91       10.2
   b18+++    746.00       51.4
   y6+       747.35       36.1
   b19+++    775.00       32.3
   b12++     776.53       25.4
   a20+++    798.26        6.9
   b20+++    808.02       36.1
   b13++     832.92       11.1
   y7+       860.33       24.3
   b14++     868.75       34.0
   a15++     911.46        6.5
   b15++     924.91       15.7
   y8+       931.54       15.4
```

```
347. MORG-339_phosphopeptide_TMT_16%ACN_032017_OT.8449.8449.3 (sEsPPAELPSLRR)

   Ion         m/z   Intensity
   y5++      314.81       16.9
   b1+       317.27       14.0
   y6++      371.35       37.3
   y7++      435.88        8.6
   b6++      439.83        7.9
   b2+       446.30       13.2
   y8++      471.24       11.1
   a7++      490.52       11.1
   b7++      504.11        9.2
   y9++      520.14       15.0
   y4+       531.31       11.5
   a8++      546.34        7.7
   b8++      561.24       16.5
   y10++     568.46      123.9
   a9++      595.07        8.8
   b3+       613.26       19.9
   y5+       628.43       19.6
   y11++     651.79        7.5
   a4+       682.02        8.1
   y12++     716.83        7.4
   y6+       741.52        9.8
   b12++     787.55       22.7
   a7+       979.56        7.3
   b7+      1007.41       25.3
   b8+      1120.54       11.5
```

```
348. MORG-339_phosphopeptide_TMT_8%ACN_032017_OT.2080.2080.3 (eQQPTSASERQTPQAPQsPR)

   Ion         m/z   Intensity
   y2+       272.14       24.0
   a1+       331.23        8.8
   b1+       359.25       10.5
   y3+       439.19        8.9
   b2+       487.38        9.6
   y4+       567.42       10.3
   b3+       615.40       20.0
   y5+       664.35       29.6
   b10++     672.54       23.8
   y11++     673.33       20.7
   y6+       735.38       19.7
   b11++     736.53       22.0
   y12++     737.42       11.0
   a12++     772.92       20.6
   y13++     781.51       16.9
   a5+       785.31       14.9
   b12++     787.14       37.2
   y14++     816.94       13.2
   a13++     821.03       14.5
   y7+       863.56       12.3
   a14++     885.55        8.9
   b14++     899.70       35.5
   b6+       900.55       18.4
   y16++     910.76       14.9
   a15++     921.43       14.1
   b15++     935.23       59.4
   y17++     959.35       13.1
   y8+       960.41       27.6
   b7+       971.57       13.2
   b16++     983.87       13.9
   a8+      1030.90       10.2
   b17++    1047.37       16.7
   y9+      1061.42       14.8
   b18++    1131.33       29.0
   b9+      1187.73       16.9
   y10+     1189.79       16.6
   b12+     1572.86       12.7
```

```
349. MORG-339_phosphopeptide_TMT_10%ACN_032017_OT.11648.11648.4 (qADVADQQTTELPAENGETENQsPAsEEEkEAkSD)

   Ion         m/z   Intensity
   b8++      543.51       27.6
   b3+       544.32       28.3
   b9++      594.02       35.8
   b4+       643.42       27.3
   b10++     644.41       19.0
   y4+       649.39       20.0
   a18+++    700.11       54.6
   b11++     709.03       25.4
   b5+       714.50       28.2
   y15+++    756.75       24.0
   b12++     765.53       31.4
   b6+       829.56       29.6
   y11++     881.25       20.5
   a7+       929.58       32.4
   b7+       957.55       32.8
   y22+++   1000.13       37.7
   y13++    1013.20       51.3
   b18++    1064.50       23.9
   a27+++   1068.25       27.0
   b8+      1085.54       38.2
   b19++    1114.75       26.5
   y15++    1133.84       39.8
   y6+      1135.64       19.2
   b9+      1186.55       29.3
   y17++    1248.87       29.9
   b10+     1287.66       19.2
   y19++    1342.01       33.3
   y20++    1398.88       16.9
   b11+     1416.69       37.4
   a24++    1417.63       34.1
   y22++    1499.56       42.9
   b12+     1529.77       62.9
   y23++    1547.54       24.5
```

```
350. MORG-339_phosphopeptide_TMT_18%ACN_032017_OT.14885.14885.2 (iDIsPSALR)

   Ion         m/z   Intensity
   y2+       288.28        8.9
   a1+       315.35        7.9
   b1+       343.23       18.6
   y3+       359.31        6.8
   b2+       458.30       48.2
   a3+       543.34       60.6
   b3+       571.39       17.4
   a4+       710.43       13.2
   b4+       738.31        8.6
   y7+       823.55       24.4
   y8+       938.53       11.1
   a7+       965.71        7.1
   b7+       993.61        8.1
   b8+      1106.58       16.7
```

```
351. MORG-339_phosphopeptide_TMT_25%ACN_032017_OT.10373.10373.4 (kNLAkPGVTSTsDsEDEDDQEGEk)

   Ion         m/z   Intensity
   y1+       376.41       31.2
   y2+       505.43       22.8
   y3+       562.38       68.7
   y4+       691.36       50.5
   y5+       819.43       37.7
   b18+++    831.43       24.4
   y12++     852.75       25.8
   b12++     861.56       77.6
   b19+++    870.01       47.7
   a20+++    903.79       37.3
   y20+++    924.10       24.8
   y6+       934.65       38.7
   a21+++    946.54       51.7
   a22+++    965.49       32.8
   a5+       985.21       39.7
   y22+++    986.07       29.9
   a14++     988.81       24.9
   b23+++   1018.04       31.3
   y23+++   1023.73       23.5
   y7+      1049.62       44.1
   y16++    1081.10       23.1
   y18++    1158.60       27.8
   y8+      1178.59       32.0
   y19++    1207.66       41.6
```

```
352. MORG-339_phosphopeptide_TMT_8%ACN_032017_OT.9871.9871.3 (eDGIDAVEVAADRPGsPR)

   Ion         m/z   Intensity
   y2+       272.15       44.5
   b1+       359.23        7.8
   y6++      375.20       16.3
   b5++      380.22       14.1
   b6++      415.50       13.8
   b7++      465.29       16.9
   y8++      468.34        9.4
   b2+       474.30       21.2
   y4+       496.34        9.7
   y9++      504.21       12.4
   b8++      529.76       17.3
   y10++     553.43        8.9
   b9++      579.33       14.2
   y5+       593.30       24.3
   b10++     614.97       12.4
   a4+       616.35        7.4
   b4+       644.40       12.2
   y12++     667.24        8.2
   a12++     693.96       10.3
   y13++     703.02       32.4
   b12++     707.87       11.5
   y6+       749.36       12.3
   b5+       759.37       29.5
   y14++     760.45       11.6
   a13++     772.37        9.6
   b13++     785.93       43.1
   a14++     820.51        7.6
   b6+       830.41       25.7
   b14++     834.53        9.3
   y16++     845.34       18.9
   b15++     862.96       15.4
   y7+       864.36       18.1
   a7+       901.32        8.0
   y17++     903.09        7.9
   b7+       929.59       15.9
   y8+       935.43       16.4
   b16++     946.05       24.5
   b17++     994.90        9.0
   y9+      1006.42       19.6
   b8+      1058.54       25.0
   y10+     1105.52       11.2
   b9+      1157.56       14.1
   b10+     1228.66       13.0
   y11+     1234.56        7.8
   b12+     1414.68       36.2
```

```
353. MORG-339_phosphopeptide_TMT_10%ACN_032017_OT.6638.6638.3 (dRAsPESLDLTR)

   Ion         m/z   Intensity
   a2++      237.22        8.3
   b2++      251.16       26.1
   a3++      272.71       12.1
   b3++      286.77       20.2
   a1+       317.30        9.1
   b1+       345.25        7.6
   a4++      356.29       11.4
   b4++      370.18       40.8
   y3+       389.26       40.3
   y8++      465.77       34.2
   a6++      469.41       18.2
   b6++      483.32       37.2
   b2+       501.14        9.6
   y4+       504.45       21.3
   a7++      512.65       22.8
   b7++      526.51       23.2
   a3+       544.37       12.8
   y9++      548.88       17.5
   b8++      582.89       10.3
   y5+       617.47       24.1
   b9++      641.00       62.5
   y11++     662.78        7.8
   b10++     697.53       21.5
   y6+       704.36       38.0
   b4+       739.44       20.6
   y7+       833.52       12.8
   y8+       930.50       86.2
```

```
354. MORG-339_phosphopeptide_TMT_30%ACN_032017_OT.4756.4756.3 (tVTPASSAkTsPAkQQAPPVR)

   Ion         m/z   Intensity
   y4+       468.43       10.1
   b3+       531.56        6.8
   y10++     661.11       24.4
   b10++     702.27        5.8
   y7+       795.57        7.3
   a7+       845.97        7.2
   a8+       916.71       31.3
   b8+       944.06       17.2
   a14++    1034.59        7.3
   b14++    1048.27        7.1
   y16++    1096.50        9.1
   b15++    1112.45       10.4
   y17++    1131.52        8.1
   y8+      1152.29        7.5
   a16++    1162.63       10.2
   b16++    1176.38        8.7
   y18++    1180.47       12.8
   b17++    1211.93       15.8
   y20++    1280.23        9.6
   b19++    1309.11        8.9
   y10+     1320.74       17.3
```

```
355. MORG-339_phosphopeptide_TMT_20%ACN_032017_OT.17774.17774.2 (sEGEFkQTSSFLV)

   Ion         m/z   Intensity
   b1+       397.26       14.1
   b2+       526.27       13.5
   b3+       583.42       12.0
   b4+       712.38       26.3
   y7+       781.47       15.0
   b5+       859.48       22.0
   y8+      1138.71       27.7
   b6+      1216.69       25.1
   y9+      1285.73       35.3
   b7+      1344.69       18.1
   y10+     1414.81       13.4
   y11+     1471.88       34.0
   y12+     1600.89       28.3
```

```
356. MORG-339_phosphopeptide_TMT_6%ACN_032017_OT.5532.5532.4 (aEEGGEsEGDAsEkDAk)

   Ion         m/z   Intensity
   a9+++     376.36       20.8
   b2+       430.53        6.9
   y2+       447.33       17.8
   y4++      460.34       19.8
   y9+++     487.24       17.0
   b12+++    503.74        6.2
   y5++      524.45        7.3
   y10+++    530.33        7.2
   a8++      535.34        9.3
   a13+++    537.68        6.9
   b13+++    546.50        6.3
   b8++      549.92        9.8
   b3+       559.24        7.3
   y3+       562.38       23.5
   a9++      564.32        7.6
   b9++      577.96       16.5
   y11+++    585.88       15.2
   a4+       588.56       11.7
   y6++      608.40       29.6
   b4+       616.19       26.1
   a10++     621.35       16.0
   y12+++    628.69       16.9
   a14+++    655.83        7.7
   a11++     657.51        7.7
   b14+++    665.58       23.2
   y14+++    666.56       12.6
   b11++     671.48       31.8
   b5+       673.49        8.1
   y8++      701.45       11.9
   b15+++    704.02       19.5
   b16+++    727.64       31.6
   y9++      729.67       19.1
   b12++     755.17        6.2
   b6+       802.37       16.1
   b14++     997.99        6.7
   b8+      1098.51        9.3
   b9+      1155.33        7.4
```

```
357. MORG-339_phosphopeptide_TMT_10%ACN_032017_OT.9663.9663.3 (lEkEEEEGISQEsSEEEQ)

   Ion         m/z   Intensity
   b5++      544.53       18.7
   b6++      608.96       32.4
   y10++     622.99       22.8
   b7++      673.60       44.2
   a9++      744.51       22.6
   b9++      758.68      136.0
   y13++     780.75       19.1
   y6+       788.31       24.2
   a3+       801.40       23.9
   b10++     802.13       51.4
   b3+       829.19       20.0
   y14++     845.28       35.0
   a11++     851.59       27.1
   b11++     865.66       59.2
   a12++     916.53       21.1
   y7+       917.32       22.5
   a4+       930.65       56.4
   a13++    1000.01       19.0
   y8+      1045.69       24.9
   y16++    1088.84       27.3
   b15++    1122.28       51.0
   y9+      1132.36       75.8
   b16++    1186.81       74.4
   b17++    1251.38       93.6
```

```
358. MORG-339_phosphopeptide_TMT_22%ACN_032017_OT.17815.17815.3 (qIsQDVkLEPDILLR)

   Ion         m/z   Intensity
   b2+       471.33       10.6
   y4+       514.46       28.9
   y9++      663.29       18.6
   b7++      669.81       14.9
   y10++     713.24       56.2
   b8++      725.68       14.2
   y6+       726.57       26.5
   a4+       737.91       29.4
   b9++      790.32       27.4
   y12++     834.85       32.8
   y7+       855.61       12.7
   b5+       881.40       25.7
   a11++     882.65       17.8
   b11++     896.75       34.3
   b12++     953.11       10.8
   y8+       968.60        9.8
   b13++    1009.57       13.7
   b14++    1066.51       22.9
```

```
359. MORG-339_phosphopeptide_TMT_18%ACN_032017_OT.21463.21463.3 (vEAkkEEsEEsEDDMGFGLFD)

   Ion         m/z   Intensity
   y5+       598.34       22.4
   b5++      622.73       39.4
   y6+       655.43       34.7
   b6++      687.08       45.9
   b7++      751.80       66.4
   y7+       786.35       85.0
   a9++      885.53       21.3
   b9++      899.74       31.8
   y8+       901.47       35.7
   a10++     950.36       32.5
   b10++     964.30       66.0
   y16++     998.77       55.6
   b11++    1047.97      424.4
   b12++    1111.91       26.5
   b13++    1169.71       74.1
   b14++    1227.39       76.6
   b5+      1243.88       64.2
   b15++    1292.83       80.7
   y11+     1312.41       33.5
   b16++    1321.39       46.1
   b17++    1395.01       57.8
   b18++    1423.47       28.5
   y12+     1441.70       22.8
   b19++    1480.06       34.6
```

```
360. MORG-339_phosphopeptide_TMT_8%ACN_032017_OT.5128.5128.2 (sFsPEER)

   Ion         m/z   Intensity
   y1+       175.25        6.6
   y2+       304.15       22.2
   b1+       317.27       11.0
   y3+       433.40        6.1
   b2+       464.15        9.8
   y4+       530.29       50.5
   b3+       631.30       10.5
   a5+       829.32        6.5
   y6+       844.29       12.5
   b5+       857.33       25.7
   b6+       986.36       14.3
```

```
361. MORG-339_phosphopeptide_TMT_25%ACN_032017_OT.19897.19897.4 (lkNDSDLFGLGLEEmGPkEssDEDR)

   Ion         m/z   Intensity
   b4++      465.50       27.0
   b6++      566.65       28.5
   b8++      696.53       21.2
   b9++      724.97       13.4
   a17+++    754.84       21.3
   b10++     781.71       27.2
   b11++     809.90       21.3
   y11++     828.55       14.9
   y19+++    838.44       32.4
   a12++     852.13       15.4
   b12++     866.77       25.2
   y6+       867.86       37.3
   a18+++    874.49       18.9
   b18+++    883.17       61.7
   y12++     892.47       24.5
   b19+++    926.88       19.3
   b4+       929.65       15.1
   b13++     930.96       16.5
   y22+++    944.67       18.7
   y13++     957.45       15.6
   a20+++    972.74       26.3
   b14++     995.86       24.9
   y14++    1013.99       18.3
   y15++    1042.83       15.4
   b15++    1069.02       15.2
   b22+++   1075.82       27.8
   b16++    1098.00       19.0
   y16++    1099.19       19.1
   a6+      1103.85       16.4
   a23+++   1109.87       13.9
   b23+++   1119.44       17.9
   b6+      1131.21       17.3
   a17++    1132.14       16.6
   b24+++   1157.61       26.0
   y18++    1201.08       13.0
   y19++    1257.69       17.6
```

```
362. MORG-339_phosphopeptide_TMT_20%ACN_032017_OT.6426.6426.3 (sATRVIGGPVTPR)

   Ion         m/z   Intensity
   y5++      285.31       11.7
   y7++      342.26        8.5
   a4++      349.66        9.3
   b4++      363.37       28.2
   b1+       397.29       14.6
   y8++      398.78       11.5
   b5++      412.81       27.6
   a6++      455.27       17.6
   b2+       468.57       25.5
   b6++      469.41       37.3
   y4+       472.31       10.2
   a7++      483.38       10.1
   b7++      497.28       19.8
   y10++     526.14       39.6
   y5+       569.46       19.8
   y11++     576.93       19.7
   a10++     609.91       11.2
   y12++     612.44       30.0
   b10++     624.13       18.0
   y6+       626.21       32.4
   a11++     660.65       12.7
   b11++     675.22       24.6
   y7+       683.52       42.2
   y8+       796.50       26.5
   y9+       895.53       26.7
```

```
363. MORG-339_phosphopeptide_TMT_18%ACN_032017_OT.9025.9025.3 (sIFREEsPLR)

   Ion         m/z   Intensity
   y1+       175.14       12.5
   b1+       317.22       17.3
   y5++      341.50        6.7
   b4++      367.35       15.7
   y3+       385.29       44.8
   a2+       402.18        6.2
   a5++      417.94        6.6
   b2+       430.36       10.6
   b5++      431.75       17.6
   a6++      482.36      160.6
   b6++      496.40       29.1
   a3+       549.63        6.3
   y4+       552.39       10.4
   y8++      557.26        8.9
   a7++      565.53        9.5
   b3+       577.41       10.3
   b7++      579.54       19.0
   a8++      614.36       13.1
   b8++      628.71        6.6
   a9++      670.97       25.3
   y5+       681.45        8.8
   b9++      685.02       36.7
```

```
364. MORG-339_phosphopeptide_TMT_12%ACN_032017_OT.5005.5005.2 (sVSGsPEPAAk)

   Ion         m/z   Intensity
   b1+       317.24        9.7
   y1+       376.33        9.3
   b2+       416.30       10.6
   b3+       503.35       12.2
   a4+       532.44        9.6
   y4+       615.42       27.2
   b5+       727.59        9.0
   y5+       744.36       31.0
   a6+       796.37        9.9
   y6+       841.49       44.2
   a7+       925.19        9.5
   b7+       953.42       20.0
   y9+      1152.66       10.8
   b10+     1192.51       10.7
```

```
365. MORG-339_phosphopeptide_TMT_16%ACN_032017_OT.9255.9255.3 (eVIEAsDkEGLsPAkR)

   Ion         m/z   Intensity
   y4++      350.77       12.1
   b7++      526.78        8.3
   y4+       700.52       15.8
   y9++      762.30       14.6
   b9++      770.17       15.0
   b5+       771.54       12.7
   y10++     819.51        8.7
   a11++     841.82       34.3
   y12++     938.62       12.2
   a13++     973.47        8.3
   y6+       980.23        9.8
   b7+      1053.50       14.2
   y14++    1060.48       13.8
   b15++    1202.06       24.0
```

```
366. MORG-339_phosphopeptide_TMT_10%ACN_032017_OT.13528.13528.3 (dFQEYVEPGEDFPAsPQRR)

   Ion         m/z   Intensity
   y4++      278.90       10.9
   y6++      397.38       14.8
   y8++      519.95       29.8
   y11++     670.10       36.0
   y12++     719.04       31.9
   y5+       723.36       10.5
   b4+       749.56       19.4
   b11++     769.41       10.4
   y6+       794.52       21.4
   a12++     829.86       21.6
   y14++     833.15       30.6
   b12++     843.67       21.7
   b13++     891.46       14.7
   b5+       912.48       17.2
   a14++     913.64       20.3
   y15++     914.74       16.2
   b6+      1011.61       17.1
   y8+      1038.61       10.2
   b7+      1140.57       39.0
   y12+     1437.01       10.4
   b11+     1538.80       31.9
   b12+     1685.83       13.7
```

```
367. MORG-339_phosphopeptide_TMT_14%ACN_032017_OT.8335.8335.4 (kFPDRLADDEGDsDSESVGQSR)

   Ion         m/z   Intensity
   y4+       447.29       31.9
   y9++      482.39       16.9
   y5+       546.40       24.4
   a14+++    615.07       14.8
   y6+       633.35       33.9
   y12++     652.52       16.6
   y18+++    668.29       58.1
   a4+       689.68       22.1
   b10++     709.14       74.8
   a17+++    715.72       30.1
   b4+       717.40       30.4
   a11++     723.76       48.4
   b17+++    725.63       27.7
   a18+++    749.09       15.0
   b18+++    758.60       23.3
   y7+       762.54       20.4
   b19+++    776.89       17.8
   y21+++    787.95       20.2
   y15++     831.36       17.0
   y8+       849.48       30.1
   a13++     864.15       20.4
   y17++     924.34       23.0
   y9+       964.46       29.8
   a15++     965.45       19.5
   y18++    1002.29       24.0
   y10+     1131.45       21.9
   a8+      1144.19       17.7
```

```
368. MORG-339_phosphopeptide_TMT_22%ACN_032017_OT.12311.12311.3 (rLksSTsFANIQENAT)

   Ion         m/z   Intensity
   y5+       562.19       13.3
   b6++      606.74       16.9
   a7++      676.28       14.5
   b8++      763.34      109.9
   y13++     765.40       25.8
   a9++      784.91       41.3
   b10++     856.09       27.5
   b3+       856.95       17.4
   y8+       860.28       22.3
   a11++     898.72       15.4
   b11++     912.82       39.0
   b12++     976.79       30.5
   y15++    1000.40       14.8
   y9+      1007.74       19.9
```

```
369. MORG-339_phosphopeptide_TMT_25%ACN_032017_OT.8738.8738.3 (tEkNHFESSPLPtSPk)

   Ion         m/z   Intensity
   y2+       473.42       52.8
   b5++      534.93       21.7
   b6++      608.57       28.9
   b7++      673.11       38.8
   b8++      716.64       25.0
   y4+       741.35       57.0
   b9++      760.18       58.0
   b3+       817.66      105.0
   y12++     818.29       65.1
   y5+       838.31       64.0
   b11++     865.17       50.7
   y13++     875.89       19.8
   a12++     899.49       19.6
   b12++     913.60       19.2
   y6+       951.64       25.8
   a13++     989.88       51.8
   b14++    1047.46       28.0
   y7+      1048.48       25.5
```

```
370. MORG-339_phosphopeptide_TMT_10%ACN_032017_OT.2944.2944.3 (sRkSPsPAGGGsSPYSR)

   Ion         m/z   Intensity
   a3++      442.24       10.3
   b4++      499.36       13.7
   y4+       522.32       19.5
   a5++      534.18       12.3
   y13++     690.07       13.1
   a8++      701.18       39.5
   b8++      715.29       21.5
   y14++     733.71       49.2
   b9++      744.09       32.2
   a10++     758.27       18.9
   b10++     772.73       42.8
   y6+       776.66       58.4
   y7+       833.52       11.2
   a12++     870.39       19.1
   a3+       882.39       14.5
   b12++     884.35       19.9
   a4+       969.39       10.6
   y10+     1018.59       12.3
   b15++    1058.15       12.1
   y11+     1115.51       16.8
```

```
371. MORG-339_phosphopeptide_TMT_18%ACN_032017_OT.5631.5631.3 (rRAsDLAGSVDTGSAGSVk)

   Ion         m/z   Intensity
   a4++      376.37       19.9
   b5++      448.61       21.6
   y3+       562.43       16.0
   y4+       619.45       25.5
   y13++     683.30       16.1
   y5+       690.58       23.0
   b11++     719.55       52.7
   y14++     739.37       16.7
   a12++     756.28       19.2
   b12++     770.32       38.7
   y6+       777.45       47.2
   b4+       780.76       48.6
   y7+       834.24       25.6
   a15++     863.82       15.0
   b15++     877.72       29.0
   y8+       935.64       36.0
   b18++     998.72       14.5
   y9+      1050.61       20.7
   a7+      1051.48       17.5
   a10+     1294.77       14.5
```

```
372. MORG-339_phosphopeptide_TMT_16%ACN_032017_OT.2804.2804.3 (sRSPVPPAEk)

   Ion         m/z   Intensity
   b3++      320.66       19.7
   y4++      337.27       20.3
   y1+       376.44       13.0
   y5++      385.83       61.7
   b1+       397.26       10.7
   a5++      404.77       67.4
   b5++      418.81       62.1
   a6++      453.31       13.9
   b6++      467.33       20.1
   y7++      483.52       19.0
   y2+       505.49       26.9
   b7++      515.68       47.3
   a2+       525.25       20.0
   y8++      527.73       46.4
   b2+       553.24       13.5
   y3+       576.56       16.5
   y9++      605.33       15.1
   b9++      616.17       16.0
   y4+       673.49       18.6
   y5+       770.46       66.1
   b5+       836.43       39.3
   y7+       966.67       10.4
```

```
373. MORG-339_phosphopeptide_TMT_16%ACN_032017_OT.12620.12620.2 (sAsADNLILPR)

   Ion         m/z   Intensity
   y2+       272.26        7.7
   b1+       317.30       17.4
   y3+       385.32       15.6
   y4+       498.38        9.9
   a3+       527.17        7.4
   y5+       611.32        9.1
   b4+       626.66        8.2
   a5+       712.83       13.3
   y6+       725.45       24.7
   b5+       741.36        8.4
   y7+       840.58       11.8
   y8+       911.56       26.1
   b7+       968.42       13.3
   y9+      1078.55       11.9
   b8+      1081.58       13.3
   b9+      1194.61       18.2
   b10+     1291.81        7.6
```

```
374. MORG-339_phosphopeptide_TMT_20%ACN_032017_OT.9072.9072.3 (qLVRGEPNVSyIcSR)

   Ion         m/z   Intensity
   b1+       358.34       15.2
   b4++      364.01       28.7
   y3+       422.21       20.0
   a2+       442.89       51.4
   b6++      456.97       84.0
   b2+       471.33       20.4
   y4+       535.42       17.6
   b8++      562.51       43.8
   b3+       570.48       18.1
   y9++      588.37       34.6
   a9++      598.28       36.0
   b9++      612.06       46.0
   y10++     652.59       26.3
   b10++     655.63       26.9
   b4+       726.46       19.9
   a5+       755.47       14.8
   b11++     777.23       31.4
   y5+       778.42       28.1
   a12++     819.64       18.2
   b12++     833.21       26.3
   y14++     865.40       48.0
   a13++     899.39       24.0
   b6+       912.31       20.7
   b13++     913.77       52.0
   b14++     957.16       32.3
   y7+       964.33       31.2
   y8+      1078.50       18.0
   y9+      1175.49       55.1
   y11+     1361.51       18.1
```

```
375. MORG-339_phosphopeptide_TMT_18%ACN_032017_OT.5405.5405.3 (lRLsPSPtSQRSR)

   Ion         m/z   Intensity
   y5++      317.28       12.8
   b4++      390.37       12.8
   y6++      407.47       12.8
   a5++      424.38       12.3
   b6++      482.11       13.9
   y8++      499.53       26.2
   y4+       546.51       24.8
   y9++      548.48       28.1
   a3+       584.10       11.6
   a8++      607.03       25.1
   b9++      664.78       17.3
   y11++     687.84       17.7
   b10++     729.13       17.3
   b4+       779.47       13.0
   b11++     806.70       11.3
   y6+       814.50       13.4
   b12++     850.77       13.7
```

```
376. MORG-339_phosphopeptide_TMT_22%ACN_032017_OT.13781.13781.3 (gVPRGLYDGPVcEVSVtPk)

   Ion         m/z   Intensity
   a7++      472.39       16.9
   y2+       473.38       76.6
   b8++      544.58       95.2
   y3+       654.33       21.5
   a11++     656.81       13.1
   b11++     670.92       18.9
   a12++     737.18       12.7
   b12++     751.15       21.1
   a6+       781.73       16.8
   y12++     798.65       18.9
   a13++     801.34       16.3
   b6+       809.37       12.8
   b13++     815.65       46.8
   y5+       840.48       26.8
   b14++     865.26       33.3
   a15++     894.68       18.8
   b15++     908.35       33.6
   y14++     936.49       14.2
   y6+       939.60       20.9
   b16++     957.92       25.2
   b8+      1087.65       39.5
   y17++    1092.28       14.4
   b9+      1144.54       16.2
   y11+     1481.77       37.9
```

```
377. MORG-339_phosphopeptide_TMT_8%ACN_032017_OT.4507.4507.3 (eRDHsPTPSVFNSDEER)

   Ion         m/z   Intensity
   y2+       304.30        9.6
   b3++      315.87       19.2
   y6++      375.19       12.7
   y3+       433.18       18.0
   b5++      467.54       15.7
   y10++     590.55       23.8
   b3+       630.40       22.4
   y11++     640.95       14.2
   y12++     689.70       24.0
   a4+       739.50       42.2
   y6+       749.23       19.9
   y14++     841.65       30.2
   b13++     882.13       16.9
   y7+       896.40       18.0
   b14++     939.79       13.5
   y8+       995.53       18.9
   b15++    1004.67       22.6
   b16++    1068.91       12.4
   y10+     1179.42       34.1
   y12+     1377.64       34.0
```

```
378. MORG-339_phosphopeptide_TMT_18%ACN_032017_OT.12747.12747.4 (eVWASDsGHEEDcLSPEPPLRPk)

   Ion         m/z   Intensity
   b2+       458.32       28.2
   a10+++    460.35       17.3
   y6++      468.99      130.1
   b10+++    470.02       27.6
   y2+       473.31       23.2
   y8++      582.04       62.1
   a9++      625.64       64.3
   b9++      640.00       35.1
   b3+       644.52       19.7
   y15+++    678.60       30.7
   y10++     682.14       47.1
   y16+++    697.60       49.1
   b10++     704.52       45.1
   b4+       715.33       32.1
   b17+++    747.11       29.5
   y17+++    753.27       43.2
   a11++     755.07       26.1
   y11++     762.11       80.2
   b11++     768.77       45.5
   a18+++    769.97      363.3
   a5+       774.55       60.1
   b18+++    778.64       29.3
   b5+       802.55       19.2
   b19+++    811.44       19.6
   a12++     812.35       33.6
   y12++     819.71       58.4
   y19+++    820.65       34.2
   b12++     826.55       49.4
   a20+++    839.47       16.2
   y20+++    843.67       22.4
   y13++     884.20       73.6
   a13++     892.43       18.1
   b21+++    901.34       26.0
   b13++     906.54       44.6
   b6+       917.53       50.5
   y6+       936.70       22.2
   a14++     948.76       82.0
   b14++     963.07       37.3
   b15++    1006.89       23.6
   a17++    1105.77       21.5
   b17++    1119.72       90.7
```

```
379. MORG-339_phosphopeptide_TMT_10%ACN_032017_OT.7225.7225.3 (eAGLsQSHDDLSNTTATPSVR)

   Ion         m/z   Intensity
   y4+       458.30       39.1
   y5+       559.40       35.0
   b8++      560.49       21.2
   b4+       600.45       18.4
   b9++      617.96       29.7
   y6+       630.43       26.1
   y12++     631.49       12.0
   b10++     675.49       38.3
   a11++     717.58       30.3
   b11++     731.66       47.2
   y14++     757.59       26.9
   b5+       767.49       18.1
   b12++     775.63       19.5
   a13++     818.57       17.6
   a14++     868.51       12.1
   b14++     882.94       22.9
   b15++     933.58       13.8
   y9+       946.56       36.3
   y17++     948.54       20.3
   b16++     969.04       26.0
   b17++    1019.56       30.0
   y10+     1033.53       48.1
   a20++    1146.65       27.4
   b20++    1161.18       11.9
   y12+     1261.59       27.7
   y13+     1376.65       22.4
```

```
380. MORG-339_phosphopeptide_TMT_14%ACN_032017_OT.11655.11655.4 (wAHDkFsGEEGEIEDDESGtENR)

   Ion         m/z   Intensity
   b6+++     415.29       11.6
   b10+++    576.15       18.3
   y10++     616.41       13.2
   b12+++    638.10       28.7
   a13+++    666.60       15.2
   y11++     672.98       18.1
   b13+++    675.84       34.0
   b14+++    718.94       43.4
   y18+++    720.97       14.7
   b4+       739.68       14.7
   y6+       743.21       19.4
   a15+++    747.36       13.4
   b15+++    757.34       36.2
   y13++     765.90       30.8
   a9++      784.90       16.5
   a16+++    785.88       54.1
   b16+++    795.38       39.9
   b9++      798.63       25.9
   a17+++    829.18       14.3
   b17+++    838.68       29.8
   b10++     863.21       15.3
   y7+       872.44       17.0
   a19+++    876.86       11.3
   b11++     892.11       16.7
   y21+++    923.27       14.3
   y22+++    947.42       16.3
   b12++     956.78       20.1
   y8+       987.41       14.7
   b21+++    989.72       21.4
   b13++    1013.21       24.4
   b22+++   1027.77       20.2
   y9+      1102.38       24.4
   b15++    1134.64       12.0
   b16++    1192.84       17.9
   y10+     1231.45       17.3
```

```
381. MORG-339_phosphopeptide_TMT_10%ACN_032017_OT.11607.11607.2 (sPsPEPIYNSEGk)

   Ion         m/z   Intensity
   b1+       397.24       21.0
   y2+       433.34       14.4
   y5+       763.43       14.7
   a5+       859.11       12.0
   b5+       887.31       26.8
   y6+       926.60       25.1
   a6+       956.35       18.5
   b6+       984.55       23.4
   a7+      1069.34       12.7
   b7+      1097.60       15.9
   y8+      1136.65       35.2
   b8+      1260.60       16.9
   y10+     1362.75       51.5
   b9+      1374.60       11.3
   y11+     1529.80       19.1
   y12+     1626.81       31.5
```

```
382. MORG-339_phosphopeptide_TMT_25%ACN_032017_OT.21175.21175.3 (aVRAAGPGNLVLPGsLTAsLEcVk)

   Ion         m/z   Intensity
   y1+       376.34       17.8
   b9++      512.19       18.8
   a10++     555.00       19.3
   b10++     569.05       42.3
   b11++     618.56       62.0
   y3+       635.42       20.6
   a12++     661.20       17.2
   a5+       670.68       14.6
   b12++     675.15       84.7
   b5+       698.51       19.6
   b6+       755.44       15.2
   y4+       764.38       23.2
   y11++     777.13       41.7
   y12++     826.08       29.2
   y5+       877.43       25.5
   y13++     882.64       23.4
   b17++     942.38       18.5
   a18++     963.73       21.9
   b18++     977.63       29.6
   b9+      1023.85       19.8
   y6+      1044.52       22.1
   y17++    1073.80       15.5
   a10+     1108.47       15.8
   y18++    1122.75       14.8
   b10+     1136.68       17.5
   y19++    1151.35       23.7
   b21++    1182.74       26.1
   y8+      1216.67       18.7
   y21++    1222.49       17.4
   b11+     1235.72       33.2
   b23++    1312.49       20.1
   y9+      1329.63       15.7
   b12+     1348.85       51.4
   y23++    1349.89       41.4
   y11+     1553.79       35.2
   y12+     1650.85       35.7
   y13+     1763.96       21.9
```

```
383. MORG-339_phosphopeptide_TMT_4%ACN_032017_OT.4918.4918.2 (sGDETPGSEAPGDk)

   Ion         m/z   Intensity
   y1+       376.36       19.2
   y2+       491.31        6.0
   y3+       548.34        6.1
   b3+       569.23       14.7
   y4+       645.45       26.4
   b4+       698.40       16.8
   y5+       716.44       22.2
   b5+       799.49       11.0
   y6+       845.51        9.5
   a6+       868.57       11.4
   b6+       896.45        8.2
   a7+       925.51       20.5
   y7+       932.29       10.5
   b7+       953.45        5.8
   y8+       989.61        9.6
   y9+      1086.54       36.3
   b9+      1169.39       16.1
   y10+     1187.57       18.1
   b10+     1240.52       20.5
   y11+     1316.65       20.5
   y13+     1488.81       13.0
   b13+     1509.69       23.4
```

```
384. MORG-339_phosphopeptide_TMT_6%ACN_032017_OT.5081.5081.3 (eGDGSATTDAAPATsPk)

   Ion         m/z   Intensity
   y3++      321.16       12.5
   y4++      371.45        8.8
   y5++      406.34       10.5
   b7++      424.30        8.2
   y6++      455.31       51.5
   a8++      460.35        8.1
   y2+       473.39       39.4
   b8++      474.42       15.3
   y7++      490.79       16.1
   a3+       502.90        7.5
   y8++      526.57        7.1
   b9++      532.47       14.7
   a4+       560.09       10.8
   b10++     567.82       45.8
   y9++      583.93        8.0
   b11++     603.38       54.9
   y10++     634.41        7.8
   y3+       640.08       26.1
   b12++     651.81       15.5
   a13++     672.84       37.5
   b5+       675.08       13.0
   y11++     684.89       15.2
   b13++     687.42       32.7
   b14++     737.89       19.2
   y4+       741.45       15.8
   b6+       746.36        7.8
   y13++     763.55       47.5
   y5+       812.57       10.0
   b15++     821.40       12.4
   b7+       847.60       14.1
   y6+       909.49       53.0
   b8+       948.40        7.0
   y7+       980.49       10.1
   y8+      1051.71        8.1
   b9+      1063.52       12.4
   b10+     1134.56       18.5
   b11+     1205.53       55.4
   b13+     1373.71        9.3
```

```
385. MORG-339_phosphopeptide_TMT_8%ACN_032017_OT.4620.4620.3 (sRTASGSsVtSLEGTR)

   Ion         m/z   Intensity
   y3+       333.24       14.3
   b6++      395.29       10.6
   b7++      438.92       12.9
   y4+       462.32       13.4
   b2+       473.35       25.1
   a3+       546.11        7.8
   a9++      557.51        7.0
   b9++      572.10       11.7
   y5+       575.38       18.3
   y10++     598.92       16.2
   a4+       617.76       43.3
   y11++     627.41       10.0
   b4+       645.35       16.3
   a10++     648.66       15.7
   a5+       704.72        9.1
   a6+       760.96        9.3
   b6+       789.63        8.0
   b13++     826.99       23.4
   y7+       843.35       16.9
   b14++     855.28       10.0
   y8+       942.39       16.9
```

```
386. MORG-339_phosphopeptide_TMT_22%ACN_032017_OT.19868.19868.4 (kTSFDQDsDVDIFPSDFTSEPPALPR)

   Ion         m/z   Intensity
   y2+       272.12       20.4
   y6++      325.76       37.4
   y3+       385.38       22.5
   y4+       456.41       15.9
   y5+       553.46       33.7
   b12+++    554.36       13.6
   y6+       650.47      109.3
   y13++     707.60       50.4
   y7+       779.46       42.3
   y14++     780.72       38.3
   b12++     830.76       14.1
   y15++     837.81       16.0
   y22+++    843.28       19.2
   a20+++    848.27       20.8
   a13++     890.00       23.7
   a22+++    913.50       15.7
   b22+++    922.41       29.8
   a23+++    936.53       25.9
   b23+++    945.66      153.7
   y9+       967.52       34.7
   a7+      1023.66       14.1
   y10+     1114.51       18.4
   y20++    1142.53       16.9
   y13+     1413.77       23.1
```

```
387. MORG-339_phosphopeptide_TMT_14%ACN_032017_OT.13746.13746.3 (skTPPSNLsPIEDAsPTEELR)

   Ion         m/z   Intensity
   b3++      388.63       11.9
   y3+       417.35       14.0
   b7++      586.17       11.0
   b8++      642.55       21.0
   y11++     670.08       13.0
   b2+       674.67       12.1
   y12++     719.09       11.2
   y6+       744.42       26.3
   b3+       775.35       16.0
   b11++     831.03       20.8
   y14++     858.83       11.2
   b4+       872.80       13.9
   b12++     895.37       23.5
   y7+       911.34       17.3
   y15++     915.65       12.1
   a13++     938.70       21.5
   a5+       942.12       23.1
   b13++     953.15       36.0
   a14++     974.48       20.2
   y8+       982.57       22.1
   b14++     988.90       20.8
   y17++    1007.85       10.8
   b6+      1056.61       15.5
   y9+      1097.49       19.6
   y19++    1106.75       12.8
   a18++    1221.41       18.7
   y10+     1226.50       15.6
   b18++    1235.94       21.0
   y11+     1339.71       14.5
   b20++    1357.03       20.0
   y12+     1436.59       27.1
```

```
388. MORG-339_phosphopeptide_TMT_4%ACN_032017_OT.11419.11419.3 (vLsDsEEEEk)

   Ion         m/z   Intensity
   y3++      317.73       11.4
   y1+       376.27       22.2
   y4++      382.23        7.2
   b2+       442.39        6.5
   y2+       505.39       24.5
   b6++      511.18        5.4
   y6++      529.73       19.5
   b7++      575.22       17.5
   a8++      625.82        4.2
   y3+       634.42       15.2
   b8++      639.80       24.4
   a9++      690.29        4.2
   b9++      704.29       21.2
   b4+       724.46        3.0
   b5+       891.32        6.4
   b6+      1020.50        8.3
   b7+      1149.43       14.2
   b8+      1278.54        5.0
```

```
389. MORG-339_phosphopeptide_TMT_4%ACN_032017_OT.12812.12812.4 (qADVADQQTTELPAENGETENQsPAsEEEkEAksD)

   Ion         m/z   Intensity
   b8++      543.52       37.0
   b3+       544.39       43.2
   b9++      594.07       56.6
   b4+       643.48       45.6
   b10++     644.53       30.4
   a18+++    700.14       86.1
   b11++     709.17       34.2
   b5+       714.54       51.1
   y4+       729.44       26.6
   b12++     765.68       48.6
   b6+       829.51       57.5
   y10++     885.57       24.8
   y11++     920.80       35.9
   y19+++    921.68       28.2
   b7+       957.61       50.1
   y20+++    959.59       25.1
   a25+++    969.47       39.6
   y13++    1052.99       57.1
   b8+      1085.59       55.0
   y14++    1116.87       32.0
   y25+++   1139.52       30.1
   b9+      1186.62       38.5
   y16++    1238.76       30.8
   b10+     1287.60       32.7
   y17++    1288.90       48.0
   y7+      1344.85       33.0
   y31+++   1353.88       25.7
   y19++    1382.38       78.8
   b23++    1383.30       51.4
   b11+     1416.75       69.0
   a24++    1417.75       61.1
   y20++    1438.92       24.4
   b12+     1529.85      109.1
   y22++    1539.45       87.1
```

```
390. MORG-339_phosphopeptide_TMT_10%ACN_032017_OT.9887.9887.3 (rVsVcAETFNPDEEEEDNDPR)

   Ion         m/z   Intensity
   y2+       272.23       30.8
   b6++      491.96       18.2
   b7++      556.22       18.0
   a9++      666.77       15.9
   y11++     672.54       18.4
   b9++      680.56       18.2
   a4+       723.31       14.0
   y12++     729.90       13.3
   b10++     737.65       51.5
   y6+       745.39       14.2
   b4+       751.49       19.5
   b11++     786.24       16.5
   b12++     843.68       42.3
   y7+       874.32       27.9
   a5+       883.34       14.1
   b13++     907.57      101.0
   b5+       911.55       13.1
   y15++     918.74       18.3
   b14++     972.40       16.4
   b6+       982.51       18.7
   y8+      1003.50       17.6
   b15++    1037.25       21.3
   a16++    1087.33       16.6
   b16++    1101.38       16.3
   b7+      1111.53       17.4
   y9+      1132.51       25.0
   b17++    1158.92       18.2
   y20++    1216.11       44.0
   y10+     1247.57       14.2
   b19++    1273.84      161.8
   y11+     1344.53       43.8
   b9+      1359.74       17.1
   y12+     1458.56       26.1
   b10+     1473.75       18.2
   y14+     1706.72       16.5
```

```
391. MORG-339_phosphopeptide_TMT_10%ACN_032017_OT.4790.4790.3 (rNsLDVSQEEk)

   Ion         m/z   Intensity
   y3++      317.25       12.9
   a4++      376.41       14.5
   b5++      448.28       36.8
   y6++      474.44        8.8
   a6++      483.83       19.5
   b6++      497.85       30.9
   y2+       505.35       18.3
   y7++      532.25       17.7
   b7++      541.43       11.7
   y8++      588.37       22.3
   a8++      591.22       27.6
   b8++      605.10       33.6
   y3+       634.48       21.2
   a9++      655.65        8.1
   b3+       667.39       10.9
   b9++      669.99       21.9
   a10++     720.15        9.0
   y10++     728.94        9.8
   b10++     734.26       16.6
   y4+       762.52       19.0
   y5+       849.47       27.0
   y6+       948.47       23.0
   b6+       994.56        7.9
   y7+      1063.60       14.9
```

```
392. MORG-339_phosphopeptide_TMT_60%ACN_032017_OT.8197.8197.4 (iQGSPTPASkkVPLPGPGsPEVk)

   Ion         m/z   Intensity
   y1+       376.39       21.2
   y2+       475.39       25.9
   y8++      539.93       39.6
   y9++      588.96       26.6
   b11+++    595.14       25.5
   b4+       615.58       18.5
   a12+++    618.89       21.5
   b12+++    628.21       52.8
   a14+++    689.05       38.9
   y11++     693.88       16.7
   b14+++    698.35       94.6
   a10++     699.39       24.4
   y4+       701.53       76.1
   a15+++    720.84       17.3
   a16+++    740.41       20.7
   y12++     743.70       16.4
   b16+++    749.60       48.1
   b17+++    782.29       26.1
   a6+       785.76       19.5
   y16+++    786.95       27.7
   a18+++    791.82       27.8
   b18+++    801.19       43.4
   y18+++    852.55       17.8
   b19+++    856.59       27.1
   a20+++    879.40       17.1
   y19+++    885.41       19.2
   b11++     892.08       29.1
   b7+       910.56       21.0
   a21+++    922.43       16.8
   y6+       925.07       18.8
   a12++     927.39       24.5
   b21+++    932.26       48.5
   y21+++    933.27       23.2
   b12++     941.73       59.4
   a8+       953.67       17.6
   a22+++    955.67       20.4
   b22+++    965.06       36.8
   b8+       981.51       29.0
   b13++     990.03       16.1
   y7+      1022.59       21.5
   a14++    1032.95       34.3
   b14++    1046.87       80.6
   y8+      1079.65       26.9
   b15++    1095.35       17.2
   b16++    1123.63       19.6
   a17++    1158.50       21.7
   y9+      1176.59       25.5
```

```
393. MORG-339_phosphopeptide_TMT_6%ACN_032017_OT.4077.4077.3 (rsPsPSPTPEAk)

   Ion         m/z   Intensity
   y4++      337.21       12.0
   y1+       376.28        7.7
   y6++      436.39       13.2
   a5++      443.87        9.5
   y2+       447.30       13.0
   y7++      480.07        6.6
   b6++      501.29       14.4
   y8++      528.47       18.9
   b7++      550.12       10.1
   b2+       553.34       13.6
   y3+       576.56       12.4
   a8++      586.12       34.4
   b8++      600.40       28.2
   y9++      611.42       17.5
   y10++     660.47       22.0
   y4+       673.43       30.4
   b10++     713.53       16.5
   b11++     749.16        9.8
   y5+       774.48       12.9
   b4+       817.43        7.2
   y6+       871.50       33.0
   b6+      1001.44       10.6
   y8+      1055.59        8.6
```

```
394. MORG-339_phosphopeptide_TMT_16%ACN_032017_OT.5064.5064.3 (sSSFSEEkGEsDDEkPR)

   Ion         m/z   Intensity
   y5++      437.62       12.7
   y3+       629.53       11.5
   y9++      670.93       12.1
   b9++      699.44        9.9
   y4+       758.57       23.1
   b10++     764.17       21.7
   a11++     833.49       12.1
   y10++     850.07       17.4
   b6+       854.62       19.4
   y5+       873.60       31.7
   y11++     914.10       20.3
   b13++     962.64       18.7
   y12++     978.54       16.0
   b7+       983.03       14.8
   y6+       988.55       18.6
   b14++    1026.57       11.4
   b15++    1205.52       19.4
```

```
395. MORG-339_phosphopeptide_TMT_14%ACN_032017_OT.8376.8376.2 (sVsNEGLTLNNSR)

   Ion         m/z   Intensity
   b1+       317.25       14.2
   y4+       490.30       10.0
   b4+       697.56        9.1
   y6+       704.45       19.1
   a5+       797.94       13.0
   y7+       817.70       27.3
   b5+       826.22       35.0
   y8+       874.54       23.9
   b6+       883.80        9.0
   a7+       968.22       17.8
   b7+       996.54       15.5
   y9+      1003.61       14.3
   y10+     1117.57       30.2
   b9+      1210.49        9.4
   y11+     1284.64        9.6
```

```
396. MORG-339_phosphopeptide_TMT_10%ACN_032017_OT.1735.1735.4 (nTEEPSsPVRkESSR)

   Ion         m/z   Intensity
   y6+++     331.36       23.2
   y3+       349.19       16.2
   b4++      352.08       17.0
   y8+++     396.69       79.5
   y5++      418.58       18.8
   b2+       445.42       15.0
   y9+++     452.46       14.6
   y4+       478.34       12.0
   y10+++    481.12       92.0
   y6++      496.73       12.8
   a7++      513.77       79.7
   b7++      527.71       30.8
   a3+       546.16       28.9
   b3+       574.36       36.6
   b11+++    588.78       14.8
   a12+++    622.74       14.0
   b9++      625.33       22.9
   b12+++    631.58       21.0
   y14+++    633.45       14.1
   a13+++    651.13       14.4
   a4+       675.36       15.6
   y9++      677.62       16.2
   a10++     690.29       29.8
   b10++     703.41       74.5
   y10++     721.60       19.4
   y11++     770.13       33.2
```

```
397. MORG-339_phosphopeptide_TMT_16%ACN_032017_OT.6468.6468.3 (lRRtESVPSDINNPVDR)

   Ion         m/z   Intensity
   y4+       486.35       17.8
   a7++      562.48       14.8
   b7++      576.17       16.3
   a10++     712.14       23.5
   y6+       714.45       17.5
   y13++     721.21       61.3
   b10++     726.36       69.4
   b11++     782.45       12.4
   y7+       827.53       36.6
   b12++     839.71       19.1
   y15++     890.06       14.0
   b13++     896.69       21.1
   y8+       942.31       14.6
   b5+       965.18       13.6
   a15++     980.96       16.5
   b15++     994.41       13.9
   b6+      1052.30       46.7
   y10+     1126.52       38.2
   b7+      1151.67       12.7
   y11+     1225.63       16.0
```

```
398. MORG-339_phosphopeptide_TMT_25%ACN_032017_OT.8164.8164.4 (aASVQPATASkSPVPSQQAsPk)

   Ion         m/z   Intensity
   y5++      420.26       26.2
   b7++      427.51       23.1
   y2+       473.38       52.1
   y11+++    478.61       31.4
   y6++      484.62       26.1
   b12+++    520.02       28.9
   y7++      527.44       35.1
   b4+       558.05       21.7
   y8++      576.48      118.4
   b14+++    585.71       33.7
   y13+++    627.42       26.6
   a16+++    637.79       20.5
   y3+       640.42       53.1
   b16+++    646.25       21.9
   y10++     674.60       46.5
   a17+++    680.31       29.8
   y15+++    684.56       30.8
   b5+       686.46       59.6
   b17+++    689.54       30.7
   y16+++    707.93       53.9
   y4+       711.06       76.1
   y11++     718.10       69.9
   b11++     736.14       65.0
   y17+++    740.68       55.1
   a6+       755.19       20.0
   b19+++    756.03       83.3
   b12++     779.27       69.3
   b6+       783.33       28.2
   b20+++    811.50       20.6
   a7+       826.56       26.1
   y5+       839.63       21.1
   b21+++    843.51       34.9
   b7+       854.47       34.1
   y21+++    869.29       40.3
   b14++     877.80      173.1
   y12++     896.70       38.1
   y13++     940.32       49.2
   y14++     975.73       30.6
   y15++    1025.84       23.1
   b9+      1026.60       35.5
   b17++    1033.90       39.0
   y17++    1110.37       37.3
   y8+      1151.65       43.1
   y10+     1347.76       22.2
   y11+     1434.75       21.6
```

```
399. MORG-339_phosphopeptide_TMT_10%ACN_032017_OT.5536.5536.3 (aETSESSGSAPAVPEAsASPkQR)

   Ion         m/z   Intensity
   y4++      379.36       22.3
   b2+       430.41       13.4
   b3+       531.54        9.2
   y8++      577.68       10.1
   y9++      641.89       24.8
   b12++     653.10       10.2
   y10++     690.51       26.5
   b13++     702.08       11.4
   a5+       719.20        8.8
   y11++     740.18       21.9
   b5+       747.39       12.6
   y12++     775.16       18.3
   y13++     824.21       23.2
   y14++     859.54       15.8
   a7+       893.84       16.5
   y15++     903.00       30.5
   y6+       915.43       19.6
   b7+       921.56        8.7
   y17++     975.23       14.3
   a9+      1037.32       12.5
   b9+      1065.55       14.9
   y19++    1082.81       10.4
   b10+     1136.51       27.1
   y9+      1282.77       21.1
   b22++    1304.94       22.2
   y10+     1379.74       24.3
   b13+     1403.66       25.3
```

```
400. MORG-339_phosphopeptide_TMT_10%ACN_032017_OT.4288.4288.2 (sVssPRTk)

   Ion         m/z   Intensity
   y3+       633.52       27.4
   a3+       635.58       12.1
   b3+       663.20        9.3
   y4+       730.63       39.1
   y5+       897.46       15.5
   a5+       899.58        9.7
   b5+       927.34        7.7
   y6+      1064.78        8.3
   b6+      1083.36        9.2
```

```
401. MORG-339_phosphopeptide_TMT_14%ACN_032017_OT.12600.12600.4 (eRLkGQEDSLASAVDATTGQEAcDsD)

   Ion         m/z   Intensity
   y3+       416.13       14.5
   b9+++     501.37       20.0
   a10+++    529.88       17.8
   b10+++    539.05       36.8
   b11+++    562.79       56.1
   y4+       576.22       22.7
   b12+++    591.88       47.0
   b13+++    615.54       54.2
   a14+++    639.05       17.4
   y5+       647.14       25.1
   b14+++    648.59      119.8
   b7++      650.70       18.9
   y12++     674.73       22.3
   b15+++    686.91       64.7
   b16+++    710.56       87.0
   a9++      737.90       25.2
   b17+++    744.26       90.8
   b9++      751.71       16.8
   y14++     760.28       14.3
   y22+++    769.55       14.7
   y6+       776.18       27.2
   b18+++    778.02       68.1
   a19+++    787.65       24.2
   a10++     793.85       26.8
   b19+++    797.07       43.4
   y15++     803.89       22.0
   b20+++    839.35       35.7
   b21+++    882.75       61.8
   y7+       904.22       17.5
   b22+++    906.42       64.1
   b13++     922.46       15.7
   b23+++    959.77       37.9
   y8+       961.31       34.3
   b24+++    997.99       27.7
   y9+      1062.39       40.7
   b16++    1065.45       16.3
   y10+     1163.33       45.7
   y11+     1234.40       29.4
   y23++    1332.55       15.7
   y12+     1349.41       47.0
   y13+     1448.57       23.8
   y15+     1606.57       15.5
   y16+     1677.75       13.3
```

```
402. MORG-339_phosphopeptide_TMT_16%ACN_032017_OT.5718.5718.4 (nSTDLDSAPEDPtSPkRk)

   Ion         m/z   Intensity
   y4+++     329.66       22.8
   b5++      380.73       13.9
   y6+++     418.77       41.1
   b6++      438.38       13.0
   y7+++     451.38       46.2
   a12+++    481.98       20.8
   y8+++     489.82       23.4
   y4++      494.09       20.2
   b8++      517.60       18.1
   y9+++     532.50       46.1
   y10+++    565.17       67.4
   b9++      566.07       16.3
   y11+++    588.86       19.7
   a10++     616.06       26.1
   y12+++    617.78       36.6
   a4+       619.32       12.9
   y6++      627.49       29.0
   b10++     630.21       15.5
   b4+       647.29       45.8
   y13+++    655.75       27.0
   a11++     673.69       38.1
   y14+++    694.06       13.1
   a16+++    722.95       13.4
   a5+       731.94       12.4
   y8++      734.16       21.4
   b5+       760.51       25.2
   b17+++    783.93       15.8
   y9++      798.20       53.4
   a13++     813.04       12.7
   a6+       847.19       85.0
   b6+       875.43       28.1
   y11++     882.67       18.2
   y12++     926.09       19.5
   b7+       962.55       18.5
   y13++     983.72       17.6
   b8+      1033.51       44.4
   b10+     1259.60       13.5
   b11+     1374.60       28.2
```

```
403. MORG-339_phosphopeptide_TMT_22%ACN_032017_OT.16414.16414.4 (eLEkRAsGQAFELILsPR)

   Ion         m/z   Intensity
   y2+       272.16       18.4
   b5+++     372.39        7.2
   y10+++    419.00       21.4
   y3+       439.36        9.9
   b7+++     451.85        9.1
   y7++      454.30       13.3
   b8+++     470.74        7.4
   a9+++     504.31       16.9
   b9+++     513.42       22.4
   b10+++    537.27       22.4
   y4+       552.39       12.8
   b5++      558.18        9.4
   y14+++    568.56        7.8
   a11+++    576.68       19.5
   b11+++    586.27       22.4
   b3+       601.46        9.8
   a12+++    619.92       12.5
   y10++     627.02       16.1
   b12+++    629.15       40.1
   y11++     655.56       17.1
   a13+++    657.75       28.9
   y15+++    688.39       10.0
   a14+++    695.26       12.2
   b14+++    704.69       33.4
   b8++      705.72        9.6
   a15+++    732.72       32.6
   b15+++    742.26       12.6
   b9++      769.73        8.1
   y6+       778.50       13.2
   b16+++    798.17       10.8
   b10++     804.84       10.6
   a11++     864.14        7.6
   b11++     878.56        9.6
   y7+       907.52        8.9
   b12++     943.25       10.9
   b13++     999.81       13.1
   y8+      1054.63       11.4
   b14++    1055.63       10.1
   y9+      1125.55       11.4
```

```
404. MORG-339_phosphopeptide_TMT_8%ACN_032017_OT.1236.1236.3 (rLETEDTSGSPSRAsR)

   Ion         m/z   Intensity
   b3++      314.92        9.9
   b5++      429.80       10.1
   b6++      487.33       47.4
   y9++      492.82       12.6
   y10++     543.51       23.1
   b9++      610.28        9.8
   b3+       628.65       17.9
   a10++     639.67       57.5
   b10++     653.42       38.0
   y5+       655.89       30.2
   y12++     664.92       17.1
   b11++     702.06       11.4
   y13++     716.21        8.7
   a12++     731.78       13.8
   y15++     836.96       10.3
   y7+       840.33       10.2
   b14++     858.60       18.5
   y8+       897.36        8.6
   b15++     942.61       11.8
   b6+       973.60       26.7
   b7+      1074.63       10.8
   y10+     1085.59       27.8
   y11+     1200.51       11.8
```

```
405. MORG-339_phosphopeptide_TMT_20%ACN_032017_OT.10596.10596.4 (dAsPPEPAsPTIGLDkkTR)

   Ion         m/z   Intensity
   a3++      277.84       20.3
   y4+++     331.13       17.5
   a9+++     404.68       21.0
   y7+++     426.17       20.0
   b6++      453.58       16.6
   y4++      496.11       15.5
   a8++      524.11       14.7
   y10+++    529.89       51.4
   b3+       583.27       21.8
   y11+++    585.58       13.9
   y12+++    608.97       67.6
   y6++      609.90       22.8
   b9++      620.84       17.0
   y7++      638.78       24.3
   y13+++    641.50       82.3
   a10++     655.78       13.7
   b10++     669.53       19.1
   b4+       679.86       21.8
   y14+++    684.41       48.0
   y8++      695.34       26.5
   y15+++    716.84       44.2
   y9++      745.57       18.2
   a5+       749.35       80.2
   y10++     794.07       35.4
   y18+++    828.87       21.6
   y11++     877.45       12.0
   a6+       878.29       13.4
   b6+       906.40       45.8
   y12++     912.79       62.0
   y13++     961.74       30.4
   y14++    1026.18       15.1
   y15++    1074.74       12.5
```

```
406. MORG-339_phosphopeptide_TMT_16%ACN_032017_OT.9744.9744.4 (kFPDRLADDEGDsDsESVGQSR)

   Ion         m/z   Intensity
   y6++      317.26       12.6
   y4+       447.30       15.8
   y5+       546.42       16.6
   a3+       574.51        8.6
   a14+++    615.35       11.9
   a9++      630.07       15.1
   y12++     692.13       14.0
   y18+++    695.10        8.6
   b10++     708.68       21.1
   a16+++    713.48       13.2
   b4+       717.87       15.6
   a11++     723.39       15.8
   y19+++    733.74      125.1
   b11++     737.46       69.3
   a12++     780.59       11.4
   b12++     794.83        8.7
   b19+++    803.51        9.5
   a5+       845.63       10.9
   y15++     871.56       13.9
   b13++     877.98       10.1
   y16++     907.58       10.4
   y8+       929.22       11.8
   a17++    1113.45        8.5
   y10+     1211.37       10.2
```

```
407. MORG-339_phosphopeptide_TMT_16%ACN_032017_OT.3835.3835.2 (kAGGANYDAQsE)

   Ion         m/z   Intensity
   y4+       514.31       12.4
   b4+       543.20        7.3
   y6+       792.41       11.7
   a7+       863.63       13.3
   y9+      1034.33        6.2
   y10+     1091.43        7.7
   y11+     1162.35       12.0
```

```
408. MORG-339_phosphopeptide_TMT_16%ACN_032017_OT.11013.11013.4 (tNNLQLHPHDAFNDEDADRLsEIsAR)

   Ion         m/z   Intensity
   y2+       246.13       18.9
   y3+       413.15       16.8
   a7++      511.79       19.1
   y4+       526.00       43.7
   y10++     639.58       27.8
   b9++      643.01       27.5
   b4+       672.42       17.0
   y11++     696.47       18.1
   b10++     700.58       32.3
   b11++     736.33       19.8
   y18+++    740.94       47.7
   y19+++    773.57       39.7
   a12++     795.26       41.8
   b5+       800.37       24.6
   b12++     809.90       27.2
   b19+++    811.40       24.4
   y13++     818.48      464.2
   y6+       822.32       28.5
   b21+++    905.56       18.3
   a14++     910.20       16.8
   b6+       913.74       27.8
   b14++     924.14       31.4
   y7+       935.24       18.3
   y23+++    937.40       16.3
   b22+++    947.80       25.9
   a23+++    976.47       68.8
   y16++     984.98       19.9
   b23+++    986.14       72.5
   b15++     988.39       24.6
   y17++    1042.09       20.1
   b7+      1050.85       34.5
   y8+      1091.07       14.8
   y18++    1111.31       33.4
   b18++    1139.21       50.5
   y19++    1159.75       31.0
```

```
409. MORG-339_phosphopeptide_TMT_18%ACN_032017_OT.7738.7738.3 (sRsLLEVPR)

   Ion         m/z   Intensity
   y1+       175.17        9.0
   a2++      223.14        6.2
   b2++      237.08       16.4
   y4++      250.21       14.3
   y2+       271.92       30.0
   a3++      306.50        9.6
   b1+       317.27       10.9
   b3++      320.75       26.5
   y3+       371.25       25.4
   b4++      377.16       16.2
   a5++      419.90       28.5
   b5++      433.83       36.0
   y7++      447.43       22.2
   b2+       473.35        8.1
   b6++      498.41       36.7
   y4+       500.33       36.4
   y8++      525.09       11.3
   a7++      533.82       13.3
   b7++      547.96       23.7
   b8++      596.50       11.8
   y5+       613.41       34.0
   y6+       726.48       29.6
```

```
410. MORG-339_phosphopeptide_TMT_10%ACN_032017_OT.7171.7171.3 (kGAEEEEEEEDDDsEEEIk)

   Ion         m/z   Intensity
   y1+       376.40       12.7
   y2+       489.45       16.6
   a8++      552.24       12.3
   a9++      616.59       17.1
   y3+       618.42       14.5
   a10++     681.62       15.4
   y4+       747.53       13.8
   b12++     810.15       27.0
   a13++     853.98       14.2
   b13++     867.72       35.3
   y5+       876.58       10.5
   a14++     937.63        9.6
   a7+       974.69       18.8
   y14++    1017.39       21.3
   y6+      1043.49       18.0
   y15++    1082.15       14.7
   b8+      1131.65       14.2
   b17++    1145.14       10.0
   y16++    1146.30       17.7
   y7+      1158.41       15.2
   a18++    1187.69       10.8
   y18++    1210.23       11.0
   b9+      1260.22       27.7
   y8+      1273.55       16.8
   y9+      1388.68       12.5
   b10+     1389.66       12.5
```

```
411. MORG-339_phosphopeptide_TMT_12%ACN_032017_OT.11097.11097.2 (sLsPLGGR)

   Ion         m/z   Intensity
   y1+       175.23       11.4
   y3+       289.15        9.5
   b1+       397.35       13.2
   y4+       402.24        6.7
   a2+       482.48        9.1
   y5+       499.33       52.8
   b2+       510.22        8.1
   y6+       666.36       11.5
   b3+       677.34        5.8
   y7+       779.45       10.6
   a5+       859.41        7.4
   b5+       887.42       19.2
   b6+       944.41       13.2
   b7+      1001.43        6.3
```

```
412. MORG-339_phosphopeptide_TMT_18%ACN_032017_OT.9592.9592.3 (aPTAALsPEPQDSkEDVk)

   Ion         m/z   Intensity
   y2+       475.35       20.2
   y6++      582.52       38.2
   y7++      640.27       12.0
   b12++     744.14       13.0
   y9++      752.69       44.5
   b6+       754.62       12.3
   y10++     817.19       18.5
   y11++     865.75       81.4
   y12++     949.16       12.1
   a14++     952.27       11.7
   b14++     966.15       11.4
   y13++    1005.83       12.3
   b15++    1031.38       11.4
   y14++    1040.71       12.6
   b16++    1088.83       26.8
   y16++    1127.27       31.3
   b17++    1137.72       12.7
   b9+      1147.54       20.6
   y17++    1175.97       21.0
   b12+     1487.74       13.7
```

```
413. MORG-339_phosphopeptide_TMT_22%ACN_032017_OT.10887.10887.4 (aAVGVTGNDITTPPNkEPPPsPEk)

   Ion         m/z   Intensity
   y5++      433.77       21.6
   b3+       471.40       21.8
   y6++      482.08       63.4
   a14+++    498.94       27.3
   b9++      508.04       39.6
   y7++      531.03      167.8
   a10++     550.51       23.3
   b10++     564.60       65.1
   y11+++    586.92       36.7
   y8++      595.38       24.8
   y3+       602.48       45.0
   b11++     615.14       49.7
   y12+++    619.37      136.3
   b5+       627.49       44.9
   y13+++    652.98       50.2
   a16+++    656.60      117.9
   b16+++    666.02       37.9
   y14+++    686.69       41.6
   b17+++    709.19       29.4
   y15+++    724.45       30.4
   b6+       728.51       33.7
   b18+++    741.52       26.2
   a7+       757.32       29.9
   a19+++    763.62       37.7
   y4+       769.35       33.0
   y9++      774.05       40.6
   y10++     831.14       23.8
   y11++     879.18       49.2
   b8+       899.85       20.6
   y12++     928.32      169.3
   y6+       963.51       24.8
   y13++     978.79       68.2
   b16++     997.74       19.1
   b9+      1014.66       39.5
   y14++    1029.35      101.2
   y7+      1060.57       38.0
   b17++    1062.91      209.4
   y15++    1085.86       30.2
   b18++    1110.96       19.1
   b10+     1127.61       29.2
   b11+     1228.77       19.0
   b12+     1329.78       54.3
```

```
414. MORG-339_phosphopeptide_TMT_10%ACN_032017_OT.12474.12474.4 (gADDAADADTAIINAEGGQNNsEEk)

   Ion         m/z   Intensity
   y3++      317.40       14.0
   a9+++     335.15       11.0
   y1+       376.33       12.4
   a14+++    505.47       11.1
   b9++      515.74       10.5
   y13+++    567.11       16.6
   b17+++    601.35        8.2
   b11++     602.46       19.7
   y8++      607.84        9.5
   y9++      636.54       19.8
   b12++     658.74       20.3
   b5+       659.40       17.7
   y16+++    661.83       11.5
   y10++     700.67       17.8
   a13++     701.55       13.3
   a6+       702.51       13.4
   b13++     715.28       20.9
   y18+++    724.17       16.4
   a21+++    729.48       23.9
   b6+       730.20       30.5
   y11++     736.59       29.6
   b21+++    738.86       18.8
   b14++     772.74       13.3
   a22+++    785.27       12.0
   y12++     793.31       18.0
   b22+++    793.99       11.1
   y4+       801.41        8.8
   b15++     808.22       21.6
   a7+       817.49       11.5
   y22+++    848.33       10.1
   y13++     850.00       25.6
   a16++     858.39        9.5
   a24+++    871.03        8.4
   a17++     887.01        8.6
   b17++     900.48       10.2
   y14++     906.68       20.5
   b8+       916.22       13.6
   b18++     929.46        9.3
   y15++     941.91        9.1
   b19++     993.08       13.5
   b20++    1050.48       16.5
   b21++    1107.95       10.3
   b11+     1203.38        9.1
   b12+     1316.68       11.4
```

```
415. MORG-339_phosphopeptide_TMT_30%ACN_032017_OT.15516.15516.4 (sksPPkVPIVIQDDSLPTGPPPQIR)

   Ion         m/z   Intensity
   b3++      421.42       16.4
   y9++      481.97       40.4
   y10++     538.42       15.0
   y5+       610.38       16.7
   b12+++    681.24       19.5
   y6+       707.50       41.0
   b13+++    720.21       27.6
   b7++      746.43       15.6
   b14+++    758.39       30.8
   y7+       764.49       31.2
   a15+++    777.52       13.9
   a8++      780.92       17.7
   b15+++    787.06       29.5
   b8++      794.64       20.8
   y20+++    800.38       17.7
   a3+       813.72       26.6
   a16+++    815.51       33.9
   y15++     816.98       19.8
   b16+++    825.17      105.0
   y21+++    833.16       21.2
   a9++      837.06       20.0
   b3+       841.61       28.2
   a17+++    847.44       19.4
   y8+       865.52       13.7
   y16++     866.57       16.3
   a10++     887.16       13.6
   b18+++    891.07       23.7
   a4+       910.21       58.6
   y23+++    921.06       15.4
   y17++     923.87       22.9
   a20+++    933.01       14.9
   b20+++    942.69       16.4
   a11++     943.64       18.8
   b11++     957.43       14.6
   y9+       962.55       71.0
   a21+++    965.69       17.7
   y18++     972.29       16.6
   b21+++    974.87       16.3
   b23+++   1049.65       15.0
   a24+++   1078.32       14.9
   b24+++   1086.86       13.9
   b16++    1236.58       36.4
   y22++    1297.66       14.1
   a6+      1364.96       18.8
```

```
416. MORG-339_phosphopeptide_TMT_16%ACN_032017_OT.8835.8835.3 (nPLGDDDAsATASk)

   Ion         m/z   Intensity
   b1+       344.23       10.6
   y1+       376.36       38.0
   y6++      437.31       10.0
   y2+       463.36       58.3
   a7++      464.31       34.2
   b7++      478.86        9.5
   b8++      514.53       16.9
   y8++      530.46       10.5
   y3+       534.36       42.1
   b3+       554.28       11.6
   y9++      587.44        8.7
   b9++      598.19       14.4
   b4+       611.51       18.7
   a10++     619.59       17.7
   a11++     670.06       22.4
   y11++     674.06       11.5
   b11++     684.07       67.5
   a5+       698.40       11.1
   a12++     705.15       15.3
   y5+       706.28       11.1
   b12++     719.59       42.8
   b5+       726.47       11.5
   a13++     749.14       13.2
   b13++     763.07       54.0
   y13++     779.08       11.4
   b6+       841.52        9.7
   b7+       956.43       12.6
   b8+      1027.50       12.6
   b10+     1265.57       10.4
```

```
417. MORG-339_phosphopeptide_TMT_25%ACN_032017_OT.13288.13288.4 (kEsPGRWGLGEDPTGVGPSLQcR)

   Ion         m/z   Intensity
   y6++      380.85       40.2
   y7++      409.38       39.6
   y3+       463.32       31.0
   b2+       487.50       22.3
   y4+       576.34       28.6
   y11++     586.43       54.4
   a16+++    649.75       33.4
   y5+       663.35       38.7
   a17+++    668.80       20.8
   b17+++    677.95       57.1
   y13++     708.06       23.9
   y20+++    713.48       26.0
   a4+       722.91       47.7
   a19+++    729.93       54.6
   y14++     737.16       54.0
   b19+++    739.24       49.5
   a11++     740.15       41.3
   b20+++    777.51       25.6
   a5+       780.73       21.9
   a21+++    810.34       32.7
   b12++     811.64       20.9
   y7+       817.42      104.1
   y16++     822.39       23.5
   a13++     845.83       22.0
   a22+++    864.00       25.1
   a14++     896.77       36.1
   y8+       916.59       42.1
   y9+       973.62       42.0
   a16++     974.61       30.8
   b17++    1017.42       23.2
   y20++    1070.33       75.3
   y10+     1074.53       23.3
   y21++    1153.62       29.0
   y11+     1171.60       62.1
```

```
418. MORG-339_phosphopeptide_TMT_18%ACN_032017_OT.17572.17572.3 (lGEDNINVVEGNEQFIsASk)

   Ion         m/z   Intensity
   y1+       376.31       29.9
   y2+       463.37       41.2
   b7++      493.43       13.0
   y3+       534.45       19.9
   b8++      543.01       26.0
   y7++      545.47       19.1
   b9++      592.57       20.9
   b4+       644.46       13.3
   b10++     656.76       13.1
   b11++     685.35       12.1
   y10++     695.63       27.0
   y4+       701.36       52.1
   b5+       758.47       18.1
   y11++     759.79       23.7
   b13++     806.89       27.5
   y12++     809.34       25.6
   y5+       814.54       27.2
   a14++     856.88       27.9
   y13++     858.47       19.7
   b6+       871.41       43.8
   y14++     916.38       13.1
   b15++     944.77       96.2
   y6+       961.50       29.4
   b7+       985.58       26.4
   a16++     986.58       20.8
   b16++    1001.39       64.8
   b8+      1084.62       45.0
   y17++    1086.67       18.3
   a18++    1105.90       20.3
   a9+      1155.65       22.9
   b19++    1163.81       27.2
   b9+      1183.69       45.0
   b10+     1312.72       37.1
   b11+     1369.76       17.2
   y10+     1389.67       14.3
   b12+     1483.89       17.8
   b13+     1612.90       24.1
   b14+     1740.93       13.8
```

```
419. MORG-339_phosphopeptide_TMT_16%ACN_032017_OT.4397.4397.3 (tGSQHGPQNAAAATFQR)

   Ion         m/z   Intensity
   a7++      473.44       13.8
   b8++      551.49       21.3
   y11++     587.79       25.0
   b9++      608.90       24.2
   y12++     616.61       27.3
   y5+       622.27       24.1
   a10++     630.00       18.5
   b10++     644.48       37.5
   a4+       655.41       38.6
   y6+       693.41       25.8
   b12++     715.69       21.9
   a13++     736.89       14.0
   y14++     748.83       16.1
   b13++     750.79       16.3
   y7+       764.39       30.6
   a14++     787.80       12.5
   y15++     792.58       17.7
   y16++     820.43       20.6
   y8+       835.48       22.5
   b6+       877.48       16.5
   y9+       949.50       24.6
   y11+     1174.60       18.2
   y12+     1231.71       17.1
```

```
420. MORG-339_phosphopeptide_TMT_4%ACN_032017_OT.2111.2111.3 (sAVRPsPsPER)

   Ion         m/z   Intensity
   y1+       175.12        5.8
   y2+       304.20        6.9
   a4++      308.17        7.2
   b1+       317.21        4.7
   b4++      322.22       23.8
   b2+       388.29        6.1
   y3+       401.29       23.9
   b6++      454.35       25.4
   a3+       459.42       34.8
   y7++      465.21       11.4
   b3+       487.37       13.9
   a7++      488.38       17.8
   y4+       568.47        8.7
   a9++      620.36        4.5
   b4+       643.52        7.3
   y5+       665.27       29.2
   a10++     685.40        6.1
   b10++     699.61       21.0
   y6+       832.37        4.7
   y7+       929.38       16.8
```

```
421. MORG-339_phosphopeptide_TMT_6%ACN_032017_OT.8607.8607.3 (sRtAsLTSAAsIDGSR)

   Ion         m/z   Intensity
   b1+       317.29        8.7
   y3+       319.22       18.2
   b4++      363.33       10.9
   y8++      428.32        8.6
   y4+       434.28       12.4
   b5++      446.73        7.0
   y10++     522.37        7.5
   y5+       547.26       18.0
   b7++      553.78       15.7
   y11++     579.62       19.2
   b8++      597.26        7.3
   a9++      618.43       75.6
   a3+       626.05        7.2
   b9++      632.85       15.1
   b3+       654.35       19.7
   y12++     662.79        7.5
   b10++     668.34       25.3
   y6+       714.35       17.4
   b11++     751.64       11.1
   y7+       785.35       20.6
   a12++     794.58       11.3
   b12++     807.95       11.3
   y8+       856.50       16.6
   b13++     865.91       28.1
   y15++     866.89       13.2
   b5+       892.24        7.8
   y9+       943.47       28.3
   y10+     1044.43       23.3
   y11+     1157.53       21.9
   b8+      1193.30        7.7
   y13+     1395.64       14.8
```

```
422. MORG-339_phosphopeptide_TMT_6%ACN_032017_OT.5138.5138.3 (eERAsSPPVEVGEPR)

   Ion         m/z   Intensity
   y2+       272.19       25.9
   b3++      323.14        6.4
   a5++      427.21       22.9
   y8++      441.84       12.2
   y4+       458.33        9.8
   a2+       460.29        8.7
   b6++      485.12        8.6
   y9++      490.25       30.4
   y5+       557.40       14.8
   a3+       616.55       20.9
   a9++      617.47       86.2
   b9++      631.95       22.7
   a10++     682.19        7.6
   y6+       686.37        8.5
   a4+       687.31        9.4
   b10++     696.41       17.0
   b4+       715.38        7.2
   b12++     774.72       12.4
   y7+       785.45       10.4
   a13++     824.84       12.0
   b13++     839.00       61.0
   a5+       854.57        8.1
   a14++     873.53       10.2
   y8+       882.46       12.9
   b14++     887.58        6.9
   b6+       969.40        9.6
   y9+       979.51       54.6
```

```
423. MORG-339_phosphopeptide_TMT_12%ACN_032017_OT.12448.12448.2 (sFDDPIVQTER)

   Ion         m/z   Intensity
   b1+       397.38        8.2
   y4+       533.27       10.0
   y5+       632.30        7.3
   b3+       659.29       13.7
   y6+       745.01        6.6
   a4+       745.97       12.2
   b4+       774.28       33.4
   y7+       842.51       42.8
   a5+       843.51       32.1
   y8+       957.51       20.3
   b6+       984.49        7.4
   y9+      1072.64        8.7
   b7+      1083.65       10.6
   b8+      1211.69        7.7
   y10+     1219.63       13.4
   b10+     1441.86        7.2
```

```
424. MORG-339_phosphopeptide_TMT_18%ACN_032017_OT.12540.12540.3 (aALsPARLGsPIRsPk)

   Ion         m/z   Intensity
   y5++      454.98       14.5
   y2+       473.37       28.2
   a7++      474.38       15.9
   b7++      488.86        9.7
   y6++      503.94       15.1
   b8++      545.52       10.4
   y3+       640.34       14.3
   b11++     705.61       35.0
   a5+       721.32       15.6
   a12++     748.15       19.4
   b5+       749.38       29.0
   y10++     750.64       28.3
   b12++     762.20       20.4
   y11++     785.68       25.3
   a6+       792.10       12.0
   b6+       820.55       12.3
   a13++     826.22       19.8
   y12++     834.73       35.2
   b13++     840.71       23.1
   y14++     974.73       10.6
   b7+       976.44       10.4
   y6+      1006.62       10.4
   y15++    1010.44        9.7
```

```
425. MORG-339_phosphopeptide_TMT_10%ACN_032017_OT.11445.11445.2 (sGEGEVSGLLR)

   Ion         m/z   Intensity
   b1+       397.22       12.3
   y4+       458.42       13.3
   y5+       545.37       17.1
   b3+       583.35       15.9
   b4+       640.04       16.5
   y6+       644.56       16.4
   b5+       769.35       17.1
   y8+       830.44       15.2
   b6+       868.44       10.1
   b8+      1012.59        9.8
   y10+     1016.48       21.3
   b9+      1125.60       14.5
   b10+     1238.77        9.6
```

```
426. MORG-339_phosphopeptide_TMT_14%ACN_032017_OT.5344.5344.3 (lRsDGAAIPR)

   Ion         m/z   Intensity
   b2++      250.21        9.1
   y2+       272.30        9.5
   a3++      319.74       13.4
   b3++      333.78       13.7
   b1+       342.87       17.7
   y3+       385.22       13.9
   b4++      391.22       38.8
   a5++      405.30        5.4
   b5++      419.77       25.3
   y8++      433.96        6.1
   a6++      441.41       21.0
   b7++      490.80       17.4
   y9++      512.17        8.5
   y5+       527.36       16.0
   a8++      533.32       15.3
   b8++      547.40       18.5
   a9++      581.72        6.9
   y6+       584.44       33.0
   b3+       666.41        5.2
   y7+       699.41       12.7
```

```
427. MORG-339_phosphopeptide_TMT_4%ACN_032017_OT.5823.5823.3 (sAEPTREPGAEAEsGSEsEPEPGPGPR)

   Ion         m/z   Intensity
   b6++      436.27        9.8
   b7++      500.91       19.3
   b3+       517.39        8.5
   y6+       580.38       17.2
   b10++     613.31        6.8
   b4+       614.26        8.1
   b11++     678.09       12.6
   y7+       709.49        8.4
   b12++     713.56       11.7
   b5+       714.90        8.3
   b13++     778.34       17.4
   y8+       806.51       18.9
   a15++     875.53        8.4
   y9+       935.42       12.6
   a17++     983.67       19.0
   b17++     998.19       12.9
   b7+      1000.54      129.6
   y20++    1049.32        7.8
   a8+      1069.33        5.6
   b8+      1097.61       19.6
   y10+     1102.50        9.5
   a20++    1180.42        5.9
   y23++    1241.61        6.1
   b21++    1258.98       12.3
   y24++    1290.26        8.8
   a23++    1321.78        7.7
   b23++    1336.52        9.2
   y13+     1375.51        9.5
   b25++    1413.46        7.9
   y19+     1999.81        5.8
```

```
428. MORG-339_phosphopeptide_TMT_22%ACN_032017_OT.12926.12926.3 (dksPSSLLEDAk)

   Ion         m/z   Intensity
   a1+       317.22       10.4
   b1+       345.19       12.4
   b2++      351.93       14.2
   y1+       376.35       12.4
   b3++      435.49       17.5
   y2+       447.42       32.3
   a4++      469.51       11.5
   b5++      527.26       12.4
   y3+       562.45       22.7
   b6++      570.73       12.2
   y9++      595.13       23.2
   a7++      613.55       16.4
   b7++      627.59       15.1
   a8++      670.25       20.0
   a2+       674.74       51.5
   y10++     677.92       28.5
   y4+       691.57       16.3
   a9++      734.52       13.2
   b9++      748.36       16.5
   y5+       804.59       23.2
   b10++     805.71       27.2
   a3+       841.69       14.8
   y11++     856.96       15.4
   y6+       917.56       15.6
   y7+      1004.67       18.6
   y9+      1188.70       30.1
```

```
429. MORG-339_phosphopeptide_TMT_16%ACN_032017_OT.18183.18183.2 (sLMssPEDLTk)

   Ion         m/z   Intensity
   b1+       317.30       12.9
   y1+       376.30       17.1
   b2+       430.42       14.2
   y2+       477.40       15.5
   b3+       561.43       16.6
   y3+       590.48       31.5
   a4+       700.38       18.1
   y4+       705.60       19.3
   y5+       834.64       14.4
   b5+       895.59       50.5
   y6+       931.56       49.0
   y7+      1098.57       20.9
   b7+      1121.42       11.1
   b8+      1236.50       29.8
   y8+      1265.52       13.6
   b9+      1349.55       15.0
   y9+      1396.69       10.3
   y10+     1509.65       10.1
```

```
430. MORG-339_phosphopeptide_TMT_22%ACN_032017_OT.9620.9620.3 (eksPELPEPSVRMk)

   Ion         m/z   Intensity
   y6++      473.99       39.0
   y8++      587.10       32.5
   b6++      612.18       19.7
   y9++      643.70       42.0
   b2+       716.88       15.4
   y11++     756.44       43.8
   a9++      759.31       23.6
   y4+       762.49       18.8
   b9++      773.38       44.1
   y5+       849.77       19.6
   a3+       855.35       16.4
   y6+       946.76       14.0
   b5+      1109.70       18.7
   y8+      1172.68       16.9
   b6+      1222.70       18.1
   b8+      1448.90       15.7
```

```
431. MORG-339_phosphopeptide_TMT_16%ACN_032017_OT.6540.6540.3 (tkPPPTYEsEEEDk)

   Ion         m/z   Intensity
   b2++      344.67       12.1
   y1+       376.37       14.0
   y2+       491.35        9.9
   y3+       620.45       15.7
   b7++      622.62       13.6
   b8++      686.97       22.6
   b2+       688.48       21.7
   y9++      719.35        9.9
   y4+       749.05       25.2
   a9++      756.21       11.8
   a3+       757.15       20.1
   y10++     768.13       49.8
   b9++      770.71       15.6
   b3+       785.68       30.1
   y11++     816.70       20.4
   y12++     865.60       21.1
   y5+       878.46       18.1
   a5+       951.43       10.0
   b12++     963.78       11.0
   b13++    1021.54       16.2
   y6+      1045.44        9.8
   y12+     1729.77        9.6
```

```
432. MORG-339_phosphopeptide_TMT_30%ACN_032017_OT.14912.14912.3 (sAVSSLRAFQTLAR)

   Ion         m/z   Intensity
   y4++      230.21       13.6
   b1+       397.27       19.2
   y4+       460.33       15.3
   y8++      481.87       13.4
   b7++      506.05       17.2
   y9++      538.60       16.6
   a3+       539.33       14.0
   b3+       567.39       75.2
   y10++     581.84       31.5
   y5+       588.36       27.9
   y11++     625.51       41.5
   a4+       626.41       12.3
   y12++     674.91       21.7
   b10++     678.90       23.8
   y13++     710.49       39.1
   a11++     715.33       13.5
   b11++     729.79       14.9
   y6+       735.46       21.7
   b12++     785.79       24.9
   y7+       806.55       24.9
   a13++     807.41       11.5
   b13++     821.52       34.9
   y8+       962.63       15.1
```

```
433. MORG-339_phosphopeptide_TMT_10%ACN_032017_OT.10157.10157.3 (sREEEWDPEYTPk)

   Ion         m/z   Intensity
   y1+       376.33       10.6
   b5++      470.79       11.3
   y2+       473.35       51.9
   y6++      482.23       33.0
   y7++      539.78        9.7
   b6++      563.66       18.5
   y3+       574.44       27.3
   b7++      621.31       65.0
   a8++      656.07       13.3
   b8++      669.67       15.6
   b3+       681.92       13.0
   a9++      720.46       27.0
   a10++     802.07       18.4
   b10++     816.01       31.5
   y5+       866.44       27.3
   y12++     904.47       12.2
   b5+       940.56       13.1
   y6+       963.54       75.1
   y7+      1078.62       16.9
   b6+      1126.45       14.2
   b7+      1241.57       35.7
```

```
434. MORG-339_phosphopeptide_TMT_12%ACN_032017_OT.5163.5163.3 (kEsGEGAEAEGATAEGAk)

   Ion         m/z   Intensity
   y1+       376.36       15.1
   y2+       447.35       16.2
   y3+       504.43       36.3
   a8++      535.42       19.2
   y4+       633.52       34.1
   a11++     664.13       36.3
   a12++     699.53       32.7
   y5+       704.49       29.2
   a13++     749.94       20.2
   y14++     760.87       20.5
   b13++     764.15       41.0
   a14++     785.71       20.6
   y15++     789.26       25.8
   b14++     799.77       38.7
   y6+       805.39       44.0
   a5+       812.49       17.4
   a6+       869.36       26.0
   y16++     872.18       17.9
   y7+       876.51       17.8
   a16++     878.66       17.3
   b16++     892.18       21.9
   a17++     914.21       25.8
   y8+       933.53       33.7
   b7+       968.05       14.0
   y9+      1062.59       19.2
   y10+     1133.61       25.4
   y13+     1390.81       15.4
```

```
435. MORG-339_phosphopeptide_TMT_14%ACN_032017_OT.2793.2793.3 (rRSESsGNLPSVADTR)

   Ion         m/z   Intensity
   y2+       276.16       52.7
   b6++      506.98       13.4
   y10++     515.01       11.5
   b8++      592.45       27.1
   y11++     598.56       13.5
   a3+       601.17       11.8
   b3+       629.41       12.2
   a9++      634.97       39.6
   y12++     642.53       53.3
   b9++      648.61      185.2
   b10++     697.76       12.8
   b11++     740.93       11.1
   y7+       745.41       41.3
   b4+       758.53       12.6
   b12++     790.40       13.2
   a5+       817.05       21.6
   b13++     826.15       28.7
   y8+       858.58       13.6
   b14++     883.64      137.6
   y10+     1029.60       14.4
   b9+      1296.69       12.2
```

```
436. MORG-339_phosphopeptide_TMT_16%ACN_032017_OT.10348.10348.2 (iVEFRAssR)

   Ion         m/z   Intensity
   b1+       343.42       12.2
   b2+       442.50       15.7
   b3+       571.44       13.2
   a4+       690.00       13.6
   b4+       718.42       35.8
   b5+       874.54       11.4
   y6+       883.47       13.9
   b6+       945.71       12.4
   a7+      1084.50        9.5
   y8+      1111.50       12.1
```

```
437. MORG-339_phosphopeptide_TMT_10%ACN_032017_OT.6775.6775.3 (eNPPsPPTSPAAPQPR)

   Ion         m/z   Intensity
   y4++      249.17        8.0
   y2+       272.20       12.8
   b1+       359.23        8.9
   y7++      368.94       21.5
   y8++      412.56       17.8
   b5++      417.40        9.6
   b2+       473.32       17.2
   y4+       497.31       36.3
   a3+       542.00       11.6
   a8++      551.18       18.0
   y11++     559.83       24.4
   b8++      565.25       15.7
   y5+       568.43       16.6
   b9++      608.59       16.2
   a4+       639.61       36.9
   a10++     642.95       25.0
   y13++     691.71       12.6
   b11++     693.15       22.7
   b12++     728.65       41.7
   y7+       736.42       35.0
   y14++     740.29       16.1
   b13++     776.72        8.3
   y8+       823.49       15.5
   b5+       834.49       13.6
   b14++     841.14       17.0
   y9+       924.63        7.7
   y10+     1021.45        8.7
   b8+      1129.58       12.3
   b9+      1216.51       14.9
   b12+     1455.73       10.0
```

```
438. MORG-339_phosphopeptide_TMT_10%ACN_032017_OT.6241.6241.3 (rGsLsPPSSAYER)

   Ion         m/z   Intensity
   b3++      305.71       13.0
   y5++      313.22       33.9
   a4++      347.76       50.6
   y6++      356.79       19.2
   y7++      405.36       13.2
   a2+       415.18       12.6
   b5++      445.52       28.9
   y8++      453.84       96.0
   y3+       467.36       12.5
   b6++      493.95       12.7
   a7++      528.26       27.0
   y4+       538.22       14.8
   a8++      572.56       33.4
   a3+       581.89       33.8
   b8++      586.24       18.8
   y5+       625.46       17.3
   b10++     665.76       12.2
   a4+       695.45       10.2
   y6+       712.47       22.5
   a11++     732.52       15.1
   b11++     747.03       13.5
   y7+       809.46       30.6
   b12++     811.42       24.5
   b5+       890.44       25.1
   y8+       906.40       96.7
   y9+      1073.43       20.0
   y10+     1186.64        9.0
```

```
439. MORG-339_phosphopeptide_TMT_4%ACN_032017_OT.6047.6047.3 (aSRVPSsDEEVVEEPQSR)

   Ion         m/z   Intensity
   a7++      484.19        5.2
   y4+       487.32        9.0
   b8++      555.57       10.3
   y5+       616.35        7.4
   b9++      619.52        5.6
   a10++     670.51        6.8
   b10++     684.67       10.6
   a5+       712.68        9.7
   a11++     719.49       11.2
   b11++     733.75       13.3
   b5+       740.35        8.6
   y12++     742.35       10.1
   y6+       745.58       19.8
   y13++     785.82        6.8
   a6+       799.34        9.7
   b6+       827.45        5.6
   y7+       844.45        7.3
   b13++     848.26        8.9
   b14++     912.26        9.3
   y8+       943.49        8.1
   a15++     947.05        7.0
   y9+      1072.53        8.5
   y10+     1201.57        8.1
   y11+     1316.55        6.1
```

```
440. MORG-339_phosphopeptide_TMT_22%ACN_032017_OT.6649.6649.4 (eEVAsEPEEAASPTTPkkk)

   Ion         m/z   Intensity
   y7+++     496.46       26.3
   y8+++     525.42       33.9
   y9+++     549.17       26.8
   y10+++    573.00       33.5
   b3+       587.40       22.2
   y4++      594.59       23.7
   y11+++    615.65       28.3
   b9++      655.20       21.2
   y12+++    658.40       23.2
   y13+++    691.21       73.5
   y6++      695.43       25.8
   a11++     712.53       27.1
   b11++     726.28       30.5
   y14+++    733.77       57.2
   y7++      744.21      114.4
   b12++     769.91       19.1
   y8++      787.67       76.1
   a13++     804.74       19.3
   y16+++    813.69       25.1
   b13++     818.25       23.6
   y9++      823.21       74.9
   b5+       825.41       19.0
   y17+++    846.64       23.4
   y10++     858.56       59.3
   b18+++    883.30       18.9
   y11++     923.39       31.4
   a16++     953.58       19.7
   b6+       954.48       28.9
   y12++     987.47       22.4
   y13++    1036.37       27.0
   b8+      1180.56       20.4
```

```
441. MORG-339_phosphopeptide_TMT_8%ACN_032017_OT.8319.8319.3 (sSGLQtPEcLsR)

   Ion         m/z   Intensity
   b1+       317.39       10.7
   y2+       342.20       10.4
   b5++      351.87        9.4
   y5++      372.36       15.9
   y6++      421.26       28.4
   a3+       433.41        5.5
   b6++      441.96       14.2
   y3+       455.14        7.4
   b3+       461.31       14.7
   b7++      490.76        8.6
   y7++      511.75       20.9
   a8++      541.39        9.7
   a4+       546.39       15.3
   b8++      555.56       18.1
   y4+       615.33       10.6
   b9++      635.41       10.6
   b10++     691.84       12.2
   b5+       702.44       20.9
   y11++     704.26        5.4
   y5+       744.43        9.6
   y6+       841.30       18.9
   b6+       883.47       12.2
   y7+      1022.31       13.4
```

```
442. MORG-339_phosphopeptide_TMT_10%ACN_032017_OT.9460.9460.4 (qADVADQQTTELPAENGETENQSPASEEEkEAksD)

   Ion         m/z   Intensity
   b3+       544.36       18.4
   b9++      593.97       21.8
   b4+       643.52       18.5
   b10++     644.39       14.4
   a18+++    699.88       34.4
   b11++     708.98       17.5
   b5+       714.44       21.9
   b12++     765.26       16.7
   b6+       829.48       22.1
   a7+       929.64       28.9
   y21+++    949.84       17.8
   b7+       957.60       21.8
   y22+++    973.24       45.5
   y23+++   1005.64       54.3
   y24+++   1043.37       39.7
   b28+++   1067.02       15.5
   b8+      1085.52       20.8
   y25+++   1086.42       19.5
   y15++    1094.19       32.5
   a19++    1100.54       14.1
   a29+++   1101.25       20.7
   y26+++   1119.40       25.6
   a9+      1158.85       14.1
   b9+      1186.59       19.6
   y17++    1209.28       32.6
   y18++    1273.91       20.2
   b10+     1287.59       16.4
   y31+++   1300.51       29.3
   y19++    1302.27       36.2
   y7+      1344.76       18.9
   y20++    1359.41       30.9
   b33+++   1414.86       13.0
   b11+     1416.74       32.6
   y22++    1458.99       35.5
   y23++    1507.96       89.6
   b12+     1529.82       44.5
   y24++    1564.14       14.2
   y25++    1629.14       16.4
```

```
443. MORG-339_phosphopeptide_TMT_18%ACN_032017_OT.8039.8039.4 (sLRRQQQPcMEPPESQLEPk)

   Ion         m/z   Intensity
   y2++      237.33       26.5
   y1+       376.32       30.2
   y6++      466.24       16.2
   y2+       473.39      151.6
   b5++      475.40       18.0
   b6++      539.40       17.4
   b11+++    575.49       17.5
   a12+++    598.10       28.1
   y3+       602.39       86.2
   b7++      603.50       46.9
   b12+++    607.87       24.5
   y9++      627.49       30.3
   b14+++    683.30       37.9
   y10++     691.65       22.2
   a15+++    702.91       16.2
   b15+++    711.82       39.0
   y4+       715.58       61.0
   y16+++    718.82       20.3
   b9++      732.74       24.9
   b16+++    755.00       59.9
   a17+++    783.12       47.6
   a10++     784.03       17.4
   b17+++    792.70      170.5
   a4+       794.42       16.7
   b10++     797.78       40.1
   a18+++    826.29       45.3
   b18+++    835.77      348.6
   y12++     837.26       33.9
   y5+       843.14       19.8
   y19+++    860.39       16.4
   b11++     862.65       75.6
   b19+++    868.00       28.3
   a12++     896.91       15.7
   b12++     911.13       18.7
   b14++    1024.19       50.8
   y7+      1059.84       17.9
   b15++    1067.78       20.8
   b16++    1131.76       56.9
   b17++    1188.33       56.9
   b18++    1252.82       72.9
   y9+      1253.69       38.6
```

```
444. MORG-339_phosphopeptide_TMT_10%ACN_032017_OT.7122.7122.4 (eTNVskEDtDQEEkASNEDVTk)

   Ion         m/z   Intensity
   y2+       477.37       14.7
   y3+       576.41       21.3
   b8++      721.77       20.9
   y5+       820.72       27.8
   b5+       840.67       46.2
   a15+++    842.04       13.2
   y11++     854.71       20.9
   a10++     855.59       21.5
   b10++     869.81      189.4
   a16+++    870.75       49.5
   b16+++    880.55       27.2
   y17+++    911.97       22.5
   y12++     918.73       16.8
   a11++     919.87       17.7
   y6+       934.61       35.4
   b18+++    961.57       25.7
   y18+++    967.66       32.7
   a19+++    990.83       14.0
   y19+++   1000.24       22.7
   b20+++   1033.12       19.5
   y14++    1066.84       24.3
   y21+++   1072.04       12.5
```

```
445. MORG-339_phosphopeptide_TMT_16%ACN_032017_OT.5769.5769.4 (vkEERASsPPVEVGEPR)

   Ion         m/z   Intensity
   b3+++     272.16       33.8
   b3++      408.48       15.5
   b7+++     420.34       13.4
   y8++      441.81       12.8
   y4+       458.29       14.8
   b8+++     475.81       21.7
   y9++      490.45       62.5
   b9+++     508.37       11.8
   a10+++    531.23       10.9
   a5++      536.44       21.3
   b5++      550.97       15.7
   y5+       557.38       18.7
   a11+++    564.70       19.3
   a6++      571.89       25.6
   y15+++    573.11       56.4
   y10++     574.25       24.1
   b6++      586.29       24.1
   y11++     616.93       33.2
   b7++      629.85       14.6
   a13+++    640.45       14.6
   b13+++    650.18       23.5
   b14+++    668.67       22.2
   b2+       686.30       21.9
   a15+++    702.63       26.6
   b15+++    711.89       68.3
   b8++      713.59       22.0
   y13++     731.09       10.7
   a16+++    734.86       10.7
   y7+       785.39       11.7
   a3+       787.38       11.7
   y14++     795.57       11.1
   b11++     860.01       13.3
   y8+       882.47       12.7
   b12++     924.62       16.5
   y9+       979.49       43.5
   b14++    1002.70       11.6
```

```
446. MORG-339_phosphopeptide_TMT_18%ACN_032017_OT.7625.7625.3 (eVSSkEEQsPVkAEVAEk)

   Ion         m/z   Intensity
   y1+       376.42       12.2
   y2+       505.41       12.0
   y7++      616.85       13.1
   y9++      715.11       45.9
   y5+       804.64       12.7
   y11++     862.73       32.8
   y6+       875.54       20.5
   y12++     927.40       13.7
   a5+       961.88       27.9
   b12++    1048.80       20.7
   b13++    1083.87       11.4
   b6+      1118.60       11.5
   b14++    1148.55       11.7
   b15++    1198.42       12.0
   b7+      1247.68       15.8
   a17++    1284.42       12.8
   b17++    1298.37       12.9
   b8+      1375.84       11.4
   y9+      1428.95       17.9
```

```
447. MORG-339_phosphopeptide_TMT_10%ACN_032017_OT.16459.16459.2 (sESsDFEVVPk)

   Ion         m/z   Intensity
   y1+       376.41       15.5
   y2+       473.33       61.6
   y3+       572.46       33.3
   b3+       613.32       11.5
   y4+       671.62       25.7
   y5+       800.59       21.5
   a5+       867.74        9.2
   b5+       895.29       26.6
   y6+       947.64       33.4
   a6+      1014.50       10.1
   b6+      1042.42       19.3
   y7+      1062.64       15.3
   b7+      1171.39       25.3
   b8+      1270.47       38.4
   a9+      1341.54       15.7
   b9+      1369.52       63.5
```

```
448. MORG-339_phosphopeptide_TMT_14%ACN_032017_OT.4952.4952.3 (rADLNQGIGEPQsPSR)

   Ion         m/z   Intensity
   b3++      286.94       16.6
   y3+       359.25       26.2
   y6++      376.19       12.6
   b5++      400.33       10.5
   b6++      464.47       15.7
   b7++      492.94       27.0
   a8++      535.36       10.8
   b8++      549.67       34.6
   b3+       572.63       12.1
   b9++      577.77       17.0
   a10++     628.03       15.8
   b10++     642.34       47.7
   y5+       654.37       48.3
   a4+       657.75       26.2
   b11++     691.05       12.0
   y13++     731.58       11.0
   y6+       751.41       30.7
   b12++     754.79       18.8
   y14++     789.70       67.9
   b5+       799.36       11.8
   a14++     872.62       15.7
   y8+       937.43       19.6
   b7+       984.68       11.0
   b8+      1097.76       11.1
   b10+     1283.75       10.8
```

```
449. MORG-339_phosphopeptide_TMT_20%ACN_032017_OT.14196.14196.3 (nGsIVsmNMkDAFAR)

   Ion         m/z   Intensity
   b2+       401.25        9.8
   y4+       464.29        9.9
   y7++      534.47        8.7
   y5+       579.43        8.0
   y8++      591.37        7.4
   b8++      604.93        7.8
   b4+       681.19        7.0
   y10++     748.59       21.3
   b5+       780.61        9.8
   y12++     854.74       13.7
   b11++     906.18       11.2
   y14++     966.68       10.4
   a13++    1001.94        6.9
   b14++    1051.08        6.7
   y8+      1181.57        6.9
   b8+      1208.48       12.0
```

```
450. MORG-339_phosphopeptide_TMT_4%ACN_032017_OT.1384.1384.3 (aLHGAQTsDEER)

   Ion         m/z   Intensity
   y1+       175.14       11.3
   a3++      262.11       10.5
   b3++      276.30       18.7
   b4++      304.46       37.9
   a5++      326.35       15.7
   b5++      340.34       30.9
   a6++      390.37       16.8
   b6++      404.18       36.2
   b2+       414.32       23.3
   y3+       433.28       22.7
   b7++      454.75       15.7
   y7++      472.85       21.4
   a3+       523.37       21.4
   a8++      524.41       12.5
   y4+       548.40       11.3
   b3+       551.38       33.3
   b9++      595.71       25.2
   y10++     605.54        8.4
   b4+       608.48       10.0
   a10++     646.32       11.8
   a5+       651.49       13.1
   b10++     660.41       37.2
   y11++     661.36       11.7
   b5+       679.46       17.2
   y5+       715.30       15.0
   b11++     725.28       23.9
   b6+       807.59        9.1
   y6+       816.33       26.6
   y8+      1015.43        8.3
   y9+      1072.52       12.2
```

```
451. MORG-339_phosphopeptide_TMT_16%ACN_032017_OT.7497.7497.3 (rAQsTDsLGTSSSLQSk)

   Ion         m/z   Intensity
   b4++      376.26       11.9
   y2+       463.37       25.6
   b6++      484.92       24.4
   y9++      561.97       11.6
   y3+       591.36       28.2
   b9++      653.44       17.6
   a10++     689.73       14.5
   y11++     702.24       14.2
   b10++     703.73       12.7
   y4+       704.56       20.5
   a4+       724.02       21.6
   a11++     733.71       13.8
   b11++     747.81       17.8
   b4+       752.71       71.0
   a12++     777.33       19.3
   a5+       825.35       17.2
   y6+       878.55       21.8
   b14++     891.07       17.1
   y14++     894.37       12.0
   a15++     941.12       15.7
   b15++     954.90       24.3
   y7+       965.52       19.8
   b6+       968.48       18.3
   a16++     984.70       12.0
   y16++     993.48       11.4
   y8+      1066.55       12.7
   y9+      1123.65       20.6
```

```
452. MORG-339_phosphopeptide_TMT_20%ACN_032017_OT.8721.8721.4 (rNEAFVckPDPSPPsPSEEEk)

   Ion         m/z   Intensity
   y6++      474.64       21.2
   y2+       505.32       22.9
   a7++      540.12       14.2
   b7++      553.78       20.1
   b10+++    559.58      135.7
   y8++      606.04       29.2
   b12+++    620.82       27.6
   y3+       634.43       19.1
   a13+++    643.24       15.0
   y9++      655.00       52.5
   y10++     698.16       35.7
   a15+++    731.69       16.2
   b8++      732.68       32.2
   y11++     747.19       67.7
   y4+       763.42       28.8
   a16+++    764.18       53.5
   a9++      766.47       36.1
   b16+++    773.48       63.8
   y16+++    774.42       33.5
   b9++      781.14       43.8
   y12++     803.97       18.4
   y17+++    823.53       17.3
   a18+++    835.99       15.8
   b10++     838.70       99.2
   b18+++    845.73       18.2
   y13++     853.01       21.5
   b11++     887.00       22.2
   b19+++    888.80       22.3
   a20+++    921.76       29.0
   b12++     930.84       51.1
   b20+++    931.64       34.5
   b6+       946.82       14.7
   y6+       947.49       39.6
   b13++     979.15       15.2
   y10+     1395.72       30.7
   b20++    1396.77       20.8
   y11+     1492.78       37.8
```

```
453. MORG-339_phosphopeptide_TMT_10%ACN_032017_OT.2536.2536.3 (sStPPRQsPsRSSSPQPk)

   Ion         m/z   Intensity
   b2+       404.23        9.7
   y2+       473.40       18.7
   b3+       585.28       12.3
   y9++      641.43       11.1
   y10++     690.81       16.2
   y4+       698.48       31.3
   a5+       751.38        9.8
   y11++     773.88        9.7
   b5+       779.00       26.9
   y5+       785.92       18.7
   a11++     811.48       48.2
   b11++     825.86       33.8
   y12++     838.24      113.6
   a12++     855.29       30.0
   a13++     899.00       20.9
   a6+       907.61       28.8
   y13++     915.69       15.5
   a14++     942.59       11.4
   b14++     956.28       16.7
   y14++     964.24       41.3
   b15++    1005.09       14.1
   y15++    1013.23       65.2
   a16++    1054.60       14.3
   b16++    1068.88       15.0
```

```
454. MORG-339_phosphopeptide_TMT_18%ACN_032017_OT.1848.1848.4 (sSRkGTENGVNGTVTSNGADsPR)

   Ion         m/z   Intensity
   y2+       272.20       27.2
   y3+       439.32       13.3
   y9++      492.62       25.3
   b11+++    530.57       21.4
   y16+++    542.47       13.6
   b12+++    549.45       15.4
   y4+       554.28       17.7
   a13+++    573.85       20.4
   b13+++    583.24       58.0
   a14+++    606.87       25.4
   b14+++    616.24       82.1
   y5+       625.29       16.3
   y19+++    637.78       14.9
   a15+++    640.76       13.3
   a8++      646.20       12.9
   b15+++    649.81       65.8
   a16+++    669.33       34.7
   a9++      674.80       38.9
   b16+++    678.46       53.9
   y6+       682.18       46.6
   b9++      688.63       50.9
   b17+++    717.04       26.5
   a10++     723.80       16.0
   a18+++    726.81       15.0
   b18+++    735.94       33.3
   b10++     738.02       26.0
   y15++     756.53       12.8
   b19+++    759.43       31.0
   b11++     795.34       22.6
   y7+       796.31       37.0
   b20+++    797.72       44.9
   b12++     823.16       20.5
   a21+++    843.70       14.3
   b21+++    853.65       23.0
   b13++     874.13       48.9
   y8+       883.33       40.9
   b14++     923.46       41.9
   b5+       974.34       29.1
   y9+       984.43       48.8
   b16++    1017.64       16.1
   y10+     1083.51       20.0
```

```
455. MORG-339_phosphopeptide_TMT_12%ACN_032017_OT.6652.6652.3 (eREIQSITDESRGsIR)

   Ion         m/z   Intensity
   y2+       288.29       14.9
   b2+       515.16        9.1
   b7++      543.34       12.4
   a3+       616.57       10.7
   b3+       644.34       14.7
   b9++      651.60       36.8
   y5+       668.44       16.3
   a10++     702.14       18.6
   y12++     714.65       33.1
   b10++     715.67       25.7
   y6+       755.39       11.3
   b11++     759.68        9.4
   b12++     837.79       13.2
   b13++     866.11       16.2
   y7+       884.39       16.6
   b5+       885.41       16.3
   b14++     949.33       11.2
   b15++    1006.31       10.6
   b9+      1301.74       12.3
```

```
456. MORG-339_phosphopeptide_TMT_14%ACN_032017_OT.2523.2523.3 (eAsPPTSADkSR)

   Ion         m/z   Intensity
   y3++      310.29       14.3
   b1+       359.16        8.7
   y6++      446.73       10.8
   y7++      497.49       15.7
   y8++      545.97       32.3
   a9++      568.92       44.3
   b9++      583.27       45.9
   y3+       619.60       20.1
   a4+       666.34       10.4
   y10++     677.48       10.6
   y4+       734.11       11.6
   b10++     762.33        9.5
   y5+       805.67       11.6
   y8+      1090.56        9.2
```

```
457. MORG-339_phosphopeptide_TMT_8%ACN_032017_OT.3699.3699.3 (rNssSPVsPASVPGQRR)

   Ion         m/z   Intensity
   y5++      307.26       20.9
   y6++      356.86       13.1
   y7++      400.30       10.3
   y9++      484.46       37.9
   b6++      509.92       10.2
   b7++      558.82       10.7
   y10++     567.90       16.2
   y5+       613.38       17.9
   y11++     616.89       22.6
   y12++     666.02       32.8
   b3+       667.22       10.2
   b9++      691.12       11.7
   y13++     709.38       12.1
   a10++     712.56       87.3
   b10++     727.23       33.1
   y14++     792.99        9.1
   b12++     820.02        7.9
   b5+       921.42       11.4
   a6+       990.59        8.2
   b16++    1038.84       12.4
   b7+      1117.59       12.7
   b10+     1452.65        9.5
```

```
458. MORG-339_phosphopeptide_TMT_14%ACN_032017_OT.8549.8549.3 (tVTmISPEDEQk)

   Ion         m/z   Intensity
   y3++      317.31       11.6
   y4++      375.09        8.0
   y1+       376.37       21.8
   b6++      479.34       12.6
   y6++      487.77       37.2
   y2+       504.39       23.6
   b2+       510.11        8.4
   b7++      528.49       10.1
   a8++      578.39       18.4
   a3+       583.58        8.8
   b8++      592.31       11.2
   b3+       611.03       13.6
   y3+       633.03       24.6
   a9++      635.98       30.3
   a10++     701.02        8.7
   b10++     714.90       26.5
   a4+       730.18       19.7
   y4+       748.39       18.8
   a11++     764.73        8.8
   b11++     778.78       14.8
   b6+       958.51       12.7
   y6+       974.75        8.3
   b8+      1184.70        8.7
   b10+     1428.71        8.7
```

```
459. MORG-339_phosphopeptide_TMT_20%ACN_032017_OT.9418.9418.3 (sETLVNAQQTPLGtPkENR)

   Ion         m/z   Intensity
   a1+       289.26       19.0
   b6++      436.92       51.4
   b3+       547.40       15.8
   y9++      660.72       33.6
   b5+       759.51       26.0
   y4+       775.44       14.9
   a6+       845.87       18.1
   a7+       916.45       20.5
   b15++     923.98       26.3
   b7+       944.84       19.5
   y15++     981.11       23.8
   b8+      1072.51       17.8
   a16++    1088.84       22.7
   b17++    1167.59       31.4
   b9+      1201.12       16.4
   b10+     1301.65       20.2
```

```
460. MORG-339_phosphopeptide_TMT_18%ACN_032017_OT.8678.8678.3 (nLAkPGVTSTsDSEDEDDQEGEk)

   Ion         m/z   Intensity
   y1+       376.31       14.5
   b4++      443.44       31.7
   b3+       528.29       18.2
   y3+       562.53       25.6
   b7++      569.89       22.2
   b8++      620.61       23.9
   y4+       691.38       15.2
   b10++     714.62       25.0
   b11++     797.77       24.8
   y5+       819.47       38.0
   a12++     841.66       14.2
   b12++     855.67       37.9
   b4+       885.64       43.7
   y6+       934.56       40.0
   a5+       954.72       14.3
   b14++     963.62       31.3
   b5+       982.45       14.5
   a6+      1011.24       18.3
   b15++    1020.58       48.5
   b6+      1039.34       17.4
   y16++    1041.27      239.7
   y7+      1049.54       33.5
   b16++    1085.56       21.0
   y18++    1118.89       28.6
   b7+      1138.68       14.9
   b17++    1143.18       45.9
   y19++    1167.72       28.9
   y8+      1178.51       37.5
   b18++    1200.77       45.5
   y9+      1293.57       20.3
   b20++    1329.41       34.3
   y10+     1422.54       26.4
   y22++    1438.41       21.4
   y11+     1509.58       34.4
   y12+     1624.72       18.1
```

```
461. MORG-339_phosphopeptide_TMT_14%ACN_032017_OT.16907.16907.4 (gADDAADADtAIINAEGGQNNSEEkkEYFI)

   Ion         m/z   Intensity
   b8++      458.73       13.2
   b4+       588.35       13.9
   b11++     642.27       30.0
   y6++      643.47       18.0
   b5+       659.30       16.7
   a12++     684.56       10.9
   b12++     698.96       41.5
   y7++      708.19       12.8
   b6+       730.37       19.3
   a21+++    755.54       24.3
   b21+++    764.86       12.2
   y9++      816.28       16.6
   b23+++    837.73       18.0
   y18+++    843.60       34.9
   b7+       845.32       21.8
   b15++     847.89       38.9
   y10++     872.75       12.2
   y19+++    881.50       12.1
   y20+++    905.10       16.1
   b16++     912.46       32.4
   b8+       916.54       18.7
   y5+       928.52       16.7
   y11++     929.95       16.0
   b17++     941.06       15.6
   y21+++    965.48       12.8
   y13++    1022.67       27.7
   y14++    1051.31       53.5
   y26+++   1112.73       13.6
   y15++    1115.76       45.1
   b26+++   1118.96       19.0
   y27+++   1151.49       33.4
   b27+++   1162.00       24.8
   y17++    1208.01       34.6
   b28+++   1216.21       31.3
   a29+++   1255.86       26.2
   b29+++   1264.83       77.7
   b11+     1283.66       14.0
   y19++    1321.40       20.9
   b12+     1396.64       32.5
   b13+     1509.71       17.7
```

```
462. MORG-339_phosphopeptide_TMT_20%ACN_032017_OT.15368.15368.2 (iDIsPSTFR)

   Ion         m/z   Intensity
   y2+       322.24        8.9
   b1+       343.46        6.2
   y3+       423.37        6.6
   b2+       458.29       24.4
   b3+       571.27       17.1
   y5+       607.35       46.6
   b4+       738.50        8.0
   y6+       774.35       10.0
   b5+       835.39        9.6
   y7+       887.45       16.5
   b6+       922.52        6.5
   y8+      1002.53       10.1
```

```
463. MORG-339_phosphopeptide_TMT_12%ACN_032017_OT.2283.2283.4 (qRRkPscsEGEEAEQk)

   Ion         m/z   Intensity
   y4++      352.77       10.7
   y1+       376.35       21.7
   a2+       486.36       13.8
   a4++      500.49       12.9
   y2+       504.42       30.0
   b4++      514.62       23.4
   b9+++     583.65       22.0
   y3+       633.44       33.1
   a11+++    636.16       10.5
   b11+++    645.48       27.3
   b6++      646.54       21.3
   y9++      658.30       30.6
   b3+       670.05       22.5
   y4+       704.44       19.7
   a7++      712.12       27.1
   a14+++    746.02       12.6
   b14+++    755.21       45.1
   b15+++    797.76       32.1
   y15+++    804.08       12.1
   y11++     822.04       17.2
   y5+       833.49       12.2
   y12++     870.54       21.1
   y7+      1019.50       13.4
   b13++    1067.42       12.6
   y8+      1148.56       12.8
```

```
464. MORG-339_phosphopeptide_TMT_25%ACN_032017_OT.17326.17326.2 (lASNsPVLPQAFAR)

   Ion         m/z   Intensity
   b1+       343.30       16.5
   b2+       414.40       15.1
   b4+       615.54       18.3
   y6+       689.46       61.7
   b5+       782.38       18.2
   y7+       802.57       24.7
   a6+       851.14       68.2
   b6+       879.53       24.1
   b7+       978.70       24.2
   y9+       998.64       72.2
   b8+      1091.65       44.6
   y12+     1366.68       17.9
   y13+     1437.70       23.6
```

```
465. MORG-339_phosphopeptide_TMT_22%ACN_032017_OT.7739.7739.3 (rVMTVTAVTTTATsDR)

   Ion         m/z   Intensity
   b7++      495.01       19.1
   b8++      544.49       34.3
   a9++      581.11       15.1
   a3+       587.99       14.8
   b9++      595.10       31.1
   b3+       615.93       17.4
   y5+       629.30       35.3
   a10++     631.32       31.4
   b10++     645.60       27.9
   y12++     652.26       39.3
   a4+       689.35       16.2
   b11++     696.04       24.3
   b4+       717.45       23.0
   b12++     731.49       20.0
   b13++     782.11       22.4
   a5+       788.11       15.1
   b5+       816.67       33.7
   y7+       831.48       24.3
   b14++     865.16       18.8
   b6+       917.68       14.3
   b15++     922.99       16.2
   y8+       932.44       23.3
```

```
466. MORG-339_phosphopeptide_TMT_22%ACN_032017_OT.10025.10025.3 (rSsLPNGEGLQLk)

   Ion         m/z   Intensity
   y1+       376.37       21.3
   b6++      482.70       14.4
   y2+       489.50       25.7
   b7++      511.39       19.9
   b8++      575.98       33.9
   a9++      590.22       17.9
   y9++      592.99       38.0
   b9++      604.00       26.9
   a3+       612.02       15.9
   y3+       617.52       35.9
   b10++     661.06       25.2
   a4+       724.94       33.7
   y4+       730.74       21.0
   y11++     732.55       17.4
   b4+       753.44       18.9
   y12++     776.43       17.7
   b12++     781.45       22.7
   y5+       787.63       32.2
   y6+       916.62       15.6
   y7+       973.70       14.7
   y9+      1184.90       17.4
```

```
467. MORG-339_phosphopeptide_TMT_10%ACN_032017_OT.13069.13069.3 (sETLVNAQQtPLGtPk)

   Ion         m/z   Intensity
   a4++      316.32        8.2
   b1+       317.34       17.0
   y5++      412.46        9.9
   y6++      461.50       16.9
   y2+       473.35       19.4
   a8++      522.95        8.0
   y7++      552.15       12.1
   b9++      600.80       12.5
   y3+       654.48       12.6
   b4+       660.22        9.0
   b10++     691.09       12.7
   y4+       711.38       11.6
   b11++     739.95       13.7
   b5+       759.26       15.7
   a12++     782.79       14.7
   b12++     796.80       20.8
   y12++     822.12       11.6
   y5+       824.13        8.1
   b6+       873.50       11.7
   b14++     915.41        8.0
   y6+       921.62        7.5
   b7+       944.59       13.7
   b9+      1200.53        7.7
```

```
468. MORG-339_phosphopeptide_TMT_6%ACN_032017_OT.5042.5042.2 (sIQEDTk)

   Ion         m/z   Intensity
   y1+       376.39        8.1
   b1+       397.22        7.0
   y2+       477.32       24.8
   y3+       592.50       11.0
   b3+       638.43        9.2
   y4+       721.42        9.4
   b4+       767.43       10.0
   y5+       849.52       12.3
   b5+       882.41       20.0
   y6+       962.53       14.3
```

```
469. MORG-339_phosphopeptide_TMT_22%ACN_032017_OT.10309.10309.3 (kRDsESEsDETPTVPQLk)

   Ion         m/z   Intensity
   y4+       714.52       42.7
   y5+       813.68       14.3
   b11++     827.05       24.4
   a12++     861.61       12.4
   a13++     912.51       17.7
   y6+       914.38       16.2
   b5+       925.77       24.1
   b13++     926.70       59.7
   y14++     935.20       24.9
   a15++    1010.75       16.6
   y7+      1011.70       34.5
   b6+      1012.60       28.0
   b15++    1024.54       13.8
   y16++    1076.25       13.2
   y8+      1112.72       16.9
   a7+      1113.60       14.0
   y9+      1241.76       20.1
```

```
470. MORG-339_phosphopeptide_TMT_25%ACN_032017_OT.6440.6440.4 (rkNLAkPGVtSTSDSEDEDDQEGEk)

   Ion         m/z   Intensity
   y3+       562.42       32.2
   y4+       691.52       26.7
   b6++      700.27       23.6
   a2+       715.25       31.6
   b14+++    741.80       29.7
   b16+++    813.67       27.4
   y5+       819.49       45.7
   b17+++    852.39       41.4
   b3+       857.21       25.7
   a18+++    886.17       24.1
   b18+++    895.19       45.8
   y20+++    898.20       34.6
   a10++     902.94       37.8
   y14++     906.79       40.5
   y21+++    921.76       33.6
   a19+++    923.74       39.5
   b4+       970.36       20.3
   b20+++    972.15       42.4
   a21+++   1004.99       23.3
   b12++    1010.70       32.5
   b21+++   1014.48       26.8
   y16++    1040.51       21.0
   y7+      1049.68       20.6
   b22+++   1057.67       34.4
   b23+++   1076.43       30.0
   a24+++   1110.28       19.9
   b24+++   1119.51       30.8
   y8+      1178.64       26.1
   y11+     1509.85       21.0
```

```
471. MORG-339_phosphopeptide_TMT_25%ACN_032017_OT.10102.10102.3 (sPTGAQPAAAkPPPLSAk)

   Ion         m/z   Intensity
   b1+       397.21       19.9
   y7++      470.06       19.5
   y8++      648.47       19.1
   y10++     719.95       29.1
   y5+       744.67       23.9
   a11++     745.76       19.0
   b11++     759.87       48.6
   a12++     794.35       45.2
   y12++     803.56       61.8
   b12++     808.42       53.7
   y6+       841.58       22.2
   a13++     842.50       24.7
   b6+       851.39       23.5
   y13++     867.72       28.4
   y14++     902.68       19.4
   b14++     905.41       24.2
   a7+       920.08       27.6
   y15++     931.62       22.8
   y7+       938.64       54.5
   a15++     947.75       21.1
   b7+       948.71       20.8
   a16++     991.69       19.5
   y17++    1030.92       60.0
```

```
472. MORG-339_phosphopeptide_TMT_10%ACN_032017_OT.10028.10028.4 (eNPPVEDssDEDDkRNPGNLYDk)

   Ion         m/z   Intensity
   y1+       376.27       15.5
   y10+++    555.13       33.3
   y11+++    593.54       23.2
   y12+++    631.70       14.8
   y9++      653.57       13.9
   y3+       654.47       17.2
   b9++      672.48       12.3
   y13+++    674.96       31.8
   y14+++    713.32       13.6
   y4+       767.58       14.2
   y15+++    768.69       18.6
   b15+++    778.08       14.9
   a16+++    806.96       12.4
   y16+++    824.42       31.9
   y10++     831.90       23.3
   a12++     838.51       22.9
   a17+++    839.38       16.4
   b17+++    848.16       35.0
   b12++     852.56       30.5
   a18+++    857.70       20.0
   y17+++    862.31       29.1
   a13++     895.39       14.9
   y18+++    905.69       13.3
   a20+++    933.60       12.5
   y6+       938.21       25.2
   y19+++    939.07       16.3
   y21+++   1003.60       27.8
   b7+      1010.54       24.4
   b22+++   1035.81       14.7
   y22+++   1041.40       18.0
   b14++    1088.17       15.5
```

```
473. MORG-339_phosphopeptide_TMT_14%ACN_032017_OT.3472.3472.4 (gLAEDVTRTtPSRR)

   Ion         m/z   Intensity
   y1+       175.06        9.5
   y4++      258.33       65.7
   b1+       287.19        7.3
   b4++      300.57        7.7
   b5++      358.42       13.5
   a2+       372.38       10.2
   y9+++     385.33       37.8
   b9+++     391.13       19.3
   y6++      399.26       12.1
   b2+       400.30       30.2
   b6++      407.49        8.9
   y10+++    423.73       32.2
   a10+++    442.71       15.0
   a3+       443.54      133.1
   b10+++    452.19        8.3
   b7++      458.13       19.3
   b3+       471.35       14.2
   y12+++    490.53       12.0
   y4+       515.36        7.8
   a8++      521.95        9.0
   y13+++    528.06       47.1
   b8++      536.60       18.6
   a13+++    555.56        8.8
   b13+++    565.66       11.6
   a9++      572.45       11.1
   y9++      577.60       33.2
   b9++      587.04       15.0
   b4+       600.34       31.1
   y10++     635.07       40.2
   a10++     663.29        7.3
   b10++     677.30       19.9
   y11++     699.68       19.5
   b5+       715.39       37.9
   y12++     734.98        9.5
```

```
474. MORG-339_phosphopeptide_TMT_10%ACN_032017_OT.12437.12437.2 (sVSELsLQGR)

   Ion         m/z   Intensity
   b1+       397.24        8.8
   y4+       473.35       10.5
   y5+       640.31       12.2
   a4+       684.01       97.3
   b4+       712.75       25.7
   y6+       753.35        7.9
   a5+       797.71        9.2
   b5+       825.37       13.9
   b8+      1233.52        6.9
```

```
475. MORG-339_phosphopeptide_TMT_14%ACN_032017_OT.2959.2959.3 (rNsSSPVsPASVPGQRR)

   Ion         m/z   Intensity
   y5++      307.36       13.2
   y9++      484.45       24.1
   b7++      519.51       11.7
   y5+       613.53       15.1
   y11++     617.00       17.2
   a3+       639.15        8.5
   a10++     673.08       11.6
   y13++     709.50       36.2
   y6+       712.38       33.9
   a11++     715.99       13.4
   b4+       754.47        9.5
   a5+       813.65       12.5
   a13++     814.76        9.2
   a14++     842.71       13.2
   b15++     921.20        9.5
   b16++     999.24       13.4
   b7+      1037.66        9.5
```

```
476. MORG-339_phosphopeptide_TMT_14%ACN_032017_OT.9180.9180.3 (sVPHFsEEEkEAE)

   Ion         m/z   Intensity
   b1+       317.16       61.5
   b4++      325.84       12.4
   b5++      399.21       12.6
   b2+       416.31       26.8
   y5++      417.35       10.8
   a8++      598.21       11.3
   b8++      611.66       17.9
   b4+       650.42       28.0
   a9++      662.11       45.3
   b9++      676.64       38.8
   y9++      703.84       10.1
   y4+       705.47       16.1
   y11++     821.01       15.4
   y5+       834.38       16.4
   b10++     855.21       11.5
   a6+       936.53       10.1
   b12++     955.46        9.7
   y6+       963.49       12.0
   y7+      1092.62        9.8
   y8+      1259.63       12.0
```

```
477. MORG-339_phosphopeptide_TMT_20%ACN_032017_OT.9428.9428.3 (vRGWsPPPEVR)

   Ion         m/z   Intensity
   y1+       175.12        6.4
   a3++      257.32        8.1
   y2+       274.15       28.3
   y5++      299.31       11.3
   y6++      347.95       31.2
   b4++      364.79        5.6
   b5++      448.33       15.9
   a6++      482.73        7.9
   y4+       499.90       15.8
   a3+       514.46       13.1
   y8++      524.32       11.5
   b7++      545.07        6.2
   y5+       597.44       16.0
   a9++      644.53       15.3
   b9++      658.58       52.8
   y6+       694.43       52.4
   b10++     708.18       13.0
   b5+       895.44       12.9
```

```
478. MORG-339_phosphopeptide_TMT_10%ACN_032017_OT.13246.13246.3 (aVDSsSEEIEVEVPVVDRR)

   Ion         m/z   Intensity
   y2+       331.26       14.6
   y6++      371.28        9.2
   y7++      420.98       10.5
   b3+       515.37       14.5
   y9++      535.05       12.2
   y10++     599.58        9.4
   b9++      614.22        9.0
   y11++     656.09       17.2
   y12++     720.38       10.0
   b11++     728.37        8.4
   y6+       741.55       16.2
   b5+       769.44       17.3
   y13++     785.42       22.8
   b12++     792.49       20.7
   y14++     828.61       16.7
   y7+       840.62        9.0
   a15++     926.34       24.8
   y16++     955.14       46.3
   b7+       985.35        9.5
   b8+      1114.40       11.1
   b18++    1125.66        8.7
   b10+     1356.56        9.6
   b13+     1683.75        9.7
```

```
479. MORG-339_phosphopeptide_TMT_14%ACN_032017_OT.13241.13241.2 (sLmssPEDLTk)

   Ion         m/z   Intensity
   b1+       317.17       10.9
   y1+       376.30       10.1
   y2+       477.47        8.7
   b3+       577.36        9.9
   y3+       590.48       21.7
   y4+       705.51       15.6
   a4+       716.45       10.1
   b5+       911.60       14.1
   y6+       931.53       25.4
   y7+      1098.54       15.4
   b7+      1137.51       13.3
   b8+      1252.40       22.0
   b9+      1365.62       11.5
```

```
480. MORG-339_phosphopeptide_TMT_14%ACN_032017_OT.6772.6772.3 (aSGPLDTSQtASSDSEDVSSSDRk)

   Ion         m/z   Intensity
   y7++      504.45       25.7
   y2+       532.50       10.2
   y8++      561.99       10.8
   y10++     669.73       26.2
   y11++     727.54       19.2
   y12++     770.44       24.3
   y5+       821.44       10.5
   a8+       930.80       12.4
   y15++     940.37       21.9
   b8+       958.73       12.3
   a17++     966.20       15.5
   y17++    1048.23       11.3
   b9+      1086.54       11.3
   y18++    1098.84       22.5
   y8+      1122.33       12.4
   y10+     1338.74       11.1
   y13+     1627.70       11.4
   b16+     1843.82       11.8
```

```
481. MORG-339_phosphopeptide_TMT_10%ACN_032017_OT.9115.9115.3 (sHsLPNsLDYAQASER)

   Ion         m/z   Intensity
   b1+       317.22        9.6
   y3+       391.36       11.3
   a6++      458.94       12.2
   y4+       462.34       17.9
   b6++      473.14       16.8
   y5+       590.39       16.9
   b8++      613.70        9.8
   b3+       621.67       20.7
   a9++      656.69       36.1
   y6+       661.35       20.0
   y11++     667.52       20.7
   b9++      670.96       26.7
   a4+       706.04       13.3
   b4+       734.42       10.9
   a10++     738.79       16.4
   b10++     752.48       19.0
   y13++     772.44       18.4
   a11++     773.46        9.7
   b11++     787.83       17.3
   a5+       802.95        9.5
   y7+       824.55       24.6
   a12++     837.47       13.9
   b12++     852.09       18.8
   y14++     856.21        9.4
   a13++     873.17       10.4
   b13++     887.46       13.8
   y8+       939.56       24.8
   y9+      1052.55       11.3
```

```
482. MORG-339_phosphopeptide_TMT_16%ACN_032017_OT.10126.10126.3 (dAPFFREsPVGR)

   Ion         m/z   Intensity
   y1+       175.12        6.5
   y2+       232.25        9.6
   a1+       317.24       14.3
   b1+       345.27       15.6
   b2+       416.33       10.8
   y4+       428.30       44.7
   b6++      482.49       18.7
   a3+       485.23        8.2
   b3+       513.39       12.2
   y8++      514.27        8.0
   a7++      532.40       25.1
   b7++      546.64       25.1
   y9++      587.64        6.6
   y5+       595.42       11.6
   a8++      616.61        8.8
   b8++      630.51       24.3
   b4+       660.42       11.5
   a9++      665.32        8.1
   y11++     672.13       20.7
   a10++     714.70       17.1
   y6+       724.41        6.8
   b10++     728.71       29.0
   b5+       807.40        9.9
```

```
483. MORG-339_phosphopeptide_TMT_10%ACN_032017_OT.12125.12125.2 (sEsPPAELPSLR)

   Ion         m/z   Intensity
   b1+       317.25       14.2
   a2+       418.31       10.0
   b2+       446.27       12.3
   y4+       472.34       14.0
   y5+       585.37       21.9
   b3+       613.34       13.0
   a4+       682.53       10.7
   a5+       779.30       27.1
   y7+       785.75       17.5
   a6+       850.31       10.6
   y8+       882.61       12.0
   y9+       979.57       72.4
   b7+      1007.41       21.8
   b8+      1120.57       15.4
   y10+     1146.52       12.0
```

```
484. MORG-339_phosphopeptide_TMT_10%ACN_032017_OT.9784.9784.4 (qADVADQQTTELPAENGETENQsPASEEEkEAk)

   Ion         m/z   Intensity
   b8++      543.50       34.9
   b3+       544.37       41.9
   y3+       576.37       24.5
   b9++      594.04       56.4
   b4+       643.43       44.3
   b10++     644.41       33.0
   y7++      661.05       24.3
   a18+++    700.12       90.9
   y8++      704.63       27.1
   b11++     709.09       37.3
   b18+++    710.05       21.8
   b5+       714.46       53.8
   b12++     765.66       54.8
   y10++     788.73       49.2
   a6+       801.36       23.8
   b6+       829.46       55.5
   y11++     872.20       72.9
   y20+++    906.09       42.3
   y4+       933.68       32.9
   b7+       957.54       45.6
   b16++     970.50       32.0
   y13++     993.19       69.9
   a27+++   1041.40       21.7
   b27+++   1050.33       25.9
   y24+++   1052.48       27.8
   a8+      1057.76       58.4
   y5+      1062.36       67.7
   y15++    1107.91       39.2
   a20++    1164.74       28.8
   y27+++   1171.77       20.6
   y16++    1172.79       43.3
   b9+      1186.61       43.0
   y17++    1200.91       46.5
   y28+++   1210.30       22.4
   a10+     1259.74       55.0
   b10+     1287.78       29.9
   b22++    1300.53       38.3
   y20++    1358.49       53.0
   y8+      1407.46      199.2
   b11+     1416.67       61.6
   a24++    1417.66       55.1
   b12+     1529.78      101.7
   b15+     1827.00       20.5
```

```
485. MORG-339_phosphopeptide_TMT_16%ACN_032017_OT.18515.18515.3 (nLLEDDsDEEEDFFLRGPSGPR)

   Ion         m/z   Intensity
   y7++      363.81        9.5
   b2+       457.35       13.6
   y5+       513.19        8.7
   y10++     567.59       43.9
   b3+       570.44       23.0
   y11++     625.16       19.4
   y12++     689.15       12.2
   b4+       699.44       15.1
   y13++     753.85       19.3
   b5+       814.51       30.2
   y14++     818.64       35.0
   a12++     843.39       11.4
   y15++     876.14       31.9
   a6+       901.41       16.1
   a13++     917.30      242.1
   b6+       929.53       29.8
   b13++     930.52       15.5
   y16++     959.10       13.1
   y17++    1017.08       24.3
   a7+      1068.07        9.9
   y18++    1074.51       17.5
   b16++    1139.12       13.4
   y20++    1195.48       14.8
   b8+      1211.29       17.2
   b20++    1287.66       14.0
   b9+      1340.72        9.8
   b10+     1469.55        9.0
   b12+     1713.93        8.7
```

```
486. MORG-339_phosphopeptide_TMT_22%ACN_032017_OT.11212.11212.4 (hVGDLGNVTAGkDGVANVsIEDR)

   Ion         m/z   Intensity
   b1+       367.18       14.8
   b5++      376.43       17.7
   b6++      404.80       21.4
   a7++      447.66       18.4
   y7++      456.83       18.9
   b7++      461.89       57.8
   y8++      492.23       19.6
   a8++      497.31       27.5
   b8++      511.44       82.4
   b12+++    536.96       32.2
   b9++      561.95       45.1
   b13+++    575.15       25.6
   y14+++    580.73       22.2
   b14+++    594.11       33.0
   b10++     597.57       34.2
   a11++     612.07       15.0
   y15+++    614.40       44.2
   a15+++    617.88       27.2
   b11++     626.15       26.7
   b15+++    627.26       51.4
   b4+       638.46       26.2
   a16+++    641.59       16.8
   y16+++    647.28       25.5
   b16+++    650.83       46.5
   y17+++    685.54       39.7
   b17+++    688.74       36.5
   y5+       699.21       52.6
   y18+++    704.79       30.1
   y19+++    742.42       14.6
   y6+       798.41       21.6
   y21+++    799.83       15.0
   y12++     806.54       16.7
   b6+       808.49       22.8
   b20+++    814.79       32.1
   y13++     835.14       65.3
   a13++     848.23       15.2
   b21+++    858.34       28.2
   b13++     862.31       26.7
   y14++     870.71       60.0
   a22+++    886.70       18.0
   b14++     890.76       25.7
   b22+++    896.45       23.1
   y7+       912.42       46.1
   y15++     921.18       96.4
   b7+       922.16       52.9
   b15++     940.32       36.7
   y16++     970.47       25.7
   b16++     975.47       19.9
   y8+       983.40       39.6
   b8+      1021.62       27.1
   y17++    1027.83       25.6
   b17++    1032.87       18.7
   y18++    1056.25       26.7
   b9+      1122.61       15.6
   y10+     1139.55       17.6
   y11+     1254.55       19.8
```

```
487. MORG-339_phosphopeptide_TMT_18%ACN_032017_OT.16647.16647.3 (wLDEsDAEMELRAk)

   Ion         m/z   Intensity
   y4++      358.86       11.3
   b1+       416.32        8.0
   y2+       447.35        7.9
   y6++      488.97       16.2
   a6++      514.30        7.2
   b2+       529.43        8.6
   y7++      553.18        6.8
   y8++      588.97       27.2
   b3+       644.40       23.4
   y9++      646.34       17.0
   y4+       716.95       16.0
   y10++     729.93       14.3
   b4+       773.35        6.8
   y11++     794.48       11.3
   b11++     815.08        6.6
   y12++     852.15        8.0
   b12++     893.09        7.1
   y13++     908.48       10.8
   a13++     914.77        9.4
   a6+      1027.68        9.0
   b6+      1055.49       12.1
   b10+     1515.59        7.1
```

```
488. MORG-339_phosphopeptide_TMT_18%ACN_032017_OT.11020.11020.3 (iTRkPASVSPttPTSPTEGEAs)

   Ion         m/z   Intensity
   y9++      479.27       28.7
   b6++      563.73       22.3
   b7++      607.19       27.8
   a8++      642.36       16.9
   b8++      656.61       55.4
   b9++      699.90       19.9
   b11++     839.31       27.9
   y15++     850.67       25.8
   y8+       857.35       28.4
   y16++     894.54       59.8
   a4+       929.27       27.5
   y17++     930.17       38.5
   b4+       957.46       25.6
   y9+       958.74       34.8
   a14++    1014.39       32.0
   b14++    1028.82       21.0
   b5+      1054.32       44.3
   y10+     1055.36       72.7
   y11+     1236.37       33.0
   y12+     1417.52       20.2
   y14+     1601.52       23.6
```

```
489. MORG-339_phosphopeptide_TMT_18%ACN_032017_OT.8127.8127.3 (tASPPGPPQYSk)

   Ion         m/z   Intensity
   y1+       376.38       13.0
   y4++      377.43        9.4
   b6++      410.61       13.1
   y5++      426.25        7.9
   a7++      445.00        8.4
   y2+       463.39       22.5
   y6++      474.96       29.8
   b8++      507.76        6.8
   a3+       541.71       13.8
   y8++      551.57       30.1
   a9++      557.67      130.6
   b3+       569.58       22.6
   b9++      572.04       25.9
   y9++      600.52       16.6
   y3+       626.42       17.4
   a10++     639.24       13.9
   b10++     653.60       33.5
   b4+       666.22        6.5
   a11++     683.19        7.2
   b11++     696.66        6.5
   y4+       754.46        6.7
   b5+       763.39       12.4
   a6+       792.64        8.0
   b6+       820.42       21.3
   y5+       851.48       11.4
   b7+       917.53        7.4
   y6+       948.52        6.7
```

```
490. MORG-339_phosphopeptide_TMT_30%ACN_032017_OT.6341.6341.4 (iRAkGTPALTSEAAQSsPPTR)

   Ion         m/z   Intensity
   y8++      462.42       20.6
   y4+       470.31       50.2
   a2+       471.29       23.9
   b10+++    489.69       18.8
   b11+++    519.24       25.1
   y10++     562.15       58.0
   y16+++    565.01       24.2
   b13+++    585.93       90.4
   y11++     605.64       31.1
   b14+++    609.55       95.9
   b8++      627.29       29.3
   y5+       637.14       48.2
   a15+++    642.90       38.0
   b15+++    652.21       99.4
   y12++     656.42       50.1
   b16+++    681.34       55.7
   b9++      684.23       21.0
   y18+++    703.46       30.3
   a10++     720.82       23.6
   y6+       724.33       47.7
   a17+++    727.23       20.5
   b10++     734.20       32.8
   b17+++    736.94       23.7
   a18+++    759.81       24.2
   b11++     778.06       28.4
   a12++     828.97       26.3
   b20+++    835.37       41.7
   b12++     842.74       37.1
   y7+       852.34       28.4
   b13++     878.27       45.8
   b14++     913.79       45.5
   y8+       923.38       37.1
   y9+       994.35       25.3
```

```
491. MORG-339_phosphopeptide_TMT_4%ACN_032017_OT.6455.6455.2 (sRTAsGssVTSLEGTR)

   Ion         m/z   Intensity
   a4+       617.36        7.0
   b4+       645.38        7.6
   b5+       812.35        6.5
   a8+      1175.60        8.0
   y11+     1253.49        6.2
   b9+      1302.55        7.0
   y14+     1592.68        6.3
```

```
492. MORG-339_phosphopeptide_TMT_10%ACN_032017_OT.12171.12171.4 (kEtEsEAEDDNLDDLERHLR)

   Ion         m/z   Intensity
   y4++      291.20       16.7
   y6++      412.58       21.0
   b9+++     470.02       28.6
   a10+++    499.53       25.9
   b11+++    546.49       10.7
   a13+++    612.79       29.0
   y10++     641.14       45.4
   a15+++    689.68       13.3
   a9++      690.48       16.2
   b15+++    698.57       30.3
   y5+       710.18      124.1
   y17+++    712.72       15.4
   b16+++    741.38       34.8
   a10++     748.54       14.7
   b11++     819.60       23.7
   y13++     820.65       30.7
   y14++     856.09       39.5
   a19+++    867.93       13.9
   b19+++    877.07       25.1
   y15++     920.43       30.5
   b5+       964.45       14.1
   b14++     991.05       12.1
   y16++    1004.11       24.7
```

```
493. MORG-339_phosphopeptide_TMT_25%ACN_032017_OT.21150.21150.3 (akQPVIGDQNsDsDEMLAVLk)

   Ion         m/z   Intensity
   y1+       376.37       19.3
   b3++      393.97       21.7
   y2+       489.31       29.0
   b5++      492.01       25.1
   b6++      548.54       28.9
   b7++      576.95       14.7
   y3+       588.52       21.8
   b8++      634.47       47.1
   b2+       658.51       17.3
   y4+       659.47       33.2
   b9++      698.50       19.6
   b10++     755.75       26.3
   y5+       772.68       27.9
   b3+       786.65       22.6
   a12++     882.52       14.9
   b12++     896.57       30.7
   y6+       903.60       25.0
   y13++     919.44       14.2
   a13++     966.04       20.8
   b13++     979.75       18.5
   b5+       982.67       22.9
   y15++    1006.23       14.8
   y16++    1062.76       16.5
   b6+      1095.71       23.0
   b15++    1101.77       17.2
   y17++    1112.27       14.7
   y8+      1147.55       19.3
   b7+      1152.85       14.4
   b16++    1167.60       34.2
   y19++    1224.34       36.9
   b18++    1259.89       38.1
   b8+      1267.89       24.0
   b19++    1308.98       24.5
   y9+      1314.78       26.9
   b20++    1365.79       23.6
   y10+     1429.64       15.9
```

```
494. MORG-339_phosphopeptide_TMT_18%ACN_032017_OT.3608.3608.3 (kQQQEPTcEPsPk)

   Ion         m/z   Intensity
   y2+       473.37       36.7
   b2+       486.51       14.1
   y7++      564.32       11.1
   y8++      613.03        8.4
   y3+       640.30       12.8
   a4+       714.23        8.8
   b10++     728.15       11.1
   y4+       737.48       33.9
   b4+       742.15       40.4
   a5+       843.64        9.3
   a12++     846.87        8.8
   b12++     860.62       10.4
   y5+       865.94       10.1
   y12++     869.58        9.4
   y8+      1224.60       11.6
```

```
495. MORG-339_phosphopeptide_TMT_10%ACN_032017_OT.7061.7061.3 (sEDDSAVPVAkAAPR)

   Ion         m/z   Intensity
   b1+       397.25        9.9
   y8++      519.99       37.6
   y9++      569.55       17.7
   b9++      605.12       13.8
   a10++     626.91       12.3
   b3+       641.20       24.5
   y11++     648.41       44.0
   y12++     706.06       30.4
   b4+       756.35       23.3
   y13++     763.56       20.7
   y5+       771.82       14.3
   b11++     819.19       17.5
   y14++     828.03       27.1
   y6+       842.57       11.4
   b12++     855.09       13.7
   b13++     890.79       27.8
   b6+       914.37       13.6
   b7+      1013.40       22.4
   y8+      1038.69       25.7
```

```
496. MORG-339_phosphopeptide_TMT_4%ACN_032017_OT.5342.5342.4 (tREQEssGEEDNDLSPEEREk)

   Ion         m/z   Intensity
   y4++      395.83        6.4
   b5++      437.28       20.8
   b2+       487.52        6.3
   a6++      506.41       18.9
   y6++      508.94       17.1
   y8++      609.20       29.5
   b8++      632.39        7.7
   y9++      666.82        8.1
   a9++      683.70        6.2
   b9++      697.40       18.5
   a16+++    712.22        7.0
   b16+++    721.64       13.7
   y10++     723.37       28.7
   a17+++    756.00       17.1
   b10++     762.13       16.6
   y11++     781.08       11.1
   y4+       790.59        6.0
   b11++     819.32       12.7
   a5+       845.80       21.9
   a12++     862.51        6.2
   b5+       873.57        5.7
   y20+++    918.09        7.8
   b13++     934.10       18.6
   y14++     938.71       20.0
   b14++     990.45        5.3
   y15++    1021.95        8.3
   b15++    1034.22        5.0
   y16++    1105.46        7.2
   b17++    1146.97        7.5
   y8+      1216.52        6.0
   b19++    1289.78        5.3
```

```
497. MORG-339_phosphopeptide_TMT_22%ACN_032017_OT.14627.14627.4 (lkQLsVPAsDEEDEVPAPIPRGR)

   Ion         m/z   Intensity
   b3++      415.22       15.1
   y8++      432.23       12.9
   y9++      482.01       22.6
   a9+++     506.07       18.5
   y10++     546.54       20.3
   b6++      603.96       13.1
   b12+++    639.28       10.5
   y12++     668.67       14.6
   a8++      674.14       11.2
   b8++      688.39       11.3
   y6+       695.26       12.3
   a14+++    711.33       10.1
   y19+++    727.87       25.8
   y13++     732.55       95.2
   a15+++    744.49       11.2
   b15+++    753.58       13.3
   a9++      758.08       31.1
   y20+++    764.91       19.1
   y7+       766.93       25.6
   y14++     790.68       10.0
   y21+++    808.06       12.9
   b10++     829.30       12.8
   y8+       863.61       11.3
   a19+++    870.69        9.4
   y16++     909.57       24.0
   b4+       941.42       10.9
   b12++     958.19       25.4
   b22+++    982.85       14.2
   y18++    1007.77       15.4
   b13++    1016.08       11.8
   a5+      1080.85       20.0
   b15++    1129.84       11.4
```

```
498. MORG-339_phosphopeptide_TMT_6%ACN_032017_OT.4566.4566.3 (rASYDSVEDSSESENsPVAR)

   Ion         m/z   Intensity
   b5++      411.83       44.0
   y4+       442.29       39.2
   b6++      455.41       15.6
   b7++      505.11       16.5
   a8++      555.59       10.6
   b8++      569.47       28.5
   y10++     578.32       10.6
   b9++      627.01       75.1
   b10++     670.26       18.0
   b4+       707.39        9.9
   b11++     714.02       24.9
   y6+       723.25       19.5
   a12++     764.25       15.4
   b12++     778.53       54.5
   y14++     793.74       65.4
   a5+       794.47       28.1
   a13++     807.91       15.6
   b13++     821.78       23.8
   b5+       822.49       23.3
   a14++     872.46       10.0
   b14++     886.66       32.3
   y8+       939.40       30.5
   b15++     943.38       17.5
   b7+      1008.51       12.2
   b16++    1026.51       14.5
   y9+      1068.48       11.8
   a18++    1111.33       16.5
   b18++    1124.90       12.7
   b8+      1137.62       19.3
   a19++    1146.41       11.1
   y10+     1155.41       15.5
   b19++    1160.46       14.4
   a9+      1224.56       11.5
   y11+     1242.56       31.1
   b9+      1252.72       18.6
   b10+     1339.70       10.3
   y12+     1357.52       13.0
   y15+     1672.73       13.2
```

```
499. MORG-339_phosphopeptide_TMT_16%ACN_032017_OT.15961.15961.3 (fSHSYLsDsDTEAkLTETSA)

   Ion         m/z   Intensity
   b6++      482.87       14.3
   y5+       508.24       13.0
   b3+       601.50       11.1
   y6+       621.38       15.3
   b8++      623.70       11.6
   b4+       688.58       12.7
   b10++     765.13       17.6
   y12++     781.85       12.1
   a11++     801.51       11.6
   b11++     815.65       13.6
   y13++     838.39       37.7
   b5+       851.45       18.1
   a12++     865.39       31.2
   b12++     880.15       20.5
   a13++     901.12       27.7
   b13++     915.58       23.3
   b6+       964.55       16.4
   y15++     978.66       18.5
   y8+      1049.50       22.5
   b14++    1094.27       30.6
   y17++    1103.66       15.0
   b15++    1151.01       30.5
   y9+      1178.69       13.0
   b8+      1246.63       10.4
   b17++    1265.87       23.5
   y10+     1279.76       15.2
   y11+     1394.58       10.8
   y12+     1561.78       10.7
```

```
500. MORG-339_phosphopeptide_TMT_18%ACN_032017_OT.10695.10695.3 (asSLNENVDHSALLk)

   Ion         m/z   Intensity
   y1+       376.35       16.9
   a5++      377.29       11.3
   a2+       440.62       11.0
   y6++      449.46       19.8
   y2+       489.49       22.5
   y3+       602.55       24.6
   y9++      613.70       18.9
   a4+       640.13       14.2
   b4+       668.54       31.4
   y4+       673.48       38.1
   a10++     674.45       71.7
   y10++     677.91       22.3
   b10++     688.54       45.6
   b11++     732.30       17.2
   y11++     735.23       16.2
   a5+       754.09       10.5
   y5+       760.45       29.3
   b12++     767.84       12.5
   b5+       782.63       10.5
   y12++     791.81       26.7
   a13++     810.75       13.3
   b13++     824.68       31.6
   a14++     866.82       12.9
   b14++     880.60       15.4
   y6+       897.58       13.2
   y7+      1012.58       11.4
   b7+      1025.49       11.0
   b8+      1124.51       11.7
   b9+      1239.57       15.3
```

```
501. MORG-339_phosphopeptide_TMT_12%ACN_032017_OT.3916.3916.4 (hRRLLEDsEsEDEAPPSRPR)

   Ion         m/z   Intensity
   y6++      355.22       20.6
   y7++      390.90       33.8
   b5++      453.61        9.8
   y8++      455.28       24.3
   y9++      513.24       14.6
   b6++      517.72       16.2
   b7++      575.56       50.6
   y14+++    577.46       19.8
   a12+++    609.85       15.7
   y5+       612.47       16.1
   y15+++    621.06       19.9
   a3+       651.39       12.4
   a13+++    653.71       25.7
   b8++      658.94       33.8
   y11++     661.15       16.8
   b13+++    662.66       11.8
   a14+++    676.64       31.0
   y6+       709.55       13.4
   b9++      723.70       13.7
   b16+++    750.86       12.5
   a4+       764.72       10.1
   a10++     792.48       12.0
   y19+++    799.96       17.7
   y13++     809.07       30.2
   a18+++    822.66       11.3
   b18+++    831.74       18.9
   b11++     870.99       13.8
   b12++     929.01       20.0
   y15++     931.24       14.4
   b13++     993.54       25.8
   b14++    1029.12       18.7
```

```
502. MORG-339_phosphopeptide_TMT_25%ACN_032017_OT.17635.17635.3 (qIkNSSLLSFDsEDENE)

   Ion         m/z   Intensity
   b3++      415.21       19.2
   b4++      471.71       13.5
   b5++      515.32       15.7
   b6++      558.76       23.3
   b7++      615.53       48.9
   b8++      672.15       94.8
   b9++      715.76       37.8
   a10++     774.95       15.2
   y13++     776.25       13.0
   b10++     789.19       47.2
   a3+       800.58       27.3
   y6+       802.54       33.0
   b11++     846.44       24.0
   y7+       917.31       16.5
   b4+       942.80       13.9
   y15++    1011.96       13.2
   y8+      1064.30       25.9
   y16++    1068.70       15.0
   b6+      1116.83       58.5
   y9+      1151.36       42.0
   b16++    1173.86       33.0
   y10+     1264.69       19.6
   y12+     1464.82       13.3
```

```
503. MORG-339_phosphopeptide_TMT_18%ACN_032017_OT.2085.2085.4 (tkkDDSHSAEDsEDEk)

   Ion         m/z   Intensity
   y1+       376.32       19.9
   b4+++     387.77       19.3
   b5+++     426.08       25.4
   b7+++     500.59       14.5
   y2+       505.37       23.9
   b3++      523.75       14.6
   b8+++     530.07       18.6
   b9+++     553.08       16.6
   b4++      580.95       17.7
   b10+++    596.47       32.4
   y3+       620.40       25.2
   b11+++    634.82       56.7
   b5++      638.73       32.0
   y9++      659.68       19.6
   a2+       660.44       19.4
   a6++      667.48       27.9
   a12+++    680.90      191.2
   b6++      681.86       57.4
   b2+       688.21       43.3
   b12+++    689.93       33.1
   a13+++    723.72       15.4
   b13+++    733.55       21.1
   a7++      736.75       15.5
   y4+       749.34       23.0
   b7++      750.40       20.3
   y11++     771.97       33.4
   b15+++    814.73       23.5
   a9++      815.56       15.6
   y15+++    829.65       14.8
   b10++     894.11       14.4
   y5+       916.38       20.3
   b11++     951.44       12.8
   y11+     1542.64       11.9
```

```
504. MORG-339_phosphopeptide_TMT_4%ACN_032017_OT.7795.7795.3 (dHsPTPSVFNsDEER)

   Ion         m/z   Intensity
   y2+       304.16       10.6
   a3++      310.89        8.6
   a1+       317.28       23.4
   b1+       345.29        7.4
   y6++      415.11       49.3
   b5++      424.13       10.7
   y3+       433.23       16.2
   a2+       454.25        7.4
   a6++      458.51        8.7
   b2+       482.38       10.8
   y7++      488.93        7.5
   b7++      516.32       10.5
   y4+       548.24       12.5
   a8++      551.37       26.0
   b8++      565.90       15.3
   y9++      581.41       22.7
   a3+       621.31       12.7
   a9++      625.45        8.7
   y10++     630.45       45.5
   b9++      639.33       21.2
   b3+       649.30        9.8
   y11++     680.40       13.8
   y12++     729.44       12.5
   b11++     779.43        7.8
   y13++     812.72       10.1
   a12++     823.71        9.9
   y6+       829.42       38.1
   b12++     837.48       18.9
   b5+       847.45       14.4
   y14++     881.51        7.6
   a13++     888.10       13.5
   b13++     901.97       21.8
   b14++     966.31       14.4
   y7+       976.35       26.3
   y8+      1075.49       16.4
   y9+      1162.63       13.7
   y10+     1259.52       24.2
```

```
505. MORG-339_phosphopeptide_TMT_6%ACN_032017_OT.12534.12534.2 (aDGYEPPVQEsV)

   Ion         m/z   Intensity
   a2+       387.76        4.7
   y3+       414.14        6.0
   b3+       473.02       11.4
   y4+       542.09       15.3
   b4+       636.50       12.4
   a5+       737.47        5.4
   y6+       738.40        7.4
   b5+       765.39       40.1
   y7+       835.37       39.4
   b6+       862.46        4.2
   b7+       959.80        5.2
   y8+       964.59        8.5
   b8+      1058.58       23.8
   b9+      1186.58       10.9
   b10+     1315.72       12.9
```

```
506. MORG-339_phosphopeptide_TMT_16%ACN_032017_OT.9451.9451.3 (aAVLsDsEDDAGNASAkk)

   Ion         m/z   Intensity
   y1+       376.38        9.2
   y4++      446.42        7.9
   b3+       471.34        8.2
   y5++      481.85        7.7
   b4+       584.46       12.2
   y8++      603.06       23.2
   y9++      660.52       18.2
   a11++     718.04       13.0
   y2+       733.56       11.4
   b12++     760.52        8.4
   y11++     782.21       16.7
   a13++     804.06       13.3
   b13++     817.61       23.5
   a6+       838.43       11.3
   b14++     852.93       17.2
   a15++     882.95        8.1
   y4+       891.66        8.8
   y13++     923.69       14.1
   y5+       962.16        8.9
   y14++    1006.97       10.1
   b7+      1033.36        8.6
   y15++    1063.71       10.5
   b17++    1111.29       11.2
   b9+      1277.66        7.9
   b10+     1392.56        7.8
```

```
507. MORG-339_phosphopeptide_TMT_25%ACN_032017_OT.21433.21433.3 (rLsLPGLLSQVsPR)

   Ion         m/z   Intensity
   y2+       272.19       25.5
   b3++      333.69       10.6
   b4++      390.27       13.7
   y3+       439.19       14.5
   b6++      467.47       12.2
   a7++      509.94       19.1
   y9++      518.33       12.5
   b7++      523.77       25.6
   y4+       538.44       10.0
   a8++      566.53       22.4
   y10++     567.42       28.3
   b8++      580.39       24.0
   b9++      624.04       15.1
   b3+       666.36        8.3
   b10++     688.08       18.4
   a11++     723.04       31.6
   b11++     737.48       26.4
   y6+       753.40       23.3
   y13++     763.41       20.5
   b4+       779.53       22.0
   y7+       866.53       16.2
   y8+       979.76        9.7
   y9+      1036.56       16.7
   y10+     1133.59       25.5
   b8+      1159.52        7.6
```

```
508. MORG-339_phosphopeptide_TMT_12%ACN_032017_OT.5153.5153.3 (rTAFYNEDDsEEEQR)

   Ion         m/z   Intensity
   y2+       303.23       12.1
   a4++      339.23        9.9
   b6++      491.99       11.5
   a7++      542.58        9.2
   b7++      556.45       21.7
   b8++      613.96       35.4
   y10++     665.75       11.1
   b9++      671.48       35.6
   y5+       690.49       22.6
   b4+       704.99       21.5
   a11++     805.81       16.6
   y12++     821.18       12.6
   y13++     856.38       12.5
   y6+       857.41       18.0
   b12++     883.88       14.3
   b13++     948.54       28.7
   y7+       972.34       22.8
   b14++    1012.33       14.9
   y8+      1087.54       10.6
```

```
509. MORG-339_phosphopeptide_TMT_16%ACN_032017_OT.11803.11803.3 (sFsLGDLSHsPQTAQHVER)

   Ion         m/z   Intensity
   b1+       317.27       16.5
   y9++      533.26       21.4
   y4+       540.38       12.9
   a9++      612.95       20.9
   y10++     616.83       13.9
   y5+       668.40       15.0
   y11++     685.31       19.0
   a10++     696.45       10.7
   b10++     711.04       15.7
   y12++     728.95       16.7
   y6+       739.44       28.0
   b4+       744.60       10.2
   y13++     785.45       36.5
   b5+       801.58       20.8
   a12++     809.47       20.4
   y7+       840.46       15.6
   y14++     842.73       11.5
   y15++     871.56       10.0
   a6+       888.00       10.1
   a14++     895.06       10.8
   b6+       916.41       25.3
   y16++     928.09       20.2
   b15++     973.65       14.6
   b7+      1029.51       13.6
   y9+      1065.53       33.5
   b17++    1091.59       15.8
   b8+      1117.01        9.6
   b18++    1156.00       19.1
   y10+     1232.62       13.0
   b9+      1253.58       10.0
```

```
510. MORG-339_phosphopeptide_TMT_20%ACN_032017_OT.10220.10220.4 (sREcSLSPkStSPPPSPEVWAESR)

   Ion         m/z   Intensity
   y8++      487.43       20.2
   b7++      525.44       16.1
   y10++     579.07       14.3
   a11+++    581.86       34.7
   b12+++    620.30       19.5
   y11++     628.09       30.1
   b13+++    652.71       14.6
   y12++     676.61       63.3
   a15+++    708.27       22.8
   y13++     720.27       14.6
   a16+++    737.45       13.9
   a9++      738.74       12.8
   b16+++    746.53       16.7
   y6+       747.38       13.3
   y18+++    750.20       15.9
   b9++      752.61       12.9
   a17+++    769.21       17.6
   b17+++    779.11      116.9
   a10++     781.77       29.6
   y19+++    787.35       34.7
   y20+++    816.49       19.6
   a11++     872.64       17.5
   y7+       876.42       13.9
   b11++     886.72       19.6
   a20+++    907.79       21.4
   b12++     930.00       20.9
   a21+++    930.87       15.2
   y8+       973.55       27.9
   a22+++    974.51       20.0
   b13++     978.05       12.5
   b22+++    983.79       28.7
   a14++    1013.44       14.5
   a15++    1061.51       19.1
   y17++    1081.26       14.6
   y10+     1157.57       25.5
   y11+     1254.75       13.3
   y12+     1351.72       46.5
```

```
511. MORG-339_phosphopeptide_TMT_20%ACN_032017_OT.12139.12139.3 (rPsLLSEFQPGSER)

   Ion         m/z   Intensity
   y5++      273.17       10.9
   a5++      424.81       17.0
   b5++      438.90       13.8
   y5+       545.34       20.7
   b7++      546.43       25.3
   y10++     575.42        8.9
   b8++      620.49       35.0
   a3+       622.55       16.7
   y11++     631.81        9.0
   y6+       673.33       12.7
   b9++      684.61       21.9
   b10++     732.43       10.3
   a4+       735.38       23.5
   b11++     761.53        8.2
   y7+       820.48       13.7
   b13++     869.90        8.4
   y8+       949.52       15.7
   y9+      1036.49       15.6
   y10+     1149.62        8.2
```

```
512. MORG-339_phosphopeptide_TMT_6%ACN_032017_OT.6845.6845.2 (cRAVTcDIsEDEED)

   Ion         m/z   Intensity
   y5+       636.41        8.6
   b5+       760.01       50.1
   y6+       803.32        7.9
   y7+       916.24       23.2
   y9+      1191.35       10.1
   y10+     1292.40       17.9
   y12+     1462.69       10.3
```

```
513. MORG-339_phosphopeptide_TMT_4%ACN_032017_OT.7631.7631.3 (aSRVPssDEEVVEEPQSR)

   Ion         m/z   Intensity
   a1+       273.02        7.6
   b4++      322.26       15.5
   b5++      371.11        6.9
   a6++      439.96       16.8
   y4+       487.28       26.2
   b8++      595.29       21.3
   y5+       616.31       24.0
   b4+       643.54       14.7
   b9++      659.97       20.1
   a5+       712.41        7.3
   b10++     724.59       36.9
   y6+       745.40       27.9
   a11++     759.71       18.2
   b11++     774.09       46.3
   b12++     823.66       25.6
   y7+       844.41       30.8
   a6+       879.01       11.7
   b13++     888.11       34.3
   a14++     938.40       11.3
   y8+       943.49       24.4
   b14++     952.64       36.1
   y9+      1072.58       18.4
   b7+      1074.40        7.9
   b8+      1189.48        8.3
   y10+     1201.58       16.9
   y11+     1316.66        9.8
   y12+     1483.59        7.6
   y13+     1650.73        9.5
   y14+     1747.66       13.7
```

```
514. MORG-339_phosphopeptide_TMT_25%ACN_032017_OT.14509.14509.4 (lkQLSVPAsDEEDEVPAPIPRGR)

   Ion         m/z   Intensity
   b8+++     432.46       41.9
   a4++      457.63       15.4
   b4++      471.50       25.9
   y9++      481.82       42.4
   b5++      514.74       25.0
   y10++     546.50       48.9
   a6++      550.12       28.8
   b6++      564.31       39.8
   y11++     604.09       25.6
   y16+++    606.48       18.8
   y17+++    638.78       21.0
   y12++     668.52       31.6
   a14+++    684.43       15.8
   b14+++    694.03       16.5
   y6+       695.44       16.3
   a15+++    717.26       20.6
   a9++      718.31       26.6
   b15+++    727.42       30.5
   b9++      731.65       44.2
   y13++     732.60       73.0
   y20+++    738.43       49.3
   y21+++    781.20       20.6
   b10++     789.32       29.8
   y14++     790.52       46.9
   a19+++    843.40       17.1
   b11++     853.97       16.7
   y8+       863.59       29.7
   y15++     873.64       15.3
   y22+++    900.95       17.3
   y16++     909.18       53.5
   b12++     918.62       23.7
   a21+++    928.28       19.7
   b22+++    956.69       23.7
   y17++     957.85       21.8
   y9+       962.55       29.1
   b13++     976.29       31.2
   b5+      1028.70       19.3
   b14++    1040.72       54.7
   y19++    1050.93       19.4
   b15++    1090.30       37.7
   b6+      1127.78       29.2
```

```
515. MORG-339_phosphopeptide_TMT_4%ACN_032017_OT.7490.7490.2 (syTPEYR)

   Ion         m/z   Intensity
   y2+       338.16       26.5
   b1+       397.36        9.7
   y3+       467.38        7.4
   y4+       564.28       60.2
   b2+       639.78       19.9
   b3+       741.31       11.5
   a4+       810.40       12.5
   y6+       908.41       16.9
   b5+       967.36       23.2
   b6+      1130.52       12.4
```

```
516. MORG-339_phosphopeptide_TMT_20%ACN_032017_OT.4965.4965.3 (aAVVtSPPPTTAPHkER)

   Ion         m/z   Intensity
   b3+       471.27       17.9
   y9++      632.98       12.8
   a11++     652.62       15.5
   y10++     682.14       19.7
   b12++     701.61       15.7
   a5+       723.50       23.1
   y11++     730.65      114.7
   a13++     736.33       28.8
   b13++     750.45       57.0
   b5+       751.41       37.6
   a14++     805.29       12.3
   b14++     819.17       12.5
   b6+       838.50       18.2
   y13++     864.77       14.2
   y14++     914.27       14.3
   y15++     963.35       12.2
   y6+       967.02       13.5
   b16++    1062.33       14.9
   y9+      1265.51       13.0
```

```
517. MORG-339_phosphopeptide_TMT_18%ACN_032017_OT.13547.13547.3 (aLDIYSAVDDAsHEk)

   Ion         m/z   Intensity
   b4++      321.83       13.3
   y1+       376.39       13.5
   b2+       414.15        9.7
   y5++      440.92       15.6
   y6++      498.22       24.7
   y2+       505.33       20.0
   b3+       529.38       19.9
   b8++      532.10       10.4
   y7++      555.72       12.2
   y8++      605.34       10.8
   y9++      640.77        9.9
   b4+       642.46       20.0
   b11++     682.25       13.7
   y10++     684.31       10.8
   b5+       805.53       16.5
   y4+       809.62       10.8
   y12++     822.31       14.1
   y5+       880.35       10.9
   b6+       892.50       11.8
   b14++     898.63       12.2
   b7+       963.52       15.5
   a8+      1034.48        9.5
   b8+      1062.68       14.4
   b9+      1177.63       25.6
   b10+     1292.66       18.5
```

```
518. MORG-339_phosphopeptide_TMT_16%ACN_032017_OT.6362.6362.4 (ssDPSLNEkWQEHGR)

   Ion         m/z   Intensity
   b1+       317.26       20.3
   y3+       369.22       15.5
   a5++      378.29       13.7
   y7+++     390.88       15.1
   y6++      406.78       12.0
   b2+       484.15       10.6
   y4+       498.28       20.0
   b9+++     499.29       24.5
   b7++      505.69       11.3
   y10+++    509.74       23.2
   a10+++    552.45       20.8
   a8++      556.58       31.8
   b10+++    562.06       23.4
   a3+       570.93       98.5
   y7++      585.58       24.3
   b3+       599.28       48.4
   b11+++    603.91       17.3
   y13+++    609.37       24.4
   y5+       626.38       20.5
   b12+++    647.75       20.0
   y8++      650.28       28.6
   a4+       668.43       15.2
   b13+++    693.26       17.2
   y9++      707.18       51.9
   b14+++    712.15       12.7
   b9++      749.10       33.7
   a5+       755.07       13.0
   y10++     763.31       16.4
   b5+       783.36       22.1
   y6+       812.41       26.8
   b6+       896.43       15.5
```

```
519. MORG-339_phosphopeptide_TMT_12%ACN_032017_OT.4536.4536.3 (kPAQETEETSSQEsAEED)

   Ion         m/z   Intensity
   a5++      378.60        7.0
   b6++      442.31       10.6
   b7++      507.49        9.5
   b9++      622.04       19.6
   b10++     665.79        8.0
   y12++     710.96        8.0
   y6+       758.87        8.6
   a13++     823.63       26.3
   y14++     825.71       34.3
   y7+       887.24        7.7
   y8+       974.18       16.1
   b7+      1013.66        6.5
   y9+      1061.37       13.1
   a17++    1071.75       11.1
   y10+     1162.46       14.5
```

```
520. MORG-339_phosphopeptide_TMT_20%ACN_032017_OT.8606.8606.3 (lSSSSATLLNsPDRAR)

   Ion         m/z   Intensity
   a2++      201.86        8.7
   y5++      307.70       17.6
   b6++      382.16       12.2
   a3+       488.91       10.5
   b9++      545.80       14.1
   y9++      561.05       17.4
   b10++     602.48       12.1
   b4+       604.38        9.2
   y5+       614.50       13.1
   y11++     647.67       31.6
   a11++     671.45        8.7
   b5+       691.50       10.3
   b6+       762.53        9.1
   a13++     777.55       14.7
   y15++     821.50       11.0
   b7+       863.66       10.3
   b14++     870.26       15.4
   y7+       895.58       16.7
   b15++     905.86       17.6
   b8+       976.75        8.7
   b9+      1089.76       10.9
   b10+     1203.76       11.7
```

```
521. MORG-339_phosphopeptide_TMT_25%ACN_032017_OT.17043.17043.4 (dIIRQPsEEEIIkLAPPPk)

   Ion         m/z   Intensity
   a1+       317.24       16.6
   y4++      334.30       21.4
   a7+++     364.28       12.1
   y1+       376.35       19.1
   a5++      414.47       12.6
   b5++      428.48       20.5
   y2+       473.43       25.2
   y3+       570.49       25.4
   b3+       571.45       17.5
   y7++      605.10       12.8
   y8++      661.14       16.7
   y4+       667.46       71.2
   a9++      675.09       13.6
   a13+++    688.41       11.7
   b9++      689.46       18.7
   y14+++    696.30       12.0
   b13+++    697.45       39.5
   y9++      717.56       38.2
   a14+++    725.36       23.8
   b4+       727.41       35.3
   a15+++    749.39       15.1
   b10++     754.09       17.3
   b15+++    759.07      129.9
   y10++     782.84       20.7
   b16+++    791.37       29.5
   b11++     810.72       14.1
   a17+++    814.35       14.0
   y11++     846.97       17.9
   a12++     852.87       18.0
   b18+++    855.57       14.3
   y18+++    866.20       11.5
   y13++     994.76       19.8
   y14++    1043.73       20.9
   b15++    1137.85       50.8
```

```
522. MORG-339_phosphopeptide_TMT_10%ACN_032017_OT.6659.6659.3 (gRTAsDSDEQQWPEEk)

   Ion         m/z   Intensity
   y1+       376.32       13.2
   b6++      449.34       10.1
   b7++      492.48       10.4
   y2+       505.33       11.1
   b8++      550.23       11.2
   b4+       614.99       16.8
   y3+       634.44       11.1
   b10++     679.01       19.2
   y9++      709.55       11.5
   a11++     728.74        9.8
   y4+       731.47       52.1
   b11++     743.06       39.5
   y10++     752.53       26.8
   b5+       782.41       44.8
   a12++     822.12       24.3
   b12++     836.08       48.1
   a13++     870.69       11.0
   b13++     884.52       12.0
   b6+       897.28       12.3
   y5+       917.46       23.7
   y13++     929.08       11.0
   a14++     934.96       12.9
   b14++     949.35       12.3
   y6+      1045.61       10.5
   y8+      1302.58       13.6
   y10+     1504.76       10.2
```

```
523. MORG-339_phosphopeptide_TMT_4%ACN_032017_OT.4802.4802.3 (sRsPLLNDR)

   Ion         m/z   Intensity
   y1+       175.11       15.6
   a2++      262.72       36.2
   b2++      277.14       11.6
   b3++      360.63       39.6
   y6++      364.24       50.8
   b1+       397.07        8.7
   y3+       404.21       23.9
   b4++      408.95        7.3
   a5++      451.51       27.1
   b5++      465.46       29.0
   a6++      508.22       11.1
   y4+       517.29       29.4
   b6++      522.39       23.9
   y8++      525.77       11.9
   b2+       553.45        7.5
   a8++      622.37       33.3
   y5+       630.36       10.1
   b8++      636.92       31.3
   b3+       720.31       14.5
   y6+       727.44       69.7
```

```
524. MORG-339_phosphopeptide_TMT_18%ACN_032017_OT.16145.16145.3 (nIQDVGAFstVk)

   Ion         m/z   Intensity
   y1+       376.28       66.1
   b5++      400.35       13.0
   b7++      464.31       14.5
   y2+       475.38       10.3
   y5++      485.69       17.0
   y6++      521.27       11.1
   b8++      537.95       29.0
   y7++      549.82       24.1
   a3+       557.46       15.2
   b3+       584.86       13.8
   y8++      599.28       19.0
   a9++      607.46       12.5
   b9++      620.84       22.3
   y9++      656.55       12.7
   b4+       700.39       13.1
   a11++     747.40       20.2
   b11++     761.30       34.1
   b5+       799.51       25.8
   y4+       823.39       15.4
   b6+       856.57       11.7
   b7+       927.52       27.3
   y5+       970.52       11.0
   b8+      1074.61        7.3
```

```
525. MORG-339_phosphopeptide_TMT_10%ACN_032017_OT.12254.12254.4 (rGPPRNYAGEEEEEGsGssEGFEPPAADGQFSGAR)

   Ion         m/z   Intensity
   y4+       390.28       20.4
   b7++      536.23       19.5
   y5+       537.32       36.7
   y11++     539.01       36.7
   y12++     587.51      161.2
   b9++      600.22       20.5
   b4+       637.39       19.5
   y13++     652.00       19.8
   b10++     664.57       27.6
   y6+       665.40       23.1
   y7+       722.37       52.7
   b11++     728.92       28.2
   b5+       793.39       22.6
   y22+++    794.13       18.8
   y8+       837.40       57.3
   b13++     857.86       25.1
   b21+++    882.54       19.5
   a14++     908.59       63.9
   b14++     922.32       34.8
   y25+++    922.95       35.6
   b15++     951.11       30.4
   a23+++    965.46       28.2
   b23+++    974.65      150.9
   y10+      979.40       43.5
   y27+++    984.76       18.4
   y18++     985.60       27.7
   a24+++    997.20       26.5
   b24+++   1006.79       76.9
   b25+++   1039.45       19.8
   y29+++   1062.91       40.3
   y11+     1076.52       68.7
   a27+++   1077.52       56.0
   b27+++   1086.80       46.5
   y20++    1097.34       19.9
   b28+++   1125.13       47.6
   y21++    1126.08       28.0
   a29+++   1134.52       17.7
   b29+++   1144.25       27.0
   y12+     1173.57       84.4
   a31+++   1226.54       21.6
   b19++    1229.92       19.5
   b31+++   1235.89       35.5
   a33+++   1274.58       21.4
   b20++    1294.38       17.7
   y13+     1302.57       22.4
   b34+++   1307.23       24.7
   b21++    1323.23       26.8
   b22++    1396.80       60.4
   b23++    1461.33      199.6
   y29++    1594.28       23.5
   b27++    1629.50       21.1
```

```
526. MORG-339_phosphopeptide_TMT_10%ACN_032017_OT.5991.5991.4 (sRsIsPcPkVDSR)

   Ion         m/z   Intensity
   y2+       262.18       40.3
   a2++      263.21       11.5
   b2++      277.29        9.6
   b3++      360.68       10.5
   y3+       377.21        8.6
   y8+++     396.62       20.1
   b1+       397.43        9.0
   a4++      402.84       47.2
   y5++      417.35        9.0
   a8+++     442.86        7.1
   b8+++     451.71       49.6
   y9+++     452.42       26.0
   y6++      465.72       18.8
   y4+       475.89       10.8
   a5++      486.71       11.0
   y10+++    489.82       12.8
   b5++      500.24       18.9
   a2+       525.44        9.2
   a6++      534.85       18.2
   a9+++     561.99       13.6
   b9+++     571.74       15.9
   a10+++    594.48       80.0
   y12+++    597.56        8.8
   b10+++    604.41       17.1
   b7++      628.97       34.2
   b11+++    642.75       69.1
   y9++      677.95       34.4
   y10++     734.42       14.3
   a4+       805.37        7.2
   y11++     817.78        7.1
```

```
527. MORG-339_phosphopeptide_TMT_4%ACN_032017_OT.10054.10054.3 (lARADsDGDLsENDDGAGDLR)

   Ion         m/z   Intensity
   y2+       288.33       25.7
   b5++      379.02       31.1
   b2+       413.85       14.0
   b7++      519.91       39.1
   y5+       531.52       14.7
   b8++      548.26       16.5
   y6+       588.38       35.8
   b9++      606.00       53.2
   a4+       613.57       17.7
   a10++     648.69       17.9
   b10++     662.63       38.2
   y7+       703.38       25.0
   b5+       756.40       13.6
   b12++     810.03       11.7
   y8+       818.77      248.3
   b14++     924.79       20.7
   y9+       932.44       27.7
   b15++     982.79       35.9
   b7+      1038.32       20.4
   y10+     1061.56       19.9
   b18++    1074.81       18.3
   b19++    1132.72       47.2
   b20++    1189.24       19.7
   b9+      1210.69       12.6
   y11+     1228.47       18.2
   b10+     1323.56       15.0
   y12+     1341.57       24.4
   y14+     1513.67       23.0
   y16+     1795.57       10.3
```

```
528. MORG-339_phosphopeptide_TMT_4%ACN_032017_OT.3104.3104.3 (eEQELHDIHsTR)

   Ion         m/z   Intensity
   a2++      230.36        7.1
   y2+       276.27       12.9
   a5++      415.55        8.3
   b5++      430.07        8.2
   y3+       443.18        9.0
   y7++      473.22       11.6
   b2+       488.39       11.1
   b6++      498.58       16.9
   y8++      530.01       21.2
   a7++      541.55        6.4
   b7++      556.05       21.2
   y4+       580.62       25.1
   y9++      594.70       17.4
   b8++      612.53       14.4
   b9++      680.71        9.7
   y5+       693.29        9.5
   a4+       717.63       10.3
   b4+       745.47       20.0
   b10++     764.81        9.5
   y6+       808.18        5.8
   b5+       858.49       13.2
```

```
529. MORG-339_phosphopeptide_TMT_12%ACN_032017_OT.4759.4759.3 (sRTASLTSAAsIDGSRSR)

   Ion         m/z   Intensity
   y5++      281.66       13.8
   y6++      339.33        8.3
   y7++      395.92       11.7
   y4+       504.92        9.1
   y5+       562.45       20.3
   a10++     614.02       12.0
   b10++     628.31        9.3
   y12++     644.21       10.1
   b3+       654.07        8.7
   y6+       677.50       14.5
   a4+       697.21       10.3
   a11++     697.92       14.1
   y13++     700.80       15.0
   b11++     711.37       21.1
   b4+       725.54       24.9
   b12++     768.17       10.5
   a5+       784.46        8.1
   y7+       790.67       10.1
   b5+       812.53        9.4
   b13++     825.97       22.2
   a14++     840.00        9.1
   b13+     1650.90       11.1
```

```
530. MORG-339_phosphopeptide_TMT_18%ACN_032017_OT.10255.10255.4 (wAHDkFSGEEGEIEDDEsGTENR)

   Ion         m/z   Intensity
   y3+       418.27       21.8
   y8++      494.11       13.6
   b10+++    549.43       20.5
   y9++      551.94       14.0
   b11+++    568.73       20.9
   y5+       576.35       22.9
   a6++      608.40       15.0
   b12+++    611.56       53.2
   y10++     616.15       15.1
   a13+++    639.89       16.7
   b13+++    649.28       66.0
   a14+++    682.93       16.2
   b14+++    692.22       80.8
   a4+       711.03       13.3
   b15+++    730.69       52.7
   b4+       739.21       14.6
   y6+       743.26       26.9
   a9++      744.46       27.3
   a16+++    759.37       28.4
   y13++     765.94       77.4
   b16+++    768.39       75.9
   b17+++    811.83       42.4
   y19+++    812.57       26.2
   b10++     823.61       21.9
   b11++     851.89       18.6
   b18+++    867.71       16.7
   y7+       872.34       17.4
   b19+++    886.89       14.4
   y15++     894.97       15.6
   b12++     916.69       51.1
   y22+++    920.35       24.2
   b21+++    963.40       42.8
   b13++     973.23       37.4
   y8+       987.39       21.6
   b22+++   1001.36       26.8
   b14++    1037.74       34.7
   b15++    1095.45       23.8
   y9+      1102.44       35.8
   b16++    1152.45       17.0
   y10+     1231.44       25.0
   y11+     1344.46       13.6
```

```
531. MORG-339_phosphopeptide_TMT_18%ACN_032017_OT.17284.17284.4 (lkQLsVPAsDEEDEVPAPIPR)

   Ion         m/z   Intensity
   y2+       272.21        9.8
   y6++      325.96       25.9
   y7++      375.26       11.1
   b3++      415.18       35.9
   b4++      471.35       12.4
   y4+       482.37       22.7
   y5+       553.45       10.1
   b6++      604.56       35.5
   b12+++    639.30       13.7
   y6+       650.42       77.3
   b7++      652.49       18.3
   a8++      674.32        8.9
   b13+++    678.02       17.6
   y12++     683.97       15.3
   b8++      688.23       43.2
   y18+++    694.17        9.8
   b2+       700.14       12.2
   a14+++    711.69       20.8
   y7+       749.57       10.4
   b15+++    753.91       63.6
   b16+++    786.46       13.3
   b17+++    810.19        8.6
   b10++     829.55       13.4
   a19+++    870.56        9.2
   y8+       878.12       12.9
   a11++     879.90       28.1
   b11++     894.00       18.9
   y16++     901.07        9.0
   a12++     944.21       10.4
   b12++     958.66       40.1
   b13++    1016.24       31.8
   a5+      1080.68       84.3
   b15++    1130.30       68.0
   b6+      1207.69       16.9
   b17++    1214.01       12.1
```

```
532. MORG-339_phosphopeptide_TMT_18%ACN_032017_OT.22320.22320.3 (iVETINSDsDsEFGIPk)

   Ion         m/z   Intensity
   b1+       343.25       12.8
   y1+       376.30       18.0
   b2+       442.34       14.3
   y2+       473.36      109.3
   b3+       571.48       22.0
   y3+       586.50       16.9
   y4+       643.53       36.8
   a4+       644.54       16.1
   b4+       672.51       12.1
   b10++     692.13        7.6
   y10++     742.50       11.0
   a11++     762.17        8.0
   b11++     775.82       10.0
   y11++     785.50       34.8
   b12++     840.44       18.8
   y12++     842.89       10.9
   a13++     899.64       24.2
   b13++     913.97       52.8
   y6+       919.74       10.9
   a14++     928.13       11.2
   b14++     942.51       89.7
   y14++     950.07       73.5
   a15++     985.04       52.6
   b7+       986.13       19.7
   b15++     999.02      145.0
   b16++    1047.72        8.2
   b8+      1101.54       13.2
   a9+      1240.65        7.5
   b10+     1383.70       11.1
   b12+     1679.71       18.7
```

```
533. MORG-339_phosphopeptide_TMT_10%ACN_032017_OT.12658.12658.3 (eAPAGGSGDGLSsPSPLEYSPR)

   Ion         m/z   Intensity
   y2+       272.21        9.5
   b1+       359.25       19.6
   b2+       430.31       16.2
   y7++      431.16        9.2
   b10++     514.74       12.3
   y4+       522.32       27.2
   y9++      523.42       22.6
   b11++     571.39       27.6
   b4+       598.40       18.8
   b12++     615.02       15.0
   y11++     650.23       17.8
   y5+       651.22       27.4
   b5+       655.34       16.6
   b13++     698.18       14.2
   b6+       712.37       10.5
   a14++     732.52       11.6
   y13++     735.56       17.3
   b14++     746.76       12.6
   y6+       764.17        9.8
   a7+       771.88       15.6
   a15++     776.13       25.1
   b15++     790.35       13.4
   y14++     792.66       17.1
   b7+       799.43       19.6
   a8+       828.20       12.6
   b8+       856.71       11.2
   y7+       861.54       14.2
   y16++     864.69       12.0
   b17++     895.41       19.9
   y8+       948.40       10.2
   b18++     959.92       16.4
   b9+       971.57       16.5
   a19++    1027.24       14.2
   b10+     1028.44       18.1
   b19++    1041.66       31.1
   y9+      1045.53       18.8
   b20++    1085.30       18.5
   b11+     1141.70       26.7
   y10+     1212.56       17.4
   b12+     1228.71       13.1
   y11+     1299.62       17.6
   b13+     1395.65       12.6
```

```
534. MORG-339_phosphopeptide_TMT_18%ACN_032017_OT.16041.16041.3 (sAFTPATATGSsPsPVLGQGEk)

   Ion         m/z   Intensity
   b1+       317.23       14.4
   y1+       376.31       16.6
   y7++      480.01       11.3
   y8++      529.10       21.9
   b3+       535.38       12.8
   y3+       562.37       27.0
   y9++      612.15       24.9
   b4+       636.42       25.4
   y10++     661.09       68.8
   y4+       690.44       13.8
   y11++     744.74       23.9
   y5+       747.45       43.2
   y12++     787.92       14.5
   b6+       804.62       15.8
   y13++     816.59       30.7
   b14++     826.53       15.3
   y6+       860.42       33.1
   a15++     861.49       22.6
   y14++     867.23       37.1
   b15++     875.67       14.2
   b16++     925.08       15.6
   y16++     953.05       16.7
   b8+       976.56       23.3
   b17++     981.74       21.2
   b18++    1009.46       10.9
   y18++    1037.42       22.8
   y8+      1056.57       13.8
   a19++    1059.65       11.3
   b19++    1074.27       12.8
   b9+      1077.58       26.2
   b20++    1102.77       13.7
   b10+     1134.57       20.6
   b21++    1167.38       14.5
   b11+     1221.59       18.7
   b12+     1388.65       17.1
```

```
535. MORG-339_phosphopeptide_TMT_8%ACN_032017_OT.3224.3224.3 (gAsASPQGRQsPsPSTRPIR)

   Ion         m/z   Intensity
   y3+       385.27       10.7
   y7++      414.01       15.4
   y8++      497.01        9.8
   y4+       541.46       10.5
   a10++     610.98       14.8
   y10++     629.03        7.4
   b11++     708.99        8.1
   a12++     743.06       14.0
   b12++     757.05       11.4
   y12++     771.13       84.3
   y13++     799.86       14.7
   a13++     826.77       17.5
   y14++     863.72       26.4
   b7+       908.63        8.0
   y15++     912.67       41.8
   b8+       965.56       12.4
   y17++     991.12        7.6
   y8+       993.62        6.9
   b17++    1061.71       12.8
   y9+      1090.49       19.8
   b19++    1166.51        7.4
```

```
536. MORG-339_phosphopeptide_TMT_22%ACN_032017_OT.5596.5596.4 (sGDPAPPDsPSWEAkkHR)

   Ion         m/z   Intensity
   b5++      329.32       16.0
   b3+       489.31       62.5
   y9+++     533.09       82.6
   y5++      549.53       25.8
   b4+       586.39       22.6
   y10+++    588.76       48.0
   y11+++    627.00       24.4
   a5+       629.31       25.4
   a11++     645.18       33.9
   b5+       657.43      107.8
   y12+++    659.00      207.3
   y3+       669.58       60.9
   y13+++    691.87       43.5
   y7++      707.17       17.0
   y14+++    715.37       40.6
   a6+       725.94       19.8
   a12++     738.05       23.3
   y15+++    747.96       79.7
   b6+       754.12       19.8
   y9++      799.15       41.8
   y17+++    804.83       18.8
   b16+++    807.02       17.6
   a7+       823.02       17.9
   b17+++    852.73       33.4
   y10++     882.47       30.0
   y11++     939.67       70.3
   b8+       966.62       35.9
   y12++     988.35       67.5
   y13++    1036.94       44.4
```

```
537. MORG-339_phosphopeptide_TMT_18%ACN_032017_OT.10582.10582.4 (sFsLGDLSHSPQTAQHVER)

   Ion         m/z   Intensity
   y6+++     247.25       11.1
   b1+       317.22       15.1
   y3+       403.36       11.8
   y11+++    430.72       24.9
   y12+++    459.83       17.6
   y13+++    497.50       30.2
   a7++      500.83       12.1
   y9++      533.44       30.7
   y14+++    535.82       21.4
   y4+       540.44       19.0
   a8++      545.04       12.3
   y15+++    554.68       21.9
   b13+++    556.31       14.2
   b8++      559.19       15.2
   y10++     576.83       32.5
   b14+++    579.58       20.1
   y16+++    592.53       39.2
   a9++      613.28        9.7
   b15+++    622.97        9.7
   b9++      627.05       17.2
   y11++     645.47       34.6
   y5+       668.55       26.5
   b10++     670.71        9.0
   y12++     689.02       27.9
   a18+++    734.82        9.4
   b4+       744.71       19.0
   b12++     783.71       18.3
   y14++     802.99       12.9
   a13++     819.59       12.8
   y7+       840.52       11.2
   b14++     869.53       11.6
   b6+       916.49       24.5
   b15++     933.84       14.8
   b7+      1029.56        9.9
```

```
538. MORG-339_phosphopeptide_TMT_14%ACN_032017_OT.12025.12025.4 (vQIPVSHPDPEPVsDNEDDsYDEEVHDPR)

   Ion         m/z   Intensity
   y2+       272.18       23.6
   b7++      496.03       14.4
   b3+       570.42       14.5
   b9++      602.05       50.6
   y10++     663.84       13.6
   b11++     715.21       35.7
   b5+       766.71       12.4
   b13++     812.95       16.0
   b6+       853.68       22.5
   y21+++    873.90       30.4
   y7+       881.03       12.0
   a21+++    895.11       28.3
   b14++     896.79       14.5
   y14++     900.54       28.4
   y22+++    905.73       27.3
   b15++     953.88       15.0
   a7+       962.25       15.4
   y24+++    980.22       12.8
   b7+       990.75       23.7
   y8+       996.47       17.4
   y25+++   1013.92       13.0
   b24+++   1028.62       12.8
   b17++    1075.95       21.6
   y27+++   1084.22       21.3
   b26+++   1107.41       40.4
   b18++    1133.22       23.9
   y18++    1139.58       19.8
   b27+++   1144.97       14.1
   a9+      1174.19       20.4
   a19++    1176.21       11.9
   b19++    1190.36       13.0
   b9+      1202.72       16.8
   y19++    1203.61       29.6
   y20++    1252.62       25.4
   y21++    1309.73       17.1
   y22++    1358.77       14.5
   a24++    1528.53       13.0
```

```
539. MORG-339_phosphopeptide_TMT_20%ACN_032017_OT.10871.10871.2 (vQsSPNLLAAGR)

   Ion         m/z   Intensity
   b1+       329.33       11.2
   y4+       374.37       12.0
   b2+       457.33       11.0
   b4+       711.63       17.2
   y7+       714.70       14.9
   b5+       808.55       11.6
   y8+       811.55       23.3
   y9+       898.52       31.0
   b6+       922.76       11.9
   b8+      1148.55       10.2
   b9+      1219.69       10.4
```

```
540. MORG-339_phosphopeptide_TMT_18%ACN_032017_OT.11049.11049.4 (aSGPLDTSQTAssDSEDVsSSDRkMSk)

   Ion         m/z   Intensity
   a12+++    466.62       17.0
   y5++      554.35       21.7
   y3+       594.61       22.1
   a16+++    633.02       23.3
   b5+       655.42       15.0
   y13+++    665.57       34.1
   a17+++    671.33       19.3
   a18+++    704.23       18.1
   a6+       742.42       17.1
   a19+++    759.30       23.8
   b6+       770.47       27.5
   a13++     783.19       14.7
   a14++     839.97       25.5
   a7+       843.23       18.5
   b14++     853.87       20.4
   b22+++    865.15       17.2
   b7+       871.41       21.2
   y18+++    872.75       86.5
   a15++     884.20       19.5
   a23+++    907.75       21.6
   y19+++    915.11       27.7
   b23+++    917.20       17.3
   y21+++    978.00       23.9
   y13++     997.03       14.6
   a17++    1006.06       16.0
   y22+++   1016.58       14.8
   a24+++   1026.81       22.9
   b24+++   1036.73       16.9
   b9+      1086.73       24.7
   y5+      1107.57       15.7
   a19++    1139.36       15.0
   a10+     1159.98       23.9
   b10+     1187.53       15.8
   y17++    1257.64       15.0
   b11+     1258.61       22.9
   b14+     1707.68       15.7
```

```
541. MORG-339_phosphopeptide_TMT_14%ACN_032017_OT.9183.9183.3 (dLLHPsPEEEk)

   Ion         m/z   Intensity
   y3++      317.19       14.3
   b1+       345.27        7.8
   b4++      354.69       15.9
   y1+       376.37       14.7
   y5++      430.84       16.2
   b2+       458.27        8.5
   a6++      472.81        6.5
   b6++      486.74       17.2
   y2+       505.32       18.2
   y6++      513.76       25.8
   b7++      535.68        7.4
   a3+       543.10        7.5
   y7++      562.88       54.4
   b3+       571.48       25.5
   a8++      585.79       16.0
   b8++      599.68       22.7
   y8++      631.16        9.9
   y3+       634.48       19.1
   b9++      664.43       19.2
   a4+       680.51       18.4
   b4+       708.42       50.1
   a10++     714.41        6.7
   b10++     728.95       14.8
   y10++     744.43        7.2
   y4+       763.44       10.3
   y5+       860.50       35.3
```

```
542. MORG-339_phosphopeptide_TMT_25%ACN_032017_OT.13916.13916.4 (sIVRQktVDIDDAQILPR)

   Ion         m/z   Intensity
   b1+       317.17       14.2
   y10+++    385.38       19.0
   b6+++     391.07       11.6
   y11+++    418.65       52.2
   a7+++     442.40       18.4
   b7+++     451.69       42.1
   a8+++     475.09       14.7
   b8+++     484.22       18.8
   y4+       498.37       17.8
   b9+++     522.75       50.4
   b3+       529.14       15.1
   b10+++    560.49       24.9
   a6++      572.23       11.7
   b11+++    598.81       41.8
   y5+       626.64       28.5
   y11++     627.82      199.6
   b12+++    637.08       45.5
   y14+++    641.30       34.1
   b4+       685.37       13.9
   a14+++    694.03       21.7
   y6+       697.43       32.9
   b14+++    703.54       32.8
   y16+++    725.81       12.1
   a15+++    731.96       13.6
   b15+++    741.40       21.2
   a9++      769.31       13.3
   b16+++    779.17       17.1
   b17+++    810.97       12.2
   y7+       812.53       28.6
   b5+       813.52       15.4
   b11++     897.40       11.7
   y8+       927.55       21.9
   y9+      1040.56       35.8
   b14++    1054.66       15.7
   y10+     1155.64       24.0
   y11+     1254.67       20.8
```

```
543. MORG-339_phosphopeptide_TMT_18%ACN_032017_OT.14958.14958.3 (gLYDGPVcEVSVtPk)

   Ion         m/z   Intensity
   b5++      368.21       12.1
   y1+       376.30       16.1
   y4++      377.45        9.4
   b2+       400.31        8.9
   b6++      416.88        9.5
   y5++      420.92       21.4
   b7++      466.31       19.9
   y6++      470.38       15.9
   y2+       473.40       63.3
   y7++      534.82        8.5
   b8++      546.51       17.4
   b3+       563.41       16.2
   b9++      610.97       38.8
   y8++      614.67       21.3
   a10++     646.61       16.4
   y3+       654.28       64.9
   b10++     660.47       62.0
   b4+       678.42       14.1
   a11++     689.52       27.1
   b11++     704.03       45.7
   a5+       707.08       13.5
   b5+       735.41       16.2
   y11++     741.38       11.0
   y4+       753.43       32.3
   y5+       840.47       40.7
   b13++     843.83       15.2
   b7+       931.53       41.1
   y6+       939.47       15.7
   b8+      1091.55       22.4
   b9+      1220.56       31.3
   b10+     1319.64       31.2
   b11+     1406.72       14.5
```

```
544. MORG-339_phosphopeptide_TMT_10%ACN_032017_OT.5870.5870.4 (sQEPISNDQkVsDDDk)

   Ion         m/z   Intensity
   b4++      336.48       20.3
   y1+       376.26       17.1
   a5++      378.73       25.3
   b5++      392.63       21.6
   b6++      436.30       22.5
   b2+       445.13       21.9
   y8+++     491.36       25.7
   y6++      494.29       29.3
   y9+++     529.54       42.4
   b8++      551.25       19.8
   a11+++    553.18       12.3
   b11+++    562.57       34.4
   y10+++    568.09       25.6
   b3+       574.36       35.2
   y11+++    597.16       56.9
   y3+       606.24       34.3
   a12+++    608.78       23.1
   b9++      615.18       84.7
   b12+++    617.60       32.9
   y12+++    634.54       37.9
   b13+++    656.62       18.4
   y13+++    667.17       27.5
   b4+       671.65       18.2
   y7++      673.00       30.3
   b14+++    695.11       15.4
   y4+       721.47       13.3
   b15+++    733.05       13.9
   y8++      737.04       20.3
   y15+++    752.90       14.4
   b5+       784.48       41.0
   b10++     793.66       42.8
   y9++      794.42       29.6
   a6+       843.17       39.6
   y10++     851.44       31.9
   b6+       871.54       18.7
   y5+       888.34       29.4
   y11++     895.13       52.7
   b7+       985.59       15.9
   y6+       987.55       21.1
   b8+      1100.49       14.5
```

```
545. MORG-339_phosphopeptide_TMT_8%ACN_032017_OT.1845.1845.3 (sRsQPcDLDAR)

   Ion         m/z   Intensity
   y1+       175.14        7.0
   y4++      237.24        6.3
   y2+       246.16       51.8
   a3++      306.65        6.7
   b1+       317.12        6.7
   b3++      320.67       13.3
   y3+       361.09       10.0
   b4++      384.66        9.7
   y7++      423.70       10.8
   b5++      433.48        8.1
   a2+       445.64        6.7
   b2+       473.03       13.5
   y4+       474.35       19.1
   y8++      487.89        7.2
   a6++      499.19       17.8
   b6++      513.29       40.4
   y9++      570.94       35.4
   y5+       589.67        7.0
   b8++      627.03       22.3
   a9++      670.46       11.0
   b9++      684.96      130.5
   b10++     720.20       16.2
   y6+       749.50        7.2
   b4+       768.41       13.7
   y7+       846.43       29.1
   y8+       974.42       11.2
```

```
546. MORG-339_phosphopeptide_TMT_22%ACN_032017_OT.20694.20694.3 (kIVETINsDsDsEFGIPk)

   Ion         m/z   Intensity
   b5++      400.43        9.2
   y2+       473.36       41.7
   y3+       586.52       11.5
   y4+       643.48       18.4
   y5+       790.55       17.1
   b5+       800.65        9.4
   a12++     866.27       10.4
   y6+       919.65       13.0
   y13++     939.17       16.2
   b13++     944.60       10.8
   y14++     989.51        9.4
   y7+      1086.47       16.4
   b16++    1103.10       10.0
   y17++    1160.80       15.3
   b8+      1195.10        9.1
   y9+      1368.57       11.1
```

```
547. MORG-339_phosphopeptide_TMT_18%ACN_032017_OT.13151.13151.4 (tEGYAAFQEDssGDEAESPSkVk)

   Ion         m/z   Intensity
   y1+       376.12       16.9
   y4++      460.21       12.4
   y2+       475.37       17.1
   y5++      509.00       44.1
   y6++      552.53       53.1
   b9++      614.13       17.9
   y7++      616.87       32.2
   y8++      652.40       29.5
   b10++     671.50       14.6
   b4+       680.52       19.5
   y14+++    685.38       17.3
   a17+++    717.13       30.3
   y15+++    728.53       25.7
   a11++     740.58       13.4
   a18+++    745.73       17.4
   b5+       751.43       36.0
   b11++     754.49       21.8
   y16+++    771.17       40.2
   y10++     774.76       14.6
   a19+++    778.25       14.5
   b19+++    787.45       40.7
   a6+       794.28       31.9
   y11++     802.97       38.9
   a20+++    807.62       22.1
   b14++     924.68       21.9
   b21+++    935.63       14.8
   y21+++    940.49       14.0
   b7+       969.44       36.2
   b15++     988.75       17.2
   b16++    1024.33       17.1
   y14++    1027.95       22.2
   b17++    1089.15       27.6
   b8+      1097.56       26.6
   b18++    1132.66       23.5
   b9+      1226.63       22.4
```

```
548. MORG-339_phosphopeptide_TMT_4%ACN_032017_OT.10037.10037.2 (gPPDFssDEER)

   Ion         m/z   Intensity
   b1+       287.12       11.2
   y2+       304.26       11.8
   y3+       433.33       12.0
   b3+       481.32        9.2
   y4+       548.20       11.1
   b4+       596.42       28.8
   a5+       714.99      114.2
   b5+       743.48       15.1
   a6+       882.34       12.2
   y7+      1029.33       12.1
   y8+      1144.58        6.1
   b8+      1192.45       13.0
   b9+      1321.62       12.2
   y10+     1338.49       14.0
   b10+     1450.49       11.8
```

```
549. MORG-339_phosphopeptide_TMT_10%ACN_032017_OT.7789.7789.2 (gEPNVSyIcSR)

   Ion         m/z   Intensity
   b1+       287.30       10.9
   a2+       388.36       15.1
   b2+       416.25       28.1
   y4+       535.42       10.5
   a5+       698.35       14.7
   b5+       726.63       18.9
   y5+       778.44       39.3
   a6+       785.77       22.5
   b6+       813.46       16.2
   y6+       865.30       22.4
   y7+       964.56       15.2
   b7+      1056.44       14.9
   b8+      1169.55       17.9
   y9+      1175.44       44.8
   b9+      1329.51       11.4
```

```
550. MORG-339_phosphopeptide_TMT_10%ACN_032017_OT.7151.7151.3 (sLDsDEsEDEDDDYQQkRk)

   Ion         m/z   Intensity
   b3+       545.36       20.7
   y6++      654.90       21.0
   y7++      712.67       25.1
   b10++     748.57       10.9
   y8++      769.79       14.8
   y9++      827.41       15.3
   y10++     892.12       23.7
   y11++     949.79       19.1
   b6+       956.56       13.9
   b14++    1002.80       13.6
   y12++    1014.24       25.6
   y13++    1097.74       11.5
   y14++    1161.85       14.2
   y16++    1303.34       24.1
   b9+      1367.65       11.2
   b12+     1726.67        9.7
```

```
551. MORG-339_phosphopeptide_TMT_16%ACN_032017_OT.9047.9047.3 (aPTAALsPEPQDSk)

   Ion         m/z   Intensity
   a1+       273.24        7.4
   y1+       376.37       11.3
   y5++      402.31       50.7
   b7++      460.98        7.1
   y2+       463.42       13.5
   y7++      515.32       34.3
   a9++      560.31        7.3
   b4+       570.46       10.6
   b9++      574.15       12.0
   y3+       578.21       19.6
   y8++      598.68       21.7
   b10++     622.93       11.5
   b5+       641.12       26.8
   b11++     687.18       11.4
   y10++     690.69        6.9
   y4+       706.30        7.1
   a12++     730.23        7.2
   b12++     744.03       13.7
   b6+       754.41       21.6
   b13++     787.83       12.4
   y5+       803.46       21.0
   y13++     825.56       10.4
   b7+       921.48       12.0
   y6+       932.33        8.6
   y7+      1029.49        7.5
   b9+      1147.55       38.1
```

```
552. MORG-339_phosphopeptide_TMT_10%ACN_032017_OT.8213.8213.2 (tAFYNEDDsEEEQR)

   Ion         m/z   Intensity
   y4+       561.39        9.8
   y5+       690.36       11.9
   b5+       826.52       11.8
   y6+       857.30       17.1
   a6+       926.99       13.7
   b6+       955.53       16.1
   y7+       972.57      117.9
   b7+      1070.50       24.7
   y8+      1087.54        9.9
   b8+      1185.59       22.2
   b12+     1739.73        9.8
```

```
553. MORG-339_phosphopeptide_TMT_6%ACN_032017_OT.5084.5084.3 (sYIETETESRER)

   Ion         m/z   Intensity
   y1+       175.11        8.8
   y3++      230.48        7.0
   y4++      274.16       15.0
   y2+       304.17        7.3
   y6++      389.34       26.4
   b1+       397.15       12.6
   y8++      504.40       35.2
   y4+       547.33        8.2
   b2+       560.43        9.6
   b7++      567.64       11.8
   y9++      568.77       18.1
   y10++     625.47       12.8
   a3+       645.28        6.6
   b3+       673.37       18.5
   y5+       676.47        7.3
   y11++     707.04       31.0
   b10++     753.63       10.7
   y6+       777.24       22.4
   b4+       802.38       22.9
   a11++     803.51       15.3
   b11++     818.06       30.2
   b6+      1032.37       13.0
   b8+      1262.55        9.1
```

```
554. MORG-339_phosphopeptide_TMT_14%ACN_032017_OT.7616.7616.4 (lEQHSQQPQLSPATsGRGLRDsPSsSEDR)

   Ion         m/z   Intensity
   b5++      412.74       11.4
   b2+       472.34       12.5
   b6++      476.79       11.4
   a7++      527.22       11.6
   y13+++    541.02       30.0
   b9++      653.73       13.5
   b10++     709.71       12.0
   b4+       737.52       13.6
   b11++     753.63       12.1
   a5+       796.79       12.2
   b12++     801.45       15.0
   y21+++    815.10       16.4
   a20+++    818.62       11.7
   b5+       824.31       14.4
   b20+++    827.50       11.7
   b13++     837.02       30.8
   y14++     839.63       34.9
   y22+++    847.78       22.8
   y7+       857.17       91.6
   b21+++    866.46       28.1
   a14++     873.70       18.4
   b14++     887.56       31.0
   y23+++    890.51       27.4
   y15++     923.43       12.3
   a6+       924.37       12.6
   a23+++    945.00       11.9
   b6+       952.57       12.6
   y25+++    962.19       12.6
   y16++     973.67       18.9
   a16++     985.72       11.9
   b16++    1000.25       13.2
   y17++    1009.20       16.4
   y27+++   1050.38       12.3
   a7+      1052.68       12.5
   y18++    1057.63       13.8
   a26+++   1058.49       15.3
   b7+      1080.60       19.0
   a18++    1092.51       14.4
   b27+++   1111.15       16.6
   y21++    1221.84       27.2
   y22++    1271.01       32.7
   a9+      1277.56       14.6
```

```
555. MORG-339_phosphopeptide_TMT_30%ACN_032017_OT.17960.17960.4 (fALLQVDsGsGsDSEPGkGkGR)

   Ion         m/z   Intensity
   a4++      323.83       12.1
   y7+++     386.75       35.0
   a5++      387.59       11.4
   b9+++     414.52       10.4
   y9+++     458.87       32.6
   y10+++    497.02       18.0
   b3+       561.22       11.5
   y12+++    571.65       17.3
   y7++      579.38       19.6
   b9++      620.32       12.3
   a4+       646.43       20.6
   b4+       674.14       12.3
   y15+++    702.31       22.1
   b10++     704.64       11.8
   y16+++    740.09       12.1
   y10++     744.99       15.0
   b5+       802.39       19.9
   y11++     828.58       11.7
   b19+++    844.51       16.2
   y19+++    853.50       11.9
   y12++     856.78       10.8
   a13++     860.28       11.8
   b6+       901.51       11.2
   a20+++    953.94       15.7
   y5+      1003.68       12.9
   b7+      1016.51       17.0
```

```
556. MORG-339_phosphopeptide_TMT_25%ACN_032017_OT.15149.15149.4 (sLNPALDGLTcGLTSHDkR)

   Ion         m/z   Intensity
   y2++      266.71       14.7
   b4++      361.30       36.8
   a5++      382.91       20.9
   b1+       396.93       42.5
   b6++      453.34       34.0
   y10+++    468.84       22.2
   y6++      487.09       16.0
   y11+++    506.23       12.9
   b7++      510.73       16.9
   y12+++    525.58       23.0
   y2+       532.42       27.0
   b8++      539.24       23.0
   y7++      543.81       12.8
   y13+++    563.87       28.8
   y8++      571.97       36.2
   b14+++    574.90       13.0
   a9++      582.00       20.3
   b9++      595.56       29.0
   a3+       596.49       20.0
   y14+++    601.49       35.1
   b15+++    603.91       14.1
   b3+       624.45       73.1
   y15+++    625.33       35.6
   a10++     632.47       18.5
   a16+++    640.49       57.2
   y10++     702.59       83.3
   b4+       721.78       12.6
   y18+++    733.27       13.9
   b12++     754.57       12.6
   y11++     759.13       36.9
   a5+       764.37       13.0
   y12++     787.64       61.0
   b5+       792.45       45.4
   b18+++    807.36       15.5
   y13++     845.20       82.2
   y14++     901.71       72.4
   b6+       905.50       25.7
   b7+      1020.51       18.5
   b17++    1031.77       27.5
   b8+      1077.59       26.3
   b9+      1190.60       13.6
```

```
557. MORG-339_phosphopeptide_TMT_22%ACN_032017_OT.12759.12759.4 (dLQsPDFTAGFHSDkVEGk)

   Ion         m/z   Intensity
   a1+       317.15       16.8
   b7+++     371.13       13.5
   y8+++     453.34       18.8
   b6++      483.56       11.9
   y9+++     502.60       19.9
   y5++      509.96       27.4
   y10+++    521.57       17.5
   y11+++    545.26       26.0
   y3+       562.49       13.8
   y6++      567.82       17.4
   b13+++    571.44       11.9
   y12+++    578.77       24.8
   b3+       586.43       14.8
   b8++      607.38       13.7
   b14+++    610.34       12.8
   y7++      611.07       27.0
   y13+++    627.97       45.2
   a9++      629.09       15.1
   b9++      642.75       14.4
   y4+       661.17       25.1
   y14+++    666.13       14.9
   b10++     671.10       18.6
   y8++      679.49       57.7
   y15+++    698.67       71.2
   a4+       725.45       14.9
   b4+       753.15       21.6
   b16+++    762.40       23.1
   y10++     781.54       34.7
   a12++     799.50       22.5
   b17+++    805.45       13.8
   b12++     813.47       17.9
   a18+++    814.39       17.9
   y11++     817.03       35.6
   a13++     842.99       13.3
   b13++     856.83       11.8
   y12++     867.37       29.8
   b14++     914.56       13.7
   y13++     941.12       16.3
   b6+       965.51       28.4
```

```
558. MORG-339_phosphopeptide_TMT_8%ACN_032017_OT.5003.5003.3 (rQEsTETLVVDR)

   Ion         m/z   Intensity
   b2++      257.82        7.5
   a3++      308.49        8.0
   b3++      322.10        9.1
   y3+       389.24        6.7
   b5++      456.22        6.5
   y4+       488.16       12.7
   b2+       514.05       13.4
   b6++      520.86       19.4
   y9++      550.46       19.5
   a7++      557.16       28.6
   b7++      571.11       22.7
   y5+       601.35        9.5
   a3+       615.13        6.6
   b8++      628.14        9.7
   b3+       643.50        8.7
   b9++      677.38       10.4
   y11++     678.46        8.6
   y6+       702.56       13.3
   b10++     727.08       14.7
   y7+       831.58       10.9
   y8+       932.45       12.0
```

```
559. MORG-339_phosphopeptide_TMT_6%ACN_032017_OT.16259.16259.3 (eAPAGGSGDGLSsPsPLEYsPR)

   Ion         m/z   Intensity
   y2+       272.21       11.1
   b1+       359.27        8.5
   a8++      415.17       10.7
   b2+       430.32       16.3
   y3+       439.26       19.8
   y7++      471.29        7.5
   b10++     514.74       12.4
   y8++      554.33       20.7
   b11++     571.39       29.6
   b4+       598.39       22.1
   y4+       602.26       34.2
   y9++      603.40       42.8
   b12++     615.01       19.1
   b5+       655.26       13.4
   y10++     686.85       13.9
   b6+       712.38       12.3
   y11++     730.32       24.1
   y5+       731.31       22.5
   a14++     732.42       11.5
   b7+       799.49       11.5
   y13++     815.43       18.1
   a15++     816.65        8.9
   a8+       828.89       82.9
   b15++     830.00       13.6
   y6+       844.41        7.6
   b8+       856.51       10.2
   y15++     901.52       10.3
   b17++     935.56       13.9
   y7+       941.74       32.5
   a9+       943.47        8.4
   b9+       971.45       20.1
   y17++     973.42        9.8
   b18++     999.87       25.2
   a10+     1000.55       16.4
   y18++    1002.02       10.3
   b10+     1028.58       18.8
   a19++    1067.62       10.6
   b19++    1081.52       23.6
   y8+      1108.41        7.6
   b11+     1141.54       33.4
   y9+      1205.46       21.1
   b12+     1228.55       18.8
   y10+     1372.42       24.3
   b13+     1395.52       16.5
   y11+     1459.46       15.7
```

```
560. MORG-339_phosphopeptide_TMT_20%ACN_032017_OT.8064.8064.4 (aAsPPASASDLIEQQQkRGR)

   Ion         m/z   Intensity
   a3++      255.82       20.3
   b3++      269.75       13.5
   a1+       273.16       11.3
   b1+       301.34       10.5
   a4++      304.18       10.6
   b4++      318.33       13.0
   a5++      353.39       17.1
   b5++      367.35       12.6
   b2+       372.30       10.4
   y7+++     377.38       18.8
   a6++      388.34       18.1
   y10+++    495.74       35.1
   y11+++    534.29       12.9
   b3+       539.31       13.7
   y12+++    563.38       11.4
   y7++      565.58       15.1
   b10++     582.76       12.3
   y13+++    586.83       17.4
   y14+++    615.92       26.1
   a16+++    625.78       14.8
   y8++      629.75       28.2
   b16+++    635.79       24.3
   b4+       636.50       24.6
   y16+++    671.93       24.4
   y9++      686.67       23.4
   y17+++    704.43      117.3
   b5+       733.79       12.1
   y10++     743.14       28.2
   a17+++    744.79       10.8
   y4+       745.85       17.0
   b17+++    754.04       22.9
   y18+++    759.43       10.8
   y19+++    783.45       12.4
   y11++     800.75       29.9
   b6+       804.55       11.6
   b18+++    806.75       19.7
   y12++     844.21       42.4
   y5+       873.54       10.5
   y13++     879.77       31.3
   y14++     923.29       54.2
   y15++     958.81       32.9
   y16++    1007.35       27.6
   b10+     1164.60       11.4
```

```
561. MORG-339_phosphopeptide_TMT_18%ACN_032017_OT.4493.4493.3 (sLGGQQGsPk)

   Ion         m/z   Intensity
   a3++      230.13        8.2
   y2++      237.30       19.9
   b1+       317.18       11.8
   y3++      320.59        9.5
   b5++      336.76       13.4
   y4++      349.26       14.1
   y1+       376.37       14.6
   b6++      400.75       21.2
   y5++      413.15       17.2
   a7++      415.45        9.6
   b7++      429.47       17.1
   b2+       430.34        9.2
   a3+       458.93        8.1
   y2+       473.45       44.3
   y6++      477.41       15.5
   b3+       487.76       21.2
   b8++      512.52        6.6
   b9++      561.15        6.8
   y9++      591.23       11.1
   y3+       640.39        9.6
   a5+       644.68        9.5
   b5+       672.45       21.5
   y4+       697.41       16.9
   b6+       800.47       17.1
   b7+       857.49       10.9
   b8+      1024.51       11.5
```

```
562. MORG-339_phosphopeptide_TMT_20%ACN_032017_OT.11882.11882.2 (lLsPQQPALQR)

   Ion         m/z   Intensity
   y3+       416.28        9.0
   b2+       456.41       13.7
   y5+       584.33       26.2
   a3+       595.51       11.5
   y6+       712.71       16.2
   b4+       720.24       13.5
   b5+       848.49        9.9
   y8+       937.62       52.4
   b6+       976.59       25.6
   y9+      1104.66        8.9
   y10+     1217.44        9.5
   b10+     1385.86        8.7
```

```
563. MORG-339_phosphopeptide_TMT_8%ACN_032017_OT.6098.6098.3 (dHsPTPSVFNSDEER)

   Ion         m/z   Intensity
   y2+       304.19       14.3
   b3++      325.09       12.0
   b1+       345.17        8.2
   y6++      375.29       12.8
   a5++      410.41        9.8
   b5++      424.30       14.5
   y3+       433.29       16.0
   a6++      458.42        9.9
   b2+       482.28       18.0
   y8++      498.31        9.6
   b7++      516.32       18.9
   y9++      541.37       10.3
   y4+       547.95       16.6
   a8++      551.46       37.5
   b8++      565.76       25.8
   y10++     590.42       82.9
   a9++      625.34       12.0
   y5+       635.37       20.1
   b9++      639.43       31.2
   y11++     641.04       22.4
   b3+       649.14       23.7
   y12++     689.52       18.7
   b10++     696.34       18.0
   y6+       749.33       38.1
   b12++     797.31       19.4
   y14++     841.36       10.3
   a13++     847.40       23.6
   b13++     861.94       28.0
   y7+       896.44       49.2
   b14++     926.58       17.8
   y8+       995.54       26.6
   y9+      1082.52       18.8
   y10+     1179.51       48.5
```

```
564. MORG-339_phosphopeptide_TMT_20%ACN_032017_OT.8350.8350.4 (eELLENEDEQNsPPkkGk)

   Ion         m/z   Intensity
   y5+++     415.56       16.2
   y6+++     448.12      105.4
   b7++      543.98       17.2
   b3+       601.41       14.4
   y10+++    627.38       17.0
   b9++      665.89       22.5
   y6++      671.65       37.1
   y12+++    709.06       16.2
   b4+       714.51       27.9
   a10++     715.54       15.7
   b10++     729.64       16.8
   y13+++    746.74      187.1
   a15+++    754.98       42.9
   y3+       790.19       15.2
   y15+++    827.12       17.5
   b5+       843.36       27.8
   a12++     856.76       16.3
   y16+++    865.29       16.6
   y9++      876.33       20.2
   a17+++    892.77       15.2
   a13++     904.79       15.5
   b13++     918.52       13.8
   a14++     953.18       18.0
   b6+       957.56       15.8
   b7+      1086.59       16.7
   b8+      1201.62       32.0
   b9+      1330.68       14.4
```

```
565. MORG-339_phosphopeptide_TMT_20%ACN_032017_OT.8722.8722.3 (vQtTPPPTIQGQk)

   Ion         m/z   Intensity
   b1+       329.28       10.4
   y4++      345.15       20.0
   y1+       376.40       27.5
   y5++      401.77       17.2
   b2+       457.36       12.8
   y2+       504.33       21.1
   y8++      549.66       12.6
   a8++      551.85       10.3
   y3+       561.43       44.9
   b8++      566.57       21.5
   y9++      597.77       45.7
   a3+       610.13       11.4
   b9++      623.12       44.3
   b3+       637.96       22.1
   a10++     672.93       14.3
   b10++     687.09       40.7
   y4+       689.46       39.4
   a4+       711.58       17.0
   b11++     715.35       25.0
   b4+       739.36       24.6
   b12++     779.73       23.5
   y12++     802.72       16.8
   b5+       836.79       11.4
   b8+      1131.63       17.5
   b9+      1244.78       17.4
```

```
566. MORG-339_phosphopeptide_TMT_14%ACN_032017_OT.9955.9955.3 (kAGPLGGssYEEEEEEEGGGGGER)

   Ion         m/z   Intensity
   y7+       589.52       13.6
   b11++     718.31       10.8
   b6+       753.18        7.6
   b7+       810.91       11.9
   a13++     833.64       10.3
   b13++     847.49        7.9
   a14++     897.92       13.9
   a8+       949.59       13.2
   a15++     963.01       11.1
   y10+      976.30        9.3
   b15++     977.10       13.7
   b8+       977.78       82.9
   y17++     995.39       20.1
   a16++    1027.65       11.0
   b17++    1105.57       11.8
   b18++    1134.14        7.4
   a22++    1234.52       10.4
   y13+     1363.67        8.2
```

```
567. MORG-339_phosphopeptide_TMT_20%ACN_032017_OT.8259.8259.3 (sPVGksPPATGSAYGSSQk)

   Ion         m/z   Intensity
   b1+       397.15       13.9
   a5++      490.00       18.6
   y2+       504.26       14.4
   b3+       593.30       12.2
   y4+       678.23       10.3
   b9++      720.25       10.6
   y5+       735.50       11.8
   y13++     740.66       47.0
   a12++     829.14       15.1
   b12++     842.68       15.3
   a13++     864.12       24.0
   y6+       898.59       13.7
   b14++     960.20       15.8
   y7+       969.67       12.7
   a15++     974.12       13.4
   a5+       979.63       13.1
   b15++     988.62       11.8
   y16++    1031.23       22.6
   y8+      1056.75       20.0
   b17++    1075.68       10.0
   y17++    1080.45       26.0
   y9+      1113.58       13.7
   y18++    1129.38       31.0
   y10+     1214.67       10.0
   y13+     1479.84       24.6
```

```
568. MORG-339_phosphopeptide_TMT_8%ACN_032017_OT.3714.3714.3 (hDtPDLsPPRR)

   Ion         m/z   Intensity
   y3++      214.58        6.3
   y4++      263.18       24.8
   y5++      346.75        8.5
   b4++      380.48        5.9
   y6++      403.27       17.2
   y3+       428.24        9.7
   b5++      438.53        8.4
   a2+       454.52        5.8
   y7++      460.42       26.9
   a6++      480.62       10.7
   b2+       482.35       14.4
   b6++      494.61       14.2
   y8++      509.34       33.7
   y4+       525.28       14.7
   b7++      578.34        5.9
   y9++      599.76        7.9
   b3+       663.25       10.4
   a10++     739.64        7.4
   b10++     753.46        9.5
   b5+       875.36       23.5
   b6+       988.38        7.2
   b7+      1155.48        8.4
```

```
569. MORG-339_phosphopeptide_TMT_30%ACN_032017_OT.15515.15515.3 (ssLVTSkLAGGQVE)

   Ion         m/z   Intensity
   y2+       247.14       21.1
   b3++      299.33       10.4
   b1+       317.23       13.6
   y3+       375.28       14.1
   y4+       432.27       19.7
   y5+       489.23       32.3
   y8++      515.74       11.5
   y9++      559.56       21.6
   y6+       560.38       22.5
   y10++     609.96       40.8
   b7++      621.24       44.0
   y11++     659.32       15.9
   a8++      664.33       16.0
   y7+       673.47       18.3
   b8++      677.98       46.4
   b4+       696.41       26.4
   b9++      713.55       77.7
   b10++     741.94       41.6
   b11++     770.56       29.2
   b5+       797.53       15.4
   b12++     834.57       75.6
   b6+       884.09       54.3
   y8+      1030.27       19.5
```

```
570. MORG-339_phosphopeptide_TMT_4%ACN_032017_OT.6054.6054.3 (tLTDEVNsPDsDRR)

   Ion         m/z   Intensity
   y2+       331.31       16.8
   y5++      364.28       29.8
   y6++      413.30       20.0
   a2+       416.20        8.4
   a7++      488.19       17.6
   y7++      496.73       14.3
   b3+       545.24       10.5
   y8++      553.69       16.6
   b8++      585.35       11.2
   y9++      603.09       13.7
   a9++      619.54        7.8
   a4+       632.75      179.1
   b9++      633.53       45.3
   b4+       660.50       18.4
   b11++     774.70        9.1
   b5+       789.48       24.4
   y6+       825.40        8.9
   y13++     832.55        8.7
   b6+       888.62       19.0
   b13++     910.49       18.4
   b7+      1002.58       19.2
   b10+     1381.60        8.8
   y13+     1663.96        8.3
```

```
571. MORG-339_phosphopeptide_TMT_12%ACN_032017_OT.18814.18814.2 (lTNLSsNsDV)

   Ion         m/z   Intensity
   b1+       343.40        8.4
   a2+       416.39        4.3
   b2+       444.34       10.0
   y4+       514.20        8.6
   b3+       558.36       13.2
   a4+       643.19        5.9
   b4+       670.98      100.8
   y5+       680.69       14.8
   y6+       768.19       21.6
   y7+       881.37        9.7
   a6+       897.50        7.9
   b6+       925.50        7.9
   y8+       995.32        7.3
   b7+      1039.47       13.5
   y9+      1096.33       11.6
   b9+      1321.54       11.2
```

```
572. MORG-339_phosphopeptide_TMT_20%ACN_032017_OT.6308.6308.4 (aGDVLEDsPkRPk)

   Ion         m/z   Intensity
   a1+       273.30       12.3
   y3++      315.30       14.8
   y4+++     329.65       17.4
   y5+++     362.14       68.2
   y1+       376.28        9.1
   y6+++     417.81       50.0
   y7+++     456.07       29.2
   b7++      465.07       14.7
   y2+       473.41       44.0
   y8+++     498.99        8.3
   a10+++    508.06        9.9
   a8++      534.99       15.9
   y9+++     537.13       57.4
   a11+++    560.56       12.4
   y10+++    569.85       65.9
   b4+       572.49       17.6
   b12+++    602.46       12.4
   y11+++    608.44       11.2
   y6++      625.72        9.0
   y12+++    627.03       11.3
   b5+       685.43       13.3
   b10++     775.76       17.2
   b6+       814.55       26.8
   y11++     911.53        8.4
   b7+       929.50       36.8
```

```
573. MORG-339_phosphopeptide_TMT_25%ACN_032017_OT.12137.12137.3 (dVINRsPTQLQLGk)

   Ion         m/z   Intensity
   a1+       317.20       20.2
   y2+       433.32       17.3
   b6++      497.84       14.0
   y3+       546.50       20.2
   b3+       557.43       17.0
   y9++      641.16       20.1
   a4+       643.23       20.1
   a9++      647.20       16.9
   b9++      661.00       26.0
   y4+       674.52       41.5
   b10++     717.45       19.0
   a11++     767.75       16.3
   y11++     775.82       16.4
   b11++     781.63       24.6
   y5+       787.58       21.7
   a5+       799.78       15.0
   a12++     824.06       14.8
   b5+       827.40       15.7
   y12++     832.68       18.6
   y13++     881.71       15.0
   y8+      1113.64       17.8
```

```
574. MORG-339_phosphopeptide_TMT_18%ACN_032017_OT.12134.12134.3 (skDVEFQDGWPR)

   Ion         m/z   Intensity
   y2+       272.17        7.7
   b2++      377.76        8.1
   b1+       397.31        6.8
   b3++      435.27       19.1
   b4++      484.93       15.3
   y4+       515.37       26.1
   b5++      549.44       14.4
   a6++      609.18        9.3
   b6++      623.00       13.4
   y5+       630.38       13.1
   b7++      686.94        9.8
   b8++      744.41       27.5
   b2+       754.74        7.7
   a9++      758.42        8.3
   b9++      773.00       11.0
   y11++     803.26       19.0
   a10++     852.25       14.4
   b10++     865.78       15.6
   y7+       905.34       15.3
   y8+      1034.45       10.8
   y9+      1133.73       19.5
```

```
575. MORG-339_phosphopeptide_TMT_8%ACN_032017_OT.3830.3830.2 (sPIVNER)

   Ion         m/z   Intensity
   y1+       175.15        9.3
   y2+       304.07        7.5
   b1+       397.24       31.4
   y3+       418.26       10.5
   y4+       517.23       12.8
   a3+       579.64        8.1
   b3+       607.42        8.8
   b4+       706.38       19.2
   y6+       727.44       45.6
   b6+       949.45       17.3
```

```
576. MORG-339_phosphopeptide_TMT_16%ACN_032017_OT.9792.9792.3 (sEPAVGPLRGLGDQSSSR)

   Ion         m/z   Intensity
   b5++      397.18       18.1
   y4+       436.14       10.9
   a2+       498.27       11.2
   b2+       526.29       18.3
   y5+       564.28       53.6
   a3+       595.48       10.7
   b9++      608.96       25.9
   b10++     637.03       18.1
   y13++     665.57       37.4
   a4+       666.65       12.8
   b4+       694.10       17.7
   y14++     715.09       10.4
   b12++     722.46       10.8
   y7+       736.42       19.8
   b13++     779.97       59.9
   b5+       793.41       25.4
   y16++     799.10       75.7
   b14++     844.00       12.0
   y8+       849.63       10.9
   b6+       850.47       18.7
   y17++     863.66       47.7
   y9+       906.49       29.5
   b7+       947.25       10.6
   y10+     1062.44       10.4
```

```
577. MORG-339_phosphopeptide_TMT_6%ACN_032017_OT.15281.15281.3 (eESEtRsESsDFEVVPk)

   Ion         m/z   Intensity
   y1+       376.24       12.7
   y2+       473.33       54.9
   y6++      474.34       26.8
   b2+       488.18        8.0
   y3+       572.42       34.6
   a8++      655.36        7.0
   y4+       671.52       27.7
   b4+       704.76        7.9
   b9++      712.99        8.3
   a10++     781.82        7.4
   y5+       800.51       15.4
   y11++     807.20        9.5
   b11++     853.64       31.4
   a12++     912.97        8.1
   b12++     927.54       21.3
   y6+       947.64       17.6
   y13++     974.96        8.5
   b13++     992.07       54.5
   a14++    1027.24       12.6
   b6+      1041.67       73.5
   a15++    1077.12       23.6
   b15++    1091.16       82.9
   y8+      1229.59        7.7
   y9+      1316.67       10.4
   b8+      1337.50        7.2
   y11+     1612.74        9.7
```

```
578. MORG-339_phosphopeptide_TMT_12%ACN_032017_OT.8041.8041.3 (kVAsVEEGDETLDSPGSQSR)

   Ion         m/z   Intensity
   b5++      397.28        8.6
   b3+       528.03        8.8
   y6+       631.32       12.2
   a10++     662.97       10.6
   y13++     674.77       12.9
   y7+       718.52       28.9
   b11++     727.69       18.4
   b12++     784.67       15.9
   b5+       794.05       14.9
   y15++     803.56        8.6
   a13++     828.02       16.5
   y8+       833.37       24.2
   b13++     841.40       30.2
   y16++     852.94       24.4
   y17++     937.10       12.5
   y18++     972.81       14.9
   b17++    1005.49        9.8
   y10+     1047.60       12.5
   y11+     1176.78       10.6
   y13+     1348.51       13.2
```

```
579. MORG-339_phosphopeptide_TMT_10%ACN_032017_OT.7796.7796.3 (eNPPSPPtsPAAPQPR)

   Ion         m/z   Intensity
   y7++      368.97        9.4
   b2+       473.29       11.4
   y4+       497.29       28.7
   a3+       542.17       11.9
   a8++      551.16        9.4
   b8++      565.28       11.1
   y5+       568.37       17.0
   b3+       570.64       10.7
   y10++     590.91       18.7
   y11++     639.58       34.2
   b9++      648.45       23.1
   b4+       666.89       31.3
   b10++     697.00        9.1
   a5+       726.73        9.7
   b11++     732.92       27.5
   y7+       736.51       22.0
   b12++     768.51       29.8
   y14++     780.21       13.5
   y15++     837.27        9.4
   b14++     880.76        9.3
   b8+      1129.57        9.2
```

```
580. MORG-339_phosphopeptide_TMT_10%ACN_032017_OT.10055.10055.3 (tSFDENDsEELEDkDSk)

   Ion         m/z   Intensity
   y2+       463.43       12.2
   y4++      468.48       21.8
   y7++      647.17       14.0
   b4+       680.35       17.6
   y8++      711.73       11.4
   y9++      776.12       16.0
   b5+       809.53       15.0
   b12++     853.23       16.2
   y10++     859.69       21.3
   a6+       895.87       29.4
   a13++     897.08       27.1
   y12++     973.69       11.5
   a7+      1010.46       12.0
   y13++    1038.56       22.3
   b14++    1089.83       18.5
   y14++    1096.27       13.7
   b15++    1147.26       23.4
   b16++    1190.24       10.2
```

```
581. MORG-339_phosphopeptide_TMT_20%ACN_032017_OT.12960.12960.3 (hssISPSTLTLk)

   Ion         m/z   Intensity
   y3++      295.72        8.3
   y1+       376.27       30.9
   y5++      402.39        8.4
   b5++      451.18        9.6
   a6++      485.24       12.0
   y2+       489.39       23.1
   y7++      495.19       24.6
   a7++      529.25        9.6
   b2+       534.49       17.6
   b7++      543.62       10.1
   a8++      579.46       12.2
   y3+       590.52       29.8
   b8++      594.11       12.5
   y9++      594.82        9.9
   a9++      636.60       12.8
   b9++      650.52       10.3
   a10++     687.14        8.3
   b3+       700.77       13.6
   y4+       703.52       18.3
   b11++     757.75       14.6
   y5+       804.43       22.7
   a6+       970.20        7.6
```

```
582. MORG-339_phosphopeptide_TMT_16%ACN_032017_OT.8721.8721.4 (kAGPLGGssYEEEEEEEGGGGGERk)

   Ion         m/z   Intensity
   b1+       358.30       11.1
   y8++      473.91       15.1
   y9++      538.46       14.2
   y11++     667.60       16.6
   y16+++    671.52       13.8
   y12++     732.03       14.9
   b17+++    737.56        9.4
   b19+++    775.40        9.2
   b12++     782.93        9.0
   y13++     796.66       23.4
   y19+++    802.25       10.5
   a21+++    804.81       10.0
   b21+++    813.99       22.0
   y20+++    821.19       18.7
   a22+++    822.99      146.3
   b22+++    832.76       12.9
   a13++     833.90       22.0
   y14++     860.76       14.3
   b23+++    875.68       11.7
   y23+++    910.18       12.8
   y15++     925.65       26.2
   b8+       977.99        9.8
   y16++    1006.84       19.2
   b16++    1041.60       13.8
   y17++    1090.66       13.8
   a17++    1091.63        9.0
   y20++    1231.09       12.9
   y21++    1287.18       15.0
```

```
583. MORG-339_phosphopeptide_TMT_22%ACN_032017_OT.8952.8952.3 (nVPGLAAPSsPTQkEGPR)

   Ion         m/z   Intensity
   b2+       443.27       19.8
   y8++      571.46       46.8
   y10++     697.96       24.6
   b5+       710.43       23.6
   y11++     747.06       34.7
   a13++     751.07       23.7
   a6+       753.43       39.1
   b13++     765.81       49.1
   y5+       815.13       18.1
   y13++     817.47       17.4
   b7+       852.45       26.2
   y14++     874.17       15.9
   y15++     902.63       29.5
   a8+       921.53       14.8
   b14++     944.20       17.8
   y16++     951.42       21.6
   a9+      1008.42       24.8
   b16++    1036.72       14.9
   y7+      1044.85       15.6
```

```
584. MORG-339_phosphopeptide_TMT_12%ACN_032017_OT.5128.5128.3 (sRsPDIQAk)

   Ion         m/z   Intensity
   a2++      263.42        7.7
   b2++      277.25        8.4
   a1+       368.91        6.1
   y1+       376.33        8.8
   y2+       447.27       10.7
   y6++      450.80       22.3
   a5++      452.41        7.6
   b5++      466.95       16.2
   a6++      509.57       27.2
   b6++      523.01       29.5
   a2+       524.79       11.6
   y7++      534.65       15.9
   y3+       575.61       16.6
   b7++      587.48       11.5
   b8++      622.39       14.8
   y4+       688.51       16.1
   b3+       720.42        8.8
   y6+       900.43       10.6
```

```
585. MORG-339_phosphopeptide_TMT_10%ACN_032017_OT.19106.19106.3 (iVEPEVVGEsDsEVEGDAWRLER)

   Ion         m/z   Intensity
   y6++      415.70       24.0
   y3+       417.34       10.7
   y7++      473.42       12.0
   y8++      501.96       25.4
   y9++      566.56       20.6
   b3+       571.43       18.9
   y10++     616.08       20.3
   y11++     680.57       18.8
   y5+       759.65        9.4
   y12++     764.06       27.1
   b5+       797.48       16.8
   y13++     821.49       11.3
   y6+       830.54       11.1
   b6+       896.57       23.9
   y14++     904.83       12.9
   y7+       945.82       11.1
   a7+       967.65       13.2
   y15++     969.70       11.6
   a15++     980.29       13.8
   b8+      1052.64       12.1
   y18++    1097.37       13.6
   y19++    1161.62       16.5
   b9+      1181.62       16.5
   y20++    1210.24       14.2
   b20++    1287.42       15.6
   a10+     1320.42        8.9
   y22++    1324.46       11.9
   y11+     1359.45       24.4
   b22++    1408.48       18.5
   a11+     1435.73        8.6
   b11+     1463.77       13.2
   b14+     1858.82        9.3
   b15+     1987.81        8.6
```

```
586. MORG-339_phosphopeptide_TMT_30%ACN_032017_OT.11699.11699.4 (kQPPVsPGTALVGSQkEPSEVPTPk)

   Ion         m/z   Intensity
   b6++      473.45       29.4
   a9++      587.57       24.8
   y4+       671.50       48.8
   b12++     742.67       19.9
   y18+++    761.91       17.7
   y5+       770.46       19.4
   y19+++    794.22       24.6
   b19+++    809.68       18.8
   y20+++    849.85       27.9
   a15++     865.40      180.4
   a21+++    876.14       39.6
   b15++     879.45       32.8
   a22+++    908.92       20.4
   y22+++    915.22       33.9
   a6+       918.65       21.8
   y13++     921.69       42.9
   y23+++    947.37       30.8
   b23+++    952.24       21.4
   y14++     971.24       35.6
   a24+++    974.68       19.2
   b24+++    983.93       17.1
   y7+       986.40       22.7
   y24+++    990.42       17.3
   y15++    1027.85       34.4
   y8+      1083.62       26.5
   b8+      1100.96       24.7
   y19++    1190.99       73.8
   y22++    1372.17       19.0
```

```
587. MORG-339_phosphopeptide_TMT_18%ACN_032017_OT.11495.11495.3 (rSsAIGIENIQEVQEk)

   Ion         m/z   Intensity
   y3++      317.57       11.9
   y1+       376.31       16.9
   a5++      398.90       13.1
   b5++      412.87       11.8
   b6++      441.12       12.7
   a7++      484.15       12.3
   y6++      495.42       13.0
   b7++      497.69       14.0
   y2+       505.39       20.9
   b8++      562.37       17.9
   b9++      619.54       20.2
   y3+       633.52       31.4
   a10++     662.23       11.0
   b10++     675.96       36.8
   b4+       711.63       11.1
   a11++     725.76       11.0
   y10++     730.33       13.0
   y4+       732.56       29.9
   b11++     740.10       35.9
   y11++     758.63       24.3
   b12++     804.56       35.4
   y12++     815.09       12.4
   b5+       824.54       16.9
   a13++     840.00       12.5
   y13++     850.82       17.8
   b13++     853.88       34.0
   y5+       861.48       24.4
   b6+       881.42       13.5
   b14++     918.09       28.9
   y14++     934.00       11.2
   a15++     968.78       12.5
   b15++     982.63       20.8
   y6+       989.57       28.5
   b7+       994.53       13.7
   y8+      1216.69       17.0
   y9+      1345.88       16.7
```

```
588. MORG-339_phosphopeptide_TMT_10%ACN_032017_OT.18215.18215.3 (tYEtPPPsPGLDPTFSNQPVPPDAVR)

   Ion         m/z   Intensity
   y3+       345.35       12.8
   b3+       623.32       14.7
   b8++      631.53       13.2
   y6+       654.41       75.0
   y7+       753.71       12.7
   y14++     763.21       43.6
   b4+       804.37       23.3
   y15++     820.69       34.6
   b12++     822.99       16.3
   y8+       850.53       38.8
   b13++     871.44       22.3
   b5+       901.57       11.7
   y18++     953.88       21.0
   a6+       970.39       16.1
   y9+       978.52       16.2
   a15++     981.76       15.6
   b15++     995.52       11.9
   a16++    1024.59       71.7
   b16++    1039.39       22.2
   y10+     1092.52       15.6
   y21++    1134.33       18.5
   b18++    1160.30       54.2
   y11+     1179.59       25.6
   y22++    1183.33       34.8
   b20++    1258.32       47.9
   b8+      1262.69       13.4
   y23++    1273.94       13.8
   y12+     1326.81       12.6
   b23++    1412.87       12.2
   b24++    1448.44       11.0
   y14+     1524.89       16.6
   b11+     1529.67       29.6
   y15+     1639.85       15.8
   b12+     1644.71       36.5
   a13+     1713.83       12.6
   b15+     1989.81       12.0
```

```
589. MORG-339_phosphopeptide_TMT_4%ACN_032017_OT.4791.4791.4 (eRERDHsPTPSVFNsDEER)

   Ion         m/z   Intensity
   y2+       304.19        7.5
   b7+++     407.36        8.0
   y6++      415.20        8.6
   y3+       433.17        8.3
   b8+++     439.93       14.1
   b5++      458.45       17.0
   b9+++     473.31        8.5
   y7++      488.97       14.5
   a10+++    496.05        7.5
   b6++      527.04       11.6
   b11+++    534.91       15.3
   y8++      538.54       11.0
   y4+       548.24       11.6
   a12+++    558.53       13.6
   b12+++    567.88       20.4
   y9++      581.61        9.2
   a7++      596.72        7.9
   a13+++    607.73        8.0
   b7++      610.69       22.2
   b13+++    616.47       24.4
   y15+++    626.26        9.3
   y10++     630.46       22.3
   a8++      644.67       19.0
   a14+++    645.57       76.1
   b14+++    654.76       19.9
   b8++      659.02       16.2
   y16+++    677.87        8.6
   y11++     680.83        7.6
   a15+++    700.55       23.0
   b9++      709.73       10.3
   b15+++    710.65        8.2
   y5+       715.21       12.4
   y12++     729.51       20.0
   b17+++    792.07       17.7
   y6+       829.24       18.1
   b12++     851.18       20.7
   y14++     881.65       16.0
   b5+       915.66       11.0
   b13++     924.79       15.5
   y7+       976.42       18.5
```

```
590. MORG-339_phosphopeptide_TMT_22%ACN_032017_OT.15102.15102.3 (sPALksPLQSVVVR)

   Ion         m/z   Intensity
   b1+       397.20       20.6
   a5++      503.90       14.2
   b5++      518.39       13.4
   a3+       537.51        9.0
   y5+       559.43       13.7
   b3+       565.32        8.8
   a7++      636.38       12.8
   a4+       650.53       10.2
   b4+       678.22       22.0
   y6+       687.65       21.7
   a8++      693.08       17.0
   y10++     711.76        8.3
   a9++      756.99       10.4
   y11++     768.18       14.0
   b9++      771.14       14.0
   y12++     803.46       27.7
   y13++     852.38       40.4
   y8+       897.60       30.1
   b12++     913.82       15.3
   b13++     963.33       12.1
   y9+      1064.58       12.8
```

```
591. MORG-339_phosphopeptide_TMT_4%ACN_032017_OT.7491.7491.2 (aGGAGGEGsDDDTSLT)

   Ion         m/z   Intensity
   b1+       301.26        9.1
   y3+       320.23       12.3
   b3+       415.18        9.1
   b4+       486.39       15.3
   b5+       543.39        8.2
   b6+       600.39        9.2
   y6+       651.20       12.4
   a7+       701.40       12.5
   b7+       729.47       19.2
   b8+       786.38       18.0
   b10+     1068.46       14.5
   y11+     1176.53       10.0
   b11+     1183.47       14.6
   y12+     1233.38       12.5
   b12+     1298.52       23.4
   y13+     1304.47        7.1
   b13+     1399.61       38.5
   y15+     1418.49       11.4
   b14+     1486.64       27.7
```

```
592. MORG-339_phosphopeptide_TMT_18%ACN_032017_OT.8784.8784.3 (tPGtPGtPSYPRTPGTPk)

   Ion         m/z   Intensity
   b1+       331.22       22.8
   y2+       473.34       32.7
   y8++      542.12       60.2
   y9++      623.68       30.3
   b4+       666.03       21.4
   y10++     667.13       21.7
   y11++     715.68      101.4
   b11++     723.13       20.7
   y5+       728.59       25.4
   b5+       763.77       32.6
   a12++     787.23       41.4
   a6+       792.38       31.3
   b12++     801.22       38.2
   b6+       820.07       17.4
   y6+       829.63       16.9
   y13++     834.56       38.0
   y14++     883.29       79.0
   a14++     886.40       20.1
   a15++     915.11       22.5
   b15++     929.16       19.7
   y15++     973.77       16.8
   b7+      1001.28       22.6
   y16++    1002.06       27.0
   y17++    1050.96       33.2
```

```
593. MORG-339_phosphopeptide_TMT_14%ACN_032017_OT.14387.14387.3 (tDGsIsGDRQPVTVADYISR)

   Ion         m/z   Intensity
   b2+       446.47       10.3
   y4+       538.36       16.6
   b9++      639.77        9.3
   y12++     703.07       28.2
   b10++     703.92       26.7
   y6+       724.50       11.9
   b11++     752.73        9.3
   a5+       755.32       10.5
   y13++     760.60       12.0
   b5+       783.68       26.2
   y14++     789.11       26.1
   b12++     801.87       12.0
   y7+       823.53       18.9
   b13++     852.29       12.8
   y15++     872.66       16.7
   b14++     902.29       10.0
   y8+       924.48       12.2
   y16++     929.13       12.1
   b16++     995.18       17.5
   y18++    1041.28       10.7
   y10+     1120.58       22.3
   b8+      1122.42       14.2
   y11+     1248.73       10.3
```

```
594. MORG-339_phosphopeptide_TMT_8%ACN_032017_OT.8079.8079.4 (eVENEQTPVsEPEEEkGSQPGPPVER)

   Ion         m/z   Intensity
   b4++      351.27       12.9
   a8+++     376.38       44.6
   y8++      440.21       12.4
   b2+       458.26       16.4
   b6++      480.13       13.3
   y4+       500.37       13.4
   b7++      530.38       43.7
   b3+       587.37       26.7
   y5+       597.46       17.0
   y15+++    622.41       36.5
   b9++      628.23       14.5
   y6+       654.39       22.0
   y11++     691.04       17.0
   b4+       701.48       26.2
   y7+       751.41       55.3
   y18+++    753.84       39.2
   y12++     755.67       14.4
   y19+++    786.61      116.3
   a16+++    789.03       13.3
   a17+++    807.92       14.2
   a12++     811.27       12.0
   b17+++    817.42       11.4
   y20+++    820.34       57.6
   b12++     824.44       16.7
   b5+       830.56       76.5
   a18+++    837.33       27.1
   b18+++    846.23       50.7
   a19+++    879.47       12.2
   b13++     889.58       13.9
   b20+++    921.62       13.3
   y15++     933.25       37.1
   b21+++    940.63       15.5
   b6+       958.42       33.5
   b22+++    972.51       12.7
   y16++     997.58       17.3
   y10+     1023.49       22.0
   b7+      1059.52       64.4
   b25+++   1081.22       32.0
   y18++    1130.45       39.5
   y19++    1179.26       83.6
   y20++    1229.95       30.3
   b9+      1255.60       17.2
   b18++    1269.48       16.5
   b19++    1333.44       67.8
   b21++    1410.63       14.7
   b11+     1551.68       17.9
```

```
595. MORG-339_phosphopeptide_TMT_4%ACN_032017_OT.3891.3891.2 (sEEEQSSASVk)

   Ion         m/z   Intensity
   y1+       376.31        8.3
   b1+       397.16        5.1
   y2+       475.43        8.1
   y3+       562.39        9.2
   y4+       633.53        5.4
   b3+       655.41        6.0
   y5+       720.53        6.7
   b4+       784.31        8.9
   y6+       807.38       14.9
   b5+       912.46        6.7
   y7+       935.55        9.9
   b6+       999.56        4.4
   y8+      1064.60        9.8
   a8+      1129.37        5.0
   y9+      1193.73        6.6
   y10+     1322.76        7.8
   b10+     1343.49        5.8
```

```
596. MORG-339_phosphopeptide_TMT_6%ACN_032017_OT.6688.6688.3 (eSLkEEDEsDDDNm)

   Ion         m/z   Intensity
   b3++      279.88        8.2
   b1+       359.26        7.6
   y3+       395.23        5.8
   b4++      459.10       15.8
   y4+       510.20        8.0
   b5++      523.27       18.3
   b6++      588.07       25.2
   b7++      645.69       20.0
   b8++      710.20       37.3
   y6+       792.31        7.7
   b9++      793.29        7.9
   b10++     851.15       27.2
   y12++     882.93        6.9
   b11++     908.63       23.9
   y7+       921.54        6.3
   y13++     926.68        9.1
   b12++     966.17       39.0
   b13++    1022.80       28.7
   y10+     1294.34        6.0
```

```
597. MORG-339_phosphopeptide_TMT_12%ACN_032017_OT.5317.5317.3 (sPSPSPTPEAk)

   Ion         m/z   Intensity
   b2++      247.35        6.6
   y4++      337.29       40.4
   y1+       376.34       10.9
   b5++      383.22        9.0
   b1+       397.35       12.8
   b6++      431.64        7.9
   y6++      436.24       20.5
   y2+       447.33       13.4
   a2+       466.13        7.7
   a7++      468.40       10.0
   y7++      480.02       17.9
   b7++      482.30       25.1
   b2+       494.68       10.4
   a8++      516.93       19.1
   y8++      528.44       31.1
   b8++      530.97       17.2
   a3+       553.55        6.4
   y3+       576.41       14.4
   a9++      581.23       14.1
   b9++      595.19       10.7
   b10++     630.71       13.0
   y4+       673.49       20.8
   a5+       737.41        7.0
   b5+       765.37       29.0
   y5+       774.47        8.2
   b6+       862.25        8.8
   b7+       963.45       28.5
```

```
598. MORG-339_phosphopeptide_TMT_16%ACN_032017_OT.3345.3345.3 (sGkNSQEDsEDsEEk)

   Ion         m/z   Intensity
   b6++      530.99        6.9
   b7++      595.30       11.5
   y7++      606.66        6.3
   y3+       634.45        8.0
   b8++      652.98       23.8
   a3+       703.34        7.2
   b3+       731.85        8.8
   a10++     787.22       19.3
   y4+       800.97       46.1
   a4+       817.70       16.5
   b4+       845.62        7.2
   b11++     858.19       14.7
   y12++     893.08       10.6
   b14++    1071.09        6.3
   y13++    1072.24        6.5
   y7+      1212.47        9.1
```

```
599. MORG-339_phosphopeptide_TMT_10%ACN_032017_OT.17760.17760.3 (sLEETLHNVDLssDDELPR)

   Ion         m/z   Intensity
   y3+       385.27       12.4
   y4+       514.37       22.9
   b7++      520.61       18.6
   a8++      563.38       10.0
   b8++      577.49       28.7
   a9++      613.21       13.6
   b9++      627.07       37.1
   y5+       629.34       21.0
   b10++     684.76       47.7
   b4+       688.47       12.0
   a11++     727.08       16.6
   b11++     741.22       34.1
   b5+       789.23       24.0
   a12++     810.30       13.3
   y13++     828.41       24.5
   b6+       902.52       18.9
   y7+       911.40       11.4
   b14++     965.71       22.8
   a7+      1011.53       12.3
   b15++    1023.17       20.8
   b7+      1039.28       18.6
   y8+      1078.29       17.3
   b16++    1087.77       19.3
   a17++    1130.31       11.4
   b17++    1143.96       18.0
   y9+      1191.49       15.1
   b18++    1192.71       10.4
   y10+     1306.48       14.9
   y12+     1519.53       10.4
```

```
600. MORG-339_phosphopeptide_TMT_14%ACN_032017_OT.5567.5567.3 (aDLNQGIGEPQsPSRR)

   Ion         m/z   Intensity
   y6++      405.35       17.8
   b2+       416.30        9.7
   y7++      454.32       17.6
   y4+       515.37       11.7
   y9++      547.33       11.6
   y10++     603.61       15.6
   b10++     612.43       18.9
   y11++     632.67       13.6
   b4+       643.39       16.1
   a11++     663.10       25.9
   a12++     746.72        9.8
   y13++     752.88       10.1
   b12++     760.48       15.8
   b5+       771.23       13.2
   b13++     808.58       14.3
   y14++     809.59       20.9
   y6+       810.54       15.0
   b6+       828.43       16.6
   y7+       907.51       14.1
   b15++     930.33       19.7
   b7+       941.55       12.0
   b9+      1127.60       24.8
```

```
601. MORG-339_phosphopeptide_TMT_16%ACN_032017_OT.9646.9646.4 (rLsGAQAELALQEESVRR)

   Ion         m/z   Intensity
   a2++      236.23       14.7
   y4++      259.23       18.9
   y5++      323.63       12.2
   b3++      333.92       20.7
   a1+       357.84       10.6
   b4++      362.30       16.0
   a5++      383.84       23.3
   b5++      397.63       30.7
   b9+++     413.04       14.0
   y11+++    443.74       14.7
   a6++      448.01       12.5
   y7++      452.53       12.7
   b6++      461.81       38.6
   b11+++    473.81       17.5
   b7++      497.50       17.8
   y4+       516.97       11.5
   y9++      544.39       11.2
   a13+++    550.12       15.6
   y15+++    552.81       78.9
   b13+++    559.59       21.2
   b8++      561.58       84.8
   y10++     601.13       38.3
   y16+++    608.51       16.1
   b9++      618.27       20.4
   y5+       646.53       11.2
   b10++     653.69       21.8
   y11++     665.58       30.6
   b3+       666.45       18.3
   y12++     701.16       57.2
   b11++     710.26       13.1
   b17+++    716.88       45.0
   b4+       723.01       13.4
   y13++     765.20       63.8
   y14++     800.83       18.3
   y15++     829.24       58.9
   b13++     839.00       19.1
   a14++     889.35       16.5
   y7+       903.62       24.1
   b15++     947.23       12.3
```

```
602. MORG-339_phosphopeptide_TMT_22%ACN_032017_OT.12597.12597.3 (kGSPVSEIGWETPPPEsPR)

   Ion         m/z   Intensity
   y5+       665.33       19.4
   a12++     736.84       20.3
   b12++     750.59       31.5
   y6+       762.40       27.9
   b6+       785.50       14.4
   y14++     831.31       68.6
   a14++     834.35       43.6
   b14++     848.07       26.2
   a15++     882.20       13.7
   y16++     929.50       13.7
   y8+       960.21       13.5
   b16++     961.42       14.3
```

```
603. MORG-339_phosphopeptide_TMT_20%ACN_032017_OT.13439.13439.4 (aLsLDNSSQVSQkGSDGDLTSGDLk)

   Ion         m/z   Intensity
   y1+       376.28       22.1
   y2+       489.42       15.4
   y3+       604.40       14.7
   b13+++    633.48       12.8
   y4+       661.55       24.0
   b10++     662.83       30.8
   b15+++    681.14       11.2
   y16+++    689.23       12.6
   y12++     697.49        9.7
   b11++     705.88       13.8
   a17+++    728.76       10.1
   b17+++    738.02       10.4
   y5+       748.51       23.8
   a12++     756.58       16.5
   y18+++    760.81        9.7
   b18+++    776.64       12.4
   a5+       781.76        9.9
   y19+++    789.70       16.7
   a19+++    805.57       36.8
   b19+++    814.33       37.9
   y6+       849.60       20.6
   y21+++    865.70       24.1
   y13++     876.10       11.1
   b22+++    896.37       19.5
   y22+++    903.80       11.6
   b6+       923.25       13.0
   a13++     934.99       28.5
   y14++     940.29       14.6
   b13++     949.32       30.5
   a24+++    962.39       13.6
   a14++     963.76       12.2
   b24+++    972.57       45.5
   y15++     983.91       13.3
   b15++    1021.16       18.7
   b16++    1078.73       16.8
   b18++    1164.86       18.9
   b19++    1220.97       16.2
   b10+     1324.70       11.5
```

```
604. MORG-339_phosphopeptide_TMT_22%ACN_032017_OT.11595.11595.2 (sLsPLSGTTDTk)

   Ion         m/z   Intensity
   y2+       477.36       21.6
   a5+       779.59       12.6
   y5+       794.49       11.6
   b5+       807.21       18.5
   y6+       851.56       39.5
   b6+       894.78       12.5
   a7+       923.05       15.5
   y7+       938.50       18.8
   b7+       950.98       13.0
   y8+      1051.26       12.0
   y9+      1148.67       37.3
   b10+     1268.63       16.1
```

```
605. MORG-339_phosphopeptide_TMT_25%ACN_032017_OT.14465.14465.3 (sGsPGRVLTTTALSTVSSGAQR)

   Ion         m/z   Intensity
   b6++      426.48       22.9
   y4+       431.31       23.6
   b7++      475.85       21.3
   a8++      518.38       28.2
   b8++      532.32       19.6
   y6+       605.39       40.7
   y7+       704.43       33.3
   a12++     705.48       22.9
   b12++     719.38       24.9
   a13++     761.90       39.7
   b13++     776.23       71.7
   y16++     796.34       36.1
   a14++     805.40       46.3
   b6+       851.24       21.0
   b15++     870.33       25.9
   y9+       892.51       50.2
   b16++     919.77       31.8
   b7+       950.50       18.1
   y19++     951.82       92.4
   b17++     962.80       23.7
   y10+     1005.56       37.5
   b18++    1006.67       40.4
   a8+      1035.29       33.0
   b20++    1070.83       29.8
   y11+     1076.57       24.6
   b21++    1134.80       22.6
   y14+     1379.88       18.4
```

```
606. MORG-339_phosphopeptide_TMT_30%ACN_032017_OT.7048.7048.4 (kVskQEEASGGPLAPk)

   Ion         m/z   Intensity
   y1+       376.39       18.3
   y2+       473.37       70.9
   y3+       544.40       29.6
   b12+++    579.73       21.6
   y13+++    590.22       25.4
   a6++      605.39       17.1
   a13+++    607.60       24.4
   b13+++    617.48       25.0
   b3+       623.93       51.9
   a14+++    632.12       98.4
   b14+++    640.63       60.0
   y11++     642.88       60.2
   y14+++    645.97       58.6
   a7++      670.40       18.1
   b7++      684.77       68.3
   y12++     706.50       23.8
   a9++      749.23       18.5
   y5+       754.35       28.4
   y6+       811.36       18.6
   b12++     868.66       18.1
   y13++     885.89       24.4
```

```
607. MORG-339_phosphopeptide_TMT_18%ACN_032017_OT.19035.19035.4 (sLAALDALNtDDENDEEEYEAWkVR)

   Ion         m/z   Intensity
   b1+       317.23       21.9
   b5++      343.62       10.8
   b2+       430.26       10.9
   b7++      436.47       20.2
   y8+++     437.25       11.3
   b8++      492.95       26.4
   b3+       501.41       21.6
   b9++      549.46       11.0
   b4+       572.41       22.9
   b10++     640.17       13.9
   y8++      655.42       12.6
   b5+       685.50       18.3
   b11++     697.80       10.4
   y9++      719.96       10.5
   y16+++    779.67        9.9
   y10++     784.65       18.8
   b6+       800.46       26.5
   b19+++    806.78       12.5
   y17+++    817.96       43.7
   b13++     819.76       13.6
   a20+++    840.36       11.3
   y11++     841.55       50.8
   a7+       843.27       22.9
   b20+++    849.95       35.5
   b7+       871.63       36.3
   b21+++    873.68       16.9
   y19+++    879.06       21.8
   y12++     899.20       43.0
   y20+++    917.53       12.6
   y21+++    954.83       19.7
   y13++     963.70       36.0
   a16++     984.67       30.1
   y14++    1021.26       40.5
   a9+      1070.50       20.6
   y15++    1078.75       31.7
   b9+      1098.68       14.0
   y16++    1168.82       21.7
   b19++    1209.53       10.4
   y17++    1226.26       44.9
   y20++    1375.84       10.9
```

```
608. MORG-339_phosphopeptide_TMT_16%ACN_032017_OT.12364.12364.2 (cVGLPSATDtS)

   Ion         m/z   Intensity
   y3+       402.07       10.1
   b3+       489.36       18.9
   y4+       503.22       25.6
   a4+       574.22       19.1
   b4+       602.69       10.9
   y6+       661.25       12.6
   b5+       698.99       25.7
   a6+       758.28       54.8
   b6+       786.52       18.3
   y8+       871.45       10.8
   y9+       928.42       22.1
   y10+     1027.43       10.7
```

```
609. MORG-339_phosphopeptide_TMT_4%ACN_032017_OT.7740.7740.2 (sEDQDLQGLR)

   Ion         m/z   Intensity
   b1+       397.23        4.3
   y4+       473.13        4.6
   b2+       526.46        4.8
   y5+       586.43       15.6
   b3+       641.36       11.1
   y6+       701.53        7.5
   y7+       829.44       14.2
   b5+       884.43       12.4
   y8+       944.67       12.5
   y9+      1073.61        9.1
   b7+      1125.66        9.5
   b8+      1182.55        7.2
   b9+      1295.78        8.2
```

```
610. MORG-339_phosphopeptide_TMT_4%ACN_032017_OT.3241.3241.2 (tRLAsESANDDNEDS)

   Ion         m/z   Intensity
   a3+       572.43        8.6
   y5+       579.19        6.8
   b3+       600.55        9.1
   b4+       671.51       11.2
   y7+       808.25       16.4
   y8+       879.40       12.9
   y10+     1095.45        6.7
   b8+      1125.58        6.7
   b9+      1239.69        9.2
   y11+     1262.50        7.5
   y12+     1333.42        6.7
   b10+     1354.67       10.5
   y13+     1446.64        8.2
   b11+     1469.66       16.6
```

```
611. MORG-339_phosphopeptide_TMT_10%ACN_032017_OT.12819.12819.3 (sAYQDYDsDsDVPEELkR)

   Ion         m/z   Intensity
   b1+       317.24       10.6
   y6++      500.80       23.5
   y7++      550.46       23.0
   b3+       551.30        8.7
   y9++      691.26       13.4
   y10++     749.12       11.2
   b5+       794.32       14.5
   y11++     832.38       14.3
   a12++     854.06        9.2
   b12++     868.68       15.5
   y12++     889.91       16.3
   a13++     902.88       10.8
   y5+       903.92       13.5
   y13++     971.42       14.5
   y14++    1028.97       11.8
   b15++    1046.26        9.3
   b7+      1072.35       12.1
   y7+      1099.55       10.1
   y16++    1174.32        9.6
   y17++    1210.26        8.8
   b11+     1636.64       10.7
   b12+     1735.64       10.6
```

```
612. MORG-339_phosphopeptide_TMT_14%ACN_032017_OT.15771.15771.3 (eVWAsDsGHEEDcLsPEPPLRPk)

   Ion         m/z   Intensity
   b2+       458.43        9.4
   y6++      468.99       34.5
   y2+       473.38       10.2
   y8++      582.10       26.7
   a9++      665.46       13.3
   b4+       715.51        9.6
   b10++     744.63       14.5
   y11++     801.74       12.4
   y13++     924.25       12.6
   y6+       936.66       28.4
   y14++     988.26       14.1
   b6+       997.45       18.4
   y15++    1057.43       13.1
   a15++    1072.83       24.6
   y16++    1085.84       21.3
   a7+      1136.28       10.6
   y8+      1162.56       16.2
   y17++    1168.85       15.8
   b17++    1199.81       16.6
   y18++    1226.55       14.3
   b18++    1247.79        9.5
   b19++    1296.72       10.3
   a9+      1330.31       12.9
   b21++    1431.09       11.4
   y21++    1438.74       12.5
   b12+     1731.65       12.3
   a14+     1976.61        9.2
```

```
613. MORG-339_phosphopeptide_TMT_18%ACN_032017_OT.2413.2413.4 (ekNGsDDDDDEkPGkR)

   Ion         m/z   Intensity
   b1+       358.84       14.5
   y6+++     391.71       14.4
   a4++      430.00       21.0
   y8+++     468.48       15.1
   y9+++     506.76       28.7
   y5++      522.56       33.1
   y10+++    545.14       26.8
   b6++      585.54       32.0
   b11+++    587.04       49.0
   a7++      628.56       13.5
   b7++      643.15       21.0
   y7++      644.38       34.6
   y13+++    658.21       21.2
   y4+       686.41       40.4
   a2+       688.33       24.0
   a9++      743.81       12.8
   b9++      757.88       22.5
   y9++      759.59       37.9
   b10++     815.29       27.8
   y10++     817.21       36.5
   b3+       830.66       23.0
   a4+       859.77       35.7
   a11++     865.47       13.6
   y11++     874.60       22.0
   b15+++    876.74       18.1
   b11++     880.01       22.4
   b12++    1058.66       15.5
```

```
614. MORG-339_phosphopeptide_TMT_20%ACN_032017_OT.7958.7958.3 (sGVAVPTsPk)

   Ion         m/z   Intensity
   y2++      236.98       13.3
   b4++      272.58        8.4
   b1+       317.20       10.0
   y3++      320.68        8.2
   b5++      322.25       13.1
   y4++      370.92        7.0
   y5++      419.79       81.2
   b7++      420.88       18.1
   y6++      469.44       20.1
   y2+       473.42       16.4
   a9++      539.24        5.2
   b4+       544.41       20.9
   y8++      554.60        5.2
   y9++      582.86        6.0
   y3+       640.48        7.5
   b5+       643.45       74.2
   a6+       712.54        7.2
   y5+       838.47       16.2
   b7+       841.60        5.1
   y7+      1008.54        5.6
```

```
615. MORG-339_phosphopeptide_TMT_14%ACN_032017_OT.7664.7664.3 (aEPRAPsPAPAVsPk)

   Ion         m/z   Intensity
   b4++      342.25        8.6
   b2+       430.33       14.0
   y2+       473.35       24.3
   y7++      489.42        8.9
   b7++      509.75       10.1
   a9++      579.88       10.1
   b9++      593.94       31.0
   a4+       655.44       12.1
   a11++     664.08       14.4
   y10++     670.42       12.7
   b11++     677.98       26.2
   b4+       683.57       26.8
   a12++     713.11       17.9
   a5+       726.41        8.4
   b12++     727.58       18.1
   y4+       739.02       13.1
   y12++     784.02       10.6
   a13++     797.23       10.1
   b13++     810.70       14.8
   a6+       823.40        8.5
   y13++     832.60       16.2
   b6+       851.89        8.8
   y6+       907.56       20.1
   y7+       978.55       10.5
   y8+      1075.63       17.4
```

```
616. MORG-339_phosphopeptide_TMT_18%ACN_032017_OT.13680.13680.3 (sLSPAATVTAELHYEQGsPR)

   Ion         m/z   Intensity
   y2+       272.16       17.9
   b1+       317.23       13.7
   a7++      415.18       10.5
   b8++      478.55       11.8
   y8++      527.25       10.9
   y9++      584.19       10.6
   b11++     629.61       20.6
   y11++     684.26       15.4
   b12++     685.53       14.3
   a13++     740.29       17.2
   y6+       753.25       14.6
   b13++     754.36       22.3
   b6+       756.59       16.9
   y13++     784.27       22.9
   b7+       857.49       12.0
   a15++     886.67       10.7
   b15++     900.79       27.9
   y16++     905.52       12.8
   y7+       916.41       20.8
   a16++     950.72       11.4
   y17++     954.00       12.7
   b8+       956.54       23.2
   b17++     992.91       14.9
   a9+      1029.85       10.8
   y8+      1053.49       25.3
   y19++    1054.35       15.2
   b9+      1057.68       13.1
   b18++    1076.55       12.4
   y9+      1166.49       11.5
   b12+     1370.73       19.6
```

```
617. MORG-339_phosphopeptide_TMT_18%ACN_032017_OT.10702.10702.3 (aLsLDNSSQVSQk)

   Ion         m/z   Intensity
   a1+       273.29        9.3
   y3++      296.21        9.4
   y4++      345.87       11.6
   y1+       376.32       27.9
   y5++      409.75       12.7
   b2+       414.26        7.8
   y6++      453.30       11.0
   b6++      462.21        9.5
   a7++      491.58       11.8
   y7++      496.92       10.1
   y2+       504.35       26.8
   b7++      505.67       27.5
   b8++      549.15       14.5
   b3+       581.21        7.6
   y3+       591.42       49.8
   a9++      598.97       14.3
   y9++      611.01       14.0
   b9++      613.62       33.7
   a10++     648.66       12.4
   b10++     662.95       69.6
   y4+       690.42       26.2
   a11++     692.49        9.4
   b4+       694.42        9.8
   b11++     706.45       28.3
   b12++     770.46       15.5
   b5+       809.63        9.6
   y5+       818.61       11.6
   b6+       923.43       15.5
   y7+       992.44        9.0
   b7+      1010.59       14.2
   b8+      1097.54        9.6
   b9+      1225.59       18.0
   b10+     1324.69        9.0
```

```
618. MORG-339_phosphopeptide_TMT_25%ACN_032017_OT.22020.22020.4 (gLNLDGTPALSTLGGFsPAskPSsPREVk)

   Ion         m/z   Intensity
   y5++      429.34       30.0
   y9+++     522.82       14.4
   b9++      535.16       14.8
   y7++      555.98       20.3
   b10++     591.58       21.3
   y11+++    601.92       36.2
   y8++      604.83       28.0
   b11++     634.52       29.3
   y13+++    690.41       13.5
   y14+++    739.23       12.4
   b5+       742.41       14.9
   y16+++    777.23       12.2
   y9++      783.44       12.2
   b6+       799.59       11.8
   y18+++    848.52       12.0
   y5+       857.81       15.8
   y10++     867.13       12.3
   a22+++    891.33       13.6
   b22+++    900.40       55.6
   y11++     902.25       19.1
   y20+++    915.27       17.2
   a23+++    920.14       32.3
   b23+++    929.35      182.7
   y21+++    938.61       23.9
   a17++     942.36       14.1
   b24+++    985.77       15.4
   b8+       997.31       11.8
   b25+++   1017.48       12.5
   y24+++   1023.59       12.4
   y6+      1024.48       13.1
   b9+      1068.60       21.9
   b26+++   1069.56       12.9
   y14++    1108.13       15.6
   b27+++   1113.35       20.0
   a28+++   1136.37       11.9
   b28+++   1146.26       18.4
   y16++    1165.24       26.4
   b10+     1181.66       13.7
   y19++    1315.99       12.0
   y20++    1371.80       11.2
   a23++    1380.01       24.7
   b13+     1482.92       12.0
   b14+     1539.87       13.1
   b16+     1743.94       12.1
```

```
619. MORG-339_phosphopeptide_TMT_10%ACN_032017_OT.2446.2446.3 (gGtPAGsTRGSPTRPNPPVR)

   Ion         m/z   Intensity
   y9++      517.33       13.5
   b3+       525.20       13.9
   y12++     667.65        8.5
   y6+       679.75       12.1
   a12++     694.27       10.6
   y13++     718.28       13.5
   a13++     744.85       30.1
   b13++     759.06       32.1
   y14++     801.89        9.5
   y15++     829.95       17.4
   y7+       835.15       10.3
   b14++     836.97        9.8
   y16++     865.08       28.4
   y17++     914.08       52.6
   b7+       917.29        8.8
   y8+       936.41        9.2
   b8+      1018.28        9.8
   b19++    1089.20       11.0
```

```
620. MORG-339_phosphopeptide_TMT_22%ACN_032017_OT.10425.10425.4 (qVDTEEAGMVTAAtASNVkASPkR)

   Ion         m/z   Intensity
   y3++      315.25       18.3
   a13+++    501.96       24.2
   b8++      530.42       40.4
   a3+       544.12       25.7
   b3+       572.34       24.1
   b9++      595.69       23.5
   a16+++    615.00       22.3
   a4+       645.50       22.8
   y14+++    647.72       24.4
   b11++     696.10       27.3
   y9++      722.58       33.3
   y16+++    724.54       19.6
   y17+++    743.55       22.3
   y10++     758.22       23.7
   b13++     767.10       25.3
   a5+       774.54       31.8
   y5+       787.63       74.6
   b20+++    838.02       22.9
   y11++     849.18       38.7
   b14++     857.26       20.8
   b21+++    867.50       17.5
   y12++     884.77       46.0
   y21+++    886.79       22.7
   a22+++    890.17       19.8
   a6+       903.90       21.2
   y13++     920.19       31.9
   a16++     922.24       37.6
   b6+       931.32       38.0
   y23+++    958.18       18.4
   y14++     970.80       56.3
   b7+      1002.45       23.8
   b23+++   1019.21       24.9
   y15++    1020.25       32.7
   b8+      1059.55       40.1
   b9+      1190.74       23.5
   b10+     1289.64       18.1
```

```
621. MORG-339_phosphopeptide_TMT_14%ACN_032017_OT.15408.15408.3 (dEkkEESEEsDDDmGFGLFD)

   Ion         m/z   Intensity
   b1+       345.26       12.7
   b5++      659.53       16.2
   b6++      724.14       23.1
   b7++      767.73       11.9
   y7+       802.46       17.2
   a3+       803.49       12.6
   b8++      832.10       22.1
   b9++      896.31       16.1
   y8+       917.35       18.5
   y16++     959.05       41.7
   b10++     980.05       11.1
   a11++    1023.29       13.8
   b12++    1094.99       21.2
   b13++    1152.78       35.8
   b14++    1226.22       15.2
   b15++    1254.60       10.6
   y11+     1314.61       11.4
   b17++    1356.76       11.2
   y19++    1381.49       18.3
```

```
622. MORG-339_phosphopeptide_TMT_6%ACN_032017_OT.10340.10340.3 (rEVLYDsEGLsADER)

   Ion         m/z   Intensity
   y2+       304.18       20.2
   b3++      307.99       10.0
   b4++      364.37       10.1
   b5++      446.11       13.5
   y4+       490.25       11.8
   b6++      503.45       33.8
   b2+       515.38        6.4
   a8++      637.29        6.1
   b8++      651.27       15.8
   y5+       657.27       14.9
   a9++      665.41        7.0
   b9++      679.49       20.8
   y11++     701.23        9.2
   a10++     722.43       15.7
   b4+       727.44       10.8
   b10++     736.22        8.0
   y12++     758.13       10.0
   y6+       770.74       13.9
   a11++     806.22        7.8
   y13++     807.03       14.2
   b11++     820.29        6.9
   y7+       827.50       10.3
   a12++     841.42        5.2
   b12++     855.25       10.4
   b13++     912.79       25.2
   y8+       956.46        6.6
   a14++     963.04        8.2
   b6+      1005.67        8.5
   y9+      1123.33       13.2
```

```
623. MORG-339_phosphopeptide_TMT_4%ACN_032017_OT.1897.1897.3 (eMsPGHGPGETR)

   Ion         m/z   Intensity
   y2+       276.26       20.5
   a3++      314.86        5.1
   b1+       359.09        8.2
   b4++      377.24       19.9
   b5++      405.71        4.6
   y9++      454.40       49.2
   a6++      460.61       16.2
   a2+       462.53       12.2
   b6++      474.80       28.0
   a7++      489.41       90.4
   b2+       490.04       32.5
   b7++      503.27       25.3
   y5+       559.34       39.4
   y11++     603.32        8.2
   y6+       616.38       28.2
   a10++     631.02        6.7
   b10++     644.78       31.6
   b3+       657.20        9.0
   y7+       753.43        5.8
   b5+       811.43       10.6
   b6+       948.43        5.1
```

```
624. MORG-339_phosphopeptide_TMT_25%ACN_032017_OT.21027.21027.3 (gMYDGPVFDLTTtPk)

   Ion         m/z   Intensity
   y1+       376.34        9.2
   y5++      428.89       11.5
   y2+       473.36       35.0
   y7++      543.16       12.9
   b8++      549.09        9.0
   b9++      605.97       12.8
   y3+       654.49       14.3
   b10++     662.55       16.1
   a4+       668.46       11.4
   b4+       696.60       22.4
   a11++     699.65       11.1
   b11++     713.72       26.1
   y10++     714.57       19.1
   a12++     749.81        9.9
   y4+       755.19       23.9
   y12++     800.44        7.0
   b13++     854.51       18.8
   y5+       856.46        9.8
   y13++     881.75        8.7
   b7+       949.56        8.0
   y7+      1084.67        8.0
   b9+      1211.87        8.5
```

```
625. MORG-339_phosphopeptide_TMT_20%ACN_032017_OT.4005.4005.3 (esEEEQPPkkTSk)

   Ion         m/z   Intensity
   b1+       359.27        6.6
   b2+       526.42        7.3
   y3+       564.44        5.5
   y6++      688.50        8.8
   y7++      737.17       29.5
   a9++      782.50       11.1
   b4+       784.29        5.8
   b9++      796.75       10.5
   y8++      801.37       16.9
   y9++      865.93       13.6
   b5+       913.42        7.9
   y4+       921.39        8.4
   y10++     930.39       11.3
   a10++     961.27        6.6
   b10++     975.42        6.6
   y11++     994.65       13.4
   a11++    1012.29        7.4
   b6+      1041.43       14.0
   y12++    1078.27        8.9
```

```
626. MORG-339_phosphopeptide_TMT_18%ACN_032017_OT.19859.19859.3 (iAsDEEIQGTkDAVIQDLER)

   Ion         m/z   Intensity
   b2+       414.48        7.0
   y3+       417.38        9.1
   b8++      598.51       14.8
   y5+       660.44        7.3
   b4+       696.36       12.8
   y6+       773.49        8.4
   y12++     787.42       10.1
   a5+       797.42        8.5
   b5+       825.64        9.6
   y13++     851.30        9.5
   b11++     856.27       16.4
   y7+       872.48       10.8
   a12++     899.40       11.0
   y14++     908.15       19.0
   b12++     913.25        9.8
   y8+       943.57       18.7
   b6+       954.25        9.4
   y15++     972.83       14.4
   a14++     984.28        6.4
   b14++     998.69       14.2
   y16++    1037.39       26.0
   b15++    1055.38       10.7
   y9+      1058.82        8.9
   b7+      1067.59        8.6
   y17++    1094.86       22.4
   b16++    1118.96        9.5
   b17++    1176.82       18.1
   y18++    1177.80       10.2
   b8+      1195.61        7.5
   b19++    1297.95       13.9
```

```
627. MORG-339_phosphopeptide_TMT_25%ACN_032017_OT.16249.16249.3 (rAsGQAFELILSPR)

   Ion         m/z   Intensity
   y2+       272.29       11.5
   b3++      313.01        8.1
   y6++      349.67       12.6
   y3+       359.21       18.6
   a5++      391.49       18.8
   b5++      405.31       30.6
   y7++      414.66        8.3
   b6++      440.86       26.0
   y4+       472.40       26.8
   a7++      500.23       16.0
   b7++      514.44       24.8
   a8++      564.76       12.9
   b8++      578.69       31.2
   y5+       585.42       28.9
   y10++     587.22       19.8
   a3+       595.96       12.3
   b9++      635.32       29.2
   a10++     677.95       14.2
   b4+       681.79       10.3
   b10++     691.96       32.5
   y6+       698.53       27.9
   y12++     699.47       17.4
   y13++     734.57       10.2
   b11++     748.53       31.4
   b12++     792.05       11.5
   b5+       809.36        8.3
   y7+       827.51       25.7
   b13++     840.65        7.9
   y8+       974.60       27.6
   y9+      1045.59       24.3
   y11+     1230.66       13.9
```

```
628. MORG-339_phosphopeptide_TMT_10%ACN_032017_OT.14524.14524.2 (sVDISLGGsPR)

   Ion         m/z   Intensity
   y2+       272.22       15.2
   b1+       397.19       12.3
   b2+       496.27        8.2
   y5+       553.22        7.6
   b3+       611.34       17.2
   y6+       666.21       10.9
   a4+       696.48        8.6
   b4+       724.38       16.5
   y7+       753.36       14.9
   b5+       811.27       11.2
   y8+       866.47       17.1
   b6+       924.53        6.4
   b7+       981.34        8.1
   y10+     1080.47       10.8
```

```
629. MORG-339_phosphopeptide_TMT_4%ACN_032017_OT.7511.7511.3 (lYERRGGGsGGGDEsEGEEVDED)

   Ion         m/z   Intensity
   y3+       378.12        8.5
   b5++      474.73        7.5
   y4+       477.22        7.3
   a3+       607.46        8.5
   b9++      643.78        8.6
   b10++     672.04        7.2
   b12++     729.07        7.2
   a13++     772.05       27.2
   b13++     786.27       22.1
   y7+       792.32        7.5
   a14++     837.28        8.7
   b14++     851.27       23.7
   y17++     892.48       14.6
   a5+       919.79        7.6
   y8+       920.98        9.8
   y19++     999.41       10.7
   b17++    1027.22        8.4
   y9+      1088.40       10.2
   b18++    1092.22       19.1
   b19++    1156.32       15.2
   b20++    1206.44       31.1
   y10+     1217.34       13.3
   b21++    1263.87       19.7
   y12+     1389.58        7.8
   y14+     1503.61       10.8
```

```
630. MORG-339_phosphopeptide_TMT_25%ACN_032017_OT.18270.18270.4 (sLsEQPVVDTATATEQAkQLVk)

   Ion         m/z   Intensity
   b1+       317.20       14.3
   y1+       376.29       31.0
   y2+       475.36       43.1
   b6++      476.39       20.7
   b7++      525.91       12.0
   y5++      537.48       26.5
   y6++      573.17       13.7
   y3+       588.60       25.3
   b10++     683.10       20.6
   y8++      701.69       22.2
   a11++     704.50       13.8
   y4+       716.56       31.0
   b11++     719.12       20.9
   y9++      752.01       41.4
   y17+++    753.17       29.7
   b12++     769.67       35.1
   y10++     787.67       23.5
   y18+++    796.22       30.9
   b18+++    799.33       51.0
   b13++     805.21       12.4
   y11++     838.31       29.8
   y19+++    839.08       17.5
   b19+++    842.04       43.3
   b5+       854.51       29.0
   b14++     855.71       22.6
   y12++     873.89       27.4
   b20+++    879.75       93.4
   a21+++    903.52       14.4
   b21+++    912.83       65.0
   y14++     981.58       12.4
   b17++    1019.72       33.3
   a7+      1022.42       19.0
   b7+      1050.58       12.9
```

```
631. MORG-339_phosphopeptide_TMT_18%ACN_032017_OT.7172.7172.4 (eQmAISGGFIRRVtNDAR)

   Ion         m/z   Intensity
   y2+       246.18       31.5
   b5+++     273.28       10.7
   b3++      317.78       39.1
   b4++      353.22       24.1
   a5++      396.01       12.4
   b5++      409.86       15.7
   b6++      453.34       22.4
   y4+       475.33       15.8
   b2+       487.33       24.4
   b8++      510.59       30.6
   b12+++    531.68       10.6
   y14+++    548.14       30.2
   a13+++    555.28       16.3
   b13+++    564.64       31.7
   a9++      569.44       22.7
   y15+++    571.59       95.1
   a3+       606.24        9.8
   b14+++    625.08       12.0
   b3+       634.32       30.0
   y17+++    663.07       14.0
   y10++     664.27       15.7
   y11++     692.51       28.7
   b16+++    701.32       98.1
   a11++     704.30       11.8
   b4+       705.39       51.2
   b11++     718.27        9.2
   y12++     721.58       45.9
   b17+++    725.12       14.1
   y6+       755.78       12.8
   y13++     765.13      101.6
   b5+       818.51       19.3
   y14++     821.69       38.4
   y15++     857.09       23.9
```

```
632. MORG-339_phosphopeptide_TMT_25%ACN_032017_OT.22017.22017.3 (gLNLDGTPALSTLGGFsPASkPsSPR)

   Ion         m/z   Intensity
   a8++      485.35       12.7
   b9++      534.45       19.0
   y8++      569.48       25.5
   y9++      618.65       30.5
   y5+       623.30       27.8
   b5+       742.42       31.1
   y13++     832.86       17.9
   b7+       900.61       20.1
   a8+       969.33       13.3
   y16++     982.98       13.6
   a18++     990.14       34.3
   b18++    1004.73       37.2
   a19++    1026.22       16.1
   b9+      1068.56       15.9
   y19++    1123.96       16.0
   a10+     1153.97       27.3
   b10+     1181.77       15.5
   y21++    1202.82       26.1
   b11+     1268.92       22.9
   b12+     1369.72       12.7
   a23++    1380.28       37.7
   b14+     1539.90       12.3
   b15+     1596.90       14.2
```

```
633. MORG-339_phosphopeptide_TMT_22%ACN_032017_OT.10723.10723.4 (eEAMNRRLASNsPVLPQAFAR)

   Ion         m/z   Intensity
   b4++      345.39       23.1
   y7++      401.90       12.8
   b9+++     434.54       10.6
   a10+++    453.87       10.6
   b10+++    463.45       16.3
   y4+       464.27       16.1
   b2+       488.30       17.9
   y9++      499.98       33.1
   b11+++    501.08       18.9
   b12+++    556.96       23.8
   b3+       559.17       11.5
   a13+++    580.34       13.0
   y5+       592.23       16.8
   a8++      601.52       14.3
   a14+++    613.10       27.1
   b8++      615.58       18.0
   b14+++    622.60       48.1
   a9++      636.55       21.2
   y11++     640.61       15.6
   b9++      650.38       35.5
   b15+++    660.10       25.9
   a10++     680.85       11.8
   y6+       689.44       56.0
   b4+       690.41       31.8
   b10++     694.71       23.8
   y18+++    703.61       11.0
   a17+++    726.06       13.1
   y19+++    727.22       25.0
   b17+++    735.08       14.7
   b11++     751.36       11.4
   b18+++    758.96       17.3
   y20+++    769.66       12.9
   y7+       802.55       30.5
   b5+       804.45       14.4
   b19+++    807.74       11.8
   b12++     834.94       12.3
   y8+       901.55       11.5
   a6+       932.17       12.5
   b14++     933.23       16.8
   b15++     989.75       29.2
   y9+       998.59       60.9
```

```
634. MORG-339_phosphopeptide_TMT_16%ACN_032017_OT.12452.12452.3 (fSSQQASTkQsNAssDVEVEEk)

   Ion         m/z   Intensity
   y2+       505.35       14.6
   y3+       634.43       16.9
   a9++      698.51       11.0
   b9++      712.75       23.9
   y4+       733.56       19.7
   b10++     776.54       29.8
   b5+       807.80       15.9
   y5+       862.59       21.4
   b6+       878.53       15.2
   b13++     952.46        9.3
   y6+       961.64       20.6
   b7+       965.60       12.8
   a14++    1021.52       15.3
   b14++    1036.07        9.3
   b8+      1066.90       60.6
   y7+      1076.53       16.7
   y15++    1175.06       14.0
   b16++    1177.39       10.0
   y16++    1218.71       13.0
   b17++    1226.96       19.1
   y8+      1243.70       11.2
   y17++    1254.40        9.7
   b18++    1291.42       25.7
   b19++    1340.95       20.8
   b20++    1405.02       10.7
   y9+      1410.62       10.8
   b9+      1423.56       10.4
   b21++    1469.87       15.9
   y11+     1595.64       12.7
```

```
635. MORG-339_phosphopeptide_TMT_16%ACN_032017_OT.1761.1761.4 (qRRkPscSEGEEAEQk)

   Ion         m/z   Intensity
   y1+       376.35       26.1
   y6++      481.27       15.2
   y2+       504.39       28.2
   b4++      514.59       17.7
   b9+++     556.75       21.8
   b10+++    575.36       12.1
   a11+++    609.62       11.7
   y9++      618.00       30.0
   b11+++    618.95       27.1
   a6++      632.50       32.2
   a3+       642.57       17.5
   b6++      646.27       32.3
   a12+++    652.09       58.4
   b12+++    661.74       87.2
   b13+++    685.63       23.3
   y4+       704.32       20.8
   y14+++    724.89       11.6
   b14+++    728.63       44.1
   a15+++    761.93       10.8
   b15+++    771.39       29.0
   y15+++    777.51       11.2
   y11++     781.66       11.8
   a9++      820.46       21.2
   y12++     830.49       14.5
   y5+       833.50       15.5
   b9++      834.58       10.2
   b12++     992.13       14.6
   b4+      1027.69       17.4
   b14++    1092.27       10.7
```

```
636. MORG-339_phosphopeptide_TMT_8%ACN_032017_OT.10692.10692.2 (tAsLTSAAsIDGSR)

   Ion         m/z   Intensity
   y3+       319.15        9.8
   y4+       434.23       10.6
   y5+       547.35       12.4
   a4+       654.47        9.9
   y6+       714.40       12.2
   a5+       755.42       13.3
   b5+       783.49       12.8
   y7+       785.30       11.1
   a6+       842.60       10.1
   a7+       913.46        9.2
   b7+       941.43       11.6
   y9+       943.38       15.9
   a8+       984.13       10.3
   b8+      1012.55       13.1
   y10+     1044.33       12.3
   y11+     1157.40       13.5
   b11+     1407.66       15.6
```

```
637. MORG-339_phosphopeptide_TMT_18%ACN_032017_OT.10606.10606.4 (qGsFSEDVISHkGDLR)

   Ion         m/z   Intensity
   y4++      230.15       10.9
   y5+++     273.18       12.5
   y2+       288.18       43.1
   y6+++     318.91       12.2
   y7+++     347.85       12.0
   a4++      350.98       12.1
   y8+++     385.54       11.6
   y5++      409.08       16.5
   a9+++     415.25       25.3
   y9+++     418.59       25.2
   b9+++     424.51       12.5
   y10+++    457.07       24.3
   y4+       460.10       18.7
   b6++      472.72       17.9
   y6++      477.95       26.1
   a11+++    490.30       10.2
   y11+++    499.99       25.1
   a7++      517.10       12.8
   y7++      521.39       46.2
   y12+++    529.05       26.5
   b7++      530.36       11.7
   a3+       554.67      140.1
   b12+++    618.78       25.1
   y9++      627.52       22.8
   y14+++    633.71       10.6
   b13+++    637.35       20.3
   b14+++    676.10       85.4
   y10++     685.08       48.5
   a4+       701.27       11.2
   b15+++    713.78       12.1
   b4+       729.22       14.7
   y11++     749.57       33.2
   y12++     792.97       12.0
   b5+       816.46       11.5
   y13++     866.45       14.7
   b12++     927.42       13.6
   b6+       945.43       15.4
   b7+      1060.39       27.7
   b8+      1159.52       14.4
```

```
638. MORG-339_phosphopeptide_TMT_14%ACN_032017_OT.6375.6375.3 (sRsPLQPTSEIPR)

   Ion         m/z   Intensity
   y2+       272.33       10.0
   b1+       317.27       14.6
   b3++      320.68       17.9
   a4++      355.07       16.0
   y3+       385.25       20.2
   a5++      411.78       25.9
   b5++      425.91       23.0
   y8++      464.56        9.0
   b2+       473.55        8.2
   a6++      475.91       18.1
   b6++      489.82       67.1
   y4+       514.33       13.5
   b7++      538.49        9.0
   y10++     569.54       33.8
   a8++      574.58       23.7
   y5+       601.14       11.2
   b3+       640.37        9.7
   a10++     683.12       11.7
   b10++     696.76       19.8
   y6+       702.66       10.4
   b4+       737.41        8.7
   a11++     739.25       10.2
   b11++     753.35       19.0
   y7+       799.44       63.3
   b12++     801.97       11.5
   b5+       850.31       11.3
   y8+       927.52       22.5
   b6+       978.50       11.9
   y10+     1137.69       19.6
```

```
639. MORG-339_phosphopeptide_TMT_10%ACN_032017_OT.17875.17875.3 (dMDEPsPVPNVEEVTLPk)

   Ion         m/z   Intensity
   a1+       317.40        9.8
   y1+       376.38       11.6
   y2+       473.35       69.5
   y3+       586.47       21.7
   y10++     677.88       17.4
   y4+       687.52       28.7
   b4+       720.34       22.3
   b11++     745.64       13.7
   y5+       786.90       18.6
   a12++     796.19        8.9
   b12++     810.78       14.8
   b13++     875.21       27.8
   y14++     908.44       14.4
   a14++     910.34       13.8
   y6+       915.67       18.5
   b14++     924.66       31.9
   b15++     974.80       17.4
   b16++    1031.71       40.4
   y7+      1044.51        9.9
   a17++    1066.24        8.4
   b8+      1180.46       30.7
   b10+     1391.53        7.9
   b11+     1490.60       10.8
   b12+     1619.68       11.9
```

```
640. MORG-339_phosphopeptide_TMT_6%ACN_032017_OT.6267.6267.3 (nSQEDsEDsEEkDVk)

   Ion         m/z   Intensity
   y1+       376.28        5.7
   y2+       475.37        7.2
   b6++      485.35        5.1
   y3+       590.42        8.0
   y6++      603.59        9.3
   y7++      686.93        6.2
   b4+       688.42        5.7
   y8++      744.38        6.4
   b5+       803.44       22.3
   b11++     820.58        8.9
   y10++     892.56        9.0
   y11++     949.83        5.1
   a12++     984.64        6.6
   b12++     999.16        5.1
   b13++    1056.29       10.7
   y6+      1205.45        8.1
   b8+      1214.51        7.6
```

```
641. MORG-339_phosphopeptide_TMT_12%ACN_032017_OT.1312.1312.3 (sPPPSPTTQRR)

   Ion         m/z   Intensity
   y1+       175.15       11.0
   y3++      230.19       12.3
   a3++      282.21       10.7
   b3++      296.29        8.2
   a1+       369.31       15.4
   y6++      379.66       18.2
   b5++      388.21       10.2
   b1+       397.22       64.0
   y3+       459.39       13.2
   a2+       466.55        8.3
   y8++      471.55       31.5
   a7++      473.08       43.4
   b7++      487.08       74.6
   b2+       494.17       29.6
   y9++      520.39       21.3
   a8++      523.81        9.0
   b8++      537.83       10.0
   y4+       560.08       35.5
   y10++     568.99      150.8
   b3+       591.39       15.4
   y5+       661.54       10.3
   b10++     679.94       24.1
   y6+       758.48       12.1
   b5+       775.41       14.2
   y8+       942.56        9.7
```

```
642. MORG-339_phosphopeptide_TMT_14%ACN_032017_OT.12553.12553.3 (kDsEEGEESFSSVQDDLSk)

   Ion         m/z   Intensity
   y2+       463.36       16.1
   y3+       576.51       21.6
   y4+       691.42       23.4
   y11++     721.69       19.6
   y5+       806.43       22.2
   b14++     925.24       15.9
   a6+       927.57       16.8
   y6+       934.56       29.6
   y15++     943.45       27.5
   y7+      1033.67       16.0
   b16++    1040.10       14.2
   y17++    1091.96       12.8
   y8+      1120.32       20.3
   y9+      1207.68       19.2
   y10+     1354.75       11.8
```

```
643. MORG-339_phosphopeptide_TMT_10%ACN_032017_OT.10227.10227.3 (gADDAADADtAIINAEGGQNNSEEkk)

   Ion         m/z   Intensity
   b4+       588.36       16.8
   b6+       730.40       11.2
   y10++     775.25       13.8
   b7+       845.46       15.9
   y12++     875.09       13.0
   b8+       916.23       15.4
   y13++     932.48       12.4
   b18++     969.38       11.0
   b9+      1031.52       18.9
   y15++    1044.97       14.8
   a20++    1076.76       16.7
   y5+      1079.05       14.2
   y16++    1080.63       30.6
   b20++    1090.91       15.1
   y17++    1171.18       13.7
   a10+     1184.92       11.5
   b10+     1212.72       10.2
   y18++    1228.30       11.2
   y19++    1264.43       23.1
   y20++    1322.05       12.0
   y22++    1393.09       18.6
   y23++    1450.01       12.1
   y10+     1548.95       10.2
```

```
644. MORG-339_phosphopeptide_TMT_14%ACN_032017_OT.1962.1962.4 (rDsFSENEkQR)

   Ion         m/z   Intensity
   y1+       175.12       11.6
   b2++      251.14       29.6
   y4+++     264.09       10.7
   a3++      320.88       11.4
   y3++      330.80       24.5
   a6+++     334.79       22.1
   a4++      393.91       16.7
   y4++      395.45       12.3
   y5++      451.99       56.5
   a2+       473.09       30.3
   y9+++     478.84       10.0
   b2+       501.07       17.0
   y10+++    516.62       34.3
   b9+++     544.67       22.4
   a7++      559.71       13.4
   y7++      560.57       47.7
   b7++      573.31       16.1
   a10+++    577.81       11.7
   b10+++    587.28       28.9
   a8++      624.17       10.7
   y8++      634.00       59.1
   b8++      637.93       17.6
   y3+       660.54       10.0
   b3+       668.44       41.0
   y9++      717.45       30.9
```

```
645. MORG-339_phosphopeptide_TMT_16%ACN_032017_OT.5887.5887.4 (rRPGPHsPDPENLLTPR)

   Ion         m/z   Intensity
   y4++      244.01        8.6
   a4++      335.15       20.7
   y3+       373.35       14.5
   y10+++    384.26        9.2
   b9+++     437.30       25.6
   b6++      466.02       11.7
   y8++      470.37       18.9
   y4+       486.30       24.0
   a11+++    503.53       12.0
   b11+++    512.72       21.6
   a2+       514.58       14.5
   y13+++    517.84       17.4
   y9++      527.49       12.7
   a7++      535.47       17.9
   y14+++    537.41       18.7
   a12+++    541.52       22.9
   b2+       542.65       29.1
   b7++      549.07       13.9
   b12+++    550.79       29.7
   a13+++    579.06       13.2
   b13+++    588.36       26.3
   y5+       599.40       13.3
   y16+++    621.71       12.7
   b14+++    626.33       15.7
   a9++      641.23       10.5
   a15+++    649.97       10.4
   b9++      655.49       30.9
   y11++     659.63       10.1
   b16+++    692.36        9.8
   y6+       713.41       15.3
   y12++     728.62       12.2
   b11++     768.49       15.3
   y13++     776.56       16.3
   b12++     825.61       16.4
   b13++     882.05       12.7
   y8+       939.52       15.6
```

```
646. MORG-339_phosphopeptide_TMT_4%ACN_032017_OT.4489.4489.3 (aASEQQADAsGGDsPk)

   Ion         m/z   Intensity
   y5++      406.73        5.6
   b7++      458.57       10.9
   y2+       473.38       17.6
   b8++      515.61        6.9
   y7++      519.04       12.0
   b9++      551.58       10.6
   y8++      554.17       11.7
   b4+       588.41        8.4
   y9++      611.70        8.2
   b10++     634.82        8.0
   y3+       640.40       13.1
   y10++     647.25       12.3
   a11++     648.97        6.6
   b11++     663.27       10.2
   a12++     677.38       10.2
   a5+       688.87       15.2
   b12++     692.03       19.5
   a13++     735.28       15.0
   y4+       755.45       10.7
   y12++     775.51        6.6
   y5+       812.59        7.2
   b14++     833.03       10.0
   b6+       844.47       10.1
   y6+       869.50        6.1
   b15++     881.49        6.1
   b7+       915.54        8.7
   b8+      1030.53       11.1
   y7+      1036.34       10.4
   b9+      1101.49        9.4
   y9+      1222.49        5.3
```

```
647. MORG-339_phosphopeptide_TMT_12%ACN_032017_OT.10895.10895.2 (sNITsLS)

   Ion         m/z   Intensity
   y2+       219.09        7.0
   b1+       317.28        8.7
   y3+       386.19       10.8
   b2+       431.29        4.9
   y4+       487.20       14.7
   b3+       544.36       13.8
   y5+       600.30        4.8
   a4+       617.32        4.9
   b4+       645.45       17.0
   y6+       714.37       21.3
   b5+       812.43       15.7
   b6+       925.49       14.1
```

```
648. MORG-339_phosphopeptide_TMT_4%ACN_032017_OT.7454.7454.3 (sAEPTREPGAEAEsGsEsEPEPGPGPR)

   Ion         m/z   Intensity
   b7++      500.77        9.1
   b3+       517.35        4.3
   y6+       580.33       11.0
   b11++     678.07       10.3
   b5+       715.46        5.0
   b13++     778.02       10.1
   y8+       806.49       12.9
   y15++     875.93        5.0
   b7+      1000.49       11.9
   a17++    1023.66        9.8
   b17++    1037.82        9.1
   y19++    1040.70        5.3
   b8+      1097.91        5.3
   y10+     1102.59        7.7
   b18++    1121.19        4.0
   b19++    1186.27        9.4
   y23++    1282.11        4.0
   b21++    1299.32       11.3
   y13+     1455.54        4.4
   y16+     1822.65        4.1
```

```
649. MORG-339_phosphopeptide_TMT_14%ACN_032017_OT.14846.14846.3 (cPEILSDEsssDEDEkk)

   Ion         m/z   Intensity
   y5++      554.02       12.4
   y2+       733.68       13.9
   y9++      861.86       18.4
   y3+       862.69       11.9
   a13++     916.97       22.1
   b13++     931.78       11.4
   y11++     984.06       12.2
   y12++    1027.36       13.4
   y13++    1083.92       10.4
   y15++    1204.53        9.5
   y16++    1253.68       13.6
```

```
650. MORG-339_phosphopeptide_TMT_25%ACN_032017_OT.14902.14902.3 (sGSPkLDPSEVYLk)

   Ion         m/z   Intensity
   b6++      554.70       18.3
   b7++      612.55       66.1
   y3+       652.65       13.1
   b9++      704.00       15.2
   y4+       751.43       36.5
   a10++     754.51       40.5
   a11++     804.80       13.0
   b11++     818.33       13.0
   y11++     874.31       55.0
   y5+       880.56       15.6
   a12++     886.16       14.3
   b12++     900.22       17.8
   y13++     946.12       13.3
   b5+       995.77       14.0
   y7+      1064.61       45.8
   y8+      1179.77       15.3
```

```
651. MORG-339_phosphopeptide_TMT_16%ACN_032017_OT.9480.9480.3 (sVSPGVTQAVVEEHcAsPEEk)

   Ion         m/z   Intensity
   y1+       376.33        9.3
   b2+       416.36        9.9
   b3+       503.40        9.9
   b9++      528.77       13.3
   y4+       731.48       33.3
   b13++     756.49       14.2
   a14++     811.60       13.9
   y11++     812.51       10.5
   b14++     825.58       19.2
   b7+       857.48       14.7
   a15++     891.34       36.1
   y13++     896.98       27.4
   y5+       898.17       22.6
   b15++     905.45       24.6
   b16++     940.97        9.7
   y6+       969.42       14.3
   b8+       985.55        9.7
   a17++    1010.13       10.8
   a9+      1028.25        9.2
   b9+      1056.67       14.8
   y17++    1089.61       11.0
   b19++    1137.58       13.2
   y18++    1138.86        9.7
   b10+     1155.73       13.1
   y19++    1182.11        9.3
   b20++    1202.13       20.5
   b11+     1254.72       14.0
   b12+     1383.81       11.4
   b13+     1512.88       14.8
```

```
652. MORG-339_phosphopeptide_TMT_10%ACN_032017_OT.18046.18046.3 (eEsDTDLEVVLEk)

   Ion         m/z   Intensity
   a1+       331.44        4.8
   y4++      359.28        5.8
   y1+       376.28       33.6
   y5++      408.92        6.6
   y6++      472.83        4.7
   b2+       488.26        7.6
   y2+       505.35       46.0
   y7++      529.48        5.2
   b7++      550.40        9.9
   a8++      600.44       13.4
   b8++      615.03       27.3
   y3+       618.39       35.8
   y9++      638.03        8.7
   a9++      650.06        8.6
   b3+       655.47       14.2
   b9++      664.54       51.3
   a10++     699.54        8.7
   b10++     714.12       74.2
   y4+       717.54       19.2
   b11++     770.67       67.9
   y5+       816.58        7.2
   a12++     821.19        6.8
   b12++     835.21       50.5
   b6+       986.45        8.8
   b7+      1099.45        8.0
   b8+      1228.42       17.1
   b9+      1327.56       12.7
   b10+     1426.67        6.2
```

```
653. MORG-339_phosphopeptide_TMT_10%ACN_032017_OT.5808.5808.4 (sGSLGSSNGDSDtEREVSDSQAGQNTk)

   Ion         m/z   Intensity
   y1+       376.32       13.6
   y2+       477.39       11.6
   b4+       574.68       12.7
   y3+       591.56       16.2
   a12++     632.85       13.6
   y4+       719.55       13.7
   a19+++    721.34       18.4
   b20+++    759.80       18.7
   y5+       776.41       20.9
   a7+       777.44       17.8
   y19+++    778.68       28.7
   a14++     788.54       27.8
   a21+++    793.10       34.9
   b21+++    801.92       37.5
   b7+       805.41       24.6
   b22+++    826.30       31.2
   y21+++    845.51       14.2
   y6+       847.48       14.6
   y22+++    874.50       20.4
   y14++     889.11       23.4
   a8+       891.12       14.3
   y23+++    893.36       12.4
   b8+       919.93       28.9
   a26+++    950.33       15.8
   y25+++    959.67       27.6
   b9+       976.09       12.4
   b17++     994.66       35.8
   b18++    1038.14       13.5
   b19++    1095.42       18.8
   b21++    1203.03       14.4
   b22++    1238.62       15.0
   y21++    1267.33       11.9
```

```
654. MORG-339_phosphopeptide_TMT_10%ACN_032017_OT.15095.15095.2 (dLEFcstEEEk)

   Ion         m/z   Intensity
   b2+       458.41       10.7
   y2+       505.39       10.1
   b3+       587.44       13.3
   y3+       634.42       11.7
   b4+       734.45       12.3
   y4+       763.43       14.3
   b5+       894.69       11.9
   y5+       944.91       26.5
   y7+      1271.77        9.4
   y8+      1418.42       11.6
   b9+      1500.46       10.4
   b10+     1629.56        9.1
```

```
655. MORG-339_phosphopeptide_TMT_22%ACN_032017_OT.21314.21314.2 (sAkPETVIDSLLQ)

   Ion         m/z   Intensity
   b3+       825.52       40.0
   y8+       888.25       19.3
   b4+       922.17       36.1
   y9+      1017.38       10.1
   a5+      1023.86       12.1
   b5+      1051.44       16.1
   y10+     1114.64       28.9
   b7+      1251.72       11.5
   b8+      1364.81       14.5
   y11+     1471.82       11.0
   b9+      1479.83       15.9
   y12+     1543.16       10.9
```

```
656. MORG-339_phosphopeptide_TMT_14%ACN_032017_OT.7265.7265.4 (sYESSEDcPEAASSPtRkVk)

   Ion         m/z   Intensity
   b1+       397.31       14.5
   y3++      416.92       14.7
   y9+++     504.54       12.9
   y10+++    528.50       13.0
   a13+++    565.45       25.4
   y11+++    571.38       34.1
   b14+++    603.95       32.0
   y13+++    656.97       19.1
   a9++      669.13       13.7
   y7++      677.09       15.6
   b3+       689.34       23.1
   y14+++    695.83       16.7
   y8++      721.13       77.4
   a10++     733.55       21.9
   y15+++    737.97       39.2
   a17+++    739.14       22.0
   b10++     747.26       24.4
   b17+++    748.63       25.9
   y9++      756.67       32.0
   y10++     791.76       15.4
   y17+++    796.84       16.4
   b12++     818.22       14.3
   a5+       835.60       17.0
   a13++     847.68       23.3
   y11++     856.23       45.2
   b13++     861.63       19.6
   b19+++    901.17       21.9
   b14++     905.34       39.4
   b6+       992.43       13.6
   b7+      1107.41       28.5
   a17++    1108.39       21.0
   y5+      1169.50       15.6
   b8+      1267.46       24.3
```

```
657. MORG-339_phosphopeptide_TMT_6%ACN_032017_OT.5172.5172.4 (eREkEIsDDEAEEEk)

   Ion         m/z   Intensity
   a2++      244.08        7.9
   a3++      308.95        7.4
   a5+++     368.45        6.8
   y1+       376.34       26.1
   b5+++     377.56       20.1
   b6+++     415.37       13.5
   y6++      482.49        7.4
   a2+       487.30        9.2
   y10+++    491.40        7.8
   b4++      501.25        8.8
   y2+       505.43       30.0
   b8+++     509.52       18.4
   b2+       515.19       32.3
   a9+++     537.94        6.7
   b9+++     547.92       31.0
   a5++      552.23        7.1
   b5++      566.23       10.9
   b10+++    590.84       29.2
   a11+++    605.20        8.4
   a6++      607.99       13.6
   b11+++    614.51       33.2
   a3+       616.34       12.9
   b6++      622.78       57.1
   y3+       634.42       35.8
   b12+++    657.29       10.4
   b13+++    700.61       42.8
   y10++     737.66        9.4
   b14+++    743.62       35.7
   y4+       763.45       24.5
   b9++      821.38        9.6
   y5+       834.49       18.3
   a10++     871.87       17.4
   b10++     885.65       14.6
   b11++     921.24       17.3
   y6+       963.49       15.6
   y12++     980.68        8.2
   b12++     985.72       11.4
   b13++    1050.26        8.8
   y7+      1078.45       12.7
   y8+      1193.58        9.4
```

```
658. MORG-339_phosphopeptide_TMT_8%ACN_032017_OT.3734.3734.2 (sIQTSPTEEGR)

   Ion         m/z   Intensity
   y3+       361.19        6.1
   y5+       591.23        5.8
   y6+       688.35       33.3
   a4+       711.33       14.2
   b4+       739.56       28.7
   b5+       826.42        8.1
   b8+      1153.66        9.9
   b9+      1282.55       12.6
   b10+     1339.52        7.0
```

```
659. MORG-339_phosphopeptide_TMT_8%ACN_032017_OT.12094.12094.3 (eESETRsESsDFEVVPk)

   Ion         m/z   Intensity
   y1+       376.34        6.9
   y2+       473.38       24.8
   y6++      474.45       13.7
   b7++      564.82        6.6
   y3+       572.46       18.1
   b8++      629.48        7.1
   y9++      659.01        8.5
   y4+       671.49       15.0
   b9++      672.89       12.0
   y10++     723.59        6.7
   b10++     756.37       11.8
   a11++     799.41       14.4
   y5+       800.40       14.0
   b5+       805.62        6.6
   b11++     814.09       12.5
   a12++     873.80        7.8
   b12++     887.14       12.8
   y6+       947.59       11.1
   b13++     952.16       25.3
   b14++    1001.51       28.9
   b15++    1051.16       35.1
   y7+      1062.56        9.6
   b16++    1099.70        6.9
   y8+      1229.77        6.4
```

```
660. MORG-339_phosphopeptide_TMT_25%ACN_032017_OT.18576.18576.3 (nLEQILNGGEsPk)

   Ion         m/z   Intensity
   y1+       376.28       10.3
   b5++      414.10       13.5
   y6++      442.50       22.6
   b6++      470.75       13.5
   y2+       473.34       27.4
   y7++      499.53       21.9
   a8++      542.28        9.5
   b8++      556.23       21.3
   a3+       558.34       10.3
   a9++      571.28        9.7
   b9++      585.05       17.4
   b3+       586.56       10.2
   a10++     635.02       27.6
   y10++     676.43       10.8
   b4+       714.46       14.6
   b11++     732.50        9.5
   y4+       769.50       15.9
   y5+       826.50       10.4
   b5+       827.57       19.8
   y6+       883.51       10.0
   b6+       940.52       16.6
   y7+       997.40       12.4
   b7+      1054.72       11.9
   b8+      1111.67       11.0
   b9+      1168.75        7.5
```

```
661. MORG-339_phosphopeptide_TMT_10%ACN_032017_OT.7070.7070.3 (sStPLPTVSSSAENTR)

   Ion         m/z   Intensity
   a5++      384.36       15.7
   b7++      497.60       19.1
   y4+       519.41       17.6
   y10++     526.41        9.5
   b8++      546.84       26.6
   a3+       557.66       10.7
   y11++     574.87       66.9
   a9++      576.31       17.2
   b3+       585.36       10.5
   b9++      590.38       25.6
   y12++     631.38       30.7
   b10++     633.84       12.8
   b11++     677.29       18.5
   y13++     679.56       11.1
   b4+       682.69       12.0
   b12++     712.48       13.1
   a13++     763.17        8.8
   y7+       764.46       19.2
   a5+       767.38       11.8
   b5+       795.39       53.0
   b14++     834.47       10.6
   y8+       851.42       34.5
   b15++     884.78       14.6
   y9+       950.53       11.2
   b7+       993.49        9.4
   b8+      1092.40        8.6
   y11+     1148.61       18.2
```

```
662. MORG-339_phosphopeptide_TMT_20%ACN_032017_OT.16171.16171.3 (vGDTSLDPNDFDFtVTGRGSPSR)

   Ion         m/z   Intensity
   y3+       359.23       20.6
   b2+       386.08        9.5
   b3+       501.30       31.4
   y10++     549.47        9.3
   a4+       574.42       14.7
   b4+       602.33        9.0
   y11++     622.99       24.3
   a11++     681.60       10.6
   y13++     754.09       22.1
   a6+       774.56        8.3
   b6+       802.52       22.5
   y14++     811.35        9.0
   y15++     868.20       59.5
   a15++     952.83       10.1
   a9+      1100.56       11.6
   y20++    1125.32       21.6
   a19++    1138.78       13.2
   y21++    1182.61       10.0
   b20++    1196.43       10.2
   y22++    1211.52        9.1
   b10+     1243.58       18.9
   y11+     1244.69       13.2
   y12+     1359.61        9.3
   b12+     1505.75       17.7
   y13+     1506.81       15.7
```

```
663. MORG-339_phosphopeptide_TMT_18%ACN_032017_OT.7422.7422.3 (aPSPAPAVsPk)

   Ion         m/z   Intensity
   y2++      237.33        6.5
   a1+       273.14        7.8
   a2+       370.36       15.7
   y1+       376.10       10.3
   b2+       397.80        7.1
   y5++      405.40       12.4
   b7++      411.42       11.1
   y6++      454.35       45.8
   a3+       457.22        7.6
   b8++      460.91       20.4
   y2+       473.44       29.4
   b3+       485.44        6.0
   y7++      490.14       21.5
   y8++      538.33       12.2
   b9++      544.16        9.1
   a10++     579.11        6.7
   a5+       625.42       11.3
   y3+       640.31       16.0
   b5+       653.41       35.4
   a6+       722.40        8.2
   y4+       739.47       11.3
   b6+       750.36        6.3
   a7+       793.09        7.3
   y5+       810.74        8.3
   b7+       821.59       16.8
   b8+       920.58        6.1
   b9+      1087.64        8.5
```

```
664. MORG-339_phosphopeptide_TMT_8%ACN_032017_OT.5274.5274.2 (sHsLDSLsR)

   Ion         m/z   Intensity
   b1+       317.25        6.6
   b2+       454.24        6.4
   y4+       542.34       10.0
   b3+       621.56        7.6
   y5+       657.32       20.4
   b4+       734.76        7.5
   y6+       770.37       11.0
   a5+       821.69       12.2
   b5+       849.36       13.9
   y7+       937.42        6.1
   y8+      1074.29        6.2
```

```
665. MORG-339_phosphopeptide_TMT_18%ACN_032017_OT.17131.17131.3 (dIDASPsPLSVQDLk)

   Ion         m/z   Intensity
   a3++      273.34       11.5
   a1+       317.20       39.9
   y4++      366.78        8.7
   y1+       376.34       13.7
   b6++      415.21       64.1
   y5++      416.25       18.9
   y6++      459.86       10.6
   y2+       489.34       12.9
   b7++      497.85        8.7
   b8++      546.96        6.8
   y8++      565.14       19.7
   b3+       573.46        7.8
   a9++      589.12       16.1
   b9++      603.52       19.4
   y3+       604.34       15.0
   a4+       616.57        8.5
   b4+       644.50       14.4
   b10++     647.22       15.2
   y9++      648.43        6.9
   a11++     682.27       13.2
   b11++     696.04       13.7
   y10++     697.09       15.3
   y4+       732.56       19.8
   y11++     740.39       10.9
   b13++     818.23       10.7
   y5+       831.77        7.5
   b14++     874.61       15.8
   y14++     890.19        7.8
   y6+       918.47       15.5
   b7+       995.34       18.8
   b9+      1205.60       16.9
   b11+     1391.76       10.3
```

```
666. MORG-339_phosphopeptide_TMT_10%ACN_032017_OT.12460.12460.3 (sEPVVAsPALAGPsPR)

   Ion         m/z   Intensity
   y2+       272.22        8.7
   b1+       317.36        9.5
   b5++      371.25        7.6
   b6++      407.01        7.3
   y8++      424.24        5.2
   y3+       439.19        6.2
   b2+       446.31        7.2
   y9++      473.33       24.4
   b7++      489.80       16.2
   y4+       536.23       23.1
   b8++      538.96        5.2
   b3+       543.35       14.8
   y10++     556.84       15.5
   a9++      560.51        8.8
   b9++      574.38       40.9
   y11++     592.41       28.6
   y5+       593.29       44.4
   a4+       614.40       17.3
   a10++     616.47       10.9
   b10++     630.91       51.5
   y6+       664.30       39.1
   b11++     666.46       45.1
   b12++     694.88       30.8
   a13++     729.45        6.0
   b5+       741.48       23.6
   b13++     743.36        6.5
   y7+       777.36       19.8
   a14++     812.49       17.0
   y8+       848.28        6.6
   y9+       945.67       11.2
   b7+       979.57        6.2
   y10+     1112.46        9.5
   b9+      1147.65        7.0
   y11+     1183.70        7.8
```

```
667. MORG-339_phosphopeptide_TMT_18%ACN_032017_OT.11076.11076.3 (sMEVLSVTDGGsPVPARR)

   Ion         m/z   Intensity
   b1+       317.20       11.3
   y6++      348.34       17.8
   y9++      488.66       15.6
   y4+       499.39       18.1
   b3+       577.30        8.6
   y12++     646.36        8.8
   a4+       648.41        7.4
   b11++     653.00        7.8
   b4+       676.43       13.8
   y13++     689.37       16.4
   y6+       695.58       20.6
   y14++     746.77       10.1
   b13++     785.31        7.4
   b5+       789.70       10.4
   y15++     796.31        8.2
   a14++     820.52        8.2
   b6+       876.32        8.4
   y8+       919.36        8.0
   b7+       975.58        9.3
   y9+       976.64        7.9
   b17++     997.11       14.2
   b8+      1076.53        8.0
   y10+     1091.45        9.0
   b9+      1191.66       26.7
   y11+     1192.62       15.6
   b14+     1668.82       10.4
```

```
668. MORG-339_phosphopeptide_TMT_10%ACN_032017_OT.2469.2469.4 (sSEPVQHEENIRksSPEPR)

   Ion         m/z   Intensity
   y2+       272.21       27.2
   y4+       498.30       35.1
   b3+       533.36       22.2
   b8++      562.27       10.7
   y12+++    584.28       11.1
   a9++      612.44       20.2
   b9++      626.94       22.4
   y13+++    629.74       12.4
   b4+       630.71       23.9
   y8++      633.14       10.9
   a13+++    656.08       17.8
   b13+++    665.50       24.4
   a10++     669.50       34.2
   y14+++    673.06       39.0
   y15+++    705.63       21.1
   a14+++    711.43       22.6
   b5+       729.39       18.9
   y10++     747.37       15.6
   b15+++    749.96       17.7
   y6+       752.48       15.2
   a16+++    772.34       15.1
   y17+++    780.64       15.0
   y18+++    810.10       10.6
   y11++     811.54       14.9
   a17+++    816.12       16.1
   b12++     818.26       17.6
   b17+++    825.34       78.7
   b6+       857.61       20.2
   y13++     944.64       22.2
```

```
669. MORG-339_phosphopeptide_TMT_22%ACN_032017_OT.14586.14586.4 (kSAEPLANTVLAsETEEEGNAQALGPTAk)

   Ion         m/z   Intensity
   y4++      323.43       16.6
   y1+       376.51       15.2
   y6++      408.33       14.2
   y7++      443.75       15.1
   y2+       447.22       15.9
   y4+       645.56       58.5
   y5+       702.41       75.6
   y18+++    704.54       24.7
   b19+++    755.70       14.9
   y6+       815.65       36.9
   a6+       827.83       33.2
   b23+++    884.09       19.9
   y7+       886.75       31.1
   a15++     897.26       31.9
   a7+       898.87       25.6
   y24+++    908.25       27.6
   b15++     911.79       15.6
   a24+++    912.57       22.4
   y16++     936.98       37.4
   a16++     961.66       34.9
   b16++     975.88       15.1
   y26+++    983.66       18.1
   a8+      1012.35       20.4
   y8+      1014.76       25.5
   a18++    1090.64       27.8
   a19++    1119.86       13.8
   y23++    1305.66       31.6
   b11+     1353.72       16.4
```

```
670. MORG-339_phosphopeptide_TMT_10%ACN_032017_OT.3491.3491.3 (aRsPDIEQGk)

   Ion         m/z   Intensity
   a3++      298.77        7.7
   b3++      312.79       15.7
   y1+       376.36        9.8
   b5++      418.79       39.9
   y2+       433.22       20.1
   y6++      459.65        9.8
   a6++      461.35       18.4
   b6++      475.36       29.4
   y7++      508.22       37.5
   a7++      526.24       35.8
   b7++      539.89       26.5
   y3+       561.36       19.8
   b8++      603.97       19.1
   b3+       624.09        6.9
   b9++      632.19        8.4
   y4+       690.42       25.0
   a4+       693.79        6.8
   y5+       803.53       24.2
   y7+      1015.57       11.5
```

```
671. MORG-339_phosphopeptide_TMT_14%ACN_032017_OT.9388.9388.4 (gAkGkQADVADQQTTELPAENGETENQsPAsEEEk)

   Ion         m/z   Intensity
   y6++      500.93       35.2
   a9+++     505.45       36.0
   y7++      549.88       29.8
   y3+       634.40       41.4
   y16+++    722.69       72.7
   y10++     754.59       31.3
   a16+++    763.49       38.9
   b4+       772.06       38.0
   b16+++    773.14       78.2
   a17+++    801.52       31.3
   b10++     807.69       33.7
   b17+++    810.85       98.3
   a18+++    833.57       31.9
   b11++     865.24      111.1
   b19+++    866.24       57.6
   b20+++    909.91       53.6
   b12++     929.32       40.8
   y5+       930.45       40.5
   y14++     962.60       39.3
   b13++     993.31       52.4
   a23+++   1000.56       67.5
   y6+      1001.54       59.4
   b14++    1043.71       51.8
   b15++    1094.30       46.1
   y7+      1098.51       68.0
   y17++    1119.43       65.0
   b26+++   1124.45       35.0
   b5+      1130.21       35.1
   b16++    1158.49       86.6
   y18++    1167.85      299.0
   y29+++   1169.10       86.0
   b17++    1215.41      165.9
   y19++    1224.89       28.1
   y8+      1265.55       40.7
   b30+++   1279.08       60.2
   b19++    1299.56       63.8
   a7+      1300.42       41.3
   y21++    1339.85       28.1
   b20++    1364.01       85.4
   b21++    1421.11       71.2
   b8+      1443.93       29.6
   y10+     1507.58       31.6
   a9+      1514.61       67.0
   b11+     1729.12       30.6
   y14+     1923.67       36.8
```

```
672. MORG-339_phosphopeptide_TMT_10%ACN_032017_OT.6220.6220.3 (rTEsVPSDINNPVDR)

   Ion         m/z   Intensity
   b5++      441.57       11.3
   y4+       486.14       18.0
   y10++     563.70       16.2
   a8++      576.94       16.4
   b8++      591.39       27.6
   y5+       600.20       17.0
   y11++     613.55       12.7
   b3+       616.38       12.8
   a9++      634.29       18.4
   y12++     696.90       17.9
   b10++     704.61       17.8
   y6+       714.73       34.5
   a4+       755.45       20.3
   b11++     761.78       18.5
   a12++     796.45       13.0
   y7+       827.42       20.4
   b13++     860.08       11.6
   b5+       882.53       13.4
   b14++     917.77       17.9
   y10+     1126.61       36.7
   y11+     1225.61       12.6
```

```
673. MORG-339_phosphopeptide_TMT_10%ACN_032017_OT.15122.15122.3 (eSSPLysPGFSDSTSAAk)

   Ion         m/z   Intensity
   y1+       376.31       17.6
   b2+       446.63       12.5
   y2+       447.41       21.0
   y8++      498.39        9.3
   y3+       518.35       15.5
   y4+       605.54       20.6
   a9++      640.27        8.8
   y11++     649.10       21.3
   y5+       706.54       16.2
   a10++     713.69       11.0
   b10++     727.61       10.2
   y12++     732.64       39.8
   y6+       793.36       10.1
   y13++     853.46       14.0
   a13++     857.95        9.9
   a14++     908.52       12.5
   b15++     966.19       14.4
   b6+       986.42       29.2
   a16++     987.42       25.9
   y8+       995.56       16.9
   y16++    1002.30       16.3
   b17++    1037.77       30.8
   b10+     1454.58       17.0
```

```
674. MORG-339_phosphopeptide_TMT_14%ACN_032017_OT.16508.16508.3 (dmDEPsPVPNVEEVTLPk)

   Ion         m/z   Intensity
   a1+       317.34       12.6
   b5++      417.29        7.9
   y2+       473.45       56.7
   y3+       586.42       17.0
   b8++      599.13        9.7
   y10++     678.20       29.3
   y4+       687.57       24.8
   b4+       736.39       19.6
   b11++     753.78       18.4
   y5+       786.54       18.3
   b12++     818.75       25.4
   b5+       833.64       23.1
   b13++     882.99       17.6
   y14++     907.71       16.6
   y6+       915.68        7.8
   b14++     932.67       33.6
   b15++     983.26       22.8
   a16++    1025.55       16.4
   b16++    1039.74       29.2
   y7+      1044.62       12.1
   b8+      1196.41       32.0
   b10+     1407.42        9.4
   b11+     1506.66       12.0
```

```
675. MORG-339_phosphopeptide_TMT_16%ACN_032017_OT.16152.16152.4 (yGATDVRTPQPcsRPAsPEPELLPELDSk)

   Ion         m/z   Intensity
   y6++      459.30       11.5
   y2+       463.37       14.7
   y3+       578.32       14.6
   b10++     660.08       19.9
   a16+++    680.37       18.2
   y10++     685.80       12.0
   b16+++    689.68        9.0
   y4+       691.48       16.6
   b5+       737.38       40.3
   b12++     788.35       27.9
   y12++     798.53       19.3
   b19+++    820.90       35.2
   y13++     882.20        9.7
   a21+++    886.70       12.0
   b21+++    896.35       14.2
   y6+       917.53       43.4
   a22+++    924.86       18.8
   b22+++    933.82      105.2
   a14++     935.91       12.3
   y22+++    946.72       10.7
   y15++     966.28       27.6
   b23+++    971.41       24.4
   a15++     984.54       11.2
   b24+++   1004.02       10.5
   a16++    1020.23        9.8
   y7+      1030.71       18.7
   y24+++   1031.71       41.7
   b16++    1034.14        9.9
   b25+++   1046.78       19.1
   y25+++   1070.00       11.7
   b26+++   1085.00       11.5
   b27+++   1123.23       12.1
   y8+      1143.60       15.7
   y28+++   1146.55       13.3
   y18++    1207.33       12.2
   b19++    1230.61       14.7
   y21++    1368.75       16.4
   y10+     1369.78       25.9
   b22++    1400.48        9.5
   b23++    1457.06       19.2
```

```
676. MORG-339_phosphopeptide_TMT_16%ACN_032017_OT.10844.10844.4 (mDTGPQSLsGkSTPSsGDVQVTEDAVRR)

   Ion         m/z   Intensity
   b4++      317.33       16.4
   b2+       476.34       18.1
   y10++     587.16       29.0
   y19+++    766.97       43.5
   y14++     800.09       21.9
   a11++     806.93       18.2
   a19+++    808.70       89.4
   b11++     820.83       24.9
   y20+++    822.42       26.5
   a6+       831.19       17.4
   y7+       846.47       38.8
   y15++     848.32       62.8
   a12++     850.07       21.2
   a20+++    850.83       21.0
   b6+       859.60       18.1
   b20+++    860.23       19.1
   b12++     864.38       57.3
   b21+++    893.64       20.7
   a7+       918.18       15.6
   y23+++    931.60       13.1
   y17++     942.26       14.7
   y8+       945.46       14.1
   b7+       946.47       48.3
   a14++     949.18       12.6
   y26+++   1017.08       29.7
   a25+++   1022.40       13.8
   a8+      1031.05       14.8
   b8+      1059.23       22.5
   b16++    1090.39       18.9
   b27+++   1117.54       18.9
   b18++    1176.75       14.8
   b23++    1454.73       12.9
   b24++    1512.52       15.0
```

```
677. MORG-339_phosphopeptide_TMT_22%ACN_032017_OT.16328.16328.3 (akQPVIGDQNsDsDEmLAVLk)

   Ion         m/z   Intensity
   y1+       376.37       13.1
   b3++      393.64       10.1
   y2+       489.50       16.5
   b6++      548.68       19.8
   y3+       588.58       14.0
   b8++      634.61       28.7
   b2+       658.71       10.8
   y4+       659.60       15.7
   b10++     755.74       13.7
   y5+       772.57       14.0
   b3+       786.73       13.6
   b12++     896.79       23.8
   y6+       919.54       10.7
   a5+       954.85       12.1
   b13++     980.11       13.2
   b5+       982.71       24.4
   y14++     985.90       12.3
   y15++    1013.57       24.9
   a14++    1023.39       29.7
   b15++    1102.30       10.3
   b7+      1152.75       15.1
   b16++    1175.66       15.1
   a17++    1218.41       13.5
   y19++    1232.27       17.2
   b8+      1267.84       20.5
   b19++    1317.06       17.4
   y9+      1330.53       11.5
   b20++    1374.11       15.9
```

```
678. MORG-339_phosphopeptide_TMT_8%ACN_032017_OT.3377.3377.4 (eRERDHsPTPSVFNSDEER)

   Ion         m/z   Intensity
   y2+       304.21       15.2
   y6++      374.95       14.0
   y3+       433.24       19.4
   b8+++     440.01       19.3
   y7++      448.92       22.6
   b5++      458.39       26.4
   b9+++     473.66       14.8
   a10+++    496.47       15.1
   b6++      527.02       18.0
   b11+++    534.74       20.2
   y4+       548.28       26.3
   a12+++    558.66       19.6
   b12+++    567.89       43.3
   y10++     590.26       29.8
   a7++      596.26       14.9
   a13+++    607.48       83.8
   b7++      609.83       12.8
   b13+++    617.01       87.8
   y5+       635.30       43.0
   b15+++    683.94       26.5
   y12++     689.76       24.4
   b9++      709.23       15.8
   b16+++    722.17       25.8
   a10++     743.52       15.1
   y6+       749.35       49.5
   b17+++    765.41       31.0
   y14++     841.82       31.5
   b12++     851.20       38.4
   y7+       896.37       38.8
   y15++     898.40       16.1
   b5+       915.72       13.6
   b13++     924.69       26.2
   y8+       995.54       13.7
   y10+     1179.62       17.4
```

```
679. MORG-339_phosphopeptide_TMT_20%ACN_032017_OT.7984.7984.3 (aRDsPPPSSNIVQGQIk)

   Ion         m/z   Intensity
   y1+       376.30        9.7
   y2+       489.55       12.6
   b3+       572.35       17.9
   y3+       617.51       11.2
   y4+       674.57       25.3
   y11++     700.83       10.6
   a11++     702.46       13.0
   a4+       711.39       10.8
   b11++     716.44       27.5
   b4+       739.54       33.0
   y12++     749.02       29.2
   a12++     751.52       36.0
   b12++     765.81       28.6
   y13++     797.61       51.7
   y5+       802.51       16.6
   b13++     829.77       23.6
   b5+       836.40       16.2
   b14++     857.97       12.7
   y14++     881.17       13.1
   y6+       901.62       21.8
   a15++     908.32       14.5
   b15++     922.61       21.8
   b16++     979.27       14.5
```

```
680. MORG-339_phosphopeptide_TMT_18%ACN_032017_OT.15940.15940.3 (kGGEFDEFVNDDtDDDLPVSk)

   Ion         m/z   Intensity
   b2+       415.14       12.2
   y2+       463.41       18.3
   y3+       562.54       11.5
   y4+       659.48       48.5
   b11++     734.60       27.3
   y5+       772.51       34.9
   b12++     791.37       19.4
   b13++     882.04       12.5
   y6+       887.55       24.3
   a15++     983.38       14.4
   b15++     997.17       18.2
   y7+      1002.57       28.0
   y15++    1008.98       16.0
   b16++    1054.65       23.9
   y16++    1066.66       14.8
   y8+      1117.63       20.1
   b19++    1209.04       12.2
   a20++    1239.44       11.5
   y9+      1298.69       13.7
   a10+     1324.47       19.9
   y10+     1413.67       13.0
   b11+     1467.89       18.1
   y12+     1642.69       11.5
```

```
681. MORG-339_phosphopeptide_TMT_18%ACN_032017_OT.13538.13538.4 (gLGLGRVSSSPAREAPAGGSGDGLSSPSPLEysPR)

   Ion         m/z   Intensity
   b15+++    556.92       16.1
   b17+++    613.11       14.9
   b21+++    698.73       12.5
   b22+++    737.31       71.0
   b23+++    756.38       17.7
   a14++     784.75       12.4
   b24+++    793.96       30.0
   y23+++    816.83       14.1
   b25+++    823.05       23.8
   b15++     834.74       18.0
   y24+++    839.95       30.5
   a26+++    842.53       12.5
   y15++     861.47       20.0
   a16++     868.57       14.5
   y25+++    872.61       19.1
   a27+++    875.04       18.6
   b16++     882.52       29.4
   b27+++    884.39       12.9
   y26+++    901.25       35.9
   a28+++    903.95       19.2
   y16++     904.98       27.2
   b28+++    913.13       28.4
   b17++     918.41       24.2
   y27+++    930.32       13.6
   b30+++    982.74       15.8
   y29+++    991.96       17.0
   a9+      1028.66       20.0
   y20++    1045.62       14.2
   b21++    1047.24       16.8
   a32+++   1097.85       35.2
   y32+++   1101.43       18.1
   b22++    1105.26       32.0
   b32+++   1107.51       24.2
   y33+++   1119.98       14.0
   b23++    1133.85       20.3
   b24++    1190.10       13.3
   y24++    1259.34       14.7
   a26++    1263.43       13.2
   y13+     1549.69       21.6
```

```
682. MORG-339_phosphopeptide_TMT_16%ACN_032017_OT.13109.13109.4 (vARPQILEPRPQsPDLcDDDVEFR)

   Ion         m/z   Intensity
   y2+       322.18       15.7
   b10+++    464.20       17.1
   b11+++    496.52       10.7
   b7++      504.51       12.2
   b12+++    539.24       17.4
   y4+       550.43       15.8
   a8++      554.88       11.7
   b3+       556.59       12.8
   b8++      569.07       44.3
   b15+++    665.71       40.1
   a10++     681.36       53.3
   y11++     691.11        8.9
   a16+++    693.43       12.0
   b10++     695.20       11.7
   b16+++    703.40       15.2
   a11++     730.39        8.7
   b11++     743.87        8.5
   a17+++    747.48        8.3
   a5+       753.44       13.6
   y18+++    756.47       17.4
   y12++     774.42       34.3
   y6+       780.42       16.6
   b5+       781.58       19.3
   b19+++    833.58       22.9
   b20+++    871.91       32.4
   y7+       895.55       13.0
   b21+++    904.53       10.5
   y22+++    921.29        9.6
   y15++     964.65       30.8
   b23+++    996.69        9.0
   b15++     997.65        9.3
   y16++    1013.64       23.8
   y8+      1055.41       11.5
   y9+      1168.44       22.5
   y11+     1380.57       28.3
   y22++    1381.55       29.7
```

```
683. MORG-339_phosphopeptide_TMT_25%ACN_032017_OT.16048.16048.4 (kPPAPPsPVQSQsPSTNWSPAVPAk)

   Ion         m/z   Intensity
   b6+++     272.79       19.6
   y1+       376.38       21.0
   y6++      406.51       29.2
   a2+       426.93       28.0
   y7++      449.90       17.8
   y3+       544.41       89.6
   a14+++    587.47       23.3
   y4+       643.52       21.5
   y5+       714.68       15.1
   b5+       720.69       15.2
   y12++     742.54       19.4
   a19+++    779.07       17.5
   a6+       789.85       15.8
   y6+       811.58       65.3
   b6+       817.18       25.6
   y20+++    818.22       32.1
   b20+++    821.11       29.4
   y13++     826.31       37.6
   a13++     831.47       40.0
   a21+++    834.65       41.8
   a22+++    867.89       19.0
   b22+++    877.34       39.2
   b14++     894.26       15.7
   y7+       898.58       45.0
   a23+++    900.44       17.5
   b23+++    909.97       19.4
   y15++     933.03       16.0
   b15++     938.25       20.4
   y24+++    939.27       23.4
   y16++     997.91       15.8
   y8+      1084.76       25.4
```

```
684. MORG-339_phosphopeptide_TMT_14%ACN_032017_OT.4877.4877.4 (tLTDEVNSPDsDRRDk)

   Ion         m/z   Intensity
   y4+++     268.56       17.6
   y1+       376.21       12.4
   a5++      381.22       11.3
   y4++      402.33       28.2
   a6++      430.56       10.7
   y9+++     461.92       15.5
   a11+++    480.46       12.6
   y2+       491.38       10.6
   a8++      531.53       23.3
   y11+++    533.18       11.9
   y6++      543.57       20.5
   a13+++    571.50       18.8
   y12+++    576.44       24.9
   a9++      580.08       33.1
   b13+++    580.72       12.1
   b14+++    632.68       13.7
   y3+       647.36       11.0
   y14+++    648.37       14.0
   b4+       660.47       23.4
   a15+++    661.59       12.7
   b15+++    671.07       13.1
   y15+++    686.44       10.7
   y9++      693.14       14.6
   y10++     749.71       22.0
   a5+       761.18       10.2
   a12++     778.39        9.5
   b5+       789.43       23.6
   y11++     799.53       20.4
   b6+       888.52       14.8
   b7+      1002.61       15.5
   b12+     1583.73       10.1
```

```
685. MORG-339_phosphopeptide_TMT_10%ACN_032017_OT.7587.7587.3 (aEsPETSAVESTQSTPQk)

   Ion         m/z   Intensity
   b1+       301.35       15.1
   y1+       376.32       14.9
   b8++      541.66       14.1
   b9++      591.32       22.9
   y3+       601.42       66.5
   b10++     655.80       24.5
   b11++     699.42       19.5
   y11++     702.53       37.0
   y12++     746.78       11.6
   b12++     749.85       23.4
   y5+       789.42       41.2
   a5+       795.84       27.4
   y13++     797.09       22.2
   b13++     813.95       28.2
   b5+       823.47       15.8
   a14++     843.83       14.7
   b14++     857.47       47.9
   a15++     894.18       18.5
   b15++     907.58       19.8
   y6+       917.53       18.1
   b6+       924.26       19.4
   a16++     942.78       16.1
   y16++     993.22       13.7
   b7+      1011.51       16.6
   b17++    1020.94       11.9
   b8+      1082.56       14.3
   b9+      1181.53       33.9
   y9+      1234.69       11.8
   b10+     1310.68       29.0
   b11+     1397.75       13.0
```

```
686. MORG-339_phosphopeptide_TMT_20%ACN_032017_OT.8002.8002.3 (sPPEDAIPkSk)

   Ion         m/z   Intensity
   b1+       397.27       28.2
   y4++      459.37       30.6
   b2+       494.28       10.9
   y5++      516.34       10.6
   y6++      551.50       36.0
   a3+       563.52       11.5
   b3+       591.47        9.9
   y7++      608.79       15.2
   y8++      673.69       13.3
   a4+       692.48       10.3
   b4+       720.59       13.4
   y9++      721.69       14.3
   b9++      737.60       12.3
   a10++     766.75        9.7
   y10++     770.64       75.9
   b5+       835.37       23.9
   b6+       906.46       10.7
   y4+       917.64       20.0
   b7+      1019.55       16.5
```

```
687. MORG-339_phosphopeptide_TMT_8%ACN_032017_OT.6113.6113.3 (sIEEGDGAFGRNPR)

   Ion         m/z   Intensity
   y2+       272.10        6.8
   y5++      300.34        5.8
   b1+       397.20        8.1
   y8++      437.82       50.0
   a2+       482.54        6.0
   y9++      495.29        7.8
   b7++      499.12        7.5
   y10++     523.85       22.4
   y11++     588.34       30.6
   a9++      594.54       14.3
   y5+       599.38       14.6
   b3+       639.32       12.5
   y12++     652.99       13.9
   y13++     709.51       21.3
   b11++     714.91        8.0
   b4+       768.27       12.4
   b12++     772.23       18.2
   b13++     820.56        9.2
   b5+       825.39        7.0
   b6+       940.37       24.3
   b8+      1068.36        6.1
```

```
688. MORG-339_phosphopeptide_TMT_30%ACN_032017_OT.11250.11250.4 (ssSPGkLLGSGLAGGSSRGPPVTPSSEk)

   Ion         m/z   Intensity
   b6++      541.98       20.7
   b7++      598.62       20.2
   b8++      655.21       22.4
   y4+       679.56       18.8
   b10++     727.10       16.8
   y12++     735.72       16.8
   b20+++    764.25       16.7
   y5+       776.52       23.9
   a12++     797.68       24.0
   y14++     807.67       21.2
   b12++     812.29       33.2
   a22+++    821.02       35.0
   b22+++    830.39       48.6
   b23+++    863.69       25.7
   y16++     872.24       31.1
   b14++     876.13       24.3
   y6+       877.35       23.0
   y23+++    881.32       18.7
   y24+++    899.63       18.2
   b15++     904.49       20.1
   b25+++    925.17       20.3
   y25+++    932.67      106.3
   a16++     933.76       30.8
   b16++     948.31       20.3
   y18++     957.35       26.3
   y19++    1000.68       22.0
   y20++    1028.91       24.7
   y21++    1085.80       27.1
   y22++    1142.02       18.8
```

```
689. MORG-339_phosphopeptide_TMT_60%ACN_032017_OT.6455.6455.4 (kWPQQVVQkkPAQETEETSSQEsAEED)

   Ion         m/z   Intensity
   a3+       613.70       20.2
   a14+++    779.29       24.0
   b15+++    822.38       30.3
   a16+++    855.84       25.2
   b16+++    865.09       46.9
   y15++     889.38       72.0
   a17+++    898.68       44.5
   b17+++    908.23       22.7
   a18+++    932.34       42.3
   b18+++    941.82       62.6
   a6+       968.35       24.3
   a20+++    990.68       30.9
   b6+       996.29      152.1
   a11++    1004.35       62.2
   b11++    1018.58       51.7
   b21+++   1042.21       30.6
   y9+      1061.19       22.1
   a22+++   1076.63       49.1
   y25+++   1114.41       27.9
   y10+     1162.46       77.5
```

```
690. MORG-339_phosphopeptide_TMT_18%ACN_032017_OT.12369.12369.3 (skTPPSNLSPIEDAsPTEELR)

   Ion         m/z   Intensity
   b7++      585.81       21.3
   a8++      628.53       15.1
   b8++      642.36       25.2
   y11++     670.51       22.6
   a9++      672.19       13.9
   b9++      686.14       58.7
   y12++     719.10       20.6
   a10++     720.18       16.0
   y6+       744.38       41.5
   b3+       775.51       20.2
   b11++     790.92       23.8
   b12++     855.76       29.9
   y15++     876.34       18.5
   a13++     898.64       26.0
   y7+       911.15       33.8
   b13++     912.84       36.5
   y16++     919.72       30.3
   b14++     948.31       16.2
   y17++     968.18       21.4
   y8+       982.39       28.9
   y18++    1016.85       13.8
   a15++    1017.86       18.0
   y9+      1097.67       19.5
   b18++    1196.06       17.5
   b19++    1260.59       16.5
   b8+      1283.80       13.5
   b20++    1316.94       17.2
   y11+     1339.73       23.3
   b9+      1370.82       24.9
   y12+     1436.63       28.1
```

```
691. MORG-339_phosphopeptide_TMT_20%ACN_032017_OT.8595.8595.3 (sEHSLFQPk)

   Ion         m/z   Intensity
   y3++      301.17        7.2
   b3++      332.20        9.9
   a4++      361.95        6.1
   a1+       369.24        5.2
   y4++      374.49        8.8
   y1+       375.89        9.4
   y5++      431.53        6.3
   b5++      432.31       10.1
   y2+       473.36       37.7
   y6++      474.29       17.1
   a6++      491.90       13.6
   a2+       498.19        7.5
   b6++      505.90       28.5
   b2+       526.46        9.2
   a7++      555.97       10.0
   b7++      569.88       44.0
   y3+       601.45       10.2
   y8++      607.85        6.1
   b3+       663.49       10.3
   y4+       748.58       14.2
   y5+       861.56        6.1
   b5+       863.44        9.5
   b6+      1010.58       11.1
```

```
692. MORG-339_phosphopeptide_TMT_12%ACN_032017_OT.10801.10801.3 (rEsDDsDVDLGSAVRQLR)

   Ion         m/z   Intensity
   b1+       386.34        5.9
   y7++      415.35        9.1
   y3+       416.65        8.2
   y9++      500.53       25.0
   y11++     607.70       17.3
   y12++     665.10        6.0
   y5+       671.32        7.9
   b3+       682.29        6.4
   y6+       742.56        6.9
   y13++     748.60       18.8
   b10++     761.63        7.4
   b11++     789.40       12.0
   b12++     833.43        7.5
   y15++     863.57        8.2
   b5+       912.42       12.4
   a15++     982.46        6.1
   b16++    1060.00        7.4
   b7+      1194.34        8.6
   b9+      1408.52        7.1
```

```
693. MORG-339_phosphopeptide_TMT_10%ACN_032017_OT.23818.23818.3 (kEEsEEsDDDMGFGLFD)

   Ion         m/z   Intensity
   y2+       281.11       16.2
   y4+       451.29       14.7
   y5+       598.36       13.7
   y6+       655.31       24.3
   b9++      720.03        8.9
   a10++     763.63       53.2
   b10++     776.85        7.9
   y13++     778.90        5.6
   y7+       786.33       13.1
   b11++     843.07        7.9
   y8+       901.45        8.5
   a14++     959.03        9.1
   y9+      1016.38        5.8
   a16++    1089.07       22.7
   y10+     1130.99       11.2
   y11+     1298.56        7.9
```

```
694. MORG-339_phosphopeptide_TMT_10%ACN_032017_OT.11051.11051.4 (tDHGAEIVYksPVVsGDtSPR)

   Ion         m/z   Intensity
   y2+       272.16       10.3
   b4++      320.59       12.7
   b5++      356.06       10.9
   a6++      406.92        8.7
   b6++      420.69       17.2
   a7++      463.51       17.9
   b7++      477.34       30.3
   y8++      489.92        9.8
   b10+++    525.21       14.1
   b8++      526.97       20.0
   b3+       583.44       11.6
   b9++      608.19        9.3
   b13+++    646.25       17.9
   y14+++    654.36       10.5
   a14+++    670.04       12.5
   y11++     671.39       16.5
   b14+++    679.25       19.3
   b5+       711.44       40.0
   y6+       712.29       17.4
   b16+++    753.80        8.6
   a17+++    782.48       10.3
   b10++     786.74       13.8
   y19+++    823.70       10.6
   y7+       879.23       13.2
   a12++     904.89        9.2
   y13++     931.80       23.6
   b13++     968.70       10.1
   y14++     981.02       16.4
   y15++    1037.71        9.7
   b8+      1052.75        8.5
   y10+     1174.69       10.0
   y11+     1341.45        8.8
```

```
695. MORG-339_phosphopeptide_TMT_12%ACN_032017_OT.4125.4125.4 (rRHSSLPTEsDEDIAPAQR)

   Ion         m/z   Intensity
   y4++      236.24       23.5
   b2++      271.80       17.9
   y3+       374.24       16.3
   b9+++     432.08       15.0
   a6++      470.03       11.4
   y4+       471.29       72.7
   b6++      483.81       22.4
   b10+++    487.23       15.8
   b11+++    526.06       18.5
   b7++      531.85       10.6
   y14+++    541.21       10.9
   b2+       542.32       49.5
   a12+++    559.27       14.7
   b12+++    569.09       42.2
   y15+++    570.40       23.4
   b8++      582.71       12.6
   y10++     591.45       10.8
   y16+++    598.76       29.7
   b13+++    607.48       71.5
   b14+++    645.14       64.0
   b9++      647.44       15.6
   y11++     655.40       26.9
   a15+++    659.54       15.7
   b15+++    668.82      104.9
   a16+++    691.85       11.8
   b16+++    701.20       12.8
   y12++     705.84       11.4
   b17+++    724.92       13.1
   y13++     754.94       16.4
   a18+++    758.19       10.3
   y7+       770.50       22.1
   b11++     788.63       18.5
   b12++     852.79       19.5
   b5+       853.59       19.2
   y8+       899.48       10.0
   b13++     910.60       38.3
   b14++     967.17       38.1
   b15++    1002.68       38.0
```

```
696. MORG-339_phosphopeptide_TMT_22%ACN_032017_OT.11540.11540.4 (vVDYSQFQEsDDADEDYGRDSGPPAkk)

   Ion         m/z   Intensity
   y5++      499.81       19.9
   y6++      528.64       18.1
   b3+       543.40       33.4
   y11+++    545.67       72.4
   y7++      572.11       52.3
   y12+++    583.90       26.2
   y13+++    627.07       59.6
   y8++      629.74       16.8
   y15+++    689.05       68.0
   b4+       706.57       21.0
   y16+++    727.48       61.1
   a5+       765.72       39.4
   b5+       793.48       15.6
   y11++     817.78       31.0
   y18+++    821.30       20.7
   b20+++    882.42       17.1
   a21+++    901.75       25.8
   y20+++    906.59       20.3
   b21+++    911.32       21.2
   a22+++    920.32       25.9
   b6+       921.58       33.2
   b22+++    930.22       37.6
   y14++     996.99       15.6
   y15++    1032.84       22.9
   a7+      1040.94       16.9
   y6+      1055.67       18.0
   b7+      1068.61       24.1
   y24+++   1082.29       42.5
   y16++    1090.49       26.2
   y17++    1147.87       16.4
   a18++    1173.17       18.6
   b18++    1186.63       16.8
   b8+      1196.62       23.3
   b20++    1322.83       30.2
   b9+      1325.72       17.7
   b22++    1394.75       21.8
   b11+     1607.82       16.5
   b12+     1722.87       17.9
```

```
697. MORG-339_phosphopeptide_TMT_25%ACN_032017_OT.18551.18551.4 (kADTTTPTPTAILAPGsPAsPPGSLEPk)

   Ion         m/z   Intensity
   y2+       473.44       22.9
   y8++      527.22       28.7
   y11++     695.21       20.7
   y4+       715.47       16.0
   y12++     778.03       14.0
   y5+       802.43       12.6
   y13++     806.65       28.6
   b22+++    821.39       21.4
   a23+++    830.59       16.1
   b15++     854.65       27.1
   y14++     855.66       66.8
   y6+       859.45       38.6
   a24+++    860.39       23.6
   a16++     869.66       35.9
   a25+++    898.12       12.3
   a26+++    940.76       13.0
   y16++     947.53       21.1
   b27+++    982.06       16.2
   y27+++    988.34       12.6
   y17++    1003.64       12.5
   a19++    1036.89       12.9
   y18++    1039.14       14.5
   y8+      1053.61       25.0
   b9+      1142.64       12.9
```

```
698. MORG-339_phosphopeptide_TMT_22%ACN_032017_OT.17179.17179.4 (aAGPGNLVLPGsLtASLEcVkEEPPRAAR)

   Ion         m/z   Intensity
   y3+       317.35       11.3
   y6++      334.51       29.2
   b7++      405.80       35.1
   b3+       429.30       14.9
   b8++      455.18       27.3
   b9++      512.17       18.7
   y11+++    514.97       11.7
   y12+++    557.86       13.3
   y13+++    595.48       16.8
   y9++      642.26       19.6
   y6+       667.61       15.4
   b6+       697.42       33.0
   y16+++    708.79       11.4
   y17+++    746.43       16.6
   a20+++    756.59       12.1
   y11++     771.78       12.8
   a7+       782.78       11.4
   y18+++    801.66       12.4
   b7+       810.54       33.2
   b14++     819.76       13.3
   y19+++    820.88       63.8
   y12++     835.61       11.7
   a15++     841.16       17.3
   y20+++    853.45       49.4
   b15++     854.48       20.3
   a21+++    876.05       26.8
   a16++     884.88       25.6
   b8+       909.68       31.8
   a23+++    962.30       14.8
   y15++     972.02       12.7
   a9+       995.06       13.5
   b18++    1019.93       12.4
   b9+      1022.68       43.0
   y16++    1062.23       13.3
   a19++    1085.07       11.9
   y19++    1230.36       56.8
   y20++    1279.39       43.7
   y22++    1385.65       17.5
   b13+     1456.95       32.5
```

```
699. MORG-339_phosphopeptide_TMT_12%ACN_032017_OT.3678.3678.2 (lcsSSSSDTSPR)

   Ion         m/z   Intensity
   y3+       359.35        7.0
   y4+       460.24       19.1
   b2+       503.37        8.6
   y5+       575.34        9.0
   y6+       662.33        6.1
   y7+       749.00       27.2
   b4+       756.88       19.4
   y9+       923.24        9.3
   b8+      1133.52       17.3
   b9+      1234.71        6.9
```

```
700. MORG-339_phosphopeptide_TMT_12%ACN_032017_OT.3828.3828.4 (tEPSGRRGSDELTVPR)

   Ion         m/z   Intensity
   y2+       272.19       12.2
   y3+       371.15       11.3
   a8+++     374.94        6.5
   b9+++     413.23       10.9
   a10+++    442.09       11.2
   y12+++    447.79        9.3
   b10+++    451.66       40.7
   y8++      458.53       10.5
   y4+       472.34       31.9
   a11+++    484.92       37.0
   y9++      487.64       14.5
   b11+++    494.70       44.9
   y14+++    509.40       15.2
   a12+++    523.15       10.3
   b12+++    532.32       26.1
   a7++      533.18       10.2
   b7++      547.05        9.3
   y10++     564.98        6.7
   b13+++    566.06        8.9
   b8++      576.07        9.5
   y5+       585.51       22.8
   b14+++    599.05        9.2
   a9++      605.29        6.7
   b9++      619.04       10.8
   b3+       637.61        6.7
   b10++     677.10        9.0
   a4+       696.51       13.1
   y6+       714.45       23.7
   y13++     715.42       11.3
   b11++     741.56       15.9
   b12++     797.86        7.8
   y7+       829.50        9.9
```

```
701. MORG-339_phosphopeptide_TMT_16%ACN_032017_OT.3146.3146.4 (gSQPGPPVERQsTPRLR)

   Ion         m/z   Intensity
   y4++      271.26       33.5
   b4++      300.31       14.3
   b8+++     317.29        8.9
   b5++      328.67       17.6
   b2+       374.40       10.0
   b6++      377.35       20.5
   b11+++    455.08        8.7
   y11+++    473.38       21.6
   a3+       474.10       15.2
   b8++      475.62       12.6
   a12+++    501.33       15.5
   b3+       502.34       44.2
   y12+++    506.17       64.1
   b12+++    510.30       27.5
   y13+++    524.66       29.2
   a9++      525.53       31.3
   a13+++    534.69       36.8
   y14+++    557.37       33.7
   a4+       571.06       13.2
   b4+       599.10       11.5
   y9++      611.49       10.7
   b10++     617.69        9.9
   a15+++    619.19        9.5
   y5+       642.29       12.4
   b5+       656.39       36.6
   a16+++    657.35       21.4
   y10++     661.07       28.6
   b16+++    666.28       14.3
   y11++     710.04       66.3
   y12++     758.59       63.7
   y13++     786.86       19.4
   y14++     835.64       20.6
```

```
702. MORG-339_phosphopeptide_TMT_18%ACN_032017_OT.15547.15547.3 (aTtPPNQGRPDsPVYANLQELk)

   Ion         m/z   Intensity
   y2+       489.45       10.8
   b3+       583.26       15.7
   b9++      617.03       17.6
   y3+       618.55       11.2
   a10++     651.48        9.8
   b10++     665.22       12.1
   y10++     702.62       11.9
   a11++     708.53       17.3
   b11++     723.07       38.1
   y5+       859.69       10.0
   a6+       863.48       16.1
   y13++     891.46       12.6
   b14++     904.65       15.5
   y14++     970.24       19.5
   a15++     971.71       14.3
   y6+       973.25      104.8
   b15++     986.21       26.9
   a7+       991.44       10.2
   b7+      1019.20        9.9
   y7+      1044.55       21.0
   y16++    1062.21       12.8
   b17++    1078.62       12.3
   y17++    1119.70       15.9
   a18++    1120.78       10.2
   b18++    1134.79       10.6
   y18++    1167.94       66.8
   b19++    1199.39       14.0
   y8+      1207.87       27.6
   y19++    1216.89       76.7
   b20++    1263.78       14.2
   y10+     1403.82       33.9
   y11+     1570.89       22.9
   y13+     1782.91       10.9
```

```
703. MORG-339_phosphopeptide_TMT_20%ACN_032017_OT.11946.11946.4 (nRPPLAAGANSkGPPDFssDEEREPTPVLGSGASVGR)

   Ion         m/z   Intensity
   b16+++    668.28       14.6
   y15++     742.27       18.5
   y9+       803.57       11.0
   a12++     804.63       11.3
   y16++     806.70       19.4
   b5+       807.37       14.0
   b12++     818.54       15.2
   a19+++    819.20       17.1
   b13++     846.93       14.9
   y17++     871.32       31.2
   y18++     928.92       15.3
   y11+      999.53       17.4
   b16++    1001.85       44.5
   b23+++   1005.25       15.1
   b8+      1006.32       15.1
   y19++    1011.91       10.9
   a24+++   1038.53       15.6
   b24+++   1047.90       19.1
   y27+++   1049.91       21.2
   a17++    1061.32      189.4
   a25+++   1070.75       17.0
   y12+     1100.13       11.0
   y29+++   1111.88       14.3
   a27+++   1136.49       14.7
   a18++    1144.33       14.8
   b27+++   1146.50       13.5
   y31+++   1154.03       12.4
   y21++    1169.38       17.5
   a28+++   1170.30       13.2
   b28+++   1179.37       12.5
   y13+     1197.73       18.2
   y33+++   1215.91       14.8
   a30+++   1226.33       14.4
   a19++    1227.90       11.0
   y34+++   1248.03       13.3
   a11+     1250.88       14.6
   a32+++   1274.21       12.3
   y23++    1275.02       11.4
   b20++    1299.96       21.8
   b33+++   1307.73       11.4
   y24++    1323.80       13.5
   y14+     1326.75       13.9
   y36+++   1331.97       12.3
   b21++    1364.38       12.1
```

```
704. MORG-339_phosphopeptide_TMT_25%ACN_032017_OT.23326.23326.2 (fVQWQSsI)

   Ion         m/z   Intensity
   y2+       299.19        9.1
   b1+       377.28        6.8
   y3+       386.18       20.9
   b2+       476.33       18.7
   y4+       514.32       15.3
   b3+       604.33       36.0
   y5+       700.29       29.3
   b4+       790.50       24.6
   y6+       828.42       24.2
   b5+       918.53       35.1
   y7+       927.52        7.1
   b6+      1005.59       21.2
```

```
705. MORG-339_phosphopeptide_TMT_8%ACN_032017_OT.3837.3837.3 (dTGENtEGkkDEDAGQDE)

   Ion         m/z   Intensity
   b1+       345.29        6.7
   y5+       519.26        6.0
   b4+       632.38        9.1
   y7+       763.37        8.1
   y10++     796.97        6.4
   y11++     825.77       17.9
   y8+       878.27        7.1
   y12++     889.76       11.9
   a11++     958.47       11.5
   b11++     971.93       11.3
   y13++     980.79        5.6
   y14++    1037.66        8.0
   b7+      1056.47        6.9
   b13++    1094.29       14.9
   a14++    1115.64        5.4
   y16++    1130.96        6.7
   y17++    1181.00       10.4
   y9+      1235.43        6.2
```

```
706. MORG-339_phosphopeptide_TMT_30%ACN_032017_OT.16371.16371.4 (kLGGDVSPtQIDVSQFGSFkEDTk)

   Ion         m/z   Intensity
   y2+       477.38       23.4
   y10+++    549.45       65.8
   y8++      685.38       13.7
   b11++     703.51       20.2
   y9++      758.83       22.9
   b12++     761.26       16.1
   a6+       771.26       14.1
   a13++     796.01       27.4
   y10++     823.18       24.5
   b14++     853.98       13.5
   y11++     866.57       41.5
   b20+++    877.55       31.4
   y19+++    884.34       32.8
   b7+       887.00       15.0
   y12++     916.25       38.5
   b21+++    920.14       15.6
   y13++     973.74       44.1
   b8+       983.82       12.7
   b23+++    992.15       14.7
   y23+++    998.59       14.2
   y14++    1029.93       31.8
   y15++    1094.33       17.7
   y16++    1184.39       22.9
   y18++    1277.12       20.1
   y19++    1326.47       26.6
   y9+      1516.80       12.6
```

```
707. MORG-339_phosphopeptide_TMT_20%ACN_032017_OT.8449.8449.4 (vNHEPEPASGASPGAtIPkSPSQLR)

   Ion         m/z   Intensity
   b6++      468.37       14.6
   b7++      517.04       12.1
   a3+       552.46       10.6
   y5+       600.37       12.8
   b10++     624.00       10.2
   b11++     659.74       10.7
   y18+++    678.65       17.9
   a4+       681.31       11.8
   y6+       687.55        9.8
   b12++     703.57       18.4
   b4+       709.48       24.6
   y19+++    710.89       25.9
   y10++     718.42       15.2
   a13++     737.42       27.5
   b13++     751.58       27.2
   y20+++    754.22       25.4
   b14++     780.42       11.6
   y12++     782.57       14.1
   y21+++    786.49       13.7
   b20+++    822.41       14.8
   y13++     831.10       21.1
   y23+++    874.63       15.4
   y15++     910.47        9.4
   b6+       935.47       15.7
   y16++     938.76       16.1
   y17++     981.86       13.9
   y18++    1017.33       12.6
   y19++    1066.35       21.4
   y21++    1179.48        9.5
```

```
708. MORG-339_phosphopeptide_TMT_20%ACN_032017_OT.12207.12207.3 (sETLVNAQQTPLGtPk)

   Ion         m/z   Intensity
   y5++      412.25       14.1
   y6++      461.43       47.9
   y2+       473.30       40.2
   y7++      512.01       27.0
   b8++      536.95       11.6
   y8++      576.08       15.9
   b9++      600.83       22.3
   a10++     637.35       10.9
   y9++      640.08       13.9
   b10++     651.56       66.8
   y3+       654.35       15.1
   b4+       660.47       12.6
   y10++     675.54       14.6
   y4+       711.56       31.5
   y11++     731.93       33.6
   b12++     756.75       14.1
   b5+       759.53       19.0
   b13++     784.94       16.2
   y5+       824.83       11.3
   a14++     861.05       12.9
   b6+       873.80       15.9
   b14++     875.77        9.5
   y6+       921.52       46.7
   b15++     923.49       14.7
   b7+       944.57       13.7
   y7+      1022.57       15.2
   b8+      1072.59       18.5
   y8+      1150.60       10.6
   b9+      1200.74       19.2
   b10+     1301.69       36.4
```

```
709. MORG-339_phosphopeptide_TMT_25%ACN_032017_OT.8713.8713.4 (nLAkPGVTsTSDSEDEDDQEGEkk)

   Ion         m/z   Intensity
   b4++      443.52       23.3
   y4++      460.46       18.9
   y6++      589.13       42.9
   a13+++    589.97       20.9
   y7++      646.60       37.4
   b9++      704.36       19.2
   y9++      768.63       33.2
   y10++     825.66       21.8
   a12++     841.93       22.5
   b19+++    842.91       23.7
   y18+++    846.20       25.9
   b12++     855.33       42.5
   a4+       857.16       29.3
   y3+       862.56       43.6
   a20+++    877.33       39.0
   a13++     884.57      141.5
   b4+       885.79       46.3
   b20+++    886.58       32.1
   b21+++    905.74       17.5
   y4+       919.52       24.1
   y12++     934.18       43.5
   b14++     963.81       19.6
   b5+       983.12       20.9
   y13++     991.82       33.1
   b15++    1020.77       18.6
   y14++    1034.62       21.6
   y5+      1048.53       22.0
   b23+++   1067.82       28.3
   y23+++   1078.34       20.6
   a17++    1129.41       20.8
   b17++    1142.85       19.8
   b18++    1200.54       19.7
   y19++    1297.81       21.3
```

```
710. MORG-339_phosphopeptide_TMT_30%ACN_032017_OT.17625.17625.4 (sTQLLQAAAAEASLNkSkSATttPsGSPR)

   Ion         m/z   Intensity
   b7++      486.48       13.7
   y13+++    582.37       19.0
   b4+       659.59       13.4
   y6+       680.29       33.4
   b18+++    834.35       20.0
   a15++     835.50       15.5
   b15++     849.63       17.1
   a6+       872.75       15.2
   y13++     873.57       17.4
   a20+++    877.81       26.2
   b6+       900.68       14.0
   a21+++    911.12       14.7
   b21+++    920.27       16.2
   b7+       971.43       17.1
   y24+++   1010.37       13.1
   b16++    1028.65       24.9
   b8+      1042.13       15.2
   b17++    1072.56       12.5
   b24+++   1073.38       11.7
   y26+++   1086.36       14.7
   b9+      1113.43       11.9
   a26+++   1139.17       23.1
```

```
711. MORG-339_phosphopeptide_TMT_22%ACN_032017_OT.8511.8511.4 (qVDTEEAGmVTAATASNVkAsPkR)

   Ion         m/z   Intensity
   b2+       457.31       16.1
   b7++      501.56       19.5
   b8++      530.46       28.7
   b3+       572.40       22.1
   b9++      604.01       33.4
   y13+++    613.75       16.2
   y14+++    647.67       19.1
   b10++     653.46       16.2
   y15+++    680.66       17.3
   b11++     703.94       26.3
   b12++     739.62       21.1
   y17+++    748.52       24.7
   a13++     761.31       21.7
   y9++      762.98       34.2
   y18+++    771.77       88.6
   b13++     774.83       18.5
   a19+++    784.03       30.5
   b19+++    793.64       58.6
   y4+       796.25      103.8
   y10++     798.12       42.7
   b5+       802.39       33.1
   y11++     849.23       33.5
   b15++     861.14       17.3
   y5+       867.88       21.5
   b21+++    872.74       17.6
   y12++     884.41       37.2
   y21+++    892.18       28.7
   a6+       903.04       18.2
   b16++     904.23       17.6
   y13++     920.27       36.6
   y22+++    929.77       18.0
   b6+       931.50       27.8
   b17++     961.78       20.9
   y14++     970.92       51.1
   b7+      1002.65       26.9
   b18++    1011.29       17.4
   y15++    1020.32       24.4
   b8+      1059.56       24.8
   b9+      1206.57       18.0
```

```
712. MORG-339_phosphopeptide_TMT_4%ACN_032017_OT.6833.6833.2 (sYIETETESR)

   Ion         m/z   Intensity
   y2+       262.12        4.4
   b1+       397.09       10.6
   y4+       492.28       10.9
   b2+       560.53        6.4
   y5+       621.11        7.2
   a3+       645.80        4.8
   b3+       673.48       10.0
   y6+       722.61       29.2
   a4+       774.34        5.2
   b4+       802.45       11.7
   y7+       851.39        8.7
   b5+       903.68        5.6
   y8+       964.07        9.7
   b6+      1032.51        4.8
   y9+      1127.55       11.1
   b8+      1262.67        9.8
```

```
713. MORG-339_phosphopeptide_TMT_25%ACN_032017_OT.20516.20516.4 (esASPTIPNLDLLEAHTkEALTk)

   Ion         m/z   Intensity
   b10+++    440.39       12.3
   y2+       477.43       17.8
   b7++      498.68       12.4
   y6++      574.63       19.0
   y3+       590.48       13.6
   y7++      624.89       13.3
   y4+       661.54       14.0
   b16+++    666.44       13.5
   y14+++    680.94       15.8
   y8++      693.33       21.4
   a11++     704.06       11.1
   y9++      728.77       17.6
   y16+++    751.50       90.5
   a5+       753.28       11.4
   a12++     760.25       13.8
   b12++     774.10       19.2
   y17+++    789.25       26.8
   y5+       790.52       22.3
   y10++     793.89       45.8
   y11++     849.84       17.5
   y19+++    855.16       44.0
   b19+++    862.58       14.0
   b20+++    886.55       19.7
   y12++     906.86       22.0
   b21+++    924.20       29.5
   y13++     964.31       25.9
   a7+       967.52       12.5
   b7+       995.48       24.6
   y14++    1020.55       15.0
   y15++    1077.58       11.6
   y16++    1126.48       46.2
```

```
714. MORG-339_phosphopeptide_TMT_8%ACN_032017_OT.10354.10354.3 (sFsIsPSRR)

   Ion         m/z   Intensity
   y3++      209.59        3.6
   a2++      258.17       10.7
   a3++      341.65       16.1
   b1+       397.29        5.7
   a4++      398.33       10.9
   b4++      413.03        4.0
   a5++      481.70        7.0
   b5++      496.00        5.3
   y4+       515.32        4.6
   y8++      555.23        4.8
   b7++      588.34        6.2
   b8++      666.38        6.9
   a3+       683.49        3.4
   b3+       711.38        4.3
   y6+       795.28        7.1
   b4+       824.46        5.2
```

```
715. MORG-339_phosphopeptide_TMT_16%ACN_032017_OT.1245.1245.4 (rSGkNSQEDsEDSEEk)

   Ion         m/z   Intensity
   y1+       376.39       18.1
   b8+++     449.18       13.8
   b9+++     487.76       42.4
   y2+       505.36       22.4
   b10+++    543.22        9.5
   b11+++    586.36       16.9
   b7++      608.85       19.7
   a12+++    615.24       41.6
   b12+++    624.56      157.2
   y3+       634.37       31.8
   b8++      673.23        9.8
   a14+++    687.47        9.3
   y9++      688.90        9.5
   b14+++    696.45       16.5
   y14+++    707.10       24.7
   y4+       721.45       28.2
   a15+++    730.28       10.2
   b9++      731.06       16.1
   b15+++    739.79       27.1
   y10++     752.35       10.8
   b10++     813.95       10.5
   y5+       836.55       15.4
   a13++     965.50       12.6
   y7+      1132.47       13.2
```

```
716. MORG-339_phosphopeptide_TMT_25%ACN_032017_OT.20039.20039.3 (lGGDVSPtQIDVSQFGSFkEDTk)

   Ion         m/z   Intensity
   y2+       477.33       22.1
   b4+       572.35       29.3
   b5+       671.71       16.4
   y9++      759.20       15.6
   y11++     866.77       25.1
   y12++     916.00       24.6
   b16++     956.41       12.9
   y13++     973.94       22.4
   a17++     985.55       20.6
   a8+      1008.79       12.9
   b8+      1036.34       49.0
   a18++    1058.89       15.4
   y16++    1184.78       27.9
   a19++    1237.91       14.6
   y18++    1276.94       16.3
   b10+     1277.74       35.6
   y19++    1326.07       28.7
   a11+     1365.12       13.8
   y8+      1369.78       17.0
   b21++    1373.95       26.8
   b11+     1392.99       17.4
   y9+      1516.96       17.4
```

```
717. MORG-339_phosphopeptide_TMT_25%ACN_032017_OT.18296.18296.4 (aEsPVkEkAVEEVITISk)

   Ion         m/z   Intensity
   a1+       273.22       12.7
   y1+       376.27       18.9
   y2+       463.36       32.4
   y3+       576.54       21.1
   b10+++    603.23       19.4
   b11+++    646.28       15.3
   y4+       677.48       30.5
   a12+++    679.41       11.9
   b12+++    689.43       33.8
   b4+       694.56       18.2
   a13+++    712.45       24.1
   y13+++    721.44       45.0
   b13+++    722.29       40.2
   a14+++    750.90       12.0
   b14+++    759.98       37.0
   a5+       765.84       10.7
   y15+++    786.71       47.8
   y5+       790.58       18.6
   b15+++    793.60       28.6
   a16+++    822.17       13.8
   b16+++    831.38       20.8
   y11++     837.61       13.2
   y16+++    842.43       12.8
   b9++      854.49       36.2
   b17+++    860.31       30.4
   y17+++    885.27       10.6
   y6+       889.68       11.2
   a10++     890.37       12.6
   y12++     902.08       13.1
   b13++    1082.59       14.3
   b14++    1139.55       13.1
```

```
718. MORG-339_phosphopeptide_TMT_30%ACN_032017_OT.12865.12865.4 (kPPAPPsPVQSQsPSTNWSPAVPAkk)

   Ion         m/z   Intensity
   y4++      451.66       24.7
   y7++      584.86       29.3
   a14+++    587.48       38.8
   y8++      628.41       19.0
   b5+       720.87       16.5
   y2+       734.03       14.1
   y10++     778.55       15.5
   a19+++    779.26       23.6
   b6+       817.79       14.8
   y11++     829.46       19.9
   y19+++    850.14       15.4
   a22+++    867.95       20.4
   y12++     872.81       16.0
   b22+++    877.24       23.8
   a23+++    900.86       22.7
   y4+       901.67       40.9
   y20+++    905.77       21.2
   y13++     921.24       76.1
   a15++     923.53       54.2
   a24+++    924.53       20.5
   a7+       956.17       20.1
   y22+++    970.03       32.6
   b16++     988.20       15.5
   y5+      1000.95       15.8
   y24+++   1026.00       26.0
   a17++    1031.57       17.8
   a25+++   1043.13       19.5
   b17++    1045.15       19.1
   y25+++   1058.87       41.7
   y15++    1068.81       17.7
   y6+      1071.77       17.0
   y16++    1112.43       23.4
   a19++    1167.86       19.8
   y7+      1168.82       33.5
   y17++    1176.23       19.6
   y18++    1225.46       21.2
   y19++    1274.46       27.9
   y22++    1455.01       16.2
```

```
719. MORG-339_phosphopeptide_TMT_6%ACN_032017_OT.8389.8389.3 (eEVAsEPEEAASPTTPk)

   Ion         m/z   Intensity
   y2++      237.23        6.1
   y1+       376.36        8.5
   y5++      386.91       17.5
   y6++      430.38       11.5
   y2+       473.33       31.3
   y8++      501.44        6.5
   y3+       574.44       13.1
   b3+       587.35        6.9
   b9++      655.62       13.3
   b4+       658.32       10.1
   y4+       675.50       13.6
   a10++     676.53        9.1
   y11++     678.80        6.2
   b10++     691.03       27.0
   a11++     712.52       10.4
   b11++     726.53       32.9
   y12++     743.36        8.0
   a12++     756.29       21.4
   y6+       859.67       16.3
   y14++     861.97        7.7
   b14++     869.31        9.0
   a15++     905.40        8.8
   b15++     919.74       17.0
   y7+       930.44        6.9
   b6+       954.43       14.8
   b8+      1180.42        6.9
   b9+      1309.59        7.4
   b10+     1380.64        7.9
   b11+     1451.63       11.3
   b12+     1538.66       12.4
```

```
720. MORG-339_phosphopeptide_TMT_4%ACN_032017_OT.18229.18229.2 (sEsDLEEAEPVVIPR)

   Ion         m/z   Intensity
   y3+       385.35       42.6
   b1+       397.29       12.0
   y4+       484.42       30.4
   b2+       526.30       17.6
   y5+       583.54       12.0
   y6+       680.53      101.5
   b4+       808.32       36.0
   y7+       809.59       55.3
   y8+       880.61       52.6
   a5+       893.41       12.8
   b5+       921.39       33.6
   y9+      1009.55       59.8
   b6+      1050.41       41.8
   y10+     1138.64       50.1
   b7+      1179.43       43.4
   b8+      1250.47       47.8
   y11+     1251.68       68.9
   a9+      1351.66       13.0
   y12+     1366.75       42.0
   b9+      1379.49       91.8
   y13+     1533.72       27.3
   b11+     1575.61       36.7
   y14+     1662.88       13.3
   b12+     1674.69       58.3
   b13+     1787.81       59.9
```

```
721. MORG-339_phosphopeptide_TMT_25%ACN_032017_OT.14753.14753.4 (eATPYVkkGSPVSEIGWETPPPEsPR)

   Ion         m/z   Intensity
   y6++      381.52       18.5
   b2+       430.18       18.0
   b10+++    583.97       19.4
   b12+++    649.30       23.5
   b13+++    678.52       21.4
   b14+++    721.49       42.8
   a15+++    749.83       18.0
   b15+++    759.20       54.5
   y6+       762.45       37.5
   a5+       763.50       30.1
   b16+++    777.97       54.1
   y19+++    786.81       19.8
   b5+       791.82       20.4
   a17+++    830.46       27.7
   b9++      831.28       29.8
   b17+++    840.12       62.7
   y7+       859.37       64.4
   a10++     861.50       20.5
   a6+       862.78       44.2
   a18+++    873.59       75.2
   b10++     874.55       54.0
   y15++     881.22       43.1
   b18+++    882.39       74.0
   b6+       890.71       32.9
   b19+++    916.92       89.3
   y21+++    938.69       20.7
   b20+++    949.11       34.0
   a12++     959.37       22.4
   y8+       960.44       25.7
   b12++     973.41       24.8
   b21+++    981.00       21.1
   b13++    1016.58       22.1
   a23+++   1047.39       23.3
   y24+++   1059.62       24.4
   b14++    1081.00       27.9
   b15++    1137.53       28.5
   b16++    1166.58       37.3
   a17++    1245.39       18.3
   b17++    1259.55       23.6
   y10+     1275.85       19.3
   b18++    1323.64       24.1
   b19++    1374.58       42.4
```

```
722. MORG-339_phosphopeptide_TMT_16%ACN_032017_OT.16555.16555.3 (tLSVAAAFNEDEDsEPEEmPPEAk)

   Ion         m/z   Intensity
   y1+       376.58       11.6
   y5++      385.86       28.1
   y2+       447.35       13.2
   a3+       503.23       13.4
   y3+       576.52       12.4
   y9++      636.99       39.5
   a5+       673.54       15.3
   b5+       701.39       13.3
   b12++     739.40       12.0
   y5+       770.49       43.5
   b6+       772.51       17.4
   y12++     842.47       10.8
   b7+       843.54       14.2
   b14++     880.85       12.3
   y13++     907.36       12.0
   y6+       917.50       19.8
   y14++     964.60       17.7
   b8+       990.62       17.9
   b16++     993.30       17.2
   y15++    1028.92       21.1
   a17++    1043.54       13.7
   y16++    1085.89       14.3
   b9+      1104.65       23.0
   b18++    1122.59       12.3
   b19++    1195.84       16.3
   a10+     1205.22       11.2
   a20++    1230.09       11.3
   b10+     1233.76       14.5
   b23++    1393.29       10.9
   b15+     1888.74       16.4
```

```
723. MORG-339_phosphopeptide_TMT_16%ACN_032017_OT.12829.12829.3 (eSSPLySPGFSDSTSAAk)

   Ion         m/z   Intensity
   y1+       376.43       26.8
   b2+       446.57       10.5
   y2+       447.27       43.2
   y3+       518.40       26.0
   y4+       605.45       24.3
   y11++     649.07       40.6
   b10++     688.30       12.1
   y12++     692.66       56.5
   a5+       715.24       15.3
   b11++     731.76       19.8
   b5+       743.47       11.7
   a12++     774.58       28.8
   b13++     832.44       21.5
   a14++     869.37       10.8
   b14++     882.69       17.0
   y7+       908.33       25.2
   y15++     919.30       26.3
   b15++     926.57       20.1
   y16++     962.39       28.3
   b6+       986.42       61.3
   y8+       995.57       17.1
   b17++     997.71       43.9
   b7+      1073.50       33.9
   b10+     1374.71       13.1
   b11+     1461.63       10.3
```

```
724. MORG-339_phosphopeptide_TMT_14%ACN_032017_OT.8280.8280.4 (gAkGkQADVADQQTTELPAENGETENQSPAsEEEk)

   Ion         m/z   Intensity
   y1+       376.31       22.1
   y6++      500.98       25.7
   a9+++     505.37       26.4
   y7++      550.03       21.7
   b11+++    577.16       22.0
   y3+       634.47       24.7
   b7++      665.11       24.8
   y10++     714.58       27.1
   b8++      722.73       43.3
   a16+++    763.48       28.2
   b4+       772.21       21.9
   b16+++    773.13       53.4
   b10++     807.75       23.6
   b17+++    810.76       73.8
   y12++     829.47       27.8
   b11++     865.27       69.8
   b19+++    866.17       38.1
   y21+++    866.97       23.9
   b20+++    909.76       38.4
   y14++     922.71       30.5
   b12++     929.15       26.8
   y5+       930.48       36.2
   b13++     993.22       37.9
   a23+++   1000.56       53.2
   y6+      1001.56       40.4
   y16++    1043.97       51.5
   a15++    1079.82       49.3
   y27+++   1080.98       43.3
   b15++    1093.87       26.9
   y7+      1098.47       49.1
   y28+++   1119.55       36.8
   b26+++   1124.56       30.8
   b5+      1129.83       38.8
   y29+++   1142.63       24.4
   a16++    1144.80       27.7
   b16++    1158.95       63.5
   y19++    1184.74       46.0
   y8+      1185.47       45.2
   a28+++   1186.54       28.3
   b17++    1215.43      123.9
   y21++    1299.42       50.2
   a7+      1300.42       29.4
   b32+++   1350.88       23.9
   b20++    1364.00       62.1
   y10+     1427.60       30.0
   b23++    1514.14       25.8
   a9+      1515.42       36.9
   y11+     1556.27       26.1
   y27++    1620.89       21.5
   b25++    1629.19       33.2
   y14+     1843.81       31.6
```

```
725. MORG-339_phosphopeptide_TMT_22%ACN_032017_OT.11124.11124.4 (ekEEPPSPVEtTPPRSLLEk)

   Ion         m/z   Intensity
   a4++      473.61       20.3
   y6++      487.92       34.5
   y8++      585.20       20.8
   y3+       618.54       14.2
   y14+++    621.45       13.4
   y15+++    653.84       19.1
   y16+++    686.36       39.9
   a2+       688.18       14.8
   a9++      712.43       20.9
   b9++      726.13       26.0
   a15+++    728.88       22.8
   y17+++    729.71       48.3
   y4+       731.95       31.9
   b15+++    738.06       34.9
   y18+++    772.84       14.2
   b10++     790.91       21.1
   a17+++    795.85       12.7
   b17+++    805.38       17.1
   a3+       817.70       13.7
   y5+       818.44       13.4
   a18+++    833.62       16.3
   y12++     840.12       13.2
   b3+       845.22       18.0
   b11++     880.95       14.7
   y13++     888.50       17.3
   y14++     932.02       15.4
   y6+       974.65       21.1
   y15++     980.42       26.8
   y16++    1029.30       25.4
```

```
726. MORG-339_phosphopeptide_TMT_20%ACN_032017_OT.6333.6333.3 (rLsGGAVPSASMTR)

   Ion         m/z   Intensity
   y4++      247.20       15.2
   b3++      333.77        9.9
   y7++      375.36       16.8
   b5++      390.86       13.4
   a6++      412.42       16.8
   b6++      426.49       30.5
   a7++      461.84       26.8
   b7++      475.88       43.1
   y10++     489.22       13.0
   y4+       494.50        9.8
   a8++      510.15       10.7
   y11++     517.67       14.6
   b8++      524.61       14.8
   a9++      553.69       15.2
   b10++     603.71       10.6
   a3+       638.28        9.9
   y6+       652.31        8.4
   y7+       749.38       42.3
   b13++     763.15        8.1
   y8+       848.42       20.4
   y9+       919.49        9.2
   b7+       950.59        9.6
   y10+      976.56        8.2
   y11+     1033.65        8.8
```

```
727. MORG-339_phosphopeptide_TMT_4%ACN_032017_OT.6144.6144.3 (aSRVPssDEEVVEEPQSRR)

   Ion         m/z   Intensity
   b3++      272.67        6.2
   b4++      322.19       10.6
   y8++      500.82        6.8
   y4+       546.19        6.2
   a4+       615.24       14.4
   b4+       643.51       20.5
   y11++     679.48       21.1
   b10++     724.45        6.5
   y12++     736.98       11.3
   b11++     774.21       11.8
   a12++     809.43       13.4
   y13++     820.76        9.4
   b12++     823.02       17.9
   b13++     887.97        7.1
   y14++     903.65       19.3
   b14++     952.52        9.1
   b15++    1000.66        7.7
   y16++    1002.32        9.7
   y17++    1079.57        6.4
   a8+      1161.45        9.1
   b18++    1186.21        6.9
   b8+      1189.42       10.6
   b9+      1318.62        6.9
   b10+     1447.62        6.7
   b13+     1774.79        6.5
```

```
728. MORG-339_phosphopeptide_TMT_22%ACN_032017_OT.9516.9516.3 (rQsFASLALR)

   Ion         m/z   Intensity
   b2++      257.88        7.6
   y2+       288.25        9.6
   b3++      341.20       11.5
   y3+       359.62        9.4
   b1+       386.66        5.1
   a4++      401.06        9.1
   b4++      415.03       15.3
   a5++      436.45       10.7
   b5++      450.15       20.6
   y8++      472.54       11.9
   y9++      536.82        8.6
   y5+       559.39       22.5
   b8++      585.75       10.2
   a9++      628.07        9.6
   y6+       630.46       16.8
   b9++      642.47        9.9
   y7+       777.53       18.7
```

```
729. MORG-339_phosphopeptide_TMT_20%ACN_032017_OT.16126.16126.3 (rIDFIPVsPAPsPTR)

   Ion         m/z   Intensity
   b3++      307.74       14.0
   y3+       373.23       16.2
   b4++      381.12       14.2
   y7++      403.41        8.7
   a5++      423.92       10.3
   b5++      437.90       29.3
   a2+       471.38        7.1
   y8++      486.47        8.6
   a7++      521.75        8.0
   y9++      536.08       14.5
   y10++     584.41       11.7
   a8++      605.16        9.0
   b3+       614.37        6.9
   b8++      619.44        8.5
   y5+       637.47       18.0
   y11++     641.48        8.9
   a9++      654.24       21.6
   b10++     703.63       26.6
   y6+       708.33       12.9
   a11++     737.94       21.5
   b4+       761.37        7.9
   y7+       805.46       24.4
   b5+       874.49       16.8
   b7+      1070.62        9.5
   y9+      1071.59       10.4
   y10+     1168.63       14.2
   y12+     1428.82        9.2
```

```
730. MORG-339_phosphopeptide_TMT_25%ACN_032017_OT.17199.17199.4 (sLQtLPTDSSAFDTDTFcPPRPkPLAR)

   Ion         m/z   Intensity
   y12+++    557.18       17.6
   y14+++    629.32       17.2
   y9++      630.74       18.1
   a16+++    654.33       15.0
   a11++     692.11       14.8
   a4+       711.66       18.0
   y16+++    716.83       24.6
   b4+       739.76       17.6
   y17+++    740.49       19.8
   a18+++    756.51       21.3
   b12++     778.98       14.2
   y11++     784.87       22.1
   b19+++    798.70       18.2
   y5+       813.88       16.0
   a20+++    821.40       14.4
   a13++     822.79       19.3
   a5+       824.56       20.3
   b20+++    830.58       21.8
   y12++     834.51       52.6
   b13++     836.42       19.9
   b5+       852.14       60.6
   a21+++    873.71       38.2
   y22+++    902.77       34.2
   a22+++    905.85       13.8
   a6+       921.10       14.6
   y23+++    940.62       22.3
   y14++     943.00       23.4
   a16++     981.74       13.6
   y15++    1000.47       36.9
   a7+      1022.20       39.7
   a17++    1054.74       19.0
   y16++    1074.03       37.5
   y17++    1109.84       33.9
   a8+      1137.86       16.0
   y18++    1153.36       26.2
   y19++    1196.78       28.3
   b19++    1197.76       14.3
   y21++    1305.14       18.0
   y22++    1353.48       31.7
```

```
731. MORG-339_phosphopeptide_TMT_25%ACN_032017_OT.19069.19069.3 (dVTNLTVGGFTPMsPR)

   Ion         m/z   Intensity
   y2+       272.18        9.8
   b3++      273.27        9.9
   a1+       317.33       17.6
   b6++      437.56       20.9
   y3+       439.29       11.0
   b7++      486.75       22.6
   b8++      515.29       24.2
   b9++      544.03       33.3
   b3+       544.90       18.8
   y10++     564.68       13.3
   y4+       570.39       19.1
   a10++     603.40       16.0
   y11++     615.71       12.0
   b10++     617.63       42.4
   a4+       630.89       14.4
   a11++     653.92       25.7
   b4+       659.01       42.2
   y13++     728.72       10.4
   y6+       768.33       27.1
   b5+       772.43       19.5
   b13++     782.20       12.4
   b6+       873.57       18.7
   y7+       915.42       20.7
   b7+       972.46       31.2
   b8+      1029.52       36.4
   b9+      1086.54       19.5
   y10+     1128.57       10.6
```

```
732. MORG-339_phosphopeptide_TMT_10%ACN_032017_OT.16253.16253.4 (iEDsEPHIPLIDDtDAEDDAPTkR)

   Ion         m/z   Intensity
   a8+++     401.40       24.6
   y6++      459.02       12.4
   y7++      516.60       10.8
   a7++      545.48        9.9
   a3+       559.41       21.5
   b3+       587.34       14.2
   a8++      601.81       14.0
   y9++      616.08       26.2
   y15+++    661.67       10.0
   y10++     674.05       19.1
   y16+++    694.59       20.1
   y17+++    732.43       13.7
   a17+++    750.91        9.9
   a11++     763.45        9.9
   y11++     764.49       18.4
   y18+++    777.70       23.5
   a18+++    789.84       13.7
   b18+++    798.68       16.0
   y5+       801.64       24.2
   y19+++    810.50       50.0
   y13++     879.87       12.1
   b5+       883.44       13.2
   b13++     892.51       13.6
   y21+++    909.01       12.9
   y6+       916.61       10.5
   a22+++    917.65       11.4
   y14++     936.01       15.8
   b14++     983.33       12.1
   y15++     992.29       36.7
   y16++    1041.25       64.9
   b18++    1198.13       12.3
   b8+      1230.58        9.4
   y9+      1231.58       10.2
   b19++    1255.74       19.1
```

```
733. MORG-339_phosphopeptide_TMT_10%ACN_032017_OT.15452.15452.2 (sAYFSDEEELsD)

   Ion         m/z   Intensity
   y2+       301.07       13.7
   b1+       317.12       14.7
   y3+       414.20       14.2
   y4+       543.17       15.5
   b3+       551.35       24.5
   y5+       672.20       17.6
   b4+       698.45       20.3
   b5+       785.51       17.1
   b6+       900.50       11.2
   y7+       916.40       12.1
   y8+      1003.37       24.7
   b7+      1029.57       25.8
   y9+      1150.38       27.7
   b8+      1158.49       24.0
   b9+      1287.57       30.6
   y10+     1313.48       12.3
   b10+     1400.73       33.5
```

```
734. MORG-339_phosphopeptide_TMT_30%ACN_032017_OT.13006.13006.4 (sVPTVDsGNEDDDSSFkIkTVAQk)

   Ion         m/z   Intensity
   b1+       317.24       14.5
   b2+       416.37       19.5
   a14+++    567.01       15.3
   y3+       575.46       13.9
   y11+++    642.18       46.0
   y4+       674.45       15.0
   y12+++    680.62       39.0
   b5+       713.25       16.2
   y13+++    719.05       30.0
   y14+++    757.44       21.8
   y5+       775.65       16.7
   a6+       800.50       14.6
   a18+++    802.41       14.1
   b13++     821.29       14.0
   b6+       828.40       29.4
   y16+++    837.71       16.3
   y17+++    857.56       51.3
   b14++     863.87       36.2
   y9++      875.41       21.5
   y18+++    912.94       24.7
   y10++     919.29       15.9
   b19+++    930.53       14.2
   y19+++    951.23       18.8
   a21+++    988.34       14.9
   b21+++    997.25       18.6
   a22+++   1011.91       20.0
   y21+++   1017.83       48.0
   b22+++   1020.53       20.4
   y22+++   1050.23       28.0
   b23+++   1063.69       20.6
   y13++    1077.81       14.8
   y14++    1135.24       15.7
```

```
735. MORG-339_phosphopeptide_TMT_6%ACN_032017_OT.6410.6410.3 (sLDSDEsEDEDDDYQQkR)

   Ion         m/z   Intensity
   y5++      476.49       13.8
   y6++      533.97       14.5
   b3+       545.38       12.7
   y7++      591.29       10.2
   b9++      644.63        5.8
   y8++      648.89       10.4
   y9++      713.49       18.8
   b5+       747.50       12.7
   b11++     766.00        5.5
   y10++     771.08       12.2
   y11++     835.61       12.1
   a13++     867.17       15.8
   b6+       876.58       12.4
   b13++     881.53        8.2
   y12++     918.46        7.6
   b15++    1027.15        8.4
   y14++    1040.77        7.8
   y6+      1066.80        5.9
   y15++    1084.62       15.0
   b9+      1287.48        7.1
   b11+     1531.72        7.4
   b12+     1646.68        8.5
```

```
736. MORG-339_phosphopeptide_TMT_25%ACN_032017_OT.14445.14445.3 (tSSESIVSVPAsSTSGSPSRVIYAk)

   Ion         m/z   Intensity
   y1+       376.27       13.2
   y8++      582.09       29.6
   y3+       610.50       17.0
   b5+       721.36       13.3
   y4+       723.60       14.6
   y5+       822.63       18.3
   b6+       834.51       19.3
   y15++     910.71       29.0
   a17++     923.90       15.3
   b7+       933.58       21.8
   b17++     937.67       16.1
   y16++     959.25       35.0
   b18++     986.16       40.3
   a8+       992.81       40.6
   b8+      1020.58       14.0
   b19++    1029.42       13.2
   y18++    1052.00       14.0
   y19++    1101.69       30.4
   b20++    1107.36       15.6
   b9+      1119.73       23.9
   b21++    1157.05       18.6
   y8+      1162.72       17.6
   b22++    1213.93       21.5
   b10+     1216.68       13.9
```

```
737. MORG-339_phosphopeptide_TMT_10%ACN_032017_OT.5839.5839.3 (aEDGAAPsPSSETPk)

   Ion         m/z   Intensity
   y3++      287.91        8.8
   y2+       473.31       27.6
   y7++      487.72       13.0
   b8++      504.33        9.8
   y8++      570.77       14.3
   y3+       574.41       18.2
   b10++     597.01       12.0
   b4+       602.25        8.3
   y9++      619.91       41.9
   a11++     626.28       11.4
   b11++     640.65       32.2
   a5+       645.64       24.9
   b5+       673.22       14.2
   y11++     691.18       15.9
   b12++     705.24       10.9
   a13++     741.62        8.4
   b6+       744.40       39.7
   b13++     755.49       28.7
   b14++     803.86       11.3
   b7+       841.36       10.2
   b12+     1408.70       11.3
```

```
738. MORG-339_phosphopeptide_TMT_4%ACN_032017_OT.8404.8404.3 (qsNAsSDVEVEEk)

   Ion         m/z   Intensity
   a1+       330.06        5.1
   b1+       358.28       15.6
   y4++      367.28       13.7
   y1+       376.40       27.9
   y5++      431.97        7.9
   y6++      481.34        7.5
   y2+       505.38       32.3
   a7++      526.15        8.7
   b7++      540.15       10.1
   a8++      575.93       18.6
   y8++      582.36        7.5
   b8++      589.87       24.8
   y3+       634.36       30.4
   b9++      654.41       43.1
   y9++      666.22        8.1
   a10++     689.72        5.9
   b10++     703.94       46.1
   y4+       733.50       19.3
   b11++     768.38       46.7
   b12++     832.94       29.9
   y5+       862.49       17.7
   b7+      1079.38       15.4
   a8+      1150.72        4.5
   b8+      1178.50       24.3
   b9+      1307.49       22.8
   b10+     1406.54        5.6
   b11+     1535.66        6.2
```

```
739. MORG-339_phosphopeptide_TMT_4%ACN_032017_OT.5571.5571.2 (qEtVEcLR)

   Ion         m/z   Intensity
   y2+       288.20        3.8
   y3+       448.38        9.6
   a2+       459.35        6.4
   b2+       487.61       10.7
   y4+       577.31       11.2
   b5+       896.47        8.9
   b6+      1056.55        8.9
   b7+      1169.60        7.1
```

```
740. MORG-339_phosphopeptide_TMT_6%ACN_032017_OT.2321.2321.4 (sSSREkQSEDTNPESk)

   Ion         m/z   Intensity
   y4++      345.30       24.1
   y1+       376.29       11.2
   y5++      402.46        5.9
   y2+       463.36       26.5
   b9+++     519.78        5.5
   b10+++    558.47       26.0
   b3+       571.62        9.4
   y8++      574.52       11.1
   a11+++    582.77        9.0
   y3+       592.28       22.2
   a6++      593.37        8.1
   b6++      607.68       12.5
   y12+++    617.39        9.9
   y9++      618.27       19.0
   a12+++    620.55      109.0
   b12+++    630.08       49.0
   a7++      657.49        6.0
   b13+++    662.36       15.4
   y13+++    669.33       12.0
   b7++      671.52        9.9
   y4+       689.41       29.7
   a14+++    696.17        9.1
   a4+       699.35       11.1
   a8++      700.86        5.6
   b14+++    705.53       38.5
   a15+++    725.42        6.4
   b15+++    734.44       10.2
   b9++      779.48        8.5
   y5+       803.48        8.9
   b10++     836.87        9.8
   b11++     887.79        9.1
   y6+       904.56       15.8
   a12++     930.74        8.8
   b12++     944.60       43.8
```

```
741. MORG-339_phosphopeptide_TMT_18%ACN_032017_OT.10610.10610.4 (eRPkTPPSNLSPIEDAsPTEELR)

   Ion         m/z   Intensity
   y2+       288.27       11.7
   y3+       417.33       11.9
   b10+++    527.23       20.3
   a11+++    546.59       17.4
   y9++      549.35       12.4
   b11+++    556.23       43.9
   b12+++    588.39       12.8
   y10++     613.84       13.2
   b13+++    626.15       43.4
   y5+       647.56       16.3
   a8++      661.97       13.2
   b14+++    669.20       38.2
   y11++     669.82       29.4
   b15+++    707.56       43.9
   y19+++    711.19       13.9
   a9++      719.00       43.8
   b16+++    731.34       53.9
   b9++      733.39       24.1
   y6+       744.42       39.3
   y13++     762.54       30.4
   b10++     790.00       12.9
   b18+++    818.62       13.1
   a11++     819.61       12.8
   b11++     833.76       59.0
   a19+++    843.43       11.7
   b19+++    852.39       13.0
   y21+++    862.83       15.6
   b12++     882.00       16.6
   b20+++    895.94       19.5
   y7+       911.40       26.0
   b13++     938.27       22.0
   b21+++    939.15       38.4
   b22+++    976.73       22.3
   y8+       982.45       17.9
   b14++    1003.34       22.9
   b15++    1060.84       18.6
   b16++    1096.43       13.1
   y9+      1097.46       20.5
   y10+     1226.60       17.1
   y12+     1436.68       17.2
   a9+      1437.67       15.1
```

```
742. MORG-339_phosphopeptide_TMT_20%ACN_032017_OT.3339.3339.4 (vGsTENIkHQPGGGR)

   Ion         m/z   Intensity
   y5++      222.29        9.5
   b3++      277.33       15.4
   y4+       346.20        9.7
   y7++      354.95       16.2
   y8+++     355.90        8.1
   b2+       386.42        8.1
   y10+++    431.82       22.6
   y5+       443.29       50.0
   b6++      449.05       10.4
   b8+++     456.42       13.9
   y11+++    474.76       12.7
   a7++      492.18       19.4
   a9+++     493.26       27.4
   b9+++     502.31       44.7
   b7++      505.77       13.1
   y12+++    508.01       41.5
   y8++      533.37       16.1
   a10+++    535.57       17.9
   b10+++    545.09       57.5
   b3+       553.43        9.8
   y13+++    563.59        8.0
   y6+       571.44       14.5
   b11+++    577.46        9.9
   y9++      589.38       13.1
   a13+++    606.39        8.1
   b13+++    615.40       10.2
   a14+++    625.16       11.3
   b14+++    634.36       11.0
   y10++     647.09       12.0
   b4+       654.48        9.5
   b8++      684.35       24.9
   y7+       708.48       13.8
   a9++      739.07        8.4
   b9++      752.88       21.1
   b5+       783.44       10.1
   b10++     817.16       16.2
```

```
743. MORG-339_phosphopeptide_TMT_25%ACN_032017_OT.13998.13998.4 (sGEGEVSGLLRkVPR)

   Ion         m/z   Intensity
   a5+++     248.21       16.8
   y4++      364.92       11.0
   b1+       397.23       17.3
   b10+++    413.25       12.7
   y9+++     419.03       23.5
   a6++      420.54       11.3
   b6++      434.50        9.8
   y10+++    452.13       76.8
   b11+++    465.38        9.5
   b7++      478.68       10.5
   a8++      492.58       15.6
   y11+++    495.03       14.9
   y6++      499.47       18.1
   b8++      506.98      204.9
   y12+++    514.12       48.0
   y7++      555.90       16.8
   y13+++    557.11       10.3
   b9++      562.85       11.8
   y14+++    576.11       50.7
   b3+       583.35       22.1
   b12+++    584.38       22.1
   a13+++    608.16       12.7
   a4+       612.20       12.6
   b13+++    617.27       38.3
   b10++     619.54       47.2
   y9++      628.05       47.2
   a14+++    640.31       22.4
   b14+++    650.20       22.3
   y10++     677.71       26.3
   a5+       741.37        9.5
   y11++     742.41       15.3
   b5+       769.39       50.7
   y12++     770.48       20.7
   y14++     863.54        9.2
   b6+       868.46       14.8
```

```
744. MORG-339_phosphopeptide_TMT_6%ACN_032017_OT.6241.6241.3 (kPAQETEETsSQEsAEED)

   Ion         m/z   Intensity
   y4+       463.17        5.4
   b7++      506.82        5.4
   a8++      557.65        8.5
   b9++      621.97       12.8
   y5+       630.33        5.3
   y11++     686.57       16.4
   a5+       755.31        5.4
   y6+       759.22        5.9
   y13++     801.09        8.3
   b12++     813.55       13.5
   a6+       856.27        7.8
   y16++     964.88        6.4
   y8+       974.45        9.6
   a17++    1111.69        5.7
   y9+      1141.29        8.1
   y10+     1242.41       10.7
   y11+     1371.55       10.5
```

```
745. MORG-339_phosphopeptide_TMT_18%ACN_032017_OT.19614.19614.4 (eQEWDsssDGTERLPEEEEIGPFSkGIk)

   Ion         m/z   Intensity
   y7++      617.45        7.8
   y8++      646.65       12.4
   a16+++    763.36       12.5
   y10++     767.62        9.2
   b4+       802.45       10.1
   a17+++    806.38        9.7
   a11++     832.15        9.3
   y19+++    859.13       31.6
   y20+++    897.49       17.4
   b5+       917.42       16.8
   a20+++    930.38        6.9
   y21+++    953.17       13.2
   b21+++    958.56        7.8
   y13++     961.15        8.9
   a13++     974.70       32.3
   a22+++    981.02        9.0
   b13++     989.00       19.9
   b22+++    990.77        6.7
   y22+++   1008.37       16.2
   y14++    1009.50       30.6
   b14++    1045.26       10.3
   a6+      1056.49        7.1
   y15++    1066.03       12.2
   b6+      1084.57        7.7
   y25+++   1164.12       11.5
   b17++    1222.68        7.8
```

```
746. MORG-339_phosphopeptide_TMT_4%ACN_032017_OT.5046.5046.3 (yRTREQEssGEEDNDLSPEER)

   Ion         m/z   Intensity
   y2+       304.31        6.4
   b4++      403.86        5.7
   y4+       530.36        8.7
   b6++      532.37        8.6
   b7++      597.08       12.7
   y5+       617.52        8.2
   a8++      665.90       10.7
   y11++     666.69        8.5
   y6+       730.45       18.0
   y7+       845.40        8.1
   b11++     857.31        8.3
   a5+       907.58       83.6
   b12++     921.53       14.0
   y15++     926.65        5.9
   y8+       959.41       15.5
   b14++    1036.02        6.9
   y9+      1074.43        8.5
   b15++    1093.28       10.3
   a16++    1135.60       10.7
   b16++    1150.22        9.2
   b17++    1193.75        7.6
   y10+     1203.45        8.2
   b18++    1242.19        5.6
   b19++    1306.76       14.3
   a8+      1331.87        8.3
   y11+     1332.67        6.7
   b20++    1371.49       10.9
   y12+     1389.55        7.0
```

```
747. MORG-339_phosphopeptide_TMT_22%ACN_032017_OT.8775.8775.4 (nLAkPGVTstsDSEDEDDQEGEkkR)

   Ion         m/z   Intensity
   y7++      667.20       15.7
   a15+++    724.73       15.8
   y10++     846.62       17.2
   a4+       857.23       10.4
   b4+       885.76       14.8
   y3+       889.54       10.3
   b19+++    896.69       12.9
   y11++     903.87       12.7
   y18+++    918.43       10.0
   a12++     921.52       12.9
   a20+++    930.26       13.8
   b20+++    939.27       14.6
   a21+++    949.50       87.9
   y19+++    951.75       22.7
   a5+       954.59       38.6
   b21+++    959.19       30.2
   b22+++   1001.24       12.6
   y21+++   1002.76       14.6
   y13++    1011.98       14.5
   a14++    1029.82       11.7
   b6+      1040.03       12.3
   b14++    1043.29       14.0
   y14++    1069.02       11.5
   y5+      1076.03       15.1
   b15++    1100.94       10.3
   b23+++   1121.17        9.5
   b7+      1139.12       11.4
   y15++    1152.94       10.3
```

```
748. MORG-339_phosphopeptide_TMT_4%ACN_032017_OT.17147.17147.4 (dDDDIDLFGsDDEEESEEAk)

   Ion         m/z   Intensity
   a1+       317.37        3.8
   b4++      345.94        5.0
   y1+       376.38       13.3
   y2+       447.33       20.6
   b2+       460.09        5.8
   y6++      461.43        5.6
   b7++      516.18       13.6
   y3+       576.48       18.7
   b13+++    588.30        4.9
   y8++      589.91       18.6
   b9++      618.34       13.1
   b14+++    630.97        5.5
   y9++      647.87        9.4
   a15+++    664.87        3.8
   y15+++    670.29        8.9
   b15+++    673.95        6.6
   a10++     687.46        6.2
   a16+++    693.66        4.7
   y17+++    746.27        8.2
   b18+++    788.92       16.3
   b5+       803.41       10.0
   b19+++    812.75       22.8
   y12++     817.46       19.5
   b13++     881.31        4.1
   y13++     891.17       12.1
   b6+       918.48       14.7
   y6+       921.25        4.3
   b14++     945.72        9.1
   y14++     947.44        8.2
   a15++     996.04        4.4
   y15++    1005.18        8.7
   b15++    1010.34       10.6
   b7+      1031.53       18.7
   b17++    1118.30        3.8
   b8+      1178.58        8.3
   y8+      1179.58        3.6
   b19++    1218.33        5.7
```

```
749. MORG-339_phosphopeptide_TMT_20%ACN_032017_OT.8538.8538.2 (aVYQGPGSsPVkS)

   Ion         m/z   Intensity
   b3+       563.45       12.6
   y4+       659.48       24.3
   b4+       691.52       20.0
   b5+       748.31       15.3
   b6+       845.64        7.4
   y7+       970.89       12.2
   b8+       989.89       10.7
   y8+      1067.63       10.6
   y9+      1124.67       11.7
   y11+     1415.79        8.5
```

```
750. MORG-339_phosphopeptide_TMT_14%ACN_032017_OT.12368.12368.2 (sLsSPTDNLELSAR)

   Ion         m/z   Intensity
   b1+       317.21       17.6
   a5+       753.44       10.3
   y7+       802.40       25.9
   a6+       853.98       10.0
   b6+       882.44       66.6
   y8+       917.69       12.5
   b7+       997.65       22.8
   y9+      1018.62       12.8
   b8+      1111.53        8.8
   y10+     1115.60       37.4
   b13+     1624.91        9.2
```

```
751. MORG-339_phosphopeptide_TMT_14%ACN_032017_OT.12844.12844.3 (lLkEGEEPTVYsDDEEPk)

   Ion         m/z   Intensity
   b4++      471.96       12.1
   y2+       473.29       14.8
   b6++      564.84       12.2
   b7++      629.63       22.2
   y4+       731.55       15.5
   b10++     777.85       18.9
   y5+       846.54       13.7
   b11++     859.61       15.2
   y13++     923.44       18.9
   a12++     929.45       17.2
   y6+       961.46       11.7
   b13++    1000.72       16.5
   b14++    1058.37       12.7
   a15++    1108.74       10.3
   b6+      1128.55       10.9
   b16++    1187.16       13.5
   b7+      1257.74       12.4
   y11+     1588.83       10.0
```

```
752. MORG-339_phosphopeptide_TMT_4%ACN_032017_OT.7686.7686.3 (nHsLEEEFER)

   Ion         m/z   Intensity
   y1+       175.21        5.8
   a2++      227.03        9.4
   b2++      241.12        6.7
   y2+       304.00        4.0
   b3++      324.58        9.8
   b1+       344.35        8.5
   a4++      367.57        7.2
   b4++      381.23        6.0
   a5++      431.48        7.9
   b5++      445.76       23.1
   y3+       451.33       14.0
   y7++      476.43       13.7
   b2+       481.33        9.8
   a6++      495.84       17.4
   b6++      510.56       27.7
   y8++      559.87        8.7
   b7++      574.79       16.5
   y4+       580.35       15.6
   a3+       620.01        5.0
   y9++      628.52        9.7
   b8++      648.50       10.3
   y5+       709.37       23.3
   b9++      712.90        9.5
   y6+       838.38       20.0
   y7+       951.46        7.0
   b6+      1019.41        5.3
```

```
753. MORG-339_phosphopeptide_TMT_16%ACN_032017_OT.16696.16696.3 (rAsGQAFELILsPR)

   Ion         m/z   Intensity
   y2+       272.18       13.4
   a5++      391.63        7.9
   b5++      405.28       11.6
   a6++      426.33        5.6
   y3+       439.33        6.5
   b6++      440.67       13.4
   y7++      454.31        8.9
   b2+       457.57        5.4
   a7++      500.33        9.6
   b7++      514.43       14.2
   y4+       552.18        8.3
   a8++      564.49        9.0
   b8++      578.75       17.3
   a9++      621.29       11.8
   b3+       624.26       11.5
   y10++     627.39       35.5
   b9++      635.44       30.5
   y5+       665.39        6.1
   b10++     691.89       19.6
   a11++     734.09       17.5
   y12++     739.62        6.9
   b11++     748.49       12.2
   y13++     774.53        6.0
   y6+       778.46        6.9
   b5+       809.33        5.7
   a13++     866.72        6.1
   b6+       880.41        6.3
   b7+      1027.69        6.9
   y8+      1054.51        9.8
   y9+      1125.57        6.7
```

```
754. MORG-339_phosphopeptide_TMT_6%ACN_032017_OT.6641.6641.4 (eAGIVHsDAEkEQEEEEQk)

   Ion         m/z   Intensity
   y3++      317.31        5.9
   y1+       376.39        8.4
   a2+       401.95        6.6
   b6++      419.14        8.9
   b3+       487.34        6.9
   y2+       504.37       10.2
   y6++      511.16        9.0
   b8++      559.53       14.9
   y7++      574.76        5.8
   a9++      581.67        7.5
   b4+       600.37        7.9
   b12+++    602.60        9.5
   y3+       633.40       10.9
   y8++      639.42        5.7
   b13+++    645.11        8.3
   a10++     645.98        9.2
   y12+++    650.81       11.6
   b10++     659.97        9.1
   a5+       671.54        5.8
   a14+++    678.78        9.5
   b14+++    688.12       14.8
   b5+       699.47       20.9
   y13+++    706.09       18.2
   a15+++    721.27       24.5
   b15+++    731.07       11.3
   y14+++    752.28       11.1
   y4+       762.48       10.2
   a16+++    764.56        7.2
   b16+++    774.22       15.6
   y15+++    784.72       11.0
   a17+++    807.94        6.7
   a6+       808.72        6.6
   b17+++    817.09       15.8
   y9++      817.93       15.4
   a18+++    850.29        5.9
   b18+++    859.95       19.4
   y10++     882.72       11.0
   y5+       891.50        9.6
   y11++     918.18       15.8
   b13++     967.07       10.9
   y12++     975.14        9.5
   y6+      1020.56        8.1
   y13++    1059.14        7.0
   b15++    1096.15        7.0
   y7+      1148.32        5.5
```

```
755. MORG-339_phosphopeptide_TMT_14%ACN_032017_OT.18183.18183.2 (nSSLLSFDsEDENE)

   Ion         m/z   Intensity
   b2+       431.30       16.1
   y4+       506.25       12.6
   b3+       518.29       22.5
   b4+       631.39       42.9
   b5+       744.50       86.5
   y6+       802.27       17.8
   a6+       803.33       17.5
   b6+       831.51       38.6
   y7+       917.26       33.1
   b7+       978.53       37.0
   y8+      1064.33       32.0
   a8+      1065.34       26.8
   b8+      1093.55       30.4
   y9+      1151.34       69.5
   y10+     1264.38       34.7
   y11+     1377.56       16.5
   b10+     1389.66       19.9
   y12+     1464.62       11.0
   b11+     1504.65       17.8
   y13+     1551.58       11.9
   b12+     1633.68       28.9
```

```
756. MORG-339_phosphopeptide_TMT_16%ACN_032017_OT.2432.2432.3 (sLRTAGVQR)

   Ion         m/z   Intensity
   y1+       175.13        6.9
   y4++      230.18        9.2
   y2+       303.27        7.4
   a3++      319.81       20.0
   b3++      333.63       38.7
   b4++      384.39       21.4
   y7++      394.40       53.0
   b1+       397.16       31.9
   y3+       402.22       25.1
   a5++      405.36       14.6
   b5++      419.79       18.6
   b6++      448.46       13.9
   y8++      450.97       59.3
   y4+       459.28       15.7
   a7++      483.94       10.0
   b7++      497.93       18.9
   y5+       530.30       17.1
   b8++      561.87       26.4
   y6+       631.40       41.0
   b3+       666.49        6.2
```

```
757. MORG-339_phosphopeptide_TMT_14%ACN_032017_OT.21520.21520.3 (eLALssPEDLTQDFEELkREER)

   Ion         m/z   Intensity
   y6++      530.64        8.5
   b4+       656.43        5.8
   y9++      733.11       18.4
   a5+       795.66       11.3
   a14++     954.54        7.7
   y13++     961.86       26.8
   y14++    1019.22       16.3
   b15++    1033.37        9.3
   a7+      1059.08        8.1
   y6+      1060.11        9.9
   y16++    1132.35       21.7
   a17++    1139.89        6.2
   y17++    1215.73       13.0
   b9+      1331.46        8.3
   b18++    1332.58        8.7
   y20++    1391.02        6.3
   y21++    1448.11       10.7
   b13+     1788.92        7.0
```

```
758. MORG-339_phosphopeptide_TMT_14%ACN_032017_OT.14460.14460.3 (sQtPPGVAtPPIPk)

   Ion         m/z   Intensity
   b1+       317.23       11.2
   y1+       376.33        7.3
   y5++      390.84       20.2
   y2+       473.40       25.8
   a7++      474.47        9.4
   b7++      488.74        6.6
   y7++      516.99        9.7
   y8++      565.96       16.1
   y9++      594.98        6.4
   a9++      600.94        8.6
   b9++      614.67        5.8
   b3+       625.79       11.1
   y10++     643.46       14.5
   a10++     648.94       11.6
   y4+       683.55        6.3
   y11++     691.91       11.2
   a11++     698.15        7.3
   b11++     711.93        7.3
   a12++     754.47       19.4
   b12++     768.44       27.8
   y5+       780.58       13.1
   a13++     802.91        6.8
   b6+       877.42        7.5
   b7+       976.47       12.1
   b8+      1047.53       15.0
   y8+      1131.61       11.2
   b9+      1228.53        8.8
```

```
759. MORG-339_phosphopeptide_TMT_30%ACN_032017_OT.16121.16121.4 (stQLLQAAAAEASLNkSk)

   Ion         m/z   Intensity
   b4+++     247.15       15.1
   b1+       317.28       17.5
   y3++      411.09       14.1
   y2+       463.43       20.1
   y4++      467.88       19.3
   y11+++    516.40       13.6
   b13+++    518.15       12.6
   y5++      524.45       14.7
   b7++      526.39       19.0
   b14+++    555.29        8.6
   b8++      562.00       22.6
   y6++      567.84       21.9
   a9++      583.70        9.4
   b9++      597.49       25.2
   a3+       598.56       17.1
   y7++      603.68       14.2
   a10++     618.61       16.1
   y14+++    620.48       36.6
   b3+       626.11       12.9
   b10++     633.01       28.0
   y8++      667.90       18.4
   b11++     697.45       13.6
   y9++      703.48       21.5
   b16+++    712.90       40.4
   b12++     732.78       13.9
   b4+       738.99       17.2
   b17+++    741.75       17.0
   y17+++    761.72        8.8
   y11++     774.67       12.4
   b13++     776.38       10.5
   a5+       824.49       15.1
   b14++     832.80       10.7
   y16++    1050.65        8.3
   b7+      1051.70       15.2
   a16++    1054.49        8.9
   b8+      1122.47        9.0
```

```
760. MORG-339_phosphopeptide_TMT_12%ACN_032017_OT.4897.4897.3 (eNSAAPSPGRPQsAsPAkEEQk)

   Ion         m/z   Intensity
   y7++      644.39        6.8
   y4+       762.48        4.3
   a14++     815.90        7.2
   b14++     830.55        4.8
   y12++     959.44        4.7
   b8+       983.23        5.4
   a17++     984.09        5.5
   b17++     998.00        8.0
   a9+      1012.79       13.9
   y14++    1065.76        7.4
   y16++    1158.45        6.5
   a18++    1162.26        5.4
   b18++    1176.76        5.4
   b10+     1196.90        6.0
   y17++    1206.98        7.2
   b19++    1241.05        4.7
   y18++    1242.00        4.3
   a11+     1265.48        5.5
   y7+      1287.78        6.5
   y12+     1917.69        4.5
```

```
761. MORG-339_phosphopeptide_TMT_20%ACN_032017_OT.8237.8237.2 (tGSsPTQGIVNk)

   Ion         m/z   Intensity
   y1+       376.28        9.2
   y3+       589.32       12.0
   y5+       759.50       26.7
   a6+       812.54        9.2
   b6+       840.16       16.6
   a7+       940.44        9.1
   b7+       968.57       15.2
   y7+       988.81        8.5
   y8+      1085.66       37.1
   b9+      1138.52       14.4
   b10+     1237.57       12.2
```

```
762. MORG-339_phosphopeptide_TMT_10%ACN_032017_OT.6586.6586.3 (sSSEGGDAGNDTRNTTADLQkAGEGT)

   Ion         m/z   Intensity
   y11++     660.47       13.3
   b4+       700.54        9.9
   b13++     772.08        9.9
   y15++     896.72       47.9
   y7+       919.70        9.4
   b7+       929.41       20.7
   b16++     930.41       12.8
   b17++     965.57       17.7
   b8+      1000.85       32.4
   a18++    1009.24       30.6
   y17++    1011.02       26.3
   y8+      1032.54       17.8
   y18++    1039.74       10.8
   b9+      1057.61       13.2
   a19++    1066.40       13.9
   y19++    1075.26       41.4
   y20++    1132.98       12.5
   b10+     1171.79       10.9
   y22++    1189.69       34.5
   y10+     1218.75       11.4
   y23++    1254.05       12.4
   b11+     1286.52       18.0
   y11+     1319.84       11.1
   a22++    1344.64       10.0
```

```
763. MORG-339_phosphopeptide_TMT_14%ACN_032017_OT.21313.21313.4 (aLAEEDsEDELPsDVDFNDPYFAEEVkk)

   Ion         m/z   Intensity
   y5++      545.72        9.9
   a13+++    583.45       10.5
   b10++     699.86       12.7
   b11++     756.44       19.6
   y9++      784.89       21.1
   y16+++    814.48       56.3
   y10++     842.79       21.3
   y17+++    847.16      115.9
   y18+++    884.57       29.8
   y11++     899.61       22.7
   b21+++    914.55       29.9
   y19+++    927.87       13.1
   a14++     931.56        8.1
   y20+++    966.23       16.7
   y12++     973.00       17.3
   a23+++    977.35       11.1
   a15++     981.69       13.6
   b23+++    986.86       13.0
   b15++     995.44       22.4
   a7+       997.83       10.0
   b24+++   1029.65       11.5
   y13++    1030.80       31.9
   y22+++   1064.78       15.5
   y14++    1080.27       14.4
   y15++    1137.91        7.9
   b8+      1154.39       10.0
   b18++    1183.80       17.7
   y26+++   1212.21        8.1
   y16++    1220.85       33.2
   y17++    1269.82       27.9
   b21++    1371.45        8.5
   b10+     1398.59       22.6
   b11+     1511.67       50.5
   y21++    1512.64       50.5
```

```
764. MORG-339_phosphopeptide_TMT_25%ACN_032017_OT.19981.19981.4 (iVEtINsDSDSEFGIPkk)

   Ion         m/z   Intensity
   a6+++     318.26        9.0
   y3++      415.85       62.7
   y4++      472.09        9.9
   y5++      501.03       33.2
   b3+       571.26       12.0
   y6++      574.67       33.5
   b13+++    609.25        9.8
   a8++      617.49        9.7
   a14+++    619.72       10.6
   b14+++    628.42        9.5
   b8++      631.63       12.7
   y7++      638.84       20.7
   y13+++    654.19       11.6
   a15+++    656.54        9.6
   a9++      660.85        9.1
   b15+++    666.29       18.7
   b9++      675.05       11.0
   y8++      682.20       14.4
   a16+++    689.12       31.9
   y14+++    692.08       10.6
   b16+++    698.85       31.4
   a10++     718.54        9.0
   y9++      740.29       11.8
   b11++     776.03       19.5
   y10++     783.48       10.2
   y16+++    796.01       12.8
   b17+++    817.83        9.9
   y3+       830.11        9.3
   y11++     840.61       24.2
   b5+       864.99       17.4
   a13++     899.56       12.5
   b13++     914.15       22.8
   a14++     927.94        9.5
   b14++     942.29       19.8
   a15++     985.37       20.7
   b15++     999.17       45.3
   y5+      1000.99       10.8
   b10+     1463.60        8.6
   b12+     1679.90        9.4
```

```
765. MORG-339_phosphopeptide_TMT_4%ACN_032017_OT.5388.5388.2 (qASPtEVVER)

   Ion         m/z   Intensity
   a1+       330.27        8.0
   b1+       358.33       10.0
   b2+       429.33        7.4
   y4+       502.36       14.1
   b3+       516.38        6.9
   y5+       631.18        5.6
   b6+       923.46       10.9
   b7+      1022.56        8.5
   y9+      1067.45       10.0
   b8+      1121.61        8.6
   b9+      1250.75       14.3
```

```
766. MORG-339_phosphopeptide_TMT_18%ACN_032017_OT.4530.4530.4 (aRVDHGAEIITQsPSRSSVASPR)

   Ion         m/z   Intensity
   b4++      336.53      137.8
   b5++      404.95       26.9
   y8++      430.26       30.3
   b10+++    430.86       38.5
   b6++      433.49       21.4
   b11+++    465.01       20.6
   b7++      469.09       31.6
   y10++     522.31       97.6
   y5+       529.49       22.7
   b8++      533.40       48.7
   b9++      590.08       50.7
   y6+       616.51       36.2
   a10++     632.77       28.4
   y18+++    641.92       21.8
   a4+       642.98       26.5
   b10++     646.49       45.5
   a16+++    667.70       43.3
   y12++     669.49       33.6
   b4+       671.34       75.6
   a17+++    696.01       24.2
   b11++     697.17       36.8
   y7+       703.45       65.4
   b17+++    705.40       21.9
   y13++     720.64       34.5
   y21+++    758.89       35.9
   b12++     761.16       35.2
   b19+++    767.84       33.1
   y14++     777.15       35.9
   a20+++    782.09       42.8
   b20+++    791.56       39.1
   b5+       808.56       29.8
   b21+++    820.38       40.7
   y15++     833.62       33.9
   b13++     844.66       23.8
   b22+++    852.81       24.6
   y8+       859.66       30.2
   a14++     878.68       26.2
   y16++     898.12       20.7
   y17++     933.70       22.3
   y18++     962.29       31.9
   y19++    1030.77      152.6
```

```
767. MORG-339_phosphopeptide_TMT_14%ACN_032017_OT.5459.5459.3 (ssGSPYGGGYGSGGGSGGYGSR)

   Ion         m/z   Intensity
   y3+       319.22       18.5
   y4+       482.27       18.5
   y12++     499.72       21.4
   b9++      530.39       27.7
   y5+       539.36       19.3
   y13++     581.68       22.0
   y6+       596.29       30.8
   a10++     597.45       24.2
   y14++     610.04       20.8
   b10++     611.94       35.7
   b4+       628.31       17.5
   y15++     638.59       20.8
   b11++     640.56       28.2
   y16++     666.72       31.4
   b12++     683.60       39.6
   a13++     698.75       15.7
   b13++     712.58       30.7
   b5+       724.82       35.1
   a14++     726.44       30.1
   y9+       797.27       36.8
   a16++     798.42       24.2
   b16++     812.80       22.1
   a17++     827.74       13.7
   y10+      854.37       29.7
   a18++     855.42       25.0
   a6+       860.77       14.8
   b18++     870.14       24.5
   b6+       888.36       32.6
   y11+      941.47       24.2
   b7+       945.44       22.2
   b19++     951.62       32.7
   y21++     952.43       21.3
   b20++     980.27       29.0
   y12+      998.39       31.7
   b8+      1002.44       15.2
   b9+      1059.48       28.4
   b10+     1222.51       28.9
   b11+     1279.45       13.3
```

```
768. MORG-339_phosphopeptide_TMT_16%ACN_032017_OT.11769.11769.3 (sPPEEPPDFccPk)

   Ion         m/z   Intensity
   y2++      237.46        8.0
   b2++      247.27        9.2
   y3++      317.24       39.9
   y1+       376.41        9.2
   y4++      397.19       18.2
   b6++      473.35       32.8
   b2+       494.50        9.1
   y7++      577.06       10.5
   b8++      579.96        8.6
   b3+       591.56        9.5
   y8++      625.45       48.6
   y3+       633.17       25.8
   a9++      639.49       14.6
   b9++      653.64       21.4
   y9++      689.58       18.6
   a4+       692.30        9.6
   a10++     719.45        9.9
   b4+       720.32        9.3
   b10++     733.41       16.7
   y4+       793.42       17.4
   b11++     813.54       17.7
   b5+       849.35       35.7
   y12++     851.56       15.7
   b6+       946.56        9.6
   b8+      1158.35        8.7
   b9+      1305.57        9.1
   b10+     1465.57       12.2
```

```
769. MORG-339_phosphopeptide_TMT_8%ACN_032017_OT.3373.3373.3 (rQPsPQPSPRDLQsSER)

   Ion         m/z   Intensity
   b6++      502.68       13.3
   b2+       514.34        9.5
   y9++      584.27        8.9
   y10++     627.59       24.4
   a9++      628.55        9.5
   y11++     676.48       18.5
   y5+       686.61        8.2
   a10++     706.62       14.8
   b10++     720.48       73.9
   y12++     740.60       28.0
   a4+       750.40       22.2
   a11++     764.38       18.4
   y6+       799.34        8.2
   b12++     834.52        9.6
   y14++     872.57       11.8
   y15++     921.18       12.1
   y16++     985.04        7.3
   b6+      1003.39       17.1
   a15++    1011.48       10.5
   b16++    1090.79       10.6
   y11+     1351.63        9.7
```

```
770. MORG-339_phosphopeptide_TMT_10%ACN_032017_OT.2905.2905.4 (yDERPGPsPLPHRDR)

   Ion         m/z   Intensity
   y5++      340.88       21.8
   b1+       393.28        8.1
   y6++      397.32       15.9
   y11+++    436.73       15.5
   y3+       445.93       54.9
   a10+++    465.51       10.3
   b10+++    474.31       10.9
   a2+       480.70       12.3
   y12+++    489.27       12.7
   a11+++    497.26       39.0
   b11+++    507.41       18.0
   a7++      508.32       58.1
   a12+++    543.53        7.6
   b12+++    552.91        8.1
   y9++      578.10       12.4
   y4+       583.69       10.1
   a13+++    595.59       18.5
   y10++     606.13       30.1
   a3+       609.19       11.1
   b3+       637.50       11.1
   b14+++    643.26       36.4
   y11++     655.20       13.9
   y5+       680.37        9.0
   a10++     697.21       10.2
   b10++     711.58       22.2
```

```
771. MORG-339_phosphopeptide_TMT_20%ACN_032017_OT.22162.22162.2 (tLsGPVTGLLATAR)

   Ion         m/z   Intensity
   b1+       411.31       11.6
   y4+       418.22       12.1
   a2+       496.41        6.6
   b2+       524.64        8.6
   y5+       531.50       13.8
   y6+       644.40       10.5
   y7+       701.55       13.9
   y8+       802.60       18.4
   b6+       944.60       15.6
   y10+      998.60       18.5
   y11+     1055.61       40.3
   b8+      1102.65        6.6
   b9+      1215.90       10.9
   y12+     1222.64       20.4
   a10+     1301.03        9.2
   b10+     1328.57       10.5
   y13+     1335.86       17.5
   b11+     1399.91       10.2
   a12+     1472.84        7.2
```

```
772. MORG-339_phosphopeptide_TMT_12%ACN_032017_OT.9514.9514.2 (sPELPEPSVR)

   Ion         m/z   Intensity
   a1+       369.24        9.4
   b1+       397.22       34.8
   y4+       458.27       38.2
   a3+       595.41       12.4
   b3+       623.38       26.1
   y6+       684.39       45.4
   b4+       736.41       34.1
   y7+       797.46       31.0
   a6+       934.70        9.3
   b6+       962.42       37.7
   y9+      1023.55       65.5
   b8+      1146.65       12.1
```

```
773. MORG-339_phosphopeptide_TMT_25%ACN_032017_OT.6975.6975.4 (nLAkPGVTSTSDsEDEDDQEGEkkR)

   Ion         m/z   Intensity
   b4++      443.18       20.8
   y5++      538.40       19.9
   b7++      569.96       16.2
   b8++      620.64       16.6
   y7++      667.06       49.6
   y8++      724.73       47.3
   b11++     758.18       15.3
   y9++      782.14       21.4
   b12++     815.44       40.5
   y10++     846.69       42.4
   a20+++    876.59       15.6
   a13++     884.68       27.4
   b4+       885.55       26.0
   b20+++    886.59       15.3
   y3+       889.51       15.2
   a21+++    895.70       31.0
   b13++     898.58       46.1
   y11++     903.96       34.5
   b21+++    905.01      111.4
   y20+++    917.08       36.9
   b14++     963.24       20.3
   y12++     968.72       21.7
   a6+      1012.10       31.8
   b15++    1020.83       26.4
   y13++    1052.19       39.5
   b23+++   1067.22       17.9
   a16++    1071.09       15.9
   y24+++   1130.07       15.9
   b17++    1142.79       27.7
   y15++    1153.27       16.1
   b18++    1200.59       27.6
   y20++    1375.36       22.7
```

```
774. MORG-339_phosphopeptide_TMT_20%ACN_032017_OT.13466.13466.3 (wRsLQQLAEER)

   Ion         m/z   Intensity
   y1+       175.08        9.1
   a2++      273.08        6.8
   b2++      286.52        8.0
   y2+       304.18       22.4
   b3++      370.35       11.2
   y3+       433.26        9.8
   a5++      476.83        6.4
   b5++      490.70       12.7
   y4+       504.28       11.2
   a6++      540.55       13.1
   a2+       544.17       21.8
   b6++      555.05       17.0
   a7++      597.71       14.4
   b7++      611.30       10.0
   y5+       617.44       13.8
   b8++      646.79        7.7
   y10++     655.67        9.3
   a9++      697.41        6.6
   b9++      711.54       37.7
   y6+       745.24       19.8
   b10++     776.04       15.9
   y7+       873.50       10.3
   y8+       986.62       14.1
```

```
775. MORG-339_phosphopeptide_TMT_25%ACN_032017_OT.9220.9220.3 (kkIPDPDSDDVsEVDAR)

   Ion         m/z   Intensity
   y6+       756.49       20.6
   a3+       800.65       24.9
   a11++     820.94       26.9
   y7+       855.42       39.9
   a13++     968.96       19.5
   y8+       970.82       19.5
   a5+      1013.07       32.4
   y16++    1033.57       22.3
   y11+     1287.58       34.8
   y12+     1384.45       33.5
```

```
776. MORG-339_phosphopeptide_TMT_16%ACN_032017_OT.6541.6541.4 (qksDAEEDGVtGSQDEEDSkPk)

   Ion         m/z   Intensity
   y4++      459.41       20.6
   y2+       473.34       12.0
   b4++      499.12       14.6
   b9+++     499.89       12.2
   y6++      581.49       11.4
   b6++      599.34       13.6
   y7++      646.25       18.9
   b7++      663.83       11.7
   y14+++    672.32       17.9
   b8++      721.56       30.6
   b15+++    722.29       11.8
   y9++      767.30       10.3
   a10++     785.71       15.6
   y17+++    796.58       11.8
   b10++     799.65       21.9
   y10++     811.22       19.2
   y18+++    820.62       30.0
   y3+       830.84       21.4
   a18+++    837.43       29.5
   y11++     839.70       38.9
   b18+++    846.60       34.6
   a3+       854.52       22.4
   b3+       882.64       14.3
   b12++     917.92       22.0
   y12++     930.17       20.5
   a20+++    986.01       11.3
   b4+       997.61       11.5
   y14++    1008.41       27.3
   b5+      1068.47       14.1
   y18++    1230.32       16.2
```

```
777. MORG-339_phosphopeptide_TMT_14%ACN_032017_OT.5497.5497.3 (sPSPAPPPPPPPPPPRR)

   Ion         m/z   Intensity
   y6++      360.29       15.3
   b1+       397.23       16.9
   y7++      408.94       18.1
   b6++      423.92       20.6
   y8++      457.43       24.3
   b7++      472.09       13.2
   y9++      505.94       28.1
   b8++      520.86       19.2
   y10++     554.50       40.7
   a9++      555.43       18.2
   b9++      569.21       14.2
   y11++     603.09       26.9
   b10++     618.06       33.2
   y5+       622.49       13.9
   a4+       650.67       36.7
   y12++     651.72      108.7
   a11++     652.60       38.3
   b11++     666.64       45.4
   y6+       719.47       18.8
   a5+       721.39       23.1
   y14++     735.68       34.5
   a13++     749.39       47.3
   y7+       816.56       24.2
   a6+       818.72       18.9
   y16++     827.74       46.3
   a15++     846.57       16.7
   y8+       913.59       19.1
   a7+       915.63       12.0
   y9+      1010.66       12.7
   a8+      1012.55       12.6
   b8+      1040.63       13.2
```

```
778. MORG-339_phosphopeptide_TMT_10%ACN_032017_OT.13929.13929.3 (vVDYSQFQEsDDADEDYGR)

   Ion         m/z   Intensity
   y3+       395.29       20.2
   b2+       428.33       12.5
   y4+       510.22       31.5
   b7++      535.08       21.1
   b3+       543.35       12.0
   b8++      599.31       15.6
   y5+       639.33       27.7
   b9++      663.29       30.1
   y11++     676.43       23.1
   b4+       706.47       28.8
   y12++     740.41       21.6
   y6+       754.38       34.9
   b5+       793.56       18.0
   b11++     804.38       17.5
   y13++     813.47       39.9
   y7+       825.50       41.4
   b12++     862.25       21.7
   y14++     877.80       20.4
   a13++     883.78       12.2
   b13++     897.65       36.7
   b6+       921.48       50.4
   y8+       940.38       24.6
   a14++     941.36       13.1
   y16++    1003.35       13.0
   b15++    1019.69       29.3
   y9+      1055.38       13.7
   b7+      1068.49       38.8
   b16++    1077.20       27.6
   y18++    1109.85       20.4
   b17++    1158.83       38.2
   b8+      1196.66       33.7
   y10+     1222.41       26.1
   b9+      1325.75       17.3
   y11+     1351.58       15.3
```

```
779. MORG-339_phosphopeptide_TMT_4%ACN_032017_OT.1456.1456.3 (dSGRGDsVSDNGSEAVR)

   Ion         m/z   Intensity
   y3+       345.34        8.6
   b6++      409.34       17.3
   b7++      493.19       18.5
   a8++      528.29        7.6
   y5+       561.68        6.6
   b9++      586.16        8.3
   y11++     600.45        5.9
   y6+       618.04       13.0
   a10++     629.19       12.6
   b10++     643.42       30.0
   b4+       645.66       18.4
   y12++     658.64       30.5
   b11++     700.73        6.5
   b5+       702.19        7.0
   b12++     729.18        7.2
   y7+       732.42       17.3
   b13++     771.91        8.0
   y16++     837.25       10.7
   y8+       847.37        8.3
   b15++     872.77       10.1
   b16++     921.75        7.6
   y9+       934.47       15.5
   y10+     1033.60       10.7
```

```
780. MORG-339_phosphopeptide_TMT_10%ACN_032017_OT.10853.10853.4 (aSGPLDTSQTAssDSEDVsSSDRk)

   Ion         m/z   Intensity
   b7++      436.67       12.7
   y7++      543.91       15.6
   y12+++    567.88       10.9
   b10++     594.47       33.5
   a11++     615.59       14.7
   a5+       627.13        9.8
   b11++     630.09       12.6
   y3+       647.64       11.6
   b5+       655.66       16.5
   y15+++    680.95       22.1
   y10++     709.91       17.6
   y16+++    723.49       24.2
   y4+       734.70       16.4
   y17+++    752.43       21.1
   a19+++    759.18       31.6
   y11++     767.67       34.0
   b19+++    769.21       35.3
   b6+       770.32       25.0
   a14++     840.83       12.0
   y12++     851.15       16.0
   b7+       871.61       18.7
   y21+++    894.04       13.6
   a8+       930.03       13.0
   y13++     934.57       32.1
   b8+       958.49       17.8
   y14++     970.06       25.0
   y15++    1020.52       29.0
   b9+      1086.49       16.4
   y7+      1087.42       13.5
```

```
781. MORG-339_phosphopeptide_TMT_14%ACN_032017_OT.12345.12345.3 (eQNcDLAPsGEVsPVkPLGR)

   Ion         m/z   Intensity
   y7++      498.51       17.9
   a3+       573.46        8.7
   b10++     691.75       10.8
   y11++     724.56       17.6
   a12++     791.50        9.3
   y12++     807.70       17.2
   a5+       848.68       14.8
   y13++     856.80       22.5
   a13++     874.95       21.3
   b5+       875.93       24.1
   b13++     889.32       25.7
   y14++     892.06       91.5
   y6+       898.29       26.6
   y15++     948.50       16.2
   y16++    1005.89       10.0
   b7+      1060.77       16.5
   b16++    1165.88       22.5
```

```
782. MORG-339_phosphopeptide_TMT_10%ACN_032017_OT.13928.13928.4 (kSPssEsEAENVDAQPQSTVRPEEIPPIPENR)

   Ion         m/z   Intensity
   a15+++    667.92       11.4
   y12++     724.22       11.0
   y7+       822.49       12.5
   a22+++    922.61       13.8
   b22+++    931.76       10.3
   a6+       977.32       18.1
   y17++     980.19       31.2
   a15++    1001.12       20.1
   a24+++   1009.00       10.4
   b15++    1014.90       20.3
   b24+++   1018.06       15.9
   y18++    1044.04       10.3
   y19++    1079.81       27.8
   a27+++   1110.72       13.0
   b27+++   1120.01       11.7
   y20++    1137.30       29.2
   b18++    1171.32       12.0
   y21++    1186.55       14.6
   y22++    1243.78       24.3
   y23++    1308.40       22.8
   y24++    1343.92       18.7
   a9+      1344.85       14.6
   y25++    1408.30       14.3
   y26++    1491.76       11.4
```

```
783. MORG-339_phosphopeptide_TMT_14%ACN_032017_OT.18373.18373.2 (dLLSDLQDIsDSER)

   Ion         m/z   Intensity
   a1+       317.34        8.2
   y3+       391.27        8.1
   y4+       506.21        7.9
   b3+       571.49        7.0
   b4+       658.34        9.8
   b5+       773.45       14.4
   y6+       786.49        7.0
   b6+       886.51        8.4
   y7+       901.13       20.0
   a7+       986.34        6.7
   b7+      1014.64        8.2
   b8+      1129.56       19.2
   y9+      1142.30        9.4
   b9+      1242.77       13.9
   y11+     1344.53        6.4
   b11+     1524.61        9.5
   y13+     1570.70        8.0
```

```
784. MORG-339_phosphopeptide_TMT_30%ACN_032017_OT.19696.19696.3 (lLkAPTLAELDsSDsEEEk)

   Ion         m/z   Intensity
   b4++      442.81        9.7
   b7++      598.89       10.5
   a8++      620.19        8.6
   b9++      698.68       13.3
   b10++     755.05       10.5
   b11++     812.69       24.8
   b3+       813.59       14.6
   a12++     881.73       12.9
   b4+       884.70        8.7
   y5+       930.49       26.7
   b13++     939.17       22.4
   a5+       953.84       22.2
   b14++     997.06       18.2
   y15++    1020.20       12.4
   y16++    1055.16       17.0
   a15++    1066.50       10.3
   b6+      1082.93       10.5
   a16++    1130.84        9.6
   b16++    1145.32       12.6
   b18++    1274.08        9.3
   y8+      1299.51       10.2
   y9+      1414.56        8.8
```

```
785. MORG-339_phosphopeptide_TMT_22%ACN_032017_OT.8455.8455.4 (tASEsISNLSEAGSVkkGER)

   Ion         m/z   Intensity
   a8+++     357.99       24.6
   a8++      536.33       16.9
   y12+++    573.95       18.5
   y6++      588.35       15.5
   b9++      607.07       23.4
   y13+++    611.90       24.3
   b4+       618.71       24.8
   y7++      631.85       19.1
   y14+++    640.89       35.9
   y8++      659.90       32.6
   y15+++    678.09       44.6
   a16+++    691.35       45.3
   y9++      695.09       74.1
   b11++     714.49       32.6
   y16+++    734.45       16.5
   b12++     750.74       21.1
   y10++     760.25       32.9
   a13++     765.31       15.1
   y17+++    777.56       23.1
   b13++     778.79       23.8
   b5+       785.23       20.4
   y11++     803.64       56.7
   y18+++    806.26       20.4
   a14++     808.12       20.3
   b17+++    819.98       21.9
   y12++     860.17       27.2
   b19+++    882.05       27.0
   y13++     917.20       26.9
   y14++     960.81       25.1
   b16++    1050.82       37.6
```

```
786. MORG-339_phosphopeptide_TMT_6%ACN_032017_OT.14130.14130.3 (fEDEGAGFEESsETGDYEEk)

   Ion         m/z   Intensity
   y3++      317.19        6.3
   y1+       376.23        8.0
   y4++      399.20        5.5
   y2+       505.36       10.5
   b8++      541.90        7.2
   y3+       634.37       15.3
   b10++     670.36       10.0
   y9++      684.13        6.1
   a11++     700.68        8.9
   b11++     714.64        9.6
   a4+       722.35        5.4
   y10++     727.63        8.2
   b4+       750.43        8.5
   b12++     797.40       15.3
   y12++     856.21       11.4
   b6+       878.29       13.0
   a14++     899.28       23.7
   a7+       907.13        8.5
   b14++     913.26       21.9
   b7+       935.41        7.0
   b15++     941.50        8.3
   y6+       969.60       21.8
   y16++    1022.82       10.2
   y7+      1070.57       11.1
   b17++    1080.63       19.4
   b8+      1082.58       11.9
   y18++    1144.43       10.1
   y8+      1199.69        6.3
   b19++    1209.78       18.6
   b9+      1211.59        8.2
   b10+     1340.56       13.6
   b11+     1427.74       10.0
   b13+     1723.80        5.5
```

```
787. MORG-339_phosphopeptide_TMT_22%ACN_032017_OT.16996.16996.4 (sLSPLSGTtDTkAEsPAGRVSDESVLPLAQk)

   Ion         m/z   Intensity
   a6++      393.65       12.6
   y7++      499.41       21.3
   y2+       504.43        9.9
   y3+       575.50       12.8
   y4+       688.58       10.4
   y5+       785.53       36.4
   a6+       786.58       25.7
   y14++     864.95       17.8
   a21+++    878.18       11.7
   b21+++    887.76       11.3
   y6+       898.66       17.8
   y15++     899.86       15.9
   y21+++    908.31       16.7
   y22+++    945.84       12.2
   y16++     948.49       29.3
   a14++     950.24       26.9
   a23+++    959.77       23.7
   b14++     964.05       10.4
   b23+++    969.55       45.6
   b8+       972.78       13.8
   a24+++    988.44        9.9
   b25+++   1031.41       21.9
   b26+++   1069.26       55.5
   y8+      1084.70       12.5
   b27+++   1101.41       11.2
   a17++    1117.58       11.0
   y27+++   1126.18       16.5
   y19++    1131.99       38.1
   b28+++   1138.97       10.5
   b18++    1159.74       11.8
   b29+++   1163.04       20.8
   b30+++   1205.76       13.7
   y9+      1213.80       17.9
   b10+     1268.62       10.9
   b26++    1602.77       10.6
```

```
788. MORG-339_phosphopeptide_TMT_10%ACN_032017_OT.8224.8224.2 (sEPAVGPLR)

   Ion         m/z   Intensity
   b1+       397.23       10.1
   y4+       442.39       15.6
   b2+       526.30        8.9
   y5+       541.76        8.3
   a3+       595.03       15.8
   b4+       694.27       10.1
   y7+       709.52       17.1
   a5+       765.40        8.2
   b5+       793.31       18.5
   y8+       838.51       14.2
   b6+       850.41       10.8
   b7+       947.43        6.3
   b8+      1060.61        8.2
```

```
789. MORG-339_phosphopeptide_TMT_10%ACN_032017_OT.6940.6940.3 (aAsPQEVTPQADEk)

   Ion         m/z   Intensity
   y1+       376.36       18.1
   b6++      447.38       22.1
   y6++      458.98       55.3
   b7++      496.93       30.8
   y2+       505.34       15.1
   y7++      509.53       31.1
   a8++      533.25       10.3
   b8++      547.58       49.0
   y8++      558.96       15.4
   y3+       620.40       24.6
   y9++      623.82       18.2
   b4+       636.56       10.9
   a10++     645.78       28.7
   b10++     659.65       21.4
   y4+       691.46       14.7
   b11++     695.60       31.4
   y11++     735.73       10.3
   a5+       736.58       10.0
   a12++     738.92       11.2
   b12++     753.17       10.9
   b5+       764.33       14.1
   b13++     817.30       12.1
   y5+       819.76       10.4
   b6+       893.43       20.9
   y6+       916.51       37.3
   b7+       992.46       27.1
   y7+      1017.58       20.6
   b8+      1093.50       31.5
```

```
790. MORG-339_phosphopeptide_TMT_14%ACN_032017_OT.8994.8994.2 (sDIsPLTPR)

   Ion         m/z   Intensity
   y2+       272.13       12.5
   b1+       317.19       13.1
   y3+       373.32        8.7
   b2+       432.37       13.7
   y4+       486.04        6.9
   b3+       545.30       11.0
   y5+       583.37       23.7
   b4+       712.19        8.8
   y7+       863.45       11.7
   b6+       922.59       14.9
   y8+       978.31        6.7
```

```
791. MORG-339_phosphopeptide_TMT_14%ACN_032017_OT.13104.13104.4 (nLAkPGVtsTSDsEDEDDQEGEk)

   Ion         m/z   Intensity
   b5+++     328.15       11.4
   y1+       376.37       25.6
   y5++      410.46       19.3
   a2+       429.46       18.5
   b4++      443.22       44.9
   y6++      467.75       13.2
   y10+++    475.19       12.4
   y2+       505.39       16.4
   y7++      525.35       13.0
   b3+       528.51       11.6
   y3+       562.48       44.6
   b7++      570.04       17.8
   a12+++    587.77       11.9
   y9++      647.32       11.3
   y4+       691.45       23.4
   a16+++    767.95       20.1
   b16+++    777.82       12.2
   y18+++    799.49       12.2
   b17+++    815.23       19.6
   y5+       819.78       44.1
   a11++     823.78       13.3
   b4+       885.89       16.5
   a19+++    887.66       14.5
   b12++     895.49       18.2
   b19+++    896.95       15.2
   a20+++    930.76       17.5
   y6+       933.99       26.3
   b20+++    939.76       67.5
   y20+++    951.19       16.6
   b21+++    959.21       24.7
   a22+++    992.61       12.0
   b22+++   1002.01       25.4
   b14++    1043.55       16.2
   b15++    1100.84       12.7
   y16++    1120.39       11.9
   b16++    1165.67       27.3
   y18++    1199.11       11.9
   b17++    1222.76       20.4
   y19++    1247.43       15.1
   b18++    1280.60       32.8
   b19++    1344.64       29.7
```

```
792. MORG-339_phosphopeptide_TMT_10%ACN_032017_OT.7544.7544.3 (sVENLPEcGITHEQR)

   Ion         m/z   Intensity
   y2+       303.31       13.6
   y6++      392.47        9.5
   b1+       397.25       11.7
   y3+       432.14       18.2
   a2+       468.25       10.5
   y8++      500.99       23.6
   y4+       569.24       23.4
   a8++      605.84       10.8
   y10++     613.88       43.3
   b3+       625.72       20.0
   a9++      633.92       10.1
   b9++      648.67       16.0
   y5+       670.33       40.4
   b4+       739.35       24.2
   a11++     741.27       14.9
   b11++     755.57       19.1
   y6+       783.44       18.3
   b12++     823.83       12.1
   a5+       824.54       12.2
   y7+       840.48       26.0
   y14++     841.38       18.5
   b5+       852.49       44.2
   a13++     874.34        9.5
   b13++     888.52       29.1
   b14++     952.54       16.0
   y8+      1000.35       22.6
   b7+      1078.52       18.9
   y9+      1129.47        9.6
   y10+     1226.60       19.1
   b9+      1295.53        9.7
```

```
793. MORG-339_phosphopeptide_TMT_4%ACN_032017_OT.7264.7264.3 (eQEssGEEDNDLSPEEREk)

   Ion         m/z   Intensity
   a1+       331.38        6.0
   y1+       376.20        7.1
   b2+       487.06        5.2
   y6++      509.09       13.0
   y7++      552.56       13.8
   y8++      608.76       14.1
   b3+       616.30        4.7
   y9++      666.41        6.4
   y10++     723.43       13.7
   a4+       755.53        7.9
   y11++     780.98        8.3
   b11++     805.23        4.7
   y12++     845.28       13.9
   a12++     847.91        9.9
   b12++     862.07        4.4
   a13++     890.89       13.3
   y13++     910.04       66.6
   y5+       919.84        9.5
   a5+       922.55       13.0
   y6+      1016.50        7.4
   b15++    1018.16        6.5
   y15++    1022.44        9.9
   a7+      1108.50        6.1
   b18++    1225.92        5.6
   b9+      1380.75        4.3
   b11+     1609.43        5.3
   b12+     1722.61        8.8
   b13+     1809.93        5.0
```

```
794. MORG-339_phosphopeptide_TMT_12%ACN_032017_OT.5654.5654.3 (lGHSEDsSEVLSR)

   Ion         m/z   Intensity
   b5++      377.24        7.4
   b2+       400.38        6.4
   b6++      434.90       12.6
   y4+       474.31        8.0
   y8++      486.87        9.1
   a7++      503.90       10.4
   a3+       509.43       22.6
   b7++      518.67        8.7
   b3+       537.07       30.3
   a8++      547.82       14.4
   y9++      551.25       30.9
   b8++      561.24       16.9
   y5+       603.51        9.0
   b9++      626.37        7.6
   a10++     661.68        6.8
   b10++     675.55       10.8
   y6+       690.48        9.0
   b11++     732.32        8.9
   b5+       753.36        7.0
   y7+       857.47        7.6
```

```
795. MORG-339_phosphopeptide_TMT_8%ACN_032017_OT.8357.8357.3 (ssSEGGDAGNDTRNTTADLQkAGEGT)

   Ion         m/z   Intensity
   b3+       651.34        9.5
   b4+       779.98        7.4
   y6+       791.47        8.3
   b5+       837.34        7.3
   a14++     855.69        7.5
   a6+       866.19        8.6
   y15++     896.77       36.2
   b15++     920.08        8.2
   y16++     954.30        7.5
   a16++     956.07        8.7
   b16++     970.69        8.8
   a7+       981.30       13.9
   a17++     992.24        8.1
   b17++    1005.70       15.0
   b7+      1008.95       34.5
   y17++    1010.69       22.6
   y8+      1032.72       28.1
   y18++    1039.10       15.4
   y19++    1075.27       29.6
   b8+      1080.46        8.6
   y20++    1132.83       11.5
   b9+      1137.20        7.4
   y9+      1147.61        8.4
   y21++    1161.19       11.5
   b20++    1184.24        8.2
   y22++    1189.92       19.7
   y10+     1218.44        7.7
   b10+     1251.35        8.0
   y11+     1319.85        8.0
   b11+     1366.52       13.6
   y25++    1381.53        8.0
```

```
796. MORG-339_phosphopeptide_TMT_4%ACN_032017_OT.5358.5358.2 (eVsDDEAEEk)

   Ion         m/z   Intensity
   b2+       458.41        4.5
   y2+       505.34        4.9
   y3+       634.47        6.3
   a4+       712.53        7.3
   b4+       740.40        9.5
   a5+       827.35       16.4
   y5+       834.50       14.7
   y6+       949.45       11.8
   b6+       984.61        6.3
   b7+      1055.50        6.1
   y7+      1064.58        9.0
   b8+      1184.57        6.7
   y8+      1231.54        5.8
   b9+      1313.55        5.8
   y9+      1330.70        4.9
```

```
797. MORG-339_phosphopeptide_TMT_8%ACN_032017_OT.13421.13421.3 (iLsDsEDcEEk)

   Ion         m/z   Intensity
   y3++      317.55       10.2
   y1+       376.27       24.5
   y4++      397.85       13.8
   y5++      455.13        5.9
   b2+       456.22        9.8
   y2+       505.32       26.9
   y6++      520.07        6.9
   a7++      560.97        7.2
   b7++      575.30        9.2
   a3+       595.22        6.6
   y7++      603.57        9.3
   b3+       623.72        7.5
   y3+       634.39       15.2
   a8++      640.81        4.7
   b8++      655.21        7.4
   a9++      705.84        6.4
   a4+       710.85        8.8
   b9++      719.84       21.4
   b4+       738.41        7.2
   b10++     784.43       22.1
   y4+       794.42        6.2
   b6+      1034.38       13.0
   b7+      1149.47       15.9
   b8+      1309.50        9.4
```

```
798. MORG-339_phosphopeptide_TMT_10%ACN_032017_OT.12199.12199.3 (gAEDAPDADTAIINAEGsQVNAEEkk)

   Ion         m/z   Intensity
   b3+       487.32        9.7
   y5++      531.87        8.2
   y6++      588.97        8.5
   b4+       602.36       15.9
   y8++      702.74       16.3
   b7+       885.42       14.3
   y4+       991.61       11.4
   b18++    1004.73       14.7
   y14++    1028.48       15.9
   a19++    1054.76        8.4
   y5+      1062.69       12.9
   b9+      1071.29       12.7
   a20++    1103.53       16.6
   b20++    1118.50       23.4
   y16++    1120.42       19.5
   y6+      1176.66       12.0
   y18++    1228.62       11.9
   b11+     1243.45       12.9
   a23++    1261.09        8.8
   y19++    1263.90       15.3
   y7+      1276.12       11.4
   y20++    1321.28        8.4
   b12+     1356.87       21.0
   y21++    1369.92       11.8
   y22++    1404.93        9.0
   b13+     1470.03        9.8
   b25++    1518.40        9.3
   y9+      1571.08        8.3
   b14+     1583.94       10.2
   b16+     1783.87       10.0
```

```
799. MORG-339_phosphopeptide_TMT_16%ACN_032017_OT.8694.8694.3 (sRsDNALNLVTER)

   Ion         m/z   Intensity
   a3++      306.88        9.7
   b1+       317.26        8.2
   b3++      320.66        7.9
   b4++      378.11       25.0
   y3+       405.29       12.0
   b5++      435.28       25.3
   a6++      456.85       11.7
   b6++      470.91       33.5
   y4+       504.33       14.5
   a7++      513.42       13.6
   b7++      527.18       20.9
   a8++      570.13       12.1
   b8++      584.07       27.5
   y5+       617.48       19.8
   b9++      641.05       25.6
   b10++     690.54       12.9
   y6+       731.47       27.6
   b12++     805.48       13.9
   y7+       844.49       25.1
   y8+       915.70       16.0
   y9+      1029.58       18.9
```

```
800. MORG-339_phosphopeptide_TMT_8%ACN_032017_OT.7533.7533.3 (sTPEsGEsDkESVGSSSLGNEGSR)

   Ion         m/z   Intensity
   y6+       619.26        8.1
   b4+       644.41        6.5
   y15++     862.11       17.0
   a11++     869.67        7.7
   b11++     883.50       10.5
   y9+       906.84        8.4
   y16++     919.65       11.1
   a13++     962.78       45.1
   a7+       969.57       43.3
   b13++     976.47       13.0
   a14++     990.97        7.8
   y10+      993.13        8.5
   a15++    1034.12        6.9
   y11+     1050.58       10.0
   y19++    1096.35       11.2
   y12+     1149.69        8.9
   y20++    1179.81        7.0
   y13+     1236.58       11.8
   y21++    1244.07        6.4
   b20++    1277.85       10.0
   b9+      1279.44        6.6
   y22++    1292.93       10.2
   b21++    1342.31        7.6
   y23++    1343.02        5.8
   y14+     1365.62        6.0
```

```
801. MORG-339_phosphopeptide_TMT_6%ACN_032017_OT.8509.8509.3 (sRVNssGESGGESDEFLQSR)

   Ion         m/z   Intensity
   b1+       317.19        9.1
   y3+       390.30        6.9
   a5++      412.94       11.3
   y8++      491.59        9.0
   a8++      589.93       11.8
   b8++      604.10       17.7
   b9++      647.48        7.1
   y5+       650.41       11.3
   y13++     720.43       13.7
   b12++     768.86       36.3
   y6+       779.49       15.6
   b13++     811.88        9.0
   a5+       825.18       13.9
   b14++     869.75        7.2
   b15++     934.11        8.5
   y8+       981.63       13.6
   a16++     994.05        9.1
   b16++    1008.22       12.4
   b17++    1064.84       11.2
   a18++    1114.34        7.1
   y10+     1167.64        8.0
   b19++    1172.35        8.4
   y11+     1224.61       11.3
   b9+      1293.87       10.1
   y12+     1311.50       20.1
```

```
802. MORG-339_phosphopeptide_TMT_10%ACN_032017_OT.3426.3426.4 (aGGRRVIENTDGsEEEMDAR)

   Ion         m/z   Intensity
   y4++      246.18       27.2
   b8+++     357.00       12.4
   b6++      413.89        9.0
   b11+++    467.11       38.5
   b7++      470.16        9.9
   b12+++    486.24        9.5
   y4+       492.28       22.4
   b8++      535.02       11.7
   y9++      552.01       27.8
   b4+       571.02       18.7
   a14+++    575.70       13.5
   a9++      577.53       24.0
   y15+++    592.00       10.4
   y10++     610.09       30.6
   a15+++    618.70       18.2
   y5+       621.22       10.6
   b16+++    670.45       18.4
   y17+++    696.64       13.6
   a5+       699.25       12.9
   b11++     700.12       21.2
   a12++     714.36       28.3
   y18+++    715.20       13.4
   a18+++    743.82       16.1
   y6+       750.25       18.0
   b18+++    752.98       69.3
   y7+       879.38       13.1
   b16++    1005.37       13.8
   y9+      1103.39       14.3
```

```
803. MORG-339_phosphopeptide_TMT_20%ACN_032017_OT.13938.13938.4 (kADNPTPQTPGTSSLSQLGAYGDSEDSDs)

   Ion         m/z   Intensity
   a5++      364.60       16.3
   y19+++    652.25       22.4
   b11++     669.15       14.2
   a18+++    675.25       16.6
   a12++     705.81       19.6
   a20+++    717.91       30.6
   b12++     719.22       16.9
   y14++     755.53       29.0
   y22+++    761.65       94.4
   b13++     762.49       25.8
   a14++     792.27       20.1
   y23+++    793.24       33.7
   y15++     812.74       14.3
   y24+++    827.74       20.8
   a6+       828.83       23.2
   y7+       834.19       34.3
   b23+++    839.59       17.3
   a15++     849.36       42.5
   y16++     855.50       21.3
   b6+       856.23       20.7
   y8+       891.25       36.7
   a16++     892.25       20.6
   b7+       953.33       15.4
   a17++     956.35       25.8
   y28+++    959.82       14.8
   a27+++    969.28       15.2
   y19++     978.07       23.1
   a18++    1013.04       14.2
   b28+++   1017.34       15.5
   b18++    1027.45       19.5
   y9+      1054.33       34.4
   b19++    1055.32       25.8
   y10+     1125.46       24.8
   y22++    1141.95       23.3
   b9+      1182.39       32.3
   y12+     1295.45       21.9
   y14+     1510.47       18.9
```

```
804. MORG-339_phosphopeptide_TMT_8%ACN_032017_OT.6030.6030.4 (sEDELRPDVDEHTQk)

   Ion         m/z   Intensity
   y7+++     362.62        8.5
   y1+       376.30       14.2
   b1+       397.18       11.2
   a5++      428.42       13.1
   y9+++     433.28        8.3
   y5++      436.28       28.4
   y2+       504.35       23.3
   b6++      520.34       11.0
   y11+++    523.10       31.4
   b2+       526.22        8.3
   y7++      543.08       16.2
   a7++      554.98       17.0
   y12+++    566.01       28.7
   b7++      569.01       13.4
   b12+++    578.02       14.9
   a13+++    602.29       10.0
   y13+++    604.62       11.5
   y3+       605.47       23.4
   a8++      611.89       20.2
   b8++      626.37       19.5
   b3+       641.30       16.9
   y14+++    647.57       18.9
   y9++      649.50       14.7
   b14+++    654.44        9.2
   b9++      676.21        9.9
   b10++     733.46       65.1
   y4+       742.31       11.5
   b4+       770.31       18.0
   y11++     783.84        7.8
   b11++     797.73       15.1
   b12++     866.34       14.1
   y13++     906.35        7.0
```

```
805. MORG-339_phosphopeptide_TMT_8%ACN_032017_OT.8605.8605.4 (eAAEAEPAEPSSPAAEAEGASASSTSsPk)

   Ion         m/z   Intensity
   y7+++     335.16       14.5
   b6++      415.23       21.4
   y2+       473.36       32.5
   b8++      499.96       10.9
   b3+       500.85       11.0
   b9++      564.13       19.7
   b4+       630.50       14.5
   y3+       640.38       11.1
   b11++     656.57       10.0
   b20+++    698.96       10.6
   b5+       701.34       14.6
   y20+++    710.38       18.9
   y4+       727.44       21.0
   b21+++    728.40       12.9
   b13++     748.80       12.7
   b22+++    752.14       11.6
   y21+++    753.36       14.4
   a23+++    771.79       21.0
   y22+++    777.09       16.2
   b23+++    780.70       15.7
   a24+++    799.90       27.1
   a6+       802.00       19.2
   a15++     805.64       20.1
   y5+       828.22       12.2
   b6+       830.40       50.2
   y15++     845.28       11.1
   a16++     870.38       10.9
   b16++     884.40       10.9
   b17++     919.21       18.1
   y17++     928.79       16.3
   y18++     972.64       12.6
   b8+       998.44       10.8
   y7+      1002.53       11.8
   y20++    1064.74       55.9
   b22++    1126.57       13.3
   b9+      1127.60       32.1
   y21++    1128.62       28.5
   y22++    1164.40       12.0
   y23++    1213.38       16.6
   b24++    1214.09       11.5
```

```
806. MORG-339_phosphopeptide_TMT_8%ACN_032017_OT.14810.14810.4 (dDDDIDLFGsDDEEESEEAkk)

   Ion         m/z   Intensity
   a1+       317.23        8.6
   y2++      367.46       14.5
   y1+       376.28       16.9
   y3++      402.55       16.1
   a9+++     403.51        8.8
   b2+       459.78       10.4
   y4++      467.28       14.4
   b10+++    468.11       10.1
   b7++      516.23       20.5
   y5++      531.99       19.2
   a8++      575.55       22.6
   b8++      590.01       28.0
   b9++      618.40       17.1
   y7++      640.07       21.4
   a15+++    664.51       17.8
   b4+       690.40       13.3
   a16+++    693.39        6.9
   y8++      704.56       27.6
   y14+++    713.62       46.5
   a11++     745.36       10.3
   y15+++    751.23       30.4
   b11++     759.70       18.4
   y9++      768.68       90.2
   a5+       775.23       25.6
   a18+++    779.82        7.8
   b5+       803.43       22.0
   y3+       804.47       14.6
   y17+++    826.61       34.1
   b13++     881.58       11.5
   y11++     884.13       20.7
   b6+       918.40       37.8
   a14++     932.06       32.2
   y4+       933.13        7.7
   b14++     945.64        7.4
   y12++     967.67       28.6
   a15++     996.15       60.6
   b15++    1010.65       11.8
   b7+      1031.47       39.0
   b16++    1054.19        7.0
   y14++    1069.71       37.7
   y15++    1126.36        7.5
   a18++    1169.32        6.9
   b8+      1178.61       24.6
   y16++    1183.27       10.3
   b19++    1218.73       17.3
   b13+     1761.76        7.1
```

```
807. MORG-339_phosphopeptide_TMT_20%ACN_032017_OT.15206.15206.2 (qADLSFSsPVEMk)

   Ion         m/z   Intensity
   y2+       507.36       10.5
   b3+       544.37       17.7
   y3+       636.42        8.8
   b4+       657.45        8.9
   y5+       832.61       27.3
   a6+       863.34       11.8
   a7+       950.56       25.8
   b7+       978.35       12.5
   y7+      1086.47       13.0
   b8+      1145.79        9.5
   y8+      1233.71        8.8
   b10+     1341.88        9.5
   y10+     1433.70       12.4
   b12+     1601.78       10.2
   y12+     1619.81        9.0
```

```
808. MORG-339_phosphopeptide_TMT_16%ACN_032017_OT.5383.5383.3 (sGkNSQEDsEDsEEkDVk)

   Ion         m/z   Intensity
   y2+       475.44       10.7
   y6++      603.66        9.6
   b8++      653.11       12.8
   y7++      686.93       13.3
   b3+       731.85        7.0
   y8++      744.53        6.7
   b10++     800.70        6.8
   y9++      808.67        7.1
   a4+       817.96       10.3
   b11++     858.58        9.7
   y10++     892.72        9.8
   b12++     942.18        7.2
   y11++     950.06        8.3
   a13++     992.09       31.5
   b13++    1006.72       23.7
   a6+      1032.54       10.1
   y15++    1178.59        6.5
   b8+      1304.68       14.2
   b16++    1306.79       11.2
   b11+     1715.90        7.5
```

```
809. MORG-339_phosphopeptide_TMT_8%ACN_032017_OT.4833.4833.3 (sVSTPSEAGSQDsGDGAVGSR)

   Ion         m/z   Intensity
   y2+       262.10        9.1
   b1+       317.21        9.9
   y3+       319.15       26.9
   b2+       416.40       10.0
   y4+       418.27       20.6
   y5+       489.35        9.4
   b8++      494.87       12.1
   b9++      523.36       12.9
   y6+       546.26       12.8
   b4+       604.41       24.5
   y12++     608.42       18.1
   y13++     636.77       16.5
   y14++     672.66       16.6
   b12++     687.94       26.3
   b5+       701.16       11.0
   y8+       718.44       15.2
   y15++     737.01       25.8
   b6+       788.35        8.9
   y17++     829.18       12.7
   b16++     886.80       13.4
   b7+       917.55       17.8
   b17++     922.25       35.4
   y19++     923.12       24.2
   a18++     957.56        9.1
   b18++     971.61       50.9
   y20++     972.44       21.6
   b8+       988.51       21.3
   y10+     1000.42       16.4
   a9+      1017.91       10.2
   b9+      1045.46       16.7
   b10+     1132.61       10.5
   b11+     1261.06        9.2
   y13+     1272.53       13.7
```

```
810. MORG-339_phosphopeptide_TMT_8%ACN_032017_OT.4688.4688.3 (rAsQSsLESSTGPSYSRS)

   Ion         m/z   Intensity
   y3+       349.13        7.3
   a5++      405.92        6.8
   b5++      420.23        7.1
   y4+       512.72        8.7
   y5+       599.14        7.6
   a8++      610.43        9.0
   b8++      624.91       25.3
   a9++      654.47       10.8
   b9++      668.13       13.0
   y6+       696.31       19.1
   a10++     698.20       10.3
   b10++     711.50       15.6
   y13++     719.05       13.2
   a4+       724.28       20.7
   a11++     748.28       20.8
   y7+       753.46       14.4
   b12++     790.91        9.6
   y8+       854.37       17.0
   y9+       941.48       14.8
   b15++     964.43        9.0
   y10+     1028.48       18.9
   y11+     1157.51       12.6
   y12+     1270.65        9.1
```

```
811. MORG-339_phosphopeptide_TMT_4%ACN_032017_OT.5313.5313.4 (lVREDENDAsDDEDDDEkR)

   Ion         m/z   Intensity
   y2++      266.90       10.9
   y3++      331.41       13.2
   y4++      388.74       15.1
   b5++      421.91       21.8
   b2+       442.36       13.0
   y5++      446.36       25.7
   b6++      486.30        9.6
   y6++      503.74       12.2
   b7++      543.56       12.0
   b12+++    557.39        6.3
   y7++      568.51       29.2
   y12+++    573.50       12.8
   b8++      601.04       35.7
   y8++      626.00       28.4
   b9++      636.62       14.7
   y14+++    654.67        9.7
   a15+++    667.20       23.7
   y9++      683.50       28.9
   y15+++    692.89       19.2
   b11++     777.54       16.5
   y11++     802.37       16.5
   b12++     835.13       16.8
   b5+       842.69        8.0
   y12++     859.49        7.3
   a18+++    868.10       16.7
   b18+++    877.27       11.3
   b13++     899.66       12.3
   b14++     956.70       14.9
   b15++    1014.33       10.6
   b16++    1071.89        8.5
   b17++    1136.38        7.3
```

```
812. MORG-339_phosphopeptide_TMT_10%ACN_032017_OT.15299.15299.3 (qLsVPASDEEDEVPAPIPR)

   Ion         m/z   Intensity
   y2+       272.26       12.1
   a7++      482.42       13.4
   y6+       650.40       48.5
   a4+       709.12        9.1
   y13++     726.97        9.4
   b4+       737.25       23.1
   b11++     741.13       12.1
   y7+       749.62        8.8
   y14++     762.61       29.3
   b12++     805.64       39.5
   a5+       806.49       16.6
   a13++     841.26       12.9
   b13++     855.16       50.1
   b15++     939.33       19.8
   b16++     987.92       11.8
   b17++    1044.49       18.8
   b9+      1236.81        8.9
   b12+     1609.81       11.7
   b13+     1708.85       20.4
```

```
813. MORG-339_phosphopeptide_TMT_4%ACN_032017_OT.5461.5461.3 (eEGtGESAGGsPARSLTR)

   Ion         m/z   Intensity
   y2+       276.30        4.1
   y7++      400.76       10.7
   y4+       476.30        7.5
   y9++      512.66        5.4
   a3+       516.97        5.8
   y10++     541.34        6.3
   y11++     576.94        9.1
   y12++     619.81        9.6
   b11++     676.45        9.8
   y13++     685.11       78.0
   a4+       698.53       22.7
   y6+       703.37        9.9
   a13++     746.02        5.4
   b13++     759.80        5.0
   a14++     824.73        6.5
   y16++     832.56        5.5
   b14++     838.53       10.2
   b17++     989.00        5.3
   b7+       999.55        4.7
   b9+      1127.46        4.2
   b13+     1519.75        3.9
```

```
814. MORG-339_phosphopeptide_TMT_12%ACN_032017_OT.12827.12827.2 (sDNALNLVTER)

   Ion         m/z   Intensity
   b1+       397.12        8.7
   y3+       405.10        6.7
   y4+       504.22        9.6
   b2+       512.27       12.2
   y5+       617.38        6.9
   b3+       626.25        6.3
   b4+       697.28       10.1
   y6+       731.02       37.7
   b5+       810.48        8.1
   y7+       844.56       10.2
   y8+       915.49       11.2
   b6+       924.49        7.2
   y9+      1029.54       19.0
   b7+      1037.60        9.1
   b8+      1136.59       10.4
   y10+     1144.51        9.2
   b10+     1366.59        6.6
```

```
815. MORG-339_phosphopeptide_TMT_10%ACN_032017_OT.13183.13183.4 (gATPAEDDEDkDIDLFGsDEEEEDkEAAR)

   Ion         m/z   Intensity
   y5++      402.46       32.9
   y6++      459.46       18.7
   y8++      589.03       20.5
   y9++      653.55       20.3
   y10++     717.83       35.9
   b6+       756.53       17.7
   y11++     775.53       33.8
   b11++     794.65       19.2
   y5+       803.37       15.5
   b12++     852.22       44.6
   b7+       871.52       31.2
   b13++     908.32       18.3
   y19+++    912.31       41.4
   y6+       918.19       21.5
   y20+++    950.82       15.3
   a14++     951.77       18.9
   a8+       958.75       31.9
   y14++     961.13       49.1
   b14++     966.09       40.4
   b22+++    973.37       32.7
   y15++    1017.68       35.6
   b15++    1022.37       20.2
   y22+++   1032.23       28.2
   y23+++   1070.20       15.3
   a16++    1082.42       28.2
   b16++    1096.26       24.8
   b9+      1115.63       18.4
   b17++    1124.51       17.6
   y17++    1131.78       23.9
   y25+++   1136.85       27.8
   a25+++   1164.58       14.4
   y26+++   1169.55       19.4
   b18++    1207.79       21.5
   b26+++   1216.91       43.3
   a27+++   1230.56       38.5
   b27+++   1240.59       18.3
   b28+++   1264.11       24.8
   b19++    1265.36       21.5
   b20++    1330.45       14.0
   b23++    1523.42       15.1
```

```
816. MORG-339_phosphopeptide_TMT_25%ACN_032017_OT.20207.20207.2 (tLSGPVTGLLATAR)

   Ion         m/z   Intensity
   b1+       411.25       10.6
   y6+       644.15       14.6
   y7+       701.39       17.6
   b5+       765.77        9.4
   y8+       802.43       25.0
   b6+       864.20       11.5
   a8+       994.50       10.2
   y10+      998.59       14.7
   y11+     1055.68       21.9
   b9+      1136.01        9.2
   y12+     1142.62       18.4
   y13+     1255.80       17.2
```

```
817. MORG-339_phosphopeptide_TMT_16%ACN_032017_OT.23602.23602.3 (eLALssPEDLTQDFEELkR)

   Ion         m/z   Intensity
   y6++      526.13        9.1
   b3+       543.35        9.7
   y3+       645.57        5.7
   y8++      647.53        6.2
   b4+       656.58        5.1
   y9++      697.86        5.4
   y10++     754.50       11.5
   a5+       795.38        6.8
   y12++     876.00       21.3
   y5+       903.44        6.0
   y13++     925.30       24.7
   y17++    1183.57        6.9
   b9+      1331.47        8.4
   b18++    1332.55        6.0
   y9+      1394.85        6.2
```

```
818. MORG-339_phosphopeptide_TMT_6%ACN_032017_OT.7106.7106.2 (sGsIIGSR)

   Ion         m/z   Intensity
   y1+       175.20        3.9
   y3+       319.13        8.7
   b1+       397.18        4.7
   y4+       432.42       11.9
   b2+       454.23        5.4
   y5+       545.40       10.5
   b3+       621.05        4.6
   b4+       734.14        5.1
   y7+       769.36        5.1
   a5+       819.27        5.4
   b5+       847.39       13.8
   b6+       904.32        5.7
```

```
819. MORG-339_phosphopeptide_TMT_12%ACN_032017_OT.12966.12966.3 (kQSQIQNQQSsDLDsDVEDY)

   Ion         m/z   Intensity
   b1+       358.15       11.7
   y3+       426.13        9.3
   y6+       807.17        9.6
   a13++     883.88       24.5
   y14++     887.77       10.1
   a6+       914.44       18.1
   y7+       922.38       27.2
   a14++     941.78       10.3
   b14++     955.27       12.9
   y8+      1035.64       14.8
   b15++    1038.74        8.7
   y10+     1317.10       16.1
   y11+     1404.41       15.4
   y14+     1774.73       13.8
   y15+     1902.43        9.0
```

```
820. MORG-339_phosphopeptide_TMT_10%ACN_032017_OT.16216.16216.3 (sLRHEAEsDFDVADLPQLER)

   Ion         m/z   Intensity
   b6++      502.30        9.9
   b7++      567.15       10.9
   y5+       642.38       21.8
   b9++      707.80       18.5
   y6+       755.56       25.4
   b10++     781.38       14.0
   y13++     793.34       13.3
   b11++     839.29       27.9
   y14++     856.99       13.1
   y7+       870.56       20.1
   a12++     874.81       39.1
   b12++     888.58       36.4
   y15++     893.04       18.7
   b5+       932.67       33.6
   y8+       941.46       25.2
   b14++     981.79       27.3
   y17++    1026.23       18.3
   b15++    1038.01       23.4
   y9+      1040.67       23.8
   y10+     1155.50       12.0
   y11+     1302.80       16.9
```

```
821. MORG-339_phosphopeptide_TMT_20%ACN_032017_OT.13227.13227.3 (sLSksDSDLLTcSPTEDATMGSR)

   Ion         m/z   Intensity
   b5++      521.19       10.4
   b6++      579.00       22.0
   b7++      622.46       21.6
   b8++      679.82       19.1
   a9++      722.14       10.7
   b9++      736.41       12.9
   y7+       737.40       15.5
   b10++     793.25       23.2
   y8+       866.11       17.0
   b4+       875.06       18.3
   b12++     923.94       11.0
   a13++     952.66       11.5
   y9+       967.22      135.7
   y18++     978.47       23.3
   y10+     1064.52       26.6
   b15++    1065.58       16.5
   b16++    1130.59       10.8
   y11+     1151.55       18.4
   b17++    1188.37       22.9
   a7+      1216.03       12.0
   y21++    1284.43       12.5
   y12+     1311.65       14.8
   y22++    1341.09       10.3
   y13+     1412.65       26.5
   y14+     1525.68       10.9
   y15+     1638.91       20.9
   y17+     1840.83       17.2
```

```
822. MORG-339_phosphopeptide_TMT_22%ACN_032017_OT.6931.6931.4 (vRGWsPPPEVRR)

   Ion         m/z   Intensity
   y1+       175.19       16.8
   y3++      215.72       47.4
   b4+++     243.41        6.7
   b3++      271.76        9.8
   a5+++     289.61        8.1
   a6+++     321.80        6.8
   b1+       328.84       14.0
   b6+++     331.27        7.1
   y8+++     340.08        6.5
   a4++      350.72        6.1
   a7+++     354.84       10.3
   b7+++     363.42        5.4
   b4++      364.73       11.1
   y6++      377.38       22.3
   a8+++     386.98        6.1
   b8+++     396.39       10.1
   y9+++     401.40        9.0
   y10+++    421.14       22.2
   y7++      425.89       91.0
   y3+       430.36        9.8
   b5++      448.11       15.4
   b6++      496.46       11.0
   y8++      509.36        8.3
   b11+++    524.45       18.2
   a7++      531.19        8.0
   b3+       542.49        6.5
   b7++      545.00       15.1
   a9++      644.30       15.0
   y5+       656.51       11.0
   b9++      658.59       66.5
   b5+       895.33        7.4
```

```
823. MORG-339_phosphopeptide_TMT_16%ACN_032017_OT.4998.4998.3 (rQPQIkQEPEsEEEEEEk)

   Ion         m/z   Intensity
   y2+       505.52        6.0
   b8++      733.80       13.3
   b9++      782.52        4.9
   b10++     847.21       10.2
   a11++     916.99        5.8
   a12++     980.52       15.0
   b12++     994.86        6.4
   y6+      1021.52        8.2
   y13++    1079.51        6.2
   y7+      1150.65        7.5
   a6+      1182.00        4.8
   b15++    1188.68        5.0
   y15++    1199.85        4.8
   y16++    1249.01        5.0
   b16++    1253.17        5.8
   b17++    1317.34        5.1
   y9+      1446.61        6.7
   y10+     1543.68        6.4
```

```
824. MORG-339_phosphopeptide_TMT_22%ACN_032017_OT.23152.23152.3 (aAGPGNLVLPGsLTAsLEcVk)

   Ion         m/z   Intensity
   a6++      335.26       14.4
   b2+       372.35       10.3
   y1+       376.30       37.4
   b7++      405.90       18.8
   b8++      455.38       24.2
   y2+       475.42       19.6
   b9++      511.83       15.4
   b11++     589.23       12.1
   y8++      608.72       10.2
   y3+       635.44       37.7
   y9++      665.58       10.5
   b6+       697.47       19.6
   y4+       764.48       32.5
   a14++     765.44       15.0
   y11++     777.04       47.7
   b14++     779.31       13.6
   b7+       810.57       23.1
   b15++     814.65       11.5
   y12++     826.03       84.4
   y5+       877.53       16.4
   a8+       881.68       10.3
   y13++     882.41       20.8
   b8+       909.58       62.6
   y14++     932.13       13.7
   y15++     988.50       11.6
   a18++    1005.05       10.9
   b18++    1019.71       28.0
   b9+      1022.66       91.0
   y6+      1044.57       26.4
   y16++    1045.62       17.3
   b19++    1099.46       17.6
   y7+      1115.57       19.1
   a20++    1135.22       10.1
   b20++    1149.18       32.7
   y19++    1151.26       12.7
   b11+     1176.54       20.7
   y8+      1216.56       15.2
   b13+     1456.88       10.0
   y11+     1553.77       12.3
   b14+     1557.89       12.1
   b15+     1628.86        9.6
   y12+     1650.78       22.9
   y13+     1763.89       10.0
```

```
825. MORG-339_phosphopeptide_TMT_30%ACN_032017_OT.9651.9651.4 (eLEkPmQSkPQSPVIQATAGsPk)

   Ion         m/z   Intensity
   y2+       473.36       34.0
   y4+       697.30       17.3
   b14+++    761.86       28.7
   b15+++    799.51       29.8
   y18+++    803.50       15.1
   a16+++    833.02       30.3
   b16+++    842.47      106.6
   y14++     845.03       40.8
   a17+++    856.97       30.2
   b18+++    899.86       37.6
   a10++     922.24       21.0
   b19+++    923.38       30.7
   a20+++    933.09       18.2
   y7+       940.65       15.1
   b20+++    942.54       26.4
   y20+++    954.92       20.2
   b4+       958.77       15.7
   a11++     986.70       20.9
   y21+++    997.81       16.9
   b11++    1000.76       15.6
   a5+      1027.29       14.9
   b22+++   1030.44       16.9
   b15++    1198.86       18.8
   b17++    1298.60       18.6
```

```
826. MORG-339_phosphopeptide_TMT_4%ACN_032017_OT.7281.7281.2 (eEQELQDIQsR)

   Ion         m/z   Intensity
   a1+       331.20        6.5
   b1+       359.22        8.1
   b2+       488.31        8.8
   y4+       583.33        7.7
   b3+       616.39        5.8
   y5+       698.49        5.2
   b4+       745.32       10.9
   a5+       830.63       11.2
   b5+       858.50        6.8
   y7+       939.46        8.6
   b7+      1101.49       21.6
   b8+      1214.60        5.0
   b9+      1342.68        6.8
```

```
827. MORG-339_phosphopeptide_TMT_22%ACN_032017_OT.10315.10315.3 (kALVVPEPEPDSDsNQER)

   Ion         m/z   Intensity
   y10++     628.58       20.2
   y11++     677.26       22.0
   a5+       712.73       17.6
   y12++     741.55       25.5
   b12++     746.60       18.6
   y13++     790.23       27.6
   b13++     803.60       22.4
   y6+       828.20       46.9
   b6+       837.16       44.0
   y14++     839.63       28.6
   y7+       915.38       16.3
   a7+       938.16       14.8
   y16++     946.00       16.0
   y8+      1030.54       21.7
   a17++    1058.76       17.2
   y9+      1127.32       16.9
   y10+     1256.56       17.6
   y11+     1353.53       18.4
   y12+     1482.75       22.5
   y13+     1579.74       18.4
```

```
828. MORG-339_phosphopeptide_TMT_4%ACN_032017_OT.4039.4039.4 (yRTREQEsSGEEDNDLSPEER)

   Ion         m/z   Intensity
   y4++      265.82        6.2
   y2+       304.30        8.1
   y11+++    445.29        7.3
   y8++      480.40        6.0
   b9+++     482.81        6.7
   y4+       530.29       26.3
   b6++      532.37       10.6
   y9++      537.78        5.4
   b11+++    545.23        9.1
   a7++      582.82        6.9
   b12+++    588.24        9.7
   b7++      597.00        8.7
   y5+       617.37       28.6
   b13+++    626.52       16.2
   b3+       650.58        6.2
   a14+++    655.28        9.0
   b14+++    664.35       29.9
   y17+++    677.37       21.1
   b8++      680.75       19.3
   a15+++    693.37       13.2
   y12++     695.74        9.8
   b15+++    702.70       16.4
   b9++      723.92        6.7
   a16+++    730.56       19.2
   b16+++    740.48       35.3
   b17+++    769.67       23.3
   b19+++    844.75        8.2
   y7+       845.68        8.3
   y15++     886.46        6.2
   b13++     939.19       15.6
   b15++    1053.76       10.1
   b16++    1110.26       14.5
```

```
829. MORG-339_phosphopeptide_TMT_18%ACN_032017_OT.14210.14210.3 (kGSPVSEIGWEtPPPEsPR)

   Ion         m/z   Intensity
   b2+       415.33       10.3
   b11++     700.67       10.4
   a6+       757.96       10.4
   y6+       762.40       17.4
   y12++     763.37       11.1
   a13++     825.32       21.5
   y13++     827.77       11.3
   b13++     839.11        8.1
   y7+       859.41       38.3
   y14++     871.44       24.2
   a14++     874.22       17.4
   b15++     936.80        9.5
   y8+      1040.15        8.6
   y18++    1041.24       11.7
   y11+     1412.60       12.4
```

```
830. MORG-339_phosphopeptide_TMT_22%ACN_032017_OT.13164.13164.4 (sRHSsIsPVRLPLNSSLGAELSR)

   Ion         m/z   Intensity
   b1+       317.20       12.2
   y3+       375.31       10.4
   y12+++    415.25       16.9
   y13+++    452.87        9.8
   a6++      474.86       27.8
   y5+       575.12       12.9
   b13+++    607.58       15.6
   y6+       632.39       29.2
   b14+++    645.55       18.2
   a9++      656.54       18.2
   y18+++    663.47       15.0
   a15+++    665.18       62.8
   a4+       669.59       31.4
   b9++      670.59       20.4
   b15+++    674.29       18.2
   y13++     678.45       10.1
   a16+++    694.30       34.2
   b4+       697.37       23.7
   b16+++    703.06       20.9
   a17+++    731.91       10.2
   a10++     735.25       20.7
   y7+       745.55       15.3
   b10++     748.84       14.3
   a18+++    750.95       11.8
   y14++     756.99       10.7
   a11++     791.80       13.6
   y21+++    794.36       12.1
   b11++     805.36       13.0
   y15++     806.60       11.9
   a20+++    817.59       11.0
   b20+++    827.02       13.9
   y8+       832.47       15.1
   y22+++    846.01       10.0
   a21+++    855.23       55.7
   b21+++    864.93       10.8
   b13++     910.56       14.6
   y9+       919.57       11.5
   y17++     938.68       17.7
   y18++     995.25       11.4
   y10+     1033.57       10.8
```

```
831. MORG-339_phosphopeptide_TMT_6%ACN_032017_OT.6737.6737.3 (aEDGAAPsPSSEtPk)

   Ion         m/z   Intensity
   y5++      435.78       10.6
   y2+       473.46       16.5
   y6++      478.95        9.3
   y7++      527.81       17.9
   b3+       544.99        9.7
   a10++     582.62       12.4
   b10++     596.53       12.3
   y8++      611.07       15.8
   a5+       644.89       11.1
   y3+       654.52       22.5
   y9++      660.01       68.2
   b5+       673.55       19.8
   y10++     695.37        6.0
   y11++     731.26        6.2
   b6+       744.32       52.1
   y12++     759.34        7.1
   y4+       783.37        8.7
   y5+       870.45       11.2
   y14++     881.36        5.4
   b8+      1008.66        8.0
   b10+     1192.40        8.2
```

```
832. MORG-339_phosphopeptide_TMT_12%ACN_032017_OT.4917.4917.3 (rNsSsPVSPASVPGQR)

   Ion         m/z   Intensity
   y9++      449.67        9.2
   y4+       457.25       23.5
   b5++      461.38       10.4
   b2+       500.48        9.1
   a7++      545.09       22.0
   y11++     548.00       23.6
   y5+       556.15        9.4
   b7++      559.41       19.8
   b8++      603.17       12.8
   a9++      637.36       16.8
   a3+       639.80       61.8
   y6+       643.27       24.5
   b9++      651.63       22.2
   b10++     686.84       12.4
   y7+       714.54       10.0
   y8+       811.51       40.0
   y15++     815.59        9.4
   y9+       898.49       28.6
   b5+       921.28       16.3
   y11+     1094.62       24.0
```

```
833. MORG-339_phosphopeptide_TMT_18%ACN_032017_OT.12454.12454.4 (eVVSYSkETQTPLATHQsEEDEEDEEMVEPk)

   Ion         m/z   Intensity
   y1+       376.37       11.4
   y2+       473.36       36.2
   y3+       602.47       24.8
   b12+++    603.39       13.5
   b8++      691.05       16.1
   y4+       701.51       23.4
   b17+++    787.13       20.9
   b10++     805.66       21.6
   y5+       832.52       24.5
   a18+++    833.50       16.9
   b11++     855.88       20.6
   b20+++    928.72       22.5
   y6+       961.41       30.4
   b21+++    966.99       24.7
   b14++     996.99       21.0
   b22+++   1010.28       23.0
   b15++    1047.26       24.7
   b23+++   1053.10       35.1
   a16++    1102.05       19.9
   b16++    1115.72       16.5
   y16++    1134.64       26.2
   y25+++   1156.41       11.4
   a26+++   1167.41       14.0
   b26+++   1177.04       21.0
   y26+++   1185.15       11.6
   y8+      1205.62       17.3
   y18++    1221.50       61.2
   a18++    1249.39       14.4
   b28+++   1254.35       52.3
   b29+++   1297.40       70.0
   y20++    1326.30       11.0
   b19++    1328.11       11.1
   y30+++   1334.64       16.2
   b21++    1450.04       12.8
   y10+     1463.78       15.7
```

```
834. MORG-339_phosphopeptide_TMT_8%ACN_032017_OT.1741.1741.4 (eAGsEAESSSADPGPGRkR)

   Ion         m/z   Intensity
   b4++      327.30       18.5
   y7+++     333.06       96.7
   b5++      391.67       10.4
   y5++      421.68       18.0
   b6++      427.63       10.1
   y6++      450.44       19.6
   a7++      478.04       11.7
   y12+++    482.14       14.1
   b3+       487.35       14.2
   y7++      499.02       30.1
   b13+++    509.46       11.1
   y13+++    525.03       14.3
   y14+++    548.84       27.6
   y8++      556.48       18.0
   b15+++    560.97       24.0
   a9++      565.41       11.6
   a16+++    571.03       15.7
   b9++      579.59       38.5
   b16+++    580.45       33.5
   y15+++    591.35       20.4
   y9++      592.29       40.8
   y10++     635.52       19.3
   y11++     679.14       27.9
   a12++     702.08       13.8
   y12++     722.33       21.3
   y4+       745.84       10.7
   b18+++    751.73        9.9
   b5+       783.49       11.5
   y13++     787.30       15.0
   b6+       854.49       14.6
   b7+       983.38       10.3
   b12+     1430.60       42.9
```

```
835. MORG-339_phosphopeptide_TMT_16%ACN_032017_OT.6524.6524.3 (ssPATTVTSPNSTPAk)

   Ion         m/z   Intensity
   y1+       376.23       12.2
   b5++      377.32       10.5
   y2+       447.22       12.4
   y7++      472.57       19.1
   y3+       544.41       28.9
   a10++     606.01       16.2
   y10++     616.20       13.6
   b10++     619.55       17.2
   a4+       624.31       14.6
   y4+       645.07       18.1
   b4+       651.99       25.5
   a11++     662.99       44.4
   y11++     666.05       39.1
   y12++     716.54       14.1
   b12++     720.54       12.8
   y5+       732.49       10.6
   y13++     752.76       13.2
   b5+       753.40       16.3
   a13++     756.91       17.8
   y14++     800.69       20.8
   a14++     805.55       17.9
   b6+       854.05       10.6
```

```
836. MORG-339_phosphopeptide_TMT_8%ACN_032017_OT.2249.2249.3 (aAEEARGSEtLGGGGGGAAGTREAR)

   Ion         m/z   Intensity
   y11++     501.37        9.2
   b9++      565.86        8.2
   b4+       630.52       11.8
   y7+       760.11        9.0
   a14++     784.42       10.4
   y17++     798.74        9.3
   a15++     813.37       26.8
   b15++     826.88       33.9
   y8+       831.13       30.6
   a16++     841.04       27.1
   y18++     842.14       25.5
   b16++     855.84       19.8
   b6+       857.08        8.5
   b7+       914.79        7.9
   b19++     955.36        7.7
   b20++     983.47        6.9
   y21++     984.73       19.4
   y22++    1049.22        7.1
   y23++    1113.61        7.0
   y24++    1148.74        9.1
   a23++    1162.90        8.9
   y14+     1173.86        8.3
   b23++    1176.73       15.2
```

```
837. MORG-339_phosphopeptide_TMT_4%ACN_032017_OT.10249.10249.2 (vVEEMsE)

   Ion         m/z   Intensity
   y2+       315.02       12.0
   b1+       329.38        3.1
   b2+       428.49       14.7
   y3+       446.20       11.6
   a3+       529.85        6.5
   b3+       557.11       24.8
   y4+       575.21        3.7
   b4+       686.40        9.6
   y5+       704.18       15.1
   y6+       803.32        5.3
   b5+       817.46       21.9
   b6+       984.39       13.9
```

```
838. MORG-339_phosphopeptide_TMT_10%ACN_032017_OT.11438.11438.3 (nPLGDDDAsAtASk)

   Ion         m/z   Intensity
   b1+       344.07        6.4
   b5++      363.93        5.4
   y1+       376.34       15.3
   y5++      393.74        6.3
   y2+       463.29       26.0
   y7++      512.90        5.7
   b8++      514.25        6.3
   y3+       534.53        9.6
   b3+       554.23        8.5
   y8++      570.61        5.7
   b4+       610.93       11.5
   y9++      628.05       52.2
   b10++     633.37       14.3
   y11++     713.70        8.1
   y4+       715.39       13.6
   b11++     723.72        5.3
   b5+       726.47        6.1
   a12++     745.60       10.5
   b12++     759.23        9.3
   a13++     788.87        7.4
   b13++     802.73       16.4
   y13++     819.05        6.7
   b7+       956.54        6.7
   b8+      1027.65        5.8
```

```
839. MORG-339_phosphopeptide_TMT_4%ACN_032017_OT.7365.7365.2 (gPPDFsSDEER)

   Ion         m/z   Intensity
   y2+       304.26       12.7
   y3+       433.28        8.3
   y4+       548.23       10.5
   b4+       596.33       17.2
   y5+       635.41       11.9
   a5+       715.02       34.1
   y7+       949.30       12.6
   a8+      1084.46        5.6
   b8+      1112.51        8.1
   a9+      1213.27        5.3
   b9+      1241.44        7.0
   a10+     1342.81       10.1
   b10+     1370.51        6.0
```

```
840. MORG-339_phosphopeptide_TMT_22%ACN_032017_OT.18345.18345.3 (aYsPTcsPTLGFk)

   Ion         m/z   Intensity
   a1+       273.30        8.2
   y1+       376.34       16.2
   b5++      415.28       12.9
   y6++      446.47       16.9
   b2+       464.23        8.6
   y2+       523.47       25.2
   y7++      530.24       12.1
   y3+       580.48       35.2
   a3+       603.39        8.0
   y8++      609.62        8.4
   b3+       630.82        9.1
   y9++      660.56       10.5
   a9++      664.16       22.4
   y4+       693.57        8.2
   a4+       700.43        7.6
   b10++     734.57       22.3
   b11++     763.15       24.0
   y11++     792.38       14.5
   y5+       794.47        8.4
   b5+       829.29       13.1
   y6+       891.73        7.1
   b6+       989.52       16.0
   y7+      1058.51       17.2
```

```
841. MORG-339_phosphopeptide_TMT_25%ACN_032017_OT.25032.25032.4 (gLNLDGTPALSTLGGFsPAskPsSPR)

   Ion         m/z   Intensity
   a7++      437.05        9.6
   b7++      450.85       12.1
   y6++      490.86       10.6
   y4+       525.74       12.2
   b9++      535.13       15.2
   b10++     591.49       34.4
   y8++      609.35        9.9
   y5+       623.25       28.2
   a17+++    628.53        8.5
   y9++      658.39       25.3
   a12++     670.96       11.0
   b19+++    693.64        8.9
   a20+++    740.13       13.4
   b5+       742.17       25.6
   y18+++    743.54       20.2
   a14++     756.77        8.6
   b14++     770.58       32.1
   a6+       771.37       17.5
   y19+++    776.26       53.0
   a15++     785.31       10.7
   b6+       799.23       12.8
   y12++     843.98       29.6
   b21+++    868.84        9.4
   a7+       872.56       68.0
   b22+++    900.51       38.6
   y14++     928.78       25.8
   a23+++    947.52       10.7
   b23+++    956.05       11.2
   y15++     979.23       13.5
   y6+       980.70       14.7
   y16++    1023.26       26.3
   a9+      1040.38       12.6
   b9+      1068.52       15.5
   y17++    1079.49       14.7
   y18++    1115.15       10.9
   y19++    1163.87       30.1
   b10+     1181.63       15.9
   y21++    1242.54        9.9
   b13+     1482.85        9.9
```

```
842. MORG-339_phosphopeptide_TMT_14%ACN_032017_OT.11236.11236.3 (sEPVVASPALAGPsPRGPTSVSATR)

   Ion         m/z   Intensity
   y11++     565.08       22.5
   y12++     648.13       10.8
   y13++     697.14       12.1
   y14++     725.61       17.0
   y15++     761.17       16.8
   a14++     813.22       10.1
   y16++     817.23       11.1
   y8+       818.40       12.3
   b5+       821.20       11.6
   y17++     852.66       23.6
   a6+       864.81        8.0
   y9+       875.52       14.1
   b6+       892.49       21.6
   y18++     901.65       42.7
   y19++     945.14        9.0
   a20++    1110.58       10.8
   a9+      1119.46       13.9
   b10+     1260.69       11.1
   b11+     1331.68        8.2
   a13+     1457.80        9.0
```

```
843. MORG-339_phosphopeptide_TMT_8%ACN_032017_OT.14710.14710.2 (sESDLEEAEPVVIPR)

   Ion         m/z   Intensity
   y3+       385.26       11.3
   y6+       680.54       31.1
   b4+       728.45       15.0
   y7+       809.56       12.1
   b5+       841.35       15.4
   y8+       880.51       17.7
   a6+       941.99       35.1
   b6+       970.26       13.4
   y9+      1009.60       17.4
   b7+      1099.39       15.4
   y10+     1138.68       11.7
   b8+      1170.47       15.5
   y11+     1251.77       26.3
   b9+      1299.56       22.6
   y13+     1453.87       15.6
   y14+     1582.97        9.9
   b12+     1594.69       16.2
   b13+     1707.81       21.5
```

```
844. MORG-339_phosphopeptide_TMT_25%ACN_032017_OT.22833.22833.4 (kVPVPEGLDLDAWINEPPsDsESEDEkPk)

   Ion         m/z   Intensity
   y2+       473.31       13.6
   y4++      480.41       11.1
   y12+++    655.87       16.5
   y9++      794.16       13.4
   a13++     811.65       28.0
   a15++     925.23       11.2
   y11++     935.14       15.1
   b7+       936.17       10.0
   y4+       959.47       14.7
   y12++     983.70       38.8
   a16++     989.60       11.2
   b24+++    998.64        9.9
   a8+      1022.14       21.8
   a25+++   1028.12       13.0
   y13++    1032.08       47.8
   b25+++   1037.49       19.7
   y22+++   1044.77       13.6
   b8+      1049.37       22.0
   y14++    1096.13       15.5
   a9+      1136.20       10.9
   y15++    1153.84       16.6
   y26+++   1172.39       10.1
   a27+++   1190.23       10.2
   y16++    1210.40       10.7
   y7+      1290.85       10.0
   y17++    1303.48       14.3
   y20++    1452.63       11.6
   a13+     1621.90       10.0
   y25++    1708.66       11.7
```

```
845. MORG-339_phosphopeptide_TMT_20%ACN_032017_OT.8485.8485.3 (nHFESSPLPTsPk)

   Ion         m/z   Intensity
   y4++      371.15       13.0
   y1+       376.22       10.1
   b4++      379.44       12.6
   y5++      419.80       46.9
   b6++      466.44       16.0
   y2+       473.38       29.5
   b7++      514.73       12.8
   y7++      524.74       26.4
   a8++      557.55       22.8
   y8++      568.49       15.5
   b8++      571.40       45.8
   a3+       600.35       15.2
   y9++      611.42       27.6
   b9++      619.90       25.8
   y3+       640.63       52.6
   b10++     670.53       11.3
   y4+       741.36       13.4
   b11++     753.65       10.9
   b4+       757.45       14.3
   y5+       838.44       36.8
   b5+       844.41       13.2
   b6+       931.48       26.2
   y6+       951.59       11.3
   y7+      1048.56       12.5
   b8+      1141.57       41.4
```

```
846. MORG-339_phosphopeptide_TMT_8%ACN_032017_OT.3130.3130.3 (eGTStPSQVPAGGTR)

   Ion         m/z   Intensity
   y6++      279.78       16.3
   a4++      288.30        6.2
   y7++      329.17        7.4
   y4+       390.30        6.2
   y8++      392.94        9.9
   b6++      441.82        9.6
   y10++     485.21       24.0
   a3+       489.22        6.3
   b3+       517.33       12.5
   a8++      535.57       11.2
   b8++      549.57       27.7
   y6+       558.33       43.9
   y11++     575.58       13.7
   a4+       576.36       34.3
   b9++      598.85       36.3
   b10++     646.90        7.5
   y7+       657.51       23.2
   b11++     682.76        6.9
   a12++     696.90        9.4
   a13++     725.42        5.5
   b13++     739.94        8.0
   a14++     776.81        5.8
   y8+       785.39       16.7
   y10+      969.43       10.0
   b8+      1097.47        9.9
   b9+      1196.50       15.5
```

```
847. MORG-339_phosphopeptide_TMT_20%ACN_032017_OT.1432.1432.4 (rRPSsPAPGPSRQSSSVGSQTLR)

   Ion         m/z   Intensity
   b2++      271.71       11.3
   b7+++     354.84       21.1
   y3+       389.61       23.2
   b5++      447.08       18.6
   y4+       517.41       25.7
   b12+++    519.49       19.1
   b7++      531.47       50.4
   y16+++    548.87       27.3
   a13+++    552.95       14.1
   y18+++    604.72       21.0
   b9++      608.43       14.6
   b15+++    620.23       11.9
   a16+++    639.67       15.7
   a10++     642.38       28.8
   b16+++    649.28       30.9
   b10++     656.92       47.5
   y19+++    660.68       26.5
   y6+       661.49       31.7
   a18+++    691.93       12.2
   a19+++    720.73       12.8
   y14++     745.56       13.8
   y7+       760.61       16.0
   a20+++    763.32       14.9
   b20+++    772.99       19.3
   y15++     774.04       27.0
   a21+++    796.65       12.6
   b21+++    806.68       19.3
   y16++     822.62       85.0
   a13++     828.90       12.9
   a22+++    834.72       13.3
   b22+++    843.88       13.4
   y8+       847.52       13.0
   y17++     858.25       17.4
   b5+       893.54       14.2
   y18++     906.70       89.2
   y9+       934.49       14.2
   b6+       990.74       11.2
   y10+     1021.60       12.7
   a7+      1033.22       24.6
   y11+     1149.58       13.7
```

```
848. MORG-339_phosphopeptide_TMT_20%ACN_032017_OT.17979.17979.4 (rRGsGSVDEtLFALPAASEPVIPSSAEk)

   Ion         m/z   Intensity
   y5++      375.45       12.9
   y1+       376.34       24.5
   y6++      424.38       47.3
   y7++      480.87       30.9
   y13+++    505.33       36.7
   y3+       576.37       31.8
   y9++      578.94       46.4
   a13+++    579.84       19.6
   b9++      627.35       19.1
   b15+++    659.65       23.3
   y4+       663.35       21.2
   b16+++    683.11       18.8
   b17+++    706.72       24.9
   y5+       750.28       31.1
   b11++     774.41       28.4
   b19+++    778.95       38.3
   y14++     806.26       19.8
   b20+++    811.28       41.0
   a21+++    834.47       34.2
   b21+++    844.15       38.1
   b12++     847.50      102.7
   b22+++    881.63       14.4
   a6+       882.36       23.1
   b13++     883.38       27.3
   y16++     898.88       16.7
   b6+       910.52       11.5
   b23+++    914.40       26.2
   a24+++    933.91       14.9
   b14++     940.16       78.9
   b24+++    943.27       20.7
   y7+       960.45       33.8
   a25+++    963.20       14.3
   b25+++    972.39       13.7
   b26+++    996.09       32.5
   a16++    1009.81       12.8
   b16++    1024.18       18.5
   y18++    1028.96       11.4
   b27+++   1039.15       38.0
   y8+      1059.76       42.7
   b18++    1103.30       22.9
   y19++    1118.77       30.2
   y9+      1156.61       47.0
   b19++    1167.80      103.1
   b21++    1265.54       37.3
   a22++    1308.53       16.4
   b22++    1322.40       93.2
   b26++    1493.70       11.6
```

```
849. MORG-339_phosphopeptide_TMT_8%ACN_032017_OT.6884.6884.3 (nEVNGTSEDIkSEGDTQsN)

   Ion         m/z   Intensity
   y8++      458.83       12.9
   y4+       529.34       16.2
   b3+       572.35       10.8
   b9++      587.79       12.0
   y9++      637.92        9.2
   b10++     644.66       11.0
   y6+       701.17       22.0
   b5+       743.44        7.9
   y11++     751.52       15.7
   a11++     809.22       10.2
   a6+       816.34       35.6
   b11++     823.77       55.3
   y7+       830.58       39.8
   b6+       844.25       27.1
   b12++     867.06       18.6
   a13++     917.38       29.0
   b13++     931.79       47.0
   b14++     959.69        8.0
   y16++     996.13       11.5
   b15++    1017.85       23.3
   b8+      1060.62        7.8
   b16++    1068.35       21.6
   b17++    1132.35       36.3
   b9+      1175.46        7.2
   y9+      1274.39        9.7
```

```
850. MORG-339_phosphopeptide_TMT_12%ACN_032017_OT.11146.11146.3 (sAYQDYDsDSDVPEELkR)

   Ion         m/z   Intensity
   b2+       387.95        6.7
   y6++      501.04       16.9
   y7++      550.32       21.1
   b3+       551.00       16.4
   y8++      607.50        6.4
   y9++      651.29       12.7
   a9++      664.21        6.9
   y10++     708.81        8.9
   y4+       774.48        7.7
   y11++     792.44       10.2
   b5+       794.41       13.3
   a12++     814.42        7.9
   b12++     828.73        9.5
   y12++     850.28       11.3
   a13++     863.06       12.5
   b13++     876.39       10.0
   y13++     931.21        9.3
   y14++     989.33        6.3
   y6+      1000.58        9.8
   b7+      1072.52        9.9
   b11+     1556.71        8.7
```

```
851. MORG-339_phosphopeptide_TMT_8%ACN_032017_OT.3047.3047.3 (aASEQQADASGGDsPk)

   Ion         m/z   Intensity
   y5++      407.07       11.4
   b6++      422.84       11.1
   y6++      435.49       25.2
   b7++      458.08       15.3
   b3+       458.87       22.1
   y2+       473.37       29.1
   y7++      478.96       29.8
   y8++      514.17       13.3
   b8++      515.85       24.6
   b9++      551.52       37.7
   a10++     580.70       11.8
   b4+       588.36       11.3
   b10++     594.94       42.4
   a11++     609.05       12.2
   a12++     637.62       48.3
   y3+       640.42       22.7
   b12++     652.16       30.1
   y11++     671.60       56.2
   b13++     709.26       16.8
   b5+       716.40       15.4
   y12++     735.42       18.5
   y4+       755.48       16.7
   b14++     792.67       17.3
   y5+       812.35       22.1
   b6+       844.19       19.4
   y6+       869.42       33.8
   b7+       915.49       19.1
   y7+       956.53       22.5
   y8+      1027.54       14.0
   b8+      1030.53       19.3
   b9+      1101.59       26.0
   y9+      1142.26       10.9
   b10+     1188.63       19.4
   b11+     1245.54       13.3
```

```
852. MORG-339_phosphopeptide_TMT_14%ACN_032017_OT.16829.16829.3 (mDsFDEDLARPSGLLAQER)

   Ion         m/z   Intensity
   b2+       476.28        7.1
   y11++     599.44        8.9
   y12++     656.21       30.2
   y6+       729.39        7.5
   b10++     745.50       15.8
   y14++     778.25       25.0
   y7+       786.09        7.8
   b11++     793.95       25.1
   b12++     837.55        5.9
   a13++     851.49        7.0
   b5+       905.27       10.4
   y16++     909.29       19.3
   b14++     922.70        9.9
   y9+       970.55       12.6
   y17++     992.77       12.8
   b16++    1014.62        6.8
   b6+      1034.35        7.1
   b18++    1143.48        8.9
   b7+      1149.44       14.8
   b9+      1333.83        5.4
```

```
853. MORG-339_phosphopeptide_TMT_22%ACN_032017_OT.16867.16867.4 (sQVETEDLILkPGVVHVIDIDRGEEk)

   Ion         m/z   Intensity
   y5++      424.33       18.7
   b6++      491.86       12.2
   y7++      538.27       17.4
   b3+       624.43       14.9
   y4+       691.44       13.1
   y10++     701.41       11.9
   y17+++    788.32       24.4
   a5+       826.28       14.9
   y5+       847.30       16.4
   y19+++    863.92       58.8
   y13++     869.72       12.9
   b19+++    870.81       12.8
   a11++     884.21       14.0
   y14++     898.27      167.9
   a20+++    899.70       19.3
   b20+++    908.38       13.1
   a21+++    938.02       14.4
   b21+++    946.78       66.0
   y22+++    978.62       17.2
   b23+++   1018.61       15.2
   y23+++   1021.82       19.3
   b14++    1024.53       11.3
   a24+++   1051.83       14.2
   a15++    1060.52       13.3
   b24+++   1061.69       20.8
   y25+++   1097.54       29.2
   b7+      1098.42       31.8
   a17++    1178.26       19.8
   y17++    1181.67       17.9
   b8+      1211.67       14.4
   b9+      1324.68       16.5
   b20++    1363.21       13.7
```

```
854. MORG-339_phosphopeptide_TMT_18%ACN_032017_OT.11755.11755.4 (hRkIVEPEVVGEsDsEVEGDAWR)

   Ion         m/z   Intensity
   y3+       432.25       22.3
   y4+       546.98       15.7
   b10+++    549.64       20.7
   b11+++    568.68       23.0
   y15+++    599.29       16.6
   y5+       604.31       35.5
   b6++      611.61       38.1
   b13+++    667.39       16.4
   b14+++    705.24       15.5
   y11++     705.97       35.5
   b8++      724.79       25.1
   y6+       733.38       40.4
   y19+++    750.70       18.9
   a15+++    752.08       30.3
   a9++      759.53       15.6
   b15+++    760.58       21.1
   y12++     770.47       17.9
   b9++      774.29       35.7
   y20+++    787.88       34.2
   a16+++    794.59       32.5
   y13++     798.54       34.7
   b10++     823.84       31.1
   a17+++    828.16       15.5
   y7+       832.52       22.4
   b17+++    837.44       30.1
   b11++     851.67       14.7
   b3+       880.58       38.5
   b12++     916.82       28.5
   b20+++    937.79       25.7
   a21+++    951.37       19.6
   y22+++    959.59       14.1
   a22+++   1013.93       28.1
   b22+++   1023.55       15.2
```

```
855. MORG-339_phosphopeptide_TMT_25%ACN_032017_OT.18224.18224.3 (aQsFDVASAPPAAEWAVPQSSRLk)

   Ion         m/z   Intensity
   y7++      523.03       12.4
   b13++     761.41       12.0
   y12++     801.16       12.0
   b5+       858.42       20.2
   y14++     885.36       13.3
   y15++     933.81       42.7
   b6+       957.36       17.6
   y16++     969.37       26.6
   a17++     990.88       19.0
   a7+      1000.66       37.8
   b17++    1004.37       25.4
   b7+      1028.53       11.5
   y7+      1044.70       25.3
   y18++    1048.17       18.9
   a8+      1087.50       11.9
   y19++    1097.79       33.1
   y8+      1144.02       16.3
   y20++    1154.71       15.0
   y9+      1214.80       15.0
   y21++    1228.96       20.1
   y10+     1400.76       13.8
```

```
856. MORG-339_phosphopeptide_TMT_12%ACN_032017_OT.6052.6052.4 (sAGSsSPEAGEDsDHEDGNYcPPVkRER)

   Ion         m/z   Intensity
   y7++      556.11        8.7
   y14+++    662.91       15.8
   a12++     718.36        9.2
   y16+++    756.85       14.3
   y17+++    795.35       10.5
   y11++     803.08       24.8
   y4+       817.31        8.2
   y19+++    857.49       11.7
   y12++     860.52       33.3
   a21+++    865.10       18.8
   b6+       866.46        8.9
   b21+++    874.17       13.7
   y20+++    880.50       51.0
   a22+++    897.54       17.7
   b22+++    907.27       17.0
   y5+       916.51       27.4
   y21+++    923.51       13.6
   y13++     925.07        9.3
   y22+++    956.24       13.4
   b7+       963.51        7.4
   y14++     993.71        9.1
   b16++    1006.37       11.9
   a8+      1064.31       10.9
   y25+++   1069.90       10.1
   y26+++   1089.08        9.3
   b18++    1092.46       10.4
   b9+      1163.54        8.8
   b27+++   1186.11        9.7
   b12+     1464.50        9.3
```

```
857. MORG-339_phosphopeptide_TMT_25%ACN_032017_OT.9807.9807.4 (kVAGSEGSSSTLVDYTSTSSTGGsPVRk)

   Ion         m/z   Intensity
   y4++      364.76       22.4
   y3+       631.36       28.1
   y17+++    688.51       20.5
   b13++     716.60       40.1
   b20+++    725.31       21.7
   y4+       728.44       23.8
   b21+++    758.75       20.2
   a14++     760.11       22.9
   a6+       773.20       36.9
   b14++     774.07       22.9
   a23+++    787.62       23.7
   a7+       830.50       27.9
   a24+++    843.64       35.2
   b24+++    852.69       27.2
   b7+       858.94       58.8
   y14++     868.51       53.0
   b25+++    885.53       20.2
   a26+++    908.66       19.5
   b26+++    917.62       24.2
   y15++     926.31       25.7
   b17++     949.85       21.5
   y6+       952.57       23.5
   a27+++    960.98       28.7
   y27+++    976.27       37.1
   b18++    1000.28       23.9
   b9+      1032.54       19.9
   b20++    1087.68       26.0
   y19++    1126.88       19.5
```

```
858. MORG-339_phosphopeptide_TMT_22%ACN_032017_OT.6205.6205.3 (sksPASTSSVNGTPGSQLSTPR)

   Ion         m/z   Intensity
   y15++     744.48       12.2
   y7+       788.52       11.4
   b11++     793.15       15.3
   a3+       813.13       11.9
   b3+       841.31       17.1
   y8+       845.29       19.9
   a13++     858.15       21.9
   b13++     871.85       50.9
   y18++     874.27       33.0
   y19++     922.43       15.4
   y9+       942.53       40.7
   b5+      1009.74       18.9
   y10+     1043.54       11.2
   b17++    1056.34       11.8
   y11+     1100.65       19.5
   b7+      1197.88       11.5
   y12+     1214.73       22.7
   y19+     1844.03       14.1
```

```
859. MORG-339_phosphopeptide_TMT_18%ACN_032017_OT.11901.11901.4 (sSkkEsEsEDsSDDEPLIk)

   Ion         m/z   Intensity
   y4++      350.42       20.5
   y1+       376.32       19.6
   y2+       489.39       24.2
   b7+++     515.24       14.3
   y3+       602.63       14.3
   b5++      624.14       11.8
   b10+++    652.39       16.7
   a11+++    698.51       17.4
   y4+       699.52       37.6
   a3+       733.34       13.7
   b12+++    736.98       13.3
   a7++      758.65       20.0
   a13+++    766.45       14.5
   b7++      772.52       18.5
   b13+++    775.59       21.5
   y11++     779.20       15.1
   a14+++    804.13       25.1
   y5+       828.48       19.3
   b15+++    856.78       46.9
   y12++     862.24       11.7
   a16+++    879.39       12.9
   b16+++    888.77       16.6
   a17+++    917.33       17.6
   b9++      920.75       14.9
   y13++     926.72       31.2
   y6+       943.63       16.7
   y17+++    955.04       13.0
   b18+++    964.62       28.0
   b10++     978.09       16.4
   y7+      1058.84       11.5
   b14++    1219.94       14.8
   b15++    1284.69       35.6
```

```
860. MORG-339_phosphopeptide_TMT_20%ACN_032017_OT.18579.18579.4 (kTsFDQDsDVDIFPSDFTSEPPALPRTGR)

   Ion         m/z   Intensity
   y9++      483.01       17.8
   y10++     547.44        6.9
   a15+++    681.56       11.4
   y19+++    702.08        7.8
   y13++     715.24       17.4
   y21+++    772.84        8.4
   b4+       773.56       12.1
   y15++     815.71        6.7
   y16++     864.73       19.5
   b12++     870.77        7.3
   a20+++    874.86        7.1
   b20+++    884.49        6.6
   a21+++    907.42       10.4
   y24+++    909.59        9.7
   b21+++    917.08        7.1
   a13++     930.74        7.7
   y17++     937.73       93.9
   a22+++    939.46        7.5
   b13++     944.38       41.9
   y18++     994.83       21.2
   b24+++   1010.32        9.4
   y27+++   1052.60       14.9
   y20++    1101.89       22.7
   y21++    1159.00       11.4
   y22++    1242.94       13.6
   y23++    1300.53        7.6
```

```
861. MORG-339_phosphopeptide_TMT_16%ACN_032017_OT.13113.13113.3 (nSkQQLsAEELDAQLDAYNAR)

   Ion         m/z   Intensity
   y4+       522.98       13.0
   b6++      579.52        9.2
   y5+       594.30       14.2
   y6+       709.38       10.0
   y13++     754.09        7.0
   b9++      763.22        7.7
   y7+       822.72       11.7
   b10++     827.58        9.5
   b11++     883.70        8.7
   a4+       888.26        8.9
   a12++     927.37       12.6
   y16++     929.78       30.1
   b12++     941.66       27.2
   y8+       950.40       29.6
   y9+      1021.48       15.8
   b14++    1040.95        8.9
   b15++    1097.45        7.9
   y10+     1136.67       10.4
   b16++    1155.39       10.6
   b17++    1190.64        7.3
   y11+     1249.50        9.3
   b18++    1271.96        8.8
   y12+     1378.58       10.2
   y14+     1578.88        7.6
```

```
862. MORG-339_phosphopeptide_TMT_18%ACN_032017_OT.5795.5795.3 (vVPQQItHTSPR)

   Ion         m/z   Intensity
   y4++      230.23        8.8
   y2+       272.12       29.7
   b4++      327.42       11.4
   y3+       359.30        9.9
   b5++      391.44       12.6
   a2+       400.45       10.0
   b2+       428.38       16.7
   b6++      447.58        9.1
   y4+       460.15       10.1
   y8++      509.97       16.7
   a7++      524.71       21.3
   b3+       525.53       53.7
   b7++      537.81       28.7
   b8++      607.27        9.5
   a4+       625.53        9.9
   a9++      643.12       37.8
   b4+       653.04       23.8
   b10++     701.22       20.9
   y6+       778.31       18.7
   b5+       781.53       15.4
   b6+       894.48       10.1
```

```
863. MORG-339_phosphopeptide_TMT_8%ACN_032017_OT.9158.9158.3 (sAsPYPTHcLsSPQR)

   Ion         m/z   Intensity
   y3+       400.27       11.2
   a7++      532.97       10.7
   y9++      583.29       12.2
   b8++      616.07       15.2
   y10++     632.00       26.3
   b3+       635.42        7.1
   y5+       654.61       38.9
   b9++      695.85       20.5
   a4+       704.12       17.7
   y11++     713.13       11.3
   a10++     738.45       11.3
   b10++     752.56       13.7
   y12++     762.16       30.3
   y6+       767.35        7.4
   a11++     822.08       17.9
   b11++     836.25        7.9
   a12++     865.13        8.2
   a5+       867.43        7.8
   b12++     879.81        7.8
   y14++     880.90        6.5
   b5+       895.31       18.2
   b13++     927.51        8.6
   b6+       992.60        5.8
```

```
864. MORG-339_phosphopeptide_TMT_4%ACN_032017_OT.4839.4839.3 (rRHsSLPtEsDEDIAPAQR)

   Ion         m/z   Intensity
   b6++      523.81        6.3
   b2+       542.42       10.8
   y11++     655.72        8.2
   b3+       679.58        5.9
   b9++      727.54       15.1
   y12++     746.32        9.4
   y7+       770.83       10.5
   a10++     797.17       11.3
   b10++     810.91        6.0
   b4+       846.42      108.0
   a11++     854.76       25.8
   b11++     868.02       10.6
   y8+       899.57       15.7
   b13++     990.33        9.7
   a14++    1032.78        8.0
   b14++    1046.57        9.7
   b15++    1082.86       12.4
   y10+     1181.59       10.9
   b18++    1230.57        6.3
   y11+     1310.74        6.5
   y13+     1588.66        6.0
```

```
865. MORG-339_phosphopeptide_TMT_14%ACN_032017_OT.14649.14649.3 (mDsFDEDLARPSGLLAQER)

   Ion         m/z   Intensity
   b1+       377.66        9.5
   b6++      525.79       10.8
   y12++     656.21       28.6
   y13++     713.79       14.4
   b10++     753.12       14.0
   a4+       778.20       37.6
   a11++     787.37       31.9
   b11++     801.34       11.8
   b4+       806.44       26.1
   y15++     835.89       20.2
   a5+       893.67       13.1
   y16++     909.21       22.8
   b5+       921.34       15.6
   b14++     930.05        8.3
   y9+       970.62       14.6
   y17++     992.74       16.7
   y10+     1126.62        8.6
   a18++    1136.87        8.2
   b18++    1150.94        8.6
   b7+      1165.48       18.0
```

```
866. MORG-339_phosphopeptide_TMT_4%ACN_032017_OT.5867.5867.2 (sLEsDNEEk)

   Ion         m/z   Intensity
   b1+       317.28        4.3
   b3+       559.45        5.5
   y3+       634.45        8.1
   a4+       697.91        4.3
   b4+       726.36        5.4
   y4+       748.50        7.8
   b5+       841.35        9.3
   y5+       863.46        6.2
   y6+      1030.41        5.4
   b8+      1213.44        4.9
```

```
867. MORG-339_phosphopeptide_TMT_20%ACN_032017_OT.10697.10697.4 (eVQTSELkAEVAsQk)

   Ion         m/z   Intensity
   b4+++     230.15       11.6
   y4+++     248.20       11.6
   b4++      344.00        8.8
   y4++      372.13       19.7
   y1+       376.25       24.6
   y5++      421.36       11.1
   b8+++     458.64       15.7
   a9+++     472.78        7.2
   b9+++     482.42       10.3
   a7++      495.27        7.9
   y9+++     504.35       10.4
   b10+++    525.34       28.5
   y10+++    547.81        8.4
   a11+++    549.26        7.5
   a3+       558.48       43.2
   b12+++    582.04       42.3
   b3+       586.19       25.6
   a4+       659.63        8.1
   y3+       671.39        8.8
   b8++      687.41       17.0
   y8++      699.98       13.2
   b9++      723.10       12.0
   y4+       742.36       11.8
   y9++      756.51        9.8
   b10++     787.55       31.7
   y10++     821.10        8.8
   b11++     837.19       32.0
   y5+       841.47       14.0
   b12++     872.68       26.5
   b6+       903.47        8.6
   y6+       970.47       11.3
   y7+      1041.60       10.7
```

```
868. MORG-339_phosphopeptide_TMT_10%ACN_032017_OT.6918.6918.4 (scAAEPEVEPEAHEGDGDkkGsAEGsSDEEGk)

   Ion         m/z   Intensity
   b5+       748.32       14.0
   y15+++    790.56        8.3
   b14++     883.53       13.1
   b20+++    942.46       15.0
   a7+       946.51        8.8
   b21+++    960.93       10.4
   b16++     969.76       21.9
   a17++     984.18       13.1
   b17++     997.74        9.8
   b22+++   1016.72       11.8
   y22+++   1022.95       24.4
   b23+++   1040.39       17.2
   a8+      1045.70       13.2
   b18++    1055.32       21.5
   y7+      1060.65       18.9
   b8+      1073.55       25.9
   a24+++   1074.54       26.2
   b25+++   1102.37        9.5
   y8+      1117.30       11.5
   a9+      1174.26       13.2
   b27+++   1186.93       10.0
   b9+      1202.60       18.0
   y27+++   1206.16       12.1
   y16++    1214.89       19.8
   a29+++   1259.53        8.2
   y18++    1300.31        9.4
   b30+++   1311.40       12.9
   b31+++   1330.69        9.1
   b11+     1429.01       10.6
   b12+     1499.65        9.2
   y22++    1533.71        8.9
   y23++    1582.25        8.6
   b16+     1937.86        9.3
```

```
869. MORG-339_phosphopeptide_TMT_4%ACN_032017_OT.7669.7669.3 (eQGsDEDkLQNsDDDEEk)

   Ion         m/z   Intensity
   y3++      317.25        4.9
   y2+       505.32        5.7
   y3+       634.42        9.4
   y4+       749.44        8.2
   b9++      771.32        7.3
   b5+       826.41        7.8
   y5+       864.32        7.8
   a11++     877.78        6.4
   b11++     892.11       12.9
   a6+       927.67       12.6
   y11++     930.15       14.9
   b6+       954.95       16.6
   a12++     961.50       12.9
   a13++    1019.20        5.7
   b13++    1032.96        5.3
   a7+      1042.13        6.3
   y13++    1052.39        8.2
   b7+      1070.52        7.9
   b14++    1090.56        6.6
   y14++    1109.84        4.9
   y16++    1221.50        5.3
   b17++    1277.38        7.1
```

```
870. MORG-339_phosphopeptide_TMT_18%ACN_032017_OT.13679.13679.3 (sSScGDAELLGQATLPVGsPSRPMSR)

   Ion         m/z   Intensity
   y7++      415.62       15.1
   y4+       490.31       17.1
   y11++     625.87       14.0
   b13++     753.81       13.2
   b6+       823.49       14.2
   a15++     846.86       12.1
   y17++     917.64       12.2
   b17++     958.20      103.7
   a18++     972.65       23.4
   y18++     973.62       28.4
   y19++    1038.28       11.2
   b19++    1070.52       11.3
   y20++    1073.71       13.0
   a9+      1108.71       11.2
   y22++    1159.69       10.2
   b22++    1241.06       20.9
   y11+     1250.60       11.8
   b23++    1288.66       10.3
   b13+     1505.98       10.4
   b15+     1719.91       10.2
   y16+     1721.01       11.6
```

```
871. MORG-339_phosphopeptide_TMT_6%ACN_032017_OT.1077.1077.3 (sSGRNsPSAASTSSNDSk)

   Ion         m/z   Intensity
   b1+       317.07        5.7
   a5++      352.24        6.0
   y1+       376.41        9.3
   b3+       461.48        6.4
   y2+       463.46        8.0
   y9++      563.42        7.7
   y3+       578.14        6.9
   y10++     598.44        7.3
   b10++     612.84       10.2
   a11++     641.82        8.0
   b11++     656.35        9.1
   y12++     691.02       19.7
   a12++     692.40       13.6
   b12++     706.98       16.0
   b5+       731.77       15.1
   a13++     736.06       31.4
   b13++     750.19       12.8
   a14++     780.32        7.2
   b15++     850.98        5.3
   y6+       866.46        9.1
   b6+       898.40        6.7
   b16++     908.65       14.7
   y7+       967.50       10.9
   y8+      1054.60       11.8
   y9+      1125.58       14.1
   y10+     1196.62       12.2
   y12+     1380.69       12.5
```

```
872. MORG-339_phosphopeptide_TMT_6%ACN_032017_OT.8335.8335.3 (lARADSDGDLsENDDGAGDLR)

   Ion         m/z   Intensity
   y2+       288.33       13.9
   b5++      379.07       11.8
   b6++      422.22        6.3
   b7++      479.96       21.7
   b9++      566.03       16.8
   y6+       588.40       15.8
   a4+       613.59        9.4
   b4+       641.36        6.8
   y7+       703.46       14.1
   b5+       756.63        7.3
   a13++     812.95       36.9
   a6+       815.23       18.8
   b14++     885.02       10.3
   b15++     942.57       20.0
   a16++     957.20        6.2
   b7+       958.38        7.0
   a18++    1020.68        7.5
   y10+     1061.46       10.6
   b19++    1092.80       23.4
   b20++    1148.59        8.3
   b10+     1243.78       10.6
   y12+     1341.65        7.2
   y14+     1513.77       13.0
```

```
873. MORG-339_phosphopeptide_TMT_10%ACN_032017_OT.11986.11986.3 (lPAPQEDTAsEAGtPQGEVQTR)

   Ion         m/z   Intensity
   y3+       404.18       15.5
   y8++      457.99       15.6
   y4+       503.35       19.4
   b3+       511.35       15.2
   b9++      577.13       16.8
   y11++     612.56       20.3
   y5+       632.56       11.4
   y6+       689.36       21.4
   b11++     724.50       10.4
   b12++     760.59       19.9
   y7+       817.15       10.4
   y15++     846.78       16.7
   b6+       865.21       25.4
   b14++     879.63       38.8
   y8+       914.50       24.4
   b7+       980.28       12.5
   b16++     991.73       12.0
   b18++    1085.18       21.9
   y9+      1095.44       16.3
   b19++    1134.77       11.3
   b9+      1152.48       26.7
   b11+     1448.66       10.4
   b12+     1519.60       15.1
```

```
874. MORG-339_phosphopeptide_TMT_14%ACN_032017_OT.8616.8616.4 (eEkESDDkPEIEDVGsDEEEEEkk)

   Ion         m/z   Intensity
   y2++      367.34       10.7
   y5++      560.88       11.4
   b10+++    625.57       14.4
   b7++      646.40       13.6
   y7++      689.96       22.8
   y8++      747.77       16.1
   a8++      810.92       12.5
   b8++      825.22       18.7
   a17+++    881.54       12.2
   y11++     908.84       16.9
   y17+++    919.69       26.5
   a10++     923.70       14.7
   b10++     938.31       16.8
   y18+++    958.28       23.2
   y12++     966.70       17.8
   b4+       974.46       15.8
   b19+++    977.11       21.4
   a11++     980.55       25.8
   y4+       991.15       24.2
   b11++     994.29       24.5
   y19+++    996.78       20.9
   b12++    1059.29       14.9
   b21+++   1062.51       11.8
   a22+++   1096.84       17.0
   a13++    1102.73       12.3
   b22+++   1105.80       12.1
   b13++    1116.88       43.3
   y5+      1120.70       14.8
   y15++    1151.61       19.9
   a14++    1152.42       13.2
   b14++    1165.84       19.0
   b6+      1176.65       23.5
   y22+++   1187.88       12.7
   b7+      1291.62       34.9
   b17++    1336.02       15.3
   y18++    1436.93       11.1
```

```
875. MORG-339_phosphopeptide_TMT_14%ACN_032017_OT.7381.7381.3 (eSDTkNEVNGTSEDIkSEGDTQsN)

   Ion         m/z   Intensity
   b3+       561.35        9.3
   b7++      632.10       11.4
   b8++      681.36        8.2
   b9++      738.50       10.6
   b10++     767.29       10.8
   y8+       917.19        8.0
   b13++     925.40        7.8
   a14++     968.83        7.0
   b14++     983.29       24.0
   a5+       991.58        6.7
   b15++    1039.71       17.5
   a6+      1105.89       18.0
   y9+      1274.92        6.9
   b18++    1326.41       13.2
   y10+     1387.71       13.3
   b20++    1412.01       13.5
   a21++    1448.53        7.2
   y22++    1454.16        6.6
   y13+     1719.03        6.7
```

```
876. MORG-339_phosphopeptide_TMT_8%ACN_032017_OT.10825.10825.3 (sDISERPSVDDVEsETGStGALETR)

   Ion         m/z   Intensity
   b6++      459.38        8.6
   y11++     600.83       10.6
   b10++     658.49       17.2
   b11++     716.00       14.5
   b12++     765.84       13.1
   b13++     830.29       17.1
   y15++     855.91        8.7
   b6+       917.44       17.8
   y17++     962.97        7.9
   y9+       971.16       41.8
   b15++     977.93       31.0
   a7+       986.32       10.4
   y19++    1055.28        8.4
   a8+      1073.48        9.4
   b18++    1100.09       10.1
   b8+      1101.34        8.0
   y20++    1133.89        8.9
   a9+      1172.37        8.3
   b19++    1191.29        7.8
   y11+     1201.34       13.4
   b21++    1255.09       11.5
```

```
877. MORG-339_phosphopeptide_TMT_10%ACN_032017_OT.6439.6439.3 (aSAAEGSEAsPPSLR)

   Ion         m/z   Intensity
   y5++      285.22       20.0
   b5++      330.12        8.8
   b6++      358.74       11.4
   b2+       388.24        8.7
   b7++      402.29       15.1
   y7++      404.21       12.2
   a8++      452.58        8.9
   b3+       459.00       10.9
   b8++      466.52       21.4
   y4+       472.27       10.6
   a4+       502.31       26.0
   y9++      512.53       13.7
   b4+       530.47       13.6
   y10++     540.47       17.7
   y5+       569.44       26.4
   a10++     571.35       44.8
   y11++     605.12       12.5
   b11++     634.42       16.3
   b5+       659.44       16.3
   b6+       716.36       19.1
   y6+       736.41       20.1
   b14++     783.17       11.7
   y7+       807.43       30.1
   b8+       932.53       17.9
   y8+       936.40       11.0
   b10+     1170.52        9.8
```

```
878. MORG-339_phosphopeptide_TMT_8%ACN_032017_OT.13026.13026.3 (gAEDAPDADTAIINAEGsQVNAEEk)

   Ion         m/z   Intensity
   y1+       376.48       12.9
   y2+       505.36       17.6
   b4+       602.39       15.1
   b11++     622.53       14.8
   y3+       634.47       23.4
   y9++      635.61       15.8
   b5+       673.37       20.5
   b12++     678.73       17.2
   y4+       705.51       18.8
   y11++     735.40       21.1
   y12++     793.07       19.4
   y5+       819.47       39.3
   b15++     828.17       32.0
   y13++     849.69       20.3
   b16++     892.93       16.9
   a17++     906.51       11.0
   y6+       918.62       16.2
   b17++     921.37       17.6
   y15++     941.44       12.2
   b8+       956.50       15.6
   a18++     990.53        9.7
   y16++     992.31       13.5
   b19++    1068.25       14.5
   b9+      1071.54       10.0
   b20++    1118.27       47.5
   b10+     1172.59       17.8
   b21++    1174.71       13.8
   y20++    1191.41       11.5
   b22++    1210.93       41.7
   y8+      1214.01       16.0
   y21++    1226.57       18.3
   b11+     1243.55       21.3
   y9+      1270.59       21.2
   b23++    1275.00       16.3
   b24++    1339.98       16.5
   b12+     1356.81       26.3
   b13+     1469.82       28.3
   y11+     1470.68       28.1
   b14+     1583.86       14.3
   y12+     1584.79       17.1
   b15+     1654.73        9.9
   b16+     1783.74       15.5
```

```
879. MORG-339_phosphopeptide_TMT_8%ACN_032017_OT.4347.4347.4 (rRHSSLPtEsDEDIAPAQR)

   Ion         m/z   Intensity
   y4++      236.26       14.0
   b2++      271.92       11.5
   y3+       374.38       10.7
   y4+       471.30       54.0
   y9++      507.30        9.4
   b2+       542.31       39.7
   a11+++    543.29       19.4
   b11+++    552.37       18.6
   a12+++    586.38       13.9
   y10++     591.51       13.2
   b12+++    595.64       34.3
   y15+++    596.58       12.3
   a8++      609.09       16.2
   a13+++    624.66       25.9
   y16+++    625.43       27.1
   y11++     655.44       17.8
   a14+++    662.69       46.5
   b14+++    671.85       62.1
   a15+++    686.21       11.3
   b9++      687.24       17.5
   b15+++    695.57       91.6
   y18+++    723.40        9.4
   y12++     746.11       10.7
   b10++     770.48       15.9
   b11++     828.54       11.4
   b12++     893.03       22.4
   b13++     950.65       37.0
   b14++    1007.19       25.0
   b15++    1042.72       30.2
```

```
880. MORG-339_phosphopeptide_TMT_16%ACN_032017_OT.18532.18532.2 (lGGPEGPGsPLLAT)

   Ion         m/z   Intensity
   y3+       304.22        8.6
   b1+       343.33        7.2
   b3+       457.30        9.4
   y5+       514.37       13.3
   a5+       655.33       10.2
   y6+       681.67        9.6
   b5+       683.50       36.9
   b6+       740.59       15.5
   y8+       835.32       12.2
   y9+       892.44       25.6
   b8+       894.56       12.0
   b9+      1061.38       12.1
   y13+     1232.43        7.0
   b11+     1271.54       22.2
   b12+     1384.75       14.6
```

```
881. MORG-339_phosphopeptide_TMT_4%ACN_032017_OT.7272.7272.3 (iAcDEEFsDSEDEGEGGRR)

   Ion         m/z   Intensity
   y6++      316.20        6.8
   b1+       343.30        4.5
   y7++      380.78       12.9
   a5++      396.11        6.0
   y8++      438.20        5.8
   y4+       445.31        6.4
   a3+       546.49       19.5
   b7++      547.32        7.2
   y11++     604.09       11.6
   y6+       631.28        6.1
   y12++     687.29        7.0
   b4+       689.48       12.2
   a10++     718.62        7.7
   y13++     760.31        8.7
   a11++     782.50       13.1
   a5+       790.66      101.7
   b11++     796.77       14.5
   b12++     854.34        9.2
   y15++     890.14       10.6
   b6+       947.46        8.1
   a15++     998.24        5.2
   a9+      1348.56        4.8
   b11+     1592.79        5.0
   b12+     1707.74        7.0
```

```
882. MORG-339_phosphopeptide_TMT_12%ACN_032017_OT.11422.11422.3 (kTsFDENDsEELEDk)

   Ion         m/z   Intensity
   y3+       620.42        9.7
   b8++      624.12        8.5
   y10++     758.60        8.1
   b4+       773.02       10.4
   a11++     822.01        8.1
   b11++     836.31        7.3
   a5+       860.11       17.8
   y5+       862.94       15.5
   a13++     943.61        7.6
   y13++     973.47        9.0
   b6+      1017.70        7.8
   b7+      1131.40        7.2
   y7+      1158.52        8.2
```

```
883. MORG-339_phosphopeptide_TMT_25%ACN_032017_OT.17696.17696.4 (lkNDSDLFGLGLEEmGPkESsDEDR)

   Ion         m/z   Intensity
   y3+       419.25       12.5
   b4++      465.45       22.5
   b6++      566.39       25.4
   b7++      623.15       17.7
   b8++      696.64       17.5
   y5+       701.06       16.3
   y17+++    725.02       21.1
   b10++     781.78       20.6
   a3+       786.60       17.8
   y19+++    812.41       18.5
   y12++     852.52       37.9
   b12++     866.78       32.3
   y21+++    879.20       24.5
   b18+++    883.26       42.0
   y22+++    917.61       12.3
   b19+++    926.70       18.8
   b13++     930.90       15.5
   a5+       988.21       12.5
   b14++     995.86       27.1
   a21+++   1001.67       12.7
   b21+++   1011.53       15.4
   b5+      1017.10       15.3
   a22+++   1040.02       12.9
   b22+++   1049.33       12.2
   y16++    1059.36       12.3
   b15++    1069.18       15.3
   y24+++   1075.13       13.5
   y17++    1087.86       15.4
   b23+++   1092.77       24.1
   b16++    1097.85       19.4
   b24+++   1131.11       24.4
   a17++    1131.85       16.4
   y19++    1217.90       15.6
   y21++    1318.98       15.6
```

```
884. MORG-339_phosphopeptide_TMT_4%ACN_032017_OT.10121.10121.2 (aVTcDIsEDEED)

   Ion         m/z   Intensity
   y2+       263.14        6.6
   b1+       301.20        6.4
   a2+       372.33        8.2
   y3+       392.37        6.9
   b2+       400.34       18.9
   b3+       501.40       12.1
   y4+       507.18       14.7
   b4+       661.27       15.3
   a5+       748.68       14.9
   b5+       776.54        9.8
   y6+       803.25       33.9
   b6+       889.52       54.8
   y8+      1031.41       14.2
   b7+      1056.40        6.5
   b8+      1185.45       29.5
   y9+      1191.36       13.2
   y10+     1292.47       19.7
   b9+      1300.53       20.6
   y11+     1391.63        9.1
   b10+     1429.65       34.5
```

```
885. MORG-339_phosphopeptide_TMT_22%ACN_032017_OT.12853.12853.3 (sQtPPGVATPPIPk)

   Ion         m/z   Intensity
   b1+       317.23       16.9
   y5++      390.83       79.1
   y6++      441.23       18.1
   y2+       473.36       45.1
   a7++      474.37       16.2
   y7++      476.80       13.5
   b7++      488.83       27.0
   b8++      524.19       24.3
   a9++      561.08       15.3
   b9++      574.94       35.4
   y3+       586.25       15.4
   y10++     603.39       14.4
   b10++     623.36       20.7
   b3+       626.33       15.6
   y4+       683.58       29.7
   a12++     714.64       15.4
   b12++     728.56       58.8
   y5+       780.58       42.2
   y13++     805.99       11.6
   a6+       849.04        9.4
   b6+       877.46       11.8
   y6+       881.66       10.8
   y7+       952.52       17.3
   b7+       976.50       22.2
   b8+      1047.76       17.1
   b9+      1148.68       51.2
```

```
886. MORG-339_phosphopeptide_TMT_20%ACN_032017_OT.16763.16763.3 (esASPTIPNLDLLEAHTk)

   Ion         m/z   Intensity
   y1+       376.29        8.6
   a5++      377.23        8.1
   y2+       477.36       11.7
   y7++      520.88        9.4
   y8++      578.23        7.5
   y3+       614.44       20.5
   y9++      635.28       12.4
   b4+       684.47        8.6
   y4+       685.52       21.5
   y10++     692.19        8.6
   b11++     717.83        8.7
   y11++     740.62       48.0
   y12++     796.77       12.3
   y5+       814.66       12.1
   y13++     847.80        8.0
   y14++     896.19       43.0
   y6+       927.51       11.8
   b15++     931.31       10.5
   a16++     985.58       10.9
   b7+       995.48       14.3
   b16++     999.85       11.3
   b17++    1049.88        9.9
   y9+      1268.75        8.1
   b10+     1319.63        8.9
   a11+     1406.49        7.4
```

```
887. MORG-339_phosphopeptide_TMT_8%ACN_032017_OT.4567.4567.3 (sEPQPEEGsPAAGQk)

   Ion         m/z   Intensity
   y1+       376.22        9.6
   y6++      400.98       12.3
   y2+       504.35        8.6
   y8++      513.13        8.3
   y3+       561.35       22.3
   y9++      577.50       10.8
   a9++      611.31        7.2
   b9++      625.48        9.1
   y4+       632.37       11.8
   y10++     641.43       14.8
   a4+       643.44        6.3
   a10++     660.61       21.8
   b4+       671.51       25.0
   b10++     674.13       16.3
   y11++     690.80        6.2
   a11++     695.51        7.6
   y5+       703.11        7.4
   b11++     710.03       16.1
   b12++     745.48       23.2
   y12++     754.45        6.9
   a13++     759.45        6.7
   y6+       800.48        8.5
   y13++     803.20       11.3
   b14++     837.89        7.7
   b6+       897.53        6.9
   y8+      1024.57        7.4
   b7+      1026.56        9.6
   y9+      1153.52        7.3
```

```
888. MORG-339_phosphopeptide_TMT_16%ACN_032017_OT.11540.11540.4 (sREcSLsPkStSPPPSPEVWAESR)

   Ion         m/z   Intensity
   b1+       317.27       30.2
   b6++      481.79        9.6
   a7++      551.71       12.4
   y10++     579.66       11.5
   a8++      599.95       16.9
   b3+       602.81       16.4
   a11+++    608.73       17.8
   y11++     627.95       15.6
   y12++     676.50       35.7
   b14+++    711.20       13.8
   b15+++    744.34       15.2
   y6+       747.21       12.7
   b4+       762.39       26.1
   a16+++    763.49       34.4
   a9++      778.43       14.7
   b9++      792.52       25.6
   a17+++    796.40       23.5
   b17+++    805.38       72.5
   y14++     810.54       14.5
   y19+++    814.42       25.8
   a19+++    872.28       19.7
   b19+++    881.63       13.0
   a11++     911.96       16.8
   b20+++    943.79       10.8
   b6+       962.77       13.7
   b12++     970.15       10.2
   y8+       973.34       15.5
   b22+++   1010.28       18.2
   b13++    1018.43       12.7
   y16++    1032.62       14.4
   y9+      1060.70       13.3
   y17++    1081.26       28.2
   y10+     1157.57       16.2
   b16++    1158.63       12.7
   y12+     1351.67       22.8
```

```
889. MORG-339_phosphopeptide_TMT_20%ACN_032017_OT.7592.7592.3 (sRsAPPNLWAAQR)

   Ion         m/z   Intensity
   a3++      307.15       11.6
   b1+       317.15       10.0
   b3++      320.78       10.1
   b4++      356.04        8.3
   a2+       445.36        7.2
   b2+       473.44        7.8
   a7++      496.63        7.4
   b7++      510.04       11.7
   y9++      527.12       29.9
   a8++      552.87        9.3
   y10++     562.61       22.4
   b8++      566.72       15.1
   y5+       631.32       14.2
   b3+       640.42        7.8
   a9++      646.29       10.5
   b9++      659.66        9.9
   a4+       683.46        8.8
   b10++     695.63       10.2
   b4+       711.39       14.4
   y6+       744.39        9.5
   b12++     795.17       15.2
   y8+       955.69        7.5
   y9+      1052.57       13.9
   y10+     1123.68        8.6
```

```
890. MORG-339_phosphopeptide_TMT_12%ACN_032017_OT.5688.5688.4 (aSRkGAEEEEEEEDDDsEEEIk)

   Ion         m/z   Intensity
   y1+       376.36       14.9
   y2+       489.34       16.6
   b12+++    602.10       11.0
   y3+       618.54       16.3
   b13+++    645.01       10.5
   b14+++    683.33       20.4
   b15+++    721.55       17.1
   y4+       747.51       15.5
   a10++     759.72       17.9
   b16+++    760.45       20.9
   y16+++    764.31       10.8
   b10++     773.69       11.0
   y17+++    787.91       14.9
   a17+++    806.66       73.3
   y18+++    807.47       13.1
   b17+++    815.56       22.0
   a18+++    849.26        9.4
   b18+++    858.82       12.4
   a4+       873.41        9.1
   y5+       876.42       11.1
   a19+++    892.36        9.6
   b19+++    901.80       25.9
   b12++     902.52       22.2
   y19+++    926.68       11.5
   a20+++    935.18       15.8
   b20+++    944.91       30.3
   a13++     953.33       10.5
   b5+       958.54        9.1
   a21+++    973.33       12.7
   b21+++    982.61       30.0
   b14++    1024.81       10.9
   b15++    1081.92        9.8
   b16++    1139.84       11.0
   b7+      1158.69        9.6
```

```
891. MORG-339_phosphopeptide_TMT_20%ACN_032017_OT.12634.12634.4 (aREADDEsLDEQASASkLSLLSR)

   Ion         m/z   Intensity
   b5++      386.83       25.9
   b6++      444.27       41.5
   a7++      494.93       18.0
   b7++      508.94       22.7
   y8++      566.68       11.7
   b3+       585.97       11.5
   a14+++    599.97       26.5
   y10++     646.11       26.4
   b9++      649.27       11.8
   a16+++    653.04       10.6
   b4+       657.35       36.2
   y11++     681.49       26.7
   a10++     692.28       16.2
   b10++     706.59       33.7
   y12++     745.46       26.5
   y13++     809.95       36.0
   y20+++    815.27       10.8
   b12++     835.01       15.8
   y21+++    858.07       10.3
   a6+       859.26       16.2
   y14++     867.74       33.7
   b13++     870.44       16.6
   y15++     924.27       46.2
   b15++     949.29       10.6
   b16++     992.52       12.6
   y16++    1007.72       21.3
   y17++    1072.33       26.0
   y18++    1129.61       16.0
   y19++    1187.20       12.4
```

```
892. MORG-339_phosphopeptide_TMT_14%ACN_032017_OT.9701.9701.4 (sETLVNAQQtPLGtPkENR)

   Ion         m/z   Intensity
   b1+       317.18       11.7
   b5++      380.20       11.3
   b6++      436.95       20.0
   b7++      472.74       16.1
   y6++      527.26       15.4
   b8++      536.88       10.0
   a13+++    540.90        9.1
   y11+++    544.07       14.4
   b3+       547.31        8.0
   y7++      555.94       17.7
   a9++      586.79       15.2
   b9++      600.76       13.4
   a14+++    601.44        9.1
   y13+++    609.99       21.5
   b14+++    610.71       12.6
   y8++      612.49        8.6
   a4+       632.31       10.2
   a15+++    633.59        9.3
   b15+++    643.05       19.1
   y14+++    647.99       26.8
   y9++      660.54       33.7
   b10++     691.17        9.4
   y16+++    719.04        7.5
   b11++     739.95       13.6
   y10++     751.64       20.8
   b5+       759.46       21.1
   a12++     782.67       10.0
   b12++     796.65       14.5
   b17+++    805.13       10.4
   a13++     810.63       11.5
   y11++     815.52       12.6
   b13++     825.13       13.7
   b18+++    843.02        8.0
   a6+       845.46        7.8
   y5+       872.56        7.8
   b6+       873.57       16.0
   y12++     879.68       12.5
   b14++     915.13       18.5
   b7+       944.51       10.0
   b15++     963.75       10.7
   y14++     972.23       23.6
   y15++    1021.83        8.3
   b8+      1072.57        9.1
```

```
893. MORG-339_phosphopeptide_TMT_14%ACN_032017_OT.10642.10642.4 (eEPLsEEEPcTSTAVPSPEkk)

   Ion         m/z   Intensity
   a4++      335.25        9.1
   y1+       376.32       10.0
   y7+++     415.21       24.1
   b2+       488.12       10.2
   y6++      572.53       52.0
   a14+++    615.08       17.5
   y13+++    620.94       48.4
   y7++      622.14       36.9
   y8++      657.60       31.3
   b16+++    689.41       12.9
   b4+       698.76       10.9
   y15+++    706.94        9.2
   y9++      708.29       17.4
   a17+++    708.95       10.6
   b17+++    718.31       13.5
   y2+       733.70       24.0
   y16+++    749.40       17.0
   b18+++    750.33       20.4
   y10++     751.65       35.3
   b10++     755.51       15.6
   y11++     802.01       10.1
   y18+++    843.51       15.4
   y3+       863.05       11.0
   y12++     882.24       19.4
   b20+++    912.95        9.7
   a14++     921.19       12.2
   y13++     930.78       19.2
   b14++     935.53       14.3
   b15++     985.19       33.6
   y14++     994.97        9.4
   b7+      1123.52       16.1
   y16++    1124.50        9.0
   b8+      1252.53       34.3
```

```
894. MORG-339_phosphopeptide_TMT_8%ACN_032017_OT.6095.6095.4 (sDAEEDGVTGsQDEEDSkPk)

   Ion         m/z   Intensity
   y2++      237.27        9.5
   b7++      467.38       11.1
   y2+       473.35       34.8
   y5++      517.04       18.1
   a14+++    567.45       17.3
   y11+++    586.81       12.2
   y12+++    620.53       20.3
   b4+       632.19        9.7
   y7++      645.96        9.2
   a16+++    649.44       10.1
   y13+++    653.49        8.5
   y14+++    672.15       20.5
   a17+++    677.86       11.2
   b17+++    687.31        7.3
   y8++      703.35       26.1
   b5+       761.35       10.2
   b13++     800.87       38.0
   b18+++    806.71       23.5
   y18+++    820.14        6.8
   y3+       830.43       13.8
   a6+       848.36        8.2
   a14++     851.28       14.5
   y19+++    859.13        8.3
   b6+       876.45        8.3
   y11++     879.73       24.4
   a7+       905.46        7.9
   y12++     929.80       19.1
   b7+       933.47       11.5
   b16++     987.58       12.4
   b17++    1030.55       11.8
   y5+      1032.51       19.4
   b9+      1133.52        9.8
   b18++    1209.52       11.3
```

```
895. MORG-339_phosphopeptide_TMT_12%ACN_032017_OT.11758.11758.3 (gAPDPRVDDDsLGEFPVSNSR)

   Ion         m/z   Intensity
   b2+       358.36        7.5
   b4+       570.35       66.2
   b9++      576.74        6.9
   b10++     634.67       13.8
   y6+       659.39       15.5
   a13++     788.39        7.0
   b13++     802.91       11.7
   y7+       806.58       16.1
   y14++     809.48        6.6
   b6+       823.70       10.0
   a14++     853.86        7.5
   b14++     867.54       12.9
   y8+       935.43        8.0
   y16++     936.75       65.5
   b15++     941.09       15.5
   y17++     985.77       89.5
   b16++     989.73        9.0
   y9+       992.51       12.6
   y10+     1105.65       11.3
   y11+     1272.57       13.2
   y12+     1387.61       12.3
   y13+     1502.91        7.0
```

```
896. MORG-339_phosphopeptide_TMT_25%ACN_032017_OT.19830.19830.4 (sPNELVDDLFkGAkEHGAVAVER)

   Ion         m/z   Intensity
   b1+       397.12       22.1
   y10++     662.99       11.8
   y15+++    690.90       53.4
   y7+       701.81       17.0
   y12++     726.86       14.3
   y16+++    729.27       43.4
   b4+       737.22       14.7
   a14+++    752.14       14.0
   b14+++    761.31       17.6
   a15+++    795.49       45.2
   y18+++    799.95       37.0
   b15+++    804.32       16.6
   y8+       838.53       23.9
   a16+++    840.40       16.3
   b5+       850.41       13.7
   b17+++    868.97       13.2
   b18+++    892.32       14.3
   y13++     905.75       17.4
   b19+++    926.06       14.4
   a13++     948.77       12.4
   b20+++    949.79       15.4
   y22+++    951.85       69.1
   y14++     978.89       11.2
   b21+++    982.89       12.0
   b22+++   1025.68       24.0
   b7+      1064.52       18.7
   y16++    1093.50       19.7
   a8+      1151.40       12.8
   b8+      1179.55       20.8
```

```
897. MORG-339_phosphopeptide_TMT_20%ACN_032017_OT.5656.5656.4 (qAsISGPAPTkASGAPPGGQQR)

   Ion         m/z   Intensity
   b7+++     317.24       11.8
   y6++      321.73       10.2
   y7++      370.19       38.5
   b6++      427.05       14.7
   b7++      475.74       14.3
   b8++      511.37       23.7
   b11+++    526.36       12.3
   y14+++    527.81       20.9
   y5+       545.09       12.6
   a9++      546.23       13.2
   b12+++    550.08       13.7
   b9++      559.54       13.8
   b13+++    579.32       19.2
   y16+++    583.73       31.5
   a14+++    589.03       39.0
   b14+++    598.06       21.0
   y17+++    602.83       29.0
   b10++     610.28       12.3
   a15+++    612.51       18.7
   b15+++    621.84       73.2
   y18+++    631.61       33.6
   y6+       642.05       61.9
   y19+++    669.81       10.1
   b17+++    686.56       12.8
   b4+       709.24       14.4
   a19+++    715.39       10.3
   y7+       739.42       47.2
   y13++     742.59       11.1
   a11++     774.56       11.5
   b11++     789.06       13.3
   y14++     791.10       50.4
   b5+       796.46       16.2
   a12++     810.45       10.7
   b12++     824.75       12.8
   a13++     853.54       23.5
   y16++     875.22       26.3
   b14++     896.33       10.3
   y17++     903.79       14.4
   b15++     932.15       62.7
   y10+      954.52       11.8
   b8+      1021.53       13.5
   y11+     1025.50       12.0
```

```
898. MORG-339_phosphopeptide_TMT_25%ACN_032017_OT.18903.18903.3 (sLSSPTVTLSAPLEGAk)

   Ion         m/z   Intensity
   y4++      317.27       19.1
   y1+       376.30       16.2
   y6++      422.57       22.8
   y2+       447.49       11.3
   y3+       504.36       27.5
   b8++      541.41       14.3
   a9++      584.61        9.8
   b9++      598.54       25.4
   y4+       633.19       29.8
   b10++     642.00       17.8
   a4+       656.63       13.0
   y11++     657.52       19.9
   a11++     663.30        9.5
   b11++     677.61       15.6
   b4+       684.52       10.6
   y12++     708.61       14.4
   a12++     711.94       27.1
   b13++     782.64       13.5
   y15++     843.54       43.7
   b14++     846.82       26.5
   b15++     875.85       23.9
   b16++     911.30       23.9
   y7+       914.64        9.6
   b7+       981.52       20.1
   y8+      1001.58       16.6
   b8+      1082.64       10.9
   b9+      1195.71       13.3
   b11+     1353.71       25.7
```

```
899. MORG-339_phosphopeptide_TMT_10%ACN_032017_OT.6892.6892.4 (eGDGSATTDAAPAtSPkAEEPSkAGDAPSEEk)

   Ion         m/z   Intensity
   y8++      531.31       13.9
   b9++      532.44        9.3
   b6+       746.32        9.3
   y5+       818.65        8.7
   y17+++    820.08       56.1
   b15++     820.92       18.1
   y12++     837.80       27.3
   y6+       889.53       29.3
   y20+++    932.71       13.3
   y21+++    965.42       21.8
   y14++     966.28       18.6
   a23+++    980.17       19.3
   b23+++    989.33       26.9
   y15++    1001.94       34.6
   y7+      1004.30       23.5
   b24+++   1012.76       11.7
   b25+++   1032.53       16.9
   y8+      1061.80       10.5
   b9+      1063.45       12.2
   b26+++   1070.64       27.9
   b18++    1083.88       10.3
   y9+      1132.98       11.9
   y27+++   1141.72        9.3
   a29+++   1145.90       22.1
   b19++    1148.85       12.3
   y29+++   1190.28       17.0
   a20++    1199.22        9.1
   b11+     1205.60       17.2
   b20++    1213.35       14.5
   b21++    1262.04        8.6
   a22++    1290.89        8.7
   y20++    1398.52       14.1
   a14+     1526.87        9.5
```

```
900. MORG-339_phosphopeptide_TMT_10%ACN_032017_OT.16943.16943.3 (sQsPsPPPLPEDLEk)

   Ion         m/z   Intensity
   b1+       317.17       11.7
   y1+       376.30       15.7
   b5++      438.72        8.7
   y6++      480.46       59.4
   y2+       505.28       24.9
   a8++      570.44       15.5
   a3+       584.65       10.5
   b3+       611.87        9.4
   y3+       618.48       19.4
   a9++      626.72       10.3
   y9++      633.65       14.7
   b9++      640.69       23.1
   a10++     675.12       10.4
   a4+       681.26       14.3
   y10++     682.61       86.1
   b4+       709.05       14.0
   y4+       733.39       28.5
   b11++     753.86        9.0
   y12++     814.69       18.7
   b13++     868.19       20.2
   b5+       876.30       32.2
   a14++     918.63        8.0
   b14++     932.24        9.4
   y6+       959.56       25.8
   y14++     961.52       12.2
   y7+      1072.89       11.0
   b9+      1280.54       34.8
```

```
901. MORG-339_phosphopeptide_TMT_12%ACN_032017_OT.9166.9166.4 (eEkESDDkPEIEDVGsDEEEEEk)

   Ion         m/z   Intensity
   y1+       376.33       10.4
   y2+       505.43       14.9
   b8+++     550.53       10.5
   b6++      589.15       21.2
   b10+++    625.76       14.8
   y3+       634.49       16.7
   b7++      646.45       27.1
   b11+++    663.62       11.6
   b12+++    706.43       17.5
   b13+++    745.04       40.6
   y4+       763.45       16.4
   b14+++    778.13       21.3
   b15+++    797.21       12.4
   a8++      810.80       10.8
   b8++      824.96       15.8
   b3+       845.50       12.1
   b16+++    852.08       14.9
   a9++      859.20       16.0
   b9++      873.80       12.5
   a17+++    881.34       14.6
   b17+++    891.03       25.6
   y5+       892.50       15.9
   y19+++    906.38       35.0
   y13++     908.75       17.1
   b10++     938.20       13.3
   y14++     973.10       16.8
   b19+++    976.88       17.8
   b11++     994.67       12.8
   b20+++   1020.13       19.2
   y6+      1021.34       21.5
   y15++    1022.42       13.2
   b12++    1058.91       10.8
   b21+++   1062.88       15.8
   b22+++   1106.22       17.4
   b13++    1117.05       14.6
   y7+      1136.63       15.2
   a6+      1148.32       10.5
   a14++    1151.66       19.2
   b14++    1165.93       15.5
   y16++    1200.69       17.1
   y17++    1258.28       12.6
```

```
902. MORG-339_phosphopeptide_TMT_10%ACN_032017_OT.5780.5780.3 (iQQQAGTVPHSQEEDsQEEEEDVSSR)

   Ion         m/z   Intensity
   b3+       599.44        8.5
   y6+       692.35        8.1
   b4+       727.49       11.5
   b5+       798.58        9.6
   b13++     817.39       12.9
   y7+       821.48       10.2
   a7+       928.45        9.7
   y15++     938.75       10.2
   b15++     939.79       28.9
   y8+       950.22        9.9
   b7+       956.57       11.5
   a8+      1027.95       10.8
   y17++    1050.15       21.0
   b8+      1055.54       30.2
   a17++    1072.63       20.7
   y18++    1099.30       16.5
   a18++    1137.78        8.2
   b18++    1151.69       12.8
   y10+     1207.55       10.2
   b19++    1216.49       11.0
   b20++    1280.61        8.6
   b21++    1345.38       11.7
   y11+     1374.52       18.0
   b22++    1402.93       19.8
   b12+     1504.41        7.7
```

```
903. MORG-339_phosphopeptide_TMT_16%ACN_032017_OT.11132.11132.3 (tSPQAPGLVAsIRSPQR)

   Ion         m/z   Intensity
   y3+       400.30       15.8
   b7++      474.97        8.4
   b2+       498.26       11.5
   b8++      531.33        7.2
   b9++      580.71        7.7
   b3+       595.51        7.8
   y10++     603.34        7.0
   y11++     632.09        6.9
   y5+       643.10       14.1
   y12++     680.94       13.7
   a11++     686.21       72.9
   a4+       695.66       14.2
   b11++     699.70       15.8
   b4+       723.63        7.6
   y14++     780.08       25.3
   b5+       794.42       14.7
   y15++     829.05       32.6
   a14++     864.26        7.7
   b14++     877.74        7.6
   y7+       923.42        8.1
   b7+       948.37        6.3
   b16++     990.44        6.5
   y8+       994.50        7.6
   b8+      1061.68        7.7
```

```
904. MORG-339_phosphopeptide_TMT_10%ACN_032017_OT.5141.5141.4 (eEENksSSEGGDAGNDTRNTTADLQk)

   Ion         m/z   Intensity
   y1+       376.39       11.4
   y2+       504.40       12.4
   b5++      544.77       11.6
   y3+       617.47       32.8
   y10++     689.08       73.7
   b4+       731.02       20.5
   y4+       732.64       13.6
   b9++      780.33       12.7
   y5+       803.55       13.2
   b10++     808.81       12.5
   a11++     823.19       15.0
   a19+++    830.19       11.4
   y13++     832.27       22.3
   b11++     836.59       22.4
   a20+++    863.40       33.3
   y14++     867.70       45.7
   b12++     894.34       19.4
   b21+++    906.42       11.3
   y22+++    930.33       25.1
   a14++     944.68       12.4
   y16++     953.33       17.0
   y23+++    968.84       27.9
   y17++     982.21       28.5
   b24+++   1006.13       18.4
   b15++    1015.22       28.6
   y18++    1046.42       17.1
   b25+++   1049.34       16.9
   a16++    1058.77       14.1
   b16++    1072.94       35.3
   b5+      1088.97       21.4
   y19++    1090.26       28.8
   b17++    1123.39       11.3
   y20++    1133.81       26.4
```

```
905. MORG-339_phosphopeptide_TMT_16%ACN_032017_OT.6100.6100.4 (lEQHSQQPQLSPATSGRGLRDSPSssEDR)

   Ion         m/z   Intensity
   b4++      369.26       14.9
   b5++      412.81       15.0
   b2+       472.38       18.8
   b6++      476.97       19.8
   a7++      526.73       16.8
   y13+++    540.77       46.3
   b3+       600.43       18.7
   b9++      653.53       20.8
   b10++     710.16       22.5
   b4+       737.44       18.4
   y21+++    788.16       26.4
   b20+++    801.29       19.0
   y22+++    821.06       30.7
   a13++     822.95       15.4
   b5+       824.16       18.6
   a21+++    829.79       15.2
   y14++     839.60       51.1
   y6+       840.33       35.9
   b22+++    868.15       24.3
   a14++     873.74       34.0
   b23+++    900.58       15.0
   b6+       952.71       24.1
   y17++     969.12       21.9
   y26+++    980.56       18.8
   b25+++    985.42       18.3
   y18++    1017.71       18.5
   y8+      1024.25       24.0
   b17++    1037.91       15.0
   y19++    1061.57       21.2
   b7+      1080.59       29.2
   b27+++   1084.22       25.2
   y20++    1117.83       13.7
   b19++    1122.71       24.3
   y21++    1181.81       46.5
   y22++    1230.77       52.1
```

```
906. MORG-339_phosphopeptide_TMT_22%ACN_032017_OT.16354.16354.4 (eELAsDLEEMATSSAkRk)

   Ion         m/z   Intensity
   y9+++     480.01       14.8
   y4++      480.78        8.4
   b2+       488.22       12.2
   y10+++    523.20       15.4
   b7++      534.54        7.5
   y11+++    566.08       16.1
   y6++      567.95       13.8
   b14+++    601.44       14.2
   y12+++    603.77       29.1
   y7++      618.54       18.5
   b15+++    625.06       13.7
   y13+++    642.22       26.1
   a4+       644.16        8.4
   a9++      649.70       12.3
   y8++      654.38       16.1
   b9++      663.39       14.9
   b4+       672.40       20.7
   y9++      719.49       24.8
   y16+++    759.27       12.4
   y10++     783.99       24.3
   b17+++    796.27       11.5
   y17+++    802.58       13.4
   b12++     815.27        9.4
   b5+       839.50       10.3
   y11++     848.69       21.2
   b6+       954.57       14.4
   b7+      1067.41       10.7
   y6+      1134.49        7.5
   b9+      1325.65        7.7
```

```
907. MORG-339_phosphopeptide_TMT_22%ACN_032017_OT.19138.19138.3 (iVETINSDsDSEFGIPk)

   Ion         m/z   Intensity
   b2+       442.48        9.4
   y2+       473.34       47.0
   a3+       543.83        7.6
   b3+       571.56        9.1
   y3+       586.55        7.1
   b9++      635.12       11.9
   y4+       643.51       15.2
   y9++      644.95        8.6
   a11++     721.46        8.5
   y11++     745.93       11.9
   a5+       757.60       11.7
   b5+       785.23       28.9
   a12++     786.23       15.2
   y5+       790.70        8.4
   y12++     802.98        6.7
   b13++     873.89       22.0
   b6+       899.57        9.2
   b14++     902.50       46.2
   y14++     909.89       15.7
   a15++     945.35       17.4
   b15++     959.07       70.5
   b8+      1101.56        8.0
   b9+      1268.57        6.9
   b10+     1383.85        6.4
   b12+     1599.82        9.9
```

```
908. MORG-339_phosphopeptide_TMT_30%ACN_032017_OT.14499.14499.4 (iVETINSDsDSEFGIPkkTTTPk)

   Ion         m/z   Intensity
   y2+       473.39       14.7
   y8+++     530.39       27.2
   y9+++     567.75       20.2
   b3+       571.27       17.9
   y10+++    586.88       17.4
   y11+++    636.00       34.5
   b4+       672.51       12.8
   y12+++    678.82       16.6
   y13+++    708.07       40.1
   a11++     722.15       15.9
   b11++     736.00       12.7
   y14+++    746.55       29.3
   y5+       776.56       12.6
   a17+++    781.97       18.3
   b5+       785.29       20.5
   a12++     786.39       18.3
   y8++      794.69       46.7
   b12++     800.07       13.1
   y15+++    801.80       27.1
   y9++      850.87       39.2
   a13++     860.35       31.9
   b6+       899.69       14.9
   b18+++    910.43       14.6
   b19+++    943.43       21.4
   a15++     944.93       18.3
   y11++     952.78       16.9
   b20+++    977.95       16.1
   y20+++    978.78       26.7
   b7+       986.98       17.0
   a21+++   1001.58       12.6
   b21+++   1011.29       23.1
   y12++    1017.56       14.4
   y13++    1060.86       16.0
   b8+      1101.74       18.5
   y14++    1119.12       13.3
```

```
909. MORG-339_phosphopeptide_TMT_16%ACN_032017_OT.3412.3412.3 (rRsPSPAPPPPPPPPPPR)

   Ion         m/z   Intensity
   y8++      427.69       13.1
   y9++      476.49       22.3
   a7++      517.41       12.9
   y10++     524.92       20.4
   b7++      531.22       16.8
   y5+       563.45       14.5
   y11++     573.52       33.4
   b8++      579.87       14.2
   b9++      628.30       11.6
   y6+       660.29       14.6
   a10++     662.52       14.3
   b10++     676.93       20.2
   a3+       681.42       13.4
   y14++     700.64       23.7
   b3+       709.51       32.4
   a11++     711.43       54.7
   b11++     724.92       36.4
   y7+       757.50       12.0
   a13++     808.77       41.3
   b13++     822.07       13.8
   y8+       854.58       17.7
   b14++     871.38       12.8
   y9+       951.50       16.9
   b6+       990.44       19.4
   y10+     1048.59       12.0
   b7+      1061.56       30.4
   y11+     1145.68       16.9
```

```
910. MORG-339_phosphopeptide_TMT_16%ACN_032017_OT.8273.8273.2 (atSPSTLVSTGPSSR)

   Ion         m/z   Intensity
   b1+       301.26       11.1
   y5+       503.35       11.1
   b3+       569.23        9.0
   y7+       691.36       19.5
   y8+       790.26       17.1
   a6+       826.29       18.4
   b6+       854.19       23.6
   y9+       903.42       13.4
   a7+       939.57       12.2
   y10+     1004.57       13.3
   b8+      1066.55       18.2
   y12+     1188.60       50.3
   y14+     1456.67       10.1
```

```
911. MORG-339_phosphopeptide_TMT_10%ACN_032017_OT.9839.9839.4 (gkQADVADQQTTELPAENGETENQsPAsEEEkEAk)

   Ion         m/z   Intensity
   b5++      479.92       31.9
   b8++      622.32       55.3
   y7++      660.95       27.2
   b9++      686.57       24.1
   a16+++    695.62       31.3
   b10++     750.66       45.5
   y9++      780.18       51.0
   b11++     801.19       41.0
   y10++     828.70       57.7
   b4+       843.52       22.5
   b12++     851.67       29.1
   a13++     902.33       26.6
   b13++     916.26      100.3
   y20+++    932.84       51.7
   y4+       933.70       23.7
   b5+       958.67       45.3
   y21+++    964.81       27.3
   b14++     972.79      144.1
   b15++    1021.26       23.3
   b6+      1057.28       36.0
   b17++    1121.03       37.9
   y15++    1147.82       31.4
   y26+++   1155.09       25.1
   b28+++   1172.75       28.0
   a19++    1192.31       58.0
   y17++    1240.92       49.4
   b8+      1243.68       46.8
   y29+++   1260.67       36.1
   a22++    1371.92       22.5
   y33+++   1397.49       24.1
   y20++    1398.46       92.0
   a33+++   1454.65       24.9
   y8+      1488.11       30.2
   b10+     1499.55       31.5
   y9+      1558.99       26.8
   b13+     1830.89       24.5
   b14+     1943.96       40.1
```

```
912. MORG-339_phosphopeptide_TMT_8%ACN_032017_OT.14124.14124.4 (sEPDRGRLtPsPDIIVLsDNEASSPR)

   Ion         m/z   Intensity
   b1+       317.26        9.0
   y3+       359.19        9.7
   b2+       446.26       12.9
   y5+       517.34       14.9
   b11+++    529.34       16.4
   y10++     578.51       17.0
   b13+++    599.99       13.8
   a14+++    627.95       16.6
   b14+++    637.86       24.4
   y6+       646.39       11.6
   b4+       658.38        9.2
   b9++      661.67       18.0
   a15+++    666.24       15.4
   b15+++    675.60       41.8
   y12++     684.23       10.6
   a16+++    699.33       11.6
   b16+++    708.61       47.3
   b10++     709.47       13.7
   a17+++    736.98       10.5
   y13++     740.41       10.6
   b17+++    746.34       31.4
   y7+       760.44       17.0
   a5+       786.33       15.4
   b11++     793.51       15.5
   y14++     798.44       11.5
   b18+++    801.41       17.3
   y20+++    803.31       23.2
   a19+++    830.90        8.0
   b19+++    840.34       17.6
   y15++     847.22       10.7
   y22+++    874.08       10.1
   y8+       875.47       11.1
   b20+++    878.03       13.5
   b13++     899.46       10.7
   y23+++    912.35       16.9
   b21+++    921.39       27.0
   y16++     930.04       16.2
   a22+++    935.11       11.2
   a14++     941.61       13.0
   b22+++    944.93       24.9
   b14++     956.16       21.9
   a23+++    964.51       11.1
   b23+++    974.26       12.5
   y17++     979.34       19.9
   a15++     998.28        9.4
   b24+++   1003.00       10.3
   b15++    1012.73       24.0
   a25+++   1025.37        8.5
   y9+      1042.34       20.2
   b16++    1062.26       29.5
   b17++    1118.99       14.4
   y10+     1155.45       17.9
   y11+     1254.52       12.7
   y23++    1367.72        8.3
```

```
913. MORG-339_phosphopeptide_TMT_18%ACN_032017_OT.13278.13278.3 (dAsPPEPAsPTIGLDkk)

   Ion         m/z   Intensity
   y2++      367.26       14.5
   y5++      510.12       21.0
   y6++      566.48       12.0
   b3+       583.12       13.6
   y8++      665.88       30.4
   y2+       733.77       14.6
   y10++     784.25       11.8
   b13++     805.74       14.1
   y11++     833.38       42.5
   y3+       848.55       27.7
   a6+       878.86       16.0
   a15++     905.16       13.6
   b6+       906.43       13.9
   y13++     946.21       30.6
   y4+       961.99       10.7
   a7+       975.40       12.9
   y14++     994.90       39.8
   y15++    1078.42       11.8
```

```
914. MORG-339_phosphopeptide_TMT_12%ACN_032017_OT.14569.14569.3 (gLVAAYSGDsDNEEELVER)

   Ion         m/z   Intensity
   y2+       304.19       19.6
   a2+       372.31        7.3
   b2+       400.33        7.4
   y3+       403.28       20.8
   b7++      446.41       11.8
   b3+       499.37       14.2
   y4+       516.35       19.5
   b4+       570.51       14.4
   a10++     601.45        7.0
   b5+       641.49       20.0
   y5+       645.44       15.4
   a11++     659.22        8.6
   b11++     673.40        8.8
   y11++     707.97        9.4
   a12++     716.63        7.6
   y12++     736.48       22.9
   y6+       774.47       19.3
   a6+       776.27       18.7
   y13++     779.93       15.0
   b13++     795.22       11.4
   b6+       804.58       27.0
   b14++     859.62        9.8
   y14++     861.69       13.4
   b7+       891.50       17.6
   y7+       903.50       10.4
   b15++     924.16       33.4
   y16++     932.75        7.1
   b8+       948.58        8.5
   b16++     980.73       38.6
   y17++     981.52       20.7
   y8+      1017.45       10.3
   b17++    1030.27       44.5
   b18++    1094.33       10.5
   y9+      1132.61        8.5
   y10+     1299.57        9.0
   b11+     1345.78        7.5
```

```
915. MORG-339_phosphopeptide_TMT_16%ACN_032017_OT.4143.4143.4 (qSRPVAGGPGAPPAARPPAsPSPQRQAGAPQAtR)

   Ion         m/z   Intensity
   b8++      491.90        8.7
   y4+       555.11        8.3
   b10++     569.21       13.9
   a11++     590.61       14.3
   b11++     604.53       19.8
   y11++     632.02       14.9
   b12++     653.32       11.6
   b19+++    656.15       14.8
   b4+       698.46        8.4
   a20+++    701.60       13.0
   y19+++    711.72        8.9
   y13++     724.23       24.7
   a21+++    734.49        9.9
   y21+++    758.88       12.0
   y14++     772.80       15.0
   y22+++    791.33       19.7
   y23+++    824.14       18.3
   b16++     850.52       15.9
   y8+       851.76       18.1
   y15++     855.88       12.3
   y25+++    866.57       20.1
   a17++     885.07       22.7
   y16++     892.25       17.2
   a7+       897.27       23.9
   b17++     898.84       23.1
   b25+++    900.18       34.9
   y17++     940.39       10.7
   y29+++    960.32       11.3
   a28+++    975.80        9.8
   y9+       979.52        8.6
   b19++     982.78       10.2
   b28+++    985.01        9.2
   y18++     989.09        9.1
   a20++    1052.65        8.1
   y19++    1067.15       10.9
   b31+++   1083.91        9.8
   a21++    1100.92       10.5
   y33+++   1106.91       12.6
   y10+     1135.84       15.4
   y21++    1137.91       15.6
   a11+     1179.58       10.7
   y22++    1186.77       21.1
   b11+     1207.75       12.2
   y23++    1235.15       14.4
   y11+     1263.20       11.2
```

```
916. MORG-339_phosphopeptide_TMT_4%ACN_032017_OT.5757.5757.3 (rASYDSVEDSsESENsPVAR)

   Ion         m/z   Intensity
   b5++      412.05       13.4
   y4+       442.37       12.4
   a7++      490.80        7.6
   b8++      569.69       11.6
   y5+       609.26        6.2
   b9++      626.83       15.2
   y11++     662.08        6.1
   b10++     670.56       10.8
   b4+       707.35        6.5
   y6+       723.42        6.8
   b11++     754.01       10.0
   b12++     818.04       11.8
   b5+       822.37        7.8
   a13++     847.37       11.4
   b14++     926.54        6.4
   y16++     934.73        7.3
   y8+       939.57        5.6
   a15++     969.15       14.2
   b15++     983.82       10.8
   b7+      1008.74        8.0
   y9+      1068.52        6.0
   b8+      1137.67        6.3
   b19++    1200.95        9.9
   y11+     1322.59        8.1
   b10+     1339.77        4.9
   y15+     1752.84        7.7
```

```
917. MORG-339_phosphopeptide_TMT_10%ACN_032017_OT.9602.9602.3 (sPsPAPPPPPPPPPPR)

   Ion         m/z   Intensity
   a3++      317.27       10.8
   b3++      330.91       12.2
   b4++      379.25       13.5
   b5++      415.25       14.1
   y8++      427.90       15.2
   b6++      463.77       11.1
   y9++      476.58       36.1
   b7++      512.42       12.9
   y10++     525.08       34.3
   b8++      560.85       19.1
   y5+       563.31       20.4
   y11++     573.59      166.7
   a9++      595.63       10.1
   y12++     608.62       12.2
   a3+       632.85       16.1
   b11++     706.50        7.2
   a4+       729.81       11.0
   y7+       757.53       16.1
   b4+       758.52       15.3
   a5+       801.46       12.7
   b5+       829.32      108.0
   y8+       854.71       12.5
   b6+       926.42       21.4
   y9+       951.72       12.5
   b7+      1023.44       19.9
   b8+      1120.55       11.4
```

```
918. MORG-339_phosphopeptide_TMT_18%ACN_032017_OT.11061.11061.2 (iYQyIQSR)

   Ion         m/z   Intensity
   a1+       315.28        7.6
   b1+       343.36       11.9
   y3+       390.37        8.0
   b2+       506.41       11.1
   a3+       606.72        8.9
   b3+       634.50       10.5
   y5+       746.30       10.3
   y6+       874.63       11.2
   b4+       877.46       11.8
   b5+       990.48       11.1
   y7+      1037.48       26.6
   b6+      1118.55       18.1
   b7+      1205.66        9.7
```

```
919. MORG-339_phosphopeptide_TMT_8%ACN_032017_OT.1443.1443.4 (qQEQTAQGTAPDAVDQQRRDsR)

   Ion         m/z   Intensity
   y7+++     342.33        8.3
   b4++      372.48        8.6
   b9+++     400.98        9.5
   y9+++     413.91       30.2
   b5++      422.81       23.5
   y6++      449.35       41.7
   a2+       458.40       21.7
   b2+       486.45       11.8
   a7++      508.47       10.6
   y7++      513.45       23.9
   b7++      522.37       12.4
   b8++      550.92       26.5
   y15+++    584.82        8.6
   b15+++    590.35       10.6
   b9++      601.59       17.2
   b3+       615.29       19.3
   y16+++    627.68       38.5
   y17+++    650.82       31.5
   y10++     655.45       21.8
   a17+++    666.93       60.0
   a11++     671.41       25.0
   b17+++    675.72       23.1
   y18+++    684.42       27.1
   b11++     685.35       31.5
   y11++     713.18       15.3
   y19+++    727.23       12.4
   b4+       743.46       18.7
   y12++     762.09       49.8
   a13++     764.74        9.8
   y13++     797.61       22.5
   y21+++    812.65       13.7
   b20+++    818.54       12.3
   b14++     827.66        8.8
   b5+       844.56       14.6
   y14++     848.04       29.9
   b21+++    874.47        7.7
   y15++     876.64       52.9
   b6+       915.55       22.8
   y16++     940.67       47.8
   y17++     976.22       16.0
   y18++    1026.75       12.9
```

```
920. MORG-339_phosphopeptide_TMT_12%ACN_032017_OT.10448.10448.2 (rLsQSsLESATSPScS)

   Ion         m/z   Intensity
   y4+       450.19       14.5
   y5+       537.23       12.5
   a3+       638.38        7.2
   b4+       794.49        7.5
   y8+       796.45       12.6
   a5+       853.44        7.6
   b5+       881.50        8.1
   y9+       925.33        8.9
   a6+      1020.75       14.5
   a8+      1262.67        9.3
   b8+      1290.61       13.9
   b10+     1448.74        7.7
   a11+     1521.78        7.7
   b11+     1549.77        8.9
   b12+     1636.79        6.9
```

```
921. MORG-339_phosphopeptide_TMT_4%ACN_032017_OT.12069.12069.2 (mGLEEssEGEAEGSVR)

   Ion         m/z   Intensity
   y3+       361.34        8.8
   y4+       418.29       13.6
   b3+       531.37        7.3
   y6+       618.42       15.1
   b4+       660.50        7.6
   b5+       789.54        8.2
   y8+       804.43       18.6
   y9+       933.53       12.0
   y10+     1100.55       15.1
   y11+     1267.54        6.9
   a10+     1410.55        8.9
   y14+     1638.62       10.7
   y15+     1695.68        7.0
```

```
922. MORG-339_phosphopeptide_TMT_18%ACN_032017_OT.10683.10683.3 (sVVFTSARATTEA)

   Ion         m/z   Intensity
   y2+       219.21        7.8
   b2++      248.14       12.7
   y3+       320.19       16.8
   b4++      371.40        6.2
   y8++      403.59        7.8
   y4+       421.46       10.5
   b5++      422.37       13.8
   b6++      465.63        8.4
   a7++      487.53        6.7
   y5+       492.42       16.2
   b7++      501.22       13.2
   y10++     527.60       17.2
   b8++      579.31       13.7
   b3+       595.39       11.3
   b9++      614.77       15.7
   y6+       648.34       10.0
   b10++     665.46       47.0
   a11++     702.33        7.7
   b11++     716.08       20.3
   y7+       719.29        8.6
   b4+       742.49       11.7
   b12++     780.48       55.2
   y8+       806.37       10.7
   b5+       843.40        6.3
   y9+       907.53        6.9
```

```
923. MORG-339_phosphopeptide_TMT_14%ACN_032017_OT.12455.12455.4 (sEkRPPSPDVIVLsDSEQPSsPR)

   Ion         m/z   Intensity
   y2+       272.29       11.2
   y3+       439.29        9.2
   b9+++     511.78       19.0
   y13+++    525.41       18.1
   a10+++    535.41       10.4
   b10+++    544.78       24.3
   b11+++    582.60       27.7
   b12+++    615.65       34.9
   y5+       623.33       47.7
   y10++     625.44        9.3
   a13+++    643.52       11.6
   b13+++    653.21       29.3
   b7++      660.93       10.0
   y11++     681.51       17.1
   y12++     731.33       12.7
   a15+++    737.52       13.7
   b15+++    747.20       28.9
   y6+       751.33       12.8
   a16+++    766.26       17.2
   b9++      767.27       22.5
   y20+++    775.29       14.5
   b16+++    776.01       23.0
   y13++     787.77       20.0
   b10++     816.39       18.7
   b17+++    819.30       26.5
   a18+++    852.66       13.6
   b18+++    861.94       73.9
   b11++     873.21       24.8
   y7+       880.47       10.2
   a19+++    885.02       11.4
   b19+++    894.34       14.8
   y15++     895.06       13.2
   a20+++    913.65       15.7
   b20+++    922.84       31.4
   y16++     943.30       14.4
   y8+       967.45       18.8
   b13++     979.28       20.1
   y9+      1082.45       10.0
   b17++    1228.00       13.9
   y10+     1249.47       16.1
   a7+      1292.36       12.3
   y12+     1461.53       10.0
```

```
924. MORG-339_phosphopeptide_TMT_8%ACN_032017_OT.24129.24129.2 (aEQYPGsPDsWV)

   Ion         m/z   Intensity
   b3+       558.32       15.6
   y5+       683.03       13.9
   b4+       721.43       39.0
   a5+       790.54       14.5
   b5+       817.97       12.1
   a6+       847.41       17.4
   y6+       850.21       18.7
   b6+       875.40       13.6
   y7+       907.31        8.6
   y8+      1004.30       39.4
   b7+      1042.56        7.7
   b8+      1139.48        9.3
   y9+      1167.33       16.3
   b9+      1254.55       12.0
```

```
925. MORG-339_phosphopeptide_TMT_20%ACN_032017_OT.11797.11797.3 (hLYISSsNPDLITR)

   Ion         m/z   Intensity
   y2+       276.19       13.2
   y6++      358.07        9.5
   a4++      364.58       11.5
   b1+       367.14        7.2
   b4++      378.77       12.1
   y3+       389.22       12.5
   b5++      422.29       11.6
   b2+       480.36        9.2
   y4+       502.46        7.3
   y9++      542.08       10.4
   y10++     585.30       19.3
   a8++      592.52       13.3
   b8++      606.59       12.2
   a3+       615.68       13.0
   y5+       617.76       23.6
   b9++      655.13        8.2
   b10++     712.55       12.9
   y6+       714.46       23.9
   y12++     723.40        7.0
   b4+       756.62       11.1
   b11++     769.05       12.6
   a12++     811.46       10.2
   b12++     825.69       25.2
   y7+       828.43       14.0
   b5+       843.71       14.3
   y8+       995.51        9.9
   y9+      1082.66        9.9
   y10+     1169.48       10.2
```

```
926. MORG-339_phosphopeptide_TMT_22%ACN_032017_OT.17301.17301.2 (sAsTLcLPEVGLAR)

   Ion         m/z   Intensity
   b1+       317.20        8.8
   y4+       416.04        7.5
   y7+       741.51       15.2
   b5+       769.47        7.2
   y8+       854.83       12.7
   b6+       929.55        9.1
   y9+      1014.71       12.3
   b7+      1042.52       10.7
   y10+     1127.54        9.8
   y11+     1228.72       13.6
   b9+      1268.76        8.3
   y12+     1395.71        8.4
   y13+     1466.67        7.2
```

```
927. MORG-339_phosphopeptide_TMT_16%ACN_032017_OT.15989.15989.3 (ikEASPEsEDEEEALPcTDWEN)

   Ion         m/z   Intensity
   b5++      494.36        7.8
   b6++      543.27        8.5
   y5+       664.35        8.3
   b9++      755.39        7.8
   b10++     812.52        9.4
   b3+       829.52        8.0
   b11++     877.70       16.2
   b4+       900.57       11.2
   y7+       921.32       99.4
   b12++     942.21       22.0
   y15++     952.26       10.6
   a5+       959.66        9.3
   b5+       987.27        9.5
   a13++     992.86       31.2
   b13++    1006.33       29.1
   y16++    1016.43       12.7
   a14++    1027.74       10.5
   b6+      1084.65       23.2
   b15++    1098.75      168.4
   y9+      1105.45       19.1
   y18++    1108.60        8.6
   y19++    1144.74        9.8
   b16++    1147.31       14.6
   y20++    1209.01       11.7
   b7+      1213.64       14.2
   b17++    1227.42        8.7
   y10+     1234.48       10.9
   b18++    1277.87        8.2
   b19++    1335.44       16.9
   y11+     1363.46        8.3
   b20++    1428.05       13.8
   b21++    1493.09       11.7
```

```
928. MORG-339_phosphopeptide_TMT_10%ACN_032017_OT.14823.14823.3 (dQNLPQILEESRsE)

   Ion         m/z   Intensity
   a1+       317.16        7.4
   a4++      337.15        6.2
   b6++      463.34        7.1
   y7++      465.14        8.0
   y3+       471.44        9.5
   b7++      519.91       21.4
   y8++      521.92        9.1
   y4+       558.36       12.4
   a3+       559.22        6.3
   b8++      576.26       20.5
   y9++      585.66        6.9
   b3+       587.34       11.7
   a9++      626.80       15.4
   y10++     634.29       19.2
   b9++      640.64       16.5
   y5+       687.30       15.0
   a10++     691.18       11.4
   b4+       700.38       13.9
   b10++     705.37       10.8
   b11++     748.83       17.8
   y6+       816.28       10.4
   b12++     827.04       20.3
   b6+       925.59       11.2
   y7+       929.25        7.8
   b7+      1038.72        5.6
   y8+      1042.23        9.3
   b8+      1151.72        8.1
   y10+     1267.45        7.0
```

```
929. MORG-339_phosphopeptide_TMT_18%ACN_032017_OT.14240.14240.3 (ktSLVIVESTDDQSQVFER)

   Ion         m/z   Intensity
   y3+       451.36       14.9
   y4+       550.68       13.5
   b9++      634.03       12.9
   y11++     656.78       17.1
   b4+       739.67       16.1
   y6+       765.50       16.0
   b12++     799.46       12.4
   b13++     863.46       11.3
   y15++     876.55       14.9
   a14++     892.48       25.7
   y7+       893.58       25.9
   b15++     970.54       17.5
   a16++    1006.09       11.5
   y8+      1008.48       19.7
   y9+      1123.55       18.3
   y10+     1224.70       15.1
   y11+     1311.62       23.1
   y12+     1440.71       15.3
   y13+     1539.77       20.2
   y14+     1652.75       19.6
```

```
930. MORG-339_phosphopeptide_TMT_10%ACN_032017_OT.8263.8263.3 (sPSPPPDGSPAATPEIR)

   Ion         m/z   Intensity
   y8++      427.85       35.4
   y9++      471.55       12.8
   y10++     499.85        9.5
   a8++      508.32       19.8
   y4+       514.32       30.2
   y11++     557.32       27.0
   b9++      566.47       10.6
   b3+       581.38       15.9
   a10++     600.71       12.7
   y12++     605.48       15.3
   y5+       615.32       28.5
   a4+       650.47       55.9
   y13++     654.40       15.3
   b4+       678.64       11.0
   b12++     685.79       15.0
   y6+       686.56       18.7
   y14++     702.93       20.8
   b13++     736.31       23.0
   y7+       757.59       14.3
   y8+       854.50       31.1
   y10+      998.65        9.7
   y11+     1113.60       22.0
   b9+      1131.43       23.6
```

```
931. MORG-339_phosphopeptide_TMT_14%ACN_032017_OT.16800.16800.4 (aLsARsLRHEAESDFDVADLPQLER)

   Ion         m/z   Intensity
   y5++      321.67        8.2
   y6++      378.52        8.6
   y3+       417.25        9.5
   a6++      474.01        7.8
   b7++      544.49        7.9
   y4+       545.40       10.9
   b13+++    600.16        9.5
   b8++      622.88        7.7
   a4+       624.49        8.4
   b14+++    638.45       23.0
   y5+       642.42       48.0
   a15+++    678.28       11.6
   b15+++    687.65       21.3
   b16+++    726.03       54.5
   b10++     755.66       33.4
   b17+++    758.87       60.4
   a18+++    772.73        8.5
   a5+       780.24       55.6
   b18+++    782.55       60.3
   b11++     791.59       15.8
   b19+++    820.94       51.1
   a20+++    849.51       18.7
   y15++     852.52       11.7
   b12++     856.14        9.5
   b20+++    858.64       75.0
   y7+       870.51       34.2
   y22+++    879.00       12.9
   b21+++    891.07       16.4
   a22+++    924.36       10.6
   b22+++    933.79       16.6
   y8+       941.50       37.4
   a14++     942.54       30.9
   b14++     957.36       14.2
   b23+++    971.29       19.5
   b24+++   1014.56       20.9
   a15++    1016.63        8.8
   b15++    1030.88        9.6
   y9+      1040.32       20.8
   b7+      1088.35       19.7
   b17++    1137.93       21.9
   y10+     1155.60       15.0
   b18++    1173.05       12.6
   b19++    1230.73       13.9
   b20++    1287.29       17.0
   y11+     1302.81       11.4
   y13+     1504.87       10.5
```

```
932. MORG-339_phosphopeptide_TMT_10%ACN_032017_OT.15624.15624.3 (lEkEEEEGIsQEssEEEQ)

   Ion         m/z   Intensity
   b3++      415.25       14.4
   b5++      544.58       11.6
   b6++      609.06       20.0
   b7++      673.36       16.1
   b9++      758.63       51.9
   y6+       868.16       14.1
   a11++     891.91       10.6
   b11++     905.56       24.6
   y14++     925.61       13.2
   a12++     956.16        9.7
   b4+       959.01        9.6
   b12++     970.44       15.5
   y7+       997.40       15.6
   b13++    1054.27       10.4
   y8+      1125.57        9.7
   y16++    1168.36       10.7
   b15++    1202.18       20.9
   b16++    1266.84       42.6
   y9+      1292.30       25.3
   b17++    1331.31       54.4
   y12+     1591.55       11.2
```

```
933. MORG-339_phosphopeptide_TMT_14%ACN_032017_OT.14028.14028.2 (lTNLSSNsDV)

   Ion         m/z   Intensity
   b1+       343.34        7.1
   a2+       416.24        8.2
   b2+       444.54        9.3
   y4+       514.17       10.1
   b3+       558.39       13.0
   y5+       601.20       19.1
   a4+       643.41        7.3
   b4+       671.43       35.0
   y6+       688.24       12.5
   b5+       758.53       20.9
   y7+       801.37       11.2
   b6+       845.64       12.2
   y8+       915.40        9.7
   a7+       931.32        6.3
   b7+       959.66       12.4
   y9+      1016.45       10.3
   b8+      1126.55        6.8
   b9+      1241.60        7.9
```

```
934. MORG-339_phosphopeptide_TMT_8%ACN_032017_OT.6799.6799.3 (dSRSLsySPVER)

   Ion         m/z   Intensity
   a1+       317.21        8.8
   b1+       345.23        8.0
   a5++      380.73        5.8
   a2+       404.40        7.0
   y6++      415.98        6.4
   b6++      478.44        6.5
   y4+       500.21       12.2
   y8++      555.62       11.6
   a3+       560.39        8.0
   a7++      585.88       27.7
   y5+       586.82       29.3
   b3+       587.84        9.1
   b8++      643.27       10.4
   a9++      677.35        5.8
   b9++      692.17        6.3
   y11++     720.82        5.5
   a10++     727.27        5.9
   b10++     741.27        5.4
   b11++     805.83        8.9
   y6+       830.37        7.6
```

```
935. MORG-339_phosphopeptide_TMT_4%ACN_032017_OT.2993.2993.3 (dEHGLEPAsPR)

   Ion         m/z   Intensity
   y2+       272.19       28.7
   y5++      304.16        9.5
   b3++      306.29        8.6
   b4++      334.67       15.9
   b1+       345.26       13.5
   a5++      377.12       12.8
   b5++      391.34       14.0
   y7++      425.27        7.6
   a6++      441.61        8.3
   a2+       446.59       11.7
   b6++      455.72       33.1
   b2+       473.77       66.3
   a7++      489.93       25.5
   y9++      522.22        7.7
   a3+       583.12        5.7
   y10++     586.82       13.6
   y5+       607.37       24.7
   b3+       611.32       17.6
   b4+       668.42        7.0
   y6+       736.55        8.7
   y7+       849.49        5.8
   y8+       906.48       15.6
   b6+       910.47       18.2
```

```
936. MORG-339_phosphopeptide_TMT_22%ACN_032017_OT.15573.15573.4 (fGIHVYQFPEcDsDEDEDFkQQDR)

   Ion         m/z   Intensity
   b4++      342.85       38.5
   b5++      392.50       38.5
   b2+       434.36       15.4
   a10+++    473.96       32.7
   a7++      524.01       22.0
   y6++      525.80       21.1
   b7++      538.01       69.5
   y4+       546.43       14.5
   b3+       547.40       35.1
   a8++      597.53       30.3
   b8++      611.57       64.5
   y8++      647.84       27.0
   b4+       684.50       37.6
   b10++     724.42       19.4
   y15+++    742.31       44.4
   y10++     770.16       31.8
   y16+++    774.92       56.7
   a17+++    783.50       36.0
   y17+++    824.13       39.9
   a12++     847.96       26.5
   b12++     861.82       44.9
   y18+++    865.87       21.5
   a19+++    871.15       17.8
   y12++     911.10       31.7
   y19+++    920.69       16.6
   b6+       946.54       24.9
   y20+++    953.81       20.3
   b20+++    999.72       26.0
   b14++    1003.27       23.7
   a21+++   1033.57       14.5
   y6+      1050.62       17.9
   y23+++   1056.17       15.1
   b7+      1074.70       27.5
   a22+++   1075.71       30.3
   y15++    1112.80       89.3
   a23+++   1114.49       26.2
   b16++    1125.28       26.3
   y16++    1161.73       86.3
   b8+      1221.72       33.9
   y17++    1235.35       44.7
   b18++    1247.48       21.3
   a9+      1290.81       41.6
   y8+      1294.78       18.3
   y19++    1380.43       18.6
```

```
937. MORG-339_phosphopeptide_TMT_6%ACN_032017_OT.7365.7365.4 (sGDETPGSEAPGDkAVEEQGDDQDSEk)

   Ion         m/z   Intensity
   y1+       376.36       27.6
   y10+++    460.27       16.9
   y2+       505.41       24.3
   a8++      506.37       16.4
   y8++      562.08       18.1
   y3+       592.31       37.8
   b10++     620.71       23.9
   b14+++    623.41       13.8
   y9++      625.95       24.1
   b15+++    647.11       20.1
   b16+++    679.88       21.5
   y10++     690.56       27.0
   b4+       698.29       27.5
   y4+       707.41       34.4
   y11++     754.83       14.2
   b13++     755.47       18.4
   y17+++    769.37       56.5
   a5+       771.22       17.4
   y18+++    793.19       36.4
   b5+       799.42       41.7
   y12++     804.58       18.9
   b20+++    827.68       34.9
   y19+++    835.44       23.4
   b21+++    865.39       23.6
   a6+       868.34       18.6
   y22+++    916.48       49.5
   b14++     934.26       28.8
   b23+++    947.08       29.3
   y6+       950.33       27.1
   b7+       953.33       20.9
   b15++     969.93       25.0
   y24+++    993.11       14.0
   a8+      1012.60       14.7
   b25+++   1014.44       30.1
   b16++    1019.19       55.0
   b26+++   1057.33       16.8
   y7+      1065.54       20.7
   b17++    1083.78       47.7
   y16++    1104.52       14.4
   y8+      1122.46       29.9
   b18++    1148.25       33.8
   y17++    1153.37       66.4
   b9+      1169.46       28.3
   y18++    1188.82       22.4
   a10+     1212.32       41.1
   b20++    1240.53       48.2
   b21++    1298.49       22.6
   y11+     1508.70       19.2
   b13+     1509.70       19.4
```

```
938. MORG-339_phosphopeptide_TMT_4%ACN_032017_OT.6532.6532.3 (eANELsDSAGEDHPAELR)

   Ion         m/z   Intensity
   y2+       288.24       11.6
   b4++      336.68        6.0
   y7++      419.22        9.0
   y9++      512.40       11.2
   y5+       585.51       16.8
   y11++     591.41       16.0
   a10++     628.35        5.5
   y12++     648.96       11.0
   b4+       673.37        9.4
   y13++     732.32       28.0
   b5+       786.54        7.1
   b13++     832.86       12.7
   y7+       837.61        9.7
   b16++     981.36       19.8
   b7+      1068.58        6.0
   y11+     1181.55        5.6
```

```
939. MORG-339_phosphopeptide_TMT_16%ACN_032017_OT.17374.17374.3 (lFSRQDccGNcsDsEEELPTRL)

   Ion         m/z   Intensity
   y4+       486.48       17.6
   b6++      488.86       11.7
   y5+       599.35       14.2
   b8++      648.76       12.0
   y6+       728.58       11.9
   b10++     734.54       12.9
   b11++     814.38        7.8
   b5+       861.44        9.0
   a13++     941.31       12.4
   b6+       976.44       18.1
   y8+       986.28       21.9
   y16++    1043.98        8.5
   b15++    1103.88        8.3
   b7+      1136.69       13.7
   a16++    1153.50       13.3
   a17++    1218.37        8.0
   b17++    1232.71       16.4
   b18++    1289.24       18.9
   a9+      1325.79        8.5
   y21++    1360.73        8.1
   y15+     1926.59        8.7
```

```
940. MORG-339_phosphopeptide_TMT_14%ACN_032017_OT.5952.5952.4 (skPAAADsEGEEEEEDTAkEk)

   Ion         m/z   Intensity
   b5++      457.60       12.7
   b6++      492.86       11.5
   y5++      518.03       26.3
   b7++      550.55       24.3
   y6++      575.75       10.8
   y7++      640.12       16.1
   y8++      704.56       14.8
   y14+++    716.70       10.1
   a3+       743.38       14.3
   y15+++    754.79       17.4
   a17+++    763.10       38.5
   y9++      769.14       22.8
   b3+       771.36       17.5
   b17+++    772.41       15.3
   a11++     777.20       26.5
   a18+++    786.00       30.3
   b11++     791.65       40.7
   b18+++    795.69       36.3
   y17+++    802.05       32.1
   y10++     833.64       17.7
   a12++     841.45       10.8
   b12++     856.11       14.0
   y3+       862.73       16.4
   b5+       913.70       13.1
   y12++     926.28       18.2
   y4+       933.16       12.8
   b20+++    957.88       13.1
   y13++     991.25       20.3
   y5+      1034.92       11.5
   b15++    1049.55       10.1
   y14++    1074.72       20.3
   b16++    1107.32       18.8
   a8+      1238.51       10.9
```

```
941. MORG-339_phosphopeptide_TMT_4%ACN_032017_OT.7974.7974.3 (eVENEQTPVsEPEEEk)

   Ion         m/z   Intensity
   y3++      317.23        5.8
   y1+       376.32        8.7
   y5++      430.60       12.4
   y2+       505.40        8.8
   b7++      530.69       11.6
   b8++      578.83        8.5
   b9++      628.51       13.4
   y3+       634.56        8.2
   a4+       673.18        6.5
   y9++      677.07       35.3
   b4+       701.30        9.7
   y10++     727.62       19.3
   y4+       763.11        6.3
   b11++     775.91       13.9
   y11++     791.61       13.3
   b12++     824.49        5.7
   b5+       830.36       13.0
   y5+       860.53       16.9
   b13++     889.27        6.3
   b6+       958.51       13.4
   y14++     977.55        6.6
   b15++    1018.65        8.6
   b7+      1059.50       25.7
   b8+      1156.40       11.0
   b9+      1255.59        9.1
   b11+     1551.74        9.8
```

```
942. MORG-339_phosphopeptide_TMT_30%ACN_032017_OT.17902.17902.4 (sAtPPPAEPASLPQEPPkAFSWASVTSk)

   Ion         m/z   Intensity
   y1+       376.40       15.8
   y2+       463.39       19.5
   y3+       564.51       26.9
   b10++     614.87       12.4
   y13+++    622.07       13.3
   y4+       663.46       13.5
   y15+++    707.88       14.1
   a5+       735.60       17.8
   y16+++    740.44       43.8
   y5+       750.50       16.7
   b18+++    778.30       14.5
   y18+++    806.81       13.6
   a14++     813.11       12.9
   a6+       832.36       16.5
   a20+++    842.17       12.6
   y20+++    862.65       20.0
   b21+++    880.22       21.7
   y12++     883.86       14.9
   b15++     891.76       24.6
   a7+       903.15       16.5
   b7+       931.48       13.7
   a22+++    932.87       25.3
   b22+++    942.48       14.0
   b23+++    966.17       31.7
   b24+++    995.24       14.5
   y25+++   1026.82       22.7
   b8+      1060.51       32.4
   b26+++   1061.65       25.7
   y16++    1109.90       55.9
   y27+++   1110.82       40.5
   b19++    1202.90       11.7
   b11+     1315.85       14.7
   b12+     1428.88       25.3
   b15+     1782.94       13.3
```

```
943. MORG-339_phosphopeptide_TMT_10%ACN_032017_OT.5448.5448.3 (gAGAGASSVGTGGGAAGGAcGGAGsEDsGDR)

   Ion         m/z   Intensity
   b6+       614.44        8.2
   b15++     644.58       11.8
   b7+       701.55       12.0
   y14++     728.28       14.4
   b18++     737.10        8.6
   y15++     756.79       11.4
   a19++     758.42        9.5
   b19++     772.75       11.7
   b8+       788.25       14.9
   y16++     792.39       15.0
   y18++     856.48        8.4
   b21++     880.65       12.3
   y19++     884.67       16.3
   b9+       887.45       24.3
   a22++     895.43        8.6
   b22++     909.48       19.3
   y20++     912.86       18.8
   y7+       925.16       75.1
   a23++     930.73        8.7
   b23++     944.59       30.5
   y8+       982.32       13.6
   y22++     992.54       12.5
   a25++    1043.32        8.9
   b11+     1045.59        9.4
   b12+     1102.81       11.0
   y10+     1110.30       12.2
   a13+     1131.33        8.6
   a27++    1164.64       11.0
   y11+     1167.31       16.6
   b27++    1179.42        8.2
   b15+     1287.63        9.8
   b16+     1358.74       13.2
   y13+     1398.47        8.2
   b17+     1415.66       10.1
   y14+     1455.54        8.7
   y15+     1512.57        8.6
```

```
944. MORG-339_phosphopeptide_TMT_4%ACN_032017_OT.4260.4260.3 (rFsHDGEEEEEDEDYGTR)

   Ion         m/z   Intensity
   y7++      428.00       13.8
   b5++      476.76        8.1
   y4+       496.35        8.2
   b7++      569.85        8.1
   y5+       611.36        4.1
   b8++      634.40        7.8
   b9++      699.06        7.8
   b3+       699.88        8.9
   y6+       740.49        8.1
   y12++     750.38        5.1
   b10++     763.50       11.6
   y13++     779.49        7.2
   a11++     813.99       21.0
   b11++     827.57       15.9
   y7+       855.27       10.4
   a12++     871.46        4.1
   b12++     885.76        7.8
   y15++     905.61        9.0
   y8+       984.43        5.3
   b14++    1007.60       16.6
   a17++    1153.53        9.0
   b17++    1168.34        7.7
   y10+     1242.53        5.1
   y13+     1557.66       11.3
   b11+     1654.51        5.7
```

```
945. MORG-339_phosphopeptide_TMT_4%ACN_032017_OT.8057.8057.3 (eALSQQLDsEDEQDQR)

   Ion         m/z   Intensity
   y2+       303.24       13.0
   b5++      379.73        9.0
   y3+       418.19       19.2
   a6++      429.25        6.7
   b2+       430.28        8.1
   b6++      443.79       16.6
   b7++      500.38       21.7
   a8++      543.41       27.7
   y4+       546.34       22.2
   b8++      557.85       10.4
   y9++      601.40       11.9
   b4+       630.42       16.8
   y10++     657.86       22.6
   y5+       675.32       16.8
   b10++     705.41       14.3
   y11++     721.64       21.9
   b5+       758.56       23.2
   b11++     763.40       14.3
   y12++     785.89       11.1
   y6+       790.35       23.2
   b12++     827.91       18.5
   y13++     829.14       11.2
   y14++     885.60        8.0
   b6+       886.53       22.3
   b13++     892.23       14.9
   y7+       919.36        7.2
   b14++     949.51       13.8
   b7+       999.65       15.2
   b15++    1013.54       17.2
   a8+      1086.47       14.0
   b8+      1114.65       16.6
   y9+      1201.41       19.8
   y10+     1314.54        6.8
   b12+     1654.86        6.9
```

```
946. MORG-339_phosphopeptide_TMT_18%ACN_032017_OT.14920.14920.4 (nGDEENPLkRGLVAAYSGDsDNEEELVER)

   Ion         m/z   Intensity
   y4+       516.35       13.2
   b13+++    628.07       14.6
   y5+       645.28       12.8
   b14+++    651.49       12.8
   b15+++    675.45       16.7
   b16+++    729.69       18.5
   b17+++    758.76       20.0
   y6+       774.47       13.3
   b19+++    816.08       36.1
   b20+++    871.16       12.2
   a12++     877.53       31.6
   y21+++    882.79       14.8
   b6+       887.97       12.2
   b12++     891.80       14.1
   y15++     897.30       17.2
   a21+++    900.16       15.2
   y7+       903.49       26.9
   b21+++    910.05       32.0
   y22+++    920.52       30.4
   y23+++    952.74       11.4
   b7+       985.07        9.6
   y24+++    990.97       18.6
   b15++    1011.87       14.8
   y8+      1017.55       18.5
   y18++    1038.26       10.8
   y26+++   1077.14       23.3
   a26+++   1105.27        9.3
   b26+++   1114.83       11.6
   y27+++   1115.57       16.2
   y9+      1132.59       11.4
   b17++    1137.00       11.6
   a27+++   1137.76        9.4
   b27+++   1147.91       15.8
   b28+++   1190.97       14.7
   y10+     1299.57       11.4
```

```
947. MORG-339_phosphopeptide_TMT_14%ACN_032017_OT.2635.2635.3 (sSNPPARSQsSDTEQPSPTSGGGk)

   Ion         m/z   Intensity
   a5++      342.97        3.9
   y7++      416.65        3.6
   y9++      509.17        4.8
   b3+       518.21        5.9
   b9++      577.45        4.1
   y5+       634.33        4.3
   b10++     661.31        4.1
   b12++     762.28        5.6
   y14++     788.87        3.9
   y7+       832.68        6.6
   a14++     863.42        5.3
   b14++     877.60        6.1
   a7+       911.71        7.5
   y8+       919.12        8.6
   y16++     936.72       12.1
   b7+       939.55       11.0
   b15++     941.63       16.3
   b16++     990.36        4.6
   a8+       998.56        5.9
   y9+      1016.58       11.9
   a17++    1019.57        5.4
   y20++    1141.74        3.6
   y10+     1144.53        4.3
   b9+      1154.68        5.0
   y21++    1190.89        8.4
   b21++    1204.50        4.1
   y23++    1291.15        3.7
   a10+     1293.51        4.1
   y12+     1374.85        5.4
   y15+     1743.87        3.9
   b15+     1881.92        4.1
```

```
948. MORG-339_phosphopeptide_TMT_8%ACN_032017_OT.7023.7023.3 (tVRPEVNAAAssDEVcDGDR)

   Ion         m/z   Intensity
   y3+       347.13        6.8
   b5++      406.78        8.5
   b7++      513.34       10.1
   a9++      570.74       10.4
   b9++      584.43        8.7
   b10++     619.98       15.4
   y5+       622.06       14.2
   a11++     688.94        8.7
   y6+       721.56       11.0
   y13++     757.02       10.1
   b12++     787.23       11.2
   b5+       812.14       13.0
   y14++     813.93       84.0
   a13++     830.69       15.4
   b14++     909.37       11.1
   b15++     958.86       14.4
   y8+       965.12       10.6
   y17++     976.82        8.0
   a7+       998.10       12.9
   b8+      1096.39       13.2
   a9+      1139.49        9.6
   b19++    1182.21        8.4
```

```
949. MORG-339_phosphopeptide_TMT_10%ACN_032017_OT.14218.14218.3 (tQsPGGcsVEAVLAR)

   Ion         m/z   Intensity
   y2+       246.20       11.0
   y3+       359.29       26.1
   y4+       458.36       15.4
   b2+       459.25       10.4
   a7++      484.83        6.6
   b7++      499.32       14.8
   y5+       529.44       21.2
   y9++      542.89       15.4
   y10++     571.42       17.2
   b8++      582.32        8.5
   y11++     599.28       10.4
   b9++      631.75       25.1
   y12++     647.94       19.3
   y6+       658.42       25.7
   b10++     696.87       18.9
   b4+       723.47       13.9
   b11++     732.20       24.3
   y7+       757.73        9.9
   a12++     767.38        5.7
   b5+       780.36       10.1
   b12++     781.96       41.8
   a13++     824.76        8.3
   b6+       837.31       15.8
   b13++     838.39       36.5
   b14++     874.04       20.3
   y8+       924.51       12.0
   b7+       997.41       10.8
```

```
950. MORG-339_phosphopeptide_TMT_20%ACN_032017_OT.14200.14200.3 (rVsFNEAALFEQSR)

   Ion         m/z   Intensity
   a1+       358.33        5.7
   y3+       390.27        6.4
   a2+       457.25        8.8
   y4+       519.26       12.9
   b6++      521.90       18.2
   a7++      543.72       14.0
   b7++      557.23       21.7
   b8++      593.07       22.4
   a3+       624.52       19.7
   a9++      635.44       14.4
   y5+       666.40       13.1
   b10++     722.95       12.6
   y6+       779.52       14.5
   b11++     787.27       10.9
   y7+       850.54       20.1
   b12++     851.53       14.5
   b13++     895.43        9.4
   y8+       921.50       17.2
   y9+      1050.48        9.5
   y11+     1311.78        6.8
```

```
951. MORG-339_phosphopeptide_TMT_6%ACN_032017_OT.5839.5839.3 (eSTESSNTtIEDEDVkAR)

   Ion         m/z   Intensity
   y4++      351.79        6.7
   y6++      473.76        8.7
   y7++      531.51        6.7
   y8++      595.79        5.7
   y9++      652.67       12.4
   a11++     731.09        5.6
   y10++     742.65        5.5
   b5+       763.39        5.3
   a12++     788.50        6.6
   b12++     802.06       10.1
   y5+       817.98       48.5
   a6+       821.93       11.4
   y14++     937.72        6.0
   y6+       946.35        4.7
   a15++     960.29        4.5
   b15++     973.55        6.1
   y16++    1052.39        5.6
   b16++    1152.99        5.1
   b17++    1187.69        6.3
   b12+     1603.64        4.7
   b14+     1847.59        4.9
```

```
952. MORG-339_phosphopeptide_TMT_25%ACN_032017_OT.16807.16807.4 (aTRLGPGLPQDGsDEEDEEWPTLEkAAk)

   Ion         m/z   Intensity
   y4++      438.47       17.5
   a8++      484.48       15.8
   b8++      498.65       21.0
   y8++      658.60       38.8
   y13+++    668.63       67.8
   y15+++    749.81       16.9
   y9++      751.36       26.3
   a6+       797.74       15.8
   a20+++    821.69       16.7
   b14++     838.17       24.5
   y4+       875.48       18.0
   y11++     880.83       34.3
   y19+++    905.68       33.8
   a23+++    926.00       32.0
   b23+++    935.07       52.1
   y20+++    937.92       51.2
   y21+++    975.68       24.6
   b8+       995.88       25.2
   y13++    1002.34       17.3
   y5+      1004.80       20.3
   b17++    1024.66       18.8
   y23+++   1027.26       20.6
   y24+++   1046.56       20.9
   y14++    1067.23       46.3
   b18++    1089.40       17.7
   y15++    1124.78       30.4
   b27+++   1145.00       17.7
   b19++    1153.13       19.5
   y17++    1236.49       24.1
   y8+      1315.95       17.2
   y19++    1358.09       20.2
   y20++    1406.97       20.8
```

```
953. MORG-339_phosphopeptide_TMT_20%ACN_032017_OT.22620.22620.3 (sEPDDELLFDFLNSSQkEPTGR)

   Ion         m/z   Intensity
   y4+       430.32       10.9
   y10++     666.79        7.7
   b4+       738.40       10.0
   y12++     797.08       28.0
   b5+       853.33       13.5
   y13++     854.49       15.5
   a13++     908.68        9.5
   y14++     927.95       13.0
   b6+       982.05       12.6
   y15++     985.02       15.5
   a15++     996.29       10.6
   b15++    1009.77       13.4
   y16++    1041.22       11.8
   y7+      1044.87        7.8
   a7+      1067.51        8.3
   y17++    1105.88       28.6
   y8+      1131.69        7.3
   y18++    1163.29       21.9
   b8+      1208.86       11.0
   y9+      1218.64        7.9
   y19++    1220.73        8.5
   y20++    1269.51       33.7
   y10+     1332.84       10.6
   y21++    1334.16       17.9
   b10+     1470.64       10.2
   y12+     1592.97        8.9
   y13+     1707.85        6.4
```

```
954. MORG-339_phosphopeptide_TMT_10%ACN_032017_OT.14889.14889.3 (rDsEsEsDETPTVPQLk)

   Ion         m/z   Intensity
   a1+       357.81       11.0
   y1+       376.20        8.8
   y2+       489.43       12.1
   y7++      506.53       19.9
   y8++      557.01       13.9
   y4+       714.52       45.5
   b9++      752.66        8.7
   y11++     762.72        9.3
   b4+       797.46       13.1
   b10++     803.15       13.7
   y5+       813.38       15.8
   a11++     837.41       15.1
   b11++     851.68       17.9
   b12++     902.22       10.7
   a13++     938.04       13.3
   b13++     952.09       22.0
   b5+       963.88        9.9
   b14++    1000.70       10.9
   y7+      1011.71       19.4
   b6+      1093.53       12.3
   y8+      1112.44        9.9
   b16++    1121.05       10.0
   b9+      1504.59        9.4
   b10+     1605.51       13.1
   b13+     1902.63        7.7
```

```
955. MORG-339_phosphopeptide_TMT_16%ACN_032017_OT.15926.15926.3 (sSScGDAELLGQATLPVGsPsRPMSR)

   Ion         m/z   Intensity
   y4+       490.32       21.0
   y9++      567.69        8.3
   b10++     625.75       10.4
   a5+       679.94       15.1
   b17++     958.27       19.0
   a18++     972.79       18.8
   a8+       995.67       11.2
   a20++    1104.52        9.5
   y20++    1114.14       13.0
   y21++    1171.49        8.0
   b11+     1307.01        8.5
   b24++    1394.89       11.1
```

```
956. MORG-339_phosphopeptide_TMT_14%ACN_032017_OT.11283.11283.3 (kVSDVTSSGVQsFDEEEGDANN)

   Ion         m/z   Intensity
   y15++     839.33       14.8
   a14++     860.29       12.9
   b14++     873.72       10.5
   y8+       877.08       11.1
   y16++     882.73       13.5
   a7+       918.79       31.4
   y17++     933.45       37.6
   b15++     938.83       14.6
   y18++     982.92        9.7
   a16++     988.90       16.8
   b16++    1002.46       10.9
   a17++    1052.99       15.3
   a11+     1289.80        9.5
```

```
957. MORG-339_phosphopeptide_TMT_22%ACN_032017_OT.16398.16398.4 (qGEVskAAsADSTTEGTPTDGFTVLSTk)

   Ion         m/z   Intensity
   y1+       376.27       17.5
   b2+       415.16       10.0
   y2+       477.39       15.0
   y3+       564.40       20.5
   a14+++    641.57       11.9
   y4+       677.59       19.6
   y13++     776.57       13.7
   b19+++    813.42       10.9
   b11++     832.25       11.7
   y14++     841.09       16.5
   a20+++    841.87       10.4
   a12++     861.48       11.9
   b21+++    869.97       20.9
   b12++     875.65       24.8
   y6+       877.45       13.6
   y15++     891.59       39.7
   b22+++    919.66       12.9
   y23+++    941.57       17.7
   b23+++    953.03       14.0
   a24+++    976.80       11.0
   b24+++    986.39       23.0
   b25+++   1024.08       45.3
   y7+      1024.98       12.6
   b15++    1040.79       21.1
   b26+++   1052.98       23.8
   y8+      1081.65       12.9
   a17++    1106.22       11.2
   b17++    1120.29       21.2
   b22++    1378.71       18.0
```

```
958. MORG-339_phosphopeptide_TMT_4%ACN_032017_OT.9112.9112.3 (mGLEESsEGEAEGSVR)

   Ion         m/z   Intensity
   y3+       361.30        5.3
   a5++      381.21        4.0
   y4+       418.34       12.2
   b6++      438.69        5.0
   y9++      467.30        6.8
   a3+       503.32        3.8
   y5+       547.34       11.1
   y11++     594.70        5.6
   a9++      601.50        6.0
   b9++      615.41       11.8
   y6+       618.46        9.7
   b10++     679.95       10.4
   b11++     715.60        9.8
   y13++     723.26        8.1
   y14++     779.93       15.5
   b5+       789.49        9.0
   y8+       804.38       10.3
   y15++     808.32        5.0
   b14++     851.77        3.6
   b6+       876.55        7.6
   b15++     901.33        5.0
   b8+      1172.43        7.1
   b10+     1358.53        6.9
```

```
959. MORG-339_phosphopeptide_TMT_4%ACN_032017_OT.2992.2992.3 (vEEHsPLER)

   Ion         m/z   Intensity
   y1+       175.07        7.3
   y2+       304.27        6.9
   b1+       329.07        5.6
   y5++      340.90        4.2
   a4++      349.10        9.3
   b4++      362.83       18.8
   y6++      409.67       10.7
   y3+       417.26       11.0
   a2+       430.36       20.1
   a5++      432.29        7.2
   b5++      446.30       31.9
   b2+       458.31       15.5
   y4+       514.34       30.9
   a7++      537.44       10.4
   b7++      551.33        8.9
   a3+       558.83        4.0
   b3+       587.37       19.0
   a8++      601.30        3.9
   b8++      615.85       11.8
   y5+       681.26       12.2
   b4+       724.47        5.5
```

```
960. MORG-339_phosphopeptide_TMT_8%ACN_032017_OT.18205.18205.2 (sFsIsPVR)

   Ion         m/z   Intensity
   y3+       371.26        7.5
   a2+       516.20        4.4
   y4+       538.21       10.2
   b2+       544.39        3.6
   y5+       651.64        7.5
   a4+       796.44        3.8
   b4+       824.40        3.9
   y7+       965.38        4.5
   b5+       991.28        4.6
   b7+      1187.39        4.2
```

```
961. MORG-339_phosphopeptide_TMT_14%ACN_032017_OT.13944.13944.3 (rGsGDTSSLIDPDTSLSELR)

   Ion         m/z   Intensity
   y2+       288.27        8.7
   b5++      391.77       12.8
   a2+       415.23       11.4
   y4+       504.40        9.7
   y9++      509.54        8.4
   b9++      585.95       15.4
   y5+       617.44        9.6
   a10++     628.60       14.0
   b10++     642.50       38.2
   y12++     679.71        9.0
   a11++     685.73        8.1
   b11++     700.04       58.4
   y6+       704.48        8.6
   b12++     748.46       11.9
   a5+       754.75        8.6
   y14++     767.25       21.1
   b5+       782.14       19.1
   a13++     791.74       12.5
   b6+       883.45        9.8
   y17++     903.00       10.0
   y8+       920.56       12.9
   b17++     999.62        9.9
   y9+      1017.47       49.2
   b18++    1064.60       13.4
   b19++    1121.21        8.3
   y10+     1132.51       32.7
   y11+     1245.66       17.4
   b11+     1398.60       12.5
```

```
962. MORG-339_phosphopeptide_TMT_18%ACN_032017_OT.15510.15510.2 (aLPLPQSStc)

   Ion         m/z   Intensity
   b1+       301.32        6.6
   y2+       360.14        8.5
   b2+       414.37       22.7
   y3+       447.19        8.8
   y4+       534.13       16.2
   b4+       624.48       49.8
   y5+       661.90       26.8
   y6+       759.24       51.0
   b6+       849.57       23.5
   b7+       936.58       10.6
   y8+       969.45       27.1
   b8+      1023.54       18.2
   y9+      1082.47        8.1
   b9+      1204.58        7.6
```

```
963. MORG-339_phosphopeptide_TMT_10%ACN_032017_OT.7947.7947.3 (sAtRVIGGPVtPR)

   Ion         m/z   Intensity
   y2+       272.24       18.2
   b1+       317.24        8.0
   y5++      325.38        6.1
   a5++      398.57        6.3
   b5++      412.85        8.7
   y8++      438.43        6.9
   y3+       453.13        6.0
   a6++      455.39        6.0
   b6++      469.07       14.4
   a7++      483.29        7.8
   b7++      498.09       10.9
   a3+       540.87       21.9
   y4+       552.30       19.2
   a9++      560.64       12.9
   b10++     624.38        9.6
   y5+       649.28        8.4
   y11++     656.83       16.9
   y6+       706.34        6.1
   b11++     714.94        9.0
   b12++     763.53        8.2
   y8+       876.47        7.7
```

```
964. MORG-339_phosphopeptide_TMT_12%ACN_032017_OT.20898.20898.3 (iAPFAsDIRtIsEDVER)

   Ion         m/z   Intensity
   y3+       403.28        7.5
   y6++      407.16        3.5
   a3+       483.00        4.0
   b7++      506.45        4.2
   b9++      640.54        9.5
   y5+       647.32        6.7
   y10++     689.75       14.9
   a10++     717.60        6.7
   b10++     731.65       39.0
   a11++     773.62        7.8
   y6+       814.37        6.2
   y12++     830.33        4.5
   y13++     865.91        6.7
   b6+       896.19        5.6
   y7+       927.52        7.1
   b13++     935.90        7.3
   b14++     993.65        6.8
   b7+      1011.60        4.4
   y8+      1108.36        4.8
```

```
965. MORG-339_phosphopeptide_TMT_4%ACN_032017_OT.9313.9313.3 (iEDVGsDEEDDSGk)

   Ion         m/z   Intensity
   y1+       376.22       12.3
   y2+       433.34       24.1
   y6++      440.30        7.5
   b2+       472.32        5.4
   y3+       520.37       15.2
   y8++      562.44        6.9
   a8++      563.25       10.9
   b8++      577.69        7.7
   b3+       587.47        6.5
   a9++      628.56        4.8
   y4+       635.41       12.5
   b9++      642.36       14.7
   y9++      645.67       25.3
   a4+       658.60        5.4
   b4+       686.47        9.7
   b10++     699.79        8.7
   b5+       743.83        6.4
   y5+       750.41        9.8
   b11++     757.28        9.9
   y12++     781.33        5.3
   a12++     786.90        6.8
   b12++     800.93       36.8
   b13++     829.33        6.8
   b7+      1025.42        5.9
   b8+      1154.44       11.8
   b9+      1283.64        9.7
   b10+     1398.63        7.9
```

```
966. MORG-339_phosphopeptide_TMT_20%ACN_032017_OT.4071.4071.4 (kRsEGLSLER)

   Ion         m/z   Intensity
   y1+       175.14       12.4
   a2++      244.06        6.5
   y2+       304.15       30.6
   y5++      308.75        7.5
   y6++      338.06        8.1
   b3++      341.05       14.7
   a7+++     347.26       29.6
   a8+++     384.59        9.9
   b8+++     394.67       59.9
   y7++      402.59        8.3
   b4++      405.59       24.7
   y3+       417.31       26.3
   a5++      420.13       30.0
   b9+++     437.54        6.2
   a2+       486.27       15.7
   y4+       504.34       46.4
   b2+       513.87       37.7
   a7++      519.89       23.5
   y9++      564.13        6.7
   y5+       617.50       22.0
   y6+       674.47       20.7
   y7+       803.50       17.5
```

```
967. MORG-339_phosphopeptide_TMT_16%ACN_032017_OT.19093.19093.3 (aASADstTEGTPTDGFTVLSTk)

   Ion         m/z   Intensity
   y1+       376.35       12.0
   y2+       477.48       11.9
   y3+       564.43       19.2
   y4+       677.46       20.0
   y11++     698.14       21.4
   y12++     748.55        8.9
   y13++     776.73       10.6
   b13++     790.45        8.8
   y14++     841.57        9.5
   b15++     876.26        8.4
   y6+       877.49       13.9
   y15++     891.62        8.9
   b16++     950.22       11.3
   y16++     982.24        6.8
   b7+       993.15        9.2
   b17++    1000.68        9.0
   b18++    1050.09       15.3
   y8+      1081.18        7.3
   b19++    1106.69       28.7
   b20++    1150.23       13.3
   b21++    1200.66       11.2
   b9+      1223.47       13.3
   b10+     1280.32        8.7
   b11+     1381.45       11.0
   b14+     1694.73        6.6
```

```
968. MORG-339_phosphopeptide_TMT_22%ACN_032017_OT.21040.21040.3 (lILTDNGSAPTFcsDDEDVAPLSAk)

   Ion         m/z   Intensity
   y1+       376.43        8.6
   b2+       456.36        8.0
   y3+       534.39       15.1
   b3+       569.49        8.1
   y4+       647.33       12.8
   y5+       744.57       39.0
   y6+       815.58       33.7
   y13++     858.74       11.7
   a14++     879.62        7.8
   b6+       899.60        9.5
   y7+       914.73       11.2
   a15++     937.90        8.4
   y15++     982.37        8.4
   y8+      1029.79       15.0
   y16++    1031.25       37.9
   b8+      1043.46       17.7
   b17++    1073.68       11.8
   y18++    1109.89       12.0
   b9+      1114.63       37.5
   a18++    1116.62       11.7
   b18++    1131.37       17.1
   y19++    1138.60       14.2
   a19++    1166.88       11.0
   b19++    1180.64       28.0
   y20++    1195.85        7.7
   a20++    1201.91       10.1
   b20++    1216.43       34.0
   b21++    1264.76        8.8
   b11+     1312.80       15.6
   b22++    1320.91       11.7
   y23++    1360.58        7.8
   b23++    1364.87        9.0
   y11+     1389.10        7.9
   y24++    1417.05        7.6
   b12+     1459.92       14.7
```

```
969. MORG-339_phosphopeptide_TMT_25%ACN_032017_OT.7925.7925.3 (rkNLAkPGVTSTsDsEDEDDQEGEk)

   Ion         m/z   Intensity
   y3+       562.38       16.1
   a6++      685.98       12.5
   b6++      699.93       33.0
   y5+       819.55       27.7
   b9++      826.53       16.2
   b10++     877.39       17.4
   y6+       934.57       32.5
   b12++     971.02       17.5
   a13++    1041.03       12.9
   b5+      1041.91       20.0
   y7+      1049.58       15.0
   b13++    1054.52       15.0
   a14++    1098.02       12.2
   b14++    1112.15       36.7
   y8+      1178.58       29.2
   a15++    1181.66       12.3
   b15++    1195.27       21.6
   y9+      1293.54       13.9
   b17++    1317.87       28.0
   b18++    1381.95       16.3
   y20++    1386.43       12.0
   b6+      1399.02       23.2
   y10+     1422.53       12.2
   b19++    1440.07       34.1
   b20++    1497.48       35.0
   y11+     1589.66       32.3
   b22++    1626.15       15.6
```

```
970. MORG-339_phosphopeptide_TMT_20%ACN_032017_OT.17983.17983.3 (aSGQAFELILsPR)

   Ion         m/z   Intensity
   y2+       272.15       17.4
   a1+       273.09        8.5
   b5++      322.80        9.3
   y6++      390.09        9.2
   b6++      396.18       12.7
   y3+       439.21       34.4
   b3+       445.29        6.9
   a7++      446.41        5.0
   y7++      454.36        9.4
   b7++      460.99       31.1
   a8++      503.21        5.1
   b8++      517.58       36.6
   y8++      528.00       20.8
   a4+       545.22        9.7
   y4+       552.30       27.6
   b9++      574.06       37.7
   y10++     627.16        5.8
   b10++     630.66       58.2
   b5+       644.41       14.8
   y5+       665.19       44.9
   y12++     699.46        6.0
   a11++     700.31        7.5
   b11++     714.25       16.7
   y6+       778.48       16.0
   b6+       791.62        8.3
   y7+       907.56        9.6
   b7+       920.50       19.0
   b8+      1033.54       15.2
   b9+      1146.63        6.4
```

```
971. MORG-339_phosphopeptide_TMT_8%ACN_032017_OT.9799.9799.3 (sDISERPsVDDVESETGSTGALETR)

   Ion         m/z   Intensity
   b6++      459.29       12.9
   b10++     698.54       12.7
   b11++     755.49        8.1
   y8+       834.47        9.3
   b13++     870.12       15.6
   y9+       891.65        8.2
   b14++     913.47       13.3
   b6+       917.83       18.1
   a15++     964.24       34.2
   y18++     966.42       16.4
   a17++    1042.69       10.8
   b17++    1057.11       18.3
   a18++    1086.58        9.6
   y11+     1121.48       12.0
   y12+     1208.78        8.5
   y23++    1258.44       10.9
   b22++    1271.77       10.5
   y15+     1551.72        8.8
```

```
972. MORG-339_phosphopeptide_TMT_16%ACN_032017_OT.9402.9402.4 (tDHGAEIVYksPVVsGDTSPR)

   Ion         m/z   Intensity
   y2+       272.25       12.3
   b4++      320.66        9.2
   b5++      356.31        9.6
   y11+++    420.74       20.8
   a7++      463.38       19.1
   b7++      477.49       28.1
   a8++      512.97        9.1
   b10+++    525.15       17.8
   b8++      527.16       19.3
   y12+++    540.20       13.0
   y10++     548.07       32.7
   a11+++    571.33        9.2
   y5+       575.34       17.5
   b11+++    580.62       15.2
   b3+       583.26       13.3
   y13+++    594.56       15.2
   b9++      608.30       13.4
   b12+++    613.24       11.5
   a13+++    636.75       10.7
   b4+       640.15        8.7
   b13+++    646.05       23.5
   y15+++    664.96       13.0
   a14+++    669.78       20.6
   b14+++    679.20       31.1
   a5+       683.51       13.1
   a15+++    725.50       12.4
   y18+++    750.94       11.0
   b10++     786.96       10.2
   y7+       799.27       19.6
   y12++     810.06        9.4
   a6+       812.00        8.3
   a18+++    816.06       12.8
   y20+++    835.17        8.9
   b6+       840.49        9.5
   y13++     891.37       18.9
   y8+       898.36       19.0
   y14++     941.16       18.6
   b7+       953.58       12.3
   b13++     968.60       10.0
   y15++     997.73       12.0
   b14++    1018.39        8.9
   b8+      1052.61       11.7
   y16++    1062.00       10.0
   y10+     1094.45       13.1
   y11+     1261.43       10.7
```

```
973. MORG-339_phosphopeptide_TMT_8%ACN_032017_OT.13643.13643.2 (eLsLDDPEVEQVR)

   Ion         m/z   Intensity
   y6+       759.69        6.1
   a5+       839.32        7.0
   y7+       856.47       34.5
   b5+       867.21       15.5
   y8+       971.53       17.3
   b6+       982.54       22.5
   y9+      1086.83        8.5
   y10+     1199.63       11.0
   b9+      1307.63        7.5
```

```
974. MORG-339_phosphopeptide_TMT_6%ACN_032017_OT.9877.9877.2 (nSTLsEEDYIER)

   Ion         m/z   Intensity
   y4+       580.36       13.6
   b4+       645.39        7.5
   y5+       695.32        6.0
   a5+       784.34        7.1
   b5+       812.11        5.6
   b6+       941.42        6.0
   y7+       953.43        6.2
   b7+      1070.58        5.4
   y8+      1120.19        9.4
   b8+      1185.57       12.9
   y10+     1334.36        7.0
   y11+     1421.62        6.5
   b11+     1590.81        5.8
```

```
975. MORG-339_phosphopeptide_TMT_4%ACN_032017_OT.4544.4544.3 (gRVsPSDADSTVSEESSER)

   Ion         m/z   Intensity
   y4+       478.30        6.4
   b7++      504.99        9.4
   a3+       514.62        9.6
   a9++      583.34        5.8
   b9++      597.93        9.2
   y5+       607.25        7.4
   a11++     677.86        6.5
   b11++     692.02       10.7
   b4+       709.46       16.4
   a12++     727.46        9.6
   y6+       736.13       71.8
   b12++     741.23       10.1
   y15++     798.11       10.5
   y7+       823.32        6.9
   a14++     835.20        9.7
   a6+       865.70        7.8
   y9+      1023.65        8.3
   b18++    1065.54        7.9
   y10+     1110.57        9.0
   y12+     1296.57        9.1
```

```
976. MORG-339_phosphopeptide_TMT_18%ACN_032017_OT.12344.12344.3 (aAsPPASASDLIEQQQk)

   Ion         m/z   Intensity
   b2+       372.05       10.2
   y1+       376.40       25.6
   y4++      380.83        9.5
   y2+       504.38       28.9
   b9++      525.58       10.6
   y7++      558.44        9.4
   b10++     582.60       16.2
   y8++      615.65       13.7
   a11++     625.54        9.6
   y3+       632.56       20.6
   b11++     639.49       34.3
   a12++     681.86       10.5
   y10++     694.55       19.4
   b12++     696.02       42.0
   b5+       733.57       22.5
   y11++     738.06       15.1
   a13++     746.49       23.2
   b6+       804.47       17.2
   b14++     824.86       12.4
   a7+       863.26       13.2
   a15++     874.35        9.8
   b15++     888.80       36.1
   y5+       889.56       32.9
   b7+       891.53       12.6
   a8+       934.88        9.1
   b16++     952.74       37.7
   b8+       962.61       12.4
   y6+      1002.67       10.3
   b10+     1164.56       15.5
   b11+     1277.66       15.5
   b12+     1390.71       21.1
   b13+     1519.80       16.7
   b14+     1647.97        9.1
```

```
977. MORG-339_phosphopeptide_TMT_8%ACN_032017_OT.17987.17987.2 (sVDIsLGGsPR)

   Ion         m/z   Intensity
   y2+       272.28        7.9
   y5+       553.35        8.7
   b3+       611.34        8.5
   y6+       666.38        9.9
   b4+       724.46       13.8
   y7+       833.33        9.7
   b5+       890.91        5.2
   y8+       946.45        8.0
   b7+      1061.31        4.7
   a8+      1090.60        6.1
   y10+     1160.51        6.5
```

```
978. MORG-339_phosphopeptide_TMT_16%ACN_032017_OT.16558.16558.3 (kIVEPEVVGEsDsEVEGDAWR)

   Ion         m/z   Intensity
   y3+       432.39        7.9
   y5+       604.27       11.2
   y11++     705.66       13.0
   y6+       733.34       10.7
   y12++     770.17       17.4
   y7+       832.34        9.4
   a13++     865.48        8.0
   b13++     879.59        7.6
   a14++     930.64       10.9
   b14++     944.50       17.4
   y8+       961.46       14.2
   y16++     962.59       16.5
   y17++    1010.67        9.2
   a16++    1044.91        8.2
   y19++    1125.39       11.2
   y9+      1128.38        7.8
   a9+      1152.90        7.9
   y11+     1410.37        8.9
   y13+     1596.49        7.1
   y14+     1695.60        6.9
```

```
979. MORG-339_phosphopeptide_TMT_8%ACN_032017_OT.10421.10421.3 (dGGEALVsPDGTVTEAPR)

   Ion         m/z   Intensity
   y2+       272.22        9.7
   y3+       343.26       12.8
   y7++      386.94        7.4
   y8++      415.26        7.0
   b7++      436.46        7.7
   y4+       472.15        7.8
   y9++      473.19        5.4
   y10++     521.93       19.3
   a4+       560.25        6.1
   y5+       573.37       16.7
   b4+       587.79        8.4
   y11++     605.53       20.7
   b5+       659.37       12.7
   y6+       672.36       13.7
   y13++     711.24        6.4
   y14++     746.37        7.7
   b13++     754.13        8.8
   b6+       772.36       15.7
   y7+       773.43       12.5
   y15++     811.71        6.6
   y8+       830.62        7.1
   y16++     840.07        5.5
   a7+       843.45        5.9
   y17++     868.41        7.6
   b15++     869.75       12.2
   b7+       871.53       21.4
   b16++     905.21       19.8
   b8+      1038.48        7.4
   y11+     1209.52        7.7
   y12+     1308.71        5.5
```

```
980. MORG-339_phosphopeptide_TMT_12%ACN_032017_OT.1574.1574.4 (kQHREsGEGEEEVADSAR)

   Ion         m/z   Intensity
   y3+       333.35       13.1
   y4+       448.24       15.4
   a11+++    516.60       12.3
   y5+       519.31       26.1
   b6++      538.15        8.3
   a12+++    559.74       23.0
   b12+++    569.63       17.5
   a13+++    592.67       15.9
   y11++     595.86       14.9
   y16+++    613.10       39.6
   a14+++    616.43       23.1
   a8++      617.38       23.1
   y6+       618.56       36.7
   y12++     625.00       23.1
   b14+++    626.01       13.3
   b8++      631.32       17.6
   a9++      646.03       26.5
   b9++      659.66        8.4
   y13++     707.98       17.0
   y7+       747.37       16.8
   a11++     774.61       11.7
   a12++     839.00        9.6
   y8+       876.46       13.8
   y16++     919.14       12.5
   y10+     1062.53       10.8
```

```
981. MORG-339_phosphopeptide_TMT_4%ACN_032017_OT.16110.16110.2 (eEGAsPLDFD)

   Ion         m/z   Intensity
   y2+       281.09       18.2
   b1+       359.41        3.5
   y3+       396.11       10.9
   a2+       460.30        2.8
   b2+       488.34        6.2
   a4+       588.37        7.3
   y5+       606.41       19.9
   b4+       616.32       18.2
   a5+       755.17       11.5
   y6+       773.24       14.0
   b5+       783.36       18.2
   y7+       844.47        5.4
   y8+       901.38        8.2
   a7+       965.45        3.1
   b7+       993.41       23.3
   y9+      1030.47        8.8
   b8+      1108.46       48.4
```

```
982. MORG-339_phosphopeptide_TMT_14%ACN_032017_OT.13198.13198.4 (eTPScPkPVQDtQPPESPVENSEQLLQTNSGAVAR)

   Ion         m/z   Intensity
   y6+       560.27       12.6
   y7+       674.43       14.8
   y8+       775.44       19.4
   a5+       776.44       16.1
   b10++     792.18       11.0
   y23+++    817.44       12.0
   b11++     849.64       14.1
   b6+       901.45       12.0
   y9+       903.50       24.7
   y18++     957.16       11.5
   b22+++    981.58       12.7
   b13++    1003.52       10.8
   y10+     1016.44       20.5
   b23+++   1025.19       21.7
   a14++    1038.71       13.8
   b14++    1052.69       16.1
   b25+++   1105.31       27.4
   y29+++   1143.79       13.7
   y22++    1161.61       19.1
   b27+++   1185.94       13.2
   b28+++   1219.52       30.7
   a7+      1231.18       15.4
   a29+++   1248.25       10.7
   y12+     1257.76       10.8
   b19++    1307.04       16.6
   b32+++   1329.09       21.1
   b33+++   1362.34       26.7
   b20++    1372.02       19.9
   y25++    1374.01       12.2
   y13+     1386.49       24.6
   b21++    1428.96       14.2
   b23++    1537.13       16.1
   b24++    1601.16       13.8
   b25++    1657.47       15.3
   b26++    1714.00       16.5
```

```
983. MORG-339_phosphopeptide_TMT_4%ACN_032017_OT.5480.5480.3 (eQAEMDDADNsEk)

   Ion         m/z   Intensity
   y1+       376.43       15.0
   y5++      451.15       11.0
   y6++      486.83       10.4
   b2+       487.45       11.3
   y2+       505.34        6.8
   y7++      543.99        9.6
   b3+       558.27       13.0
   y8++      601.81        9.2
   a9++      603.81        9.1
   b4+       687.48       10.1
   y4+       786.51        6.5
   b5+       818.43       11.2
   y5+       901.71       10.4
   b6+       933.34        8.1
   b7+      1048.53        7.0
   b8+      1119.51       13.7
```

```
984. MORG-339_phosphopeptide_TMT_8%ACN_032017_OT.19229.19229.3 (eVDDILGEGsDDsDIEk)

   Ion         m/z   Intensity
   y1+       376.21       13.1
   b2+       458.20        8.9
   y2+       505.32       25.4
   b3+       573.34        4.8
   y3+       618.49        6.9
   y8++      649.23        5.5
   y9++      678.11       14.7
   b11++     720.24        5.8
   y4+       733.21        8.0
   y10++     742.27        7.8
   y11++     771.08       21.2
   b5+       801.28       21.6
   b6+       914.52       22.6
   b14++     919.13       12.4
   y14++     941.48        7.6
   a15++     961.66       15.6
   b7+       971.55        7.4
   b15++     975.53       21.6
   b16++    1040.35       15.7
   b8+      1100.72       16.1
   y7+      1130.36        7.4
   b14+     1836.77        8.4
```

```
985. MORG-339_phosphopeptide_TMT_22%ACN_032017_OT.14302.14302.4 (dRAsPAAAEEAVPEWAScLkSPR)

   Ion         m/z   Intensity
   a7+++     317.35        9.9
   y5++      415.17       11.9
   b6++      454.15       16.8
   b7++      489.78       11.9
   a13+++    516.34       10.0
   y11+++    520.95        9.8
   b13+++    525.27       13.8
   b14+++    568.88       11.0
   b15+++    630.65       12.5
   a10++     640.04       12.4
   a16+++    645.28       12.2
   b16+++    654.73       16.8
   y9++      667.54       15.8
   b17+++    683.48       18.1
   a4+       711.20       20.3
   y4+       716.63       11.5
   a18+++    727.83       19.0
   y10++     731.51       16.3
   y18+++    734.62       20.4
   b18+++    736.48       18.9
   b12++     739.30       23.9
   y19+++    766.50       14.5
   b19+++    774.19       12.8
   y11++     780.37       30.5
   y12++     830.21       13.4
   y21+++    846.21       14.4
   a6+       879.30       11.0
   a20+++    883.80       11.2
   y22+++    898.47       10.0
   a21+++    913.08       11.0
   y14++     929.88       14.7
   y6+       989.62       10.1
   y15++     994.71       19.6
   y16++    1030.21       23.0
   y17++    1065.71       13.3
   b18++    1104.34       10.7
   y19++    1149.83       25.2
   a9+      1150.59       16.6
   y11+     1560.04       10.5
```

```
986. MORG-339_phosphopeptide_TMT_4%ACN_032017_OT.14629.14629.4 (mESEAGADDsAEEGDLLDDDDNEDRGDDQLELk)

   Ion         m/z   Intensity
   y1+       376.34       30.8
   y2+       489.39       42.1
   y3+       618.46       47.6
   y10++     709.41       18.6
   b4+       722.35       35.8
   y4+       731.56       29.1
   b12++     760.00       17.8
   b5+       793.42       33.9
   b13++     824.30       16.0
   y12++     831.09       25.2
   y20+++    840.70       19.8
   b6+       850.49       21.2
   b14++     852.50       17.5
   y5+       859.57       18.4
   y21+++    883.66       34.5
   y13++     888.63       26.1
   b15++     910.60       35.2
   b7+       921.45       44.0
   y22+++    926.63       28.2
   a24+++    946.20       33.6
   y23+++    950.42       18.2
   b16++     967.16       45.8
   y6+       974.68       28.8
   y15++    1003.75       47.8
   b25+++   1007.64       20.1
   a17++    1009.29       27.6
   b17++    1023.67       40.0
   y16++    1061.20       78.2
   b18++    1080.70       24.8
   y26+++   1082.85       21.9
   y17++    1117.75       69.3
   y28+++   1125.55       20.4
   b29+++   1145.92       25.1
   y8+      1146.57       22.6
   b9+      1151.27       40.7
   a30+++   1174.30       37.5
   b30+++   1183.65       77.6
   y31+++   1220.82       40.5
   b31+++   1226.81       62.5
   y19++    1231.85       38.7
   y20++    1260.28       59.4
   b32+++   1264.57       49.6
   y21++    1324.92       38.9
   b25++    1510.91       30.6
   b12+     1518.56       38.0
   b27++    1596.92       28.0
   b13+     1647.60       29.7
   b28++    1654.45       24.3
   b29++    1718.51       32.7
   b15+     1819.67       21.3
   b31++    1839.57       19.1
```

```
987. MORG-339_phosphopeptide_TMT_12%ACN_032017_OT.6254.6254.4 (sLDsDESEDEDDDYQQkRk)

   Ion         m/z   Intensity
   b1+       317.23       11.8
   y6+++     437.17       27.1
   y7+++     475.43       22.3
   y8+++     513.74       24.5
   b3+       545.41       15.2
   y9+++     552.46       14.9
   a13+++    578.06       17.7
   y10+++    595.26       23.6
   y11+++    633.71       19.6
   y6++      654.99       14.2
   a15+++    675.32       11.4
   y12+++    676.53       12.0
   y13+++    705.59       30.2
   b10++     708.48       12.6
   y7++      712.38       16.8
   a16+++    718.64       31.2
   y14+++    748.25       54.7
   a11++     752.09       23.4
   y15+++    786.81       22.1
   b12++     824.27       10.7
   y9++      827.59       18.2
   y16+++    842.43       18.3
   a13++     867.64       10.2
   y10++     892.18       19.2
   y11++     949.41       19.4
   b6+       956.58       12.1
   y12++    1013.80        9.8
   b13+     1761.58       12.5
```

```
988. MORG-339_phosphopeptide_TMT_4%ACN_032017_OT.4966.4966.3 (qGEDNsHTQDTEDLEk)

   Ion         m/z   Intensity
   y1+       376.19        9.6
   y2+       505.37       10.7
   a3+       516.44        5.9
   y7++      540.08        8.2
   y3+       618.27       13.2
   a4+       631.62        7.7
   b4+       659.48       11.8
   b10++     711.65       15.2
   y10++     723.27       10.5
   y4+       733.56        7.9
   a11++     747.96       12.2
   b5+       773.09       15.8
   y11++     806.47        8.6
   b12++     826.76        9.0
   y5+       862.51       11.0
   y12++     863.58        9.4
   b13++     883.90       10.1
   b14++     940.68       12.7
   y6+       963.42       13.0
   y14++     985.52        7.8
   b15++    1005.28       15.3
   y7+      1078.77       10.5
   y9+      1307.93        6.6
```

```
989. MORG-339_phosphopeptide_TMT_8%ACN_032017_OT.6782.6782.3 (eYVSNDATQsDDEEk)

   Ion         m/z   Intensity
   y3++      317.26        9.6
   y1+       376.34       10.2
   y2+       505.17       10.9
   b8++      555.19        8.0
   y7++      580.43        8.8
   b3+       621.51       11.3
   y8++      630.82        9.1
   y3+       634.45        8.5
   y9++      666.61        7.7
   a4+       680.38        7.8
   a10++     688.44        8.7
   b10++     702.41        8.1
   b4+       707.94       15.2
   a5+       794.89        6.3
   b12++     818.25        7.7
   b5+       822.56        9.1
   y12++     824.58       13.4
   y5+       864.64        7.1
   y13++     873.61       10.1
   b13++     882.73       13.8
   a6+       909.56        6.8
   b6+       937.63        9.7
   b14++     947.20       12.9
   y14++     955.70        7.8
   b7+      1008.24       11.3
   y6+      1031.46        9.8
   b8+      1109.46        8.3
   b9+      1237.76        5.7
```

```
990. MORG-339_phosphopeptide_TMT_6%ACN_032017_OT.10070.10070.2 (sAEDLLER)

   Ion         m/z   Intensity
   y2+       304.22        6.1
   b1+       397.25        6.0
   y3+       417.31        7.6
   y4+       530.44       13.0
   a3+       568.95        7.8
   b3+       597.35        8.3
   y5+       645.68        7.0
   b4+       712.31       16.0
   a5+       797.23        4.1
   b5+       825.40        7.3
   y7+       845.47        9.7
   b6+       938.44        5.3
   b7+      1067.53        7.8
```

```
991. MORG-339_phosphopeptide_TMT_8%ACN_032017_OT.11750.11750.3 (aVPVSPsAVEEDEDEDGHTVVATAR)

   Ion         m/z   Intensity
   b1+       301.24        8.4
   b2+       400.32       19.7
   y4+       418.35       12.1
   y10++     513.94        9.4
   y5+       517.40       15.4
   y11++     578.50       10.7
   b4+       596.39        9.3
   y6+       616.48       12.1
   y12++     635.98       12.3
   b5+       683.33        8.6
   y13++     700.57       12.9
   y7+       717.52       16.1
   y15++     822.63       13.4
   y8+       854.74        9.4
   y16++     887.22       18.1
   y9+       911.38       16.0
   a7+       919.61       17.8
   b15++     932.70       15.0
   y17++     936.61       23.0
   b7+       947.36       18.8
   y18++     972.34       11.3
   a17++    1004.47        8.3
   y10+     1026.72       10.7
   y19++    1055.27       17.6
   b18++    1087.21       10.8
   a9+      1089.41       13.0
   b9+      1117.62        9.2
   b19++    1137.33        8.8
   y21++    1147.46       10.7
   y11+     1155.55       12.8
   b20++    1187.32       16.9
   y22++    1196.86       22.8
   b21++    1236.31       11.4
   y23++    1245.91       21.8
   b10+     1246.69       14.8
   y12+     1270.55       10.0
   b22++    1271.58        9.2
   y24++    1295.43        8.9
   b24++    1358.28        9.1
   y14+     1514.74       13.4
   y15+     1643.78        9.1
   b16+     1978.75        8.4
```

```
992. MORG-339_phosphopeptide_TMT_20%ACN_032017_OT.9303.9303.4 (rSsAIGIENIQEVQEkR)

   Ion         m/z   Intensity
   y2++      266.76       17.6
   a3++      307.14       28.4
   b3++      320.96       16.0
   b4++      356.28        9.8
   a5++      398.63       48.0
   b9+++     412.80       41.3
   a10+++    441.46       30.0
   a2+       444.94       27.5
   a11+++    483.81       22.2
   b7++      497.78       30.2
   y6++      509.18       12.4
   a12+++    527.19       10.3
   y2+       532.14       13.1
   y12+++    557.56       27.2
   b8++      562.54       25.2
   y7++      573.54       24.6
   y13+++    595.88       55.7
   a9++      604.84       50.5
   b14+++    612.46       17.4
   b10++     675.98       12.4
   y9++      687.18       34.1
   b11++     739.70       13.9
   y10++     751.63       51.4
   b16+++    774.50       16.1
   y4+       789.57       14.0
   a5+       796.23       11.1
   b12++     804.60       26.7
   y11++     808.24       39.8
   y12++     836.70      108.5
   y13++     893.32       41.3
   y14++     928.76       43.5
   b15++     982.91       14.1
```

```
993. MORG-339_phosphopeptide_TMT_20%ACN_032017_OT.13528.13528.4 (aSIGNIALsPsPVRksPR)

   Ion         m/z   Intensity
   y2+       272.26       14.1
   a1+       273.19        8.6
   b5++      336.77        7.6
   b6++      393.21       15.3
   y4++      398.88       11.0
   b7++      428.95       17.7
   y8+++     439.15       14.6
   y9+++     471.68       37.5
   y5++      477.16       11.1
   b12+++    499.91       11.5
   b3+       501.46        8.5
   a13+++    523.60        8.9
   a4+       530.29        7.5
   b13+++    532.59       20.5
   b4+       558.40       11.2
   y11+++    564.58       17.7
   y7++      575.02       13.4
   b14+++    584.86       12.9
   y12+++    589.04       34.7
   a10++     603.13        7.5
   b10++     617.54        9.0
   y13+++    626.07       15.2
   y8++      658.44       18.8
   y14+++    664.51        8.8
   b5+       672.43       24.7
   a15+++    694.62       24.4
   b15+++    704.26       10.7
   y9++      706.81       24.6
   b16+++    759.79       10.0
   a13++     784.88        9.0
   b6+       785.52       17.4
   y10++     790.18       11.7
   b17+++    792.13        8.2
   b13++     798.98       10.5
   y11++     847.15       27.1
   b7+       856.62       16.3
   b14++     877.31       13.3
   y12++     882.66       28.3
   y13++     939.35        6.9
   b8+       969.68        9.9
```

```
994. MORG-339_phosphopeptide_TMT_16%ACN_032017_OT.10653.10653.4 (tGIRTDSREDEIsPPPPNPVVk)

   Ion         m/z   Intensity
   b6+++     318.22       12.1
   y4++      336.27       12.6
   y2+       475.42       18.9
   y7++      490.42       10.8
   y8++      538.89       11.7
   y9++      587.52       25.6
   b13+++    617.20       17.0
   a14+++    640.36        8.8
   b14+++    649.47       16.6
   y4+       671.29       19.8
   a15+++    672.28       15.5
   b15+++    681.89       20.3
   y16+++    690.85       10.2
   a16+++    704.75       10.5
   a10++     706.72        8.1
   a4+       709.13       11.6
   b16+++    714.57       11.8
   y11++     727.72       32.0
   y17+++    729.41       16.6
   b4+       737.39       56.7
   b17+++    746.55       21.3
   a11++     771.71        8.3
   b18+++    784.83       24.8
   y5+       785.74       12.0
   y12++     792.35        9.8
   a19+++    807.72       10.4
   b19+++    817.52       20.8
   a12++     827.61       13.1
   b5+       838.68       10.0
   a20+++    840.93       10.6
   b12++     841.80       14.2
   b20+++    850.17       22.7
   a21+++    874.05        9.2
   b21+++    882.86       15.6
   a6+       925.09       21.2
   b14++     973.92       13.4
   b15++    1022.31        8.9
   b16++    1071.27        8.0
   b18++    1176.65       18.7
```

```
995. MORG-339_phosphopeptide_TMT_22%ACN_032017_OT.15685.15685.3 (nssPSISPNTSFASDGSPSPLGGIkR)

   Ion         m/z   Intensity
   y7++      485.49        8.7
   y9++      577.15       12.5
   y11++     649.37       21.6
   a3+       649.98       11.5
   a10++     674.08        8.5
   y16++     902.89       10.9
   y17++     953.54        9.4
   a16++     955.68        8.5
   a17++     999.51       11.2
   y18++    1010.59       12.6
   b17++    1013.58       10.6
   a7+      1034.76       25.1
   a18++    1047.46       16.1
   y20++    1102.57       14.5
   b20++    1153.52       16.9
   b8+      1159.28        9.5
   a21++    1196.29       16.5
   b9+      1273.32        9.5
   y11+     1297.63       14.5
```

```
996. MORG-339_phosphopeptide_TMT_16%ACN_032017_OT.15000.15000.3 (sRsPVDSPVPASMFAPEPsSPGAAR)

   Ion         m/z   Intensity
   b1+       317.26       15.0
   y5+       471.33       20.5
   b6++      476.45       22.4
   b8++      568.76       10.6
   b9++      618.15       25.7
   b11++     702.09       11.7
   b13++     811.16       14.6
   y8+       822.38       20.5
   y16++     826.86       13.6
   b5+       836.37       13.0
   y17++     876.46       11.7
   b14++     884.65       24.3
   b15++     919.78       38.5
   a6+       923.23       15.0
   y18++     924.48       47.5
   b6+       951.47       30.7
   b17++    1033.17       29.0
   b7+      1038.47        9.0
   y10+     1048.46       36.0
   y11+     1119.53       19.0
   y22++    1123.81       12.2
   a19++    1150.91       26.3
   b9+      1234.59       17.7
   y12+     1266.60       13.2
   y14+     1484.26        9.8
   y15+     1555.84        9.7
   y16+     1652.65       15.3
   b15+     1838.82       10.6
```

```
997. MORG-339_phosphopeptide_TMT_14%ACN_032017_OT.11741.11741.3 (gcGVktPPSsPPEVsEDEDAk)

   Ion         m/z   Intensity
   y2+       447.40        7.2
   a9++      697.39        8.7
   y10++     714.00        8.4
   y11++     763.20        9.4
   y14++     938.08       18.8
   b13++     957.28       10.6
   b5+       960.71        8.5
   y15++     987.14       23.0
   a14++     992.57        7.8
   b14++    1006.14       52.1
   y16++    1077.74        8.9
   y7+      1102.10        7.4
   a6+      1113.27       10.0
   y19++    1334.56        9.3
   y10+     1427.85        9.4
   y11+     1524.72       13.1
```

```
998. MORG-339_phosphopeptide_TMT_10%ACN_032017_OT.1543.1543.4 (sTsATDTHHVELARER)

   Ion         m/z   Intensity
   y5++      322.92       39.5
   y8+++     337.52       13.3
   y9+++     383.04       41.4
   y6++      387.39       14.2
   a2+       390.35       14.6
   y10+++    416.81       89.9
   y7++      436.91       36.0
   y11+++    455.10       25.9
   a7++      473.39       13.8
   b7++      487.46       20.9
   y12+++    488.67       50.6
   b11+++    492.83       30.7
   y8++      505.30       80.9
   y13+++    512.48       40.0
   a12+++    520.85       18.6
   a8++      541.58       21.3
   b8++      555.95       19.3
   y14+++    568.21       17.1
   y9++      574.00       49.1
   b3+       585.17       20.3
   y15+++    601.16       13.3
   b14+++    606.06       16.7
   a9++      610.68       21.5
   y10++     624.54       45.2
   a4+       628.40       12.2
   a15+++    639.65       18.6
   y5+       644.32       17.3
   b15+++    649.24       29.9
   b4+       656.38       22.4
   a10++     659.35       15.8
   b10++     674.22       16.3
   y11++     682.11       51.0
   a11++     724.42       12.9
   a5+       729.57       19.4
   y12++     732.56       49.4
   b11++     738.23       31.3
   b5+       757.41       14.9
   y13++     768.14       32.4
   y7+       872.40       50.0
   b7+       973.41       12.2
```

```
999. MORG-339_phosphopeptide_TMT_10%ACN_032017_OT.6607.6607.3 (rsPsPAPPPPPPPPPPR)

   Ion         m/z   Intensity
   y6++      330.72       13.6
   y7++      379.36       19.5
   y8++      427.74       13.3
   b5++      457.50       11.7
   y9++      476.41       46.0
   a6++      479.24       17.0
   b6++      493.21       28.4
   a2+       525.04       24.7
   b7++      541.75       16.2
   y5+       563.37       28.1
   y11++     573.46       89.0
   a8++      576.09       12.0
   b8++      590.15       13.4
   a3+       622.39       13.6
   a9++      624.86       11.6
   b9++      638.64       18.4
   b3+       650.65       13.0
   y13++     657.44       21.8
   y6+       660.05       20.0
   b10++     687.12       26.8
   b11++     735.67       40.4
   y7+       757.56       15.5
   y15++     789.33       24.6
   b4+       817.27       14.8
   a13++     818.63       25.9
   y8+       854.55       14.3
   a14++     867.47       27.4
   y9+       951.61       23.7
   b6+       985.38       70.7
   y10+     1048.66       18.9
   b7+      1082.39       18.4
   y11+     1145.65       29.9
```

```
1000. MORG-339_phosphopeptide_TMT_12%ACN_032017_OT.7253.7253.3 (tASGSsVTSLEGTR)

   Ion         m/z   Intensity
   b5++      317.17       12.1
   y3+       333.26       13.0
   y7++      382.27        9.8
   b6++      400.87        7.4
   b2+       402.03        8.0
   y8++      431.60       13.1
   b7++      449.99       19.6
   y4+       462.24       18.2
   b3+       488.94       11.2
   b8++      500.60       20.0
   y9++      515.53       32.1
   a4+       518.09        8.4
   a9++      529.78       28.4
   b9++      544.57       20.2
   b4+       546.34       44.6
   y5+       575.37       11.2
   b10++     601.02       16.0
   a11++     651.25       10.7
   y6+       662.48       14.3
   b11++     665.41       18.6
   y13++     666.16        9.7
   b12++     693.99       12.3
   b13++     744.64       12.7
   y7+       763.40       16.7
   b7+       899.50        9.7
   b8+      1000.44        8.1
```

```
1001. MORG-339_phosphopeptide_TMT_8%ACN_032017_OT.5998.5998.3 (aSVASDPEsPPGGNEPAAASGQR)

   Ion         m/z   Intensity
   y4+       447.22        8.8
   y5+       518.35       11.7
   b4+       558.47        6.2
   y6+       589.39       11.9
   b5+       645.49        8.2
   y14++     654.93       18.5
   y7+       660.74        6.5
   a12++     689.29        8.9
   b13++     731.78        6.7
   y15++     738.49       16.9
   y8+       757.35       17.6
   b6+       760.46       17.6
   a14++     774.75        9.8
   b14++     788.54       63.3
   y16++     802.38       19.7
   y17++     851.54       20.3
   b15++     853.02       23.2
   a16++     887.62       10.4
   y18++     909.26        7.9
   a17++     923.39        8.8
   b17++     937.16        8.3
   y19++     952.54        6.8
   b18++     972.63       11.0
   b8+       986.60       12.8
   y20++     987.57        9.0
   a19++     994.23        7.5
   b19++    1007.91       10.6
   b20++    1051.63        9.2
   y11+     1057.44        8.7
   y13+     1211.53        6.1
   b15+     1704.80        7.7
```

```
1002. MORG-339_phosphopeptide_TMT_8%ACN_032017_OT.10184.10184.3 (sPsPPPDGSPAATPEIR)

   Ion         m/z   Intensity
   y2+       288.17       10.2
   a3++      317.19       19.1
   b1+       397.23        8.4
   b5++      427.97       42.6
   y9++      471.57        7.0
   y10++     499.79        7.2
   y4+       514.32       25.5
   a8++      548.33       16.6
   y11++     557.56        9.3
   b8++      562.48       11.0
   a9++      592.02        7.3
   y12++     605.64        7.1
   b9++      606.47       15.4
   y5+       615.37       22.7
   a3+       633.32        7.9
   a10++     640.27       11.7
   b10++     654.55       14.4
   b3+       661.28       19.5
   y14++     703.19       36.3
   b12++     726.07       21.0
   y7+       757.45       12.4
   b4+       758.48       20.6
   b13++     776.11        9.4
   y8+       854.51       29.1
   b5+       855.46       23.6
   y9+       941.67        6.9
   b7+      1067.44        9.6
   b9+      1211.44       19.4
```

```
1003. MORG-339_phosphopeptide_TMT_4%ACN_032017_OT.3469.3469.3 (dDEkEPEEGEDDRDsANGEDDS)

   Ion         m/z   Intensity
   b2+       460.28        5.8
   b6++      586.85        4.3
   b7++      651.47        6.0
   y7+       706.96        7.0
   b8++      716.25       13.0
   a9++      730.38        5.0
   b10++     808.77        5.5
   b11++     866.77       16.0
   y16++     910.09        5.7
   a4+       918.66        6.9
   b12++     924.28       18.6
   b4+       946.04        9.4
   y17++     959.10       19.2
   a13++     987.70        7.1
   b14++    1059.81       13.3
   y10+     1145.44        6.3
   y11+     1260.41        7.4
   y20++    1266.91        6.0
   y21++    1324.00        6.1
   y14+     1561.39        5.3
   y16+     1819.82        4.8
```

```
1004. MORG-339_phosphopeptide_TMT_18%ACN_032017_OT.16024.16024.4 (eAQVNVRMDsFDEDLARPSGLLAQER)

   Ion         m/z   Intensity
   a9+++     415.32       18.3
   b7++      513.81       13.5
   b8++      579.16        9.9
   b9++      637.05       46.5
   y12++     655.88       43.6
   b4+       657.59       15.5
   y13++     713.61       26.6
   a17+++    753.62       13.0
   b17+++    762.90       21.7
   b5+       771.46       14.8
   y14++     778.15       45.6
   a11++     780.10       11.8
   y7+       786.09       60.0
   b11++     793.95       16.7
   b18+++    795.07       60.2
   y20+++    796.49       41.5
   y21+++    829.37       17.5
   y15++     835.64       16.9
   a6+       842.50       11.2
   b12++     851.68       26.6
   y22+++    867.30       19.2
   b6+       870.61       13.7
   a21+++    871.81       10.5
   y8+       873.69       10.5
   b21+++    880.58        9.6
   y23+++    899.83       12.5
   a22+++    909.25       40.1
   b13++     916.01       18.8
   b22+++    918.87       11.9
   y24+++    942.57       14.1
   y9+       970.50       25.4
   b14++     973.68       35.5
   y17++     992.69       26.7
   b25+++   1028.13       23.3
   b15++    1030.35       12.8
   b16++    1065.10       10.4
   y19++    1115.76       10.2
```

```
1005. MORG-339_phosphopeptide_TMT_14%ACN_032017_OT.7746.7746.3 (hGSPtAPIGAGSPEFTEQGR)

   Ion         m/z   Intensity
   y8++      482.44       16.0
   a8++      522.09       11.9
   b8++      535.85       13.9
   b9++      564.55       12.9
   y11++     589.48       12.2
   y5+       590.35       12.1
   b10++     599.92       20.3
   b11++     628.47       13.4
   b12++     671.59       15.7
   y13++     674.90       12.2
   b13++     720.73       16.9
   y14++     723.65       37.7
   y6+       737.59       22.1
   a5+       761.23       13.1
   b5+       789.67       14.1
   b15++     858.15       16.8
   b6+       860.37       22.4
   y17++     897.97       13.2
   y8+       963.46       23.9
   y9+      1050.54       11.6
   a19++    1051.51       11.6
   y10+     1107.58       15.5
   y12+     1235.60       12.0
```

```
1006. MORG-339_phosphopeptide_TMT_8%ACN_032017_OT.17400.17400.3 (nPLGDDDAsATAskTETsQVAPA)

   Ion         m/z   Intensity
   b3+       554.50        6.9
   y10++     670.67        7.1
   b5+       726.41        7.0
   y7+       753.65        9.5
   y14++     875.69        6.6
   y8+       882.65       11.8
   a7+       928.84        9.3
   b7+       956.45       11.0
   y15++     959.24       10.5
   a14++     967.38        6.9
   y16++     994.92        9.8
   a8+       999.78        9.4
   b8+      1027.67        6.6
   y17++    1052.81        9.3
   b16++    1096.87       12.2
   b17++    1146.93        7.4
   y21++    1252.65        6.2
   b20++    1344.43       50.9
   b21++    1379.89      102.9
```

```
1007. MORG-339_phosphopeptide_TMT_20%ACN_032017_OT.13783.13783.3 (sRtHDGLAEVLVLEGSk)

   Ion         m/z   Intensity
   b1+       317.30        8.1
   y1+       376.38        8.5
   b5++      453.77        8.6
   y2+       463.40        9.8
   y3+       520.37       23.0
   b8++      574.54        7.2
   b9++      639.07       16.6
   y4+       649.44       19.4
   a10++     674.46       10.1
   b10++     688.48       18.5
   b11++     745.10       37.4
   y5+       762.67       30.9
   a4+       763.65       24.4
   b12++     794.58       19.4
   a13++     837.28       10.3
   y14++     848.35       11.4
   b13++     851.20       29.7
   y6+       861.67       20.5
   a14++     901.71        7.7
   b5+       906.67        9.6
   b14++     915.59       35.8
   a15++     929.88        7.8
   y15++     938.81        8.7
   b15++     944.06       21.8
   y7+       974.66       12.1
   b16++     987.71       13.5
   y12+     1443.88        8.3
```

```
1008. MORG-339_phosphopeptide_TMT_18%ACN_032017_OT.9203.9203.4 (aAkPcPTEASPPAASPSGDsSPPATAPYDPR)

   Ion         m/z   Intensity
   y5+       647.33       53.9
   b9++      692.72       20.8
   b17+++    693.47       17.9
   y6+       718.46       25.4
   y13++     727.76       16.2
   b10++     736.35       23.9
   a12++     819.40       19.6
   b4+       826.97       14.9
   b12++     833.77       16.0
   y16++     847.58       18.2
   a13++     855.21       20.9
   a22+++    859.07       23.9
   y26+++    868.75       46.2
   b13++     869.47       65.5
   b14++     904.81       35.5
   b24+++    924.44       23.6
   y18++     926.56       15.0
   a15++     933.54       21.3
   a25+++    948.30       45.2
   y28+++    954.28       14.4
   a5+       958.12       37.7
   y19++     962.50       13.6
   b26+++    981.42       33.4
   a16++     982.43       28.0
   y9+       987.54       15.8
   b17++    1040.16       19.4
   b18++    1068.44       26.6
   y10+     1084.50       16.3
   b29+++   1106.73       19.4
   b19++    1126.42       17.3
   y25++    1253.45       20.6
   y13+     1453.68       18.0
   b10+     1471.89       27.3
   y14+     1510.61       13.9
```

```
1009. MORG-339_phosphopeptide_TMT_14%ACN_032017_OT.8249.8249.4 (qSNASsDVEVEEkETNVSkEDTDQEEk)

   Ion         m/z   Intensity
   y2+       505.40       12.6
   y3+       634.42       12.9
   b16+++    762.59       26.5
   a17+++    786.39       13.9
   y15+++    823.42       27.0
   y16+++    866.55       27.7
   y5+       877.07       15.4
   y11++     883.51       14.8
   y17+++    909.24       18.9
   a19+++    934.55       12.3
   y12++     940.57       14.1
   y18+++    942.50       35.7
   a13++     957.90       12.8
   b13++     971.30       26.5
   a20+++    977.30       18.6
   y6+       978.50       47.1
   y19+++    984.80       87.9
   b20+++    986.99       33.1
   y13++     990.83       43.3
   b7+       999.49       28.4
   y20+++   1018.45       21.4
   b14++    1036.28       12.9
   y14++    1055.05       15.1
   y21+++   1056.75       31.0
   b22+++   1059.06       26.1
   a8+      1070.92       17.5
   y7+      1093.46       13.8
   b23+++   1097.39       40.6
   y22+++   1112.42       17.9
   b24+++   1140.08       16.8
   y23+++   1140.90       14.9
   b16++    1143.29       13.2
   y24+++   1164.51       11.8
   b25+++   1183.17       14.9
   a9+      1199.67       18.6
   a18++    1222.30       12.5
   b26+++   1226.06       15.2
   y26+++   1232.30       15.0
   y16++    1298.60       17.9
   b10+     1326.71       12.9
```

```
1010. MORG-339_phosphopeptide_TMT_8%ACN_032017_OT.6414.6414.3 (sAGEDNDGPVLTDEQkSR)

   Ion         m/z   Intensity
   y5++      438.97        8.4
   y6++      496.57        7.9
   y7++      546.94        6.7
   y3+       619.40        6.8
   b4+       654.14        8.5
   y10++     701.77       10.4
   y11++     730.10       28.2
   b5+       769.45       15.5
   a13++     776.76       10.4
   y12++     787.11       53.9
   b13++     790.76       14.9
   y13++     844.73       15.5
   y14++     902.16       10.2
   a15++     905.36        6.8
   y15++     966.75        8.9
   y6+       991.76        6.5
   y16++     995.36        8.9
   b7+       998.49       12.9
   y17++    1030.75       12.0
   y7+      1092.61       12.4
```

```
1011. MORG-339_phosphopeptide_TMT_18%ACN_032017_OT.16304.16304.3 (rAEAQALLQDYVSTQsAEE)

   Ion         m/z   Intensity
   b5++      393.38       12.5
   a6++      415.09       21.1
   y7++      416.14        7.8
   a2+       429.08       15.3
   b2+       457.39        8.0
   a7++      471.43        7.7
   b7++      485.46       34.2
   y4+       515.11       10.9
   b8++      542.00       35.6
   b3+       586.33       11.4
   b9++      606.07       39.3
   y5+       643.16       11.5
   b4+       657.38        7.4
   b10++     663.62       24.8
   y11++     669.19        7.9
   y12++     725.52       10.1
   y6+       744.30       15.5
   b11++     745.21       38.0
   a12++     779.98       15.6
   y13++     782.18       13.1
   b5+       785.67       16.6
   b12++     794.82      103.4
   y7+       831.29       52.0
   b13++     838.26       23.6
   b6+       856.55        8.8
   b14++     888.82       20.3
   y8+       930.37       20.7
   b15++     952.45       15.8
   y9+      1093.39       13.9
   y10+     1208.44       20.7
   y11+     1336.50       19.4
   y12+     1449.58       14.6
   y13+     1562.73        8.5
```

```
1012. MORG-339_phosphopeptide_TMT_8%ACN_032017_OT.8202.8202.2 (sEDDSAVPVAk)

   Ion         m/z   Intensity
   b1+       397.30        6.2
   y2+       447.39        7.8
   b3+       641.34        6.8
   y4+       643.48       24.7
   y5+       742.44        7.9
   b4+       756.27       10.6
   y6+       813.49       15.2
   a5+       815.43        7.7
   y7+       900.56       13.4
   b6+       914.34        6.7
   b7+      1013.43       20.9
   y8+      1015.52       11.4
```

```
1013. MORG-339_phosphopeptide_TMT_10%ACN_032017_OT.8557.8557.4 (aSGPLDTSQTAssDSEDVSSSDRk)

   Ion         m/z   Intensity
   y5++      411.52       11.5
   b7++      436.49       17.7
   y6++      454.99       19.3
   y7++      504.53       10.3
   y8++      561.79       14.1
   b10++     594.53       30.2
   y9++      626.56       13.2
   b11++     629.59       12.0
   y15+++    654.01       26.0
   b5+       655.26       21.9
   y10++     670.04       20.8
   a17+++    671.05        9.9
   b17+++    679.84       14.2
   y16+++    696.96       18.8
   a12++     699.27       10.1
   b12++     713.45       27.2
   y17+++    725.59       20.6
   y11++     727.56       21.1
   a19+++    732.77       14.7
   y4+       734.53       12.6
   b19+++    741.83       26.4
   a6+       742.92       27.7
   b6+       770.36       16.5
   b20+++    771.71       12.4
   a13++     783.04       11.7
   b13++     796.62       24.8
   y19+++    797.71       13.9
   y12++     811.21       16.1
   a14++     839.89       12.9
   b14++     853.89       11.3
   b7+       871.62       20.9
   a23+++    881.11       24.4
   y13++     894.62       26.9
   a8+       930.13       26.6
   b8+       958.45       15.8
   y15++     980.57       36.1
   b18++    1069.79       11.8
   b9+      1086.57       16.3
   y17++    1087.60       16.4
```

```
1014. MORG-339_phosphopeptide_TMT_25%ACN_032017_OT.13949.13949.4 (kIQPQLPDEDGNHsDkEDEQPQVVVLk)

   Ion         m/z   Intensity
   y2+       489.45       21.0
   y3+       588.51       24.5
   y4+       687.53       29.9
   y5+       786.71       21.9
   y17+++    820.99       21.2
   y18+++    859.47       15.8
   y12++     935.83       35.9
   a20+++    939.17       31.6
   y20+++    940.42       23.5
   b20+++    948.48       17.0
   a21+++    971.63       17.5
   y21+++    972.79       29.9
   a15++     978.18       25.1
   b21+++    980.28       19.8
   y13++     992.77       74.8
   a7+      1006.66       29.8
   y22+++   1010.53       15.9
   y7+      1011.70       31.0
   y23+++   1052.78       14.9
   a24+++   1080.07       15.6
   y24+++   1085.48       16.4
   b25+++   1122.09       15.6
   y25+++   1127.74       17.1
   y8+      1139.24       16.8
   b26+++   1160.22       19.5
   y26+++   1165.99       39.0
   y17++    1230.37       17.5
   b17++    1235.90       16.4
   y9+      1268.74       21.5
   y20++    1410.01       22.6
   a12+     1536.53       24.4
```

```
1015. MORG-339_phosphopeptide_TMT_12%ACN_032017_OT.15952.15952.2 (sGFFVEsPR)

   Ion         m/z   Intensity
   y2+       272.16        4.3
   y3+       439.29        4.6
   y4+       568.28        5.6
   a3+       573.32        7.4
   b3+       601.30        8.5
   a4+       720.39        4.5
   b4+       748.35        6.2
   y6+       814.54        4.5
   b5+       847.45        7.5
   y8+      1018.48        7.0
```

```
1016. MORG-339_phosphopeptide_TMT_16%ACN_032017_OT.11496.11496.3 (sASPDDDLGSSNWEAADLGNEERkQk)

   Ion         m/z   Intensity
   b8++      555.27       10.8
   y5++      573.93       11.2
   b5+       767.29       15.7
   y9++      780.58       49.9
   y10++     838.38       12.5
   y11++     873.81       11.2
   b6+       882.44       15.7
   y12++     909.22       25.4
   b7+       997.36       21.1
   y4+      1017.87       10.8
   b18++    1070.53       10.8
   b19++    1098.97       10.7
   y15++    1124.08       24.2
   y5+      1146.98       16.3
   b9+      1167.83       19.2
   a21++    1206.58       11.0
   y17++    1210.85       11.4
   y19++    1296.01       44.9
   y21++    1411.00       38.3
   y23++    1517.11       56.4
   y24++    1560.00       16.6
   b25++    1606.27       11.6
```

```
1017. MORG-339_phosphopeptide_TMT_4%ACN_032017_OT.3432.3432.3 (rIADSSVQtDDEEGEGR)

   Ion         m/z   Intensity
   b4++      343.37        9.7
   y7++      396.60        5.3
   y4+       418.36        7.4
   b6++      430.35        5.8
   a7++      466.15        7.7
   y9++      544.34       12.2
   y5+       547.20        8.8
   b9++      634.24        8.6
   y6+       676.51       21.9
   a10++     677.48       13.6
   b4+       685.49       19.3
   b10++     691.73       22.3
   y12++     701.63       20.8
   b11++     749.83        7.3
   b5+       772.65        9.3
   y7+       791.38        5.1
   b12++     813.92        9.2
   b13++     878.41        6.6
   y8+       906.31        5.4
   b7+       958.65        6.5
   b15++     971.52        5.7
```

```
1018. MORG-339_phosphopeptide_TMT_14%ACN_032017_OT.12050.12050.3 (sTsPPPSPEVWAESR)

   Ion         m/z   Intensity
   y2+       262.16        6.6
   b1+       317.29       13.2
   y6++      374.33        7.4
   a5++      376.13        7.4
   y3+       391.26        7.8
   a6++      424.24        7.7
   y4+       462.30       14.1
   a7++      468.24       10.0
   b7++      482.26       11.4
   y8++      487.34       38.4
   y9++      530.59        6.2
   a3+       557.48        5.8
   y10++     579.56       20.2
   b3+       585.24       17.8
   b9++      595.60       18.8
   y11++     627.43       38.1
   a10++     630.33        7.0
   y12++     676.45       28.7
   b11++     738.06       14.4
   y6+       747.35       11.8
   b12++     773.31        9.1
   b5+       779.47       12.3
   a13++     824.11        5.7
   b13++     838.18        8.2
   y7+       876.42        7.2
   b14++     881.46        8.6
   b7+       963.49       27.2
   y8+       973.53       21.5
   y9+      1060.54        7.8
   y10+     1157.41        9.1
   b9+      1189.66        6.1
   b10+     1288.65        6.3
```

```
1019. MORG-339_phosphopeptide_TMT_8%ACN_032017_OT.9097.9097.3 (ssSPVLVEEEPERSLR)

   Ion         m/z   Intensity
   y4++      266.12        5.9
   b1+       317.23        7.3
   y6++      379.30        6.5
   y7++      443.74        5.8
   y8++      508.33       10.7
   y4+       531.42        7.8
   a3+       542.96        5.1
   b3+       571.16        5.5
   b9++      619.26        6.1
   y10++     622.18        6.2
   a4+       640.66        6.5
   y5+       660.47        7.0
   b4+       667.79        5.5
   a10++     669.90       22.1
   y11++     678.76       15.3
   b10++     683.83        7.9
   y6+       757.53        7.6
   b5+       767.63        9.8
   y13++     777.07       33.7
   a12++     782.87        5.5
   b12++     797.05        7.1
   y14++     820.42        6.6
   b13++     875.03        5.1
   b6+       880.52        7.1
   b15++     974.86        8.8
   y9+      1144.62        4.9
   b9+      1237.48        6.2
```

```
1020. MORG-339_phosphopeptide_TMT_8%ACN_032017_OT.14252.14252.4 (sASPDDDLGSSNWEAADLGNEER)

   Ion         m/z   Intensity
   y2+       304.14        6.6
   y6++      359.17        6.9
   y7++      416.95        8.2
   y3+       433.37        6.9
   b6++      441.36        8.2
   y8++      452.38        9.1
   y9++      487.85        8.1
   b7++      499.12       11.8
   y14+++    526.38        8.6
   a8++      541.74        7.5
   y4+       547.35       14.4
   b13+++    548.25        6.7
   b8++      555.83       31.1
   a14+++    581.73        6.7
   y16+++    583.63        9.6
   b9++      584.44       17.1
   b14+++    591.05        7.8
   y5+       604.29       29.8
   a15+++    605.42       13.6
   a10++     613.89       10.4
   b15+++    614.79        7.3
   y17+++    621.99        7.4
   b10++     627.95       11.5
   b16+++    638.16       14.1
   y11++     645.51        6.7
   y18+++    659.99        9.2
   a17+++    667.50        7.7
   b11++     671.09       10.2
   b17+++    676.68       11.4
   y12++     702.49        9.6
   b18+++    714.47       14.4
   y6+       717.37       14.5
   a19+++    723.78        6.5
   b19+++    733.32        8.9
   y13++     746.04       19.8
   b5+       767.38       11.0
   y14++     789.42       24.8
   b21+++    814.37       10.7
   y15++     818.00       66.8
   b13++     821.42       10.9
   y7+       832.45       11.9
   a14++     872.14        6.3
   y16++     874.68       17.7
   b6+       882.40        8.1
   b14++     885.81       12.4
   y8+       903.57       11.8
   b15++     921.54       12.4
   y17++     931.96       12.6
   b16++     956.70        9.3
   y9+       974.41        9.1
   y18++     989.41        9.4
   b7+       997.39       37.4
   b17++    1014.63       19.5
   b8+      1110.55       10.1
```

```
1021. MORG-339_phosphopeptide_TMT_8%ACN_032017_OT.9066.9066.3 (tEsVPSDINNPVDR)

   Ion         m/z   Intensity
   y2+       290.22        5.0
   a3++      300.07        6.9
   y5++      300.89        4.6
   y6++      357.68        6.6
   y3+       389.22        4.3
   b5++      412.27        6.3
   y7++      414.32        4.7
   b6++      455.22        4.6
   b2+       460.25        5.7
   y8++      471.93        7.8
   y4+       486.27       16.1
   b7++      513.46       11.2
   a8++      555.29       15.8
   y10++     563.88       30.8
   b8++      569.91       21.7
   a3+       599.33       23.9
   y5+       600.29       34.7
   b9++      626.62        8.1
   a10++     669.95        5.2
   b10++     683.99       17.6
   y6+       714.32       23.6
   b4+       726.34       20.3
   b11++     732.34        4.8
   b12++     782.29        6.7
   y7+       827.54        7.2
   b13++     839.67        6.9
   b6+       910.45        9.0
   b7+      1025.44       18.0
   y9+      1029.44        5.4
   b8+      1138.58        7.7
   b10+     1366.63        4.5
```

```
1022. MORG-339_phosphopeptide_TMT_22%ACN_032017_OT.22701.22701.3 (sVstSPSILPAYLk)

   Ion         m/z   Intensity
   b1+       317.30       19.7
   y1+       376.29       19.9
   y5++      410.83       37.7
   a5++      411.98        9.7
   b2+       416.36       10.0
   y6++      467.30       24.0
   y2+       489.37       22.1
   y7++      524.25        5.6
   a8++      560.52        5.9
   b8++      574.82       14.6
   b3+       583.37        6.2
   y9++      616.07        7.5
   a9++      617.32       11.3
   b9++      631.44       37.0
   y3+       652.58       21.3
   y10++     659.34        6.3
   a10++     666.30       10.3
   b10++     679.74       10.9
   b11++     715.58       14.0
   y4+       723.52       11.9
   y11++     749.99        6.1
   b12++     797.01       23.9
   y5+       820.53       39.2
   b5+       851.23        6.1
   b13++     853.37       20.2
   y6+       933.75       11.3
   b8+      1148.51       18.4
   b9+      1261.58       35.4
```

```
1023. MORG-339_phosphopeptide_TMT_10%ACN_032017_OT.14508.14508.3 (dEILPtTPISEQk)

   Ion         m/z   Intensity
   a1+       317.30       10.3
   y4++      360.84        5.3
   y1+       376.30       10.2
   a2+       446.25        5.1
   y6++      465.96       11.5
   b6++      489.35        6.7
   y2+       504.47       14.9
   y7++      516.42        6.2
   b7++      540.60        5.4
   b3+       587.44        8.7
   b8++      588.40        5.4
   a9++      631.61       30.0
   y3+       633.51       10.2
   b9++      645.49       18.7
   y9++      655.23       24.3
   b10++     688.60        5.7
   b4+       700.52       23.6
   y10++     711.87        5.8
   y4+       720.51       15.5
   b12++     817.50        5.6
   y6+       930.77        5.4
   b9+      1289.78        6.7
```

```
1024. MORG-339_phosphopeptide_TMT_12%ACN_032017_OT.13660.13660.3 (lLPEGEETVEsDDDk)

   Ion         m/z   Intensity
   b1+       343.21        5.8
   y1+       376.32        6.6
   b2+       456.37       10.0
   y2+       491.44       10.6
   b8++      549.92       11.9
   y7++      558.55       10.1
   b9++      599.33        4.4
   y3+       606.36       13.2
   y8++      609.36       12.6
   b10++     663.99       15.2
   b4+       682.62        5.5
   y4+       721.02       28.1
   b6+       868.45        5.9
   y13++     880.23       12.8
   y5+       888.32       16.4
   b14++     920.28        7.3
   b7+       997.42       13.6
   y6+      1017.42       10.8
   b8+      1098.60        8.3
   y7+      1116.52        5.3
   b9+      1197.78       10.5
```

```
1025. MORG-339_phosphopeptide_TMT_4%ACN_032017_OT.11365.11365.2 (tASGSsVtsLEGTR)

   Ion         m/z   Intensity
   y3+       333.16        6.5
   y4+       462.21        6.2
   y6+       742.37        8.0
   a7+       871.37        8.1
   b11+     1489.61        7.4
```

```
1026. MORG-339_phosphopeptide_TMT_8%ACN_032017_OT.12289.12289.3 (vADPDHDHtGFLTEyVATR)

   Ion         m/z   Intensity
   y4+       446.33        6.0
   b3+       515.35       20.2
   y8++      516.36       10.0
   a8++      545.01        5.5
   b8++      559.06        6.1
   b9++      648.88        6.2
   y5+       689.66        7.0
   y11++     709.16        5.8
   b5+       727.41        9.8
   b11++     751.02        5.8
   y12++     777.98       11.8
   a12++     794.11        5.5
   b12++     807.91       11.9
   y6+       818.23       17.6
   y13++     835.54        6.1
   b13++     858.54        6.2
   b6+       864.50        5.8
   y14++     904.20        9.8
   a14++     909.27        8.1
   y7+       919.52        6.4
   y15++     961.36       17.4
   y16++    1010.24       20.3
   b16++    1094.32        6.5
   a17++    1115.04        7.1
   b8+      1116.51        6.9
```

```
1027. MORG-339_phosphopeptide_TMT_4%ACN_032017_OT.4792.4792.3 (sTSPcQtPEPDNEAHIR)

   Ion         m/z   Intensity
   y3+       425.24        6.1
   y8++      476.50       17.6
   y9++      540.34        6.2
   b3+       585.71        7.0
   y10++     589.25       14.6
   a9++      675.57        5.5
   y11++     679.42        8.4
   a10++     723.38       10.7
   b10++     738.11       16.6
   y6+       739.41       10.5
   y12++     744.01       40.9
   y13++     823.58        8.3
   b5+       842.38        7.8
   y14++     872.44       17.2
   a13++     903.30        4.8
   a14++     938.21        5.7
   y8+       951.53       16.2
   b14++     952.58        8.7
   b6+       970.45       10.2
   b15++    1020.89        9.7
   b16++    1077.81        8.0
   y10+     1177.69       10.1
   b9+      1377.68        8.3
```

```
1028. MORG-339_phosphopeptide_TMT_10%ACN_032017_OT.6060.6060.4 (sFVPEEEkHEERVPR)

   Ion         m/z   Intensity
   y4++      264.26       12.7
   y5++      328.84       21.5
   b1+       397.15        8.7
   a5++      421.50       12.2
   y8+++     427.37       21.8
   y9+++     470.28       17.7
   a6++      485.62       10.7
   y10+++    513.41       33.8
   y4+       527.18       11.3
   b9+++     541.56        9.9
   b2+       544.39        8.3
   a7++      550.69       10.6
   y11+++    556.55       13.5
   b7++      564.00       15.8
   a10+++    574.87       18.3
   b10+++    583.89       39.0
   y12+++    588.73       36.2
   a3+       615.49       12.6
   y13+++    621.83       16.2
   b3+       643.33       20.7
   y14+++    670.88       30.0
   b12+++    679.59       12.1
   a4+       712.04       24.2
   a8++      728.54        8.8
   b8++      743.35       16.9
   b14+++    744.71       20.1
   y10++     769.44        8.3
   b5+       869.43       16.6
   b10++     876.04       25.9
   y12++     882.63       18.9
   b11++     940.59       10.9
```

```
1029. MORG-339_phosphopeptide_TMT_14%ACN_032017_OT.11816.11816.2 (aLGLEEsPEEEGk)

   Ion         m/z   Intensity
   y2+       433.35        8.7
   b4+       584.56        9.8
   b5+       713.54       10.5
   y5+       820.51        9.0
   b6+       842.56       11.0
   y6+       917.58       23.9
   a7+       981.86        9.1
   b7+      1009.60        8.4
   y7+      1084.52        9.8
   b8+      1106.72       11.8
   b9+      1235.14        9.0
   b11+     1493.82        9.7
```

```
1030. MORG-339_phosphopeptide_TMT_10%ACN_032017_OT.7272.7272.3 (akEVENEQTPVsEPEEEk)

   Ion         m/z   Intensity
   b8++      694.08       15.3
   y10++     727.07        9.1
   b9++      744.90       16.5
   a3+       759.59        9.9
   b11++     842.61       12.8
   y12++     855.51       11.2
   y5+       860.86       16.1
   b4+       886.57       10.6
   a12++     911.53       24.0
   b12++     926.13       40.7
   y14++     977.24       16.8
   b13++     990.16        8.4
   a14++    1025.41       16.4
   b14++    1039.14       10.7
   y16++    1091.35       11.5
   a6+      1101.52       13.8
   y7+      1156.25       11.9
   b16++    1168.12       11.4
   y8+      1255.40       10.6
   b9+      1487.82       10.5
```

```
1031. MORG-339_phosphopeptide_TMT_16%ACN_032017_OT.2328.2328.4 (eEEENSSNDTASQSTSPsQPRkk)

   Ion         m/z   Intensity
   y2++      367.08        7.6
   y4++      494.08       18.3
   a9++      618.50        9.0
   y12+++    624.04       10.1
   b16+++    642.95       10.9
   y13+++    647.52        7.7
   a17+++    665.28       10.1
   b17+++    674.82        8.5
   y14+++    680.94       12.4
   b10++     682.96       14.7
   y15+++    719.53       11.8
   y2+       733.42        8.3
   b4+       746.51       11.4
   a12++     748.18        9.2
   y16+++    757.39       11.1
   b12++     761.52       12.7
   a19+++    763.93        7.9
   b19+++    772.84       10.9
   y9++      784.36       18.6
   y17+++    786.45        8.9
   a20+++    796.49       15.5
   b20+++    805.55       19.4
   y18+++    815.08       20.7
   a21+++    848.34        8.0
   b5+       860.40       16.1
   b14++     869.66       10.8
   y12++     935.26       20.9
   b6+       947.54       16.4
   y13++     970.68       15.8
   b22+++    977.01        7.8
   y4+       986.60        7.9
   y14++    1021.22       10.0
   b7+      1034.41       11.4
   b8+      1148.49        8.8
   b9+      1263.58       11.5
   b11+     1435.60       12.3
```

```
1032. MORG-339_phosphopeptide_TMT_16%ACN_032017_OT.8759.8759.4 (dSSEATGWEEPsPPSIRRk)

   Ion         m/z   Intensity
   a1+       317.17        8.5
   y7+++     361.60       14.2
   y1+       376.38        7.9
   y8+++     417.11       16.2
   y9+++     449.71       16.6
   b12+++    529.59        7.7
   y2+       532.17       12.6
   y7++      541.97       19.7
   b13+++    561.78        8.0
   a14+++    584.16       10.9
   b14+++    593.98        7.1
   y12+++    597.70       11.4
   a15+++    613.45        7.6
   y13+++    616.54       15.1
   y8++      625.15       31.1
   a10++     646.92       20.8
   b4+       648.10       17.8
   y14+++    650.31       19.1
   a16+++    651.35       15.1
   y15+++    673.76       10.1
   b17+++    712.73       10.1
   b5+       719.46       14.3
   y10++     738.27       10.7
   y17+++    746.33        7.3
   a18+++    755.10        9.2
   y11++     802.90       17.4
   b6+       820.44       11.6
   b7+       877.47       17.1
   y12++     896.04       10.7
   y13++     924.52       12.6
   b8+      1063.46       11.0
   b9+      1192.57       10.3
   b10+     1321.62       13.7
```

```
1033. MORG-339_phosphopeptide_TMT_10%ACN_032017_OT.3313.3313.3 (sSNPPARsQSSDTEQPsPTSGGGk)

   Ion         m/z   Intensity
   y8++      499.97        4.5
   b3+       518.34        5.0
   a9++      603.91        4.5
   y6+       735.26        4.5
   b12++     762.61       12.0
   y14++     828.63        7.1
   y7+       832.63       10.8
   b14++     877.58       10.3
   a7+       911.94        5.4
   y16++     936.72        6.3
   b15++     941.59       19.7
   a16++     976.12       11.4
   y8+       999.56        6.5
   y9+      1096.58       10.4
   y18++    1097.54       11.3
   y19++    1133.38        4.3
   b19++    1172.85        4.5
   y20++    1182.18        7.1
   b20++    1215.90        4.4
   y10+     1224.66        4.8
   b23++    1301.91        4.3
   y12+     1454.71        4.4
   y13+     1569.75        5.0
```

```
1034. MORG-339_phosphopeptide_TMT_22%ACN_032017_OT.19482.19482.3 (atLSDSEIETNSATSAIFGk)

   Ion         m/z   Intensity
   y1+       376.29       13.8
   y2+       433.34       35.2
   y3+       580.44       42.7
   y4+       693.46       31.5
   y12++     728.26        9.8
   b11++     736.26       10.9
   y5+       764.55       16.1
   a12++     765.60       12.8
   b12++     779.75       15.2
   y13++     784.17       10.2
   b5+       797.58       12.1
   a13++     800.69       10.3
   b13++     815.36       15.1
   y14++     849.33       25.0
   y6+       851.65       25.7
   b14++     865.70       13.1
   b6+       884.79       12.5
   a15++     894.59        8.6
   b16++     944.81       20.5
   y7+       952.46       22.2
   a7+       985.29       11.5
   a17++     986.75        9.7
   y17++     993.04       12.4
   b17++    1001.24       62.3
   b7+      1013.34       13.8
   y8+      1023.58        9.9
   b18++    1074.76       52.7
   b19++    1103.37       22.4
   b8+      1126.55       23.4
   b9+      1255.61       18.0
   b10+     1356.59        8.8
   b11+     1470.78        8.2
```

```
1035. MORG-339_phosphopeptide_TMT_6%ACN_032017_OT.5394.5394.4 (gPSEAPQEAEAEEGAtSDGEkkR)

   Ion         m/z   Intensity
   y5+++     359.40        8.1
   y6+++     397.59        7.8
   b8++      513.28        8.2
   a9++      534.93        7.0
   y5++      538.43       15.7
   b9++      548.92       14.5
   y6++      596.14        6.8
   b4+       600.39        9.2
   b10++     613.60        9.6
   b11++     649.20       11.9
   b17+++    650.83        8.1
   b5+       671.55       20.8
   a18+++    679.71        8.6
   y15+++    705.99       10.2
   b19+++    708.37       11.4
   b12++     713.58        9.7
   y8++      729.70       13.7
   a6+       739.98        7.1
   a20+++    741.83       10.7
   y16+++    749.30       15.3
   b20+++    751.37       10.9
   a13++     764.27       13.5
   y9++      765.22        8.7
   b6+       768.41       23.3
   y10++     793.93        8.2
   y20+++    890.33        6.8
   b7+       896.59        8.3
   y21+++    919.83       11.1
   b17++     975.44        8.6
   b8+      1025.35       14.1
   b9+      1096.49        6.7
   a20++    1112.76        8.0
```

```
1036. MORG-339_phosphopeptide_TMT_16%ACN_032017_OT.8143.8143.4 (kGAEEEEEEEDDDsEEEIkAIR)

   Ion         m/z   Intensity
   y7++      544.49       16.3
   a8++      552.59       10.5
   y8++      608.95       17.8
   a9++      616.59       11.2
   y14+++    663.33        8.6
   a10++     681.64       14.3
   y9++      692.58       13.9
   a16+++    710.93       12.9
   y4+       716.44       13.4
   b5+       744.13       12.3
   y10++     749.85       19.5
   b17+++    763.41        9.9
   y17+++    792.19       10.8
   b18+++    801.20       17.9
   y11++     807.40       18.9
   b12++     810.11       19.1
   y12++     865.15        9.2
   b13++     868.11       19.1
   y21+++    920.36       14.3
   a21+++    972.34        9.3
   b7+      1002.99       11.8
   y15++    1058.51       19.3
   b16++    1080.70        9.7
```

```
1037. MORG-339_phosphopeptide_TMT_8%ACN_032017_OT.11724.11724.2 (tLsPtPSAEGYQDVR)

   Ion         m/z   Intensity
   b1+       331.27        7.7
   b2+       444.42        8.0
   b3+       611.28        7.7
   y6+       737.48       17.0
   y7+       866.49       15.2
   b5+       889.27       12.6
   y8+       937.25       15.1
   a6+       958.59       13.1
   y9+      1024.26        8.6
   b9+      1273.56       13.0
   a10+     1302.67       14.1
   y12+     1399.65       19.7
   a11+     1465.53        8.4
   b13+     1736.81       13.2
   b14+     1835.65        8.9
```

```
1038. MORG-339_phosphopeptide_TMT_8%ACN_032017_OT.22246.22246.3 (aEAkEEsEEsDEDmGFGLFD)

   Ion         m/z   Intensity
   y2+       281.07        7.7
   y3+       393.98        7.1
   a4++      415.48        6.2
   b4++      429.60        8.0
   y4+       451.36       10.6
   y5+       598.35        6.0
   y6+       655.42       11.7
   b9++      771.33       11.3
   y13++     793.84       13.0
   y7+       802.32        6.7
   b4+       858.66       11.7
   y14++     877.17        8.8
   y8+       917.26       16.2
   b12++     977.01       16.1
   b13++    1034.61       17.3
   b14++    1108.15       23.1
   b15++    1136.29        9.6
   y10+     1161.57       14.3
   a16++    1195.93        9.4
   b16++    1210.22       53.1
   b17++    1238.25       23.6
   b18++    1295.38       50.1
```

```
1039. MORG-339_phosphopeptide_TMT_22%ACN_032017_OT.6778.6778.4 (rMsPkPELTEDQk)

   Ion         m/z   Intensity
   b3++      342.60       15.7
   a5+++     370.89       18.9
   y1+       376.32       19.4
   b5+++     380.28       33.2
   b7+++     455.44       29.4
   b8+++     493.40       21.6
   y2+       504.34       26.3
   y9+++     516.27       15.0
   a9+++     518.06       16.7
   b9+++     526.82       14.0
   y7++      546.13       15.2
   y10+++    548.34       27.4
   a5++      556.15       14.0
   a10+++    560.53       31.9
   b10+++    570.33       45.8
   y8++      594.64       12.4
   y11+++    603.59       14.1
   a6++      604.61       12.7
   b11+++    608.48       38.1
   b6++      618.28       15.9
   y3+       619.45       23.3
   a12+++    641.59       19.1
   b12+++    651.21       29.8
   b3+       684.53       13.0
   b8++      739.32       14.7
   y4+       748.53       19.7
   b4+       781.14       15.1
   y10++     822.17       47.6
   y5+       849.54       15.6
   b10++     854.60       15.0
   y6+       962.51       14.9
   y8+      1188.55       16.2
```

```
1040. MORG-339_phosphopeptide_TMT_10%ACN_032017_OT.11112.11112.3 (sNSSDAPGEEssSETEkEIPVEQk)

   Ion         m/z   Intensity
   y8++      715.16       14.2
   b5+       720.37       28.1
   b6+       791.45       19.9
   y10++     829.63       31.8
   y11++     894.96       12.7
   y6+       942.71       12.4
   y7+      1071.93       11.0
   b9+      1074.19       17.5
   y14++    1105.52       99.9
   b17++    1171.29       11.4
   a10+     1175.96       11.0
   b10+     1203.58       12.4
   y16++    1234.20       12.1
   b18++    1235.16       14.3
   y17++    1262.36       22.3
   b19++    1292.24       25.9
   y18++    1310.99       17.9
   y19++    1346.52       13.7
```

```
1041. MORG-339_phosphopeptide_TMT_18%ACN_032017_OT.22574.22574.3 (iVTVAAISQDsNPAtPNVSTNmEEF)

   Ion         m/z   Intensity
   y9++      543.48        9.3
   b4+       642.52       15.9
   y11++     682.41        8.8
   a5+       685.30       11.8
   b5+       713.50       16.3
   b12++     754.81       11.5
   b6+       784.54       18.8
   y6+       786.34       13.0
   b13++     803.71       17.8
   b14++     839.09       18.9
   y7+       873.37       32.3
   b7+       897.61       40.1
   a15++     915.39       12.2
   a8+       956.90        9.6
   a16++     964.01        9.3
   y8+       972.39       16.0
   b16++     977.46       15.4
   b8+       984.72       16.7
   a18++    1070.21        9.9
   a9+      1084.82       49.0
   y9+      1086.58       10.0
   b9+      1112.73       10.5
   y10+     1183.49       42.9
   y21++    1199.66        9.5
   b10+     1227.69       11.8
   b22++    1309.54        9.2
   y11+     1364.48       24.9
   a11+     1366.56        8.7
   b23++    1373.88        9.3
   b12+     1508.89       12.7
   y13+     1532.67       17.0
```

```
1042. MORG-339_phosphopeptide_TMT_30%ACN_032017_OT.16169.16169.3 (rNsELFQSLISk)

   Ion         m/z   Intensity
   y1+       376.29       12.6
   a5++      441.22        9.7
   b5++      455.26        9.9
   y2+       463.34       17.5
   a6++      514.91        8.2
   b6++      528.74        9.0
   y3+       576.41       19.3
   a7++      579.18       10.6
   b7++      592.72       18.6
   a8++      622.45       11.8
   b8++      636.46       31.9
   a3+       639.08       14.4
   b3+       667.39        9.7
   b9++      692.83        8.1
   a10++     735.65        9.2
   b10++     749.31       17.8
   y5+       776.59        9.3
   a11++     778.55        8.0
   b11++     793.07        7.9
   y6+       904.58        8.9
```

```
1043. MORG-339_phosphopeptide_TMT_10%ACN_032017_OT.19628.19628.3 (vEEEDTGDPFGFDsDDESLPVSSk)

   Ion         m/z   Intensity
   y5++      373.68       15.2
   y1+       376.30       10.3
   y3+       550.37       20.0
   b3+       587.28        9.4
   y10++     653.66        8.7
   b4+       716.48       10.4
   y5+       746.52       63.6
   b5+       831.55       10.8
   y6+       859.58       16.2
   y14++     896.68       25.3
   b6+       932.46       18.5
   y7+       946.58       16.2
   b15++     975.68        8.9
   b7+       989.30       11.0
   a16++    1018.60       31.4
   b16++    1032.77       11.7
   a8+      1076.53       10.0
   a17++    1083.13       19.3
   b17++    1097.60       28.8
   b8+      1104.51       35.5
   b18++    1140.55       21.5
   y9+      1190.40        8.8
   b19++    1197.32       18.1
   y20++    1212.67       17.1
   b20++    1245.58        8.6
   b21++    1295.82       21.5
   b22++    1339.39        9.1
   b10+     1348.72       27.7
   y23++    1405.67        9.4
   b12+     1552.77       15.8
```

```
1044. MORG-339_phosphopeptide_TMT_10%ACN_032017_OT.11960.11960.4 (eRPkTPPSNLsPIEDAsPTEELR)

   Ion         m/z   Intensity
   y3+       417.32       10.3
   b2+       515.35        9.7
   b11+++    582.29       13.7
   b7++      632.46       10.2
   a13+++    643.73       12.6
   b13+++    652.81       24.6
   b14+++    695.91       28.9
   a9++      719.21       16.0
   b15+++    734.27       21.4
   y6+       744.43       22.1
   a16+++    748.36       32.8
   b16+++    757.99       26.4
   a10++     776.41       24.0
   b17+++    813.59       10.2
   a18+++    836.75       11.8
   y20+++    857.51       15.2
   b11++     873.84       11.4
   y21+++    889.78       13.1
   a12++     907.80       14.9
   y7+       911.42       10.5
   a20+++    912.88       12.8
   b20+++    922.81       15.9
   a4+       941.19       14.9
   a13++     964.33       15.4
   b21+++    965.68       22.2
   b13++     978.81       14.3
   y8+       982.46       13.0
   b22+++   1003.39       11.1
   y9+      1097.44       11.5
   b15++    1100.95       15.7
   a16++    1121.71       14.7
   y10+     1226.55       12.3
```

```
1045. MORG-339_phosphopeptide_TMT_8%ACN_032017_OT.6437.6437.3 (rAsISEPSDTDPELR)

   Ion         m/z   Intensity
   y2+       288.17       10.9
   b3++      312.70        7.7
   a4++      354.76        7.3
   b6++      477.26       20.6
   y4+       514.32       29.4
   y9++      515.43       17.7
   b7++      525.56        7.8
   b8++      569.06        8.9
   y10++     580.10        8.8
   a3+       596.15       13.6
   a9++      612.46       16.8
   y11++     622.80       24.7
   b3+       624.62       10.1
   b9++      626.60       17.8
   b10++     676.99       11.8
   y6+       730.46       10.7
   b11++     734.91       34.3
   y14++     799.05       11.6
   y7+       845.47        7.0
   b13++     847.71       12.7
   b14++     904.37        6.3
   y8+       932.39        6.9
   b6+       953.50       13.2
   y9+      1029.47       27.0
   y10+     1158.54        9.7
   y11+     1245.57       11.1
```

```
1046. MORG-339_phosphopeptide_TMT_16%ACN_032017_OT.13635.13635.4 (lkDLGHPVEEEDEsGDQEDDDDEIDDGDkDQDI)

   Ion         m/z   Intensity
   a11+++    559.95       11.3
   b6++      562.01       16.2
   b9++      724.33       13.4
   b10++     789.26       13.4
   y5+       847.48       17.6
   y20+++    849.98       16.6
   b11++     853.38       14.0
   b19+++    887.74       13.1
   y21+++    893.22       12.7
   b12++     911.22       25.3
   a20+++    916.85       14.8
   b20+++    926.56       14.3
   b21+++    964.48       14.6
   b13++     975.84       20.4
   y15++     976.64       13.4
   b22+++   1003.04       22.8
   y7+      1019.49       20.9
   a23+++   1036.01       15.1
   b23+++   1045.75       28.7
   b14++    1059.18       16.6
   y25+++   1060.26       12.8
   b24+++   1083.86       45.6
   b15++    1087.84       20.8
   y26+++   1093.40       16.7
   a6+      1095.11       14.5
   y17++    1104.61       20.4
   y8+      1134.52       13.5
   b16++    1145.31       24.9
   b26+++   1160.35       21.7
   a7+      1191.31       12.6
   b17++    1209.41       16.4
   b28+++   1217.91       22.9
   y9+      1249.61       29.5
   y31+++   1266.01       13.4
   b18++    1273.95       13.7
   y21++    1339.35       11.6
   y10+     1362.67       14.7
   b20++    1388.68       11.1
   b31+++   1417.99       14.8
   a32+++   1446.52       14.0
   b22++    1503.94       12.4
   b10+     1576.66       15.8
   b24++    1624.98       18.6
   y30++    1841.92       11.9
```

```
1047. MORG-339_phosphopeptide_TMT_4%ACN_032017_OT.2253.2253.2 (sGSLGSSNGDsDTER)

   Ion         m/z   Intensity
   y3+       405.21        7.4
   b3+       461.61        4.5
   y4+       520.46        5.9
   b4+       574.39        7.7
   b5+       631.39        5.0
   y5+       687.46        5.3
   b7+       805.40        9.1
   b8+       919.33        7.1
   y8+       973.21        5.5
   b9+       976.79        7.0
   y9+      1060.24        4.4
   b10+     1091.53        8.9
   y11+     1204.49        5.8
   b12+     1373.57        4.9
```

```
1048. MORG-339_phosphopeptide_TMT_10%ACN_032017_OT.10691.10691.4 (esDDkPEIEDVGsDEEEEEkk)

   Ion         m/z   Intensity
   b5++      557.54       14.5
   b7++      670.25       10.5
   y7++      690.09       18.6
   a15+++    746.35       11.8
   y8++      747.56       14.0
   b4+       756.29       14.7
   b9++      791.25       13.2
   y16+++    800.71       12.8
   y9++      830.44       41.1
   a10++     834.72       21.9
   b10++     848.87       20.7
   y10++     859.58       36.8
   y3+       862.27       27.9
   b11++     898.26       12.1
   y11++     909.05       17.6
   a19+++    918.22       12.9
   y17+++    920.19       24.4
   b19+++    927.51       13.8
   y18+++    958.46       19.0
   y12++     966.37       17.1
   y19+++    996.86       11.3
   b13++    1010.34       12.3
   y14++    1087.49       10.7
   y15++    1151.74       14.5
   y16++    1200.32       10.4
```

```
1049. MORG-339_phosphopeptide_TMT_10%ACN_032017_OT.18820.18820.3 (sSsLEsLQTAVAEVR)

   Ion         m/z   Intensity
   y2+       274.25        7.3
   b1+       317.21        6.8
   y7++      373.21        5.5
   a5++      393.28        6.1
   y3+       403.29       14.4
   y8++      437.28       13.3
   y4+       474.32       24.5
   b6++      491.04        6.3
   y9++      493.60        7.9
   b7++      547.36       10.1
   b3+       571.37        5.1
   y5+       573.42       21.1
   b8++      611.47       21.2
   y6+       644.35       35.8
   a9++      647.59       12.4
   b4+       684.28        5.6
   b10++     697.45       21.5
   y12++     698.36        7.7
   y7+       745.43       22.2
   b11++     746.88       34.4
   b12++     782.45       23.8
   b5+       813.44       13.3
   b13++     847.06       13.7
   y8+       873.47       13.0
   a14++     882.42       12.8
   b14++     896.56       19.4
   b6+       980.37        6.5
   b7+      1093.87        5.2
   b8+      1221.43        8.5
   b10+     1393.53        5.5
```

```
1050. MORG-339_phosphopeptide_TMT_14%ACN_032017_OT.13447.13447.4 (sSSETALSSLAGDDIVkDHEDTcIR)

   Ion         m/z   Intensity
   a10+++    415.20       12.6
   a7++      478.97        9.9
   b2+       484.56       11.4
   b7++      493.58       17.2
   b13+++    506.28        9.5
   b8++      536.73        8.2
   y4+       549.36        8.1
   b3+       571.26        8.1
   y12+++    577.49       11.5
   y14+++    634.72       13.3
   y15+++    658.33        9.4
   y5+       664.33        8.5
   y9++      701.91       10.1
   y17+++    725.13        9.2
   b17+++    734.52        9.5
   a13++     744.09        8.1
   y10++     751.39       18.8
   y18+++    754.47       13.9
   a18+++    762.91       13.2
   b18+++    772.68       12.5
   a5+       773.48       13.4
   y19+++    791.84       20.9
   y6+       793.39       29.9
   b5+       800.94       10.9
   a14++     802.23       15.9
   y21+++    849.47       10.0
   y12++     865.78       12.1
   b15++     872.38       18.1
   y23+++    920.75       14.1
   y13++     923.25       15.9
   a22+++    924.26        9.6
   y7+       930.56       11.9
   y14++     951.77       39.2
   y15++     987.15       39.7
   y16++    1043.74       19.3
   a8+      1044.61       15.9
   y17++    1087.09       17.6
   b17++    1100.98       10.6
   y18++    1130.94       20.0
   a9+      1132.00       14.8
```

```
1051. MORG-339_phosphopeptide_TMT_8%ACN_032017_OT.6125.6125.4 (tLTDEVNsPDsDRRDk)

   Ion         m/z   Intensity
   b5++      395.19        7.1
   y7+++     400.66       10.0
   y4++      402.39       11.3
   y8+++     433.37       17.9
   b6++      444.28        6.4
   y5++      460.01        6.1
   b10+++    461.60        6.2
   a7++      487.55        8.3
   y9+++     489.12        8.4
   a3+       517.18        6.4
   y10+++    527.29       13.5
   y6++      543.69        6.9
   a12+++    545.45        7.3
   y11+++    560.09       10.8
   a13+++    598.26       19.7
   y7++      600.29       17.7
   y12+++    602.61       14.4
   b13+++    607.47       27.1
   y13+++    641.23        6.4
   a14+++    649.46       11.3
   b14+++    659.61        5.8
   b4+       660.43       10.7
   b10++     691.73       10.2
   b15+++    697.73        8.5
   y9++      732.95        7.7
   y10++     789.58       21.8
   b6+       888.49       13.2
   b7+      1002.63        7.8
```

```
1052. MORG-339_phosphopeptide_TMT_4%ACN_032017_OT.7008.7008.2 (tLtDEVNSPDSDR)

   Ion         m/z   Intensity
   y3+       377.26        5.1
   y5+       589.26       13.9
   b5+       869.22        7.4
   b7+      1082.80        5.0
   b10+     1381.57        9.9
   b12+     1583.79        7.3
```

```
1053. MORG-339_phosphopeptide_TMT_18%ACN_032017_OT.18522.18522.4 (lTVEEATASVSEGGGLQGIsmkDSDEEDEEDD)

   Ion         m/z   Intensity
   b10++     615.99       12.3
   b11++     659.70       16.3
   y6+       751.20       12.4
   b13++     752.37       11.5
   b15++     809.34       13.5
   b16++     865.69       13.2
   b6+       872.53       14.6
   y7+       880.57       14.7
   b17++     930.35       14.7
   y13++     935.10       24.6
   b23+++    939.40       18.1
   a18++     944.18       11.5
   a7+       945.07       11.9
   y24+++    951.22       16.7
   b18++     958.32       12.9
   a24+++    959.07       15.8
   a25+++    996.84       12.4
   b25+++   1006.60       13.2
   b19++    1015.28       20.8
   y15++    1020.33       12.5
   y27+++   1033.07       30.5
   b8+      1044.72       22.4
   b26+++   1049.86       25.3
   a20++    1084.59       11.1
   b27+++   1092.86       37.8
   a9+      1103.59       17.8
   y29+++   1119.16       19.7
   b9+      1131.18       28.7
   y18++    1169.02       15.5
   b29+++   1174.14       40.6
   y19++    1197.80       11.2
   b30+++   1217.16       38.0
   y20++    1226.26       24.3
   b10+     1230.57       13.1
   b31+++   1255.55       12.8
   y21++    1290.79       13.0
   b11+     1317.23       11.4
   y22++    1334.23       24.5
   y24++    1427.53       12.8
```

```
1054. MORG-339_phosphopeptide_TMT_6%ACN_032017_OT.11372.11372.3 (tSFDENDsEELEDk)

   Ion         m/z   Intensity
   y4++      367.30        4.2
   y1+       376.24       14.8
   b2+       418.46        4.2
   y5++      431.99        5.4
   y2+       491.31       24.8
   a7++      505.57        5.8
   b7++      520.00        5.2
   b3+       565.37        5.8
   y7++      579.34        4.8
   b8++      603.43        4.8
   y3+       620.39       20.5
   y8++      637.24        4.4
   b9++      668.12        5.6
   b4+       680.44        7.3
   y9++      694.41       19.6
   a10++     718.39       11.3
   a11++     774.70        5.9
   a5+       781.06        5.5
   b11++     789.08       31.3
   b5+       809.43        8.3
   b12++     853.55       27.7
   y5+       862.32       12.7
   a13++     897.32        5.1
   b13++     911.19       10.3
   b6+       923.38        8.0
   b7+      1038.57        7.3
   b9+      1334.47        7.2
   b10+     1463.62       13.9
```

```
1055. MORG-339_phosphopeptide_TMT_4%ACN_032017_OT.9541.9541.2 (aNRADsEEEGAVELAA)

   Ion         m/z   Intensity
   b3+       571.41        6.6
   b5+       757.48       12.4
   y8+       759.52        7.1
   y9+       888.53        6.3
   b6+       923.97        7.4
   b7+      1053.48        5.5
   b8+      1182.53        8.6
   y11+     1184.50        5.6
   b9+      1311.57       11.0
   y13+     1370.84        4.9
```

```
1056. MORG-339_phosphopeptide_TMT_8%ACN_032017_OT.20965.20965.4 (eQEWDSssDGtERLPEEEEIGPFSk)

   Ion         m/z   Intensity
   y4++      354.60       20.8
   y1+       376.24       11.0
   y5++      382.64       13.4
   y6++      439.47       13.7
   y2+       463.42       20.8
   b10+++    504.42        8.7
   y3+       610.41       13.4
   y14+++    630.59        6.2
   y15+++    690.69        6.7
   y4+       707.50       31.7
   b9++      727.32       11.0
   y5+       764.47       26.4
   a4+       774.38        6.9
   b11++     846.49       10.3
   a18+++    849.59       11.1
   y19+++    859.33       10.6
   y6+       877.61       17.3
   y13++     880.30       17.6
   y21+++    926.47       11.9
   b20+++    939.67       19.4
   b23+++   1039.73       10.3
   b14++    1045.56       31.9
   a24+++   1059.64        8.4
   b24+++   1068.70       15.2
   b15++    1093.78       14.3
   b16++    1158.60       12.2
   b17++    1223.24       14.4
   b18++    1287.76       14.0
   y19++    1288.62        9.6
   b19++    1351.86       22.3
   b20++    1408.82       42.0
   b21++    1437.40       36.8
```

```
1057. MORG-339_phosphopeptide_TMT_6%ACN_032017_OT.13857.13857.2 (sYEDLTELEDR)

   Ion         m/z   Intensity
   y4+       532.32        8.1
   y5+       661.36        6.1
   b3+       689.34        6.7
   y6+       762.31        9.0
   b4+       804.44       10.6
   y7+       875.54       12.0
   b5+       917.74       12.2
   y8+       990.54        8.8
   b6+      1018.47        4.5
   y9+      1119.65        6.6
   y10+     1282.61        7.6
   b9+      1389.68        8.4
   b10+     1504.70        5.3
```

```
1058. MORG-339_phosphopeptide_TMT_4%ACN_032017_OT.6467.6467.3 (sEAAAPHAAGGGLssDEEEGTSSQAEAAR)

   Ion         m/z   Intensity
   b1+       317.20        6.9
   b2+       446.43        8.4
   b7++      447.27        7.9
   b8++      482.89       10.6
   b3+       517.20        7.5
   b11++     575.32        6.0
   b4+       588.48       13.0
   a13++     646.69        9.3
   b5+       659.41       11.1
   b13++     660.57       15.4
   a14++     729.37       11.0
   y7+       732.42       10.0
   y8+       819.47       12.5
   b16++     885.20       15.9
   b7+       893.45        9.2
   b17++     949.42       10.6
   y10+      977.57       15.6
   a18++    1000.06       14.8
   a9+      1007.35        9.6
   y19++    1020.63       14.4
   b9+      1035.52        6.3
   b19++    1078.40       10.5
   b10+     1092.25        6.0
   b20++    1106.48       11.6
   y22++    1120.34        7.0
   a11+     1121.09        6.5
   b11+     1150.04       12.0
   b21++    1157.51        6.0
   y23++    1189.05        8.3
   b22++    1200.92        7.2
   y12+     1235.56        9.7
   y24++    1237.76       12.3
   b24++    1308.96        8.0
   b25++    1344.36       12.2
   y13+     1364.88        9.2
   y15+     1646.70        6.2
   y16+     1813.52        6.3
```

```
1059. MORG-339_phosphopeptide_TMT_10%ACN_032017_OT.3636.3636.3 (gAGAGASSVGTGGGAAGGAcGGAGsEDSGDR)

   Ion         m/z   Intensity
   b14++     609.28        9.5
   b6+       614.75        9.8
   b16++     680.04       12.0
   y15++     716.92       12.6
   b18++     736.98        7.4
   y16++     752.66       12.7
   b19++     772.63       12.9
   b8+       788.76       10.7
   y18++     816.38        7.0
   y7+       845.71        8.0
   a9+       859.55       10.0
   y20++     872.82       15.1
   b21++     881.30       14.2
   b9+       887.50       17.7
   a22++     895.57        7.8
   y8+       902.40       15.6
   b22++     909.31       11.7
   b24++     973.28        8.9
   a11+     1017.11        7.6
   b11+     1045.63       10.0
   y11+     1087.50       10.2
   b13+     1159.35        6.6
   y12+     1247.37        7.0
   b15+     1287.76        7.3
   b16+     1358.75       12.9
   b17+     1415.77        7.9
   y15+     1432.51       10.8
```

```
1060. MORG-339_phosphopeptide_TMT_4%ACN_032017_OT.6401.6401.2 (eEEEGISQESsEEEQ)

   Ion         m/z   Intensity
   y3+       405.39        6.1
   b2+       488.41        6.3
   a3+       589.41        6.2
   b3+       617.30        9.4
   b4+       746.30       16.4
   y6+       788.26        7.2
   b5+       803.37        6.6
   a6+       888.57        7.2
   b6+       916.48       41.6
   y7+       917.38       25.7
   a7+       975.53       40.2
   b7+      1003.56       16.9
   y8+      1045.40        7.3
   b8+      1131.49       17.6
   y9+      1132.33       36.5
   b9+      1260.57       15.4
   y11+     1302.39       14.7
   y12+     1431.56       10.8
   y13+     1560.69        7.1
   b12+     1643.72       16.3
   y14+     1689.56        6.0
   b13+     1772.74       17.1
   b14+     1901.66       13.0
```

```
1061. MORG-339_phosphopeptide_TMT_14%ACN_032017_OT.19104.19104.3 (gLVAAYSGDsDNEEELVERLESEEEk)

   Ion         m/z   Intensity
   b2+       400.35        8.4
   b3+       499.47        7.8
   y2+       505.49        6.8
   b4+       570.46        8.3
   y8++      624.82        9.1
   b5+       641.46       10.0
   b12++     730.23        7.0
   y11++     795.62        9.9
   b6+       804.53       13.7
   y5+       850.47       10.8
   b7+       891.51        9.7
   y13++     924.72       11.2
   y15++    1046.36       24.8
   b9+      1063.54       13.4
   y7+      1092.66       12.1
   b18++    1094.61        7.6
   y16++    1103.57       17.9
   y17++    1187.12       13.2
   a20++    1214.63        7.1
   b20++    1228.59        7.2
   y18++    1244.75        6.6
   y8+      1248.64       10.0
   y19++    1273.27        9.8
   b21++    1293.67       13.5
   a11+     1317.58       10.9
   b11+     1345.65       11.4
   a12+     1432.08        7.0
   b13+     1588.62        8.0
```

```
1062. MORG-339_phosphopeptide_TMT_18%ACN_032017_OT.12988.12988.3 (sRSPVDSPVPASMFAPEPSSPGAAR)

   Ion         m/z   Intensity
   y9++      436.41       13.7
   b6++      476.40       31.9
   y11++     520.37       14.1
   a9++      603.80       15.9
   b9++      618.02       28.4
   b11++     702.15       18.2
   y8+       742.41       29.7
   y16++     787.07       28.9
   b13++     810.85       20.7
   b5+       836.73       16.0
   a14++     870.83       29.3
   y9+       871.45       34.2
   y18++     884.54       39.2
   a15++     905.80       14.7
   b15++     920.19       71.6
   a6+       923.49       29.2
   y19++     927.97       14.5
   b6+       951.49       29.2
   y10+      968.49       64.0
   b17++    1033.23       25.5
   y11+     1039.57       31.7
   y22++    1083.82       15.1
   y12+     1186.58       25.9
   b21++    1217.46       12.5
   b9+      1234.57       17.7
   y13+     1317.46       16.5
   y14+     1404.76       13.0
   y16+     1572.77       23.3
   y19+     1855.89       17.4
```

```
1063. MORG-339_phosphopeptide_TMT_18%ACN_032017_OT.11588.11588.4 (rLsEEAcPGVLSVAPTVTQPPGREEQR)

   Ion         m/z   Intensity
   b5++      462.95       19.7
   a6++      483.99       15.6
   a11+++    498.18       29.0
   b2+       499.08       21.5
   y15+++    555.57       14.9
   a7++      564.48       21.0
   b13+++    569.57       15.2
   b7++      578.48       49.3
   y18+++    655.60       30.0
   a10++     691.00       30.2
   b10++     705.09       51.1
   y20+++    707.14       42.4
   a11++     747.11       26.7
   y13++     748.13       68.3
   y21+++    760.14       28.8
   b11++     761.53       49.8
   a4+       767.30       24.4
   b19+++    768.62       14.2
   y6+       774.16       20.8
   y22+++    783.70       63.1
   a12++     790.57       33.0
   a20+++    791.80       30.3
   b4+       794.92      126.0
   b20+++    801.46       30.5
   b12++     805.17       30.3
   y15++     833.25       44.0
   b21+++    834.04       23.4
   a13++     840.81       26.2
   b13++     854.65       16.0
   y16++     876.78       68.0
   b14++     889.86       38.5
   y17++     933.28       73.9
   a24+++    938.04       19.4
   y18++     982.76       24.6
   b6+       995.47       18.9
   y19++    1011.34       18.3
   y20++    1059.87       78.3
   y21++    1139.90       33.3
   b7+      1155.49       29.1
   y23++    1239.90       19.0
```

```
1064. MORG-339_phosphopeptide_TMT_8%ACN_032017_OT.2523.2523.3 (sPPPSPTTQR)

   Ion         m/z   Intensity
   b2++      247.23        6.1
   a3++      282.27       13.5
   b3++      296.29        9.8
   y5++      301.83       22.9
   a4++      330.79        8.1
   y6++      344.92        7.8
   a5++      374.20        7.7
   b5++      388.23       18.1
   y7++      394.16        9.5
   b1+       397.19       16.6
   y3+       404.20       10.3
   b6++      436.34        6.9
   y8++      441.85       17.3
   a7++      473.36        4.7
   b7++      487.44        7.7
   y9++      490.51        6.6
   b2+       494.37        4.9
   y4+       505.46       12.6
   b3+       591.35       14.5
   b9++      601.36        8.5
   y5+       602.34       19.2
   b5+       775.34       17.8
```

```
1065. MORG-339_phosphopeptide_TMT_14%ACN_032017_OT.7030.7030.3 (gTPALTSEAAQSsPPTR)

   Ion         m/z   Intensity
   a7++      415.35        8.7
   y4+       470.28       21.1
   b8++      493.78       10.1
   b9++      529.23       27.3
   b4+       556.58       11.4
   y10++     562.04       16.3
   b10++     564.91       26.3
   y11++     605.72       15.4
   a11++     614.85       11.2
   b11++     629.10       17.6
   y5+       636.85       26.8
   a5+       641.02       29.6
   b12++     672.55       23.8
   y6+       724.39       20.4
   y14++     748.37        8.2
   b13++     756.13        9.5
   b6+       770.42        9.4
   a14++     790.12        9.1
   y7+       852.35       11.4
   a16++     889.04        9.8
   y8+       923.48       13.4
   b9+      1057.56        8.3
   b10+     1128.59       11.7
```

```
1066. MORG-339_phosphopeptide_TMT_18%ACN_032017_OT.12111.12111.3 (sPGTATLALSQQQSLQER)

   Ion         m/z   Intensity
   y2+       304.22       15.2
   b1+       397.24       14.0
   y3+       432.20       28.3
   b8++      504.68       24.2
   y8++      508.92       11.7
   y4+       545.32       18.8
   y9++      552.36       15.4
   b9++      561.57       29.3
   b10++     605.00       22.3
   y10++     609.15       16.3
   y5+       632.40       33.9
   a11++     654.91       14.6
   b11++     668.74       14.0
   y12++     700.51       12.5
   a12++     718.55       21.0
   b5+       723.09       34.8
   b12++     732.59       36.2
   y6+       760.52       26.0
   a6+       796.23       12.0
   b13++     797.19       41.6
   b6+       824.42       16.2
   b14++     840.72       30.0
   y7+       888.33       45.2
   b15++     897.37       16.4
   b7+       937.49       25.2
   a16++     946.87       11.4
   b16++     960.81       17.3
   b8+      1008.44       24.2
   y8+      1016.53       22.1
   b17++    1025.82       25.0
   y9+      1103.59       22.3
   b9+      1121.58       26.6
   b10+     1208.76       12.3
   b11+     1336.69       15.6
```

```
1067. MORG-339_phosphopeptide_TMT_18%ACN_032017_OT.8973.8973.4 (rLSESSALkQPATPtAAEsSEGEGEEGDDGGETESR)

   Ion         m/z   Intensity
   b13+++    610.20       11.5
   b9++      715.79       14.6
   y7+       735.32       11.6
   y15++     762.14        8.8
   a10++     766.45       17.4
   b10++     780.17       30.2
   y8+       850.48       12.1
   b12++     863.93       13.9
   b13++     914.69       17.5
   a14++     948.58        9.4
   y27+++    951.46        9.0
   a23+++    973.35        8.9
   b23+++    983.21       13.6
   a24+++    992.60        9.1
   y19++    1018.41       21.2
   y10+     1022.63       41.7
   a25+++   1035.19       12.4
   a15++    1039.56       19.1
   b25+++   1045.16       13.7
   a8+      1046.10       19.6
   b15++    1053.17       17.1
   y20++    1054.19       15.7
   b28+++   1145.44       14.0
   y11+     1151.16       11.9
   y22++    1179.90       13.1
   y23++    1228.86       10.0
   a19++    1258.93       12.2
   b32+++   1264.87       10.8
   b19++    1272.35        9.1
   y35+++   1299.59        8.9
   y25++    1314.22       14.6
   y13+     1337.55       14.4
   y26++    1363.29       15.0
   y15+     1523.52        8.8
```

```
1068. MORG-339_phosphopeptide_TMT_16%ACN_032017_OT.11791.11791.4 (tAPPVLPTGYDsEEEEESRPMSYDEkR)

   Ion         m/z   Intensity
   b2+       402.27       12.5
   y10+++    499.76       10.6
   y12+++    586.06       11.0
   a15+++    623.26       10.2
   y13+++    629.08       14.2
   y14+++    672.13       10.9
   b5+       695.43       12.8
   a17+++    709.35       10.3
   a18+++    738.14       15.4
   y10++     749.57       18.7
   b12++     755.29        9.7
   y4+       776.22       11.4
   b19+++    799.52       11.8
   b6+       808.49       25.6
   y17+++    809.46        9.2
   y11++     813.73       14.5
   a7+       877.42       48.3
   y12++     878.41       17.8
   y19+++    881.95       21.9
   b14++     883.48       10.0
   a22+++    895.44       81.9
   y20+++    916.48        9.2
   y13++     942.93       13.8
   y22+++    986.24       12.3
   b16++    1012.98       12.1
   y24+++   1051.24        9.5
   y25+++   1084.19       22.2
   y26+++   1107.06       10.5
   a21++    1299.04        9.2
   b11+     1341.73        9.6
   y20++    1373.52       19.7
   b24++    1495.74       12.3
   y23++    1528.12       12.6
```

```
1069. MORG-339_phosphopeptide_TMT_4%ACN_032017_OT.9717.9717.3 (eEEEGISQEssEEEQ)

   Ion         m/z   Intensity
   a2++      230.15        5.8
   y2+       276.11        8.4
   y3+       405.22        7.5
   b6++      459.11       15.3
   b2+       488.28        8.1
   b7++      502.56       12.5
   y8++      563.23        6.0
   y9++      606.93        9.3
   b3+       617.31        9.1
   b9++      630.64        7.2
   y11++     691.44       14.8
   a10++     699.96       16.4
   b4+       746.32       10.5
   b5+       803.48       13.2
   y6+       868.37       10.7
   y14++     884.99        7.5
   b6+       916.53       31.8
   b13++     927.18       22.6
   b14++     991.63       32.8
   y7+       997.38        9.4
   b7+      1003.48       11.1
   b8+      1131.68        9.8
   y9+      1212.37       10.8
   b9+      1260.65        5.9
```

```
1070. MORG-339_phosphopeptide_TMT_22%ACN_032017_OT.15461.15461.3 (dVTNLTVGGFTPmsPR)

   Ion         m/z   Intensity
   y2+       272.30        9.4
   b3++      273.20        8.9
   a1+       317.33       15.0
   a2+       416.12        8.7
   b6++      437.46       15.5
   y7++      466.37        8.8
   b7++      487.01       11.0
   b8++      515.41       27.8
   y9++      523.31       13.6
   b9++      543.91       31.3
   b3+       545.04       10.7
   y10++     572.71       12.1
   y4+       586.36       11.7
   a10++     603.28       12.7
   b10++     617.50       32.7
   y11++     623.46       12.1
   a11++     654.17       19.4
   b4+       659.05       25.3
   y5+       683.28       29.4
   a5+       744.27        7.6
   b5+       772.49       22.5
   y6+       784.37       33.5
   b13++     790.27        7.5
   b6+       873.56       19.7
   y7+       931.43       18.5
   b7+       972.59       22.9
   y8+       988.42       22.3
   b8+      1029.55       16.8
   y9+      1045.42       31.3
   b9+      1086.63       17.8
   y10+     1144.49        8.2
   b11+     1334.76        7.4
```

```
1071. MORG-339_phosphopeptide_TMT_16%ACN_032017_OT.16101.16101.3 (qAAGAELLSGASAGASsAGGIsVEER)

   Ion         m/z   Intensity
   y2+       304.13        7.4
   y3+       433.29       11.4
   b8++      492.64        7.2
   a5+       600.29        8.1
   b12++     643.44        8.5
   y5+       699.30       23.2
   b14++     707.65       11.7
   b15++     743.30       21.2
   b6+       757.41        8.7
   b16++     786.51       12.4
   y6+       812.48        9.2
   y17++     833.05        6.7
   a17++     855.72        7.7
   b17++     869.61       17.6
   b7+       870.53       15.7
   y18++     876.42       14.3
   a18++     891.56       11.2
   y8+       926.51        8.0
   b19++     934.41        6.7
   a20++     948.63        9.6
   b20++     962.27        9.3
   b8+       983.59       15.2
   y9+       997.46        9.2
   a21++    1005.47       10.2
   b21++    1019.19       25.0
   y21++    1054.26       10.4
   b9+      1070.60        9.3
   y22++    1090.38        8.3
   b23++    1152.45        7.3
   y10+     1164.43        8.4
   b11+     1198.72       12.2
   b24++    1216.51        9.3
   y11+     1251.42        7.1
   y13+     1379.57        8.6
```

```
1072. MORG-339_phosphopeptide_TMT_12%ACN_032017_OT.14255.14255.3 (dVEMGNsVIEENEMk)

   Ion         m/z   Intensity
   a1+       317.26       19.7
   y3++      318.19        6.4
   a5++      366.84        9.1
   y1+       376.28       12.4
   a2+       416.17        9.2
   y6++      505.23       12.6
   a7++      507.32       16.8
   y7++      561.26        6.7
   b3+       573.41        9.6
   b9++      628.18       16.8
   y3+       636.41       11.8
   b10++     692.72       14.7
   y9++      694.44        8.9
   a5+       733.42        7.8
   a11++     742.45       11.2
   b6+       875.39        6.8
   b13++     878.79       27.6
   y5+       879.44       15.2
   y13++     910.39        8.3
   y6+      1008.54       11.6
   b8+      1141.63        7.0
   b9+      1254.57       14.7
```

```
1073. MORG-339_phosphopeptide_TMT_10%ACN_032017_OT.12111.12111.4 (sVsPGVTQAVVEEHcAsPEEk)

   Ion         m/z   Intensity
   y3++      317.27       11.1
   y4++      366.30       19.5
   y1+       376.35       12.8
   b6++      418.85        7.8
   y5++      449.86       21.2
   b7++      469.56        9.5
   y2+       505.37       19.3
   y7++      565.24        7.5
   b9++      568.96       26.4
   y13+++    599.09        8.4
   b10++     618.60       19.5
   y3+       634.17       15.2
   y14+++    641.64       13.8
   b16+++    654.18       11.5
   b11++     667.81       16.6
   y15+++    675.09       22.3
   b4+       680.46       11.9
   y9++      698.39       25.1
   a17+++    700.65       19.0
   y16+++    707.59        7.6
   y17+++    726.72        7.9
   y4+       731.40        8.1
   b12++     732.30        9.6
   b5+       737.49        8.2
   b18+++    742.52        9.9
   y10++     762.85       10.1
   b19+++    785.67       12.5
   b13++     796.75        7.9
   y11++     812.60       29.7
   b20+++    828.73       13.1
   b6+       836.40       19.7
   y12++     862.14       22.6
   b14++     865.65       13.5
   y13++     897.71       21.0
   y5+       898.67       12.5
   a7+       909.83        8.8
   b15++     945.76       10.7
   y14++     961.61       11.9
   a16++     966.68        9.2
   b16++     981.26       46.5
   y15++    1012.16        8.9
   b17++    1064.85       12.3
   b8+      1065.48        9.4
```

```
1074. MORG-339_phosphopeptide_TMT_4%ACN_032017_OT.6866.6866.3 (aGEAPTENPAPATEQSsAE)

   Ion         m/z   Intensity
   b7++      443.48        9.3
   y4+       473.16        6.8
   b3+       487.29        5.4
   b8++      500.29       18.6
   a4+       530.52        6.5
   y10++     535.21        9.2
   b9++      548.84       22.9
   b4+       558.39       13.3
   a10++     570.39        8.9
   b10++     584.44       41.7
   b11++     632.65       14.1
   y12++     641.15       36.5
   b12++     668.45       19.4
   a13++     705.12       15.5
   y13++     705.97       12.6
   y6+       730.18        5.6
   b6+       756.30       12.6
   b14++     783.20        9.0
   y7+       831.37       14.6
   b15++     847.53       15.0
   b7+       885.52       19.1
   b16++     891.14       19.3
   y8+       902.33        5.8
   y17++     905.14        6.3
   b8+       999.39       41.5
   b18++    1010.02        7.3
   b10+     1167.52       23.6
   b11+     1264.55        5.9
   b12+     1335.87        7.1
```

```
1075. MORG-339_phosphopeptide_TMT_8%ACN_032017_OT.6263.6263.3 (tASEGSEAETPEAPk)

   Ion         m/z   Intensity
   a4++      335.25        8.0
   y5++      385.97       16.3
   y6++      436.71        9.1
   y2+       473.34       16.1
   b8++      521.60        7.0
   y9++      600.78        8.4
   a10++     622.99       12.9
   b10++     637.05       19.7
   a4+       670.71       18.5
   y11++     672.85       21.3
   y4+       673.61       10.9
   b12++     749.88        6.5
   y5+       770.46       13.6
   a13++     771.58       11.9
   b13++     785.65       17.7
   y6+       871.58        6.9
   b7+       971.49        6.7
   b8+      1042.72        5.9
   b9+      1171.60        8.3
   b10+     1272.51       11.6
```

```
1076. MORG-339_phosphopeptide_TMT_20%ACN_032017_OT.11446.11446.4 (sRkADNPTPQTPGTSSLSQLGAYGDsEDSDS)

   Ion         m/z   Intensity
   b8+++     443.76       16.7
   b5++      508.71       11.2
   a10+++    509.50       10.3
   b10+++    518.73       15.9
   a11+++    543.16       11.1
   b11+++    552.41       27.8
   a14+++    628.50        9.1
   b14+++    637.54       12.0
   b8++      664.83       15.0
   b15+++    666.58       10.3
   b16+++    695.58       14.3
   b17+++    733.41       30.1
   b18+++    762.29       16.5
   b10++     777.32        9.6
   y23+++    793.31       11.6
   b19+++    805.13       28.3
   y15++     812.26       10.1
   b11++     828.11       11.5
   b20+++    842.65       41.8
   a21+++    852.08        9.4
   y16++     855.74       16.3
   b21+++    861.72       22.4
   a12++     862.64       11.0
   a4+       873.39       17.8
   b12++     876.14       13.0
   b22+++    885.40       32.0
   y8+       891.44       23.5
   y26+++    897.70       16.5
   y17++     899.43       19.6
   b4+       901.46       19.7
   b13++     904.73       21.0
   b23+++    939.86       14.7
   b24+++    958.61       14.4
   a15++     985.02       10.9
   a25+++    987.95       13.9
   b25+++    996.52       14.8
   b15++     998.79       10.0
   y20++    1026.49       11.4
   b16++    1042.54        9.9
   y9+      1054.28       17.0
   y21++    1077.04       13.0
   y11+     1182.37       18.9
   y12+     1295.40       14.3
   y14+     1510.56       14.1
```

```
1077. MORG-339_phosphopeptide_TMT_18%ACN_032017_OT.9829.9829.3 (rFSVSPASPSsQQTPPPVtPR)

   Ion         m/z   Intensity
   y2+       272.25        9.7
   a7++      474.28       10.0
   y5+       649.46       10.5
   y6+       746.43       15.0
   b12++     771.03       10.3
   a5+       778.61       10.0
   b5+       806.61       20.8
   y14++     819.62       19.3
   a13++     820.79       11.5
   b13++     835.28       22.7
   y7+       843.42       32.0
   y15++     855.39       27.2
   y16++     904.21       11.9
   b15++     933.61       13.4
   a7+       946.67       13.0
   y17++     947.57       10.1
   a16++     968.83       10.7
   b7+       974.38       15.4
   b17++    1031.16       11.8
   b8+      1061.41       11.0
   b18++    1080.80       14.7
   y20++    1113.83       14.0
```

```
1078. MORG-339_phosphopeptide_TMT_8%ACN_032017_OT.6892.6892.3 (dSGLEDGREsPSFDTPSQR)

   Ion         m/z   Intensity
   y4+       487.21        7.7
   y5+       588.30       18.3
   b9++      595.00        7.8
   y11++     665.31        7.2
   b10++     678.23        9.9
   a11++     712.49       10.8
   y12++     743.78       14.6
   a12++     755.88       10.4
   b12++     769.88       12.9
   y13++     772.28       22.9
   a6+       818.33        7.9
   a13++     829.86        6.9
   b13++     844.04        7.3
   b6+       846.49       11.7
   y7+       850.42        7.1
   b14++     901.67       28.3
   a15++     937.66       13.5
   b15++     951.52        7.1
   y9+      1034.51       19.0
   b8+      1059.55        9.4
   y10+     1201.57        9.5
```

```
1079. MORG-339_phosphopeptide_TMT_20%ACN_032017_OT.16867.16867.3 (lIRDFQEYVEPGEDFPAsPQRR)

   Ion         m/z   Intensity
   y8++      519.74        7.5
   y4+       556.34        6.5
   b8++      648.03        6.9
   y11++     669.90        9.8
   b9++      697.73        6.7
   y12++     718.99        8.6
   b10++     762.05       12.0
   y6+       794.52        6.7
   b5+       874.65        6.4
   y15++     914.74       11.8
   a14++     946.71       11.8
   b14++     960.69       28.1
   a6+       974.30        8.3
   y8+      1038.54       12.5
   y18++    1116.29        7.7
   y9+      1153.21       13.9
   a21++    1379.12        6.7
   b10+     1522.77       13.5
   b14+     1921.19        6.9
```

```
1080. MORG-339_phosphopeptide_TMT_22%ACN_032017_OT.17384.17384.3 (sRLsAIEIDIPVVSHST)

   Ion         m/z   Intensity
   b1+       317.30       10.6
   y3+       344.15        7.6
   y7++      363.67       10.7
   a6++      455.33       10.2
   b6++      469.44        6.8
   y10++     534.00       14.6
   a8++      576.36        7.5
   b8++      590.20       13.4
   b9++      647.93       24.4
   y12++     655.52        9.9
   y13++     690.59       11.5
   b10++     704.42       10.5
   y7+       726.46        8.1
   b11++     752.74        9.4
   b12++     802.87        7.8
   a13++     837.71       12.6
   y8+       839.44       10.5
   b13++     852.33       19.3
   b14++     895.94        8.5
   b6+       937.23        7.5
   y9+       954.42        9.7
   y11+     1196.64       12.3
```

```
1081. MORG-339_phosphopeptide_TMT_4%ACN_032017_OT.16034.16034.3 (vVcDADDsDsDVVsDk)

   Ion         m/z   Intensity
   y3++      329.23        4.6
   y1+       376.48        8.0
   y4++      379.15        6.1
   y5++      428.56        9.9
   y2+       491.34        6.0
   a3+       560.32        5.7
   b8++      585.82        8.0
   a9++      629.46        5.8
   y3+       658.37       13.7
   a4+       675.20        4.1
   y9++      710.94        7.3
   y4+       757.48       11.7
   y10++     768.68        9.7
   b5+       774.23       11.6
   b11++     785.25       10.1
   a12++     820.44       14.8
   b12++     834.17       13.3
   a13++     869.44        7.7
   b13++     883.86       18.5
   b6+       889.48        9.2
   b14++     966.94        4.9
   y6+       971.43        4.8
   b7+      1004.67        8.1
   b15++    1025.22       10.4
   y7+      1138.60        5.6
   b9+      1286.35        5.1
   b10+     1453.46        6.3
   b11+     1568.48        9.5
   b12+     1667.67        7.2
```

```
1082. MORG-339_phosphopeptide_TMT_12%ACN_032017_OT.5368.5368.4 (qkFHDsEGDDTEETEDYR)

   Ion         m/z   Intensity
   y2+       338.18       19.9
   y5++      342.39       11.0
   y3+       453.18       13.4
   b9+++     528.30       14.4
   b5++      558.11        9.4
   b10+++    566.65       16.1
   y4+       582.30       16.1
   b11+++    600.44       17.1
   y15+++    626.08       14.3
   a12+++    634.00       16.6
   y10++     636.59       19.9
   b6++      641.70       16.5
   b12+++    643.58       32.0
   y11++     665.02       43.4
   y16+++    675.55       28.7
   a13+++    676.70       45.5
   b7++      706.27        9.5
   a8++      720.16       15.9
   a15+++    753.87       10.1
   b15+++    763.19       17.5
   a16+++    791.68       16.5
   y17+++    794.41        9.0
   b16+++    801.63       26.1
   y6+       812.48       22.4
   y13++     813.42       14.2
   b10++     849.59       18.7
   y14++     871.06        8.0
   b11++     900.02       12.7
   y7+       941.41       13.8
   b12++     964.64       16.7
   b13++    1029.30       12.5
```

```
1083. MORG-339_phosphopeptide_TMT_8%ACN_032017_OT.17087.17087.3 (gLVAAYsGDsDNEEELVER)

   Ion         m/z   Intensity
   y2+       304.38       12.8
   y3+       403.36        9.9
   a3+       471.41        7.6
   y4+       516.25       11.6
   b4+       570.26        8.0
   a10++     641.47       10.4
   y5+       645.39        7.3
   a11++     698.93        6.8
   y12++     736.29        9.9
   b12++     770.16       14.9
   y6+       774.22       10.1
   a6+       776.00       23.7
   b6+       804.56       14.6
   a13++     820.40        7.6
   b13++     834.87       12.5
   b14++     899.52       13.4
   y14++     901.66        9.5
   y7+       903.02        6.9
   y15++     936.98        9.1
   a7+       943.22        8.4
   b15++     963.72       13.8
   b7+       971.77       15.5
   y16++     972.77        7.8
   a16++    1006.82        9.9
   b16++    1020.64       26.3
   y17++    1021.71        9.7
   b17++    1070.10       18.4
   y9+      1132.79        6.8
   b18++    1134.37       11.4
   b9+      1143.59        8.7
   y10+     1299.59        6.7
   y12+     1471.43        6.1
   b13+     1668.71       10.6
```

```
1084. MORG-339_phosphopeptide_TMT_6%ACN_032017_OT.3163.3163.3 (sRtPPAIR)

   Ion         m/z   Intensity
   y4++      228.73        8.3
   y5++      277.27       64.5
   y2+       288.38        8.1
   a3++      353.67       12.6
   y3+       358.84       17.7
   y6++      367.67       67.0
   a1+       368.76       11.1
   b1+       397.03      182.4
   b4++      416.21        6.9
   y7++      445.77       10.4
   y4+       456.40       26.3
   b6++      500.32       13.1
   y5+       553.32      103.6
   b7++      556.71       13.7
   y6+       734.28       22.3
```

```
1085. MORG-339_phosphopeptide_TMT_12%ACN_032017_OT.12512.12512.2 (sSTsFANIQENAT)

   Ion         m/z   Intensity
   b1+       317.32        6.4
   y5+       562.12       12.9
   y6+       675.31        6.3
   y7+       789.41        9.1
   a5+       791.02       13.9
   b5+       819.61       12.7
   y8+       860.58        6.0
   b6+       890.34       11.3
   b7+      1004.46        9.7
   y9+      1007.47        6.3
   b8+      1117.53       20.8
   y10+     1174.57        7.1
   b9+      1245.64       12.3
   y12+     1362.44        8.6
   b10+     1374.78        6.6
```

```
1086. MORG-339_phosphopeptide_TMT_18%ACN_032017_OT.8950.8950.4 (aRPSQLPEQSSSAQQNGSVSDIsPVQAAk)

   Ion         m/z   Intensity
   y1+       376.32       16.7
   y6++      422.07       15.6
   y10+++    441.86       12.9
   y2+       447.35       22.0
   y7++      505.28       16.0
   y3+       518.33       24.7
   y9++      619.07       18.6
   y16+++    646.40       21.3
   y10++     662.98       17.0
   b17+++    666.13       12.1
   b11++     705.89       13.2
   y19+++    728.21       12.5
   y5+       745.50       14.0
   b12++     749.37       13.3
   b20+++    756.77       12.7
   b13++     785.08       21.8
   a21+++    786.17       12.5
   y21+++    800.27       12.4
   a22+++    824.05       17.5
   b22+++    833.31       28.2
   a14++     834.53       20.7
   y14++     841.52       23.1
   y22+++    842.57       26.7
   b14++     848.56       56.5
   a6+       854.47       23.5
   y24+++    913.10       23.2
   b24+++    921.59       16.7
   b25+++    954.33       18.2
   b16++     970.12       20.9
   y26+++    985.19       16.5
   b26+++    997.02       22.7
   b17++     998.98       12.1
   y17++    1005.47       12.0
   y7+      1009.44       27.4
   b27+++   1020.88       51.2
   a28+++   1034.40       14.7
   b18++    1041.98       15.9
   b28+++   1044.43       24.5
   y19++    1091.78       35.9
   y8+      1122.44       12.2
   y20++    1135.29       18.1
   b21++    1192.88       32.3
   b22++    1249.38       43.8
   y10+     1324.66       13.0
   b25++    1430.97       24.8
```

```
1087. MORG-339_phosphopeptide_TMT_4%ACN_032017_OT.2600.2600.3 (aPDRtPPSEEDSAEAER)

   Ion         m/z   Intensity
   b1+       301.28        4.6
   a5++      411.44        5.2
   b5++      425.67       17.3
   b6++      473.87        4.6
   b3+       513.31       11.5
   b8++      566.41        5.9
   y11++     610.35        6.3
   b9++      631.25        6.9
   y12++     658.96       34.8
   y6+       662.21       12.7
   b4+       669.11        4.5
   a10++     681.36       11.6
   b10++     695.22       11.9
   b11++     752.60       24.5
   y7+       777.46        6.3
   y14++     827.73       11.8
   b5+       850.39        6.7
   y15++     884.72        8.4
   b14++     896.57        6.3
   y8+       906.24        5.6
   y16++     933.54        7.1
   b6+       947.94        5.4
   a16++     982.48        8.6
   b16++     996.65        5.0
   y9+      1035.20        4.8
   y11+     1219.56        5.5
   y12+     1316.60       23.3
```

```
1088. MORG-339_phosphopeptide_TMT_16%ACN_032017_OT.19730.19730.3 (gGNVFEALIQDDsEEEEEEEENR)

   Ion         m/z   Intensity
   b6++      417.38       13.8
   y3+       418.25       22.0
   b7++      452.65       16.8
   b3+       458.38       10.9
   b8++      509.59       22.9
   a4+       529.29       13.7
   y4+       547.29       29.8
   b4+       557.33       23.4
   b9++      565.80       18.9
   a5+       676.30       40.5
   b11++     687.31       13.2
   b5+       704.45       20.5
   a6+       805.36       36.7
   b6+       833.47       28.5
   y13++     860.03       17.0
   b7+       904.50       37.0
   y14++     924.04       22.8
   y7+       934.40       36.0
   b15++     957.61       18.8
   y15++     980.28       13.7
   b8+      1017.52       46.2
   b16++    1022.22       12.4
   y16++    1037.24       12.6
   y8+      1063.41       33.9
   a17++    1072.71       10.8
   b17++    1086.76       17.9
   a9+      1102.58       12.1
   b9+      1130.64       63.7
   a18++    1137.44       11.1
   b18++    1151.32       18.3
   y9+      1192.51       21.5
   y19++    1210.42       11.1
   b19++    1215.94       19.2
   b10+     1258.63       37.3
   y20++    1259.68       33.3
   b20++    1280.33       18.5
   y22++    1345.07       14.7
   b11+     1373.84       14.1
   b12+     1488.62       15.9
   y12+     1603.60       11.5
   y13+     1718.63       16.4
```

```
1089. MORG-339_phosphopeptide_TMT_16%ACN_032017_OT.10760.10760.4 (tNNLQLHPHDAFNDEDADRLsEISAR)

   Ion         m/z   Intensity
   b6++      457.31       15.4
   y8++      506.54       18.7
   b7++      525.89       21.8
   y5+       575.37       16.2
   y10++     599.65       16.4
   b4+       672.51       16.8
   b10++     700.58       19.8
   y18+++    714.04       26.6
   a11++     721.91       20.8
   b11++     735.89       12.7
   y6+       742.43       15.1
   y19+++    746.78       23.0
   a18+++    750.16       11.5
   b18+++    759.22       21.5
   y20+++    791.95       29.2
   a12++     795.85       18.6
   b5+       800.62       24.7
   b12++     809.36       18.5
   b19+++    811.65       25.1
   a20+++    839.65       15.7
   b20+++    849.64       16.2
   y22+++    872.72       33.6
   a6+       885.30       14.2
   a21+++    896.20       19.8
   a14++     909.53       15.7
   b6+       913.65       26.2
   b14++     924.20       31.8
   y16++     944.54       20.0
   y24+++    948.21       17.5
   a15++     974.54       15.6
   a23+++    976.82       17.3
   y25+++    986.86       13.4
   b15++     988.43       13.4
   y17++    1002.92       17.8
   y8+      1011.13       12.8
   a7+      1022.21       14.6
   b25+++   1038.64       15.1
   b16++    1046.26       30.6
   b7+      1050.65       20.3
   y18++    1070.71       42.5
   a8+      1119.63       24.7
   b18++    1139.39       20.8
```

```
1090. MORG-339_phosphopeptide_TMT_14%ACN_032017_OT.9046.9046.4 (dLEGLHRAEQsLHDLQER)

   Ion         m/z   Intensity
   y4++      273.29        9.3
   a1+       317.19       14.0
   y6++      399.34       17.0
   y3+       432.27        7.7
   y7++      455.80       16.0
   a7++      512.24        8.9
   y12+++    521.19       10.8
   y8++      539.15        8.6
   y4+       545.40       16.3
   a8++      547.62       19.0
   b12+++    553.73        9.0
   b8++      561.53        9.5
   y13+++    566.61       11.5
   b3+       587.62       15.3
   a13+++    590.25      123.9
   b13+++    599.78       14.9
   y9++      603.77       20.4
   y14+++    604.94       35.3
   y15+++    623.80        6.9
   b9++      625.90       10.6
   b14+++    637.74       12.9
   y5+       660.39       14.0
   y16+++    666.19        9.1
   a10++     675.76        7.7
   b10++     689.88        8.5
   y11++     703.75        6.7
   a16+++    708.48       12.4
   b16+++    718.02        7.9
   b5+       757.92        7.1
   b17+++    761.06        8.4
   y12++     781.16       19.8
   y6+       797.65        8.7
   b6+       894.62       10.1
   b14++     955.98        6.6
```

```
1091. MORG-339_phosphopeptide_TMT_8%ACN_032017_OT.20679.20679.3 (iVEPEVVGEsDsEVEGDAWR)

   Ion         m/z   Intensity
   y2+       361.15        9.7
   y3+       432.28       13.0
   b2+       442.26        6.6
   a8++      513.27        6.3
   y4+       547.38        5.6
   b3+       571.40       17.2
   y5+       604.37       22.0
   b4+       668.41        8.4
   b10++     675.05        8.1
   y6+       733.31       26.5
   b5+       797.60       13.8
   y13++     799.00       25.3
   y7+       832.08       66.6
   y14++     848.38       17.7
   b13++     880.61       11.5
   b6+       896.56       22.4
   y15++     897.75       12.1
   b14++     930.31       17.6
   y8+       961.24        9.1
   a7+       967.37       10.7
   b15++     994.59       15.8
   b7+       995.60       27.2
   b8+      1052.63        9.1
   b17++    1080.90       15.9
   b18++    1116.10       14.8
   y9+      1128.49        7.8
   b9+      1181.71       14.9
   a19++    1195.45        7.9
   b19++    1209.38       15.8
   y11+     1410.48        6.7
   y12+     1539.45        6.2
   y13+     1596.67        8.4
   b14+     1858.75        5.8
```

```
1092. MORG-339_phosphopeptide_TMT_16%ACN_032017_OT.14542.14542.3 (sLskSDSDLLTcsPTEDATMGSR)

   Ion         m/z   Intensity
   b1+       317.28       10.4
   b7++      622.54        7.9
   b8++      679.86        7.8
   b9++      736.33        7.6
   a10++     778.88        8.0
   a12++     909.74       13.8
   b12++     923.15        8.2
   b4+       954.89       18.1
   y17++     960.83       20.3
   y9+       967.19       12.8
   b13++    1007.45       29.3
   y10+     1064.49       11.8
   a7+      1215.48        9.7
   y20++    1240.50        8.4
   b18++    1263.62        8.8
   a8+      1331.05        8.0
   y12+     1391.46        7.7
   y13+     1492.38        9.1
   y15+     1718.72        7.1
```

```
1093. MORG-339_phosphopeptide_TMT_8%ACN_032017_OT.3428.3428.3 (nSGATADAGSIsPR)

   Ion         m/z   Intensity
   y2+       272.23        7.3
   a1+       316.16        5.7
   b5++      330.92        8.6
   b6++      366.13       10.2
   y7++      384.18        8.1
   b7++      423.93       10.1
   b2+       431.72       15.5
   y3+       439.28       18.1
   y8++      441.20        5.2
   b8++      459.25       25.8
   a3+       460.05        6.2
   a9++      473.45        9.4
   b3+       487.90       22.3
   a10++     517.27       12.8
   y10++     527.48       11.1
   a4+       531.02       15.7
   y4+       552.35       15.6
   b4+       559.14       11.6
   b11++     587.81       11.0
   y5+       639.32       13.6
   b5+       660.51       10.6
   b12++     671.38       11.1
   y6+       696.38       22.0
   b6+       731.43        9.6
   b7+       846.35       13.0
   y8+       882.50        9.3
```

```
1094. MORG-339_phosphopeptide_TMT_12%ACN_032017_OT.10759.10759.3 (sHsSPsLNPDAsPVTAk)

   Ion         m/z   Intensity
   b1+       317.25        7.7
   y2+       447.42        6.2
   y3+       548.57       12.1
   y8++      549.47        6.2
   y9++      597.74       11.4
   b8++      600.39        9.9
   y4+       647.62        6.3
   y10++     654.99       12.0
   b10++     706.34        8.8
   y11++     711.68        6.7
   b11++     741.88        9.2
   y5+       744.50       16.0
   a5+       777.40       11.9
   b12++     825.66        7.1
   a13++     859.95        8.7
   a14++     909.10        8.8
   y6+       911.65        7.8
   b6+       972.05        7.9
   y7+       982.55        8.9
   y16++    1039.04        7.0
   b7+      1085.53        7.2
   y8+      1097.62        6.2
   b8+      1199.49       10.0
```

```
1095. MORG-339_phosphopeptide_TMT_20%ACN_032017_OT.20549.20549.2 (sIEMMDIVVEk)

   Ion         m/z   Intensity
   y2+       505.38        4.3
   y3+       604.60        5.8
   b3+       639.32        4.5
   y4+       703.47        7.4
   b4+       770.39        5.3
   y5+       816.61        8.0
   a5+       873.75        8.0
   b5+       901.09        9.1
   y6+       931.52        6.7
   b6+      1016.46        6.9
   y7+      1062.77        5.2
   a7+      1101.61        6.7
   y8+      1193.65        9.2
   b8+      1228.55        6.7
   y9+      1322.84        5.7
   y10+     1435.79        4.9
```

```
1096. MORG-339_phosphopeptide_TMT_18%ACN_032017_OT.15851.15851.4 (rNLLEDDsDEEEDFFLRGPSGPR)

   Ion         m/z   Intensity
   y4+       416.13       12.5
   b6++      486.00       24.7
   b7++      543.44       24.3
   y10++     567.57       37.7
   a13+++    614.36        9.8
   y11++     625.21       14.2
   b8++      627.23       13.3
   b9++      684.39       23.1
   y12++     689.61       20.5
   a15+++    712.33       18.4
   y18+++    716.30       12.4
   b15+++    722.40       30.7
   a10++     734.99       16.4
   a11++     798.85       13.3
   a17+++    802.48        8.6
   b11++     813.38       16.4
   y14++     818.63       33.2
   b18+++    830.97       10.8
   y21+++    835.20       11.3
   y15++     876.08       25.1
   b12++     877.98       15.0
   a21+++    901.79       12.6
   b21+++    910.65       22.4
   b13++     935.57       22.5
   y16++     959.36       17.0
   y17++    1017.36       11.9
   b7+      1085.54        9.8
```

```
1097. MORG-339_phosphopeptide_TMT_10%ACN_032017_OT.11331.11331.3 (aAVLsDsEDDAGNASAk)

   Ion         m/z   Intensity
   y1+       376.38       16.9
   y6++      388.79        7.2
   y2+       447.35       14.5
   y3+       534.37       21.6
   y4+       605.49       13.9
   y11++     687.38        9.6
   b10++     697.01       13.0
   a11++     717.90       16.2
   y5+       719.49       11.0
   a5+       723.62       14.2
   b11++     731.84       24.8
   b12++     760.87        7.4
   y6+       776.51       14.6
   a13++     803.37        7.4
   a14++     838.88       16.0
   y7+       847.80       14.2
   b14++     853.50       45.5
   b6+       866.43       10.7
   a15++     883.25        7.0
   b15++     896.71       16.2
   b16++     932.17        7.6
   y16++     970.26        7.1
   b10+     1392.43        8.4
   b11+     1463.79        9.3
```

```
1098. MORG-339_phosphopeptide_TMT_4%ACN_032017_OT.12505.12505.4 (dLGHPVEEEDEsGDQEDDDDEIDDGDkDQDI)

   Ion         m/z   Intensity
   y7++      510.58       11.5
   b9++      618.31       17.1
   y9++      625.37       20.1
   b10++     675.97       14.7
   y10++     682.04       20.5
   b11++     740.60       17.4
   y18+++    775.19       10.5
   b19+++    807.85       14.3
   y5+       847.61       21.1
   b6+       848.46       10.6
   y13++     861.01        9.9
   b21+++    889.27       25.0
   b14++     910.11       26.4
   a22+++    917.31       17.0
   y14++     919.14       10.9
   b22+++    927.04       28.8
   y6+       962.49       15.0
   b23+++    964.79       11.9
   y23+++    974.08       25.1
   y15++     976.79       16.3
   b7+       977.57       12.7
   y7+      1019.58       14.1
   b16++    1038.71       32.0
   b26+++   1060.86       10.3
   b17++    1096.18       20.8
   y17++    1104.80       10.5
   y8+      1134.59       16.8
   b18++    1153.21       15.7
   b27+++   1180.14       60.4
   b19++    1210.74       13.0
   b28+++   1218.49       44.6
   y9+      1249.67       27.6
   b29+++   1261.07       46.0
   b20++    1268.70       27.1
   y20++    1274.82       13.8
   b30+++   1299.54       33.9
   b21++    1333.26       52.7
   y10+     1362.71       13.2
   b22++    1389.72       39.5
   y22++    1396.58        9.8
   b23++    1447.53       25.7
   b24++    1505.03       20.6
```

```
1099. MORG-339_phosphopeptide_TMT_4%ACN_032017_OT.15434.15434.4 (atEEEDSDEDAVLAPSGVTGAGTGDEGDGQAPGST)

   Ion         m/z   Intensity
   y4+       361.17       35.6
   y5+       432.31       18.8
   b9++      658.02       20.5
   y15++     660.39       11.6
   a17+++    666.82       28.4
   b10++     715.94       32.0
   b19+++    742.52       20.2
   b11++     751.54       83.0
   a20+++    752.45       40.3
   a12++     787.08       20.2
   y9+       789.35       26.6
   b12++     801.01      103.1
   a13++     843.71       18.3
   y28+++    844.57       21.8
   b13++     857.60      107.0
   a25+++    885.72       20.1
   y21++     909.44       16.2
   y10+      918.24       12.1
   a15++     927.14       26.3
   y22++     945.08       27.4
   b16++     984.50       23.4
   y23++    1001.75       39.1
   y11+     1033.26       17.9
   a7+      1043.34       16.5
   a18++    1049.16       14.4
   y24++    1050.92       12.9
   b30+++   1057.75       20.3
   b18++    1063.16       13.2
   y25++    1086.40       14.9
   y12+     1090.42       18.5
   b19++    1113.69       11.9
   b21++    1177.86       12.5
   b8+      1186.42       13.6
   y14+     1248.49       18.2
   y31++    1431.65       15.0
   b11+     1501.56       20.5
   a30++    1571.66       13.9
   b12+     1600.64       15.7
   b13+     1713.76       15.1
   b34++    1741.73       12.8
   b14+     1784.78       20.2
```

```
1100. MORG-339_phosphopeptide_TMT_30%ACN_032017_OT.16093.16093.3 (iVETINsDsDSEFGIPkkTTTPk)

   Ion         m/z   Intensity
   b5+       785.76       11.9
   y8++      794.75       24.2
   y10++     880.01        8.8
   y11++     953.32       13.0
   y13++    1061.36       28.8
   y14++    1119.12       17.2
   y6+      1133.41       12.2
   a8+      1153.93       18.9
   b8+      1181.28       16.6
   y15++    1202.12       10.7
   a9+      1320.43       11.7
   y17++    1343.47        8.4
   b10+     1463.91       11.1
   a20++    1491.69        9.0
   b20++    1505.95       10.6
   a21++    1542.15        8.5
   y21++    1571.42        7.6
   b12+     1679.24       13.6
```

```
1101. MORG-339_phosphopeptide_TMT_14%ACN_032017_OT.12234.12234.4 (sEPDRGRLTPSPDIIVLSDNEAsSPR)

   Ion         m/z   Intensity
   b1+       317.27       15.0
   y14+++    532.29        8.5
   b13+++    546.85       13.9
   b14+++    584.59       11.6
   y5+       597.15       10.8
   b15+++    622.19       17.4
   y11++     627.68       10.1
   a16+++    646.03       10.8
   b16+++    655.22       32.5
   b17+++    692.93       31.7
   b11++     712.99       13.9
   b18+++    721.74       22.1
   y6+       726.01       10.4
   y13++     740.70       14.6
   a12++     747.66       20.7
   y20+++    749.56       18.1
   a19+++    750.45       18.7
   b19+++    760.53       32.1
   y21+++    768.46       21.3
   y14++     798.66       13.9
   a21+++    832.21        7.7
   y7+       840.36       11.6
   b21+++    841.47       26.4
   a14++     861.93        7.8
   b22+++    865.26       19.7
   b14++     876.23       10.0
   y24+++    891.62       14.2
   a23+++    911.22       18.9
   a15++     918.16        9.2
   b23+++    920.51        8.6
   b15++     932.38       13.2
   y25+++    934.36        8.5
   y8+       955.39       12.8
   a16++     967.91        8.1
   b16++     982.25       19.6
   b17++    1039.07       14.6
   y9+      1042.36       19.2
   y10+     1155.42       13.7
   y11+     1254.55        7.7
```

```
1102. MORG-339_phosphopeptide_TMT_4%ACN_032017_OT.22579.22579.2 (iVsAQsLAEDDVE)

   Ion         m/z   Intensity
   b1+       343.40       10.0
   y3+       362.25        9.0
   b2+       442.36       26.5
   y4+       477.19       15.2
   y5+       606.34       25.2
   y6+       677.36       38.4
   b4+       680.42       11.0
   a5+       780.71        7.2
   y7+       790.47       16.2
   b5+       808.53       42.4
   y8+       957.36       20.7
   b6+       975.73        8.3
   y9+      1085.48       13.8
   b7+      1088.47       29.8
   b8+      1159.60       62.0
   b9+      1288.61       48.5
   y11+     1323.60       18.7
   b10+     1403.64       29.8
   y12+     1422.32       12.7
   b11+     1518.67       40.3
```

```
1103. MORG-339_phosphopeptide_TMT_18%ACN_032017_OT.3656.3656.4 (qGPVSHSATQQPVTADkQQSHDPVsPR)

   Ion         m/z   Intensity
   y2+       272.16       11.8
   b7++      461.90       14.6
   b8++      497.45       12.8
   y4+       538.29       10.9
   b9++      547.86        9.1
   a16+++    602.93       13.8
   b16+++    611.63       19.1
   y5+       635.26       10.7
   b11++     675.95       31.3
   y16+++    691.01        9.3
   b5+       698.37       11.4
   y18+++    776.83       11.0
   y11++     794.30       20.3
   a19+++    806.99       12.7
   a6+       807.64       14.8
   a14++     810.24       16.5
   b19+++    816.10       16.7
   b14++     824.47       17.2
   b6+       835.59       24.3
   a20+++    836.50       25.6
   a15++     846.00        8.6
   y22+++    908.65       15.9
   b16++     917.79       14.0
   b22+++    929.85       39.3
   y23+++    937.34       13.9
   y14++     938.28        9.2
   b23+++    961.81       12.4
   y15++     987.16       31.1
   b8+       993.63        9.7
   y16++    1036.23       40.4
   b9+      1094.59        8.6
   y17++    1099.69       10.1
   y19++    1215.02       12.7
   y20++    1250.33       10.5
   y21++    1294.09        8.3
   b11+     1350.80       12.7
```

```
1104. MORG-339_phosphopeptide_TMT_6%ACN_032017_OT.9072.9072.3 (sAGEDNDGPVLTDEQk)

   Ion         m/z   Intensity
   y3++      317.35        6.7
   y1+       376.28       13.4
   b1+       397.19        6.9
   b2+       468.16        6.5
   y6++      481.75       10.7
   y2+       504.35       15.6
   y8++      580.05        8.4
   a10++     612.14       10.9
   a4+       626.49       25.8
   y3+       633.33        6.9
   b4+       654.18        7.5
   b11++     683.08       30.5
   a12++     719.43       14.9
   y4+       748.40       13.6
   b5+       769.42        7.2
   y13++     844.86        7.6
   y5+       849.49       21.1
   b14++     855.57       14.2
   b15++     919.29       13.3
   y6+       962.68       13.5
   b7+       998.49        7.8
   b8+      1055.39        6.2
   y7+      1061.12        9.2
   b10+     1251.58       15.9
```

```
1105. MORG-339_phosphopeptide_TMT_10%ACN_032017_OT.4379.4379.4 (nYQQNYQNSEsGEkNEGSESAPEGQAQQR)

   Ion         m/z   Intensity
   y13+++    448.55        8.6
   y8++      457.55       23.9
   y4+       502.33       11.1
   a6++      506.66        9.5
   y11++     601.27       11.6
   b3+       635.33        9.0
   y6+       687.44       19.7
   a14+++    727.30        8.9
   b10++     749.86        8.8
   b4+       763.12       11.3
   y19+++    767.15        9.9
   y7+       816.52       15.1
   b17+++    836.93       11.3
   a18+++    856.53       14.9
   b12++     861.44       15.4
   b5+       877.15       24.4
   a19+++    899.44       16.3
   b19+++    909.29       15.3
   y8+       913.47       46.4
   y23+++    919.39       14.5
   b13++     926.70       20.8
   a20+++    928.66       20.6
   b20+++    938.00       20.7
   a22+++    984.58       13.3
   y25+++   1011.87        8.9
   b6+      1040.56       14.2
   y10+     1071.35       11.3
   b14++    1105.26       18.0
   b26+++   1122.91       17.2
   a7+      1140.09       10.8
   y28+++   1151.44       11.8
   b15++    1162.31        9.9
   b7+      1168.69       11.8
   b28+++   1207.96        9.2
   b16++    1226.87       23.4
   y21++    1257.78        9.0
   b8+      1282.67       11.0
   b19++    1362.92        8.7
   b20++    1406.63       10.0
   b21++    1442.42       31.1
```

```
1106. MORG-339_phosphopeptide_TMT_18%ACN_032017_OT.23111.23111.2 (sPNELVDDLFk)

   Ion         m/z   Intensity
   b1+       397.26        8.5
   y3+       636.51        9.7
   b4+       737.48        8.0
   y4+       751.59       10.0
   b5+       850.66       15.0
   y5+       866.72       11.1
   b6+       949.66        5.3
   y6+       965.67        6.5
   b7+      1064.60        9.2
   y7+      1078.62        7.5
   b8+      1179.52        7.7
   y10+     1418.89       12.7
```

```
1107. MORG-339_phosphopeptide_TMT_4%ACN_032017_OT.12351.12351.3 (sDIsERPSVDDVESEtGsTGALETR)

   Ion         m/z   Intensity
   y3+       405.17        6.2
   b3+       545.38        9.2
   y6+       646.49        7.4
   b10++     698.60       15.9
   y7+       747.43        7.8
   b11++     755.98       11.0
   a12++     791.79        6.7
   b12++     805.76        6.8
   b13++     870.10       17.3
   y9+       971.47       33.2
   b15++     978.01        5.7
   b6+       997.40       18.5
   y18++    1007.15       10.3
   y20++    1133.33        6.4
   y10+     1151.98        9.1
   a8+      1153.67        7.6
   b18++    1180.08        6.9
   y21++    1197.51       12.5
   a9+      1252.38        5.6
   b21++    1295.14        6.4
   b22++    1351.77        9.5
   y12+     1368.37        7.3
   a11+     1482.69        6.8
   y15+     1711.68        6.3
```

```
1108. MORG-339_phosphopeptide_TMT_8%ACN_032017_OT.11311.11311.3 (rSAsPDDDLGSSNWEAADLGNEER)

   Ion         m/z   Intensity
   b6++      462.22       15.4
   b7++      519.75       20.4
   b8++      577.39       22.1
   y5+       604.36       19.2
   b9++      633.80       12.3
   b10++     662.45       11.6
   b4+       711.40       11.0
   y6+       717.38       35.9
   b13++     806.56       19.7
   y7+       832.42       26.4
   a14++     885.51       16.9
   b14++     899.60       24.8
   y8+       903.46       25.5
   b6+       923.46       23.5
   y17++     932.07       10.4
   a15++     949.43       29.0
   y9+       974.45       20.6
   b16++     999.56       21.6
   b17++    1034.84       22.2
   b7+      1038.49       15.7
   b18++    1092.60       35.7
   y20++    1095.63       29.4
   y10+     1103.50       26.9
   b19++    1149.17       24.3
   b8+      1153.50       19.4
   b20++    1177.69       15.1
   y11+     1289.64       22.3
   b22++    1299.27       17.3
   b23++    1363.75       16.7
   y12+     1403.70       13.3
   y13+     1490.70       13.0
   y14+     1577.59       12.9
   y15+     1634.73       17.5
   y16+     1747.78       20.7
   y17+     1862.74       18.9
   y18+     1977.76       12.8
```

```
1109. MORG-339_phosphopeptide_TMT_20%ACN_032017_OT.14706.14706.3 (rENQVVELHYNAIcQYMGPGDsDED)

   Ion         m/z   Intensity
   b5++      428.90        9.1
   b6++      478.36        9.6
   b7++      542.79       18.7
   b8++      599.45       16.1
   b3+       629.41        8.1
   b9++      667.95       19.3
   y6+       717.37        7.6
   b11++     806.64       21.0
   y7+       814.28       11.7
   b12++     842.20       21.7
   y8+       871.35       20.6
   b13++     898.70       31.8
   b6+       955.95       10.5
   a14++     964.78       12.7
   y9+      1002.37       11.3
   a15++    1028.80       10.8
   b15++    1042.77       25.6
   a7+      1056.74       19.7
   b16++    1123.90       15.7
   y10+     1165.38       11.6
   b17++    1189.92       45.9
   b18++    1218.29       29.2
   b21++    1353.14       20.3
   b22++    1435.68        7.6
   y12+     1453.44       14.9
   y13+     1566.63        8.2
   y15+     1751.74        9.9
```

```
1110. MORG-339_phosphopeptide_TMT_4%ACN_032017_OT.9131.9131.3 (aERGGssGEELEDEEPVk)

   Ion         m/z   Intensity
   y1+       376.34       10.5
   a6++      419.91        6.9
   y3+       572.40       26.6
   b3+       586.39        7.0
   b9++      610.98        8.8
   b10++     675.41       12.8
   y4+       701.49       11.6
   b11++     732.11       13.9
   a12++     782.55        9.0
   b12++     796.26       14.6
   y5+       830.44       19.7
   y12++     834.95       11.6
   b13++     853.75        6.9
   y13++     918.54       16.6
   y6+       945.50       10.5
   a15++     968.81        8.1
   y15++     975.55        7.5
   b15++     983.10       40.9
   b16++    1031.77        8.8
   y16++    1054.43        7.1
   y7+      1074.61       10.5
   b17++    1081.15       15.9
   y8+      1187.63        9.4
```

```
1111. MORG-339_phosphopeptide_TMT_16%ACN_032017_OT.21051.21051.2 (cQTLFGFDsDDESA)

   Ion         m/z   Intensity
   y7+       818.28       16.0
   b6+       879.46       27.1
   y8+       965.43        5.8
   y9+      1022.32       20.1
   y10+     1169.38       20.7
   a9+      1280.32        6.1
   y11+     1282.45       10.3
   y12+     1383.56        6.9
```

```
1112. MORG-339_phosphopeptide_TMT_4%ACN_032017_OT.10365.10365.2 (dIISSDGDsS)

   Ion         m/z   Intensity
   a1+       317.23        5.0
   b1+       345.42        3.4
   y3+       388.20        5.0
   a2+       430.34        2.9
   y4+       445.18        6.0
   y5+       560.28        7.5
   b3+       571.52       15.2
   a4+       630.21        4.9
   y7+       734.24       13.5
   y8+       847.28        3.9
   b6+       860.56        9.1
   b7+       917.64        3.3
   y9+       960.38        2.4
   b8+      1032.73        4.5
```

```
1113. MORG-339_phosphopeptide_TMT_20%ACN_032017_OT.16868.16868.3 (tRTPAsINATPANINLADLTR)

   Ion         m/z   Intensity
   a5++      404.86        9.4
   b5++      419.01       10.9
   b2+       566.94       11.4
   y11++     599.67        8.6
   b8++      615.55       11.9
   b9++      651.55       14.1
   b10++     702.13        8.1
   a4+       736.96        8.3
   b4+       765.47       10.3
   y7+       802.09       19.1
   a5+       808.62        9.1
   a13++     828.80       39.0
   b5+       836.56       20.1
   b13++     842.41       16.6
   b14++     899.74        9.3
   y8+       915.62       13.0
   y17++     918.18       10.5
   b15++     956.48        8.0
   y18++     966.45        7.8
   a6+       975.90        8.2
   a16++     999.09       13.3
   y19++    1016.83       10.2
   y9+      1029.79       16.8
   a18++    1091.55        7.6
   y11+     1197.73       28.0
   a20++    1198.64       24.0
   y12+     1298.91        9.5
```

```
1114. MORG-339_phosphopeptide_TMT_10%ACN_032017_OT.20231.20231.3 (ekEPAEGNEsSEAPsPVNAMNSFV)

   Ion         m/z   Intensity
   b4++      471.76        6.7
   b8++      657.55        6.1
   y7+       782.20        8.5
   b12++     913.42       21.2
   a4+       914.16       17.6
   b13++     949.17       37.1
   y9+       978.48        9.0
   a5+       985.50       15.4
   b14++     997.82       20.7
   y18++     998.80        8.7
   b17++    1179.23       18.6
   b18++    1235.99       10.8
   y11+     1242.61       20.3
   b19++    1271.77        8.4
   a8+      1285.88        6.5
   b8+      1313.62       12.1
   a21++    1380.33       14.5
   a22++    1423.87       13.5
```

```
1115. MORG-339_phosphopeptide_TMT_20%ACN_032017_OT.11704.11704.3 (aAGALGSASsGSMPNLAAR)

   Ion         m/z   Intensity
   a1+       273.24        6.7
   y3+       317.24       15.2
   b6++      335.51        9.0
   b3+       429.16        9.3
   y9++      458.56        7.3
   y11++     585.49        6.9
   a12++     599.59       12.3
   b12++     613.59       10.5
   y12++     621.66       12.8
   y6+       641.47       20.5
   a6+       642.39       21.5
   b14++     727.71        6.7
   b7+       757.30        6.5
   y7+       772.55        6.6
   a8+       800.18       11.9
   y17++     814.31        7.4
   b8+       828.37       10.2
   b16++     841.71       18.3
   b18++     912.75        8.4
```

```
1116. MORG-339_phosphopeptide_TMT_6%ACN_032017_OT.7920.7920.3 (sRTsVQtEDDQLIAGQSAR)

   Ion         m/z   Intensity
   b1+       317.23        5.6
   b6++      484.95        8.7
   y5+       518.28        7.4
   y6+       589.44       13.4
   b9++      697.54       11.0
   y7+       702.50       10.3
   b4+       741.14        5.9
   y13++     742.41        6.2
   b10++     754.96       11.1
   a11++     805.27       12.3
   y14++     806.11       13.0
   b5+       840.43        5.7
   b12++     875.56       17.2
   a13++     917.77        6.8
   b13++     931.81        8.9
   y9+       943.57        8.3
   b14++     967.88       10.8
   y10+     1058.51        9.1
   b16++    1059.52        6.0
   y11+     1173.61        6.3
```

```
1117. MORG-339_phosphopeptide_TMT_14%ACN_032017_OT.16279.16279.4 (sAEPLANTVLAsETEEEGNAQALGPTAk)

   Ion         m/z   Intensity
   b1+       317.24       11.0
   y1+       376.32       10.3
   y6++      408.29        8.3
   y2+       447.41        7.2
   y8++      507.36        7.8
   b3+       517.31        7.3
   y3+       547.96       13.0
   b9++      556.98        8.8
   y15+++    582.02        9.7
   b10++     613.40        9.6
   y11++     628.76        9.0
   y4+       645.47       22.5
   b11++     649.04       14.8
   a17+++    684.51        7.5
   y12++     693.32        8.9
   y5+       702.48       22.4
   b18+++    713.19        9.2
   b5+       727.69        8.2
   y19+++    742.45       11.6
   b19+++    750.91        8.1
   y13++     757.92       10.1
   a20+++    765.87        8.2
   a13++     782.73        8.0
   b13++     797.29       12.3
   b6+       798.44       14.4
   a21+++    808.55       19.9
   y6+       815.48       21.2
   b21+++    817.43       24.6
   y14++     822.21       12.0
   b22+++    841.47        7.8
   b14++     847.72       11.6
   y15++     873.12       19.6
   b23+++    879.18       16.2
   a7+       884.28        7.1
   b24+++    898.33       15.1
   b7+       912.52        8.1
   y25+++    940.50       10.7
   a26+++    954.58        8.5
   b16++     976.75        8.5
   b27+++    987.73        9.7
   b8+      1013.73        8.3
   y17++    1020.69       10.3
   b17++    1041.26        8.9
   a9+      1084.63        7.5
   b20++    1161.88        8.0
   b21++    1225.75       11.1
   b22++    1261.79       10.6
   b11+     1296.79        8.4
   b23++    1318.28       12.6
   b24++    1346.90       13.7
```

```
1118. MORG-339_phosphopeptide_TMT_18%ACN_032017_OT.20045.20045.4 (eLALSsPEDLTQDFEELkREER)

   Ion         m/z   Intensity
   y5+++     316.18        5.8
   b4++      329.20        6.4
   y7+++     397.20        6.5
   a11+++    480.18        5.8
   y9+++     488.86       18.5
   b11+++    489.53       20.4
   y6++      530.15        6.6
   y13+++    641.49       18.2
   b4+       656.38       12.3
   y14+++    679.81       10.3
   a16+++    695.86        8.8
   b11++     733.07       11.2
   a17+++    734.11        8.0
   b17+++    742.71       12.1
   b5+       743.43        9.8
   y16+++    755.26       36.7
   a12++     783.12       11.6
   a6+       882.16        6.0
   y20+++    901.52        9.2
   y12++     904.90        8.9
   a14++     914.57        5.8
   y21+++    938.88        7.1
   b20+++    957.57       10.4
   y13++     961.53        7.2
   y14++    1019.31       10.2
   y15++    1083.75        9.0
```

```
1119. MORG-339_phosphopeptide_TMT_10%ACN_032017_OT.13299.13299.3 (sRTLSQssESGtLPSGPPGHTMEVSc)

   Ion         m/z   Intensity
   a9++      669.55       10.7
   y13++     678.59       19.0
   b12++     845.69       12.3
   y16++     853.62       18.4
   a13++     888.61       11.5
   b6+       902.65       15.8
   b15++     994.44       18.6
   a16++    1009.08       30.8
   y9+      1017.40       18.2
   a7+      1041.41       24.1
   y10+     1114.41       12.3
   y20++    1128.54       13.0
   a19++    1134.32       15.4
   y11+     1171.69       16.5
   a8+      1208.86       18.9
   y13+     1355.63       30.2
   y14+     1468.68       13.5
   a11+     1482.01       14.1
```

```
1120. MORG-339_phosphopeptide_TMT_25%ACN_032017_OT.15780.15780.4 (gLYDGPVcEVSVTPktVTPASSAkTsPAk)

   Ion         m/z   Intensity
   a15+++    678.46       14.4
   b16+++    748.44       15.7
   y15+++    774.64       18.0
   b17+++    780.75       18.7
   y11++     792.27       16.4
   y5+       812.28       14.2
   a19+++    837.85       14.0
   b21+++    900.20       14.8
   a7+       903.09       13.0
   b22+++    928.77       14.8
   b7+       931.28       25.1
   y20+++    935.68       17.2
   a23+++    942.98       20.4
   b23+++    952.86       18.5
   y21+++    978.14       38.6
   y14++     982.59       61.3
   b24+++   1071.09       17.3
   b8+      1091.76       13.1
   b25+++   1104.84       18.7
   y25+++   1115.97       15.4
   y26+++   1154.83       12.9
   y15++    1160.72       17.8
   y6+      1170.16       17.7
   y16++    1209.36       12.9
   b18++    1221.84       15.7
```

```
1121. MORG-339_phosphopeptide_TMT_4%ACN_032017_OT.9286.9286.3 (aGPLGGsSYEEEEEEEGGGGGER)

   Ion         m/z   Intensity
   y3+       361.36        8.0
   y4+       418.30        9.3
   y5+       475.13       13.6
   y6+       532.27       11.4
   b4+       568.38        8.1
   y7+       589.37       23.8
   b6+       682.22       12.8
   y8+       718.40       16.1
   b12++     743.99       15.6
   y14++     746.58       10.5
   y9+       847.51       24.7
   b14++     872.75        9.8
   b15++     937.53       12.7
   y10+      976.43       12.5
   a16++     987.88        9.4
   b16++    1001.95       17.0
   b17++    1030.44       14.2
   b18++    1059.34       10.7
   a19++    1073.55        7.5
   y11+     1105.47        9.6
   b21++    1144.54        7.9
   b22++    1209.01        9.2
   b10+     1228.59       15.4
   b11+     1357.75        9.4
   b12+     1486.62        9.9
   b13+     1615.69       14.8
   b14+     1744.68        9.4
```

```
1122. MORG-339_phosphopeptide_TMT_18%ACN_032017_OT.16387.16387.3 (kADNPTPQTPGTSSLSQLGAYGDsEDsDS)

   Ion         m/z   Intensity
   y3+       387.79       14.4
   b3+       544.36       21.9
   b7+       953.19       13.9
   y8+       971.17       42.6
   a18++    1012.79       24.9
   a20++    1076.74       29.3
   y21++    1117.18       29.7
   y9+      1134.25       49.8
   a21++    1158.81       22.0
   y11+     1262.31       54.1
   b10+     1279.38       26.2
   y12+     1375.36       40.7
   y13+     1503.56       21.9
   y14+     1590.48       36.6
   a28++    1591.46       40.5
   y15+     1703.71       15.9
   y16+     1790.73       18.5
   y17+     1877.60       16.6
```

```
1123. MORG-339_phosphopeptide_TMT_4%ACN_032017_OT.19834.19834.2 (sAyFSDEEELsD)

   Ion         m/z   Intensity
   y2+       301.04        4.9
   b2+       388.20        5.7
   y3+       414.21       12.8
   y4+       543.26       16.8
   y5+       672.26        8.7
   b6+       980.47        7.1
   b7+      1109.38       14.4
   b8+      1238.55       15.3
   b9+      1367.58       24.7
   y11+     1464.70        4.9
   b10+     1480.59       21.4
```

```
1124. MORG-339_phosphopeptide_TMT_18%ACN_032017_OT.14307.14307.3 (fSVSPASPSSQQtPPPVtPR)

   Ion         m/z   Intensity
   y7++      422.22       12.7
   b7++      453.15       14.8
   b3+       563.41       16.0
   y5+       649.38       10.1
   y11++     684.69       10.7
   b12++     716.85       13.5
   y6+       746.40       13.1
   y13++     776.28       16.4
   a6+       790.39       20.1
   a13++     793.60       17.1
   b13++     807.31       11.6
   y7+       843.50       17.2
   a7+       877.29        9.5
   b15++     904.29       12.2
   b7+       905.58       16.7
   a8+       974.84        9.8
   a17++     988.73       16.9
   y18++     997.33        9.1
   b8+      1002.84       17.0
   y8+      1024.41       11.2
```

```
1125. MORG-339_phosphopeptide_TMT_14%ACN_032017_OT.20961.20961.3 (kEEsEESEDDMGFGLFD)

   Ion         m/z   Intensity
   y2+       281.09       10.1
   y4+       451.14       12.7
   y5+       598.26       10.7
   y6+       655.31       17.1
   b10++     744.35       10.6
   y13++     745.44        6.3
   b11++     810.04       15.6
   a12++     824.59        8.7
   y8+       901.48        6.4
   b14++     940.52        6.2
   y9+      1016.46       10.0
   a16++    1056.03       17.1
   y11+     1232.35        6.1
```

```
1126. MORG-339_phosphopeptide_TMT_8%ACN_032017_OT.17320.17320.4 (emLLEDVGsEEEPEEDDEAPFQEk)

   Ion         m/z   Intensity
   b1+       359.23        9.0
   y1+       376.33       11.1
   y5++      439.42       29.7
   y6++      474.64       10.7
   y2+       505.27       19.7
   b2+       506.46       10.4
   b7++      538.41       12.5
   y12+++    555.29       10.8
   b8++      567.06       17.6
   a13+++    586.27       10.0
   y8++      596.87        7.4
   y3+       633.42       16.6
   y9++      654.78       10.2
   b10++     715.27       15.8
   y10++     718.67        7.6
   b16+++    719.85        9.2
   b4+       732.40       16.8
   a17+++    749.13        8.3
   b11++     779.38       15.0
   y4+       780.61       14.8
   y18+++    791.83       11.6
   b18+++    801.39       10.0
   b19+++    824.36       12.3
   b12++     843.62        8.9
   b5+       861.54       18.4
   y5+       877.47       32.3
   a13++     878.54       22.9
   a21+++    896.69       35.6
   b22+++    948.46       15.7
   y14++     961.14       22.0
   b6+       976.52       13.3
   b7+      1075.50       22.6
   y7+      1077.58       14.1
   b16++    1078.70       10.3
   b8+      1132.59       13.5
   y17++    1137.68       14.9
   y8+      1192.57        9.7
   b18++    1201.22       10.4
   b19++    1236.17       11.7
   b10+     1428.67        8.2
   b12+     1686.85       16.5
```

```
1127. MORG-339_phosphopeptide_TMT_6%ACN_032017_OT.11396.11396.2 (eVVDSTTSsL)

   Ion         m/z   Intensity
   y3+       386.20        8.3
   b2+       458.31        9.2
   y4+       487.16        6.8
   b3+       557.28       13.1
   y5+       588.30        9.1
   y7+       790.40       10.1
   b6+       860.58       10.1
   y8+       889.49        7.7
   b7+       961.51        9.7
   b8+      1048.52        9.8
   b9+      1215.63        6.2
```

```
1128. MORG-339_phosphopeptide_TMT_18%ACN_032017_OT.22187.22187.3 (iVTVAAISQDsNPATPNVSTNmEEF)

   Ion         m/z   Intensity
   y11++     642.50       12.1
   a5+       685.21        9.8
   b5+       713.57       14.1
   y14++     783.60       12.0
   b6+       784.61       16.9
   y6+       786.35       10.4
   b13++     803.76       13.1
   b14++     839.13       20.8
   y15++     867.14        9.8
   y7+       873.35       25.8
   a15++     875.40        9.8
   b15++     889.73       37.3
   b7+       897.63       32.7
   a16++     923.66        7.6
   y16++     925.33        8.2
   a8+       956.15       13.4
   b18++    1044.79       37.9
   b9+      1112.94        8.3
   y10+     1183.52       45.1
   b10+     1227.76       10.1
   b22++    1269.59       10.1
   y11+     1284.53       19.3
   y13+     1452.61       12.7
   b12+     1508.80        9.1
   b15+     1777.98        7.4
```

```
1129. MORG-339_phosphopeptide_TMT_20%ACN_032017_OT.5198.5198.4 (vRLRsDGAAIPR)

   Ion         m/z   Intensity
   y1+       174.94        7.3
   y3++      193.19        6.3
   b4+++     252.26        7.5
   y2+       272.21       27.1
   y8+++     289.56        6.1
   a5+++     298.91        7.7
   b5+++     307.96       11.1
   b6+++     346.37       47.9
   a7+++     356.29       17.8
   b7+++     365.40       28.5
   b4++      378.08        4.8
   a8+++     379.65       26.4
   y3+       385.23       17.3
   b8+++     388.99       35.5
   b9+++     412.83       18.4
   y11+++    431.33        6.5
   a10+++    441.14       19.5
   a5++      447.56        5.2
   b10+++    450.37       11.3
   y4+       456.31       31.5
   a2+       457.33       16.8
   b5++      461.07        5.3
   a11+++    473.17        4.3
   b2+       485.46        4.4
   b6++      519.01        9.7
   y5+       527.36       13.5
   b7++      547.37       14.1
   a8++      569.24       11.8
   a3+       569.93        7.0
   b8++      583.04       23.3
   y6+       584.31       30.5
   b9++      618.59       38.8
   b10++     675.20        7.6
   y7+       699.49        9.1
```

```
1130. MORG-339_phosphopeptide_TMT_16%ACN_032017_OT.15335.15335.3 (mAPTPIPTRsPsDSSTASTPIAEQIER)

   Ion         m/z   Intensity
   b9++      597.96        7.9
   a4+       601.99        8.2
   b10++     681.64       11.0
   a11++     715.52        7.6
   y6+       745.49       11.5
   b6+       840.50        8.9
   b13++     871.30       15.5
   y8+       955.50       15.6
   b15++     957.61        9.4
   a16++     994.23        8.0
   y18++    1018.54        9.0
   a17++    1030.07        7.9
   b8+      1038.87       36.1
   y9+      1056.83       15.7
   y10+     1143.74       12.0
   y20++    1146.85        8.3
   a20++    1172.69       11.5
   b20++    1187.01        7.3
   b9+      1194.63        7.2
   y21++    1195.55       13.4
   y11+     1214.69        8.2
   y22++    1251.95        9.5
   y12+     1315.60       11.2
   y13+     1402.68        7.5
   y14+     1489.76        9.2
   y16+     1771.98       11.9
   y17+     1868.69       12.2
```

```
1131. MORG-339_phosphopeptide_TMT_12%ACN_032017_OT.17697.17697.3 (sSQFEGTFLGsEsDEDSVR)

   Ion         m/z   Intensity
   b1+       317.30        8.6
   y4+       476.23        6.5
   b9++      614.15        9.0
   b10++     642.56       10.7
   y11++     676.97        6.0
   a11++     711.34        6.0
   b12++     790.06       30.4
   b5+       808.42        5.5
   a6+       837.70        7.2
   b6+       865.58        7.1
   y7+       887.39        8.1
   y15++     894.63        5.8
   b14++     931.41        5.1
   b7+       966.36        9.5
   y18++    1075.29        8.1
   b8+      1113.67        7.3
   b18++    1146.51       10.5
   y10+     1240.56       12.5
   b10+     1283.61        8.3
   y11+     1353.49        6.8
```

```
1132. MORG-339_phosphopeptide_TMT_20%ACN_032017_OT.17903.17903.3 (ekPDsDDDLDIESFVTAk)

   Ion         m/z   Intensity
   b1+       359.16        7.9
   y3+       548.65       10.8
   y9++      619.51        6.7
   y4+       647.49       10.9
   b7++      663.47        7.1
   b8++      721.14        9.9
   b9++      777.36        8.7
   y5+       794.57       11.9
   b10++     834.81       16.0
   a11++     877.43        7.5
   y6+       881.05       12.5
   a4+       900.10       21.9
   b12++     956.18       14.5
   b13++     999.78        8.3
   y7+      1010.48        9.0
   a5+      1067.27        7.6
   b15++    1122.77       13.6
   y8+      1123.63       13.3
   b6+      1210.14        9.7
```

```
1133. MORG-339_phosphopeptide_TMT_8%ACN_032017_OT.24186.24186.3 (eAAGGNDssGATSPINPAVALE)

   Ion         m/z   Intensity
   y3+       332.22        9.9
   y6+       599.39       14.6
   b10++     618.35       11.5
   b11++     653.93       36.4
   a12++     690.12       26.2
   a6+       700.97       12.0
   b12++     704.38       29.0
   y7+       713.69       37.4
   b6+       729.66       34.5
   b13++     747.94       13.8
   a14++     781.83       13.9
   b14++     796.75       10.9
   b7+       844.06       15.7
   b15++     852.99       11.5
   y17++     901.89       10.5
   b16++     910.07       15.2
   y9+       923.60       35.3
   a8+       983.45       10.3
   y20++     994.09       24.6
   y10+     1010.65       24.9
   b8+      1011.57       16.1
   y21++    1029.87       21.8
   b19++    1043.53       32.6
   b20++    1079.11       44.1
   y11+     1111.53       16.8
   a21++    1121.49       11.0
   b21++    1135.74       15.9
   b11+     1306.38       12.9
   b12+     1407.60       11.8
   b13+     1494.53       15.0
   b16+     1818.64       11.0
```

```
1134. MORG-339_phosphopeptide_TMT_16%ACN_032017_OT.11061.11061.3 (aAPPPPPPPPPLEsSPR)

   Ion         m/z   Intensity
   b4++      283.61       21.8
   b5++      332.19       18.4
   a2+       344.35        8.3
   b2+       372.23       23.2
   b6++      380.67       25.7
   a7++      415.18       19.2
   b7++      429.19       17.3
   b3+       469.41       10.6
   y8++      481.83       12.5
   b9++      526.38       11.3
   y9++      529.94       10.3
   a4+       538.39       11.5
   y10++     579.11       14.5
   b11++     623.09       21.0
   y11++     627.19       27.0
   a5+       635.60       20.9
   y5+       655.04        7.6
   b5+       663.39       19.8
   y13++     724.47       21.5
   a6+       732.66       10.0
   b13++     744.63       13.5
   b6+       760.41       12.6
   y15++     821.62       22.6
   a7+       829.42        9.9
   y7+       865.40        9.0
   y8+       962.67        9.8
   y9+      1059.49       19.4
   y10+     1156.49       29.3
   y11+     1253.74       18.6
   y12+     1350.80        7.7
```

```
1135. MORG-339_phosphopeptide_TMT_14%ACN_032017_OT.14764.14764.2 (gLSAStVDLSSSS)

   Ion         m/z   Intensity
   y3+       280.19       13.3
   y4+       367.16       15.6
   y5+       480.33       16.2
   b4+       558.40        9.8
   y6+       595.31       23.4
   b5+       645.06       11.3
   y7+       694.19       12.6
   b6+       826.54        9.6
   b7+       925.47       35.5
   y9+       962.42       14.4
   y10+     1033.38        7.6
   b8+      1040.46       21.8
   b9+      1153.53       49.9
   b10+     1240.52       31.2
```

```
1136. MORG-339_phosphopeptide_TMT_8%ACN_032017_OT.12501.12501.3 (vGEcSGPAAVNNSsDtESVPsPR)

   Ion         m/z   Intensity
   b9++      530.11       12.0
   y4+       536.38       20.8
   y5+       635.35       11.6
   b11++     636.49        7.4
   b12++     693.39        9.6
   a14++     805.90        8.5
   y13++     815.69       11.5
   b6+       819.35        8.6
   y14++     864.33       16.1
   b15++     877.44        7.9
   y15++     900.46       18.3
   b7+       916.03        9.5
   b16++     968.59        8.5
   y17++     984.22       16.4
   b8+       987.46       12.9
   a17++    1018.66       17.3
   b17++    1033.18       17.7
   y19++    1056.13        9.1
   y9+      1147.14       15.0
   b10+     1157.61       15.4
   b21++    1257.87       10.1
   b11+     1271.46       10.8
   y12+     1515.45        7.6
   y13+     1629.62        9.8
```

```
1137. MORG-339_phosphopeptide_TMT_14%ACN_032017_OT.18018.18018.3 (dkEETEGNESSEATsPVNAIYSLA)

   Ion         m/z   Intensity
   b6++      596.04        8.3
   b9++      745.91       12.6
   b11++     833.17       14.9
   b12++     897.69       23.5
   a13++     919.30        8.5
   b13++     933.18       18.2
   y9+       947.78       16.9
   b4+       960.97        8.5
   a14++     969.46       10.3
   b14++     983.62       30.8
   y19++    1009.49       31.4
   b15++    1066.81        7.1
   y10+     1114.49       14.7
   b16++    1115.46       13.7
   b17++    1165.33       24.9
   y22++    1189.85       11.0
   b6+      1190.51        7.2
   y11+     1215.68        8.4
   b18++    1222.33       16.0
   b19++    1257.96       22.8
   y12+     1286.64       11.2
   b20++    1314.41       36.9
   b21++    1395.92       25.3
   b22++    1439.07        8.5
```

In [ ]:
